# Bar-seq strategies for the LeishGEdit toolbox

**DOI:** 10.1101/2020.03.19.998856

**Authors:** Tom Beneke, Eva Gluenz

## Abstract

The number of fully sequenced genomes increases steadily but the function of many genes remains unstudied. To accelerate dissection of gene function in *Leishmania* spp. and other kinetoplastids we developed previously a streamlined pipeline for CRISPR-Cas9 gene editing, which we termed LeishGEdit [1]. To facilitate high-throughput mutant screens we have adapted this pipeline by barcoding mutants with unique 17-nucleotide barcodes, allowing loss-of-function screens in mixed populations [2]. Here we present primer design and analysis tools that facilitate these bar-seq strategies. We have developed a standalone easy-to-use pipeline to design CRISPR primers suitable for the LeishGEdit toolbox for any given genome and have generated a list of 14,995 barcodes. Barcodes and oligos are now accessible through our website www.leishgedit.net allowing to pursue bar-seq experiments in all currently available TriTrypDB genomes (release 41). This will streamline CRISPR bar-seq assays in kinetoplastids, enabling pooled mutant screens across the community.

**Highlights:**

- Developing tools for pooled bar-seq mutant screens across the kinetoplastid community
- Development of a standalone script to design primers suitable for the LeishGEdit toolbox
- Generation of 14,995 barcodes that can be used for bar-seq strategies in kinetoplastids
- Bar-seq primers for all TriTrypDB genomes (release 41) can be obtained from www.leishgedit.net

## Article

Since the first reports of genetic manipulations using gene replacement strategies in *Leishmania* parasites nearly three decades ago [3] about 200 genes have been subjected to gene deletion in *Leishmania* spp. by researchers around the globe as of April 2018 [4]. This number has been dramatically increased since the introduction and use of CRISPR-Cas9 technologies in the field of kinetoplastids [1, 5-9]. Our approach for generation of CRISPR null mutants, which we termed LeishGEdit, was designed to facilitate high-throughput mutant screens [1, 10]. The first step involves engineering a cell line expressing Cas9 nuclease and T7 RNA polymerase constitutively. Using this cell line, linear sgRNA and donor DNA constructs can be transfected in parallel, allowing for single guide RNA (sgRNA) transcription *in vivo* and integration of donor DNA constructs with 30 nt homology flanks (HF) identical to the target locus. This method does not involve any cloning procedures, PCR purifications or *in vitro* transcription prior to transfections and we have previously used it to generate 100 null mutants in order to dissect flagellar function in *Leishmania mexicana* [2].

To study such a large cohort of mutants, we have recently adapted the LeishGEdit method further and introduced a barcode analysis by sequencing (bar-seq) strategy for *Leishmania mexicana* [2]. Bar-seq methods are powerful scalable approaches that are applied to track the phenotype of several mutants at once [11, 12]. The technique was originally developed to analyze libraries of thousands of *Saccharomyces cerevisiae* [13] or *Schizzosaccharomyces pombe* [14] gene deletion mutants, but has since been applied for other genome-wide screens in multiple parasites, including *Plasmodium berghei* [15, 16], *T. brucei* [17] and *Toxoplasma gondii* [18]. These studies have shown that bar-seq strategies are powerful for the analysis of phenotypes that result in differential cell growth or survival, or enrichment of a sub-population in a particular place from which DNA can be isolated.

In a typical bar-seq assay, each mutant is tagged with a unique barcode to allow quantitation of individual mutants within a mixed population. The barcode might consist of a short DNA sequence optimized for Illumina sequencing or also a sequence that is actually required for targeting the gene itself (e.g. in bar-seq RNAi or CRISPR libraries as explained in (1) and (2) further below). For the bar-seq analysis, mixed populations can either be generated by pooling individually generated mutants, or mutants can already be generated as pool, e.g. if a method is available that requires only a single target vector to silence (e.g. RNAi) or delete (e.g. some CRISPR methods) a gene of interest. The pool of barcoded mutants is then subjected to the experimental conditions of interest and DNA samples are extracted at the beginning of the experiment and at defined intervals thereafter. This allows to track each barcode over the experimental time-course. Using next-generation sequencing, each barcode sequence is counted from PCR amplicons, and the relative abundance of each barcode within the entire pool can be calculated, giving a measure of fitness for each mutant. There are at least four different types of bar-seq libraries, which may be grouped as follows: (1) bar-seq RNAi libraries, in which RNAi vector inserts are used both for knockdown of the target gene and barcode read-out [17], (2) bar-seq CRISPR libraries, in which sgRNAs in vectors are used for deletion of the target ORF and also serve as a barcode for the read-out of the assay [18], (3) bar-seq pooled knockout libraries, in which mutants are generated in pools by transfecting barcoded vectors [15, 16] and (4) bar-seq individual knockout libraries, in which each mutant is generated individually and barcoded in the process of gene deletion before pooling of mutants [13, 14]. The latter can be achieved by adapting the LeishGEdit toolbox [2] (Fig. 1A).

**Figure 1.**
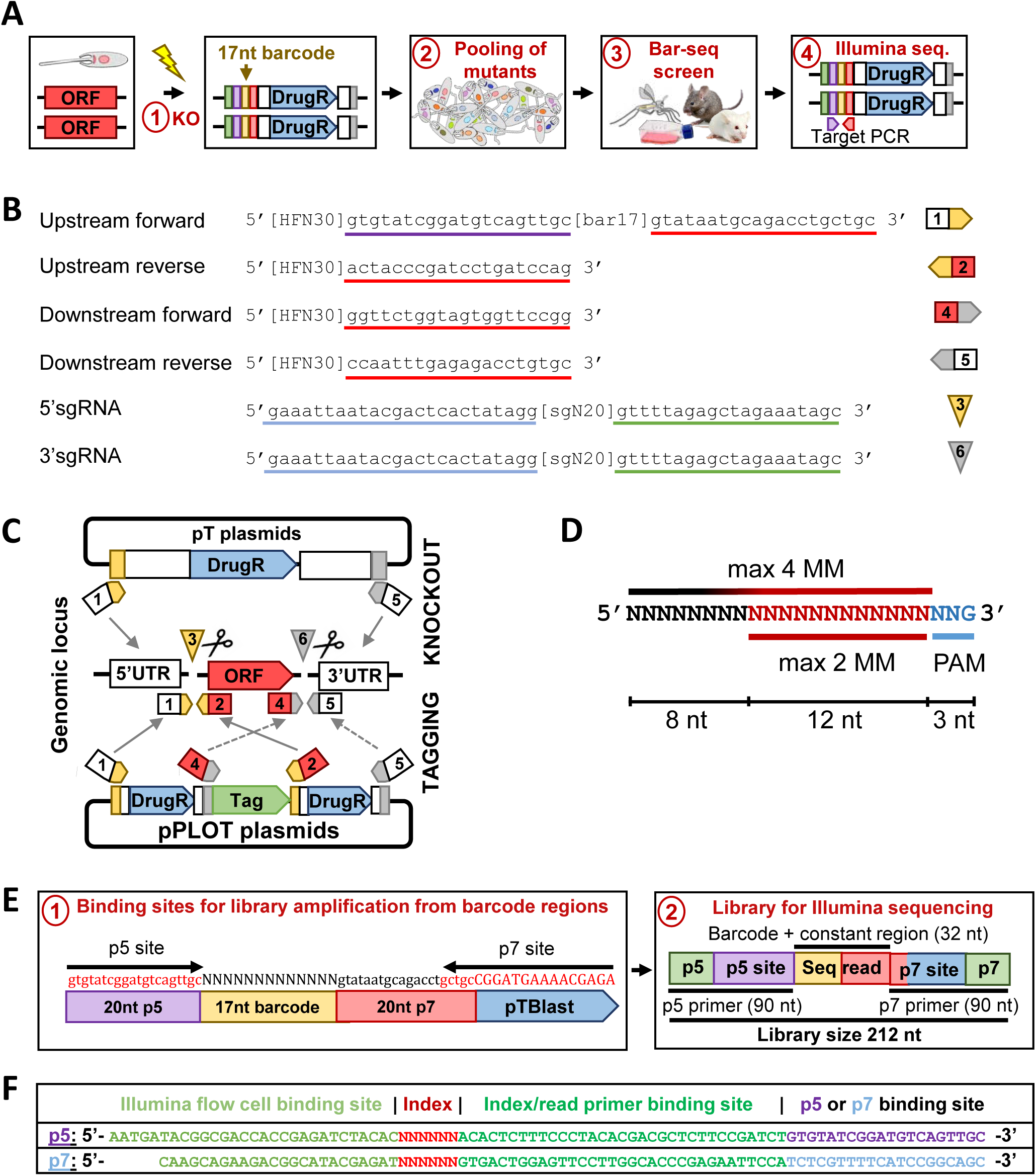
Overview of primer design and a bar-seq strategy for the LeishGEdit toolbox. **(A)** Overview of the previously designed strategy for bar-seq phenotyping [2]. (1) A cell line expressing Cas9 and T7 RNAP is subjected for a double allele deletion using two different drug selectable markers. A 17 nt barcode (yellow), surrounded by two constant regions (purple and red) is inserted into the target locus by using donor DNA with 30 nt HF (green and gray). (2) Barcoded mutants are pooled and (3) analyzed in an *in vitro* or *in vivo* screen. (4) Barcode abundance is read out by amplifying barcodes using their surrounding constant regions. **(B and C)** Figure adapted from Beneke and Gluenz [10]. Shown is the PCR strategy for donor DNA amplification from pT and pPLOT plasmids. **(B)** Overview of constant and variable sequences in the LeishGEdit primers: Forward and reverse primers for donor DNA amplification contain target-gene specific 30 nt homology flanks ([HFN30]) adjacent to pT and pPLOT plasmid primer binding sites (underlined in red). The upstream forward primer can be barcoded ([bar17]). An additional primer binding site is required for the read out by sequencing (underlined in purple). Primers for sgRNA template amplification contain the T7 promotor sequence (underlined in blue), the 20 nt target sequence ([sgN20]) and an overhang sequence to the sgRNA backbone sequence (underlined in green). **(C)** pT plasmids consist of a *L. mexicana* derived 5’ and 3’UTR and a drug resistance marker gene to allow gene replacement by drug selection. pPLOT plasmids contain drug resistance markers and *Crithidia* and *T. brucei* UTRs, as well as myc epitope tags in-frame with various protein tags. pPLOTs can be used for amplification of tagging cassettes, allowing generation of fusion proteins with epitope tags fused at the N- or C-terminus. **(D)** Criteria used by CCTop to identify suitable sgRNA target sites. sgRNA target sites may have up to 2 MM in the first 12 nt upstream of the PAM site or up to 4 MM in the entire sgRNA target sequence [22]. **(E and F)** Illumina sequencing strategy for reading out barcode abundances. **(E)** (1) Two long primers (p5 and p7 primers; specified in (F)) bind to constant regions adjacent to the 17 nt barcode (binding sites in red font). (2) Library size and expected sequencing read length of the amplicon is indicated. **(F)** Primer sequences of p5 and p7 primers used for Illumina sequences. Long p5 and p7 primers contain flow cell binding sites, additional indices for Illumina sequencing and an index/read Illumina sequencing binding site.

The chosen bar-seq strategy often depends on available tools in the organism to be screened. For example most *Leishmania* spp. lack the RNAi machinery [19] and as yet no single vector-based CRISPR libraries have been reported for any kinetoplastids, possibly because of the challenge of reliably targeting two alleles of a gene in one go for a knockout library.

Here we report how we have adapted the LeishGEdit toolbox to enable bar-seq fitness screens of individual knockout libraries: A new feature has been added to our CRISPR primer design website www.leishgedit.net, allowing to design primers for barcoding of numerous kinetoplastid species. Additionally, we provide a standalone easy-to-use pipeline that can be used locally to design CRISPR primers with enhanced sgRNA design, and compatible with the LeishGEdit toolbox for any given genome.

The LeishGEdit primer design pipeline designs in total six primer sequences for each given ORF in the genome to enable CRISPR-Cas9 gene editing, allowing to tag a gene of interest at its N- or C-terminus with numerous available tags and to delete both alleles of the ORF (Fig. 1B and C). Two sgRNA primers are designed, one targeting the 5’UTR of the target gene and one the 3’UTR. sgRNA primers consist of a T7 RNAP promotor sequence, a 20 nt sgRNA target sequence to introduce the DSB at a locus of interest and a 20 nt overlap to the CRISPR-Cas9 backbone sequence allowing generation of sgRNA templates by PCR. An additional universal primer, containing the entire sgRNA backbone sequence [20] is needed to amplify both sgRNAs. Four primers are designed for pPLOT and pT plasmid amplification and can be used in different combinations to produce donor DNA. These primers include the: upstream forward primer (#1), upstream reverse primer (#2), downstream forward primer (#3) and downstream reverse primer (#4). An additional primer can be optionally designed to allow for CRISPR-Cas9 mediated gene editing using donor constructs amplified from pPOT plasmid templates [21]. Donor DNA primers contain the 30 nt HF sequence immediately adjacent to the sgRNA target sequence and its PAM site, as well as primer binding sites compatible with pT, pPLOT and pPOT plasmids. While the upstream forward primer and downstream reverse primer position is always variable depending on the chosen sgRNA, the upstream reverse primer (#2) and downstream forward primer (#4) are designed at the same positions for each gene.

To produce a standalone pipeline for LeishGEdit primer design, we produced a bash script that designs sgRNA primers using CCTop [22] for the prediction of target sites, and then designs donor DNA primers with sequences in close proximity to the sgRNA target sequence (Supplementary file 1). The sgRNA target sequence is selected from a 130 nt search window upstream of the start codon or downstream of the stop codon. The highest scoring sgRNA within this window is chosen based on the CCTop scoring pipeline. The number of alignments to the genome with mismatches (MM) for any given sgRNA sequence is the main scoring criterion: Specifically, CCTop finds potential sgRNA target sites that have up to 2 MM in the first 12 nt upstream of the PAM site or up to 4 MM in the entire sgRNA target sequence and sorts these by least MM for the highest scoring sgRNA [22] (Fig. 1D). There is experimental evidence that Cas9 sgRNA complexes are functional to introduce double-strand breaks (DSBs) when their target site has up to 4 MM [23-26]. Additionally, the number of perfect matches of a selected sgRNA sequence (23 nt, including the protospacer adjacent motif ‘NGG’) within the genome is determined. Since this is computed independently from the CCTop pipeline, this count gives an additional indication on potential sgRNA off-target sites.

Thus, to provide an indication for sgRNA specificity the primer design outcome gives both these outputs to designed oligo sequences: (1) the number of perfect sgRNA matches in the target genome and (2) the number of imperfect sgRNA matches with 1-4 MM in the target genome.

To perform bar-seq assays using the LeishGEdit toolbox, the upstream forward primer (#1) can be barcoded [2]. The upstream forward primer is modified by inserting a 17 nt barcode and a 20 nt constant region in-between the existing 30 nt HF and 20 nt pT-pPLOT-pPOT primer binding site. Using the additional 20 nt constant region and 20 nt pT-pPLOT-pPOT primer binding site allows reading out barcode abundance by using Illumina amplicon sequencing strategies. Amplicon libraries can be produced as previously shown in a single PCR step library preparation protocol using custom designed p5 and p7 primers [2] (Fig. 1E and F). Barcodes were initially generated by using barcode generator 2.8 [27] and then customized to select barcodes with 40-60% GC content (for unbiased Illumina sequencing), at least 3 MM between barcodes and allowing no blast hit in *Lutzomyia longipalpis* (BioProject: PRJNA20279), *Mus musculus* (BioProject: PRJNA169) and multiple kinetoplastid genomes available on TriTrypDB (release 41) [28], including *Blechomonas ayalai* B08-376, *Crithidia fasciculata* CfCl, *Endotrypanum monterogeii* LV88, *Leishmania aethiopica* L147, *Leishmania arabica* LEM1108, *Leishmania braziliensis* MHOMBR75M2903, *Leishmania braziliensis* MHOMBR75M2904, *Leishmania donovani* BPK282A1, *Leishmania enriettii* LEM3045, *Leishmania gerbilli* LEM452, *Leishmania infantum* JPCM5, *Leishmania major* Friedlin, *Leishmania major* LV39c5, *Leishmania major* SD75.1, *Leishmania mexicana* MHOMGT2001U1103, *Leishmania panamensis* MHOMCOL81L13, *Leishmanaia panamensis* MHOMPA94PSC1, *Leishmania pyrrhocoris* H10, *Leishmania seymouri* ATCC30220, *Leishmania* spp. MARLEM2494, *Leishmania tarentolae* ParrotTarII, *Leishmania tropica* L590, *Leishmania turanica* LEM423, *Paratrypanosoma confusum* CUL13, *Trypanosoma brucei* Lister427, *Trypanosoma brucei* TREU927, *Trypanosoma brucei gambiense* DAL972, *Trypanosoma congolense* IL3000, *Trypanosoma cruzi* CLBrenerEsmeraldo-like, *Trypanosoma cruzi* CLBrenerNon-Esmeraldo-like, *Trypanosoma cruzi* CLBrener, *Trypanosoma cruzi* Dm28c, *Trypanosoma cruzi* SylvioX10-1-2012, *Trypanosoma cruzi* marinkelleiB7, *Trypanosoma evansi* STIB805, *Trypanosoma grayi* ANR4, *Trypanosoma rangeli* SC58, *Trypanosoma theileri* Edinburgh and *Trypanosoma vivax* Y486. This yielded a total of 14,995 barcodes (Supplementary file 2), which can be used for fitness screens of many kinetoplastid species in culture and in commonly used laboratory models for *in vivo* infections (sand fly and mouse). Thus, with these parameters a minimum barcode length of 17 nt was found to be sufficient for generating enough barcodes for the number of ORFs in available kinetoplastid genomes.

Finally, to facilitate the analysis of bar-seq read out data, we have produced a bash script that allows counting of all 14,995 barcodes across de-multiplexed Illumina sequencing samples (Supplementary file 3). Barcodes are counted by determining the total number of occurrences of the 17nt barcode sequence within each sequencing sample (allowing 0 nt MM). The total of reads within one sample is also determined, which allows to normalize barcode counts across one sample by calculating their relative proportion within the pool. Subsequently, the relative proportion of each mutant can be used to calculate the “mutant fitness” of each mutant in the pool as determined previously [2]. Depending on the design of experiment “mutant fitness” can be calculated by dividing the barcode counts for a given time point by the barcode counts at the start of a bar-seq screen, the previous time point or a respective control at an identical time point (e.g. treated vs. non-treated culture in drug treatment screens).

We have used this new LeishGEdit primer design pipeline to design primers for all currently available TriTrypDB genomes as listed above (for DB release 41) (Supplementary file 4) and allocated a barcode to every single gene in each genome. The analysis of sgRNA MM counts, as well as perfect match counts show that the large majority of designed sgRNAs have only one perfect match within the genome and no additional imperfect matches with 1-4 MM (Fig. 2A and B). To allow easy-to-use access to these resources we have made these primer designs and scripts for analysis available on our CRISPR primer design website www.leishgedit.net. This will help other researchers in the community to perform CRISPR bar-seq assays and contribute to the standardization of methods, for example enabling direct comparisons of mutants generated in different laboratories.

**Figure 2.**
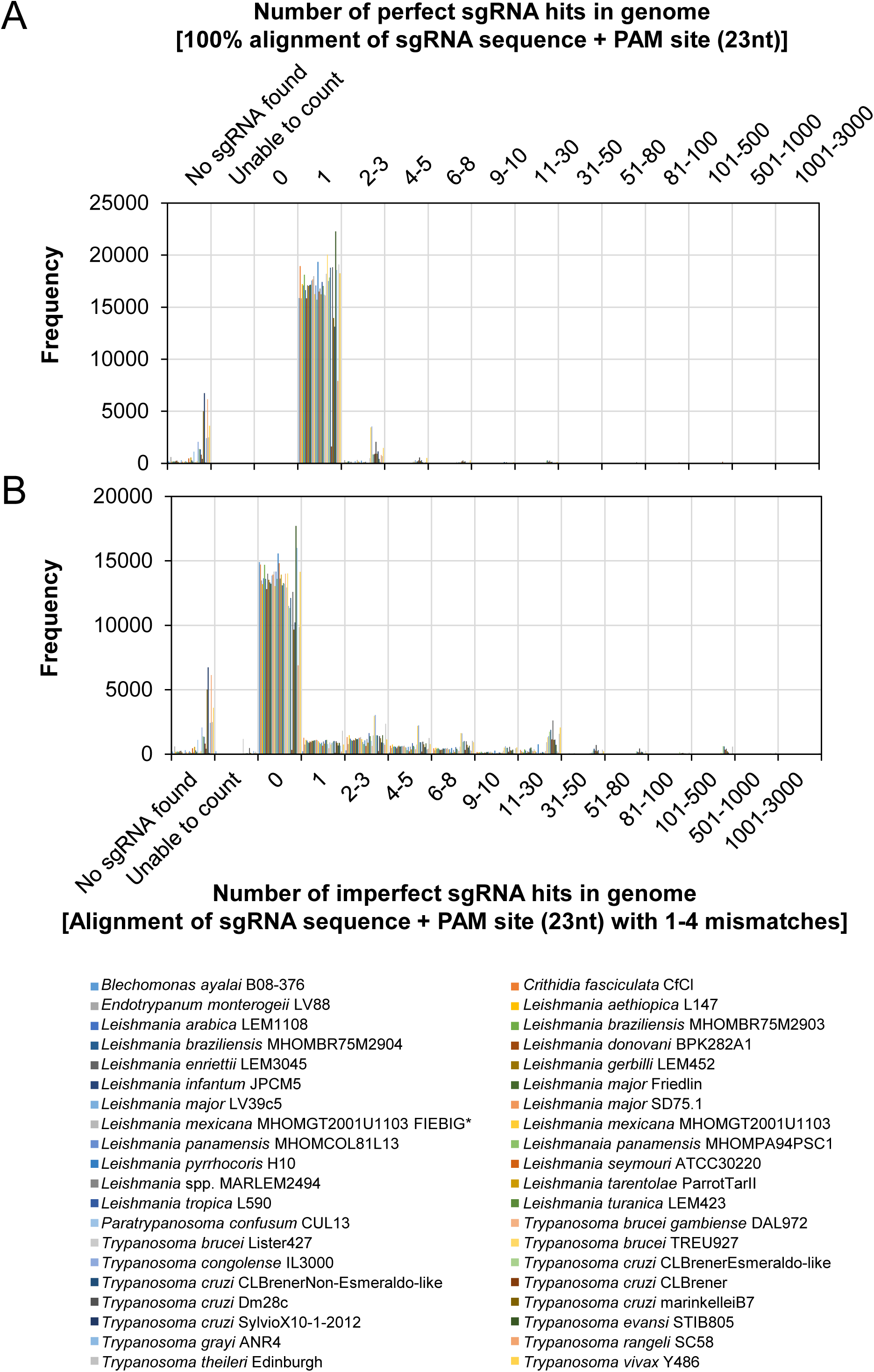
Number of perfect and imperfect sgRNA alignments for TriTrypDB genomes (release 41). The LeishGEdit standalone primer design pipeline has been used to design primers for all TriTrypDB genome annotations (DB release 41). Histograms show the number of **(A)** perfect and **(B)** imperfect sgRNA alignments (as defined in Fig. 1D) to the target genome. X axis: Categories of histogram, showing number of perfect or imperfect hits, number of instances where no sgRNA was found and sgRNAs where the number of alignments could not be determined. Y axis: Frequency of X axis categories. Asterisk: sgRNAs for *L. mexicana* were designed using gene models from Fiebig, Kelly [29] and are termed *Leishmania mexicana* MHOMGT2001U1103 FIEBIG.

## Author contributions

EG was supported by a Royal Society University Research Fellowship (UF100435 and UF160661). TB was supported by a studentship (MRC reference 15/16_MSD_836338) and a PhD to first post-doc position award from the Oxford-MRC Doctoral Training Partnership.

## Acknowledgement

We like to thank Edward Hookway for useful advice on design and read-out strategies of bar-seq assays.

## Supporting information

**Supplementary file 1. A standalone pipeline for LeishGEdit primer design.**

Automated primer design bash script to generate all six primers needed for LeishGEdit gene editing. Donor DNA primers, including upstream forward primer (#1), upstream reverse primer (#2), downstream forward primer (#4), downstream reverse primer (#5), contain the 30 nt HF sequence adjacent to the sgRNA target sequence and its PAM site, as well as primer binding sites compatible with pT, pPLOT and pPOT plasmids. Additionally, two sgRNA primers are designed, one targeting the 5’UTR of the target gene (#3) and one the 3’UTR (#6). The script output gives further information for sgRNAs, including the imperfect CCTop sgRNA counts (potential sgRNA target sites that have up to 2 MM in the first 12 nt upstream of the PAM site or up to 4 MM in the entire sgRNA target sequence), as well as sgRNA perfect match counts within the entire genome. Instructions for the usage of the script are contained within the “Readme” file.

**Supplementary file 2. List of generated Barcodes.**

This file contains 14,995 17nt barcodes for bar-seq experiments. Barcodes were generated as described in the main text.

**Supplementary file 3: A barcode counter script for all generated barcodes.**

An automated bash script for counting barcodes in de-multiplexed bar-seq samples. The script is set to count barcodes with 0 MM over the 17nt barcode, but can be modified if desired to count also barcodes with MM. Instructions for the usage of the script are contained within the “Readme” file.

**Supplementary file 4. LeishGEdit primer design for all available genomes on TriTrypDB.**

The LeishGEdit pipeline (Supplementary file 1) has been used to design primers for all currently available TriTrypDB genomes (for DB release 41). All files are available on www.leishgedit.net.

