## Supplementary_file_1 for "Bar-seq strategies for the LeishGEdit toolbox": bedInterval.py

```
'''
Created on Mar 20, 2014

@author: juan
'''


class MyInterval(object):
    def __init__(self,start,end,value):
        self.start = start
        self.end = end
        self.value = value

class BedInterval( object ):
    '''
    classdocs
    '''


    def __init__(self):
        '''
        Constructor
        '''
        self.chroms = {}
        
    def insert( self, chrom, start, end, gene_id, gene_name ):
        from bx.intervals.intersection import Interval
        from bx.intervals.intersection import IntervalNode 
        from bx.intervals.intersection import IntervalTree
        if chrom in self.chroms:
            self.chroms[chrom].insert( start, end, MyInterval(start,end,[gene_id, gene_name]) )
        else:
            self.chroms[chrom] = IntervalTree()
            self.chroms[chrom].insert( start, end, MyInterval(start,end,[gene_id, gene_name]) )

    def loadFile(self, file):
        bedFile = open(file,'r')
        for line in bedFile:
            line = line.rstrip('\n').split('\t')
            self.insert(line[0], int(line[1]), int(line[2]), line[3], line[4])
                   
    def overlaps(self, chrom, start, end):
        if not chrom in self.chroms:
            return False
        overlapping = self.chroms[chrom].find(start, end)
        if len(overlapping)>0:
            return True
        else:
            return False
        
    def closest(self,chrom, start, end):
        if not chrom in self.chroms:
            return ['NA', 'NA', 'NA']
        
        #first checking if this site overlaps any from the loaded file
        overlapping = self.chroms[chrom].find(start, end)
        if len(overlapping)>0:
            return [overlapping[0].value[0],overlapping[0].value[1],0]
        
        #to the left
        #it finds features with a start > than 'position'
        left = self.chroms[chrom].before(start, max_dist=1e5)
        
        #to the right
        #it finds features with a end < than 'position'
        right = self.chroms[chrom].after(end, max_dist=1e5)
        
        if len(left)>0:
            if len(right)>0:
                distLeft = max(0,1 + start - left[0].end)
                distRight = max(0,1 + right[0].start - end)
                if distLeft < distRight:
                    return [left[0].value[0],left[0].value[1],distLeft]
                else:
                    return [right[0].value[0],right[0].value[1],distRight]
            else:
                distLeft = max(0,1 + start - left[0].end)
                return [left[0].value[0],left[0].value[1],distLeft]
        else:
            if len(right)>0:
                distRight = max(0,1 + right[0].start - end)
                return [right[0].value[0],right[0].value[1],distRight]
            else:
                return ['NA', 'NA', 'NA']
        
        
```
