## Supplementary_file_1 for "Bar-seq strategies for the LeishGEdit toolbox": CCTop.py

```
# -*- coding: UTF-8 -*-

'''
Created on 16 Jun 2015

@author: juanlmateo
'''

import sys
import argparse
import textwrap
import os
# from subprocess import Popen, PIPE
import re
import math

from bedInterval import BedInterval

iupac_code = {'R':'[AG]', 'Y':'[CT]', 'S':'[GC]', 'W':'[AT]', 'K':'[GT]', 'M':'[AC]', 'B':'[CGT]', 'D':'[AGT]', 'H':'[ACT]', 'V':'[ACG]', 'N':'[ACGT]'}

def build_expression(seq):
    result = ''
    for c in seq:
        if c in iupac_code:
            result = result + iupac_code[c]
        else:
            result = result + c
            
    return result


def reverse_complement(sequence):
    rev_comp = []
    
    for idx in range(len(sequence) - 1, -1, -1):
        if sequence[idx] == 'A':
            rev_comp = rev_comp + ['T']
        elif sequence[idx] == 'C':
            rev_comp = rev_comp + ['G']
        elif sequence[idx] == 'G':
            rev_comp = rev_comp + ['C']
        elif sequence[idx] == 'T':
            rev_comp = rev_comp + ['A']
        else:
            rev_comp = rev_comp + ['N']
    return "".join(rev_comp)

def reverse_complementPAM(sequence):
    # wilcard characters are replaced by '*', treated as mismatches 
    rev_comp = []
    
    for idx in range(len(sequence) - 1, -1, -1):
        if sequence[idx] == 'A':
            rev_comp = rev_comp + ['T']
        elif sequence[idx] == 'C':
            rev_comp = rev_comp + ['G']
        elif sequence[idx] == 'G':
            rev_comp = rev_comp + ['C']
        elif sequence[idx] == 'T':
            rev_comp = rev_comp + ['A']
        else:
            rev_comp = rev_comp + ['*']
    return "".join(rev_comp)

def getOffTargetsNGG(sequence, bowtiePath, indexPath, exons, genes, coreMismatches, coreRange, totalMismatches, revPAM):
    
    if (coreMismatches == "NA" or coreRange == "NA"):
        command = bowtiePath + os.path.sep + "bowtie " + " -a " + indexPath + " -n " + str(2 + 1)  # plus 1 mismatch in the PAM, maximum 3!
        command = command + " -l " + str(2 + len(revPAM))  # minimum 5, additional 3 bases from the PAM
        command = command + " -e " + str(totalMismatches * 30 + 30)
        command = command + " -y "  # try hard to find all mismatched seeds
        command = command + " --quiet "  # just print alignments, no info
        command = command + " -c " + revPAM + reverse_complement(sequence[:-len(revPAM)])
    else:
        command = bowtiePath + os.path.sep + "bowtie " + " -a " + indexPath + " -n " + str(coreMismatches + 1)  # plus 1 mismatch in the PAM, maximum 3!
        command = command + " -l " + str(coreRange + len(revPAM))  # minimum 5, additional 3 bases from the PAM 
        command = command + " -e " + str(totalMismatches * 30 + 30)
        command = command + " -y "  # try hard to find all mismatched seeds
        command = command + " --quiet "  # just print alignments, no info
        command = command + " -c " + revPAM + reverse_complement(sequence[:-len(revPAM)])
    bowtieOutput = os.popen(command, "r")
    offtargets = []
    
    while True:
        line = bowtieOutput.readline()
        if not line: break
        
        columns = line.rstrip('\n').split('\t')
        mismatches = columns[7].split(':')
        # mismatches in the PAM are not allowed
        # bowtie shows the mismatches wrt. the input sequence, independently of the orientation of the alignment
        if int(mismatches[0]) < (len(revPAM) - 1):
            continue
        
        newOfftarget = Offtarget(False, columns[2], columns[1], int(columns[3]), columns[7], columns[4], len(sequence), 3, coreRange)
        newOfftarget.setGeneInfo(exons, genes)
        offtargets.append(newOfftarget)
    
    return offtargets

# Cpf1
def getOffTargetsTTTN(sequence, bowtiePath, indexPath, exons, genes, totalMismatches, PAM):
    
    
    command = bowtiePath + os.path.sep + "bowtie " + " -a " + indexPath + " -n " + str(2 + 1)  # plus 1 mismatch in the PAM, maximum 3!
    command = command + " -l 6"  # minimum 5, additional 4 bases from the PAM
    command = command + " -e " + str(totalMismatches * 30 + 30)
    command = command + " -y "  # try hard to find all mismatched seeds
    command = command + " --quiet "  # just print alignments, no info
    command = command + " -c TTTN" + sequence[len(PAM):]
    bowtieOutput = os.popen(command, "r")
    offtargets = []
    
    while True:
        line = bowtieOutput.readline()
        if not line: break
        
        columns = line.rstrip('\n').split('\t')
        mismatches = columns[7].split(':')
        # mismatches in the PAM are not allowed
        # bowtie shows the mismatches wrt. the input sequence, independently of the orientation of the alignment
        if int(mismatches[0]) < (len(PAM) - 1):
        # if mismatches[0] == '0' or mismatches[0] == '1':
            continue
        
        newOfftarget = Offtarget(True, columns[2], columns[1], int(columns[3]), columns[7], columns[4], len(sequence), len(PAM), "NA")
        newOfftarget.setGeneInfo(exons, genes)
        offtargets.append(newOfftarget)
    
    return offtargets

def getOffTargetsOtherPAMs(sequence, bowtiePath, indexPath, exons, genes, totalMismatches, PAM):
    # NNGRRT, NNNNGATT, NNAGAAW, NAAAAC 
    # A**C**, AATC****, *TTCT**, GTTTT*
    revPAM = reverse_complement(PAM)
    countWildcard = revPAM.count('N')
    countWildcardInSeed = revPAM[0:5].count('N')
    bowtieOutput = os.popen(bowtiePath + os.path.sep + "bowtie " + " -a " + indexPath + " -n " + str(countWildcardInSeed) + 
                            " -l 5" + 
                            " -e " + str(totalMismatches * 30 + countWildcard * 30) + 
                            " -y " +  # try hard to find all mismatched seeds
                            " --quiet " +  # just print alignments, no info
                            " -c " + revPAM + reverse_complement(sequence[:-len(PAM)]), "r")
    offtargets = []
    
    while True:
        line = bowtieOutput.readline()
        if not line: break
        columns = line.rstrip('\n').split('\t')
        mismatches = columns[7].split(',')
        # we need to check the the matched sequence has a valid PAM 
        if PAM == "NNGRRT":  # A**C**
            # mismatches substituting the 'R' wildcard (first two) must be 'A' or 'G'
            if (columns[1] == '-' and not (mismatches[0][2] in 'AG' and mismatches[1][2] in 'AG')) or (columns[1] == '+' and not (mismatches[0][2] in 'TC' and mismatches[1][2] in 'TC')):
                continue
        if PAM == "NNNNGATT":  #AATC****
            # all mismatches in the seed (l 5) allowed
            pass
        if PAM == "NNAGAAW":  # *TTCT**
            # mismatches substituting the 'W' wildcard (first one) must be 'A' or 'T'
            if not (mismatches[0][2] in 'AT'):
                continue
        if PAM == "NAAAAC":  # GTTTT*
            # all mismatches in the seed (l 5) allowed
            pass
        
        newOfftarget = Offtarget(False, columns[2], columns[1], int(columns[3]), columns[7], columns[4], len(sequence), len(PAM), "NA")
        newOfftarget.setGeneInfo(exons, genes)
        offtargets.append(newOfftarget)
    
    return offtargets

def getOffTargets(sequence, bowtiePath, indexPath, exons, genes, coreMismatches, coreRange, totalMismatches, pamType) :
    
    if pamType == 'NRG' or pamType == 'NGG':
        offtargets = getOffTargetsNGG(sequence, bowtiePath, indexPath, exons, genes, coreMismatches, coreRange, totalMismatches, "CCN")   
        if pamType == 'NRG': 
            offtargets = offtargets + getOffTargetsNGG(sequence, bowtiePath, indexPath, exons, genes, coreMismatches, coreRange, totalMismatches, "CTN")
    elif pamType == 'TTTN':
        offtargets = getOffTargetsTTTN(sequence, bowtiePath, indexPath, exons, genes, totalMismatches, pamType)
    else:
        offtargets = getOffTargetsOtherPAMs(sequence, bowtiePath, indexPath, exons, genes, totalMismatches, pamType)
    
    offtargets.sort(key=lambda offtarget: (offtarget.score, offtarget.distance))
    
    return offtargets

class Offtarget:
    # New offtarget site coming for forward search, TTTN
    def __newFwd(self, chromosome, strand, start, substitutions, sequence, lengthSeq, lengthPAM, coreRange):
        self.chromosome = chromosome
        self.strand = strand
        if strand == "+":  # the search is done with the forward sequence!
            self.sequence = list(sequence)  # to make the string modifiable
        else:
            self.sequence = list(reverse_complement(sequence))  # to make the string modifiable
        self.start = start  # assuming bed coordinates
        self.end = start + lengthSeq
        
        tmp = substitutions.split(",")
        
        self.mismatches = len(tmp) - 1
        self.alignment = ['|'] * (lengthSeq - lengthPAM)
        self.score = 0
        for substitution in tmp:
            [idx, nt] = substitution.split(':')
            idx = int(idx)
            if strand == "+":
                self.sequence[idx] = nt[0]
            else:
                self.sequence[idx] = reverse_complement(nt[0])
            if idx < lengthPAM:  # The mismatch in the PAM is not considered for score calculation of alignment 
                continue
            self.score = self.score + pow(1.2, idx - lengthPAM + 1)
            self.alignment[idx - lengthPAM] = '-'
        if coreRange > 0 and coreRange != "NA":
            self.alignment = "PAM[" + "".join(self.alignment[:coreRange]) + "]" + "".join(self.alignment[coreRange:])
        else:
            self.alignment = "PAM" + "".join(self.alignment)
        self.sequence = "".join(self.sequence)
        
    # New offtarget site coming for reverse search, NGG and other PAMs
    def __newRev(self, chromosome, strand, start, substitutions, sequence, lengthSeq, lengthPAM, coreRange):
        self.chromosome = chromosome
        if strand == "+":  # the search is done with the reverse complemented sequence!
            self.strand = "-"
            self.sequence = list(sequence)  # to make the string modifiable
        else:
            self.strand = "+"
            self.sequence = list(reverse_complement(sequence))  # to make the string modifiable
        self.start = start  # assuming bed coordinates
        self.end = start + lengthSeq
        
        tmp = substitutions.split(",")
        
        self.mismatches = len(tmp) - 1
        self.alignment = ['|'] * (lengthSeq - lengthPAM)
        self.score = 0
        for substitution in tmp:
            [idx, nt] = substitution.split(':')
            idx = int(idx)
            if strand == "+":
                self.sequence[idx] = nt[0]
            else:
                self.sequence[idx] = reverse_complement(nt[0])
            if idx < lengthPAM:  # The mismatch in the PAM is not considered for score calculation of alignment 
                continue
            self.score = self.score + pow(1.2, lengthSeq - idx)
            self.alignment[lengthSeq - 1 - idx] = '-'
        if coreRange > 0 and coreRange != "NA":
            self.alignment = "".join(self.alignment[:-coreRange]) + "[" + "".join(self.alignment[-coreRange:]) + "]PAM"
        else:
            self.alignment = "".join(self.alignment) + "PAM"
        self.sequence = reverse_complement("".join(self.sequence))
        
        
    def __init__(self, forward, chromosome, strand, start, substitutions, sequence, lengthSeq, lengthPAM, coreRange):
        if(forward):
            self.__newFwd(chromosome, strand, start, substitutions, sequence, lengthSeq, lengthPAM, coreRange)
        else:
            self.__newRev(chromosome, strand, start, substitutions, sequence, lengthSeq, lengthPAM, coreRange)
       
    def setGeneInfo(self, exons, genes):
        closest = exons.closest(self.chromosome, self.start, self.end)
        
        self.geneID = closest[0]
        self.geneName = closest[1]
        self.distance = closest[2]
        self.intragenic = genes.overlaps(self.chromosome, self.start, self.end)
            
    def getGenomicCoordinates(self):
        return [self.chromosome, str(self.start + 1), str(self.end)]
    def getBedCoordinates(self):
        return [self.chromosome, str(self.start), str(self.end)]

########################################################################
# CRISPRater score calculation
########################################################################
model_weight = [0.14177385, 0.06966514, 0.04216254, 0.03303432, 0.02355430, -0.04746424, -0.04878001, -0.06981921, -0.07087756, -0.08160700]
model_offset = 0.6505037

patternCG = re.compile("[CG]")
def getGCFreq(seq):
    cg = len(patternCG.findall(seq))
    return(float(cg)/len(seq))

def calcFeatures(seq):
    feat = [0]*10
    feat[0] = getGCFreq(seq[3:13])
    if(seq[19]=="G"):
        feat[1] = 1
    if(seq[2]=="T" or seq[2]=="A"):
        feat[2] = 1
    if(seq[11]=="G" or seq[11]=="A"):
        feat[3] = 1
    if(seq[5]=="G"):
        feat[4] = 1
    if(seq[3]=="T" or seq[3]=="A"):
        feat[5] = 1
    if(seq[17]=="G" or seq[17]=="A"):
        feat[6] = 1
    if(seq[4]=="C" or seq[4]=="A"):
        feat[7] = 1
    if(seq[13]=="G"):
        feat[8] = 1
    if(seq[14]=="A"):
        feat[9] = 1
    return(feat)

def getScore(seq):
    features = calcFeatures(seq)
    
    score = 0
    for idx in range(0,len(features)):
        score = score + features[idx]*model_weight[idx]
    score = score + model_offset
    return(score)

def getScoreText(score):
    text = "{0:.2f}".format(score)
    if score < 0.56:
        return text + ' (LOW)' #low score -> red
    elif score > 0.74:
        return text + ' (HIGH)' #high score -> blue
    else:
        return text + ' (MEDIUM)' #medium score -> grey

########################################################################

class sgRNAbindingSite:    
    def __init__(self, targetSeq, sequence, position, strand, fwdPrimer, revPrimer):
        self.sequence = sequence
        if(len(targetSeq)==20):
            self.effi_score = getScore(targetSeq)
        else:
            self.effi_score = None
        self.position = position  # leftmost coordinates, including the PAM if 5'
        self.strand = strand
        self.score = None
        self.label = None
        self.oligo1 = ""  # leading GG, forward
        self.oligo2 = ""  # leading GG, reverse
        self.oligoAfwd = ""  # adding, forward
        self.oligoArev = ""  # adding, reverse
        self.oligoSfwd = ""  # substituting, forward
        self.oligoSrev = ""  # substituting, reverse
        # oligos
        if fwdPrimer == "TAGG":  # T7
            if targetSeq[0] == 'G' and targetSeq[1] == 'G':
                self.oligo1 = 'TA' + targetSeq
                self.oligo2 = 'AAAC' + reverse_complement(targetSeq[2:])
            elif targetSeq[0] == 'G' and not targetSeq[1] == 'G':
                self.oligoAfwd = 'TAg' + targetSeq
                self.oligoArev = 'AAAC' + reverse_complement(self.oligoAfwd[4:])
                self.oligoSfwd = 'TAGg' + targetSeq[2:]
                self.oligoSrev = 'AAAC' + reverse_complement(self.oligoSfwd[4:])
            else:
                self.oligoAfwd = 'TAgg' + targetSeq
                self.oligoArev = 'AAAC' + reverse_complement(self.oligoAfwd[4:])
                self.oligoSfwd = 'TAgg' + targetSeq[2:]
                self.oligoSrev = 'AAAC' + reverse_complement(self.oligoSfwd[4:])
        elif fwdPrimer == "CACCG":
            if targetSeq[0] == 'G':
                self.oligo1 = 'CACC' + targetSeq
                self.oligo2 = 'AAAC' + reverse_complement(targetSeq)
            else:
                self.oligoAfwd = 'CACCg' + targetSeq
                self.oligoArev = 'AAAC' + reverse_complement('G' + self.oligoAfwd[5:])
                self.oligoSfwd = 'CACCg' + targetSeq[1:]
                self.oligoSrev = 'AAAC' + reverse_complement('G' + self.oligoSfwd[5:])
        else:
            self.oligo1 = fwdPrimer + targetSeq
            self.oligo2 = revPrimer + reverse_complement(targetSeq)

    def addOffTargets(self, offTargets, coordinates, isThereGeneInfo):
        self.offTargets = offTargets
        # score computation
        # if the query is plasmid not score computation
        # the real target is not considered as an off-target for score computation
        # more off-targets => lower score
        # less mismatches => lower score
        # closer to exon => lower score
        averMismatches = 0
        totalMismatches = 0.0
        averDistance = 0
        for idx in range(len(offTargets)):
            offTarget = offTargets[idx]
            
            if not coordinates is None:
                # checking if the "off-target" site is in fact the on-target
                if offTarget.chromosome == coordinates[0] and offTarget.start >= coordinates[1] and offTarget.start <= coordinates[2]:
                    continue
            if offTarget.distance == 'NA' and isThereGeneInfo:
                continue
            totalMismatches = totalMismatches + 1
            averMismatches = averMismatches + offTarget.score
            
            if isThereGeneInfo:
                if offTarget.distance != 0:
                    try:
                        averDistance = averDistance + math.log10(offTarget.distance)
                    except ValueError:
                        print(offTarget.distance)
                else:
                    averDistance = averDistance + 0
            
        if totalMismatches > 0:
            self.score = averMismatches / totalMismatches + averDistance / totalMismatches - totalMismatches
        else:
            self.score = 100
                

            
class sgRNAbindingSites:
    def __init__(self):
        self.sites = []
    def add(self, targetSeq, sequence, position, strand, fwdPrimer, revPrimer):
        newSite = sgRNAbindingSite(targetSeq, sequence, position, strand, fwdPrimer, revPrimer)
        self.sites.append(newSite)

def getSeqCoords(seq, bowtiePath, indexPath):
    # We use bowtie to determine if the query sequence was extracted from the target genome
    if len(seq) < 500:
        # returns 0-index coordinates, bowtie uses 0-index
        bowtieOutput = os.popen(bowtiePath + os.path.sep + "bowtie " + indexPath + " --quiet -c " + seq, "r")
        
        line = bowtieOutput.readline()
        if not line: return None
        columns = line.split('\t')
        # [chromosome, start, end, strand]
        return [columns[2], int(columns[3]), int(columns[3]) + len(seq), columns[1]]
    else:
        # 5 prime
        bowtieOutput = os.popen(bowtiePath + os.path.sep + "bowtie " + indexPath + " --quiet -c " + seq[0:100], "r")
        line = bowtieOutput.readline()
        if not line: return None
        columns5 = line.split('\t')
        
        # 3 prime
        bowtieOutput = os.popen(bowtiePath + os.path.sep + "bowtie " + indexPath + " --quiet -c " + seq[-100:], "r")
        line = bowtieOutput.readline()
        if not line: return None
        columns3 = line.split('\t')
        
        if columns5[2] != columns3[2]:
            return None
        if columns5[1] != columns3[1]:
            return None
        if (int(columns3[3]) + 100 - int(columns5[3])) != len(seq):
            return None
        
        # [chromosome, start, end, strand]
        return [columns5[2], int(columns5[3]), int(columns3[3]) + 100, columns5[1]]

def getFormattedCoords(coords):
    return coords[0] + ":" + coords[1] + "-" + coords[2]
    
def getPlainOTPosition(distance, intragenic):
    if (distance == 0):
        return "Exonic"
    elif(intragenic):
        return "Intronic"
    else:
        return "Intergenic"

def addCandidateTargets(pam, target_size, sgRNA5, sgRNA3, query, strand, candidates, fwdPrimer, revPrimer):
    reg_exp = build_expression(pam)
    sgRNA5_re = '^' + build_expression(sgRNA5)
    sgRNA3_re = build_expression(sgRNA3) + '$'
    if pam == 'TTTN':
        indices = [m.start() for m in re.finditer('(?=' + reg_exp + ')', query, re.I)]
        for index in indices:
            if (index + target_size + len(pam)) > len(query):
                continue
            candidate_sequence = query[index + len(pam):index + len(pam) + target_size]
            pam_sequence = query[index:index + len(pam)]
            if (not re.search(sgRNA5_re, candidate_sequence) is None) and (not re.search(sgRNA3_re, candidate_sequence) is None):
                # we need to transform the index from the reversed sequence to the forward sequence
                if strand == '+':
                    candidates.add(candidate_sequence, pam_sequence + candidate_sequence, index, strand, fwdPrimer, revPrimer)
                else:
                    candidates.add(candidate_sequence, pam_sequence + candidate_sequence, len(query) - (index + target_size + len(pam)), strand, fwdPrimer, revPrimer)
    else:    
        indices = [m.start() for m in re.finditer('(?=' + reg_exp + ')', query, re.I)]
        for index in indices:
            if (index - target_size) < 0:
                continue
            candidate_sequence = query[index - target_size:index]
            pam_sequence = query[index:index + len(pam)]
            if (not re.search(sgRNA5_re, candidate_sequence) is None) and (not re.search(sgRNA3_re, candidate_sequence) is None):
                # we need to transform the index from the reversed sequence to the forward sequence
                if strand == '+':
                    candidates.add(candidate_sequence, candidate_sequence + pam_sequence, index - target_size, strand, fwdPrimer, revPrimer)
                else:
                    candidates.add(candidate_sequence, candidate_sequence + pam_sequence, len(query) - (index + len(pam)), strand, fwdPrimer, revPrimer)

def valid_dinucleotideIUPAC(string):
    validChars = ['A', 'C', 'G', 'T', 'N'] + list(iupac_code.keys())
    string = string.upper()
    if string != ''.join(c for c in string if c in validChars) or len(string) != 2:
        msg = "%r is not a valid dinucleotide sequence" % string
        raise argparse.ArgumentTypeError(msg)
    return string

def valid_overhang(string):
    validChars = ['A', 'C', 'G', 'T', 'N']
    string = string.upper()
    if string != ''.join(c for c in string if c in validChars) or len(string) > 5:
        msg = "%r is not a valid overhang sequence (up to 5 nt)" % string
        raise argparse.ArgumentTypeError(msg)
    return string

def doSearch(name, query, pamType, targetSize, totalMismatches, coreLength, coreMismatches, sgRNA5, sgRNA3, fwdPrimer, revPrimer, outputFolder, bowtiePath, indexPath, exonsFile, genesFile, maxOT):
     
    if pamType not in ["NGG", "NRG"]:
        coreLength = "NA"
        coreMismatches = "NA"
    totalSeqSize = args.targetSize + len(pamType) 
    
    # exons and genes
    exons = BedInterval()
    genes = BedInterval()
    if exonsFile is not None and genesFile is not None:
        try:
            from bx.intervals.intersection import Interval
            exons.loadFile(exonsFile)
            genes.loadFile(genesFile)
        except ImportError:
            sys.stderr.write('The bx-python module is not available. Ignoring exon and gene files!\n')
    
    coordinates = getSeqCoords(query, bowtiePath, indexPath)
    if not coordinates is None:
        # What is the input sequence is in the reverse strand???
        # so we use the reverse complement
        if coordinates[3] == "-":
            query = reverse_complement(query)
    
    candidates = sgRNAbindingSites()
    addCandidateTargets(pamType, targetSize, sgRNA5, sgRNA3, query, '+', candidates, fwdPrimer, revPrimer)
    addCandidateTargets(pamType, targetSize, sgRNA5, sgRNA3, reverse_complement(query), '-', candidates, fwdPrimer, revPrimer)
    
    
    if(len(candidates.sites) < 1):
        sys.stderr.write('No candidates found in the query sequence named %s.' % name)
        return
    
    maxScore = -1e10
    minScore = 1e10
    for idx in range(len(candidates.sites)):       
        ot = getOffTargets(candidates.sites[idx].sequence, bowtiePath, indexPath, exons, genes, coreMismatches, coreLength, totalMismatches, pamType)
        if maxOT < float("inf"):
            candidates.sites[idx].addOffTargets(ot[:maxOT], coordinates, len(exons.chroms) > 0)
        else:
            candidates.sites[idx].addOffTargets(ot, coordinates, len(exons.chroms) > 0)
        
        if candidates.sites[idx].score > maxScore:
            maxScore = candidates.sites[idx].score
        if candidates.sites[idx].score < minScore:
            minScore = candidates.sites[idx].score
        sys.stderr.write("Sequence %s: Done with candidate %i out of %i.\n" % (name, idx + 1, len(candidates.sites)))

    # scaling scores to the range [0, 1000]
    if len(candidates.sites) > 1:
        for idx in range(len(candidates.sites)):
            if maxScore > minScore:
                candidates.sites[idx].score = (candidates.sites[idx].score - minScore) / (maxScore - minScore) * 1000
            else:
                candidates.sites[idx].score = 1000
    elif len(candidates.sites) == 1:
        candidates.sites[0].score = 1000
    else:
        return
        
    # sorting candidates by score
    candidates.sites.sort(key=lambda site: (site.score), reverse=True)
    for idx in range(len(candidates.sites)):
        candidates.sites[idx].label = 'T' + str(idx + 1)
    
    # reporting
    bedFile = open(outputFolder + os.path.sep + name + '.bed', 'w')
    if coordinates is not None:
        for idx in range(len(candidates.sites)):
            bedFile.write(coordinates[0] + '\t' + str(coordinates[1] + candidates.sites[idx].position) + '\t' + str(coordinates[1] + candidates.sites[idx].position + totalSeqSize) + '\t' + candidates.sites[idx].label + '\t' + str(int(candidates.sites[idx].score)) + '\t' + candidates.sites[idx].strand + '\n')
    else:
        for idx in range(len(candidates.sites)):
            bedFile.write(name + '\t' + str(candidates.sites[idx].position) + '\t' + str(candidates.sites[idx].position + totalSeqSize) + '\t' + candidates.sites[idx].label + '\t' + str(int(candidates.sites[idx].score)) + '\t' + candidates.sites[idx].strand + '\n')
    bedFile.close()
    
    output = open(outputFolder + os.path.sep + name + '.xls', 'w')
    fasta = open(outputFolder + os.path.sep + name + '.fasta', 'w')
    
    output.write("Input:\t" + query + "\n")
    output.write("PAM:\t" + pamType + "\n")
    output.write("Target site length:\t" + str(targetSize) + "\n")
    output.write("Target site 5' limitation:\t" + sgRNA5 + "\n")
    output.write("Target site 3' limitation:\t" + sgRNA3 + "\n")
    output.write("Core length:\t" + str(coreLength) + "\n")
    output.write("Core MM:\t" + str(coreMismatches) + "\n")
    output.write("Total MM:\t" + str(totalMismatches) + "\n\n")    
    
    for idx in range(0, len(candidates.sites)):
        fasta.write('>' + candidates.sites[idx].label + '\n')
        fasta.write(candidates.sites[idx].sequence + '\n')
        
        if candidates.sites[idx].effi_score is None:
            output.write(candidates.sites[idx].label + '\t' + candidates.sites[idx].sequence + '\t' + str(int(candidates.sites[idx].score)) + '\n')
        else:
            output.write(candidates.sites[idx].label + '\t' + candidates.sites[idx].sequence + '\t' + str(int(candidates.sites[idx].score)) + '\tCRISPRater score\t' + getScoreText(candidates.sites[idx].effi_score) + '\n')
            
        if candidates.sites[idx].oligo1 != '':            
            output.write('Oligo fwd\t' + str(candidates.sites[idx].oligo1) + '\n')
            output.write('Oligo rev\t' + str(candidates.sites[idx].oligo2) + '\n')
        else:
            output.write('Oligo adding fwd\t' + str(candidates.sites[idx].oligoAfwd) + '\n')
            output.write('Oligo adding rev\t' + str(candidates.sites[idx].oligoArev) + '\n')
            if candidates.sites[idx].oligoSfwd != "" and candidates.sites[idx].oligoSrev != "":
                output.write('Oligo substituting fwd\t' + str(candidates.sites[idx].oligoSfwd) + '\n')
                output.write('Oligo substituting rev\t' + str(candidates.sites[idx].oligoSrev) + '\n')
        
        if(pamType == 'TTTN'):
            output.write('Chromosome\tstart\tend\tstrand\tMM\tPAM\ttarget_seq\talignment\tdistance\tposition\tgene name\tgene id\n')
        else:
            output.write('Chromosome\tstart\tend\tstrand\tMM\ttarget_seq\tPAM\talignment\tdistance\tposition\tgene name\tgene id\n')
        for idx2 in range(0, len(candidates.sites[idx].offTargets)):
            # in html output only top 20 offtarget sites
            offTarget = candidates.sites[idx].offTargets[idx2]
            
            output.write("\t".join(offTarget.getGenomicCoordinates()))
            output.write("\t" + offTarget.strand)
            output.write("\t" + str(offTarget.mismatches))
            if(pamType == 'TTTN'):
                output.write("\t" + offTarget.sequence[:len(pamType)] + "\t" + offTarget.sequence[len(pamType):])
            else:
                 output.write("\t" + offTarget.sequence[:-len(pamType)] + "\t" + offTarget.sequence[-len(pamType):])
            output.write("\t" + offTarget.alignment + "\t" + str(offTarget.distance) + "\t" + getPlainOTPosition(offTarget.distance, offTarget.intragenic))
            output.write("\t" + offTarget.geneName + "\t" + offTarget.geneID + "\n")
        output.write("\n")
        
    
    output.close()
    fasta.close()
        

if __name__ == "__main__":
    parser = argparse.ArgumentParser(formatter_class=argparse.RawTextHelpFormatter, description="CCTop is the CRISPR/Cas9 Target online predictor.", epilog=textwrap.dedent('''\
        If you use this tool please cite it as:
        
        Stemmer, M., Thumberger, T., del Sol Keyer, M., Wittbrodt, J. and Mateo, J.L.
        CCTop: an intuitive, flexible and reliable CRISPR/Cas9 target prediction tool.
        PLOS ONE (2015). doi:10.1371/journal.pone.0124633
        
        Have fun using CCTop!
        '''))
    parser.add_argument("--input", metavar="<file>", type=argparse.FileType('r'), help="Fasta file containing the sequence(s) to be scanned for sgRNA candidates.", required=True)
    parser.add_argument("--index", metavar="<file>" , help="Path to the bowtie index files including the name of the index.", required=True)
    parser.add_argument("--bowtie", metavar="<folder>", help="Path to the folder where the executable bowtie is.", default="." + os.path.sep)
    parser.add_argument("--targetSize", metavar="<int>", help="Target site length. (default: %(default)s)", default=20, type=int)
    parser.add_argument("--pam", help="PAM type. (default: %(default)s)", default="NGG", choices=['NGG', 'NRG', 'TTTN', 'NNGRRT', 'NNNNGATT', 'NNAGAAW', 'NAAAAC'])
    parser.add_argument("--sgRNA5", metavar="<sequence>", type=valid_dinucleotideIUPAC, help="Filter candidates target sites with the most 5 prime nucleotides defined by this sequence. IUPAC code allowed. (default: %(default)s)", default="NN")
    parser.add_argument("--sgRNA3", metavar="<sequence>", type=valid_dinucleotideIUPAC, help="Filter candidates target sites with the most 5 prime nucleotides defined by this sequence. IUPAC code allowed. (default: %(default)s)", default="NN")
    parser.add_argument("--fwdOverhang", metavar="<sequence>", type=valid_overhang, help="Sequence of the 5 prime forward cloning oligo. (default: %(default)s)", default="TAGG")
    parser.add_argument("--revOverhang", metavar="<sequence>", type=valid_overhang, help="Sequence of the 5 prime reverse cloning oligo. (default: %(default)s)", default="AAAC")
    parser.add_argument("--totalMM", metavar="<int>", help="Number of total maximum mismatches allowed in the off-target sites. (default: %(default)s)", default=4, type=int)
    parser.add_argument("--coreLength", metavar="<int>", help="Number of bases that enclose the core of the target site. (default: %(default)s)", default=12, type=int)
    parser.add_argument("--coreMM", metavar="<int>", help="Number of maximum mismatches allowed in the core of the off-target sites. (default: %(default)s)", default=2, type=int)
    parser.add_argument("--maxOT", metavar="<int>", help="Maximum number of off-target sites to be reported. (default: %(default)s)", default=float("inf"), type=int)
    parser.add_argument("--output", metavar="<folder>", help="Output folder. (default: %(default)s)", default="." + os.path.sep)
    parser.add_argument("--exonsFile", metavar="<file>", help="Path to the pseudo-bed file containing the coordinate of exons in the target genome. (default: NotUsed)", default=None)
    parser.add_argument("--genesFile", metavar="<file>", help="Path to the pseudo-bed file containing the coordinate of genes in the target genome. (default: NotUsed)", default=None)
    args = parser.parse_args()
    
    
    # check that the file is a proper multifasta file
    inputFile = args.input.read()
    args.input.close()
    inputFile = inputFile.replace("\r", "\n").replace("\n\n", "\n")
    lines = inputFile.split("\n")
    
    validChars = '-_.() abcdefghijklmnopqrstuvwxyzABCDEFGHIJKLMNOPQRSTUVWXYZ0123456789'
    
    fileContent = []
    seqLine = ""
    for line in lines:
        line = line.strip()
        if len(line) > 1 and line[0] == ">":  # header
            if seqLine != "":
                fileContent[len(fileContent) - 1].append(seqLine)
                name = ''.join(c for c in line if c in validChars)
                fileContent.append([name])
                seqLine = ""
            elif len(fileContent) == 0:  # first header
                name = ''.join(c for c in line if c in validChars)
                fileContent.append([name])
            else:  # wrong format
                sys.stderr.write("It looks that your input file is not (multi)fasta format. Please check it and try again.")
                sys.exit(1)
        elif re.match('[ACGTNacgtn]+', line) is not None:
            if fileContent == "":  # we find sequence without header
                sys.stderr.write("It looks that your input file is not (multi)fasta format. Please check it and try again.")
                sys.exit(1)
            else:
                seqLine = seqLine + line
        
    if seqLine != "":
        fileContent[len(fileContent) - 1].append(seqLine)
    else:
        sys.stderr.write("It looks that you file is not (multi)fasta format. Please check it and try again.")
        sys.exit(1)
    
    for sequence in fileContent:
        doSearch(sequence[0], sequence[1], args.pam, args.targetSize, args.totalMM, args.coreLength, args.coreMM, args.sgRNA5, args.sgRNA3, args.fwdOverhang, args.revOverhang, args.output, args.bowtie, args.index, args.exonsFile, args.genesFile, args.maxOT)
    
```
