## Supplementary_file_1 for "Bar-seq strategies for the LeishGEdit toolbox": LeishGEdit_CRISPR_search_v4.2.2.sh

```
#!/bin/bash
#-------------------------------------------------------------------------------------------------------------------------------------------------------------------------------------------------------------------------
#README:
#This script was written by Tom Beneke and is designed to pipe bed ORF files from any genome into CCTop's sgRNA design tool (Stemmer et al., PLoS One 2015) and output LeishGEdit suitable primers.

#Dependencies:
#This script uses Bowtie, Bedtools, gawk and CCTop standalone script
#install Bedtools with: $ sudo apt install bedtools
#install Bowtie with: $ sudo apt install bowtie
#install gawk with: $ sudo apt install gawk
#CCTop doesn't need to be installed, just copy bedInterval.py and CCTop.py into the respective directory

#Instructions on usage:
#Place this script "LeishGEdit_CRISPR_search_v4.2.sh", bedInterval.py and CCTop.py in a single folder together with:
#(1) your reference genome file (chromosomes or contigs in single file fasta format).
#(2) your ORF bed file (file format per line: "LmxM.01	201	2220	ID=LmxM.01.0030	1	-"). Your file should be tab-delimted and 6-columned. OTHER FORMATS ARE NOT ALLOWED!
#Navigate to your directory and execute the script by "sh LeishGEdit_CRISPR_search_v4.2.sh"

#Output:
#You can chose between three output formats:
#(1) Design primers for pT and pPLOT vectors
#(2) Design primers for pPOT vectors
#(3) Design primers for both vector systems
#Depending on the chosen output vector system or systems LeishGEdit will create one or two .csv files with all primers for your reference genome (termed: "reference-genome"pPOT_primers.csv or "reference-genome"pT-pPLOT_primers.csv).
#These files will contain the LeishGEdit primers required to target your gene of interest in the upstream or downstream region. The script output will also provide: 
#(1) The number of sgRNA off-target counts (1-4 MM) (determined by CCTop).
#(2) The number of sgRNA perfect matchtes (0 MM) (determined independently from CCTop).
#The file "All_primers_counts.csv" contains all homology arm and sgRNA target sequences without additional sequences (if you would like to use other plasmid or sgRNA systems).
#Files are in tab-delimited format!
#Additionally, you will find two folders that contain informations for all scored and ranked sgRNAs found in one 130nt search window (termed: 3_off-target_sgRNA_files and 5_off-target_sgRNA_files).
#For computing reasons these sgRNA ranking files are not named by GeneIDs, although each file corresponds to one GeneID. However, you can find the sgRNA corresponding search window coordinates in the ".Csg.bed" and ".Nsg.bed" files that have also been created.

#Testing the script on your system
#To test the script we have provided two test files:
#(1) A ORF bed test file ("script-test-bed-file.bed")
#(2) A genome reference test file ("script-test-genome-reference-file.fasta")
#Testing the script with these two files will take approximately 10 minutes and allows you to see whether the script is working before starting to analyse larger genome files. 
#You will also see if you get any error messages on your installed version of Bowtie, Bedtools, gawk or CCTop.

#Program start
#-------------------------------------------------------------------------------------------------------------------------------------------------------------------------------------------------------------------------

echo "
.o+++++++++++++++++++++++++++++++++++++++++++++++++++++++++++++++++++++++++++++++++++++++++o. 
-h-        .::::::::..                                                                    h(.
-h-    .:::-........-:::::::--.                                                           h(.
-h-   ::......................-:::::-.                                                    h(.
-h-  ::..............................-::::-.                                              h(.
-h-   .+...............::((((:::...........-::::.                                         h(.
-h-    .(:............+:-------:))......-:......-:::.                                     h(.
-h-      .(:..........+:---------:)...:(((.-......-.-::-                                  h(.
-h-         -::........((:------:)-..:(:(..):-.   ......(-                                h(.
-h-            -:::......:::((:::....+:+...........-. :(:o.                               h(.
-h-                -::::.............:(+.(.......-....:::-                                h(.
-h-                    .-::::-........-...........-..::-.-::::-.                          h(.
-h-                          .-:::::-......-.......::::(-::::..-::::.                     h(.
-h-                                 .--:::::-......::::.     .::::-.-:::-.                h(.
-h-                                           -----               ::..:::-                h(.
-h-                                                                      :-..(:.          h(.
-h-  :h(            uu        dm         zyhhhy. hhyyyy       m+ uu  ym     .(:.-(.       h(.
-h-  (m+     eezu       .(+(. dm        my       sm           m+    dzkd       .+-.::     h(.
-h-  (m+    um  mm  hm  my--. dmmnsm+  dm  oooo  smhyyy  myzzym+ om  sm          (:.-(    h(.
-h-  (m+    hmsss   hm  (sdh- dm   ms  ym     mo sm     om    m+ om  om           (-.-(   h(.
-h-  (myooo  doooo  hm  o(om+ dm   ms   omho+smo smsooo  ddhuhm+ om  my            +..o.  h(.
-h-                                                                               .+.+.   h(.
-h-                                                           --:.   (oo:oo:+(::--s((.    h(.
-h-                                                  ...    :s+-o+yoo+ss.os.+o:os(.       h(.
-h-                                       .        (+oo+ys(o+ssos-(ss .-:(+::((-          h(.
-h-                                 -::-    -+:((-+oo:oo(oo.   ..---                      h(.
-h-                                    .-:+sss.so+::(o(o+-                                h(.
-h-                                      : .:((so.                                        h(.
-d++++++++++++++++++++++++++++++++++++++++++++++++++++++++++++++++++++++++++++++++++++++++d(.
"
echo "
                       Welccome to LeishGEdit primer design version 4.2!"
echo "
     This script was written by Tom Beneke. Please cite Beneke et al., 2017 and Beneke and Gluenz, 2020 when using it!" 
echo "
     Please make sure you have bowtie, bedtools, gawk and python higher than 2.7 installed!"
echo "
              [1] Type in 1 to run LeishGEdit v4.2 pipeline for pT and pPLOT vectors"
echo "
              [2] Type in 2 to run LeishGEdit v4.2 pipeline for pPOT vectors"
echo "
              [3] Type in 3 to run LeishGEdit v4.2 pipeline for both vector systems"
echo "
              [4] Type in 4 to exit LeishGEdit v4.2"
echo "
--------------------------------------------------------------------------------------------------"


pause(){
   read -p "Press [Enter] key to continue..." fackEnterKey
}

pauseRUN(){
   read -p "Press [Enter] key to start primer design with LeishGEdit v4.2..." fackEnterKey
}

read -p "Please Select An Option: " vector
if [ $vector -eq 1 ]; then
	echo "You have selected design for pT and pPLOT vectors!"	
	pause
elif [ $vector -eq 2 ]; then
	echo "You have selected design for pPOT vectors!"	
	pause
elif [ $vector -eq 3 ]; then
	echo "You have selected design for pT, pPLOT and pPOT vectors!"	
	pause
else
	echo  "
________________________________________________________
< If you exit LeishGEdit this cow will be slaughtered! >
<                  Please don't go!                    >
 -------------------------------------------------------
             \   ^__^
              \  (oo)\_______
                 (__)\       )---
                     ||----w |
                     ||     ||
	"
    	pause
	echo "

________________________________________________________
<              You just killed this cow...             >
 -------------------------------------------------------
                      \_______
                ^__^  \       )---
                (##)  ||----w |
                (__)  ||     ||
                     
                     
	"
	exit 1
fi


#-------------------------------------------------------------------------------------------------------------------------------------------------------------------------------------------------------------------------


#Input files
#-------------------------------------------------------------------------------------------------------------------------------------------------------------------------------------------------------------------------
read -p 'Please enter your reference genome (chromosomes or contigs in single file fasta format): ' REFERENCE_GENOME
	echo "You are uploading <"$REFERENCE_GENOME"> as a reference genome."
	pause
read -p 'Please enter your ORF bed (file format per line: "LmxMC9T7.01.1	201	2220	ID=LmxM.01.0030	1	-"). 
Your file should be tab-delimted and 6-columned. OTHER FORMATS ARE NOT ALLOWED! Your upload: ' BED_ORF_INPUT
	echo "You are uploading <"$BED_ORF_INPUT"> as your ORF bed file."
	pause
	if perl -ne '(s/\t//go)==5 or exit -1' $BED_ORF_INPUT
	then
	    	echo "Your bed ORF input has the corrected format" 
	else
	    	echo "Your bed ORF input is not tab-delimited or 6-columned"
		exit
	fi
read -p 'Please enter your path to bowtie. You can get the path path to bowtie by command "type bowtie" (on most systems it will be: "/usr/bin/"): ' BOWTIE_LOCATION
	echo "Your bowtie location is: <"$BOWTIE_LOCATION"> "
	pauseRUN
echo "Thanks for your input. Your genome reference file and bed ORF will be now processed! This might take a while...."
#-------------------------------------------------------------------------------------------------------------------------------------------------------------------------------------------------------------------------


#check for dependencies
#-------------------------------------------------------------------------------------------------------------------------------------------------------------------------------------------------------------------------
if [ ! -f bedInterval.py ]; then
    	echo "bedInterval.py file not found!"
	exit 1
fi

if [ ! -f CCTop.py ]; then
    	echo "CCTop.py file not found!"
	exit 1
fi
#-------------------------------------------------------------------------------------------------------------------------------------------------------------------------------------------------------------------------
#Clean up possible windows formating issues on bed file and genome file
sed -i -e 's/\r$//' $BED_ORF_INPUT
sed -i -e 's/\r$//' $REFERENCE_GENOME

WORKING_GENOME=working_genome_sgl.fasta

#Making fasta genome file single line file
awk '/^>/ {printf("\n%s\n",$0);next; } { printf("%s",$0);}  END {printf("\n");}' $REFERENCE_GENOME | tail -n +2 > $WORKING_GENOME


#Index reference genome
echo "Starting with indexing the Genome!"
$BOWTIE_LOCATION"bowtie-build" -r -f $WORKING_GENOME $WORKING_GENOME


#-------------------------------------------------------------------------------------------------------------------------------------------------------------------------------------------------------------------------
#Extract search regions
#create file folders for buffer
echo "Creating buffer folders for bedtools calculations"
mkdir ORFinput
mkdir Nsgbuffer
mkdir Nhombuffer
mkdir Csgbuffer
mkdir Chombuffer
mkdir Bedtools

#Modify ORF input file for vlookups later on
awk '{printf "%s\t%s\n", NR,$0}' $BED_ORF_INPUT > ORFinput/Genome-ORF.bed.2.csv
awk 'BEGIN { OFS = "\t" } {($5 = "ABCDEFG"$1"HIJKLM"$5"OPQRST")} {$1="";sub("\t","")} 1' ORFinput/Genome-ORF.bed.2.csv > ORFinput/Genome-ORF.bed.3.csv
sed -e 's/\(HIJKLM\).*\(OPQRST\)/\1\2/' ORFinput/Genome-ORF.bed.3.csv > Genome-ORF.bed.input.csv

#Nsg window
awk 'BEGIN { OFS = "\t" } {if ($6== "+") ($3 = $2 + 0)} 1' Genome-ORF.bed.input.csv > Nsgbuffer/Genome-ORF.Nsg1.bed
awk 'BEGIN { OFS = "\t" } {if ($6== "+") ($2 = $2 - 130)} 1' Nsgbuffer/Genome-ORF.Nsg1.bed > Nsgbuffer/Genome-ORF.Nsg2.bed
awk 'BEGIN { OFS = "\t" } {if ($6== "-") ($2 = $3 + 130)} 1' Nsgbuffer/Genome-ORF.Nsg2.bed > Nsgbuffer/Genome-ORF.Nsg3.bed
awk 'BEGIN { OFS = "\t" } {if ($6== "-") ($3 = $3 - 0)} 1' Nsgbuffer/Genome-ORF.Nsg3.bed > Nsgbuffer/Genome-ORF.Nsg4.bed
awk '$2 > $3 { temp = $3; $3 = $2; $2 = temp } 1' OFS='\t' Nsgbuffer/Genome-ORF.Nsg4.bed > Nsgbuffer/Genome-ORF.5Nsg.bed
awk '$2 > 0' Nsgbuffer/Genome-ORF.5Nsg.bed > Genome-ORF.Nsg.bed


#Nhom window
awk 'BEGIN { OFS = "\t" } {if ($6== "+") ($3 = $2 + 30)} 1' Genome-ORF.bed.input.csv > Nhombuffer/Genome-ORF.Nhom1.bed
awk 'BEGIN { OFS = "\t" } {if ($6== "+") ($2 = $2 - 160)} 1' Nhombuffer/Genome-ORF.Nhom1.bed > Nhombuffer/Genome-ORF.Nhom2.bed
awk 'BEGIN { OFS = "\t" } {if ($6== "-") ($2 = $3 + 160)} 1' Nhombuffer/Genome-ORF.Nhom2.bed > Nhombuffer/Genome-ORF.Nhom3.bed
awk 'BEGIN { OFS = "\t" } {if ($6== "-") ($3 = $3 - 30)} 1' Nhombuffer/Genome-ORF.Nhom3.bed > Nhombuffer/Genome-ORF.Nhom4.bed
awk '$2 > $3 { temp = $3; $3 = $2; $2 = temp } 1' OFS='\t' Nhombuffer/Genome-ORF.Nhom4.bed > Nhombuffer/Genome-ORF.5Nhom.bed
awk '$2 > 0' Nhombuffer/Genome-ORF.5Nhom.bed > Genome-ORF.Nhom.bed


#Csg window
awk 'BEGIN { OFS = "\t" } {if ($6== "+") ($2 = $3 - 3)} 1' Genome-ORF.bed.input.csv > Csgbuffer/Genome-ORF.Csg1.bed
awk 'BEGIN { OFS = "\t" } {if ($6== "+") ($3 = $3 + 127)} 1' Csgbuffer/Genome-ORF.Csg1.bed > Csgbuffer/Genome-ORF.Csg2.bed
awk 'BEGIN { OFS = "\t" } {if ($6== "-") ($3 = $2 - 127)} 1' Csgbuffer/Genome-ORF.Csg2.bed > Csgbuffer/Genome-ORF.Csg3.bed
awk 'BEGIN { OFS = "\t" } {if ($6== "-") ($2 = $2 + 3)} 1' Csgbuffer/Genome-ORF.Csg3.bed > Csgbuffer/Genome-ORF.Csg4.bed
awk '$2 > $3 { temp = $3; $3 = $2; $2 = temp } 1' OFS='\t' Csgbuffer/Genome-ORF.Csg4.bed > Csgbuffer/Genome-ORF.5Csg.bed
awk '$2 > 0' Csgbuffer/Genome-ORF.5Csg.bed > Genome-ORF.Csg.bed


#Chom window
awk 'BEGIN { OFS = "\t" } {if ($6== "+") ($2 = $3 - 33)} 1' Genome-ORF.bed.input.csv > Chombuffer/Genome-ORF.Chom1.bed
awk 'BEGIN { OFS = "\t" } {if ($6== "+") ($3 = $3 + 157)} 1' Chombuffer/Genome-ORF.Chom1.bed > Chombuffer/Genome-ORF.Chom2.bed
awk 'BEGIN { OFS = "\t" } {if ($6== "-") ($3 = $2 - 157)} 1' Chombuffer/Genome-ORF.Chom2.bed > Chombuffer/Genome-ORF.Chom3.bed
awk 'BEGIN { OFS = "\t" } {if ($6== "-") ($2 = $2 + 33)} 1' Chombuffer/Genome-ORF.Chom3.bed > Chombuffer/Genome-ORF.Chom4.bed
awk '$2 > $3 { temp = $3; $3 = $2; $2 = temp } 1' OFS='\t' Chombuffer/Genome-ORF.Chom4.bed > Chombuffer/Genome-ORF.5Chom.bed
awk '$2 > 0' Chombuffer/Genome-ORF.5Chom.bed > Genome-ORF.Chom.bed


#empty buffer
echo "Deleting buffers"
rm -r Nsgbuffer
rm -r Nhombuffer
rm -r Csgbuffer
rm -r Chombuffer

#extract N sgRNA design windows
bedtools getfasta -fi $WORKING_GENOME -bed Genome-ORF.Nsg.bed -s -name > Bedtools/NsgRNA.fasta

#extract C sgRNA design windows
bedtools getfasta -fi $WORKING_GENOME -bed Genome-ORF.Nhom.bed -s -name > Bedtools/Nhom.fasta

#extract N homology design windows
bedtools getfasta -fi $WORKING_GENOME -bed Genome-ORF.Csg.bed -s -name > Bedtools/CsgRNA.fasta

#extract C homology design windows
bedtools getfasta -fi $WORKING_GENOME -bed Genome-ORF.Chom.bed -s -name > Bedtools/Chom.fasta

echo "Bedtools sequence cut-extraction done"

#-------------------------------------------------------------------------------------------------------------------------------------------------------------------------------------------------------------------------
#sgRNA search

#NsgRNA search
echo "Starting 5'sgRNAs search. This might take a while..."
#run sgRNA search on NsgRNA window
mkdir NsgRNA.search
python CCTop.py --input Bedtools/NsgRNA.fasta --index $WORKING_GENOME --bowtie $BOWTIE_LOCATION --output NsgRNA.search

echo "Finished 5'sgRNAs search. Summarizing now the off-target counts for 5'sgRNAs!"
#count off-targets NsgRNA
cd NsgRNA.search
mkdir buffer
mkdir targetcount
for f in ./*.xls; do
	grep -v 'Oligo' "$f" > buffer/"$f".1.xls
	grep -v 'Chromosome' buffer/"$f".1.xls > targetcount/"$f".tc.xls
	echo 	"$f""\t""$(awk '$1 == "T1" {print $2}' "targetcount/"$f".tc.xls")\t$((($(sed -s -n '/^T1/,/^T2/p;/^T2/q' "targetcount/"$f".tc.xls" | wc -l)) - 4))" >> offtargetcounts.txt
done
cd ..

#CsgRNA search
echo "Starting 3'sgRNAs search. This might take a while..."
#run sgRNA search on CsgRNA window
mkdir CsgRNA.search
python CCTop.py --input Bedtools/CsgRNA.fasta --index $WORKING_GENOME --bowtie $BOWTIE_LOCATION --output CsgRNA.search

echo "Finished 3'sgRNAs search. Summarizing now the off-target counts for 3'sgRNAs!"
#count off-targets NsgRNA
cd CsgRNA.search
mkdir buffer
mkdir targetcount
for f in ./*.xls; do
	grep -v 'Oligo' "$f" > buffer/"$f".1.xls
	grep -v 'Chromosome' buffer/"$f".1.xls > targetcount/"$f".tc.xls
	echo 	"$f""\t""$(awk '$1 == "T1" {print $2}' "targetcount/"$f".tc.xls")\t$((($(sed -s -n '/^T1/,/^T2/p;/^T2/q' "targetcount/"$f".tc.xls" | wc -l)) - 4))" >> offtargetcounts.txt
done
cd ..


#-------------------------------------------------------------------------------------------------------------------------------------------------------------------------------------------------------------------------
#Run sgRNAs back against genome to define HF regions and count for on-targets

#-------------------------------------------------------------------------------------------------------------------------------------------------------------------------------------------------------------------------
#Start 5'sgRNA processing
echo "Start 5'sgRNA processing"

#for vlookup: clean up CCTop output, NsgRNA input, Nhom input
sed -e 's/\(HIJKLMOPQRST\).*\(.xls\)/\1\2/' NsgRNA.search/offtargetcounts.txt > NsgRNA.search/offtargetcounts.2.txt
awk '{ gsub("./","",$1); gsub(".xls","",$1); print}' OFS='\t' NsgRNA.search/offtargetcounts.2.txt > NsgRNA.search/offtargetcounts.3.txt
sed -e 's/HIJKLMOPQRST.*$/HIJKLMOPQRST/' Bedtools/NsgRNA.fasta > NsgRNA.search/NsgRNA.1.txt
sed -e 's/HIJKLMOPQRST.*$/HIJKLMOPQRST/' Bedtools/Nhom.fasta > NsgRNA.search/Nhom.1.txt

#transform NsgRNA and Nhom fasta input file into column file to match with CCTop output file
awk 'BEGIN{RS=">";OFS="\t"}NR>1{print $1,$2}' NsgRNA.search/NsgRNA.1.txt > NsgRNA.search/NsgRNA.2.txt
awk 'BEGIN{RS=">";OFS="\t"}NR>1{print $1,$2}' NsgRNA.search/Nhom.1.txt > NsgRNA.search/Nhom.2.txt

#Merge CCTop output, NsgRNA input, Nhom input
awk 'NR == FNR{a[$1] = $0;next}; {print $0, $1 in a?a[$1]: "@@@@NA@@@@"}' OFS='\t' NsgRNA.search/offtargetcounts.3.txt NsgRNA.search/NsgRNA.2.txt > NsgRNA.search/merge1.csv
awk -F'[[:space:]]' '!$4{ $4="@@@@NA@@@@"}; !$5{ $5="@@@@NA@@@@"}1' OFS='\t' NsgRNA.search/merge1.csv > NsgRNA.search/merge2.csv
awk 'NR == FNR{a[$1] = $0;next}; {print $0, $1 in a?a[$1]: "@@@@NA@@@@"}' OFS='\t' NsgRNA.search/Nhom.2.txt NsgRNA.search/merge2.csv > NsgRNA.search/merge3.csv
awk -F'[[:space:]]' '!$6{ $6="@@@@NA@@@@"}; !$7{ $7="@@@@NA@@@@"}1' OFS='\t' NsgRNA.search/merge3.csv > NsgRNA.search/merge4.csv

#reverse complement sgRNAs for match-up with homology arms
cat NsgRNA.search/merge4.csv | awk '{print $4}' | rev | tr "[ATGC]" "[TACG]" > NsgRNA.search/NsgRNAs_reverse.csv
paste -d'\t' NsgRNA.search/merge4.csv NsgRNA.search/NsgRNAs_reverse.csv > NsgRNA.search/merge5.csv
awk 'BEGIN { OFS = "\t" } {$0=gensub(/\s*\S+/,"",3)} {$0=gensub(/\s*\S+/,"",5)} { temp = $3; $3 = $5; $5 = temp } {gsub("@@@@NA@@@@","No_sgRNA_found",$5) && gsub("@@@@TN@@@@","No_reverse_sgRNA",$6)} 1' NsgRNA.search/merge5.csv > NsgRNA.search/merge6.csv

#match sgRNA to define specific homology arms
echo "Matching 5' sgRNAs to define upstream homology arms"
awk 'BEGIN { OFS = "\t" } {IGNORECASE=1; ret = index($3, $5); printf ret "\t"} 1' NsgRNA.search/merge6.csv > NsgRNA.search/NsgRNAvshom.csv
awk 'BEGIN { OFS = "\t" } {IGNORECASE=1; ret = index($4, $7); printf ret "\t"} 1' NsgRNA.search/NsgRNAvshom.csv > NsgRNA.search/NsgRNAvshom.2.csv
awk '$1 > 0 || $2 > 0' NsgRNA.search/NsgRNAvshom.2.csv > NsgRNA.search/NsgRNAvshom.2.1.csv
awk 'BEGIN { OFS = "\t" } {if ($1>$2) ($2 = $1); else ($1 = $2)} {$0=gensub(/\s*\S+/,"",2)} {$1 = $1 -30} {printf substr($4,$1,30)"\t"} 1' NsgRNA.search/NsgRNAvshom.2.1.csv > NsgRNA.search/NsgRNAvshom.3.csv
awk '$2 > 0' NsgRNA.search/NsgRNAvshom.3.csv > NsgRNA.search/NsgRNAvshom.3.1.csv
awk 'BEGIN { OFS = "\t" } {printf substr($5,161,30)"\t"} 1' NsgRNA.search/NsgRNAvshom.3.1.csv > NsgRNA.search/NsgRNAvshom.4.csv
cat NsgRNA.search/NsgRNAvshom.4.csv | awk '{print $1}' | rev | tr "[ATGC]" "[TACG]" > NsgRNA.search/Nupstream_reverse.csv
paste -d'\t' NsgRNA.search/Nupstream_reverse.csv NsgRNA.search/NsgRNAvshom.4.csv > NsgRNA.search/NsgRNAvshom.4.1.csv
awk 'BEGIN { OFS = "\t" } {$0=gensub(/\s*\S+/,"",2)} {$0=gensub(/\s*\S+/,"",3)} { temp = $1; $1 = $3; $3 = temp } 1' NsgRNA.search/NsgRNAvshom.4.1.csv > NsgRNA.search/NsgRNAvshom.5.csv

#Check for on target counts forward
echo "Checking on targets for 5' sgRNAs (forward)"
awk 'BEGIN {print "#!/bin/bash"; print "mkdir NsgRNA.search/NsgRNA_count_f"}' > NsgRNA.search/NsgRNAs_genome_count.sh
echo "WORKING_GENOME="$WORKING_GENOME >> NsgRNA.search/NsgRNAs_genome_count.sh
cat NsgRNA.search/NsgRNAvshom.5.csv | awk '{print "grep -i -c "$7" $WORKING_GENOME > NsgRNA.search/NsgRNA_count_f/"$1".txt"}' >> NsgRNA.search/NsgRNAs_genome_count.sh
sh NsgRNA.search/NsgRNAs_genome_count.sh
awk '{print $0, FILENAME}' NsgRNA.search/NsgRNA_count_f/*.txt > NsgRNA.search/NsgRNA_count_f.txt
rm -r NsgRNA.search/NsgRNA_count_f
{
sed -e's/[[:space:]]\{1,\}/\t/g' NsgRNA.search/NsgRNA_count_f.txt
} > NsgRNA.search/NsgRNA_count_f.csv
awk 'BEGIN { OFS = "\t" } {gsub("NsgRNA.search/NsgRNA_count_f/","",$2)} {gsub(".txt","",$2)} { temp = $1; $1 = $2; $2 = temp } 1' NsgRNA.search/NsgRNA_count_f.csv > NsgRNA.search/NsgRNA_count_f.2.csv
awk 'NR == FNR{a[$1] = $0;next}; {print $0, $1 in a?a[$1]: "@@@@NA@@@@"}' OFS='\t' NsgRNA.search/NsgRNA_count_f.2.csv NsgRNA.search/NsgRNAvshom.5.csv > NsgRNA.search/pre-final-Ntagging.csv

#Check for on target counts reverse
echo "Checking on targets for 5' sgRNAs (reverse)"
awk 'BEGIN {print "#!/bin/bash"; print "mkdir NsgRNA.search/NsgRNA_count_r"}' > NsgRNA.search/NsgRNAs_genome_count.sh
echo "WORKING_GENOME="$WORKING_GENOME >> NsgRNA.search/NsgRNAs_genome_count.sh
cat NsgRNA.search/NsgRNAvshom.5.csv | awk '{print "grep -i -c "$8" $WORKING_GENOME > NsgRNA.search/NsgRNA_count_r/"$1".txt"}' >> NsgRNA.search/NsgRNAs_genome_count.sh
sh NsgRNA.search/NsgRNAs_genome_count.sh
awk '{print $0, FILENAME}' NsgRNA.search/NsgRNA_count_r/*.txt > NsgRNA.search/NsgRNA_count_r.txt
rm -r NsgRNA.search/NsgRNA_count_r
{
sed -e's/[[:space:]]\{1,\}/\t/g' NsgRNA.search/NsgRNA_count_r.txt
} > NsgRNA.search/NsgRNA_count_r.csv
awk 'BEGIN { OFS = "\t" } {gsub("NsgRNA.search/NsgRNA_count_r/","",$2)} {gsub(".txt","",$2)} { temp = $1; $1 = $2; $2 = temp } 1' NsgRNA.search/NsgRNA_count_r.csv > NsgRNA.search/NsgRNA_count_r.2.csv
awk 'NR == FNR{a[$1] = $0;next}; {print $0, $1 in a?a[$1]: "@@@@NA@@@@"}' OFS='\t' NsgRNA.search/NsgRNA_count_r.2.csv NsgRNA.search/pre-final-Ntagging.csv > NsgRNA.search/pre-final-Ntagging.2.csv
echo "Preparing for final outcome 5'sgRNA"

#Clean-up outcome and add original names (NA should be for all sgRNA that have not been found compared to input)
awk 'BEGIN { OFS = "\t" } {if ($10>$12) ($12 = $10); else ($10 = $12)} {$0=gensub(/\s*\S+/,"",12)} {$0=gensub(/\s*\S+/,"",11)} {$0=gensub(/\s*\S+/,"",9)} {$0=gensub(/\s*\S+/,"",4)} {$0=gensub(/\s*\S+/,"",4)} { temp = $4; $4 = $5; $5 = temp } {$0=gensub(/\s*\S+/,"",6)} { if($6 == 0) ($6 = "@@@@NA@@@@")} 1' NsgRNA.search/pre-final-Ntagging.2.csv > NsgRNA.search/pre-final-Ntagging.3.csv
awk 'BEGIN { OFS = "\t" } {($1 = "ABCDEFG"$1"HIJKLMOPQRST")} 1' ORFinput/Genome-ORF.bed.2.csv > ORFinput/Genome-ORF.bed.finalmatch.csv
awk 'NR == FNR{a[$1] = $0;next}; {print $0, $1 in a?a[$1]: "@@@@NA@@@@"}' OFS='\t' NsgRNA.search/pre-final-Ntagging.3.csv ORFinput/Genome-ORF.bed.finalmatch.csv > NsgRNA.search/pre-final-Ntagging.4.1.csv
awk -F'[[:space:]]' '!$9{ $9="@@@@NA@@@@"}; !$10{ $10="@@@@NA@@@@"}; !$11{ $11="@@@@NA@@@@"}; !$12{ $12="@@@@NA@@@@"}; !$13{ $13="@@@@NA@@@@"} 1' OFS='\t' NsgRNA.search/pre-final-Ntagging.4.1.csv > NsgRNA.search/pre-final-Ntagging.4.csv

echo "End 5'sgRNA processing"
#End 5'sgRNA processing
#-------------------------------------------------------------------------------------------------------------------------------------------------------------------------------------------------------------------------

#-------------------------------------------------------------------------------------------------------------------------------------------------------------------------------------------------------------------------
#Start 3'sgRNA processing
echo "Start 3'sgRNA processing"

#for vlookup: clean up CCTop output, CsgRNA input, Chom input
sed -e 's/\(HIJKLMOPQRST\).*\(.xls\)/\1\2/' CsgRNA.search/offtargetcounts.txt > CsgRNA.search/offtargetcounts.2.txt
awk '{ gsub("./","",$1); gsub(".xls","",$1); print}' OFS='\t' CsgRNA.search/offtargetcounts.2.txt > CsgRNA.search/offtargetcounts.3.txt
sed -e 's/HIJKLMOPQRST.*$/HIJKLMOPQRST/' Bedtools/CsgRNA.fasta > CsgRNA.search/CsgRNA.1.txt
sed -e 's/HIJKLMOPQRST.*$/HIJKLMOPQRST/' Bedtools/Chom.fasta > CsgRNA.search/Chom.1.txt

#transform CsgRNA and Chom fasta input file into column file to match with CCTop output file
awk 'BEGIN{RS=">";OFS="\t"}NR>1{print $1,$2}' CsgRNA.search/CsgRNA.1.txt > CsgRNA.search/CsgRNA.2.txt
awk 'BEGIN{RS=">";OFS="\t"}NR>1{print $1,$2}' CsgRNA.search/Chom.1.txt > CsgRNA.search/Chom.2.txt

#Merge CCTop output, CsgRNA input, Chom input
awk 'NR == FNR{a[$1] = $0;next}; {print $0, $1 in a?a[$1]: "@@@@NA@@@@"}' OFS='\t' CsgRNA.search/offtargetcounts.3.txt CsgRNA.search/CsgRNA.2.txt > CsgRNA.search/merge1.csv
awk -F'[[:space:]]' '!$4{ $4="@@@@NA@@@@"}; !$5{ $5="@@@@NA@@@@"}1' OFS='\t' CsgRNA.search/merge1.csv > CsgRNA.search/merge2.csv
awk 'NR == FNR{a[$1] = $0;next}; {print $0, $1 in a?a[$1]: "@@@@NA@@@@"}' OFS='\t' CsgRNA.search/Chom.2.txt CsgRNA.search/merge2.csv > CsgRNA.search/merge3.csv
awk -F'[[:space:]]' '!$6{ $6="@@@@NA@@@@"}; !$7{ $7="@@@@NA@@@@"}1' OFS='\t' CsgRNA.search/merge3.csv > CsgRNA.search/merge4.csv

#reverse complement sgRNAs for match-up with homology arms
cat CsgRNA.search/merge4.csv | awk '{print $4}' | rev | tr "[ATGC]" "[TACG]" > CsgRNA.search/CsgRNAs_reverse.csv
paste -d'\t' CsgRNA.search/merge4.csv CsgRNA.search/CsgRNAs_reverse.csv > CsgRNA.search/merge5.csv
awk 'BEGIN { OFS = "\t" } {$0=gensub(/\s*\S+/,"",3)} {$0=gensub(/\s*\S+/,"",5)} { temp = $3; $3 = $5; $5 = temp } {gsub("@@@@NA@@@@","No_sgRNA_found",$5) && gsub("@@@@TN@@@@","No_reverse_sgRNA",$6)} 1' CsgRNA.search/merge5.csv > CsgRNA.search/merge6.csv

#match sgRNA to define specific homology arms
echo "Matching 3' sgRNAs to define downstream homology arms"
awk 'BEGIN { OFS = "\t" } {IGNORECASE=1; ret = index($3, $5); printf ret "\t"} 1' CsgRNA.search/merge6.csv > CsgRNA.search/CsgRNAvshom.csv
awk 'BEGIN { OFS = "\t" } {IGNORECASE=1; ret = index($4, $7); printf ret "\t"} 1' CsgRNA.search/CsgRNAvshom.csv > CsgRNA.search/CsgRNAvshom.2.csv
awk '$1 > 0 || $2 > 0' CsgRNA.search/CsgRNAvshom.2.csv > CsgRNA.search/CsgRNAvshom.2.1.csv
awk 'BEGIN { OFS = "\t" } {if ($1>$2) ($2 = $1); else ($1 = $2)} {$0=gensub(/\s*\S+/,"",2)} {$1 = $1 +23} {printf substr($4,$1,30)"\t"} 1' CsgRNA.search/CsgRNAvshom.2.1.csv > CsgRNA.search/CsgRNAvshom.3.csv
awk '$2 < 162' CsgRNA.search/CsgRNAvshom.3.csv > CsgRNA.search/CsgRNAvshom.3.1.csv
awk 'BEGIN { OFS = "\t" } {printf substr($5,1,30)"\t"} 1' CsgRNA.search/CsgRNAvshom.3.1.csv > CsgRNA.search/CsgRNAvshom.4.csv
cat CsgRNA.search/CsgRNAvshom.4.csv | awk '{print $2}' | rev | tr "[ATGC]" "[TACG]" > CsgRNA.search/Cdownstream_reverse.csv
paste -d'\t' CsgRNA.search/Cdownstream_reverse.csv CsgRNA.search/CsgRNAvshom.4.csv > CsgRNA.search/CsgRNAvshom.4.1.csv
awk 'BEGIN { OFS = "\t" } {$0=gensub(/\s*\S+/,"",3)} {$0=gensub(/\s*\S+/,"",3)} { temp = $1; $1 = $3; $3 = temp } 1' CsgRNA.search/CsgRNAvshom.4.1.csv > CsgRNA.search/CsgRNAvshom.5.csv

#Check for on target counts forward
echo "Checking on targets for 3' sgRNAs (forward)"
awk 'BEGIN {print "#!/bin/bash"; print "mkdir CsgRNA.search/CsgRNA_count_f"}' > CsgRNA.search/CsgRNAs_genome_count.sh
echo "WORKING_GENOME="$WORKING_GENOME >> CsgRNA.search/CsgRNAs_genome_count.sh
cat CsgRNA.search/CsgRNAvshom.5.csv | awk '{print "grep -i -c "$7" $WORKING_GENOME > CsgRNA.search/CsgRNA_count_f/"$1".txt"}' >> CsgRNA.search/CsgRNAs_genome_count.sh
sh CsgRNA.search/CsgRNAs_genome_count.sh
awk '{print $0, FILENAME}' CsgRNA.search/CsgRNA_count_f/*.txt > CsgRNA.search/CsgRNA_count_f.txt
rm -r CsgRNA.search/CsgRNA_count_f
{
sed -e's/[[:space:]]\{1,\}/\t/g' CsgRNA.search/CsgRNA_count_f.txt
} > CsgRNA.search/CsgRNA_count_f.csv
awk 'BEGIN { OFS = "\t" } {gsub("CsgRNA.search/CsgRNA_count_f/","",$2)} {gsub(".txt","",$2)} { temp = $1; $1 = $2; $2 = temp } 1' CsgRNA.search/CsgRNA_count_f.csv > CsgRNA.search/CsgRNA_count_f.2.csv
awk 'NR == FNR{a[$1] = $0;next}; {print $0, $1 in a?a[$1]: "@@@@NA@@@@"}' OFS='\t' CsgRNA.search/CsgRNA_count_f.2.csv CsgRNA.search/CsgRNAvshom.5.csv > CsgRNA.search/pre-final-Ctagging.csv

#Check for on target counts reverse
echo "Checking on targets for 3' sgRNAs (reverse)"
awk 'BEGIN {print "#!/bin/bash"; print "mkdir CsgRNA.search/CsgRNA_count_r"}' > CsgRNA.search/CsgRNAs_genome_count.sh
echo "WORKING_GENOME="$WORKING_GENOME >> CsgRNA.search/CsgRNAs_genome_count.sh
cat CsgRNA.search/CsgRNAvshom.5.csv | awk '{print "grep -i -c "$8" $WORKING_GENOME > CsgRNA.search/CsgRNA_count_r/"$1".txt"}' >> CsgRNA.search/CsgRNAs_genome_count.sh
sh CsgRNA.search/CsgRNAs_genome_count.sh
awk '{print $0, FILENAME}' CsgRNA.search/CsgRNA_count_r/*.txt > CsgRNA.search/CsgRNA_count_r.txt
rm -r CsgRNA.search/CsgRNA_count_r
{
sed -e's/[[:space:]]\{1,\}/\t/g' CsgRNA.search/CsgRNA_count_r.txt
} > CsgRNA.search/CsgRNA_count_r.csv
awk 'BEGIN { OFS = "\t" } {gsub("CsgRNA.search/CsgRNA_count_r/","",$2)} {gsub(".txt","",$2)} { temp = $1; $1 = $2; $2 = temp } 1' CsgRNA.search/CsgRNA_count_r.csv > CsgRNA.search/CsgRNA_count_r.2.csv
awk 'NR == FNR{a[$1] = $0;next}; {print $0, $1 in a?a[$1]: "@@@@NA@@@@"}' OFS='\t' CsgRNA.search/CsgRNA_count_r.2.csv CsgRNA.search/pre-final-Ctagging.csv > CsgRNA.search/pre-final-Ctagging.2.csv
echo "Preparing for final outcome 3'sgRNA"

#Clean-up outcome and add original names (NsgRNA merge should bring in all original search requests)
awk 'BEGIN { OFS = "\t" } {if ($10>$12) ($12 = $10); else ($10 = $12)} {$0=gensub(/\s*\S+/,"",12)} {$0=gensub(/\s*\S+/,"",11)} {$0=gensub(/\s*\S+/,"",9)} {$0=gensub(/\s*\S+/,"",4)} {$0=gensub(/\s*\S+/,"",4)} { temp = $4; $4 = $5; $5 = temp } {$0=gensub(/\s*\S+/,"",6)} { if($6 == 0) ($6 = "@@@@NA@@@@")} 1' CsgRNA.search/pre-final-Ctagging.2.csv > CsgRNA.search/pre-final-Ctagging.3.csv
awk 'NR == FNR{a[$1] = $0;next}; {print $0, $1 in a?a[$1]: "@@@@NA@@@@"}' OFS='\t' CsgRNA.search/pre-final-Ctagging.3.csv NsgRNA.search/pre-final-Ntagging.4.csv > CsgRNA.search/pre-final-C_N.1.csv
awk -F'[[:space:]]' '!$15{ $15="@@@@NA@@@@"}; !$16{ $16="@@@@NA@@@@"}; !$17{ $17="@@@@NA@@@@"}; !$18{ $18="@@@@NA@@@@"}; !$19{ $19="@@@@NA@@@@"}1' OFS='\t' CsgRNA.search/pre-final-C_N.1.csv > CsgRNA.search/pre-final-C_N.csv


echo "End 3'sgRNA processing"
#End 3'sgRNA processing
#-------------------------------------------------------------------------------------------------------------------------------------------------------------------------------------------------------------------------

#-------------------------------------------------------------------------------------------------------------------------------------------------------------------------------------------------------------------------

#Make final table and add plasmid binding sites
awk 'BEGIN { OFS = "\t" } { temp = $5; $5 = $8; $8 = temp } {$0=gensub(/\s*\S+/,"",14)} {$0=gensub(/\s*\S+/,"",7)} {$0=gensub(/\s*\S+/,"",6)} {$0=gensub(/\s*\S+/,"",5)} {$0=gensub(/\s*\S+/,"",4)} {$0=gensub(/\s*\S+/,"",3)} {$0=gensub(/\s*\S+/,"",2)} {$1="";sub("\t","")} {gsub("@@@@NA@@@@","No_sgRNA_found",$2)} {gsub("@@@@NA@@@@","No_sgRNA_found",$3)} {gsub("@@@@NA@@@@","No_sgRNA_found",$4)} {gsub("@@@@","",$5)} {gsub("-1","unable_to_define",$5)} {gsub("@@@@NA@@@@","No_sgRNA_found",$6)} {gsub("@@@@NA@@@@","No_sgRNA_found",$7)} {gsub("@@@@NA@@@@","No_sgRNA_found",$8)} {gsub("@@@@NA@@@@","No_sgRNA_found",$9)} {gsub("@@@@","",$10)} {gsub("-1","unable_to_define",$10)} {gsub("@@@@NA@@@@","No_sgRNA_found",$11)} 1' CsgRNA.search/pre-final-C_N.csv > All_primers_counts.csv

#Cleaning up search files
echo "Cleaning up files and saving off-target searches..."

mkdir 5_off-target_sgRNA_files
mkdir 3_off-target_sgRNA_files
mkdir YOUR_INPUTS
mv NsgRNA.search/*.xls 5_off-target_sgRNA_files
mv CsgRNA.search/*.xls 3_off-target_sgRNA_files
mv $REFERENCE_GENOME YOUR_INPUTS
mv $BED_ORF_INPUT YOUR_INPUTS
cp ORFinput/Genome-ORF.bed.finalmatch.csv 5_off-target_sgRNA_files
cp ORFinput/Genome-ORF.bed.finalmatch.csv 3_off-target_sgRNA_files

rm -f $WORKING_GENOME.*
rm -r Genome-ORF.bed.input.csv
rm -r Bedtools
rm -r ORFinput
rm -r NsgRNA.search
rm -r CsgRNA.search

#-------------------------------------------------------------------------------------------------------------------------------------------------------------------------------------------------------------------------

#-------------------------------------------------------------------------------------------------------------------------------------------------------------------------------------------------------------------------
#adding primer binding sites and modifying sgRNAs for T7 transcription
#Modify final output for pT and pPLOT plasmids
if [ $vector -eq 1 ]; then
	awk 'BEGIN {print "GeneID\tUpstream forward primer\tUpstream reverse primer\tUpstream sgRNA primer\tUpstream sgRNA Off-target counts (1-4 MM)\tUpstream sgRNA perfect match counts\tDownstream forward primer\tDownstream reverse primer\tDownstream sgRNA primer\tDownstream sgRNA Off-target counts (1-4 MM)\tDownstream sgRNA perfect match counts"}' > $REFERENCE_GENOME"pT-pPLOT_primers.csv"
	awk 'BEGIN { OFS = "\t" } {if ($2 != "No_sgRNA_found") ($2 = $2"gtataatgcagacctgctgc")} {if ($3 != "No_sgRNA_found") ($3 = $3"actacccgatcctgatccag")} {if ($4 != "No_sgRNA_found") ($4 = substr($4,1,20))} {if ($4 != "No_sgRNA_found") ($4 = "gaaattaatacgactcactatagg"$4"gttttagagctagaaatagc")} {if ($7 != "No_sgRNA_found") ($7 = $7"ggttctggtagtggttccgg")} {if ($8 != "No_sgRNA_found") ($8 = $8"ccaatttgagagacctgtgc")} {if ($9 != "No_sgRNA_found") ($9 = substr($9,1,20))} {if ($9 != "No_sgRNA_found")($9 = "gaaattaatacgactcactatagg"$9"gttttagagctagaaatagc")} 1' All_primers_counts.csv >> $REFERENCE_GENOME"pT-pPLOT_primers.csv"

elif [ $vector -eq 2 ]; then
	awk 'BEGIN {print "GeneID\tUpstream forward primer\tUpstream reverse primer\tUpstream sgRNA primer\tUpstream sgRNA Off-target counts (1-4 MM)\tUpstream sgRNA perfect match counts\tDownstream forward primer\tDownstream reverse TAG primer\tDownstream reverse KO primer\tDownstream sgRNA primer\tDownstream sgRNA Off-target counts (1-4 MM)\tDownstream sgRNA perfect match counts"}' > $REFERENCE_GENOME"pPOT_primers.csv"
	awk 'BEGIN { OFS = "\t" } {if ($2 != "No_sgRNA_found") ($2 = $2"gtataatgcagacctgctgc")} {if ($3 != "No_sgRNA_found") ($3 = $3"actacccgatcctgatccag")} {if ($4 != "No_sgRNA_found") ($4 = substr($4,1,20))} {if ($4 != "No_sgRNA_found") ($4 = "gaaattaatacgactcactatagg"$4"gttttagagctagaaatagc")} {if ($7 != "No_sgRNA_found") ($7 = $7"ggttctggtagtggttccgg")} {if ($8 != "No_sgRNA_found") ($8 = $8"ccaatttgagagacctgtgc\t"$8"ccggaaccactaccagaacc")} {if ($9 != "No_sgRNA_found") ($9 = substr($9,1,20))} {if ($9 != "No_sgRNA_found") ($9 = "gaaattaatacgactcactatagg"$9"gttttagagctagaaatagc")} 1' All_primers_counts.csv >> $REFERENCE_GENOME"pPOT_primers.csv"

elif [ $vector -eq 3 ]; then
	awk 'BEGIN {print "GeneID\tUpstream forward primer\tUpstream reverse primer\tUpstream sgRNA primer\tUpstream sgRNA Off-target counts (1-4 MM)\tUpstream sgRNA perfect match counts\tDownstream forward primer\tDownstream reverse primer\tDownstream sgRNA primer\tDownstream sgRNA Off-target counts (1-4 MM)\tDownstream sgRNA perfect match counts"}' > $REFERENCE_GENOME"pT-pPLOT_primers.csv"
	awk 'BEGIN { OFS = "\t" } {if ($2 != "No_sgRNA_found") ($2 = $2"gtataatgcagacctgctgc")} {if ($3 != "No_sgRNA_found") ($3 = $3"actacccgatcctgatccag")} {if ($4 != "No_sgRNA_found") ($4 = substr($4,1,20))} {if ($4 != "No_sgRNA_found") ($4 = "gaaattaatacgactcactatagg"$4"gttttagagctagaaatagc")} {if ($7 != "No_sgRNA_found") ($7 = $7"ggttctggtagtggttccgg")} {if ($8 != "No_sgRNA_found") ($8 = $8"ccaatttgagagacctgtgc")} {if ($9 != "No_sgRNA_found") ($9 = substr($9,1,20))} {if ($9 != "No_sgRNA_found")($9 = "gaaattaatacgactcactatagg"$9"gttttagagctagaaatagc")} 1' All_primers_counts.csv >> $REFERENCE_GENOME"pT-pPLOT_primers.csv"
	awk 'BEGIN {print "GeneID\tUpstream forward primer\tUpstream reverse primer\tUpstream sgRNA primer\tUpstream sgRNA Off-target counts (1-4 MM)\tUpstream sgRNA perfect match counts\tDownstream forward primer\tDownstream reverse TAG primer\tDownstream reverse KO primer\tDownstream sgRNA primer\tDownstream sgRNA Off-target counts (1-4 MM)\tDownstream sgRNA perfect match counts"}' > $REFERENCE_GENOME"pPOT_primers.csv"
	awk 'BEGIN { OFS = "\t" } {if ($2 != "No_sgRNA_found") ($2 = $2"gtataatgcagacctgctgc")} {if ($3 != "No_sgRNA_found") ($3 = $3"actacccgatcctgatccag")} {if ($4 != "No_sgRNA_found") ($4 = substr($4,1,20))} {if ($4 != "No_sgRNA_found") ($4 = "gaaattaatacgactcactatagg"$4"gttttagagctagaaatagc")} {if ($7 != "No_sgRNA_found") ($7 = $7"ggttctggtagtggttccgg")} {if ($8 != "No_sgRNA_found") ($8 = $8"ccaatttgagagacctgtgc\t"$8"ccggaaccactaccagaacc")} {if ($9 != "No_sgRNA_found") ($9 = substr($9,1,20))} {if ($9 != "No_sgRNA_found") ($9 = "gaaattaatacgactcactatagg"$9"gttttagagctagaaatagc")} 1' All_primers_counts.csv >> $REFERENCE_GENOME"pPOT_primers.csv"
fi

#-------------------------------------------------------------------------------------------------------------------------------------------------------------------------------------------------------------------------

#-------------------------------------------------------------------------------------------------------------------------------------------------------------------------------------------------------------------------
#End of script
echo "LeishGEdit v4.2 pipeline using CCTop sgRNA design tool (Stemmer et al., PLoS One 2015) is done! Please cite our publications when using these primers: Beneke et al., Roy Soc Open Sci 2017. and Beneke and Gluenz, MBP 2020" 
```
