## Supplementary_file_3 for "Bar-seq strategies for the LeishGEdit toolbox": agrep_count_14995_barcodes.sh

```
#!/bin/bash
#--------------------------------------------------------------------------------------------------------------------------------------------------------------------------------------------------------------

#README
#This script is combining bash commands to count barcodes with 0 mismatches in a given fastq files and was written by Tom Beneke.
#If you would like to count your barcodes with mismatches (which we do not recommend!), you can modify the "agrep" command in the script below.
#Please cite Beneke and Gluenz, MBP 2020 when using it.

#Dependencies:
#Please install "Pandas" to run this script.

#Instructions on usage:
#To run this script copy it into a directory containing all fastq files and the python script 'pivot_table.py'
#Your fastq files will need to be de-multiplexed and unzipped. You can unzip your file e.g. by using "gunzip".
#Navigate to your directory and execute the script with "sh agrep_count_14995_barcodes.sh"

#Output:
#Once the script started it will prompt you to type in your Illumina sequencing machine ID, which it will need to determine the total read count. This will be the "total_read_count.txt" file.
#Additionally, you will find two files termed: "all_counts.csv" and "all_counts.txt". Both output files will contain the barcode counts across all fastq files, but the table is not further formatted.
#The final output file with all counts in an appropriate format will be called "0mismatch_all_barcodes_count.csv". This file will contain all raw barcode counts and should be used for further analysis. 
#
#

#--------------------------------------------------------------------------------------------------------------------------------------------------------------------------------------------------------------

pause(){
   read -p "Press [Enter] key to continue..." fackEnterKey
}

read -p 'Please enter your Illumina sequencing machine ID. Usually this is something like: "@M04023". You will find this above each read in any of your fastq files.: ' MACHINE_ID
	echo "You are uploading <"$MACHINE_ID"> as your Illumina sequencing machine ID."
	pause

#create file folder
mkdir count

#total read count
echo "Counting total number of reads in all fastq files!"
grep -c $MACHINE_ID *.fastq > total_read_count.txt
echo "Count of reads done!"

#count all barcodes
echo "Starting to count all barcodes...this will take a while."

agrep -0 -c 'TCACGGCTGTAACGGAC' *.fastq > count/B4_Fm0.txt
agrep -0 -c 'CGAGTTCGCTATACCGG' *.fastq > count/B5_Fm0.txt
agrep -0 -c 'TACACGAGGTCCGAAAT' *.fastq > count/B7_Fm0.txt
agrep -0 -c 'CTATACATACCCCAGAG' *.fastq > count/B10_Fm0.txt
agrep -0 -c 'ACACCAATAGTGTGATC' *.fastq > count/B11_Fm0.txt
agrep -0 -c 'ATGGCTGCTGCACACCT' *.fastq > count/B14_Fm0.txt
agrep -0 -c 'AAGTCCCGAATGATATC' *.fastq > count/B15_Fm0.txt
agrep -0 -c 'CGTGTCTATGAGGCCGT' *.fastq > count/B16_Fm0.txt
agrep -0 -c 'GATTCCACGCGCTCTTC' *.fastq > count/B17_Fm0.txt
agrep -0 -c 'AGGTCCCAGCATTCTAG' *.fastq > count/B18_Fm0.txt
agrep -0 -c 'GTATGATACGTACCCAA' *.fastq > count/B19_Fm0.txt
agrep -0 -c 'TGCGCAAGGTTAAGTTA' *.fastq > count/B21_Fm0.txt
agrep -0 -c 'TATCCACTTTCTCGACG' *.fastq > count/B22_Fm0.txt
agrep -0 -c 'CCGCCCCTTAAATAAAG' *.fastq > count/B23_Fm0.txt
agrep -0 -c 'GCTTAGATCCATGTCGG' *.fastq > count/B24_Fm0.txt
agrep -0 -c 'CAAGTCAGGGTTGTAGT' *.fastq > count/B25_Fm0.txt
agrep -0 -c 'GTCCACTCACAATCTGA' *.fastq > count/B26_Fm0.txt
agrep -0 -c 'CGTTGTCTTAAGTGCGC' *.fastq > count/B27_Fm0.txt
agrep -0 -c 'CAGGATTACTATAGGCT' *.fastq > count/B29_Fm0.txt
agrep -0 -c 'AGTACTTGTGCTGGCAA' *.fastq > count/B30_Fm0.txt
agrep -0 -c 'TGTTTGTAACGGTTTGC' *.fastq > count/B33_Fm0.txt
agrep -0 -c 'ACATATGCATGGCACCC' *.fastq > count/B34_Fm0.txt
agrep -0 -c 'ATCTGCGGCAGTAGGTC' *.fastq > count/B35_Fm0.txt
agrep -0 -c 'GACACGTGGCAGGTATT' *.fastq > count/B36_Fm0.txt
agrep -0 -c 'TCGACCGTACGTTAGGT' *.fastq > count/B37_Fm0.txt
agrep -0 -c 'GTATATGAGATAGACGG' *.fastq > count/B39_Fm0.txt
agrep -0 -c 'TGACTTTTACCGCGTGA' *.fastq > count/B40_Fm0.txt
agrep -0 -c 'CAATACTCCCGCCGAAT' *.fastq > count/B42_Fm0.txt
agrep -0 -c 'GATCTCTATGCTCTAGT' *.fastq > count/B44_Fm0.txt
agrep -0 -c 'CATAGTACCCAGCCTGA' *.fastq > count/B45_Fm0.txt
agrep -0 -c 'TACACCTAGACGGCTAG' *.fastq > count/B46_Fm0.txt
agrep -0 -c 'GTAGTATCGTATCGGAA' *.fastq > count/B48_Fm0.txt
agrep -0 -c 'TTGCGACCGGATAGCCA' *.fastq > count/B49_Fm0.txt
agrep -0 -c 'TAATCGGGCACTGCCAC' *.fastq > count/B50_Fm0.txt
agrep -0 -c 'CACTCACGCTATTCGTA' *.fastq > count/B51_Fm0.txt
agrep -0 -c 'TACCGCCGGGTGTCTAA' *.fastq > count/B52_Fm0.txt
agrep -0 -c 'CTTGGATATTAGCGTCA' *.fastq > count/B53_Fm0.txt
agrep -0 -c 'CAAACGTCGATACATTG' *.fastq > count/B54_Fm0.txt
agrep -0 -c 'TAAGGACAAGCGTACAA' *.fastq > count/B55_Fm0.txt
agrep -0 -c 'AGATCGAATGCTCACGC' *.fastq > count/B56_Fm0.txt
agrep -0 -c 'CAGCGTTCGAACACACA' *.fastq > count/B57_Fm0.txt
agrep -0 -c 'TAGACGGGCAAACCAGA' *.fastq > count/B58_Fm0.txt
agrep -0 -c 'TATAGCGGCACCATCGC' *.fastq > count/B60_Fm0.txt
agrep -0 -c 'TCATAATGACAGGACCT' *.fastq > count/B61_Fm0.txt
agrep -0 -c 'CCGCACAATTCTACAGG' *.fastq > count/B62_Fm0.txt
agrep -0 -c 'CTCTAAGGGCGGTTTGG' *.fastq > count/B63_Fm0.txt
agrep -0 -c 'AAGAGGAGATTAACGAG' *.fastq > count/B64_Fm0.txt
agrep -0 -c 'CGAAATCCTTAGTTCCC' *.fastq > count/B65_Fm0.txt
agrep -0 -c 'GACCATAAACCCACTCG' *.fastq > count/B66_Fm0.txt
agrep -0 -c 'GGTAGTTGTCAAGCATA' *.fastq > count/B67_Fm0.txt
agrep -0 -c 'GGACGTATTCAGTCGAC' *.fastq > count/B68_Fm0.txt
agrep -0 -c 'CTCCTTACTCTCCCGGT' *.fastq > count/B69_Fm0.txt
agrep -0 -c 'TTCAATTCCCTATTCGC' *.fastq > count/B71_Fm0.txt
agrep -0 -c 'CTGACTGTTCCCACAAC' *.fastq > count/B72_Fm0.txt
agrep -0 -c 'GATCACAGCGTAGTACC' *.fastq > count/B73_Fm0.txt
agrep -0 -c 'TGTGTTGTATGAGCGGA' *.fastq > count/B75_Fm0.txt
agrep -0 -c 'CGAGCGGCAAATATTAT' *.fastq > count/B76_Fm0.txt
agrep -0 -c 'ACCGCGAAGAGAGGACT' *.fastq > count/B77_Fm0.txt
agrep -0 -c 'TCACAGGGCGTAAAGTC' *.fastq > count/B78_Fm0.txt
agrep -0 -c 'ACCTTAGTTTCCCAAAC' *.fastq > count/B79_Fm0.txt
agrep -0 -c 'TACGGGGATTTGTTCCC' *.fastq > count/B80_Fm0.txt
agrep -0 -c 'CGGTCTTTCGCGTCAAG' *.fastq > count/B81_Fm0.txt
agrep -0 -c 'GCCATGTAGTTAGCCTT' *.fastq > count/B82_Fm0.txt
agrep -0 -c 'GACAGCCTCGAACCTCA' *.fastq > count/B83_Fm0.txt
agrep -0 -c 'TTGATCTGGCTGCAAAC' *.fastq > count/B84_Fm0.txt
agrep -0 -c 'AGGTCTACCTTTGATGG' *.fastq > count/B85_Fm0.txt
agrep -0 -c 'GGATATCAGTTGACAAC' *.fastq > count/B87_Fm0.txt
agrep -0 -c 'CAACTTCGCAACCAGTT' *.fastq > count/B88_Fm0.txt
agrep -0 -c 'CACCGTAGCGTGCCTTA' *.fastq > count/B89_Fm0.txt
agrep -0 -c 'TTCTTTTCCACCCGGAG' *.fastq > count/B90_Fm0.txt
agrep -0 -c 'GGCTTTCACATTGGTTA' *.fastq > count/B91_Fm0.txt
agrep -0 -c 'GGTAAAAGTGGCAGAGT' *.fastq > count/B92_Fm0.txt
agrep -0 -c 'CACGACACATTACTGGC' *.fastq > count/B93_Fm0.txt
agrep -0 -c 'ACAAGTACACCCTACAT' *.fastq > count/B94_Fm0.txt
agrep -0 -c 'TTGTAGTGTCGTGGGGA' *.fastq > count/B95_Fm0.txt
agrep -0 -c 'AAGTACGCCTATCTACA' *.fastq > count/B96_Fm0.txt
agrep -0 -c 'CTTGGCCGGGACAGATT' *.fastq > count/B97_Fm0.txt
agrep -0 -c 'CCAGCCCTGAGCTTACA' *.fastq > count/B98_Fm0.txt
agrep -0 -c 'AGTCTTCGCAAAGCTCA' *.fastq > count/B99_Fm0.txt
agrep -0 -c 'AGCGGGCCATATTTTAC' *.fastq > count/B100_Fm0.txt
agrep -0 -c 'TAAGGCTGGTTCCGTCA' *.fastq > count/B103_Fm0.txt
agrep -0 -c 'ACTATCTTCCGTCCCCC' *.fastq > count/B104_Fm0.txt
agrep -0 -c 'GAGAATTCGCAGTTGGC' *.fastq > count/B105_Fm0.txt
agrep -0 -c 'ACAGCAAATCGATTACC' *.fastq > count/B106_Fm0.txt
agrep -0 -c 'TACACCAATAGCTGATG' *.fastq > count/B107_Fm0.txt
agrep -0 -c 'ATTCCAAGTCTCACGAG' *.fastq > count/B108_Fm0.txt
agrep -0 -c 'CATCCGCTGTTGTGAGC' *.fastq > count/B109_Fm0.txt
agrep -0 -c 'GAGCGCATGGTCTGATC' *.fastq > count/B110_Fm0.txt
agrep -0 -c 'CGGGACCTCGTTCTTAT' *.fastq > count/B112_Fm0.txt
agrep -0 -c 'GTCGACGGGTTTCTGTA' *.fastq > count/B113_Fm0.txt
agrep -0 -c 'TTGAATGAACCCCGCTC' *.fastq > count/B114_Fm0.txt
agrep -0 -c 'TCATGGGACGTACGCAA' *.fastq > count/B115_Fm0.txt
agrep -0 -c 'ATAAGAATCGGCCTTGA' *.fastq > count/B116_Fm0.txt
agrep -0 -c 'CGAGACGATACAGTGTG' *.fastq > count/B117_Fm0.txt
agrep -0 -c 'GCAAACTAGAGAGTCAA' *.fastq > count/B118_Fm0.txt
agrep -0 -c 'GACTTCGGACGAGTCAA' *.fastq > count/B120_Fm0.txt
agrep -0 -c 'GACACAGCCTCAATATT' *.fastq > count/B121_Fm0.txt
agrep -0 -c 'CGTGGGTTTCGGTTTTG' *.fastq > count/B122_Fm0.txt
agrep -0 -c 'AGACCCGTCGATCAATG' *.fastq > count/B123_Fm0.txt
agrep -0 -c 'GTCAGGAAGTATCCGGA' *.fastq > count/B124_Fm0.txt
agrep -0 -c 'ATGTAGGCTCGCAAAGG' *.fastq > count/B125_Fm0.txt
agrep -0 -c 'GCGGTGGCTAAAAATGC' *.fastq > count/B126_Fm0.txt
agrep -0 -c 'GGTGTACCAAACTTGTA' *.fastq > count/B128_Fm0.txt
agrep -0 -c 'GCCTGGTGGATTTTCAC' *.fastq > count/B129_Fm0.txt
agrep -0 -c 'ATAGGAAGAATCACACG' *.fastq > count/B131_Fm0.txt
agrep -0 -c 'ACATATCTCGGTATTCG' *.fastq > count/B5446_Fm0.txt
agrep -0 -c 'AGTATAACCGAGGGAAC' *.fastq > count/B8570_Fm0.txt
agrep -0 -c 'GCGAACTACTCCAGTGC' *.fastq > count/B269_Fm0.txt
agrep -0 -c 'AAGCTACGACTACTTAG' *.fastq > count/B1313_Fm0.txt
agrep -0 -c 'GTGGATTACGTAGACCG' *.fastq > count/B160_Fm0.txt
agrep -0 -c 'GTCACCACAGTTAATTG' *.fastq > count/B167_Fm0.txt
agrep -0 -c 'TCATCTGGACACTGCAA' *.fastq > count/B259_Fm0.txt
agrep -0 -c 'GCCGGTGCACATATTCG' *.fastq > count/B305_Fm0.txt
agrep -0 -c 'ATCCTATACGACAGGAA' *.fastq > count/B306_Fm0.txt
agrep -0 -c 'GTCATAAATCGCTTGTC' *.fastq > count/B374_Fm0.txt
agrep -0 -c 'CCGACCATCGTTGTCAC' *.fastq > count/B4161_Fm0.txt
agrep -0 -c 'TTACGGGCACTTCCTAT' *.fastq > count/B5277_Fm0.txt
agrep -0 -c 'GTCCATCTGGTCATTGT' *.fastq > count/B5328_Fm0.txt
agrep -0 -c 'CAGTATGAAGGCTCCTG' *.fastq > count/B5744_Fm0.txt
agrep -0 -c 'CTCGTACGTCCCGCATT' *.fastq > count/B7238_Fm0.txt
agrep -0 -c 'CATACTCCCCGTTTTGA' *.fastq > count/B7310_Fm0.txt
agrep -0 -c 'ACCTGCTCGAACGGTAT' *.fastq > count/B8804_Fm0.txt
agrep -0 -c 'TAAGAAGAGACTGATGC' *.fastq > count/B8863_Fm0.txt
agrep -0 -c 'CCATGGACGTATTGTGT' *.fastq > count/B9338_Fm0.txt
agrep -0 -c 'CTATCCCACTCACGTTA' *.fastq > count/B9783_Fm0.txt
agrep -0 -c 'GCAGGTCTTCTCATACG' *.fastq > count/B9788_Fm0.txt
agrep -0 -c 'GCTAACGGAGTGAGCGT' *.fastq > count/B1058_Fm0.txt
agrep -0 -c 'GGCAGAAATCTGCTTGT' *.fastq > count/B1065_Fm0.txt
agrep -0 -c 'GGCCCGCGTATCTAACT' *.fastq > count/B132_Fm0.txt
agrep -0 -c 'TGGCTTAGCGCGAGACA' *.fastq > count/B133_Fm0.txt
agrep -0 -c 'AATACTAGGCAACGAGG' *.fastq > count/B134_Fm0.txt
agrep -0 -c 'GTGTTTCACATCTACGC' *.fastq > count/B137_Fm0.txt
agrep -0 -c 'GCAGTCTCAATGCGGGT' *.fastq > count/B139_Fm0.txt
agrep -0 -c 'TACCAGCGGAAAGGAGA' *.fastq > count/B141_Fm0.txt
agrep -0 -c 'TACAGATGCGTAGTCGG' *.fastq > count/B142_Fm0.txt
agrep -0 -c 'ACCAATATGTCTGCTCG' *.fastq > count/B144_Fm0.txt
agrep -0 -c 'AATCGGTGGTCGATCTT' *.fastq > count/B146_Fm0.txt
agrep -0 -c 'AGAATATCGAGCAGGGC' *.fastq > count/B148_Fm0.txt
agrep -0 -c 'GTAGTTCGCTCCAGACG' *.fastq > count/B149_Fm0.txt
agrep -0 -c 'ATGACCTTCGGGGTCTA' *.fastq > count/B150_Fm0.txt
agrep -0 -c 'GTCGAATATGTCAACGA' *.fastq > count/B151_Fm0.txt
agrep -0 -c 'ACATTATGGAGCTACGA' *.fastq > count/B152_Fm0.txt
agrep -0 -c 'CTCCGACTACGACGCAA' *.fastq > count/B153_Fm0.txt
agrep -0 -c 'GGTTTACACATAGCCTG' *.fastq > count/B155_Fm0.txt
agrep -0 -c 'GCCCTACCCTGTGGATT' *.fastq > count/B156_Fm0.txt
agrep -0 -c 'GGTAGGTACGGCAAATA' *.fastq > count/B159_Fm0.txt
agrep -0 -c 'CAAAGTCCATACCCGAC' *.fastq > count/B161_Fm0.txt
agrep -0 -c 'GTTTGGTCGTCGTTCCT' *.fastq > count/B162_Fm0.txt
agrep -0 -c 'CGAGAAACTTGGACGCC' *.fastq > count/B163_Fm0.txt
agrep -0 -c 'TGTAGGTACTATCTCCT' *.fastq > count/B164_Fm0.txt
agrep -0 -c 'GGTCACACATGACGGTT' *.fastq > count/B165_Fm0.txt
agrep -0 -c 'CGTCGTAAAACTAGACA' *.fastq > count/B166_Fm0.txt
agrep -0 -c 'CATGCAAGAATTCAGAG' *.fastq > count/B168_Fm0.txt
agrep -0 -c 'TAATATCCCCGAGCTTT' *.fastq > count/B169_Fm0.txt
agrep -0 -c 'CGAAGCACTGGACAATC' *.fastq > count/B170_Fm0.txt
agrep -0 -c 'TTAGCATCGAACACGTT' *.fastq > count/B171_Fm0.txt
agrep -0 -c 'TCGAGCGACTTCATGCA' *.fastq > count/B172_Fm0.txt
agrep -0 -c 'TTGGCCTTTGGCCATTC' *.fastq > count/B173_Fm0.txt
agrep -0 -c 'ACGCGAGTGATGTGTTC' *.fastq > count/B174_Fm0.txt
agrep -0 -c 'AGTGAGCTCAGAATTCA' *.fastq > count/B175_Fm0.txt
agrep -0 -c 'GTTCTTAAACCGGTCCC' *.fastq > count/B176_Fm0.txt
agrep -0 -c 'CATCCCGAAGCACTCTT' *.fastq > count/B177_Fm0.txt
agrep -0 -c 'TTGACCCAAACTTCGAA' *.fastq > count/B179_Fm0.txt
agrep -0 -c 'TCTGATGAAACACGACA' *.fastq > count/B180_Fm0.txt
agrep -0 -c 'GCAGTACATGTAGGGAT' *.fastq > count/B183_Fm0.txt
agrep -0 -c 'ACGGATTAGGCCACTTA' *.fastq > count/B184_Fm0.txt
agrep -0 -c 'GTTAAACCTTCACGTTC' *.fastq > count/B185_Fm0.txt
agrep -0 -c 'TACCGTAAAGCTCTTTG' *.fastq > count/B186_Fm0.txt
agrep -0 -c 'TGGCCGGAACACAACCT' *.fastq > count/B187_Fm0.txt
agrep -0 -c 'GCCCCAGCAACCTAGAT' *.fastq > count/B188_Fm0.txt
agrep -0 -c 'GTATGCAGTAGCGGATC' *.fastq > count/B189_Fm0.txt
agrep -0 -c 'GTGTACAACGAAGTCTC' *.fastq > count/B190_Fm0.txt
agrep -0 -c 'GGGACTCTGATTCGAAA' *.fastq > count/B191_Fm0.txt
agrep -0 -c 'AACACTGGGGTCGACTG' *.fastq > count/B192_Fm0.txt
agrep -0 -c 'CCATAACGAAAACACTG' *.fastq > count/B194_Fm0.txt
agrep -0 -c 'TCGTAAGACTGGTCGTT' *.fastq > count/B195_Fm0.txt
agrep -0 -c 'ATGGTGCGGCTTAGAAA' *.fastq > count/B196_Fm0.txt
agrep -0 -c 'TCAAACCCACTAACACC' *.fastq > count/B197_Fm0.txt
agrep -0 -c 'AACACAAATGCCCTCGC' *.fastq > count/B198_Fm0.txt
agrep -0 -c 'ATGTGGCTCAATCACTT' *.fastq > count/B199_Fm0.txt
agrep -0 -c 'TGCAGGACGTGCAACAT' *.fastq > count/B200_Fm0.txt
agrep -0 -c 'TACGCAACAAGCTGATT' *.fastq > count/B201_Fm0.txt
agrep -0 -c 'AGGGCAAACGTGGAAGT' *.fastq > count/B202_Fm0.txt
agrep -0 -c 'AACAACGTCGTACCCGC' *.fastq > count/B205_Fm0.txt
agrep -0 -c 'GTTCACCAGTACCAGTC' *.fastq > count/B206_Fm0.txt
agrep -0 -c 'TCGCGGGCTCACCATTT' *.fastq > count/B207_Fm0.txt
agrep -0 -c 'GTCGTAGTAAAATGACC' *.fastq > count/B208_Fm0.txt
agrep -0 -c 'CACAAGAGTCGTGTACC' *.fastq > count/B209_Fm0.txt
agrep -0 -c 'AGCCTCTACACGTTGCG' *.fastq > count/B210_Fm0.txt
agrep -0 -c 'TCTACACCACGGGTAAA' *.fastq > count/B211_Fm0.txt
agrep -0 -c 'AAGCAAGCATTCGTTGA' *.fastq > count/B213_Fm0.txt
agrep -0 -c 'AACTCCATAGGACACGC' *.fastq > count/B214_Fm0.txt
agrep -0 -c 'ACAAGACACGACACATA' *.fastq > count/B215_Fm0.txt
agrep -0 -c 'GCGTTAGTACCGAACAC' *.fastq > count/B216_Fm0.txt
agrep -0 -c 'AGCTCCTTTGGCACAAA' *.fastq > count/B217_Fm0.txt
agrep -0 -c 'CACTACGAGCGAAAACA' *.fastq > count/B219_Fm0.txt
agrep -0 -c 'TTGCTGAAGATTTGGTC' *.fastq > count/B220_Fm0.txt
agrep -0 -c 'ATAAGAGACCCCCCTTT' *.fastq > count/B221_Fm0.txt
agrep -0 -c 'CTCATAACTTTGCCCGA' *.fastq > count/B222_Fm0.txt
agrep -0 -c 'TAACAGCCGTAGTGTTA' *.fastq > count/B223_Fm0.txt
agrep -0 -c 'AGACGTGCTGGCAGATC' *.fastq > count/B224_Fm0.txt
agrep -0 -c 'GTTGGTGTTAGGACTCT' *.fastq > count/B225_Fm0.txt
agrep -0 -c 'CGCTTATCCCAATGACT' *.fastq > count/B226_Fm0.txt
agrep -0 -c 'TCATCTGGGGGAACACC' *.fastq > count/B227_Fm0.txt
agrep -0 -c 'CGAATCGGATATCTGGA' *.fastq > count/B228_Fm0.txt
agrep -0 -c 'CCTCCTCTAGTACCCGA' *.fastq > count/B229_Fm0.txt
agrep -0 -c 'TACAGTTACCAGGCGAT' *.fastq > count/B230_Fm0.txt
agrep -0 -c 'CTGGCCCATAGAACCAT' *.fastq > count/B231_Fm0.txt
agrep -0 -c 'CGGCTTTACTTTTACCA' *.fastq > count/B233_Fm0.txt
agrep -0 -c 'CAATGTCGCATTCACTT' *.fastq > count/B234_Fm0.txt
agrep -0 -c 'GTCAACGTATATCAGGC' *.fastq > count/B235_Fm0.txt
agrep -0 -c 'GTCGCTAACTCCAAATC' *.fastq > count/B236_Fm0.txt
agrep -0 -c 'GGTGGTTTAACATGCAG' *.fastq > count/B238_Fm0.txt
agrep -0 -c 'TGAAACATAGGCGAACA' *.fastq > count/B239_Fm0.txt
agrep -0 -c 'TTATCCTCTATCGAACG' *.fastq > count/B240_Fm0.txt
agrep -0 -c 'TATCAGTGTCCAGCAAC' *.fastq > count/B241_Fm0.txt
agrep -0 -c 'TTCGTCGACATCTGCCC' *.fastq > count/B242_Fm0.txt
agrep -0 -c 'GGAGCCACTTGACGAAC' *.fastq > count/B243_Fm0.txt
agrep -0 -c 'TAAGCATTGCTGCCATC' *.fastq > count/B244_Fm0.txt
agrep -0 -c 'GCTTAGTTCCCTCTAAT' *.fastq > count/B245_Fm0.txt
agrep -0 -c 'CGTGGTTCTGTGTATCT' *.fastq > count/B247_Fm0.txt
agrep -0 -c 'GTGTTGCGTTGGCGTGA' *.fastq > count/B248_Fm0.txt
agrep -0 -c 'AAGGTTGCGCGTAACCG' *.fastq > count/B249_Fm0.txt
agrep -0 -c 'GCGTTCCACTTTAGTAA' *.fastq > count/B251_Fm0.txt
agrep -0 -c 'CATGAAAGGGTACGAAC' *.fastq > count/B252_Fm0.txt
agrep -0 -c 'TGTGAGGTCGGGGGTAT' *.fastq > count/B253_Fm0.txt
agrep -0 -c 'AAGTTTAAGTCACCGCA' *.fastq > count/B254_Fm0.txt
agrep -0 -c 'CTCCCGACAGATGCTTA' *.fastq > count/B255_Fm0.txt
agrep -0 -c 'TGATCGTAACCACGATT' *.fastq > count/B257_Fm0.txt
agrep -0 -c 'GGGAGTGTCGTGATGAC' *.fastq > count/B258_Fm0.txt
agrep -0 -c 'ATATCTGCGAAGTCCGA' *.fastq > count/B260_Fm0.txt
agrep -0 -c 'CCGGTCACAGATGAAAT' *.fastq > count/B261_Fm0.txt
agrep -0 -c 'CGAATGACCCACGGAAA' *.fastq > count/B262_Fm0.txt
agrep -0 -c 'GTCTTAACCCCGTTTGA' *.fastq > count/B263_Fm0.txt
agrep -0 -c 'ATCAGGTCACCCCATAC' *.fastq > count/B264_Fm0.txt
agrep -0 -c 'GAGAGCTGTCAGGTGTA' *.fastq > count/B267_Fm0.txt
agrep -0 -c 'GTAAGGGTTACTGTCAG' *.fastq > count/B268_Fm0.txt
agrep -0 -c 'TGTCCATCGGATTGGTA' *.fastq > count/B270_Fm0.txt
agrep -0 -c 'CAGAACCGGCAAACCAA' *.fastq > count/B271_Fm0.txt
agrep -0 -c 'GGGAGCACATACATTGG' *.fastq > count/B273_Fm0.txt
agrep -0 -c 'GGAAGTTTGGTCTGTGC' *.fastq > count/B274_Fm0.txt
agrep -0 -c 'AGCTCGAAACGCCCAGT' *.fastq > count/B276_Fm0.txt
agrep -0 -c 'CCAAGTGGTAGCGAGTG' *.fastq > count/B277_Fm0.txt
agrep -0 -c 'GCGTAAGTTTCCACAGA' *.fastq > count/B278_Fm0.txt
agrep -0 -c 'GACTCTTATCCGGATGG' *.fastq > count/B279_Fm0.txt
agrep -0 -c 'ATTACCACGTGGATGTC' *.fastq > count/B281_Fm0.txt
agrep -0 -c 'CGTGGATGGGATAGTGT' *.fastq > count/B282_Fm0.txt
agrep -0 -c 'ATACACAGGTGGCGTGT' *.fastq > count/B283_Fm0.txt
agrep -0 -c 'ATAGGTGAAGACCGGGC' *.fastq > count/B284_Fm0.txt
agrep -0 -c 'AGTAACGGTTTGTGTAC' *.fastq > count/B286_Fm0.txt
agrep -0 -c 'AGACCTCGTCAGTCATC' *.fastq > count/B288_Fm0.txt
agrep -0 -c 'TAGATGCGTACTTGGAG' *.fastq > count/B289_Fm0.txt
agrep -0 -c 'ATCCCAATTCCGCCTAG' *.fastq > count/B290_Fm0.txt
agrep -0 -c 'TGCTCGGGTCAGATGAA' *.fastq > count/B291_Fm0.txt
agrep -0 -c 'AGCTTTCTTCGCTGGAA' *.fastq > count/B293_Fm0.txt
agrep -0 -c 'AGGGTCAACTCGGGGTT' *.fastq > count/B294_Fm0.txt
agrep -0 -c 'GCGGTCTTAGGCTAGCT' *.fastq > count/B295_Fm0.txt
agrep -0 -c 'TGTGAAGAAGAGCTCAA' *.fastq > count/B296_Fm0.txt
agrep -0 -c 'CAAATCAGGCTCGATAA' *.fastq > count/B297_Fm0.txt
agrep -0 -c 'CAGCCATGTCCCTCAAA' *.fastq > count/B299_Fm0.txt
agrep -0 -c 'ACACGGGTAGATGCCCA' *.fastq > count/B300_Fm0.txt
agrep -0 -c 'AAGAATCGACTCGTAGT' *.fastq > count/B301_Fm0.txt
agrep -0 -c 'CTATTCATCGTGGGATT' *.fastq > count/B302_Fm0.txt
agrep -0 -c 'TAGGCAATCAGTCATCT' *.fastq > count/B303_Fm0.txt
agrep -0 -c 'TCGGACGCTATTAAGGG' *.fastq > count/B304_Fm0.txt
agrep -0 -c 'GGCAACCGATAGATTCA' *.fastq > count/B307_Fm0.txt
agrep -0 -c 'TTCAAACGGTTAGGAGG' *.fastq > count/B309_Fm0.txt
agrep -0 -c 'ACCCTCAGTAGGTATAC' *.fastq > count/B311_Fm0.txt
agrep -0 -c 'GTATCGGGTCATCGTTT' *.fastq > count/B312_Fm0.txt
agrep -0 -c 'GCGTCGAGAGTACGTAG' *.fastq > count/B313_Fm0.txt
agrep -0 -c 'CCTCAATCGGGATAAAG' *.fastq > count/B314_Fm0.txt
agrep -0 -c 'TACCTGTAGTGTGCTCG' *.fastq > count/B315_Fm0.txt
agrep -0 -c 'CCGCGACAGAGAATTCT' *.fastq > count/B316_Fm0.txt
agrep -0 -c 'CAGGTGACGGCACTGAT' *.fastq > count/B317_Fm0.txt
agrep -0 -c 'GCATCTGTGGTGACTTA' *.fastq > count/B318_Fm0.txt
agrep -0 -c 'AGTCACTGCGCTCCGAT' *.fastq > count/B322_Fm0.txt
agrep -0 -c 'GTTTTGTGCTGCCCGAC' *.fastq > count/B323_Fm0.txt
agrep -0 -c 'TACATATGCCGTAGGGT' *.fastq > count/B324_Fm0.txt
agrep -0 -c 'TGTTGGTCGTAAGGGAA' *.fastq > count/B325_Fm0.txt
agrep -0 -c 'AACCCATATTTTGGCGC' *.fastq > count/B327_Fm0.txt
agrep -0 -c 'AGTGATCGTAGTTCCAG' *.fastq > count/B329_Fm0.txt
agrep -0 -c 'AGCACAAGACAGATTGT' *.fastq > count/B331_Fm0.txt
agrep -0 -c 'AGGTGGTGAATGCTCTA' *.fastq > count/B332_Fm0.txt
agrep -0 -c 'GCCGCTTACCCTTCGAA' *.fastq > count/B333_Fm0.txt
agrep -0 -c 'AACACGGATAAGCCGGG' *.fastq > count/B335_Fm0.txt
agrep -0 -c 'TCACTAAAGGCGTACCG' *.fastq > count/B336_Fm0.txt
agrep -0 -c 'GGCGGACCTAGGAAGTA' *.fastq > count/B337_Fm0.txt
agrep -0 -c 'TTACAGACACGAGTCCT' *.fastq > count/B338_Fm0.txt
agrep -0 -c 'GCGTATACCAGGACTGC' *.fastq > count/B340_Fm0.txt
agrep -0 -c 'TAAAGTGGCCCCACAAG' *.fastq > count/B341_Fm0.txt
agrep -0 -c 'ACTACCCAAGTTGCCCG' *.fastq > count/B342_Fm0.txt
agrep -0 -c 'TTACGTGCTGATGCACA' *.fastq > count/B344_Fm0.txt
agrep -0 -c 'GCATCGTACTAGGTAGG' *.fastq > count/B346_Fm0.txt
agrep -0 -c 'CTCCAAAGGTAGCGCCA' *.fastq > count/B347_Fm0.txt
agrep -0 -c 'AATTCGTGGGTCCGGCT' *.fastq > count/B349_Fm0.txt
agrep -0 -c 'GCGTCTAAATTAGGGAC' *.fastq > count/B350_Fm0.txt
agrep -0 -c 'AATGTAGACGAGGCATC' *.fastq > count/B351_Fm0.txt
agrep -0 -c 'CAGGTCACCATGTTACC' *.fastq > count/B353_Fm0.txt
agrep -0 -c 'TGTTAGAGAGGAGCCCT' *.fastq > count/B354_Fm0.txt
agrep -0 -c 'GTTAGTCTCTGCTGCAG' *.fastq > count/B355_Fm0.txt
agrep -0 -c 'AAGGAGTGATTACTCAC' *.fastq > count/B357_Fm0.txt
agrep -0 -c 'ATAGAGGACGTGTATCT' *.fastq > count/B358_Fm0.txt
agrep -0 -c 'AATCGGGTCCTCCTTAC' *.fastq > count/B360_Fm0.txt
agrep -0 -c 'AAGCTATTGGAGCGCGT' *.fastq > count/B361_Fm0.txt
agrep -0 -c 'TGACATCCGGCATAAAT' *.fastq > count/B362_Fm0.txt
agrep -0 -c 'GGAGTCTGATAGTCACG' *.fastq > count/B363_Fm0.txt
agrep -0 -c 'TTACACGACGAACGGCC' *.fastq > count/B364_Fm0.txt
agrep -0 -c 'CGCCTCCAATTCTAAAG' *.fastq > count/B365_Fm0.txt
agrep -0 -c 'TGCGTAATCAAATGCTC' *.fastq > count/B366_Fm0.txt
agrep -0 -c 'AGAACGAGATGCACGTA' *.fastq > count/B367_Fm0.txt
agrep -0 -c 'TGTAATGTTTCCGGCCT' *.fastq > count/B369_Fm0.txt
agrep -0 -c 'AACAGCCGAAGCGTCTC' *.fastq > count/B370_Fm0.txt
agrep -0 -c 'ATGTCTTCTCATGTCGG' *.fastq > count/B372_Fm0.txt
agrep -0 -c 'TCTTACGACGGTCTAAT' *.fastq > count/B375_Fm0.txt
agrep -0 -c 'TACAGAAGCCTCATCTA' *.fastq > count/B376_Fm0.txt
agrep -0 -c 'GGGTGGAGTTCCTATGT' *.fastq > count/B377_Fm0.txt
agrep -0 -c 'TGCGTAAGTTCGCTTTG' *.fastq > count/B378_Fm0.txt
agrep -0 -c 'CTGTTGTCGAATCATTG' *.fastq > count/B379_Fm0.txt
agrep -0 -c 'AACACCATAGAACGGAC' *.fastq > count/B381_Fm0.txt
agrep -0 -c 'GGCATACCGCCTTTCGA' *.fastq > count/B382_Fm0.txt
agrep -0 -c 'ATGGAATTACCGCGACT' *.fastq > count/B383_Fm0.txt
agrep -0 -c 'GCGTATAGAACGAGATC' *.fastq > count/B384_Fm0.txt
agrep -0 -c 'ACGCCTACCTCCAAATG' *.fastq > count/B385_Fm0.txt
agrep -0 -c 'CATCTGTCAAGCTGCTC' *.fastq > count/B386_Fm0.txt
agrep -0 -c 'AAGCTGGCTCTAGCTGT' *.fastq > count/B387_Fm0.txt
agrep -0 -c 'GCACCCCAAGGTAACCA' *.fastq > count/B388_Fm0.txt
agrep -0 -c 'GAGGTTGTGATGTCGCT' *.fastq > count/B389_Fm0.txt
agrep -0 -c 'CGATTCATTAACCCACA' *.fastq > count/B390_Fm0.txt
agrep -0 -c 'ATGAAAGACTGGACGAT' *.fastq > count/B392_Fm0.txt
agrep -0 -c 'GGCGCATTCAGGTTTTC' *.fastq > count/B393_Fm0.txt
agrep -0 -c 'ACGCTAAGCCCCGACTA' *.fastq > count/B394_Fm0.txt
agrep -0 -c 'GTGCTATGTACAGTGGC' *.fastq > count/B395_Fm0.txt
agrep -0 -c 'TATGTCCCGCTTCGACT' *.fastq > count/B396_Fm0.txt
agrep -0 -c 'GACAGTTTAATTGGGGC' *.fastq > count/B397_Fm0.txt
agrep -0 -c 'GCTATATAAGTCCAAGG' *.fastq > count/B398_Fm0.txt
agrep -0 -c 'GACGTACACAAAAGACC' *.fastq > count/B399_Fm0.txt
agrep -0 -c 'ACATGGAATAAGTTCGG' *.fastq > count/B400_Fm0.txt
agrep -0 -c 'TCGGTGAACTCATCATC' *.fastq > count/B401_Fm0.txt
agrep -0 -c 'GGAGTGTCGCTCGAGAA' *.fastq > count/B402_Fm0.txt
agrep -0 -c 'ATAGTAGAAGCGCAGTT' *.fastq > count/B403_Fm0.txt
agrep -0 -c 'CATCTCTCACCGATATC' *.fastq > count/B404_Fm0.txt
agrep -0 -c 'TAGGGTACATAGGGGGC' *.fastq > count/B405_Fm0.txt
agrep -0 -c 'CATGTCATGAAGGGCGC' *.fastq > count/B407_Fm0.txt
agrep -0 -c 'CTGACATATCTTCGCCA' *.fastq > count/B408_Fm0.txt
agrep -0 -c 'GGCAATGCTTGCTGAGT' *.fastq > count/B409_Fm0.txt
agrep -0 -c 'ATTTCAGACCTACAGCC' *.fastq > count/B411_Fm0.txt
agrep -0 -c 'GTGCTAGCGCGTATGAA' *.fastq > count/B412_Fm0.txt
agrep -0 -c 'CTGCCGTTGACTACAGA' *.fastq > count/B413_Fm0.txt
agrep -0 -c 'ACGCGCCCTATAATCAA' *.fastq > count/B414_Fm0.txt
agrep -0 -c 'CATAACTACGCCAGCGA' *.fastq > count/B416_Fm0.txt
agrep -0 -c 'GACTCTATTTCGCCCCT' *.fastq > count/B417_Fm0.txt
agrep -0 -c 'TTAGATAGGACCGAGTG' *.fastq > count/B418_Fm0.txt
agrep -0 -c 'AACGCAACGTTAGAGTC' *.fastq > count/B419_Fm0.txt
agrep -0 -c 'CAGTGTGGCGAATATTC' *.fastq > count/B420_Fm0.txt
agrep -0 -c 'GAGAGCCTTGTCATGGT' *.fastq > count/B421_Fm0.txt
agrep -0 -c 'ATGATCCGTTGGACAGG' *.fastq > count/B422_Fm0.txt
agrep -0 -c 'GCCAATTCCCAAAATCG' *.fastq > count/B424_Fm0.txt
agrep -0 -c 'CAAGTGGTCCCTTCATC' *.fastq > count/B429_Fm0.txt
agrep -0 -c 'CAGTATCTCAGAATACG' *.fastq > count/B430_Fm0.txt
agrep -0 -c 'ACCTCGAAAATCCAGGT' *.fastq > count/B431_Fm0.txt
agrep -0 -c 'GTCGGGGGAGATTTTCC' *.fastq > count/B432_Fm0.txt
agrep -0 -c 'GCAAACGATTTTCGCTA' *.fastq > count/B433_Fm0.txt
agrep -0 -c 'ATGATCCTTCGTGTCGT' *.fastq > count/B434_Fm0.txt
agrep -0 -c 'CCGAACTAGTAGCTTCG' *.fastq > count/B435_Fm0.txt
agrep -0 -c 'TGAATGTGCGCCTCTCA' *.fastq > count/B436_Fm0.txt
agrep -0 -c 'TGTGGTCATGCTGTCGC' *.fastq > count/B437_Fm0.txt
agrep -0 -c 'GCCGGTAAGGTACCAAT' *.fastq > count/B438_Fm0.txt
agrep -0 -c 'CCAGCATTGTTCGCGAC' *.fastq > count/B439_Fm0.txt
agrep -0 -c 'AAGAGATATCGCGCTGA' *.fastq > count/B440_Fm0.txt
agrep -0 -c 'GAGGACGGCGAAAGTAT' *.fastq > count/B441_Fm0.txt
agrep -0 -c 'ATGAGAGCCACGTGGAG' *.fastq > count/B442_Fm0.txt
agrep -0 -c 'TACAGGCGGCACTTGCT' *.fastq > count/B443_Fm0.txt
agrep -0 -c 'ATGCACCTTAGCACTGT' *.fastq > count/B444_Fm0.txt
agrep -0 -c 'TAGGTTGTAAGACCATG' *.fastq > count/B445_Fm0.txt
agrep -0 -c 'TCGGCGTGTTTGCTCCT' *.fastq > count/B446_Fm0.txt
agrep -0 -c 'CTAACGGGACAGCGATA' *.fastq > count/B447_Fm0.txt
agrep -0 -c 'ATACGCAACACCACCAG' *.fastq > count/B448_Fm0.txt
agrep -0 -c 'GATATCAAAGGCCGGAG' *.fastq > count/B449_Fm0.txt
agrep -0 -c 'CACTGTACAGGGCGATT' *.fastq > count/B450_Fm0.txt
agrep -0 -c 'TACTACAGTCCGAACTC' *.fastq > count/B451_Fm0.txt
agrep -0 -c 'AGCTGCATTGCCGTGAA' *.fastq > count/B452_Fm0.txt
agrep -0 -c 'TGCGTTGGATCGGCAAG' *.fastq > count/B453_Fm0.txt
agrep -0 -c 'ACCGCAGGATAGAGCAA' *.fastq > count/B454_Fm0.txt
agrep -0 -c 'ACCTCCTAGTCGAGAGG' *.fastq > count/B455_Fm0.txt
agrep -0 -c 'CACAGTACCGAAGCAGT' *.fastq > count/B456_Fm0.txt
agrep -0 -c 'TCGCTTGAGTTGCGACT' *.fastq > count/B457_Fm0.txt
agrep -0 -c 'ATCTCGTCCAAGGTGTG' *.fastq > count/B458_Fm0.txt
agrep -0 -c 'TCACCCCACCCGGTTTA' *.fastq > count/B459_Fm0.txt
agrep -0 -c 'TAAGCCTAAACCACGCA' *.fastq > count/B460_Fm0.txt
agrep -0 -c 'ACCAGTTGGACCCCACA' *.fastq > count/B461_Fm0.txt
agrep -0 -c 'GTATCCACTCTTCCAAT' *.fastq > count/B462_Fm0.txt
agrep -0 -c 'TGCTACTTTTGAGGGCG' *.fastq > count/B463_Fm0.txt
agrep -0 -c 'GTATTGTCTCACCCAAA' *.fastq > count/B464_Fm0.txt
agrep -0 -c 'CCTGCTCGACACCAACT' *.fastq > count/B465_Fm0.txt
agrep -0 -c 'TCGAATTTGTGTCGGAA' *.fastq > count/B467_Fm0.txt
agrep -0 -c 'CGCACTCCCTCATAGAT' *.fastq > count/B468_Fm0.txt
agrep -0 -c 'TACAACCGGTTCGAAGT' *.fastq > count/B469_Fm0.txt
agrep -0 -c 'GGAACAGAGGGTGATGT' *.fastq > count/B471_Fm0.txt
agrep -0 -c 'GTGCTTCGTCGTTCGAA' *.fastq > count/B472_Fm0.txt
agrep -0 -c 'ACGATCATGGCTGGCTA' *.fastq > count/B473_Fm0.txt
agrep -0 -c 'GACTAGGTTCAGTGAGA' *.fastq > count/B474_Fm0.txt
agrep -0 -c 'ATCAACCCGTCGTCAGG' *.fastq > count/B475_Fm0.txt
agrep -0 -c 'CGGGACTGGAAGCTGTT' *.fastq > count/B476_Fm0.txt
agrep -0 -c 'AGATGAATCTCGTGACG' *.fastq > count/B477_Fm0.txt
agrep -0 -c 'GATTCACTGTTGACCAG' *.fastq > count/B478_Fm0.txt
agrep -0 -c 'GCGCGCCTAAGCATGTT' *.fastq > count/B479_Fm0.txt
agrep -0 -c 'TTAGCGAACTGTAAAGC' *.fastq > count/B482_Fm0.txt
agrep -0 -c 'AATTACCGGAGGAAGCT' *.fastq > count/B483_Fm0.txt
agrep -0 -c 'ATGCTATCAGAAGAGAC' *.fastq > count/B484_Fm0.txt
agrep -0 -c 'ATCCCGTATGCACCAAC' *.fastq > count/B485_Fm0.txt
agrep -0 -c 'GTTGTTCAAGATACGGG' *.fastq > count/B486_Fm0.txt
agrep -0 -c 'GGGGTGGTTCTACTAGA' *.fastq > count/B487_Fm0.txt
agrep -0 -c 'GTGAGCCTGTAGCATAC' *.fastq > count/B488_Fm0.txt
agrep -0 -c 'GCTAGGCAACCTCTGCT' *.fastq > count/B489_Fm0.txt
agrep -0 -c 'CTTCTGGCGCTGTTTCG' *.fastq > count/B490_Fm0.txt
agrep -0 -c 'ATCAAACTTAGGTCGGA' *.fastq > count/B491_Fm0.txt
agrep -0 -c 'TTCGGGACAAACGTTAA' *.fastq > count/B492_Fm0.txt
agrep -0 -c 'AGTAGGAACATCCTGGC' *.fastq > count/B493_Fm0.txt
agrep -0 -c 'CTAGAAGTTATGGCCTT' *.fastq > count/B494_Fm0.txt
agrep -0 -c 'CGAGTCTGTAGTTACTT' *.fastq > count/B495_Fm0.txt
agrep -0 -c 'CTTAACGCTAAGGAGTT' *.fastq > count/B496_Fm0.txt
agrep -0 -c 'CCAGTTTCACGGGTCTG' *.fastq > count/B497_Fm0.txt
agrep -0 -c 'GCAACGCCACAGTGTTA' *.fastq > count/B498_Fm0.txt
agrep -0 -c 'TGCCGAGCATTCTTGAC' *.fastq > count/B499_Fm0.txt
agrep -0 -c 'TCTCTAGGGTACTTGAG' *.fastq > count/B500_Fm0.txt
agrep -0 -c 'CACATTCGTCACTCACC' *.fastq > count/B501_Fm0.txt
agrep -0 -c 'GTTAGATGCTCATTGAC' *.fastq > count/B502_Fm0.txt
agrep -0 -c 'GATCGCAATAGGATTCC' *.fastq > count/B504_Fm0.txt
agrep -0 -c 'TATCTTGTGCACAAGGG' *.fastq > count/B505_Fm0.txt
agrep -0 -c 'CCTGAGTCTGTTGGAAA' *.fastq > count/B506_Fm0.txt
agrep -0 -c 'GTTGCTGCGCGTCAAGT' *.fastq > count/B507_Fm0.txt
agrep -0 -c 'CCACACTAAAGTCTTCT' *.fastq > count/B508_Fm0.txt
agrep -0 -c 'CACGGTTATAGGGAAGT' *.fastq > count/B510_Fm0.txt
agrep -0 -c 'TTAAGTGCCGAGGCTAC' *.fastq > count/B511_Fm0.txt
agrep -0 -c 'AGCCGACTGCATTCGCT' *.fastq > count/B512_Fm0.txt
agrep -0 -c 'GTTACCGTGCACTTGCT' *.fastq > count/B513_Fm0.txt
agrep -0 -c 'TACTTGAGCGGCGATGA' *.fastq > count/B514_Fm0.txt
agrep -0 -c 'CCGATTGGTGAGAACCT' *.fastq > count/B515_Fm0.txt
agrep -0 -c 'ACTATATGGTCCGTCGT' *.fastq > count/B517_Fm0.txt
agrep -0 -c 'ATCCTTCGGCGAAATTT' *.fastq > count/B518_Fm0.txt
agrep -0 -c 'GCTCGAAGTTAAACACT' *.fastq > count/B519_Fm0.txt
agrep -0 -c 'GATCCAGATTCCCGCAA' *.fastq > count/B520_Fm0.txt
agrep -0 -c 'GCCCCATCAGAAATTTA' *.fastq > count/B521_Fm0.txt
agrep -0 -c 'CTCCTACGGCTTATAAT' *.fastq > count/B522_Fm0.txt
agrep -0 -c 'GCTCGGTGGACAGAAGA' *.fastq > count/B523_Fm0.txt
agrep -0 -c 'GACTTGGGTATGTGTAC' *.fastq > count/B524_Fm0.txt
agrep -0 -c 'CCATGATACCGCATACG' *.fastq > count/B525_Fm0.txt
agrep -0 -c 'TGCGCGGAGAATCGGAT' *.fastq > count/B526_Fm0.txt
agrep -0 -c 'ACGAGGGTTCCTCTGTA' *.fastq > count/B527_Fm0.txt
agrep -0 -c 'GATCAGATCGAGGACTG' *.fastq > count/B528_Fm0.txt
agrep -0 -c 'AGCCGAAGCGGCATTAA' *.fastq > count/B529_Fm0.txt
agrep -0 -c 'ACGGTCGCTCATTGGAT' *.fastq > count/B530_Fm0.txt
agrep -0 -c 'AAGACGGAAAGTGGCAC' *.fastq > count/B533_Fm0.txt
agrep -0 -c 'TTAGGGCTGGCCACATT' *.fastq > count/B534_Fm0.txt
agrep -0 -c 'GTCACTAGGTTGAGGGG' *.fastq > count/B537_Fm0.txt
agrep -0 -c 'GACTTGCATCCCTGCTA' *.fastq > count/B538_Fm0.txt
agrep -0 -c 'CCGGAGAACTTTATTCG' *.fastq > count/B539_Fm0.txt
agrep -0 -c 'GTTATAGTTGGCAGTCG' *.fastq > count/B540_Fm0.txt
agrep -0 -c 'GCCAAATACGCTGTAAC' *.fastq > count/B541_Fm0.txt
agrep -0 -c 'CCTACAAATCAAAGGTG' *.fastq > count/B543_Fm0.txt
agrep -0 -c 'TTGTAGGACTCGTTTGC' *.fastq > count/B544_Fm0.txt
agrep -0 -c 'GCGTCGTAACACTATTC' *.fastq > count/B545_Fm0.txt
agrep -0 -c 'CAACCGGTTGAGTTCGG' *.fastq > count/B546_Fm0.txt
agrep -0 -c 'CATGGACGAATTCCGTA' *.fastq > count/B547_Fm0.txt
agrep -0 -c 'TGTGTATGCGTCGAAAA' *.fastq > count/B548_Fm0.txt
agrep -0 -c 'CACCTGCGTCAACAAAT' *.fastq > count/B549_Fm0.txt
agrep -0 -c 'ATCCGGTGGAGCTAAGT' *.fastq > count/B550_Fm0.txt
agrep -0 -c 'TACCTTGCGGTTGGAGG' *.fastq > count/B553_Fm0.txt
agrep -0 -c 'AAGCATACTGTCGCTGC' *.fastq > count/B554_Fm0.txt
agrep -0 -c 'CACGACGAATGTACGCG' *.fastq > count/B556_Fm0.txt
agrep -0 -c 'CAGTATCCAGTCCGTAC' *.fastq > count/B557_Fm0.txt
agrep -0 -c 'CGGAGTCACTTGCGCAT' *.fastq > count/B558_Fm0.txt
agrep -0 -c 'GTGTAGCAAGAACTCGA' *.fastq > count/B559_Fm0.txt
agrep -0 -c 'GAGTGCAAAAGCGACCT' *.fastq > count/B560_Fm0.txt
agrep -0 -c 'TTCAACACAGGGAGCGT' *.fastq > count/B561_Fm0.txt
agrep -0 -c 'CTAAGCCCCAAGGCAGA' *.fastq > count/B563_Fm0.txt
agrep -0 -c 'TAGTACTAGGCGGAATC' *.fastq > count/B564_Fm0.txt
agrep -0 -c 'CGGTGTGGTGCTCCTTT' *.fastq > count/B565_Fm0.txt
agrep -0 -c 'ATTAGGGAGGTGTTAGA' *.fastq > count/B567_Fm0.txt
agrep -0 -c 'ATTTACGCCACGCTTGG' *.fastq > count/B568_Fm0.txt
agrep -0 -c 'GGTGGCAGGTCCTATTT' *.fastq > count/B569_Fm0.txt
agrep -0 -c 'GCGTTCAAGCCATAAAG' *.fastq > count/B570_Fm0.txt
agrep -0 -c 'GATTAAGAGGGGCATCG' *.fastq > count/B571_Fm0.txt
agrep -0 -c 'GTGACTGCTAAATCCGG' *.fastq > count/B572_Fm0.txt
agrep -0 -c 'GTCCCGTCAAGATACGA' *.fastq > count/B573_Fm0.txt
agrep -0 -c 'CATGAACTTCCGAAAGC' *.fastq > count/B574_Fm0.txt
agrep -0 -c 'ACGCTAGAAGAGCAGAT' *.fastq > count/B575_Fm0.txt
agrep -0 -c 'GCTTCTACATCACGGAA' *.fastq > count/B576_Fm0.txt
agrep -0 -c 'TGTTAGATGATCACCAG' *.fastq > count/B577_Fm0.txt
agrep -0 -c 'ACTTAATCGCTCCCGCA' *.fastq > count/B579_Fm0.txt
agrep -0 -c 'ATTAACGGGCATCTTTG' *.fastq > count/B580_Fm0.txt
agrep -0 -c 'CCGGATAATCCATCATA' *.fastq > count/B581_Fm0.txt
agrep -0 -c 'TAAGATGCCGTTGTCCT' *.fastq > count/B582_Fm0.txt
agrep -0 -c 'TATCTGTAACCCATGGG' *.fastq > count/B583_Fm0.txt
agrep -0 -c 'CCAAATTCCCACCTACC' *.fastq > count/B584_Fm0.txt
agrep -0 -c 'TAAAGAAGACCCACAGT' *.fastq > count/B586_Fm0.txt
agrep -0 -c 'AGGAGGTTAAACCAAGC' *.fastq > count/B587_Fm0.txt
agrep -0 -c 'CCAGTTTGTGAATAACC' *.fastq > count/B588_Fm0.txt
agrep -0 -c 'TGTCCTACGATGGCGTT' *.fastq > count/B591_Fm0.txt
agrep -0 -c 'TGGCTGGCCTAAGTCTC' *.fastq > count/B592_Fm0.txt
agrep -0 -c 'AACTTCACCTATTGGCG' *.fastq > count/B593_Fm0.txt
agrep -0 -c 'TTCCAGTCTTAGTGGCA' *.fastq > count/B595_Fm0.txt
agrep -0 -c 'GGCGAGTGTGTCATCTG' *.fastq > count/B596_Fm0.txt
agrep -0 -c 'AGCGCTGAAAGCCATCA' *.fastq > count/B597_Fm0.txt
agrep -0 -c 'TAAGGGATTAGCAACGC' *.fastq > count/B598_Fm0.txt
agrep -0 -c 'TGAGTAAGGGTCCATAC' *.fastq > count/B599_Fm0.txt
agrep -0 -c 'ACTACCGAGAACCAGAG' *.fastq > count/B600_Fm0.txt
agrep -0 -c 'GGCATGTGAAAGATCTC' *.fastq > count/B601_Fm0.txt
agrep -0 -c 'ATTTCAGGGAGCACGCG' *.fastq > count/B602_Fm0.txt
agrep -0 -c 'ATTCCAAAGGCACAACT' *.fastq > count/B603_Fm0.txt
agrep -0 -c 'GCTGTACGGGATCATTC' *.fastq > count/B604_Fm0.txt
agrep -0 -c 'ATCTCTCTTTTGAGGGC' *.fastq > count/B605_Fm0.txt
agrep -0 -c 'ATCGAAATTTCCGGAGG' *.fastq > count/B606_Fm0.txt
agrep -0 -c 'GTCTACCTAGCACTCAT' *.fastq > count/B607_Fm0.txt
agrep -0 -c 'CTAAGTCGCAGATGTAT' *.fastq > count/B608_Fm0.txt
agrep -0 -c 'CTGACGATAAATGCTAG' *.fastq > count/B609_Fm0.txt
agrep -0 -c 'TTCACCTACTACCCTAA' *.fastq > count/B610_Fm0.txt
agrep -0 -c 'CCTTCTAAGTGCTACCA' *.fastq > count/B611_Fm0.txt
agrep -0 -c 'GATACATTACCAACCAG' *.fastq > count/B614_Fm0.txt
agrep -0 -c 'CTGCGATCCCTGACGAT' *.fastq > count/B615_Fm0.txt
agrep -0 -c 'ACACTATTACGCATTGC' *.fastq > count/B616_Fm0.txt
agrep -0 -c 'GCGGTAATAGCGTGTTT' *.fastq > count/B617_Fm0.txt
agrep -0 -c 'CACGACCGCATAGGTGT' *.fastq > count/B618_Fm0.txt
agrep -0 -c 'CTCTTGATCGACGGGCT' *.fastq > count/B619_Fm0.txt
agrep -0 -c 'AGTTGGGTAGATCACAG' *.fastq > count/B620_Fm0.txt
agrep -0 -c 'CATGTAGGATTCGTCAA' *.fastq > count/B621_Fm0.txt
agrep -0 -c 'TCGCTGTGTCTCGAATT' *.fastq > count/B622_Fm0.txt
agrep -0 -c 'ATGGGTAACACGAACGC' *.fastq > count/B623_Fm0.txt
agrep -0 -c 'TAGAACCTACGGCCTCT' *.fastq > count/B626_Fm0.txt
agrep -0 -c 'CTCGTGAGGAGACCATG' *.fastq > count/B627_Fm0.txt
agrep -0 -c 'AGGGTCATATGACTTTG' *.fastq > count/B628_Fm0.txt
agrep -0 -c 'CACTAAAGCAAGAACGT' *.fastq > count/B629_Fm0.txt
agrep -0 -c 'CATCGTGGTGCGAGTCA' *.fastq > count/B632_Fm0.txt
agrep -0 -c 'GTTGGACTGTGATTGCA' *.fastq > count/B633_Fm0.txt
agrep -0 -c 'CGGATACTGCGCACACT' *.fastq > count/B634_Fm0.txt
agrep -0 -c 'TGTTCGTTTCCTGGAAT' *.fastq > count/B636_Fm0.txt
agrep -0 -c 'CCACATGTTGACGAGCA' *.fastq > count/B637_Fm0.txt
agrep -0 -c 'GAGGAACGTAGGATTTT' *.fastq > count/B638_Fm0.txt
agrep -0 -c 'GATCCGAATCGCTTTGG' *.fastq > count/B639_Fm0.txt
agrep -0 -c 'TGGCGTGTGGTACGATG' *.fastq > count/B640_Fm0.txt
agrep -0 -c 'CCAACACCTACCGCTTT' *.fastq > count/B642_Fm0.txt
agrep -0 -c 'CGGGGTTGTTTTCGCAG' *.fastq > count/B645_Fm0.txt
agrep -0 -c 'CATTCATTGAAGAGGAG' *.fastq > count/B647_Fm0.txt
agrep -0 -c 'ATTGCTGACGCGCGTAA' *.fastq > count/B648_Fm0.txt
agrep -0 -c 'CTCTAGTCATATCCGCC' *.fastq > count/B649_Fm0.txt
agrep -0 -c 'CTAGTAGTATAACGGAG' *.fastq > count/B650_Fm0.txt
agrep -0 -c 'TGAAATAATTGGGCGTC' *.fastq > count/B651_Fm0.txt
agrep -0 -c 'GGTACTAGGTGCGTTTT' *.fastq > count/B652_Fm0.txt
agrep -0 -c 'AACAAGCTCTCGCCAAT' *.fastq > count/B653_Fm0.txt
agrep -0 -c 'GCGAAAGGGTCGGTTCT' *.fastq > count/B655_Fm0.txt
agrep -0 -c 'GGAATATGCGATACCAC' *.fastq > count/B657_Fm0.txt
agrep -0 -c 'AACGAGACTCGAAGCTT' *.fastq > count/B658_Fm0.txt
agrep -0 -c 'GCTCTACGTCGCCTCTA' *.fastq > count/B659_Fm0.txt
agrep -0 -c 'GGTTATCACTCTCACCG' *.fastq > count/B661_Fm0.txt
agrep -0 -c 'CACTTAAGACCCAAGCG' *.fastq > count/B663_Fm0.txt
agrep -0 -c 'TCGCGGCATGTCCTAAC' *.fastq > count/B664_Fm0.txt
agrep -0 -c 'TAACCAGTTACACGAGG' *.fastq > count/B666_Fm0.txt
agrep -0 -c 'AGAAGCTACATGATAGG' *.fastq > count/B667_Fm0.txt
agrep -0 -c 'TGACTAGCCTCGTCGTC' *.fastq > count/B669_Fm0.txt
agrep -0 -c 'TGAGTCGTGAACACGAG' *.fastq > count/B670_Fm0.txt
agrep -0 -c 'CTGCACTGAATAGTTAC' *.fastq > count/B671_Fm0.txt
agrep -0 -c 'AATTTGCGCCTTGTCGA' *.fastq > count/B672_Fm0.txt
agrep -0 -c 'TATTCGCTGTGCTCCAG' *.fastq > count/B673_Fm0.txt
agrep -0 -c 'GACTAATCTGGGTACGA' *.fastq > count/B674_Fm0.txt
agrep -0 -c 'CAGTACTACCTAGACGC' *.fastq > count/B675_Fm0.txt
agrep -0 -c 'CTGACTCGCAGAATGTT' *.fastq > count/B676_Fm0.txt
agrep -0 -c 'GGTACACACGCAATGTT' *.fastq > count/B677_Fm0.txt
agrep -0 -c 'AGTGTACAACTCTACGG' *.fastq > count/B678_Fm0.txt
agrep -0 -c 'GCCATTCAGACCGAGTC' *.fastq > count/B679_Fm0.txt
agrep -0 -c 'TGAACTCAGTGCTATAG' *.fastq > count/B680_Fm0.txt
agrep -0 -c 'ATCATCCTACTGCAGGG' *.fastq > count/B682_Fm0.txt
agrep -0 -c 'CCGAACTCAACTGGTTG' *.fastq > count/B683_Fm0.txt
agrep -0 -c 'CATGGCGTGAGACGATC' *.fastq > count/B686_Fm0.txt
agrep -0 -c 'GGCGGCGATCGTAATTA' *.fastq > count/B687_Fm0.txt
agrep -0 -c 'AAGTGCTGCACTTGGGG' *.fastq > count/B688_Fm0.txt
agrep -0 -c 'TATTTGACCGACGCCCG' *.fastq > count/B689_Fm0.txt
agrep -0 -c 'TCAATTCCCTGATAAGC' *.fastq > count/B690_Fm0.txt
agrep -0 -c 'TTGCAAACAGGAGGTAC' *.fastq > count/B691_Fm0.txt
agrep -0 -c 'CTTCGTCCAATCCGCGA' *.fastq > count/B692_Fm0.txt
agrep -0 -c 'CCACCGTGAGTAATTAT' *.fastq > count/B693_Fm0.txt
agrep -0 -c 'GGAACGAGTAGCTCAGG' *.fastq > count/B694_Fm0.txt
agrep -0 -c 'ATGCCGATAAGTAGGTT' *.fastq > count/B695_Fm0.txt
agrep -0 -c 'ACACTGACACATCACGA' *.fastq > count/B696_Fm0.txt
agrep -0 -c 'CACGATCGATAGGTTCA' *.fastq > count/B698_Fm0.txt
agrep -0 -c 'GTTCAAGCTTTGGAGTG' *.fastq > count/B699_Fm0.txt
agrep -0 -c 'CACTCGCTGTGTACTAG' *.fastq > count/B701_Fm0.txt
agrep -0 -c 'GCTACGGAATACACTGT' *.fastq > count/B702_Fm0.txt
agrep -0 -c 'CAAACGAACGATAACCT' *.fastq > count/B704_Fm0.txt
agrep -0 -c 'GCTCTAAGGTAGAAGTC' *.fastq > count/B705_Fm0.txt
agrep -0 -c 'TAGGTAGCATGATGTGT' *.fastq > count/B706_Fm0.txt
agrep -0 -c 'GCTTATCGTCGAAACAT' *.fastq > count/B707_Fm0.txt
agrep -0 -c 'ATGCGTACTACCCTGGT' *.fastq > count/B708_Fm0.txt
agrep -0 -c 'AAGTCGATTCACGGGCC' *.fastq > count/B709_Fm0.txt
agrep -0 -c 'GCGCGCTTATATGGATA' *.fastq > count/B710_Fm0.txt
agrep -0 -c 'AAGGTCAAACACCGTGA' *.fastq > count/B711_Fm0.txt
agrep -0 -c 'GTCTATACGTCAGATCA' *.fastq > count/B712_Fm0.txt
agrep -0 -c 'GCCAGACCCAGGTGTAA' *.fastq > count/B715_Fm0.txt
agrep -0 -c 'GATCGCGATGCGAGCAA' *.fastq > count/B717_Fm0.txt
agrep -0 -c 'TGAATGTCTAGATCCCT' *.fastq > count/B718_Fm0.txt
agrep -0 -c 'AGGAGGGTCTATCGTTA' *.fastq > count/B719_Fm0.txt
agrep -0 -c 'CCTACAGGCAACAGGAC' *.fastq > count/B720_Fm0.txt
agrep -0 -c 'TAGTGCTATTTCCATCG' *.fastq > count/B721_Fm0.txt
agrep -0 -c 'TCTTCCGGTTGACTGAT' *.fastq > count/B722_Fm0.txt
agrep -0 -c 'CGCCGAACGGACTCTAT' *.fastq > count/B723_Fm0.txt
agrep -0 -c 'AGAGGAAGTTAGGTCCG' *.fastq > count/B724_Fm0.txt
agrep -0 -c 'ATATCCGCCAGACCAAT' *.fastq > count/B725_Fm0.txt
agrep -0 -c 'AAGTTCTCGACTGTCTT' *.fastq > count/B726_Fm0.txt
agrep -0 -c 'CTGCTTCGACGGATACA' *.fastq > count/B727_Fm0.txt
agrep -0 -c 'GTACGGTACTCATTTGT' *.fastq > count/B728_Fm0.txt
agrep -0 -c 'GTAGCTCTCGTCGGTGA' *.fastq > count/B729_Fm0.txt
agrep -0 -c 'CGCCGGGTTTACTCTCA' *.fastq > count/B730_Fm0.txt
agrep -0 -c 'AAGCTATACAGCTTCCT' *.fastq > count/B731_Fm0.txt
agrep -0 -c 'TAACTGTGAGGCCTTGC' *.fastq > count/B732_Fm0.txt
agrep -0 -c 'TCAAAACTACAGCGTTG' *.fastq > count/B733_Fm0.txt
agrep -0 -c 'GGTGGCTCGATTGATAC' *.fastq > count/B734_Fm0.txt
agrep -0 -c 'AATACCACCGACGCTAG' *.fastq > count/B735_Fm0.txt
agrep -0 -c 'GGTCTCCTGTTCGTAGT' *.fastq > count/B736_Fm0.txt
agrep -0 -c 'ACTTTTGTATCGCGCCC' *.fastq > count/B737_Fm0.txt
agrep -0 -c 'ACCTTTATGCCAAGTGG' *.fastq > count/B738_Fm0.txt
agrep -0 -c 'GGGTTACAGCACACAGT' *.fastq > count/B739_Fm0.txt
agrep -0 -c 'TGGCTAGGATTCGTCTG' *.fastq > count/B740_Fm0.txt
agrep -0 -c 'GTCAGTCAAATGCCAAC' *.fastq > count/B741_Fm0.txt
agrep -0 -c 'GCAGCTGTCTTATCAAT' *.fastq > count/B742_Fm0.txt
agrep -0 -c 'CTCAAAAGCAAGTTCTG' *.fastq > count/B743_Fm0.txt
agrep -0 -c 'CGCTCAAGCGGATGTTC' *.fastq > count/B744_Fm0.txt
agrep -0 -c 'GTGTTAACGAGGATGCT' *.fastq > count/B745_Fm0.txt
agrep -0 -c 'GTACCTGCCCGTAGTGT' *.fastq > count/B746_Fm0.txt
agrep -0 -c 'ACTGATAGGAAACAGCA' *.fastq > count/B747_Fm0.txt
agrep -0 -c 'ATTACCAGTACAGCGCC' *.fastq > count/B748_Fm0.txt
agrep -0 -c 'ACGCTCTACGTCAAGGC' *.fastq > count/B749_Fm0.txt
agrep -0 -c 'TTGGTATGCCCCCGATA' *.fastq > count/B750_Fm0.txt
agrep -0 -c 'TCTACGTGCCTCACGCT' *.fastq > count/B752_Fm0.txt
agrep -0 -c 'TGGAGCTGGGCCAAAGT' *.fastq > count/B753_Fm0.txt
agrep -0 -c 'GTTCTCCGCGTAGACAA' *.fastq > count/B754_Fm0.txt
agrep -0 -c 'ATCGTTGCCGGGGATGT' *.fastq > count/B755_Fm0.txt
agrep -0 -c 'ACGTCAAAGCTGCTCAC' *.fastq > count/B756_Fm0.txt
agrep -0 -c 'ATTCGGTGGTTACCGTA' *.fastq > count/B758_Fm0.txt
agrep -0 -c 'GTAGGCACTGTAAAGTA' *.fastq > count/B759_Fm0.txt
agrep -0 -c 'GGCTAATCGACGACAGG' *.fastq > count/B760_Fm0.txt
agrep -0 -c 'GCACGTAACTCTCCCTC' *.fastq > count/B761_Fm0.txt
agrep -0 -c 'TAAAACTCCTCGCGCTG' *.fastq > count/B762_Fm0.txt
agrep -0 -c 'GCGAAAATCTCGGCAAC' *.fastq > count/B763_Fm0.txt
agrep -0 -c 'GGGACGATTGCCTTTAC' *.fastq > count/B765_Fm0.txt
agrep -0 -c 'GATACTTCCTTCCTACC' *.fastq > count/B767_Fm0.txt
agrep -0 -c 'AGCCGCCGTGCTAGTAA' *.fastq > count/B768_Fm0.txt
agrep -0 -c 'TTCCGATTCTGCGGGCT' *.fastq > count/B769_Fm0.txt
agrep -0 -c 'TGCTCACATTGCAGCCC' *.fastq > count/B770_Fm0.txt
agrep -0 -c 'TGGAAATAAGTCCTACG' *.fastq > count/B771_Fm0.txt
agrep -0 -c 'ATTCGCAAGAATCACGC' *.fastq > count/B772_Fm0.txt
agrep -0 -c 'ACGATAAACACTACCCC' *.fastq > count/B773_Fm0.txt
agrep -0 -c 'ACTCTCGAGTACTTCAT' *.fastq > count/B774_Fm0.txt
agrep -0 -c 'AGGCCCGCATTACCTAT' *.fastq > count/B775_Fm0.txt
agrep -0 -c 'CTTGCGAGGACTGGTCA' *.fastq > count/B776_Fm0.txt
agrep -0 -c 'TTAGAGCCGGGATTACT' *.fastq > count/B777_Fm0.txt
agrep -0 -c 'TTCCGTCGGTCCGTAAG' *.fastq > count/B778_Fm0.txt
agrep -0 -c 'CCAAGTATCCTCGCTAG' *.fastq > count/B779_Fm0.txt
agrep -0 -c 'ATGGCTCGGGACTCCTT' *.fastq > count/B781_Fm0.txt
agrep -0 -c 'CTGGATCTTATAGTGCT' *.fastq > count/B782_Fm0.txt
agrep -0 -c 'CGATCTTGACCTCTCAC' *.fastq > count/B783_Fm0.txt
agrep -0 -c 'TAGCGATGACCCCGAAG' *.fastq > count/B784_Fm0.txt
agrep -0 -c 'CAATTCCAAGCCATGCT' *.fastq > count/B785_Fm0.txt
agrep -0 -c 'GTTATCGGCCTGGCGAT' *.fastq > count/B787_Fm0.txt
agrep -0 -c 'AACATCCGACCCACGGA' *.fastq > count/B788_Fm0.txt
agrep -0 -c 'CATCCGCACCAGCGTAA' *.fastq > count/B789_Fm0.txt
agrep -0 -c 'GTTAGGACGTGTCACAT' *.fastq > count/B790_Fm0.txt
agrep -0 -c 'ACTCACTAAAGGGCATT' *.fastq > count/B791_Fm0.txt
agrep -0 -c 'ATGGACGTAATGCGGAT' *.fastq > count/B792_Fm0.txt
agrep -0 -c 'GTCTGTTAAGCCTAGGC' *.fastq > count/B793_Fm0.txt
agrep -0 -c 'CTGGGTGCTTGCACGTT' *.fastq > count/B795_Fm0.txt
agrep -0 -c 'AGTTCGAACACCCGCTG' *.fastq > count/B800_Fm0.txt
agrep -0 -c 'TAGGGAGGGATTAGGAC' *.fastq > count/B801_Fm0.txt
agrep -0 -c 'TTAGTCAAGGCAAACGG' *.fastq > count/B802_Fm0.txt
agrep -0 -c 'CCTCACTAGTCGAGGTG' *.fastq > count/B804_Fm0.txt
agrep -0 -c 'TATGGCCATATGAGAGT' *.fastq > count/B806_Fm0.txt
agrep -0 -c 'GACTCCTGTTCGTTTCA' *.fastq > count/B807_Fm0.txt
agrep -0 -c 'GTCTTCGTTCTAGGAAC' *.fastq > count/B808_Fm0.txt
agrep -0 -c 'TTGCAACCGATGTATCG' *.fastq > count/B809_Fm0.txt
agrep -0 -c 'GGTGTGTTTCAGACGCT' *.fastq > count/B812_Fm0.txt
agrep -0 -c 'ACTCTTAATCCGTGCAC' *.fastq > count/B813_Fm0.txt
agrep -0 -c 'CACCTGCAGGTAACGAT' *.fastq > count/B814_Fm0.txt
agrep -0 -c 'CCTCTTGGAATCTGCGT' *.fastq > count/B815_Fm0.txt
agrep -0 -c 'TGATCATCGCAAATCGT' *.fastq > count/B816_Fm0.txt
agrep -0 -c 'TTATTGGCCAGCGTATG' *.fastq > count/B817_Fm0.txt
agrep -0 -c 'CAAAGACGCCTCGCAAT' *.fastq > count/B818_Fm0.txt
agrep -0 -c 'GGTTTGCGATAGCTCGA' *.fastq > count/B819_Fm0.txt
agrep -0 -c 'ACACTCGTTTCTACATG' *.fastq > count/B821_Fm0.txt
agrep -0 -c 'ACTAGTTGGCTAGCGAA' *.fastq > count/B822_Fm0.txt
agrep -0 -c 'TGACTCGATGGGGCTCT' *.fastq > count/B823_Fm0.txt
agrep -0 -c 'AAGTTCAATTCGGGTTC' *.fastq > count/B824_Fm0.txt
agrep -0 -c 'GTTTTTGTGGAGTTGCT' *.fastq > count/B825_Fm0.txt
agrep -0 -c 'CCTGATCCTTGGTATTG' *.fastq > count/B826_Fm0.txt
agrep -0 -c 'TGTGAAGCCACCGCTTC' *.fastq > count/B828_Fm0.txt
agrep -0 -c 'ATAGTGTGGTCGGTTCT' *.fastq > count/B829_Fm0.txt
agrep -0 -c 'AAGACAACCGTCGCCGT' *.fastq > count/B830_Fm0.txt
agrep -0 -c 'CCACGGAGTACGTAATG' *.fastq > count/B832_Fm0.txt
agrep -0 -c 'TCAATGCGTAACACTGA' *.fastq > count/B833_Fm0.txt
agrep -0 -c 'AGCTGCGTCATGTGCTA' *.fastq > count/B834_Fm0.txt
agrep -0 -c 'TGACCCAGGTAGCTATG' *.fastq > count/B836_Fm0.txt
agrep -0 -c 'AGCCCTGACTAATACGA' *.fastq > count/B837_Fm0.txt
agrep -0 -c 'TGATCACCAGTACCCAT' *.fastq > count/B838_Fm0.txt
agrep -0 -c 'CATAAGGGCAGAGCATG' *.fastq > count/B839_Fm0.txt
agrep -0 -c 'AGCAAGAGGCCGAAGGA' *.fastq > count/B840_Fm0.txt
agrep -0 -c 'GAGGGGTCGAAGATAAG' *.fastq > count/B841_Fm0.txt
agrep -0 -c 'AGACGAGCAGTGCAGAA' *.fastq > count/B842_Fm0.txt
agrep -0 -c 'GTCCAGTGGTAGAACTT' *.fastq > count/B843_Fm0.txt
agrep -0 -c 'GACATGGTGGCATAGTA' *.fastq > count/B846_Fm0.txt
agrep -0 -c 'GACACGACAGGGATCAA' *.fastq > count/B848_Fm0.txt
agrep -0 -c 'TGTAGGCGCCATCTAGC' *.fastq > count/B850_Fm0.txt
agrep -0 -c 'GCGGTGAGAAGTTAATC' *.fastq > count/B852_Fm0.txt
agrep -0 -c 'CGCAGCGCTGGGTATAA' *.fastq > count/B853_Fm0.txt
agrep -0 -c 'CATCGAACACGATCTAG' *.fastq > count/B854_Fm0.txt
agrep -0 -c 'GTTGTATGTTGCTCAGT' *.fastq > count/B855_Fm0.txt
agrep -0 -c 'ACCATCGACGACAGACG' *.fastq > count/B856_Fm0.txt
agrep -0 -c 'TAAACAGAAACGCCACC' *.fastq > count/B857_Fm0.txt
agrep -0 -c 'GATAGATCAATAGCGCT' *.fastq > count/B858_Fm0.txt
agrep -0 -c 'GTCGTGGTGCTCATGAA' *.fastq > count/B859_Fm0.txt
agrep -0 -c 'TGCTTCCAACTATGGAG' *.fastq > count/B860_Fm0.txt
agrep -0 -c 'TCGTTTCCTTCTTACAG' *.fastq > count/B861_Fm0.txt
agrep -0 -c 'CGTAGTACCTGCGAACT' *.fastq > count/B862_Fm0.txt
agrep -0 -c 'TTGACCCCCCATGGAAA' *.fastq > count/B863_Fm0.txt
agrep -0 -c 'CAGACGCTCCGTTAAGT' *.fastq > count/B864_Fm0.txt
agrep -0 -c 'GGGGTCCGACTATCTTA' *.fastq > count/B865_Fm0.txt
agrep -0 -c 'TCTGGAAGAGACAAGTT' *.fastq > count/B866_Fm0.txt
agrep -0 -c 'AGAATAAGCTTCGTCCA' *.fastq > count/B867_Fm0.txt
agrep -0 -c 'GTCCCTAAGCATAGTGG' *.fastq > count/B868_Fm0.txt
agrep -0 -c 'CAAGCCACAAACTACGT' *.fastq > count/B869_Fm0.txt
agrep -0 -c 'GTTCAAGCGCTAGGGAT' *.fastq > count/B870_Fm0.txt
agrep -0 -c 'AGACTTGCCCACCGGTA' *.fastq > count/B871_Fm0.txt
agrep -0 -c 'TGCGGGCGGTTTATACC' *.fastq > count/B872_Fm0.txt
agrep -0 -c 'CTATTATACTACGCGCT' *.fastq > count/B873_Fm0.txt
agrep -0 -c 'CCAACTCTTTAGCCCGC' *.fastq > count/B875_Fm0.txt
agrep -0 -c 'AGAAAGTCACTCCGGCA' *.fastq > count/B877_Fm0.txt
agrep -0 -c 'AGTAACCAGCATTAGCC' *.fastq > count/B880_Fm0.txt
agrep -0 -c 'TACGTCGAGAGCAGGAA' *.fastq > count/B881_Fm0.txt
agrep -0 -c 'AGTTTCCAAAATCGCCC' *.fastq > count/B882_Fm0.txt
agrep -0 -c 'TCGGACTGTCAGTACAG' *.fastq > count/B883_Fm0.txt
agrep -0 -c 'GTCCACTACCGTCTGTC' *.fastq > count/B884_Fm0.txt
agrep -0 -c 'CGATATACTGCAGGTAG' *.fastq > count/B885_Fm0.txt
agrep -0 -c 'TGCGCTAGCCTAAATAT' *.fastq > count/B886_Fm0.txt
agrep -0 -c 'CGACAACTTCACGCTGG' *.fastq > count/B887_Fm0.txt
agrep -0 -c 'TCGTACTTCGTCAGTTT' *.fastq > count/B888_Fm0.txt
agrep -0 -c 'CATTACGGCCGCGATGA' *.fastq > count/B889_Fm0.txt
agrep -0 -c 'CTGCTTGCCTGGTTGAA' *.fastq > count/B890_Fm0.txt
agrep -0 -c 'AATCTCAGCGGATCGAC' *.fastq > count/B891_Fm0.txt
agrep -0 -c 'CGTTACAGCCACAGCAC' *.fastq > count/B892_Fm0.txt
agrep -0 -c 'TTAATCTAGGGACGGCG' *.fastq > count/B893_Fm0.txt
agrep -0 -c 'CAGTATATCGCCATGTA' *.fastq > count/B894_Fm0.txt
agrep -0 -c 'GCGACCCTTCACGTTTG' *.fastq > count/B895_Fm0.txt
agrep -0 -c 'AACTGTTCAATGCCCAC' *.fastq > count/B896_Fm0.txt
agrep -0 -c 'TAGGGGTCGGCATTCAA' *.fastq > count/B897_Fm0.txt
agrep -0 -c 'GCAAAAGGCTCAGTCCG' *.fastq > count/B898_Fm0.txt
agrep -0 -c 'ATCGTAGTATCGCTGCC' *.fastq > count/B899_Fm0.txt
agrep -0 -c 'TAATACGTGCCGACAGT' *.fastq > count/B900_Fm0.txt
agrep -0 -c 'CGTGATATCGCCGGTAT' *.fastq > count/B901_Fm0.txt
agrep -0 -c 'TGCAAGTTGCAGGGTTA' *.fastq > count/B902_Fm0.txt
agrep -0 -c 'AGCCCAAATAGTTCTTC' *.fastq > count/B903_Fm0.txt
agrep -0 -c 'CCTGGCAGAAATTGGCT' *.fastq > count/B905_Fm0.txt
agrep -0 -c 'GTTACCCGGTTCTAACT' *.fastq > count/B906_Fm0.txt
agrep -0 -c 'GCAACTGAACACCACAG' *.fastq > count/B907_Fm0.txt
agrep -0 -c 'TCAGGAGAGAGCGCTTT' *.fastq > count/B908_Fm0.txt
agrep -0 -c 'TGCGTGTTAAGAGCGTC' *.fastq > count/B909_Fm0.txt
agrep -0 -c 'GATAGCAACGTTGCCGC' *.fastq > count/B910_Fm0.txt
agrep -0 -c 'TCTACCTTTGGGCCGGT' *.fastq > count/B911_Fm0.txt
agrep -0 -c 'ATTCACTTGATGGGCCC' *.fastq > count/B912_Fm0.txt
agrep -0 -c 'TCTGTACGTTTGGCCTG' *.fastq > count/B913_Fm0.txt
agrep -0 -c 'GGAATTCGGAGAAAGAG' *.fastq > count/B915_Fm0.txt
agrep -0 -c 'TGGAAGACACGAAGCTA' *.fastq > count/B916_Fm0.txt
agrep -0 -c 'CGATACGATACAAAGAG' *.fastq > count/B918_Fm0.txt
agrep -0 -c 'CTCCCCGTCAATCTCCT' *.fastq > count/B919_Fm0.txt
agrep -0 -c 'TACCAGGCTTATTACGC' *.fastq > count/B920_Fm0.txt
agrep -0 -c 'GACGCTTAGTCCAGCTA' *.fastq > count/B921_Fm0.txt
agrep -0 -c 'AGTCTCACATAACTTCG' *.fastq > count/B922_Fm0.txt
agrep -0 -c 'TTGGAGGATTTATGCCT' *.fastq > count/B923_Fm0.txt
agrep -0 -c 'GTTATAGAGGGTACTAC' *.fastq > count/B924_Fm0.txt
agrep -0 -c 'ATGGCCGTATATCGCGA' *.fastq > count/B925_Fm0.txt
agrep -0 -c 'ATCTGGACTGGATTGGT' *.fastq > count/B926_Fm0.txt
agrep -0 -c 'GGGCGCACCTTGTTTAA' *.fastq > count/B927_Fm0.txt
agrep -0 -c 'GACGCGTATACTTCAGT' *.fastq > count/B928_Fm0.txt
agrep -0 -c 'GCTGGAGGTAACTTATT' *.fastq > count/B929_Fm0.txt
agrep -0 -c 'TCGCTACGGCGTGTAGT' *.fastq > count/B930_Fm0.txt
agrep -0 -c 'TGTGGGCGTGACTTTGA' *.fastq > count/B932_Fm0.txt
agrep -0 -c 'ACCCAGTTGTTTGTGTT' *.fastq > count/B933_Fm0.txt
agrep -0 -c 'CGCATCCTAGTAGGTAA' *.fastq > count/B934_Fm0.txt
agrep -0 -c 'AACTATCTCAACCGTTC' *.fastq > count/B935_Fm0.txt
agrep -0 -c 'CGCTTGGAGACAAATGC' *.fastq > count/B936_Fm0.txt
agrep -0 -c 'CTTCCAAATGGCGCATA' *.fastq > count/B938_Fm0.txt
agrep -0 -c 'GAGTAACTGACGGAGAC' *.fastq > count/B939_Fm0.txt
agrep -0 -c 'ATTGCGCCAAATTGGAG' *.fastq > count/B941_Fm0.txt
agrep -0 -c 'TGCGCAAACACCTTGAA' *.fastq > count/B942_Fm0.txt
agrep -0 -c 'ATATAGAGAGAACAGCC' *.fastq > count/B943_Fm0.txt
agrep -0 -c 'TATGAAGGATAAGCCCG' *.fastq > count/B945_Fm0.txt
agrep -0 -c 'CCATGCACGGTTCCAGT' *.fastq > count/B947_Fm0.txt
agrep -0 -c 'TACTATGCTCCAACCCC' *.fastq > count/B948_Fm0.txt
agrep -0 -c 'ATCCCATGGGTTTTAAG' *.fastq > count/B949_Fm0.txt
agrep -0 -c 'GATAGCAATAGCGACGA' *.fastq > count/B950_Fm0.txt
agrep -0 -c 'CGCAAACACAAAACGAA' *.fastq > count/B951_Fm0.txt
agrep -0 -c 'CCTGCTCATGTAGTTAG' *.fastq > count/B952_Fm0.txt
agrep -0 -c 'CGTTTAGTGCATGCGTC' *.fastq > count/B953_Fm0.txt
agrep -0 -c 'TGGACGTAGCTGGGTAC' *.fastq > count/B954_Fm0.txt
agrep -0 -c 'AAGGACTATCTTTCACC' *.fastq > count/B955_Fm0.txt
agrep -0 -c 'GATAGCTTGGTACGATC' *.fastq > count/B956_Fm0.txt
agrep -0 -c 'GCGGATATGTTTGCATT' *.fastq > count/B957_Fm0.txt
agrep -0 -c 'AATCCAAAGATCACGCG' *.fastq > count/B958_Fm0.txt
agrep -0 -c 'AGGCACAAGTTAGAGGC' *.fastq > count/B960_Fm0.txt
agrep -0 -c 'GAGGAACCCATATCGCG' *.fastq > count/B961_Fm0.txt
agrep -0 -c 'GGACATCACGCGGAATA' *.fastq > count/B962_Fm0.txt
agrep -0 -c 'GTTAGTCCAATCTGTGA' *.fastq > count/B963_Fm0.txt
agrep -0 -c 'GCTGACGAAAGAACCAG' *.fastq > count/B964_Fm0.txt
agrep -0 -c 'ACCCAGGTAAGTTTGAC' *.fastq > count/B965_Fm0.txt
agrep -0 -c 'GGGTAATTTAAGGGCTG' *.fastq > count/B966_Fm0.txt
agrep -0 -c 'CTACCGGGAGCGTACTA' *.fastq > count/B969_Fm0.txt
agrep -0 -c 'AGACAAGGCCGGGATGA' *.fastq > count/B971_Fm0.txt
agrep -0 -c 'GTGAAGAATCTTCTAGC' *.fastq > count/B972_Fm0.txt
agrep -0 -c 'GCGTCATTGTTGACGAC' *.fastq > count/B973_Fm0.txt
agrep -0 -c 'CTGTAACGTGTCATAAG' *.fastq > count/B974_Fm0.txt
agrep -0 -c 'GATACGCGCTACTCATC' *.fastq > count/B976_Fm0.txt
agrep -0 -c 'ACTACAACTTCGGCTAA' *.fastq > count/B977_Fm0.txt
agrep -0 -c 'CCTAGACCGCAATACTG' *.fastq > count/B979_Fm0.txt
agrep -0 -c 'GCTGTGGTAAGCATCAC' *.fastq > count/B982_Fm0.txt
agrep -0 -c 'TCTGGGTTTTGCTATGA' *.fastq > count/B983_Fm0.txt
agrep -0 -c 'ACTCAGCAATGTCACGC' *.fastq > count/B985_Fm0.txt
agrep -0 -c 'CTAACTGACCCAACTCA' *.fastq > count/B986_Fm0.txt
agrep -0 -c 'TTCCGTTTAAACTGCCC' *.fastq > count/B987_Fm0.txt
agrep -0 -c 'TCGGTAACGGTGCTTTA' *.fastq > count/B988_Fm0.txt
agrep -0 -c 'AGACCATAAAAGATGGG' *.fastq > count/B990_Fm0.txt
agrep -0 -c 'AATGTTCGATGTGAGAG' *.fastq > count/B991_Fm0.txt
agrep -0 -c 'AACGTAAGGAGAACCAA' *.fastq > count/B992_Fm0.txt
agrep -0 -c 'CCAGTGCTCTGTCAAAA' *.fastq > count/B993_Fm0.txt
agrep -0 -c 'GGTTCAGCGCGAATCTT' *.fastq > count/B994_Fm0.txt
agrep -0 -c 'CTCTCCTGAAAGACGTT' *.fastq > count/B995_Fm0.txt
agrep -0 -c 'GTTATGGTCCCTTACCC' *.fastq > count/B996_Fm0.txt
agrep -0 -c 'GTTGGAGTGTGAAGCAC' *.fastq > count/B997_Fm0.txt
agrep -0 -c 'TCTGCTTCGGACCAACA' *.fastq > count/B998_Fm0.txt
agrep -0 -c 'AGTTGGCGGGCTGTTTT' *.fastq > count/B999_Fm0.txt
agrep -0 -c 'CTGCTTTTATGACTGCT' *.fastq > count/B1000_Fm0.txt
agrep -0 -c 'CGGCCTAACGGGTTGAT' *.fastq > count/B1001_Fm0.txt
agrep -0 -c 'AATAGAGAAGGCCGTTG' *.fastq > count/B1002_Fm0.txt
agrep -0 -c 'CAAACAAGGGACACAAC' *.fastq > count/B1003_Fm0.txt
agrep -0 -c 'CTTCTGCACAGAACGTT' *.fastq > count/B1004_Fm0.txt
agrep -0 -c 'CTTAACCGACCGTTTCC' *.fastq > count/B1005_Fm0.txt
agrep -0 -c 'TGCCTAGCTAAGACTGG' *.fastq > count/B1006_Fm0.txt
agrep -0 -c 'CTCGGGCTCTGTTATGG' *.fastq > count/B1007_Fm0.txt
agrep -0 -c 'GCTCCCATATATGTGTT' *.fastq > count/B1010_Fm0.txt
agrep -0 -c 'GCCTAAGATCGTCGACA' *.fastq > count/B1012_Fm0.txt
agrep -0 -c 'CACCTTGAGTAATGAAC' *.fastq > count/B1013_Fm0.txt
agrep -0 -c 'ACTATCCGCCGTAACGC' *.fastq > count/B1014_Fm0.txt
agrep -0 -c 'AGCCCTGGTCCGGAATA' *.fastq > count/B1015_Fm0.txt
agrep -0 -c 'ATGGTTGCTCGCTTTTT' *.fastq > count/B1016_Fm0.txt
agrep -0 -c 'CTCGTATTTGAACTCTG' *.fastq > count/B1017_Fm0.txt
agrep -0 -c 'CAATCCTACGATCATGA' *.fastq > count/B1018_Fm0.txt
agrep -0 -c 'CAACTGTGTCAAACAGA' *.fastq > count/B1019_Fm0.txt
agrep -0 -c 'AAGACCGTGCACTACAG' *.fastq > count/B1020_Fm0.txt
agrep -0 -c 'CGCTTGGGTCCACCATA' *.fastq > count/B1023_Fm0.txt
agrep -0 -c 'ACTGGATCAAGCTCAGG' *.fastq > count/B1024_Fm0.txt
agrep -0 -c 'ATATGTGCACCGACAAG' *.fastq > count/B1025_Fm0.txt
agrep -0 -c 'TAGTCGTCCAGTTCGAT' *.fastq > count/B1026_Fm0.txt
agrep -0 -c 'TTCGGTCTTTTCGCTTG' *.fastq > count/B1027_Fm0.txt
agrep -0 -c 'CTCAAACCAAGTCCCTT' *.fastq > count/B1028_Fm0.txt
agrep -0 -c 'CACTACGTATCGAAGAC' *.fastq > count/B1029_Fm0.txt
agrep -0 -c 'GTGTATAGAGGGATGAC' *.fastq > count/B1032_Fm0.txt
agrep -0 -c 'GTCCTACTCCAGCCGTT' *.fastq > count/B1033_Fm0.txt
agrep -0 -c 'CTTCGTGCGCAAGAGTA' *.fastq > count/B1034_Fm0.txt
agrep -0 -c 'CAAGCATCAGTAGACCA' *.fastq > count/B1035_Fm0.txt
agrep -0 -c 'TGTGAACGAAGCAGATT' *.fastq > count/B1036_Fm0.txt
agrep -0 -c 'ATCGTCCTCGTTTGCAA' *.fastq > count/B1037_Fm0.txt
agrep -0 -c 'CACTGACGACATTGGCG' *.fastq > count/B1038_Fm0.txt
agrep -0 -c 'CACCTAATAACATGGGG' *.fastq > count/B1039_Fm0.txt
agrep -0 -c 'AGTTGTTAACGCCCAAC' *.fastq > count/B1040_Fm0.txt
agrep -0 -c 'TGCTAGCGATCCATGGA' *.fastq > count/B1041_Fm0.txt
agrep -0 -c 'CGTGCCGAGTATATATG' *.fastq > count/B1042_Fm0.txt
agrep -0 -c 'CGTTTCGCTCTAGATGA' *.fastq > count/B1045_Fm0.txt
agrep -0 -c 'AATACTACACAGTCAGG' *.fastq > count/B1046_Fm0.txt
agrep -0 -c 'GGCCTTCCACGCAATTT' *.fastq > count/B1047_Fm0.txt
agrep -0 -c 'GACGTAGTCGTGTTCTT' *.fastq > count/B1048_Fm0.txt
agrep -0 -c 'GTGCACTATCAAACGGG' *.fastq > count/B1049_Fm0.txt
agrep -0 -c 'CCTGAACATGACTGGGG' *.fastq > count/B1050_Fm0.txt
agrep -0 -c 'TCACCAGTAATTGTGCT' *.fastq > count/B1051_Fm0.txt
agrep -0 -c 'CGCCAATATAGATCGAT' *.fastq > count/B1052_Fm0.txt
agrep -0 -c 'GTGAGGCTAATCACGAG' *.fastq > count/B1054_Fm0.txt
agrep -0 -c 'CAGTTGAGGAGGTTCGC' *.fastq > count/B1056_Fm0.txt
agrep -0 -c 'ACCGTCGCTAACTATGT' *.fastq > count/B1057_Fm0.txt
agrep -0 -c 'GACATACACAGGACGGC' *.fastq > count/B1059_Fm0.txt
agrep -0 -c 'TACATATCTAGCCGCAA' *.fastq > count/B1061_Fm0.txt
agrep -0 -c 'TGGAAGCCAGTCCGCAA' *.fastq > count/B1062_Fm0.txt
agrep -0 -c 'CATACTACTCCCGCAAA' *.fastq > count/B1063_Fm0.txt
agrep -0 -c 'ACTACTCTCTTATGCGC' *.fastq > count/B1066_Fm0.txt
agrep -0 -c 'ACAACGAATGTCAAGCT' *.fastq > count/B1067_Fm0.txt
agrep -0 -c 'CCGTACAGGAAGGTTGC' *.fastq > count/B1068_Fm0.txt
agrep -0 -c 'GCCAGGAGTCCTATAAG' *.fastq > count/B1069_Fm0.txt
agrep -0 -c 'ATTGTTCCATGGTGCTT' *.fastq > count/B1070_Fm0.txt
agrep -0 -c 'TGCAACGATGGGTTAGC' *.fastq > count/B1071_Fm0.txt
agrep -0 -c 'GTTCCCGATATGACAGT' *.fastq > count/B1072_Fm0.txt
agrep -0 -c 'ATGGTATCCAGAAGGTA' *.fastq > count/B1074_Fm0.txt
agrep -0 -c 'ACCGCGGAAACAAATTT' *.fastq > count/B1075_Fm0.txt
agrep -0 -c 'GGCATATGTCTACACCT' *.fastq > count/B1076_Fm0.txt
agrep -0 -c 'TCGCCTAAATCATGCTC' *.fastq > count/B1078_Fm0.txt
agrep -0 -c 'GACCAACCGAACCATTT' *.fastq > count/B1080_Fm0.txt
agrep -0 -c 'GGCCATTGGCGTCTCTA' *.fastq > count/B1082_Fm0.txt
agrep -0 -c 'GTAAAGGACCTATGATC' *.fastq > count/B1083_Fm0.txt
agrep -0 -c 'ACCGAGGGCATCTCTAT' *.fastq > count/B1084_Fm0.txt
agrep -0 -c 'TTGACGAGGAGAATTCT' *.fastq > count/B1086_Fm0.txt
agrep -0 -c 'GCTGTGGCGTATTCGAA' *.fastq > count/B1087_Fm0.txt
agrep -0 -c 'TGACGCCAAGACAGGTA' *.fastq > count/B1088_Fm0.txt
agrep -0 -c 'TACTGAGCCGCCAGTTC' *.fastq > count/B1089_Fm0.txt
agrep -0 -c 'TGCGTGTCCGTTTGTAT' *.fastq > count/B1090_Fm0.txt
agrep -0 -c 'AGGGTAATCGCAAAGAC' *.fastq > count/B1092_Fm0.txt
agrep -0 -c 'CACGAATATGCGGCCTT' *.fastq > count/B1093_Fm0.txt
agrep -0 -c 'TTACATGGCATGATCGA' *.fastq > count/B1094_Fm0.txt
agrep -0 -c 'CGTAGTTGAACCTCTCC' *.fastq > count/B1095_Fm0.txt
agrep -0 -c 'TATTAGACCGGCGCTGT' *.fastq > count/B1097_Fm0.txt
agrep -0 -c 'GCGCTCGTTCGATTCTG' *.fastq > count/B1098_Fm0.txt
agrep -0 -c 'ATTATGGCGGAAACCAT' *.fastq > count/B1099_Fm0.txt
agrep -0 -c 'CTGGCTGTATGACGACG' *.fastq > count/B1102_Fm0.txt
agrep -0 -c 'CATCATAGTCAGTACTC' *.fastq > count/B1103_Fm0.txt
agrep -0 -c 'AACAATAGAGCAACGTC' *.fastq > count/B1104_Fm0.txt
agrep -0 -c 'TAATTTTCCCCTCCGCG' *.fastq > count/B1105_Fm0.txt
agrep -0 -c 'TGGAGTATTCAGGGAAG' *.fastq > count/B1106_Fm0.txt
agrep -0 -c 'AACCCCTGCGCTATGCA' *.fastq > count/B1107_Fm0.txt
agrep -0 -c 'GGAGAAGGCTATGGGTG' *.fastq > count/B1109_Fm0.txt
agrep -0 -c 'TATGCAGGGCGCGGAAT' *.fastq > count/B1111_Fm0.txt
agrep -0 -c 'CGGCAATTATGACCCAC' *.fastq > count/B1112_Fm0.txt
agrep -0 -c 'GCCATGTTTCATTACGC' *.fastq > count/B1113_Fm0.txt
agrep -0 -c 'AGTGCCCAAATCATCTG' *.fastq > count/B1114_Fm0.txt
agrep -0 -c 'TGCACTATCAGGGGTGC' *.fastq > count/B1116_Fm0.txt
agrep -0 -c 'AGGACAGCCACTAATTT' *.fastq > count/B1121_Fm0.txt
agrep -0 -c 'TTAGGCCGCCTAGGACT' *.fastq > count/B1122_Fm0.txt
agrep -0 -c 'GGAACAGATATTACGCT' *.fastq > count/B1123_Fm0.txt
agrep -0 -c 'CTGTTTACCCCCGAAAC' *.fastq > count/B1124_Fm0.txt
agrep -0 -c 'TAGCGACTCCGTTCAAA' *.fastq > count/B1125_Fm0.txt
agrep -0 -c 'TTGTGAGAACATCTACG' *.fastq > count/B1126_Fm0.txt
agrep -0 -c 'TCGAGTCGGTAAGACAA' *.fastq > count/B1127_Fm0.txt
agrep -0 -c 'GTGCCATCGCTACTTGA' *.fastq > count/B1129_Fm0.txt
agrep -0 -c 'TGGACGCGAACCGTTGA' *.fastq > count/B1130_Fm0.txt
agrep -0 -c 'GATAACGGCTGTTACAC' *.fastq > count/B1131_Fm0.txt
agrep -0 -c 'CGCTAGGTTCCATAGAA' *.fastq > count/B1132_Fm0.txt
agrep -0 -c 'CAGCCCTCAGGTTCGTT' *.fastq > count/B1135_Fm0.txt
agrep -0 -c 'ATATAGGGTCGACTGGG' *.fastq > count/B1137_Fm0.txt
agrep -0 -c 'TGCGTGAGCGATCTGTG' *.fastq > count/B1138_Fm0.txt
agrep -0 -c 'CCGGTTAATCACAACAT' *.fastq > count/B1139_Fm0.txt
agrep -0 -c 'CCATAAGATCGGCTTCC' *.fastq > count/B1140_Fm0.txt
agrep -0 -c 'TTACCATTGTCGCGATG' *.fastq > count/B1143_Fm0.txt
agrep -0 -c 'TATTTCCCGTCTCACAT' *.fastq > count/B1145_Fm0.txt
agrep -0 -c 'AGTTCACCCCCTTTGCT' *.fastq > count/B1146_Fm0.txt
agrep -0 -c 'ATCGTATGCCCCTGAAG' *.fastq > count/B1148_Fm0.txt
agrep -0 -c 'GGCCATAGAGGGCCTTT' *.fastq > count/B1149_Fm0.txt
agrep -0 -c 'TAGCAGTCCTCGGTTCG' *.fastq > count/B1150_Fm0.txt
agrep -0 -c 'ATCTATTACGGGGCTTA' *.fastq > count/B1151_Fm0.txt
agrep -0 -c 'AGAAGCTTAAGTCAAGC' *.fastq > count/B1152_Fm0.txt
agrep -0 -c 'GCGACTTTATGCTGACT' *.fastq > count/B1153_Fm0.txt
agrep -0 -c 'AGCTAAGGACGAAGGCC' *.fastq > count/B1154_Fm0.txt
agrep -0 -c 'CTAGGTGTTCAGTTCGT' *.fastq > count/B1155_Fm0.txt
agrep -0 -c 'CACGTAGGGCTCGAAAA' *.fastq > count/B1156_Fm0.txt
agrep -0 -c 'ATACAGACTATAGCGAC' *.fastq > count/B1158_Fm0.txt
agrep -0 -c 'CTCCCGGTAGGGTTAGA' *.fastq > count/B1159_Fm0.txt
agrep -0 -c 'TAATACATACGAGGGCC' *.fastq > count/B1160_Fm0.txt
agrep -0 -c 'TCGTAGACGATGGTTAT' *.fastq > count/B1161_Fm0.txt
agrep -0 -c 'CAACTTCCATTATCCTC' *.fastq > count/B1162_Fm0.txt
agrep -0 -c 'CAGATGTTCATAGTGGT' *.fastq > count/B1163_Fm0.txt
agrep -0 -c 'GAGACATGACTGCCCAT' *.fastq > count/B1164_Fm0.txt
agrep -0 -c 'ACGGGGGACATCCAAAC' *.fastq > count/B1165_Fm0.txt
agrep -0 -c 'AGCATGATGGCTGTTGC' *.fastq > count/B1167_Fm0.txt
agrep -0 -c 'ACCATGTAGACGCGCCT' *.fastq > count/B1168_Fm0.txt
agrep -0 -c 'GTTAGCGTGATCTTGTC' *.fastq > count/B1169_Fm0.txt
agrep -0 -c 'CTACGAACTATGGGAAG' *.fastq > count/B1170_Fm0.txt
agrep -0 -c 'TACTAAACTGCCCTTGA' *.fastq > count/B1171_Fm0.txt
agrep -0 -c 'CTCTCTGAACGCCTAAA' *.fastq > count/B1172_Fm0.txt
agrep -0 -c 'GCCACAGCGATTCTTAA' *.fastq > count/B1174_Fm0.txt
agrep -0 -c 'GCGTCTAGTCGTGATCT' *.fastq > count/B1175_Fm0.txt
agrep -0 -c 'GTCAAGAGAACCACAAT' *.fastq > count/B1176_Fm0.txt
agrep -0 -c 'AGACGGCACGCAGTTTG' *.fastq > count/B1177_Fm0.txt
agrep -0 -c 'CGCGGATGTCAAGCATT' *.fastq > count/B1178_Fm0.txt
agrep -0 -c 'GGACCATAAATCTCGCG' *.fastq > count/B1180_Fm0.txt
agrep -0 -c 'AAGTGCTCCTCCCCTAC' *.fastq > count/B1181_Fm0.txt
agrep -0 -c 'CATCTCCTATCCCTTAC' *.fastq > count/B1183_Fm0.txt
agrep -0 -c 'CATGATTCAAGCTTACC' *.fastq > count/B1184_Fm0.txt
agrep -0 -c 'TATCAGCGTACGAGAGG' *.fastq > count/B1185_Fm0.txt
agrep -0 -c 'CATAGAGCCGATACGGC' *.fastq > count/B1187_Fm0.txt
agrep -0 -c 'TCGAGCCTTGGTTAGCG' *.fastq > count/B1188_Fm0.txt
agrep -0 -c 'CAGTCTATCCGGCTGTG' *.fastq > count/B1189_Fm0.txt
agrep -0 -c 'AGTCAGTCGTGTCCGCA' *.fastq > count/B1190_Fm0.txt
agrep -0 -c 'CCATTATTCAGTGCCAG' *.fastq > count/B1192_Fm0.txt
agrep -0 -c 'CGGCTTGTGGGAACAGA' *.fastq > count/B1193_Fm0.txt
agrep -0 -c 'CGGTGCTTTGCTAAGGT' *.fastq > count/B1195_Fm0.txt
agrep -0 -c 'GATAGGGGCGGTTGAAA' *.fastq > count/B1198_Fm0.txt
agrep -0 -c 'CAATCGTTCCTGGGTGA' *.fastq > count/B1199_Fm0.txt
agrep -0 -c 'ACGTTCTGTCGCAACAG' *.fastq > count/B1203_Fm0.txt
agrep -0 -c 'GATTAGAACACGGGAAG' *.fastq > count/B1204_Fm0.txt
agrep -0 -c 'GCTGCGATCCAGAATAC' *.fastq > count/B1206_Fm0.txt
agrep -0 -c 'CACCTAGCGACCTGTGA' *.fastq > count/B1207_Fm0.txt
agrep -0 -c 'TGTTCATACAGTGTGCC' *.fastq > count/B1208_Fm0.txt
agrep -0 -c 'CTTAACCTCCGCCATTA' *.fastq > count/B1211_Fm0.txt
agrep -0 -c 'ATTCCTTTATTACGGCC' *.fastq > count/B1212_Fm0.txt
agrep -0 -c 'TGACACGACCTGAATGA' *.fastq > count/B1213_Fm0.txt
agrep -0 -c 'ACGTTGTAAAATCGGCA' *.fastq > count/B1214_Fm0.txt
agrep -0 -c 'CGCCGAGTATACCTCTT' *.fastq > count/B1216_Fm0.txt
agrep -0 -c 'GATACGAGCTAGAAACC' *.fastq > count/B1217_Fm0.txt
agrep -0 -c 'GTTACAAGACGGTCCTG' *.fastq > count/B1218_Fm0.txt
agrep -0 -c 'AATCGTCTACGCCATAA' *.fastq > count/B1219_Fm0.txt
agrep -0 -c 'ATGACGTGGTGAATGTG' *.fastq > count/B1222_Fm0.txt
agrep -0 -c 'AGGACAAATGCAGACGT' *.fastq > count/B1223_Fm0.txt
agrep -0 -c 'CGTGGAAGACTATCCTC' *.fastq > count/B1224_Fm0.txt
agrep -0 -c 'ACTCTCTCTCGAATCAT' *.fastq > count/B1225_Fm0.txt
agrep -0 -c 'CTAAGCTATACCGAAGC' *.fastq > count/B1226_Fm0.txt
agrep -0 -c 'ATCTGAGGGCACAGACT' *.fastq > count/B1227_Fm0.txt
agrep -0 -c 'AACAAGTGTCAGTACTC' *.fastq > count/B1231_Fm0.txt
agrep -0 -c 'AGCGAAGAGCATATGGG' *.fastq > count/B1232_Fm0.txt
agrep -0 -c 'AGGGCATACGCCGAACA' *.fastq > count/B1233_Fm0.txt
agrep -0 -c 'AGTATTCAGTCGCATAC' *.fastq > count/B1234_Fm0.txt
agrep -0 -c 'CCTATGTGTCAAGTTGT' *.fastq > count/B1235_Fm0.txt
agrep -0 -c 'AATAAGCGACCTCTCTT' *.fastq > count/B1237_Fm0.txt
agrep -0 -c 'CAGCTCGTTGTACGTCT' *.fastq > count/B1238_Fm0.txt
agrep -0 -c 'GCCAACGAATCTCCTTA' *.fastq > count/B1239_Fm0.txt
agrep -0 -c 'TAACGGCACAGGGATGG' *.fastq > count/B1240_Fm0.txt
agrep -0 -c 'ATGCGACCATACTGTGT' *.fastq > count/B1241_Fm0.txt
agrep -0 -c 'ACGGGATTGAGTACCTG' *.fastq > count/B1243_Fm0.txt
agrep -0 -c 'ATCTGGGGGGAGTCCTA' *.fastq > count/B1244_Fm0.txt
agrep -0 -c 'GGGATCTCACGTAGAAG' *.fastq > count/B1245_Fm0.txt
agrep -0 -c 'AATCGGAGCCTGCGCAT' *.fastq > count/B1246_Fm0.txt
agrep -0 -c 'GCACCTTAGCGATCGAC' *.fastq > count/B1247_Fm0.txt
agrep -0 -c 'TCGGTTACGAACTGATA' *.fastq > count/B1248_Fm0.txt
agrep -0 -c 'GCTTCAACTCGTGGAAC' *.fastq > count/B1250_Fm0.txt
agrep -0 -c 'TAAGTGCTAAGTTCCCT' *.fastq > count/B1251_Fm0.txt
agrep -0 -c 'ATGGTCAAAACTGGGAG' *.fastq > count/B1252_Fm0.txt
agrep -0 -c 'TTCCGTCACTGAGTCGC' *.fastq > count/B1253_Fm0.txt
agrep -0 -c 'CCTCATATAGGATCGGA' *.fastq > count/B1254_Fm0.txt
agrep -0 -c 'ATAGTGGTTGACTCACC' *.fastq > count/B1255_Fm0.txt
agrep -0 -c 'CAGGAACTTCTGCGAGT' *.fastq > count/B1256_Fm0.txt
agrep -0 -c 'GACGGTTTCTGGCATAA' *.fastq > count/B1257_Fm0.txt
agrep -0 -c 'CACTTGTGGTACGTAGC' *.fastq > count/B1258_Fm0.txt
agrep -0 -c 'TAAAGTCGCCGATCAAC' *.fastq > count/B1259_Fm0.txt
agrep -0 -c 'CTCCGATTTGAAATGGA' *.fastq > count/B1260_Fm0.txt
agrep -0 -c 'CACTGCATGGGAGTTAG' *.fastq > count/B1261_Fm0.txt
agrep -0 -c 'ACAGATGCAATCGCTTA' *.fastq > count/B1263_Fm0.txt
agrep -0 -c 'GGGGGTAGAAGTTCAGG' *.fastq > count/B1264_Fm0.txt
agrep -0 -c 'GTCAGATATGTGCCATT' *.fastq > count/B1266_Fm0.txt
agrep -0 -c 'GGTGATAGGCCAGCATG' *.fastq > count/B1267_Fm0.txt
agrep -0 -c 'ACTCTGTTCCGGTCCGT' *.fastq > count/B1268_Fm0.txt
agrep -0 -c 'CACTCACCGTTGACCAT' *.fastq > count/B1270_Fm0.txt
agrep -0 -c 'TGATGTTGGGACGTAAG' *.fastq > count/B1272_Fm0.txt
agrep -0 -c 'CGGTAATGCAGTGGTTT' *.fastq > count/B1273_Fm0.txt
agrep -0 -c 'CGAGCAAACCTATATCA' *.fastq > count/B1274_Fm0.txt
agrep -0 -c 'CTAGTACGCTCGTTCAC' *.fastq > count/B1275_Fm0.txt
agrep -0 -c 'CAACTACAGCATTTACG' *.fastq > count/B1277_Fm0.txt
agrep -0 -c 'TATCACCAAGAGTGACC' *.fastq > count/B1278_Fm0.txt
agrep -0 -c 'GTGGGTAAAATTCCGAA' *.fastq > count/B1280_Fm0.txt
agrep -0 -c 'TGCGAGGGGAGATTATG' *.fastq > count/B1281_Fm0.txt
agrep -0 -c 'AGGAACTTGACGATAGA' *.fastq > count/B1283_Fm0.txt
agrep -0 -c 'TGTTGGAATTGGATGCA' *.fastq > count/B1284_Fm0.txt
agrep -0 -c 'TACACGAGATGCATATC' *.fastq > count/B1285_Fm0.txt
agrep -0 -c 'GAGATACTCGAGTTACA' *.fastq > count/B1286_Fm0.txt
agrep -0 -c 'GCGACTAACCAGGTCAA' *.fastq > count/B1287_Fm0.txt
agrep -0 -c 'TATCTACCCCATCCTCC' *.fastq > count/B1288_Fm0.txt
agrep -0 -c 'CTCTTGGCTTGTTCGGT' *.fastq > count/B1289_Fm0.txt
agrep -0 -c 'GCGTTGTCACCCCTTTA' *.fastq > count/B1290_Fm0.txt
agrep -0 -c 'CTGGATAACGCATTAGT' *.fastq > count/B1291_Fm0.txt
agrep -0 -c 'TACCGTGCATTCCACCT' *.fastq > count/B1292_Fm0.txt
agrep -0 -c 'CTCCTCGTCATGACACA' *.fastq > count/B1293_Fm0.txt
agrep -0 -c 'TTGAACAGGCCGAATCT' *.fastq > count/B1294_Fm0.txt
agrep -0 -c 'TTAAAAGCCCGTTACCT' *.fastq > count/B1295_Fm0.txt
agrep -0 -c 'TTGTCTACTTCAGGGCG' *.fastq > count/B1296_Fm0.txt
agrep -0 -c 'CGTTATATCGTAGGGTT' *.fastq > count/B1297_Fm0.txt
agrep -0 -c 'CTGCGTTCACGTGGCTA' *.fastq > count/B1298_Fm0.txt
agrep -0 -c 'GGTTAGGGCATACTTAC' *.fastq > count/B1299_Fm0.txt
agrep -0 -c 'CCTACAATTTGATACGG' *.fastq > count/B1300_Fm0.txt
agrep -0 -c 'AGGATAACCCAAAGCCA' *.fastq > count/B1301_Fm0.txt
agrep -0 -c 'TGCAGATTCCCGCCAGA' *.fastq > count/B1302_Fm0.txt
agrep -0 -c 'GATGAGTCGTCTCAGAA' *.fastq > count/B1303_Fm0.txt
agrep -0 -c 'CTAGCCTGTTTTTGCAG' *.fastq > count/B1305_Fm0.txt
agrep -0 -c 'TCGTGTCATTGTCCACT' *.fastq > count/B1306_Fm0.txt
agrep -0 -c 'GCGACATCTCATTACTC' *.fastq > count/B1307_Fm0.txt
agrep -0 -c 'GTGATAGGGCACACACT' *.fastq > count/B1308_Fm0.txt
agrep -0 -c 'GTAGAGAGATCGAGTCA' *.fastq > count/B1309_Fm0.txt
agrep -0 -c 'ACGATATAAGTCGACCC' *.fastq > count/B1310_Fm0.txt
agrep -0 -c 'CACACATTCGAAAGGCA' *.fastq > count/B1311_Fm0.txt
agrep -0 -c 'TGACCCATTCGAATAGC' *.fastq > count/B1312_Fm0.txt
agrep -0 -c 'ACGGGAAAGTGTAACTA' *.fastq > count/B1314_Fm0.txt
agrep -0 -c 'GCTAGGGAGCAGTTGGT' *.fastq > count/B1315_Fm0.txt
agrep -0 -c 'CAGCGTATGTTGAACTT' *.fastq > count/B1316_Fm0.txt
agrep -0 -c 'TAAGATAATCGCTCGGC' *.fastq > count/B1317_Fm0.txt
agrep -0 -c 'ACTATCGGATTGTACAG' *.fastq > count/B1319_Fm0.txt
agrep -0 -c 'TATTTGCACGGCTCCCA' *.fastq > count/B1321_Fm0.txt
agrep -0 -c 'TTATACGTCTGTTCGAG' *.fastq > count/B1322_Fm0.txt
agrep -0 -c 'CAATCCTCATGTCCCCC' *.fastq > count/B1323_Fm0.txt
agrep -0 -c 'AAGTGGTGCTGAAGGCC' *.fastq > count/B1324_Fm0.txt
agrep -0 -c 'ATGTGTCCTCATGGGGG' *.fastq > count/B1325_Fm0.txt
agrep -0 -c 'TGCATCCCGCATAGAGG' *.fastq > count/B1326_Fm0.txt
agrep -0 -c 'ATACGCTGGACCAGCGT' *.fastq > count/B1327_Fm0.txt
agrep -0 -c 'CGCAAGCATATAGACGT' *.fastq > count/B1328_Fm0.txt
agrep -0 -c 'AATCTAAGCGCCCAGGA' *.fastq > count/B1329_Fm0.txt
agrep -0 -c 'TGCTTTAACTTGGTTCC' *.fastq > count/B1330_Fm0.txt
agrep -0 -c 'CGTCTCTTTAAACGCGG' *.fastq > count/B1332_Fm0.txt
agrep -0 -c 'ACATCGCCAATCCTAAA' *.fastq > count/B1333_Fm0.txt
agrep -0 -c 'CTGTAATACGACAACGA' *.fastq > count/B1336_Fm0.txt
agrep -0 -c 'TGTACCCGAATCACAAC' *.fastq > count/B1338_Fm0.txt
agrep -0 -c 'AAGTAGTCCGGTTGATA' *.fastq > count/B1339_Fm0.txt
agrep -0 -c 'ACCGATCAAGCAGCCTC' *.fastq > count/B1340_Fm0.txt
agrep -0 -c 'CAACGATCGTCTATATC' *.fastq > count/B1342_Fm0.txt
agrep -0 -c 'ACCAGCCACTTAACCCT' *.fastq > count/B1343_Fm0.txt
agrep -0 -c 'CGCACAGAATCAAGAAC' *.fastq > count/B1344_Fm0.txt
agrep -0 -c 'TTAGATTAGCCCCCTTT' *.fastq > count/B1345_Fm0.txt
agrep -0 -c 'TATGTGGGGAAAACTCG' *.fastq > count/B1346_Fm0.txt
agrep -0 -c 'TGACATTCGACTTTGAC' *.fastq > count/B1347_Fm0.txt
agrep -0 -c 'CTCTCAACTCCTACTCG' *.fastq > count/B1348_Fm0.txt
agrep -0 -c 'ACCTATCACAGGTCTAT' *.fastq > count/B1349_Fm0.txt
agrep -0 -c 'ATGCTCGGTGCGTTTAT' *.fastq > count/B1350_Fm0.txt
agrep -0 -c 'GGTCGCGAACTGGACAT' *.fastq > count/B1351_Fm0.txt
agrep -0 -c 'AGGGACGTTAACTTCAG' *.fastq > count/B1352_Fm0.txt
agrep -0 -c 'AAGCGATTAGGCCCAGT' *.fastq > count/B1353_Fm0.txt
agrep -0 -c 'TACAAATCATGGCCTCC' *.fastq > count/B1355_Fm0.txt
agrep -0 -c 'CGCAAACTATATCGTGG' *.fastq > count/B1357_Fm0.txt
agrep -0 -c 'TGGCCGCTGAGTAGAAT' *.fastq > count/B1358_Fm0.txt
agrep -0 -c 'CACACAGGGTTAAGCTG' *.fastq > count/B1359_Fm0.txt
agrep -0 -c 'CCTCCTTATAGCAGGCC' *.fastq > count/B1361_Fm0.txt
agrep -0 -c 'TAGTGGCGGATCAGACA' *.fastq > count/B1362_Fm0.txt
agrep -0 -c 'CGTAATACGGGGATACG' *.fastq > count/B1363_Fm0.txt
agrep -0 -c 'ACGGCATGTTCTTATGG' *.fastq > count/B1365_Fm0.txt
agrep -0 -c 'AGGCACGTCAGAAACTG' *.fastq > count/B1366_Fm0.txt
agrep -0 -c 'CGCTGAGATTGCGACTC' *.fastq > count/B1367_Fm0.txt
agrep -0 -c 'TCGACGAGGACAGAGTG' *.fastq > count/B1368_Fm0.txt
agrep -0 -c 'CGGTTATCACTTTTGAG' *.fastq > count/B1369_Fm0.txt
agrep -0 -c 'AGGACCAAGGAGCTGAT' *.fastq > count/B1370_Fm0.txt
agrep -0 -c 'TAGTGTCTGGGGATATA' *.fastq > count/B1371_Fm0.txt
agrep -0 -c 'CGATACAACCTAACACG' *.fastq > count/B1374_Fm0.txt
agrep -0 -c 'GTGAGTTACGAATCGAA' *.fastq > count/B1375_Fm0.txt
agrep -0 -c 'GTTCTCCGCTGATAAAC' *.fastq > count/B1378_Fm0.txt
agrep -0 -c 'CATCTCGTGGTGTCTAG' *.fastq > count/B1380_Fm0.txt
agrep -0 -c 'GACCCTGAGCTGCTAAA' *.fastq > count/B1381_Fm0.txt
agrep -0 -c 'GCGCAATGAACTGCTCA' *.fastq > count/B1383_Fm0.txt
agrep -0 -c 'TCTGTAGGCGGTAGATT' *.fastq > count/B1384_Fm0.txt
agrep -0 -c 'CAAGAGGGACGAAGCAT' *.fastq > count/B1385_Fm0.txt
agrep -0 -c 'CACGCGATCCGTTACGA' *.fastq > count/B1386_Fm0.txt
agrep -0 -c 'ATAGGAAGTCTTATGGC' *.fastq > count/B1387_Fm0.txt
agrep -0 -c 'GGGCCTATAAAGTTCCA' *.fastq > count/B1388_Fm0.txt
agrep -0 -c 'GTGGAGGGTAACTTAAA' *.fastq > count/B1389_Fm0.txt
agrep -0 -c 'GTCTGAGACACCTCACC' *.fastq > count/B1391_Fm0.txt
agrep -0 -c 'GGCAGTAAGCTCGCAAT' *.fastq > count/B1392_Fm0.txt
agrep -0 -c 'GTACCCGGTGCCTTATG' *.fastq > count/B1393_Fm0.txt
agrep -0 -c 'GCATTCCATCGCTGCTG' *.fastq > count/B1394_Fm0.txt
agrep -0 -c 'AGTTTGGCACGAAGGCT' *.fastq > count/B1395_Fm0.txt
agrep -0 -c 'GTCACGGTCGCATGATC' *.fastq > count/B1397_Fm0.txt
agrep -0 -c 'TCTGAGTGGTCGGACAC' *.fastq > count/B1398_Fm0.txt
agrep -0 -c 'GGACCCATAGTTTTGAA' *.fastq > count/B1399_Fm0.txt
agrep -0 -c 'TAAAAGGGCAACCAACC' *.fastq > count/B1400_Fm0.txt
agrep -0 -c 'CAATCCACCTAGTTTGC' *.fastq > count/B1401_Fm0.txt
agrep -0 -c 'CTGCTACACAGTTGCCA' *.fastq > count/B1402_Fm0.txt
agrep -0 -c 'ATAGAGGTGGAGTCTAC' *.fastq > count/B1403_Fm0.txt
agrep -0 -c 'GCATTACTCTCCGAAAT' *.fastq > count/B1404_Fm0.txt
agrep -0 -c 'CGTTTAGGGACAATTGC' *.fastq > count/B1405_Fm0.txt
agrep -0 -c 'TCGCTGTCCTAAGCGGA' *.fastq > count/B1406_Fm0.txt
agrep -0 -c 'CTAAGAGGGCACAACAG' *.fastq > count/B1407_Fm0.txt
agrep -0 -c 'CCTAAACGCCCTATGCC' *.fastq > count/B1408_Fm0.txt
agrep -0 -c 'AGCTGGTGATCTGTAGG' *.fastq > count/B1409_Fm0.txt
agrep -0 -c 'CGAAGTCAGGATTCCGG' *.fastq > count/B1410_Fm0.txt
agrep -0 -c 'CAAGGTCCCTCATTAAT' *.fastq > count/B1412_Fm0.txt
agrep -0 -c 'TACGCGCAATACCTCAT' *.fastq > count/B1413_Fm0.txt
agrep -0 -c 'AGTAGCCTAAGAGATGC' *.fastq > count/B1414_Fm0.txt
agrep -0 -c 'TGCAGCGCATACCTTTC' *.fastq > count/B1415_Fm0.txt
agrep -0 -c 'AGCTCGACGGGTACTAT' *.fastq > count/B1416_Fm0.txt
agrep -0 -c 'ATTAACGCCGGTAATTG' *.fastq > count/B1417_Fm0.txt
agrep -0 -c 'GTAGGGTCGCTGCTATC' *.fastq > count/B1418_Fm0.txt
agrep -0 -c 'TCATCCAGGGAAGCGTT' *.fastq > count/B1419_Fm0.txt
agrep -0 -c 'GGAGTACCGGATCATGG' *.fastq > count/B1420_Fm0.txt
agrep -0 -c 'ACGGTGGAAATTTATGG' *.fastq > count/B1422_Fm0.txt
agrep -0 -c 'TCGCATTAGTGTCCCTT' *.fastq > count/B1423_Fm0.txt
agrep -0 -c 'CAAGTTCTGCGAATACG' *.fastq > count/B1424_Fm0.txt
agrep -0 -c 'GGTAGCCGCAACTACGA' *.fastq > count/B1425_Fm0.txt
agrep -0 -c 'AGTTCTAATTCCCCCGG' *.fastq > count/B1427_Fm0.txt
agrep -0 -c 'GACGTAGGTAGCCTGTT' *.fastq > count/B1429_Fm0.txt
agrep -0 -c 'AGCAAGGGTATGTAATC' *.fastq > count/B1430_Fm0.txt
agrep -0 -c 'GGCTAAAAGAGCTGGGC' *.fastq > count/B1431_Fm0.txt
agrep -0 -c 'AGTCGAATGTACTGACC' *.fastq > count/B1433_Fm0.txt
agrep -0 -c 'GATTGAGAAAGTAGCCG' *.fastq > count/B1435_Fm0.txt
agrep -0 -c 'TTGCCAACTATCCCATC' *.fastq > count/B1436_Fm0.txt
agrep -0 -c 'TCGCCAGCAATAGACCG' *.fastq > count/B1437_Fm0.txt
agrep -0 -c 'GTCCTGTTTCAGTGTGC' *.fastq > count/B1439_Fm0.txt
agrep -0 -c 'GTGTCGTCAGACGGTTG' *.fastq > count/B1440_Fm0.txt
agrep -0 -c 'TTGTCAACTGGTTCCTG' *.fastq > count/B1441_Fm0.txt
agrep -0 -c 'GTATGTCGTTCCTAAGC' *.fastq > count/B1442_Fm0.txt
agrep -0 -c 'GGGTTGGGATTGCTAGC' *.fastq > count/B1443_Fm0.txt
agrep -0 -c 'GCAATCTTCCGTGCAGC' *.fastq > count/B1444_Fm0.txt
agrep -0 -c 'CGAATAGGGATTTCGCT' *.fastq > count/B1445_Fm0.txt
agrep -0 -c 'CTAAGCATCGGCCCTCA' *.fastq > count/B1446_Fm0.txt
agrep -0 -c 'CATCGATATTGGTGGCG' *.fastq > count/B1447_Fm0.txt
agrep -0 -c 'CGATTCATAGCGCCGTG' *.fastq > count/B1448_Fm0.txt
agrep -0 -c 'AGGAGTATGGTTATGTG' *.fastq > count/B1449_Fm0.txt
agrep -0 -c 'CTTACGGCGTTACATGA' *.fastq > count/B1450_Fm0.txt
agrep -0 -c 'GTCTCTATTCCCGCCCA' *.fastq > count/B1451_Fm0.txt
agrep -0 -c 'ATTACGCTTGGTGGGAC' *.fastq > count/B1452_Fm0.txt
agrep -0 -c 'TTCCGACTTCGGCACAC' *.fastq > count/B1453_Fm0.txt
agrep -0 -c 'ACTAATGGGTAGTTCGT' *.fastq > count/B1454_Fm0.txt
agrep -0 -c 'CTAGAACTTTCGGGGCC' *.fastq > count/B1455_Fm0.txt
agrep -0 -c 'GGACTGCCCTCCAATTG' *.fastq > count/B1457_Fm0.txt
agrep -0 -c 'TCGGATTTGATCTTGTG' *.fastq > count/B1458_Fm0.txt
agrep -0 -c 'TACTGTCCATTAACGCA' *.fastq > count/B1459_Fm0.txt
agrep -0 -c 'ATCCAATGCTGGTGCTA' *.fastq > count/B1461_Fm0.txt
agrep -0 -c 'CTACTCGAGCCCTGTAC' *.fastq > count/B1463_Fm0.txt
agrep -0 -c 'AGTACTGTCTCGCTCTT' *.fastq > count/B1464_Fm0.txt
agrep -0 -c 'GCCGTATATCTTCGGTC' *.fastq > count/B1465_Fm0.txt
agrep -0 -c 'GTGAAATCCCTGTTTGC' *.fastq > count/B1468_Fm0.txt
agrep -0 -c 'GCCTGTATCGGCACATA' *.fastq > count/B1470_Fm0.txt
agrep -0 -c 'GCCGTAACGAGAAGCGA' *.fastq > count/B1471_Fm0.txt
agrep -0 -c 'ACATGGGTAAAGCCAAT' *.fastq > count/B1472_Fm0.txt
agrep -0 -c 'GTACTGCATCATACTCT' *.fastq > count/B1473_Fm0.txt
agrep -0 -c 'GGAACCTAGACTACCCG' *.fastq > count/B1474_Fm0.txt
agrep -0 -c 'TCTCCCTTCGACAGACA' *.fastq > count/B1475_Fm0.txt
agrep -0 -c 'CGAATCTTGGACGTTTG' *.fastq > count/B1476_Fm0.txt
agrep -0 -c 'AGTCGGAGAGATCGACC' *.fastq > count/B1477_Fm0.txt
agrep -0 -c 'ATAGCCTCGTGGAAGCA' *.fastq > count/B1479_Fm0.txt
agrep -0 -c 'CTGTGTGATTAGCCGGG' *.fastq > count/B1480_Fm0.txt
agrep -0 -c 'CAAACTCTTTGAATCGC' *.fastq > count/B1481_Fm0.txt
agrep -0 -c 'AGAAGTTACCTCCGTAA' *.fastq > count/B1483_Fm0.txt
agrep -0 -c 'CAGGCGTTACGATCTAG' *.fastq > count/B1484_Fm0.txt
agrep -0 -c 'GGGATCATACCTTGAAC' *.fastq > count/B1485_Fm0.txt
agrep -0 -c 'CATCTAGCTTGGAAGGG' *.fastq > count/B1486_Fm0.txt
agrep -0 -c 'TGCGAATGAGAGCGCTG' *.fastq > count/B1487_Fm0.txt
agrep -0 -c 'AAGGTGACCAGATCAAG' *.fastq > count/B1488_Fm0.txt
agrep -0 -c 'CTTGGTAATTGTGGAGC' *.fastq > count/B1489_Fm0.txt
agrep -0 -c 'TGTTTGGACGTCGTTAC' *.fastq > count/B1491_Fm0.txt
agrep -0 -c 'CAGAACCGCTATGGCTT' *.fastq > count/B1492_Fm0.txt
agrep -0 -c 'CCTCCGGGTTGTTCAAG' *.fastq > count/B1494_Fm0.txt
agrep -0 -c 'CGCGGTTACTCCAAATC' *.fastq > count/B1495_Fm0.txt
agrep -0 -c 'GGGATGTACGGATTGTA' *.fastq > count/B1496_Fm0.txt
agrep -0 -c 'GCATAGTACTACTACGT' *.fastq > count/B1497_Fm0.txt
agrep -0 -c 'TATTTGTGGCTGGCGTA' *.fastq > count/B1498_Fm0.txt
agrep -0 -c 'CGACGCGGTACATTCTG' *.fastq > count/B1500_Fm0.txt
agrep -0 -c 'TCTGCACGTAGCAAGCC' *.fastq > count/B1501_Fm0.txt
agrep -0 -c 'CGTTCTTCGACGCTCCT' *.fastq > count/B1502_Fm0.txt
agrep -0 -c 'ACTGGCTATGTGGGATG' *.fastq > count/B1503_Fm0.txt
agrep -0 -c 'TCGGTAACTACGCCACT' *.fastq > count/B1504_Fm0.txt
agrep -0 -c 'ATTGCAGTAGAGTCCGA' *.fastq > count/B1505_Fm0.txt
agrep -0 -c 'ACTTCACGTGTTGATGC' *.fastq > count/B1506_Fm0.txt
agrep -0 -c 'TTGATCCGGGTTTGTTT' *.fastq > count/B1507_Fm0.txt
agrep -0 -c 'ACGAGCCTTATGGAAGT' *.fastq > count/B1509_Fm0.txt
agrep -0 -c 'TGTAATGAAGGCACAAC' *.fastq > count/B1511_Fm0.txt
agrep -0 -c 'ATAACCCCGTTTGGCCT' *.fastq > count/B1512_Fm0.txt
agrep -0 -c 'TCATATGCCGATGCCGC' *.fastq > count/B1513_Fm0.txt
agrep -0 -c 'TTGCTTCGAGCAATTCT' *.fastq > count/B1514_Fm0.txt
agrep -0 -c 'CCTGATGGTGGTTAGTC' *.fastq > count/B1515_Fm0.txt
agrep -0 -c 'CGGGGTATTACCTTCTA' *.fastq > count/B1516_Fm0.txt
agrep -0 -c 'CTCGAGCTAAACACATT' *.fastq > count/B1517_Fm0.txt
agrep -0 -c 'TGCTTACACGGTTCGTT' *.fastq > count/B1518_Fm0.txt
agrep -0 -c 'ACGGATCGGAGCGTTTC' *.fastq > count/B1519_Fm0.txt
agrep -0 -c 'TGCGCCGATTACATACT' *.fastq > count/B1521_Fm0.txt
agrep -0 -c 'CCTCAAAACGGTAGCTG' *.fastq > count/B1522_Fm0.txt
agrep -0 -c 'GATTGCGACTACCATGG' *.fastq > count/B1523_Fm0.txt
agrep -0 -c 'CCGAAGGCCACCAATAA' *.fastq > count/B1524_Fm0.txt
agrep -0 -c 'TAAGTCTACGCTCCCCA' *.fastq > count/B1525_Fm0.txt
agrep -0 -c 'GTTCTGAAGCAAGAACC' *.fastq > count/B1526_Fm0.txt
agrep -0 -c 'TTCAAAATCCGGCGATC' *.fastq > count/B1527_Fm0.txt
agrep -0 -c 'ATATAAACCTCGACGGC' *.fastq > count/B1528_Fm0.txt
agrep -0 -c 'TCACTACTAACCCCCAT' *.fastq > count/B1529_Fm0.txt
agrep -0 -c 'ATATCCAACTTTCGCCT' *.fastq > count/B1530_Fm0.txt
agrep -0 -c 'TTCCGTAAGCTTTCAAC' *.fastq > count/B1531_Fm0.txt
agrep -0 -c 'TTGACTCTGTATGCAGT' *.fastq > count/B1532_Fm0.txt
agrep -0 -c 'TGAGAGATGACCGGCGT' *.fastq > count/B1535_Fm0.txt
agrep -0 -c 'AATGAACCGGCCACACT' *.fastq > count/B1536_Fm0.txt
agrep -0 -c 'AGATAACCGACTGTTTC' *.fastq > count/B1538_Fm0.txt
agrep -0 -c 'GTGGAGTTAGCGTCGCA' *.fastq > count/B1540_Fm0.txt
agrep -0 -c 'TAGCAAAAGTGTAGGAG' *.fastq > count/B1541_Fm0.txt
agrep -0 -c 'TGTTTACCTGGACAGGC' *.fastq > count/B1542_Fm0.txt
agrep -0 -c 'TGTCTCCCGGAACCTGT' *.fastq > count/B1544_Fm0.txt
agrep -0 -c 'CGTGCACGGGCTCAAAA' *.fastq > count/B1545_Fm0.txt
agrep -0 -c 'GGCCAATAGCGCGTTCA' *.fastq > count/B1546_Fm0.txt
agrep -0 -c 'GTTTCCTGACAGTACCC' *.fastq > count/B1547_Fm0.txt
agrep -0 -c 'ATTTCTTTCTACGCGCA' *.fastq > count/B1548_Fm0.txt
agrep -0 -c 'GGCCTGAACCCGAATTA' *.fastq > count/B1549_Fm0.txt
agrep -0 -c 'ACACTGAATCAGCTTCT' *.fastq > count/B1550_Fm0.txt
agrep -0 -c 'GGATTCCGGCCAGCAAA' *.fastq > count/B1551_Fm0.txt
agrep -0 -c 'CTAATTGCTCAACCCCT' *.fastq > count/B1552_Fm0.txt
agrep -0 -c 'GCCGATTTAGAGCGCCA' *.fastq > count/B1553_Fm0.txt
agrep -0 -c 'ACACTTGGAGGGTCTTG' *.fastq > count/B1554_Fm0.txt
agrep -0 -c 'GACCACCGAATATCTTT' *.fastq > count/B1555_Fm0.txt
agrep -0 -c 'CAGTAGAGATCGCATGC' *.fastq > count/B1556_Fm0.txt
agrep -0 -c 'AACGGAGAGGTATGTAT' *.fastq > count/B1557_Fm0.txt
agrep -0 -c 'GATATTGGATCGGGATC' *.fastq > count/B1558_Fm0.txt
agrep -0 -c 'CCATTTCACAGCAGGAG' *.fastq > count/B1559_Fm0.txt
agrep -0 -c 'TTCCGACATGGCATCAC' *.fastq > count/B1561_Fm0.txt
agrep -0 -c 'GGACAATTCCCCATATG' *.fastq > count/B1562_Fm0.txt
agrep -0 -c 'TGACGGAATACCCATGC' *.fastq > count/B1563_Fm0.txt
agrep -0 -c 'TCAAACCGTGCACGAAG' *.fastq > count/B1564_Fm0.txt
agrep -0 -c 'CTACTTCGGACACGGTT' *.fastq > count/B1565_Fm0.txt
agrep -0 -c 'TCACTCTCTAGCTCGAT' *.fastq > count/B1566_Fm0.txt
agrep -0 -c 'AAGCATCTGGCCCCCTT' *.fastq > count/B1567_Fm0.txt
agrep -0 -c 'GGTAAATGCGACCGGGT' *.fastq > count/B1568_Fm0.txt
agrep -0 -c 'GCCAGTGCTAGTAGTGC' *.fastq > count/B1569_Fm0.txt
agrep -0 -c 'GTAGTGAAAACGCGCAA' *.fastq > count/B1570_Fm0.txt
agrep -0 -c 'GCTCGGCCAAAATCCCT' *.fastq > count/B1571_Fm0.txt
agrep -0 -c 'GTTTCGCTCTTACCATT' *.fastq > count/B1572_Fm0.txt
agrep -0 -c 'CGCAGTAAAGGGGTGTT' *.fastq > count/B1573_Fm0.txt
agrep -0 -c 'CGTATGAGGTGGGGTAC' *.fastq > count/B1574_Fm0.txt
agrep -0 -c 'TGCATGTCATGCTTGCC' *.fastq > count/B1575_Fm0.txt
agrep -0 -c 'TTCACGATTCGATTCCC' *.fastq > count/B1576_Fm0.txt
agrep -0 -c 'ATCAACCCAAGCTGTTA' *.fastq > count/B1577_Fm0.txt
agrep -0 -c 'ATGCACGTGAACTGTGT' *.fastq > count/B1580_Fm0.txt
agrep -0 -c 'CGGCACTAGATATCACG' *.fastq > count/B1582_Fm0.txt
agrep -0 -c 'CAAAATGCACTGCGAAC' *.fastq > count/B1584_Fm0.txt
agrep -0 -c 'CCACGGGATAAGATTCT' *.fastq > count/B1585_Fm0.txt
agrep -0 -c 'GATGTCAGGATGAATGT' *.fastq > count/B1586_Fm0.txt
agrep -0 -c 'TGCATCTTACCGAACCC' *.fastq > count/B1587_Fm0.txt
agrep -0 -c 'AACTTCTGGCCAGTTCC' *.fastq > count/B1588_Fm0.txt
agrep -0 -c 'GCTACGAACAGGGCAGT' *.fastq > count/B1589_Fm0.txt
agrep -0 -c 'AGTTGGTAGGTGTCCAG' *.fastq > count/B1590_Fm0.txt
agrep -0 -c 'ACAATAAGGGTGGCGTT' *.fastq > count/B1592_Fm0.txt
agrep -0 -c 'GCAATGTCGTGTGGAAA' *.fastq > count/B1593_Fm0.txt
agrep -0 -c 'TTCCGCTGTACTGAATG' *.fastq > count/B1594_Fm0.txt
agrep -0 -c 'ATAAACCGCTACCGCCC' *.fastq > count/B1595_Fm0.txt
agrep -0 -c 'GACTGTCCTCCCGAGTA' *.fastq > count/B1596_Fm0.txt
agrep -0 -c 'CACAACGTAAACACTAG' *.fastq > count/B1597_Fm0.txt
agrep -0 -c 'AGCATCGACGTAGCAGC' *.fastq > count/B1598_Fm0.txt
agrep -0 -c 'GGATCGAGTACGATACC' *.fastq > count/B1599_Fm0.txt
agrep -0 -c 'GCGTTAAGTCCGCGTAA' *.fastq > count/B1600_Fm0.txt
agrep -0 -c 'ACAACAGTATGGGATCA' *.fastq > count/B1602_Fm0.txt
agrep -0 -c 'ACGAATTCGCCGCGAAG' *.fastq > count/B1603_Fm0.txt
agrep -0 -c 'ATGAACTGCATCTTGCA' *.fastq > count/B1605_Fm0.txt
agrep -0 -c 'ATCCAGTGGCCGTGTGA' *.fastq > count/B1606_Fm0.txt
agrep -0 -c 'TAGTACCCTCATAACGC' *.fastq > count/B1607_Fm0.txt
agrep -0 -c 'ACTCAGTGCGCTAAGCG' *.fastq > count/B1608_Fm0.txt
agrep -0 -c 'GGTATTCCCTAACTATC' *.fastq > count/B1609_Fm0.txt
agrep -0 -c 'GGATGTTCACGCCAGCT' *.fastq > count/B1610_Fm0.txt
agrep -0 -c 'AGGTGGTCAAGAAGGGT' *.fastq > count/B1611_Fm0.txt
agrep -0 -c 'CTCAGGCGCAACGATCT' *.fastq > count/B1612_Fm0.txt
agrep -0 -c 'GGCCGTGGTGAACAAGA' *.fastq > count/B1613_Fm0.txt
agrep -0 -c 'TGCCATCATGAATTCTC' *.fastq > count/B1615_Fm0.txt
agrep -0 -c 'TCTATCCTCCTGCTTAC' *.fastq > count/B1616_Fm0.txt
agrep -0 -c 'GGATTGACGTTATGCCG' *.fastq > count/B1617_Fm0.txt
agrep -0 -c 'GTTTATCACGGGATCTC' *.fastq > count/B1618_Fm0.txt
agrep -0 -c 'GCCCGCGTAATTTCATA' *.fastq > count/B1619_Fm0.txt
agrep -0 -c 'AGTCGTCAAGACAGGCA' *.fastq > count/B1620_Fm0.txt
agrep -0 -c 'TTGTTATCGGATGAGGG' *.fastq > count/B1621_Fm0.txt
agrep -0 -c 'TTGCAGAGAGTATCGAA' *.fastq > count/B1622_Fm0.txt
agrep -0 -c 'GCTACTTGAGCTCTGTC' *.fastq > count/B1625_Fm0.txt
agrep -0 -c 'CCTTGTATCGACCATTG' *.fastq > count/B1626_Fm0.txt
agrep -0 -c 'TCTACACATGACCTGGC' *.fastq > count/B1627_Fm0.txt
agrep -0 -c 'AGGGCCACATTTAGGCG' *.fastq > count/B1628_Fm0.txt
agrep -0 -c 'TCACCCCGTATGAAAGG' *.fastq > count/B1629_Fm0.txt
agrep -0 -c 'AAGTGGGTGTTCTCTTC' *.fastq > count/B1630_Fm0.txt
agrep -0 -c 'AGACTGGCCCTTTCGAG' *.fastq > count/B1631_Fm0.txt
agrep -0 -c 'GTCCAACGATGAGGGGA' *.fastq > count/B1632_Fm0.txt
agrep -0 -c 'CGTCATAAGGCGAAAAT' *.fastq > count/B1633_Fm0.txt
agrep -0 -c 'CGTGTCTGAGAGAGACT' *.fastq > count/B1634_Fm0.txt
agrep -0 -c 'ATTGTCTGGGGTCTTGC' *.fastq > count/B1635_Fm0.txt
agrep -0 -c 'CCTTCGCCAGATTCCAC' *.fastq > count/B1636_Fm0.txt
agrep -0 -c 'CCGTATACCACAAAAAC' *.fastq > count/B1637_Fm0.txt
agrep -0 -c 'ACCTGTATCCGTCACGA' *.fastq > count/B1638_Fm0.txt
agrep -0 -c 'ATCTGCGTGGATAAAAG' *.fastq > count/B1639_Fm0.txt
agrep -0 -c 'AAGGCACTTTTGCGAAG' *.fastq > count/B1640_Fm0.txt
agrep -0 -c 'TACTAACGCCCATTCGA' *.fastq > count/B1641_Fm0.txt
agrep -0 -c 'CGAGGATCCGGATTCAA' *.fastq > count/B1642_Fm0.txt
agrep -0 -c 'ATTTTGGCGGCCTTCGG' *.fastq > count/B1643_Fm0.txt
agrep -0 -c 'TGCGCATTTCTGGGCCT' *.fastq > count/B1644_Fm0.txt
agrep -0 -c 'CTCTAAGACCTGGTTAC' *.fastq > count/B1645_Fm0.txt
agrep -0 -c 'CCGCCAAGTTTCACTGG' *.fastq > count/B1646_Fm0.txt
agrep -0 -c 'TTCCGGATAATGTCCCT' *.fastq > count/B1648_Fm0.txt
agrep -0 -c 'CCAATCCCTGATGGCGA' *.fastq > count/B1649_Fm0.txt
agrep -0 -c 'CATGCTAGGGCAACCGA' *.fastq > count/B1650_Fm0.txt
agrep -0 -c 'TATTCCACGTCACAGAA' *.fastq > count/B1651_Fm0.txt
agrep -0 -c 'GCTGGATCTTTCGAGAG' *.fastq > count/B1652_Fm0.txt
agrep -0 -c 'ATCTATGTATGGTCGAC' *.fastq > count/B1653_Fm0.txt
agrep -0 -c 'CGATCAGGTACTTTGCA' *.fastq > count/B1655_Fm0.txt
agrep -0 -c 'CCACTCGTATTGCTAAT' *.fastq > count/B1656_Fm0.txt
agrep -0 -c 'TAATCACCAGTAGCCGC' *.fastq > count/B1657_Fm0.txt
agrep -0 -c 'AGGTCCTCACAGTGAGG' *.fastq > count/B1658_Fm0.txt
agrep -0 -c 'AACACAAACGCGTCCGA' *.fastq > count/B1659_Fm0.txt
agrep -0 -c 'AGGTATGCATCGCGGGA' *.fastq > count/B1660_Fm0.txt
agrep -0 -c 'CACGAGATATGGCGCCA' *.fastq > count/B1662_Fm0.txt
agrep -0 -c 'GTTGCAAGCCAGGAGTT' *.fastq > count/B1663_Fm0.txt
agrep -0 -c 'AATTGAAGCAGTAGTCG' *.fastq > count/B1664_Fm0.txt
agrep -0 -c 'GGATTGCCTAAAATCGA' *.fastq > count/B1665_Fm0.txt
agrep -0 -c 'TTGGGGTCTCAATGCAC' *.fastq > count/B1666_Fm0.txt
agrep -0 -c 'GGCTCCCAACGTTTGTT' *.fastq > count/B1667_Fm0.txt
agrep -0 -c 'GTAACCTGACATGTAGT' *.fastq > count/B1668_Fm0.txt
agrep -0 -c 'TGATCCCGGAAGGTTCT' *.fastq > count/B1669_Fm0.txt
agrep -0 -c 'GCAAAAGCCTAAAACCA' *.fastq > count/B1670_Fm0.txt
agrep -0 -c 'TACCAACCGCCTGTGGA' *.fastq > count/B1671_Fm0.txt
agrep -0 -c 'TGCGGAATCAGTCTCTC' *.fastq > count/B1672_Fm0.txt
agrep -0 -c 'GACCTCCTAGAAGTGCG' *.fastq > count/B1673_Fm0.txt
agrep -0 -c 'GCTACTGTAGTATATCC' *.fastq > count/B1674_Fm0.txt
agrep -0 -c 'AACGCGGTCTCCAGTGT' *.fastq > count/B1675_Fm0.txt
agrep -0 -c 'GTAATTGTTACGCCGTT' *.fastq > count/B1676_Fm0.txt
agrep -0 -c 'TAGGCGTGCTGTTAGGT' *.fastq > count/B1677_Fm0.txt
agrep -0 -c 'GGATTAGATGCCTGTAC' *.fastq > count/B1678_Fm0.txt
agrep -0 -c 'CTTCGGACAGAAGAAGC' *.fastq > count/B1679_Fm0.txt
agrep -0 -c 'ACGTCCTTGCGAGTTCA' *.fastq > count/B1680_Fm0.txt
agrep -0 -c 'TAGTGACTTGTGCGGCG' *.fastq > count/B1681_Fm0.txt
agrep -0 -c 'CTGTTCTATAGATAGCG' *.fastq > count/B1682_Fm0.txt
agrep -0 -c 'AACTAACGACCAGCTAT' *.fastq > count/B1683_Fm0.txt
agrep -0 -c 'AGCGGCAGATATGCGCT' *.fastq > count/B1684_Fm0.txt
agrep -0 -c 'TCACGTGGGTAGTCTAA' *.fastq > count/B1685_Fm0.txt
agrep -0 -c 'ATGACCTGGTATAACCC' *.fastq > count/B1686_Fm0.txt
agrep -0 -c 'ACTAACTATCGGCTAGG' *.fastq > count/B1687_Fm0.txt
agrep -0 -c 'GCGAGAACTTTTGTTGG' *.fastq > count/B1688_Fm0.txt
agrep -0 -c 'TAGACCGCACGGGATAT' *.fastq > count/B1689_Fm0.txt
agrep -0 -c 'GTGGGCTCACTGTCAGT' *.fastq > count/B1690_Fm0.txt
agrep -0 -c 'AATACGATGAGGTCCGG' *.fastq > count/B1691_Fm0.txt
agrep -0 -c 'CCATCTTCTAGCGGGCT' *.fastq > count/B1692_Fm0.txt
agrep -0 -c 'AATTGTCCCTGGCGTGT' *.fastq > count/B1694_Fm0.txt
agrep -0 -c 'ATGGACCTATTTGCGTC' *.fastq > count/B1695_Fm0.txt
agrep -0 -c 'GTGTGGGGTGAAGATCA' *.fastq > count/B1697_Fm0.txt
agrep -0 -c 'CATACTATGCCTCGGCC' *.fastq > count/B1698_Fm0.txt
agrep -0 -c 'CGCTCATATGCCAAAAG' *.fastq > count/B1699_Fm0.txt
agrep -0 -c 'GCATTGTGACCCATTAA' *.fastq > count/B1700_Fm0.txt
agrep -0 -c 'CTTGAACAATCGGGTAC' *.fastq > count/B1701_Fm0.txt
agrep -0 -c 'GCGCATATTCAAAGCGT' *.fastq > count/B1702_Fm0.txt
agrep -0 -c 'TCGACATCTTTCCGAAA' *.fastq > count/B1703_Fm0.txt
agrep -0 -c 'TATCCTATGCTCCGCAA' *.fastq > count/B1704_Fm0.txt
agrep -0 -c 'TGTGCAATGCGCAGTAC' *.fastq > count/B1705_Fm0.txt
agrep -0 -c 'TATCGACACGCTATCAT' *.fastq > count/B1707_Fm0.txt
agrep -0 -c 'CTAAAGTGCAGCGAGAC' *.fastq > count/B1708_Fm0.txt
agrep -0 -c 'GATGCCCTACTCTCAGT' *.fastq > count/B1710_Fm0.txt
agrep -0 -c 'ACATGTTAACAACCCGC' *.fastq > count/B1711_Fm0.txt
agrep -0 -c 'CGCTTAACTCCTTTACA' *.fastq > count/B1712_Fm0.txt
agrep -0 -c 'AGACAACAGAGCACGGT' *.fastq > count/B1714_Fm0.txt
agrep -0 -c 'CAGCAAGTTCTAAGGTC' *.fastq > count/B1715_Fm0.txt
agrep -0 -c 'GGCCTGGGTGATTCTTC' *.fastq > count/B1718_Fm0.txt
agrep -0 -c 'CAGGGAAATCCAATCAC' *.fastq > count/B1719_Fm0.txt
agrep -0 -c 'AGAATCGGTAGGGTGGC' *.fastq > count/B1720_Fm0.txt
agrep -0 -c 'CTGGATGTAAATTGCAC' *.fastq > count/B1721_Fm0.txt
agrep -0 -c 'TATCGCGAGCGGTCCTT' *.fastq > count/B1722_Fm0.txt
agrep -0 -c 'ACCCAAGGGCCATTACG' *.fastq > count/B1723_Fm0.txt
agrep -0 -c 'GTTAAACATCGGCGGTG' *.fastq > count/B1724_Fm0.txt
agrep -0 -c 'AGAGAACGCCGGTATAC' *.fastq > count/B1725_Fm0.txt
agrep -0 -c 'CCGCACGTACAAATGTC' *.fastq > count/B1726_Fm0.txt
agrep -0 -c 'TAAACTGAGTCCGTCAC' *.fastq > count/B1728_Fm0.txt
agrep -0 -c 'TAACGACTCTGGTCTTA' *.fastq > count/B1729_Fm0.txt
agrep -0 -c 'GTAATTCGATGGGTCTT' *.fastq > count/B1730_Fm0.txt
agrep -0 -c 'TCGCTACTCCCGTTTTA' *.fastq > count/B1731_Fm0.txt
agrep -0 -c 'ATAAACCTCGGCTTCGG' *.fastq > count/B1732_Fm0.txt
agrep -0 -c 'GACTGCATCGCGCATGA' *.fastq > count/B1733_Fm0.txt
agrep -0 -c 'GTAGTGCGATCTCTAAA' *.fastq > count/B1734_Fm0.txt
agrep -0 -c 'TACGATCTCATCGGGTC' *.fastq > count/B1735_Fm0.txt
agrep -0 -c 'CAGCCGAGGTCGAAACA' *.fastq > count/B1736_Fm0.txt
agrep -0 -c 'GCCATGTCTCTATCGCG' *.fastq > count/B1737_Fm0.txt
agrep -0 -c 'ACCCGAACCTGGTACTC' *.fastq > count/B1738_Fm0.txt
agrep -0 -c 'GCACTCCAATCTCACCC' *.fastq > count/B1739_Fm0.txt
agrep -0 -c 'CTCAGGGAAATCTTCTG' *.fastq > count/B1740_Fm0.txt
agrep -0 -c 'GGAATAGTGAACCGAAA' *.fastq > count/B1741_Fm0.txt
agrep -0 -c 'ACCCGGCAAGTTTAGAG' *.fastq > count/B1743_Fm0.txt
agrep -0 -c 'AGAGCGGTCAACTGTTT' *.fastq > count/B1744_Fm0.txt
agrep -0 -c 'CCGAATTTATATTCGCG' *.fastq > count/B1747_Fm0.txt
agrep -0 -c 'AGTCCAGAACTATACGC' *.fastq > count/B1748_Fm0.txt
agrep -0 -c 'CATATGGGGAGGAGTAC' *.fastq > count/B1749_Fm0.txt
agrep -0 -c 'GTTACGAGGCCTTAGCT' *.fastq > count/B1750_Fm0.txt
agrep -0 -c 'TTCCTACGTTACATTCC' *.fastq > count/B1751_Fm0.txt
agrep -0 -c 'GACCGAATAACTAGGGC' *.fastq > count/B1752_Fm0.txt
agrep -0 -c 'GCTGTACAATGAGCTAT' *.fastq > count/B1754_Fm0.txt
agrep -0 -c 'TAAGGTCAGAGTACGAG' *.fastq > count/B1756_Fm0.txt
agrep -0 -c 'CTACAGACCAGCGAGCT' *.fastq > count/B1757_Fm0.txt
agrep -0 -c 'CTTGATCACGCATTCTG' *.fastq > count/B1758_Fm0.txt
agrep -0 -c 'GCTCCGTTCGTTTTCCT' *.fastq > count/B1759_Fm0.txt
agrep -0 -c 'CGCACCCGACGCAATTA' *.fastq > count/B1760_Fm0.txt
agrep -0 -c 'CTATCCACTTTGTGAAG' *.fastq > count/B1762_Fm0.txt
agrep -0 -c 'TGCAGATAGCAATAGCG' *.fastq > count/B1763_Fm0.txt
agrep -0 -c 'CCTAGGCTGTATCTGCC' *.fastq > count/B1766_Fm0.txt
agrep -0 -c 'TCAGGCTCAACTCTTAG' *.fastq > count/B1767_Fm0.txt
agrep -0 -c 'CGATGCCCCGAAAGTTC' *.fastq > count/B1768_Fm0.txt
agrep -0 -c 'ACGGTGCTGTAAACGTG' *.fastq > count/B1769_Fm0.txt
agrep -0 -c 'TATCCATACGTGAAGGC' *.fastq > count/B1770_Fm0.txt
agrep -0 -c 'AGAAACGGTCGTCTCTT' *.fastq > count/B1771_Fm0.txt
agrep -0 -c 'GGTGGCAACGTGTTTAG' *.fastq > count/B1772_Fm0.txt
agrep -0 -c 'TCTTGTGTCCTTTGGGA' *.fastq > count/B1773_Fm0.txt
agrep -0 -c 'ACTTACTCAGGGGTTGT' *.fastq > count/B1774_Fm0.txt
agrep -0 -c 'TACTGCGTCGTCTGTCG' *.fastq > count/B1775_Fm0.txt
agrep -0 -c 'CAGACAAGTGAGCGAGG' *.fastq > count/B1777_Fm0.txt
agrep -0 -c 'TACACCGGCTCAACGGT' *.fastq > count/B1778_Fm0.txt
agrep -0 -c 'AATGCCTGTTCCGTATT' *.fastq > count/B1779_Fm0.txt
agrep -0 -c 'TTAGCAGCGGATCTGGT' *.fastq > count/B1780_Fm0.txt
agrep -0 -c 'CTGTATAGGTTAATCGC' *.fastq > count/B1781_Fm0.txt
agrep -0 -c 'GGGGTAGTGAGTAATGG' *.fastq > count/B1782_Fm0.txt
agrep -0 -c 'CCTATAAGTCTGGCTTA' *.fastq > count/B1783_Fm0.txt
agrep -0 -c 'TGCACTAGGAGTTTCCG' *.fastq > count/B1784_Fm0.txt
agrep -0 -c 'GTTGGGAAACAGAGTAA' *.fastq > count/B1785_Fm0.txt
agrep -0 -c 'CACGTTCGCCTACGTGA' *.fastq > count/B1786_Fm0.txt
agrep -0 -c 'ACAGAGGCGATGGTATT' *.fastq > count/B1787_Fm0.txt
agrep -0 -c 'ATCCGCCATTCGAAGGC' *.fastq > count/B1789_Fm0.txt
agrep -0 -c 'TCAACACCGGCGACTAT' *.fastq > count/B1790_Fm0.txt
agrep -0 -c 'CCTGCTCTATTGTGATG' *.fastq > count/B1791_Fm0.txt
agrep -0 -c 'ATCGCCGTTAAGATGTT' *.fastq > count/B1792_Fm0.txt
agrep -0 -c 'GACCCATCGTTGACGTT' *.fastq > count/B1794_Fm0.txt
agrep -0 -c 'CGGTGTTACATTCGAAC' *.fastq > count/B1795_Fm0.txt
agrep -0 -c 'CCAAAAGTTACCCGCCC' *.fastq > count/B1798_Fm0.txt
agrep -0 -c 'TGCGAGGAGCACGAATA' *.fastq > count/B1799_Fm0.txt
agrep -0 -c 'TAACACTGAGCTCGCTA' *.fastq > count/B1801_Fm0.txt
agrep -0 -c 'GTCTACGTAAGTATGTC' *.fastq > count/B1802_Fm0.txt
agrep -0 -c 'CCTGTTTGGATAAGCGG' *.fastq > count/B1803_Fm0.txt
agrep -0 -c 'TTCGTTACCCGGCGTTA' *.fastq > count/B1804_Fm0.txt
agrep -0 -c 'TGGCCTAAGTGGCTTTC' *.fastq > count/B1806_Fm0.txt
agrep -0 -c 'ATGGAACGTCCTCCGCT' *.fastq > count/B1807_Fm0.txt
agrep -0 -c 'CTCAGCGAATCTGTGAA' *.fastq > count/B1808_Fm0.txt
agrep -0 -c 'ATGCCCGTAACGTATGT' *.fastq > count/B1809_Fm0.txt
agrep -0 -c 'ACAGTAGACGGAATACG' *.fastq > count/B1810_Fm0.txt
agrep -0 -c 'CTGTAGTCAGGGTCTTC' *.fastq > count/B1811_Fm0.txt
agrep -0 -c 'GCCCAGTCCCCTAATAA' *.fastq > count/B1812_Fm0.txt
agrep -0 -c 'AGGGCTGATTTGTTCCC' *.fastq > count/B1813_Fm0.txt
agrep -0 -c 'TTATTAGGCGAGTAACC' *.fastq > count/B1814_Fm0.txt
agrep -0 -c 'GGACCGAGAGTACGAAC' *.fastq > count/B1815_Fm0.txt
agrep -0 -c 'ACAGTAGGTGACGAGTG' *.fastq > count/B1816_Fm0.txt
agrep -0 -c 'GTAGAGTCGCTTCTAAT' *.fastq > count/B1817_Fm0.txt
agrep -0 -c 'TGCCGAGATTACGAGTT' *.fastq > count/B1818_Fm0.txt
agrep -0 -c 'CTAACGTGTCTCGGTAT' *.fastq > count/B1819_Fm0.txt
agrep -0 -c 'GCTACCAGCACTCGATA' *.fastq > count/B1820_Fm0.txt
agrep -0 -c 'ATCGTCGTGGCTAAGCC' *.fastq > count/B1821_Fm0.txt
agrep -0 -c 'CGTGAGCAGCGTCCTAT' *.fastq > count/B1822_Fm0.txt
agrep -0 -c 'AACTCCCGCTACAAAAC' *.fastq > count/B1823_Fm0.txt
agrep -0 -c 'CTGATCTTATACGCCGT' *.fastq > count/B1824_Fm0.txt
agrep -0 -c 'AGGTGTTCGTAAGAACA' *.fastq > count/B1825_Fm0.txt
agrep -0 -c 'GGCAACGAAGATTACTT' *.fastq > count/B1826_Fm0.txt
agrep -0 -c 'CCATGTCAACCACTTAG' *.fastq > count/B1827_Fm0.txt
agrep -0 -c 'TGAATGCCATCATTGCG' *.fastq > count/B1828_Fm0.txt
agrep -0 -c 'TCATTACCGGTAGACGT' *.fastq > count/B1830_Fm0.txt
agrep -0 -c 'TTCTACGTACTCTCAGT' *.fastq > count/B1831_Fm0.txt
agrep -0 -c 'ACCTGTTATTACCCCTG' *.fastq > count/B1832_Fm0.txt
agrep -0 -c 'GTCTGGGACTATTTCCC' *.fastq > count/B1833_Fm0.txt
agrep -0 -c 'AGCTCGTTTGGAGGGGA' *.fastq > count/B1834_Fm0.txt
agrep -0 -c 'TAAACCGTCTTTGGCCG' *.fastq > count/B1835_Fm0.txt
agrep -0 -c 'CCGAGCGGTTAGGAAAG' *.fastq > count/B1837_Fm0.txt
agrep -0 -c 'ATTCCTACTGTCGCAAA' *.fastq > count/B1838_Fm0.txt
agrep -0 -c 'GCCGACTCAAATTTTGT' *.fastq > count/B1839_Fm0.txt
agrep -0 -c 'AGCTCAAGACCAGAGAA' *.fastq > count/B1840_Fm0.txt
agrep -0 -c 'AATTCTGGGGGAGGCAC' *.fastq > count/B1841_Fm0.txt
agrep -0 -c 'TATGACGTCTCCACAGA' *.fastq > count/B1842_Fm0.txt
agrep -0 -c 'ACTTCCGTCCATATCTT' *.fastq > count/B1843_Fm0.txt
agrep -0 -c 'TAAATCCCTCTCACGGA' *.fastq > count/B1844_Fm0.txt
agrep -0 -c 'TATCTGATGACACCCGG' *.fastq > count/B1845_Fm0.txt
agrep -0 -c 'TTCTACTTACGAACTGC' *.fastq > count/B1846_Fm0.txt
agrep -0 -c 'GTTGCCGACCAAACACG' *.fastq > count/B1847_Fm0.txt
agrep -0 -c 'AGGCAACACGTTTACGC' *.fastq > count/B1848_Fm0.txt
agrep -0 -c 'AGGGCCCGATTGCCTAT' *.fastq > count/B1850_Fm0.txt
agrep -0 -c 'CTTGGCCTATTCATGAC' *.fastq > count/B1851_Fm0.txt
agrep -0 -c 'TCTAACCTCGGTAAGGG' *.fastq > count/B1853_Fm0.txt
agrep -0 -c 'TTAGGGTTTCCGAAAAG' *.fastq > count/B1854_Fm0.txt
agrep -0 -c 'CTTGATATTCCGTGACG' *.fastq > count/B1855_Fm0.txt
agrep -0 -c 'ACAGCATGCTATCGACG' *.fastq > count/B1856_Fm0.txt
agrep -0 -c 'TACTATCCTCTGCCTAT' *.fastq > count/B1857_Fm0.txt
agrep -0 -c 'GTGTATATCCGTAATCG' *.fastq > count/B1858_Fm0.txt
agrep -0 -c 'CATTGAGTTAAGTCCAC' *.fastq > count/B1859_Fm0.txt
agrep -0 -c 'TCTGGAGTGCGCCCAAT' *.fastq > count/B1860_Fm0.txt
agrep -0 -c 'TCGGGATTGGATCTGCA' *.fastq > count/B1862_Fm0.txt
agrep -0 -c 'TAGGGGTAATTGAGGCC' *.fastq > count/B1863_Fm0.txt
agrep -0 -c 'TACGAGAAGGGTCGTAT' *.fastq > count/B1864_Fm0.txt
agrep -0 -c 'CTCACATTTGGGCCCTT' *.fastq > count/B1865_Fm0.txt
agrep -0 -c 'GGTGGGTAAAGAAGGCG' *.fastq > count/B1867_Fm0.txt
agrep -0 -c 'TCGACGCTCGTAAGCGT' *.fastq > count/B1868_Fm0.txt
agrep -0 -c 'GGCGCGTCGTAATACAC' *.fastq > count/B1869_Fm0.txt
agrep -0 -c 'CTTGACTTGTGGTGGCG' *.fastq > count/B1871_Fm0.txt
agrep -0 -c 'TACATTACGGGATGACT' *.fastq > count/B1873_Fm0.txt
agrep -0 -c 'GATTCGACCGATTTGCA' *.fastq > count/B1874_Fm0.txt
agrep -0 -c 'TAAACGAACAGGTCTAC' *.fastq > count/B1875_Fm0.txt
agrep -0 -c 'AGCTGGGAGTTAAGCCC' *.fastq > count/B1876_Fm0.txt
agrep -0 -c 'GCAACCGCCGGTTATTC' *.fastq > count/B1878_Fm0.txt
agrep -0 -c 'TGACGTTTGGGACATAG' *.fastq > count/B1879_Fm0.txt
agrep -0 -c 'GGCTGCGGAAGAAAAAT' *.fastq > count/B1881_Fm0.txt
agrep -0 -c 'CTACCACATCCAACACG' *.fastq > count/B1882_Fm0.txt
agrep -0 -c 'CGCTAAAAGGGTAGCTA' *.fastq > count/B1883_Fm0.txt
agrep -0 -c 'GGATCGTACTACACAAA' *.fastq > count/B1884_Fm0.txt
agrep -0 -c 'AGACGATCGTTTGTGTA' *.fastq > count/B1885_Fm0.txt
agrep -0 -c 'TTGAAGTAGTAGCGATC' *.fastq > count/B1886_Fm0.txt
agrep -0 -c 'TGAGTACACCCATGCGC' *.fastq > count/B1887_Fm0.txt
agrep -0 -c 'CCTGGGCCCCTTATGAT' *.fastq > count/B1888_Fm0.txt
agrep -0 -c 'GGCGCTTTTTGACCGTC' *.fastq > count/B1889_Fm0.txt
agrep -0 -c 'CAGAGAATTGTAATCGG' *.fastq > count/B1890_Fm0.txt
agrep -0 -c 'AGACCGCCTGGGTAGAT' *.fastq > count/B1891_Fm0.txt
agrep -0 -c 'TCTCCCATCTGACTGTA' *.fastq > count/B1892_Fm0.txt
agrep -0 -c 'TATCGAATAAGCCTGCA' *.fastq > count/B1893_Fm0.txt
agrep -0 -c 'AGGAAGCTCAGTTCTCG' *.fastq > count/B1894_Fm0.txt
agrep -0 -c 'CCGGAATCTGGTTTAAT' *.fastq > count/B1895_Fm0.txt
agrep -0 -c 'CACGTGAGTCTTACGGC' *.fastq > count/B1896_Fm0.txt
agrep -0 -c 'TAAGCTTGGCGAAATCT' *.fastq > count/B1897_Fm0.txt
agrep -0 -c 'GCAACGGCTATACTAAG' *.fastq > count/B1898_Fm0.txt
agrep -0 -c 'GCACACTATGCTATTCA' *.fastq > count/B1900_Fm0.txt
agrep -0 -c 'CGCCATCATTTAGGCTG' *.fastq > count/B1901_Fm0.txt
agrep -0 -c 'GTTATAAGCGTGACGCT' *.fastq > count/B1903_Fm0.txt
agrep -0 -c 'TGGAGACACGCTGACTC' *.fastq > count/B1904_Fm0.txt
agrep -0 -c 'GACACTACCACTTCCGT' *.fastq > count/B1905_Fm0.txt
agrep -0 -c 'CGCCACGGTAGTAAACT' *.fastq > count/B1906_Fm0.txt
agrep -0 -c 'CGACCTATCATAAAGTC' *.fastq > count/B1907_Fm0.txt
agrep -0 -c 'CCGGAGTCAAAAGGGTG' *.fastq > count/B1908_Fm0.txt
agrep -0 -c 'AAGAGGAGCGGACTTGA' *.fastq > count/B1910_Fm0.txt
agrep -0 -c 'ACTTCGGTCTGTCATTG' *.fastq > count/B1911_Fm0.txt
agrep -0 -c 'CGGTAAGACAGTCTTGT' *.fastq > count/B1912_Fm0.txt
agrep -0 -c 'CTCCCGCAGATTGGATA' *.fastq > count/B1913_Fm0.txt
agrep -0 -c 'GTCGCGGATAGTAATTT' *.fastq > count/B1914_Fm0.txt
agrep -0 -c 'GATATGACGGCTTACGT' *.fastq > count/B1915_Fm0.txt
agrep -0 -c 'CAATGCGGCACGAGCTA' *.fastq > count/B1916_Fm0.txt
agrep -0 -c 'TTACCATCTCTCCATAG' *.fastq > count/B1917_Fm0.txt
agrep -0 -c 'CACAACGGTAGAGACTC' *.fastq > count/B1918_Fm0.txt
agrep -0 -c 'CATCGTAGTCGACGGGA' *.fastq > count/B1919_Fm0.txt
agrep -0 -c 'TCTATGCCTAGGTCTGA' *.fastq > count/B1920_Fm0.txt
agrep -0 -c 'ACTGGCGGAGCATAGAT' *.fastq > count/B1921_Fm0.txt
agrep -0 -c 'ATGTCCACGGAATTTGT' *.fastq > count/B1923_Fm0.txt
agrep -0 -c 'AAGATGGCCGAACTAGA' *.fastq > count/B1924_Fm0.txt
agrep -0 -c 'GACACCCAAGCACGTGA' *.fastq > count/B1926_Fm0.txt
agrep -0 -c 'AAGGATCGGGACAAGGA' *.fastq > count/B1927_Fm0.txt
agrep -0 -c 'TCTCAATCAAACACGCC' *.fastq > count/B1928_Fm0.txt
agrep -0 -c 'ACGCGTTTAGAGGAGCC' *.fastq > count/B1929_Fm0.txt
agrep -0 -c 'ACAGTCCTCGCGAGTAC' *.fastq > count/B1930_Fm0.txt
agrep -0 -c 'CGTGCAAATAACGCATT' *.fastq > count/B1933_Fm0.txt
agrep -0 -c 'CAATGTTGTCTTACGCA' *.fastq > count/B1935_Fm0.txt
agrep -0 -c 'CACCAATTGGTTTAGGA' *.fastq > count/B1936_Fm0.txt
agrep -0 -c 'GTATGGCGTACATCGTT' *.fastq > count/B1938_Fm0.txt
agrep -0 -c 'GCGCTAAGCAGCAGGTA' *.fastq > count/B1939_Fm0.txt
agrep -0 -c 'GACCTAAGAAAAAGGCG' *.fastq > count/B1941_Fm0.txt
agrep -0 -c 'ATTTGTCGACGGTTTCT' *.fastq > count/B1942_Fm0.txt
agrep -0 -c 'GCCTCAACGCGTCAATT' *.fastq > count/B1943_Fm0.txt
agrep -0 -c 'CGTCTGGCCGTAGATTA' *.fastq > count/B1944_Fm0.txt
agrep -0 -c 'CCGTCGGCATGAGTCTA' *.fastq > count/B1945_Fm0.txt
agrep -0 -c 'CACTTTGGCCACGAACT' *.fastq > count/B1947_Fm0.txt
agrep -0 -c 'CCGTTCATAGCATGTCC' *.fastq > count/B1948_Fm0.txt
agrep -0 -c 'ACGTAAATTGACGGCGC' *.fastq > count/B1949_Fm0.txt
agrep -0 -c 'GTTCACAATTCTGGTCG' *.fastq > count/B1950_Fm0.txt
agrep -0 -c 'CGTAAGTCTATGGTGGC' *.fastq > count/B1952_Fm0.txt
agrep -0 -c 'TAATATACGCGAACGGC' *.fastq > count/B1953_Fm0.txt
agrep -0 -c 'ACCCGGTATGTCTTTTC' *.fastq > count/B1956_Fm0.txt
agrep -0 -c 'CTTAGCTATCACCCATT' *.fastq > count/B1957_Fm0.txt
agrep -0 -c 'ACGCAAACTAAGAGGTA' *.fastq > count/B1958_Fm0.txt
agrep -0 -c 'ATGCATCAGGTCCTGTT' *.fastq > count/B1960_Fm0.txt
agrep -0 -c 'ACAACAATGAGCATGCA' *.fastq > count/B1961_Fm0.txt
agrep -0 -c 'CACGCGTGCTGTCATAA' *.fastq > count/B1962_Fm0.txt
agrep -0 -c 'GAGCTCCTACGCAACAC' *.fastq > count/B1963_Fm0.txt
agrep -0 -c 'CGTAATATGCCACGTCT' *.fastq > count/B1964_Fm0.txt
agrep -0 -c 'GAGATATCTATTCCTCG' *.fastq > count/B1965_Fm0.txt
agrep -0 -c 'TGTATGGCCTCGAAGAA' *.fastq > count/B1967_Fm0.txt
agrep -0 -c 'GCTCTTTATAACGTCGC' *.fastq > count/B1968_Fm0.txt
agrep -0 -c 'CTATCGTATGCAGAGGC' *.fastq > count/B1971_Fm0.txt
agrep -0 -c 'CTGAATATCTTGTACCG' *.fastq > count/B1972_Fm0.txt
agrep -0 -c 'GGCATAAAGAGCTGCTA' *.fastq > count/B1973_Fm0.txt
agrep -0 -c 'TGGATAACCTGCCGGAA' *.fastq > count/B1974_Fm0.txt
agrep -0 -c 'ATTCCATCGTGAAGTGG' *.fastq > count/B1975_Fm0.txt
agrep -0 -c 'ATTCCTTTCGGCTGTAG' *.fastq > count/B1977_Fm0.txt
agrep -0 -c 'GACGGGTGATTATACCC' *.fastq > count/B1978_Fm0.txt
agrep -0 -c 'AAGTTCCTAGCGCTAAG' *.fastq > count/B1979_Fm0.txt
agrep -0 -c 'TGCCATCCGGATAGTTT' *.fastq > count/B1980_Fm0.txt
agrep -0 -c 'CAGCCGCAATGAATCGC' *.fastq > count/B1982_Fm0.txt
agrep -0 -c 'GTGAAATAGCGATCACG' *.fastq > count/B1983_Fm0.txt
agrep -0 -c 'AGCGGTAATGCCTGATG' *.fastq > count/B1984_Fm0.txt
agrep -0 -c 'GTCGTTGCAGAATCGCC' *.fastq > count/B1985_Fm0.txt
agrep -0 -c 'CGAGGAAAGCATCCTTG' *.fastq > count/B1986_Fm0.txt
agrep -0 -c 'TGCAGGCAAATTAGTAC' *.fastq > count/B1987_Fm0.txt
agrep -0 -c 'GTGCACTAGACCAATTC' *.fastq > count/B1988_Fm0.txt
agrep -0 -c 'AATATCTCTCGCCCGTT' *.fastq > count/B1989_Fm0.txt
agrep -0 -c 'CGGACTAGTTATACCGA' *.fastq > count/B1990_Fm0.txt
agrep -0 -c 'CAATAGCATTGCAGACA' *.fastq > count/B1992_Fm0.txt
agrep -0 -c 'AAGATCCCCTTTCGTGA' *.fastq > count/B1993_Fm0.txt
agrep -0 -c 'TGAGGAGTTATGCGGTG' *.fastq > count/B1994_Fm0.txt
agrep -0 -c 'CACCGATCCCCTACTAA' *.fastq > count/B1995_Fm0.txt
agrep -0 -c 'CGTTCGTTTCGATCAGG' *.fastq > count/B1996_Fm0.txt
agrep -0 -c 'TAACTGCCCGGCATTTG' *.fastq > count/B1997_Fm0.txt
agrep -0 -c 'TACACATGCGTCGGTGG' *.fastq > count/B1998_Fm0.txt
agrep -0 -c 'CGTCAAAGTAGAAGCGG' *.fastq > count/B2000_Fm0.txt
agrep -0 -c 'AGATCGGCCATAGGTAG' *.fastq > count/B2001_Fm0.txt
agrep -0 -c 'TCTTGGAGTGAGTATAC' *.fastq > count/B2002_Fm0.txt
agrep -0 -c 'GCTTCTACAGGCTCTTT' *.fastq > count/B2004_Fm0.txt
agrep -0 -c 'GTCAAGCACGCTACGAG' *.fastq > count/B2005_Fm0.txt
agrep -0 -c 'CGTGTATGTGATGGTGA' *.fastq > count/B2006_Fm0.txt
agrep -0 -c 'TGGATACCGCTTGAAGC' *.fastq > count/B2008_Fm0.txt
agrep -0 -c 'AGCTCGGCTCGATATGC' *.fastq > count/B2009_Fm0.txt
agrep -0 -c 'AATGGAGCCCAACCCTC' *.fastq > count/B2011_Fm0.txt
agrep -0 -c 'GTCTTACCATCGCGTCC' *.fastq > count/B2012_Fm0.txt
agrep -0 -c 'AGCGGCTCATGGTCAGA' *.fastq > count/B2014_Fm0.txt
agrep -0 -c 'GACTACTCTACTCTGGG' *.fastq > count/B2016_Fm0.txt
agrep -0 -c 'AAGCGCAGAAACTGGTT' *.fastq > count/B2017_Fm0.txt
agrep -0 -c 'GACATAGTGCGAAGCGT' *.fastq > count/B2019_Fm0.txt
agrep -0 -c 'CCAGTTGCCGCCAAGTT' *.fastq > count/B2020_Fm0.txt
agrep -0 -c 'GTATTAGGCTTAGCACT' *.fastq > count/B2022_Fm0.txt
agrep -0 -c 'GCCGTTGAACCTCCCAA' *.fastq > count/B2023_Fm0.txt
agrep -0 -c 'GTGAGATAACACGGGAC' *.fastq > count/B2024_Fm0.txt
agrep -0 -c 'AACCAACCGGGAATATA' *.fastq > count/B2026_Fm0.txt
agrep -0 -c 'CAGCCCTGGTTGAACTC' *.fastq > count/B2027_Fm0.txt
agrep -0 -c 'CGTGCTATGGCTTCGAC' *.fastq > count/B2028_Fm0.txt
agrep -0 -c 'AGTCTACACTCGGGAGA' *.fastq > count/B2029_Fm0.txt
agrep -0 -c 'ATTGTGACTGGGGTTTA' *.fastq > count/B2030_Fm0.txt
agrep -0 -c 'ACCTCTGACCGTCGAAC' *.fastq > count/B2031_Fm0.txt
agrep -0 -c 'TGACGACGTTTTACCAG' *.fastq > count/B2032_Fm0.txt
agrep -0 -c 'TAATGTACGGCGCGAGA' *.fastq > count/B2033_Fm0.txt
agrep -0 -c 'TGCGCCCAACCACGATA' *.fastq > count/B2034_Fm0.txt
agrep -0 -c 'TTATCACTAGAGGGCTA' *.fastq > count/B2036_Fm0.txt
agrep -0 -c 'CACGGATGACTTCGATT' *.fastq > count/B2037_Fm0.txt
agrep -0 -c 'CGAGATCCTTGCCTTCA' *.fastq > count/B2038_Fm0.txt
agrep -0 -c 'CCTTGTGTTTGTCCGCA' *.fastq > count/B2039_Fm0.txt
agrep -0 -c 'CGTTGTTATTCGTGTAG' *.fastq > count/B2040_Fm0.txt
agrep -0 -c 'TGCTGCGGTCTAAGCTC' *.fastq > count/B2041_Fm0.txt
agrep -0 -c 'TAGGGTGAGCCTTAGCA' *.fastq > count/B2042_Fm0.txt
agrep -0 -c 'CGTACCGAATCTCTCTA' *.fastq > count/B2043_Fm0.txt
agrep -0 -c 'CTTGTGCGGTAACTGCA' *.fastq > count/B2044_Fm0.txt
agrep -0 -c 'TCTTCAAACCGTCTGGG' *.fastq > count/B2045_Fm0.txt
agrep -0 -c 'CAGGTGATAGCCGTAGC' *.fastq > count/B2046_Fm0.txt
agrep -0 -c 'TAGCGCACATGCTCGAT' *.fastq > count/B2049_Fm0.txt
agrep -0 -c 'ATGATAGGCGTCCTGTC' *.fastq > count/B2050_Fm0.txt
agrep -0 -c 'AGTTACGATATTCTCCC' *.fastq > count/B2051_Fm0.txt
agrep -0 -c 'CCTCTTTTGAATTGGGT' *.fastq > count/B2052_Fm0.txt
agrep -0 -c 'CGCTTAGTTGCCTAGAG' *.fastq > count/B2053_Fm0.txt
agrep -0 -c 'GACCGCTTAGAGATCCA' *.fastq > count/B2054_Fm0.txt
agrep -0 -c 'ATTACACAGCAGCTGTG' *.fastq > count/B2055_Fm0.txt
agrep -0 -c 'GGTACATTCCGGAAGAA' *.fastq > count/B2056_Fm0.txt
agrep -0 -c 'GTACCGAGGATGCAAGA' *.fastq > count/B2057_Fm0.txt
agrep -0 -c 'TTCATCCGCCAAGGATC' *.fastq > count/B2058_Fm0.txt
agrep -0 -c 'CCATTACACTTCCGTCG' *.fastq > count/B2059_Fm0.txt
agrep -0 -c 'AATGTTACGCGTACAGT' *.fastq > count/B2060_Fm0.txt
agrep -0 -c 'AGAGATTAGGCGAAGAT' *.fastq > count/B2061_Fm0.txt
agrep -0 -c 'TTAGCTTACCTGCGGCG' *.fastq > count/B2064_Fm0.txt
agrep -0 -c 'GCGGTACTATCTGATTT' *.fastq > count/B2067_Fm0.txt
agrep -0 -c 'CGCGTAGAGCCATACTT' *.fastq > count/B2068_Fm0.txt
agrep -0 -c 'TTCTTCAAGCTAATGCG' *.fastq > count/B2069_Fm0.txt
agrep -0 -c 'CAATGGAGTTCAACTTC' *.fastq > count/B2070_Fm0.txt
agrep -0 -c 'CGTAGTATTGCGTTGAA' *.fastq > count/B2071_Fm0.txt
agrep -0 -c 'TGTCACTGACGTCACGA' *.fastq > count/B2074_Fm0.txt
agrep -0 -c 'CCTATCGACACTCCGCA' *.fastq > count/B2076_Fm0.txt
agrep -0 -c 'TCTTGAGGGTAAGAGGT' *.fastq > count/B2077_Fm0.txt
agrep -0 -c 'TCTGCCTGCCCCGATTA' *.fastq > count/B2078_Fm0.txt
agrep -0 -c 'TGCTCAGTGAGCGTAGG' *.fastq > count/B2079_Fm0.txt
agrep -0 -c 'ATAGCTGCCAGCCGCAT' *.fastq > count/B2080_Fm0.txt
agrep -0 -c 'GACCCAAACGAGACGAA' *.fastq > count/B2081_Fm0.txt
agrep -0 -c 'AAGGAGTAGATGCAGTT' *.fastq > count/B2084_Fm0.txt
agrep -0 -c 'TAACCCGCGCTGGAATC' *.fastq > count/B2085_Fm0.txt
agrep -0 -c 'TTCCTATTCGACGCGTT' *.fastq > count/B2087_Fm0.txt
agrep -0 -c 'TGAGGGCCATACCCTGT' *.fastq > count/B2088_Fm0.txt
agrep -0 -c 'ACTCCGGTTATTACCGG' *.fastq > count/B2089_Fm0.txt
agrep -0 -c 'TGATAGTGCCCCAGCGT' *.fastq > count/B2090_Fm0.txt
agrep -0 -c 'GTCCGCATTAAAGCGTA' *.fastq > count/B2091_Fm0.txt
agrep -0 -c 'ACGTTGTCCCTTGAGTA' *.fastq > count/B2092_Fm0.txt
agrep -0 -c 'CGGGTGTTAGCACTTCT' *.fastq > count/B2093_Fm0.txt
agrep -0 -c 'TCGACTATGCACGTCGA' *.fastq > count/B2094_Fm0.txt
agrep -0 -c 'GTCTCAAACGGCTTCGC' *.fastq > count/B2095_Fm0.txt
agrep -0 -c 'ATTCACAAAGCGTCGCG' *.fastq > count/B2096_Fm0.txt
agrep -0 -c 'TTGATCCCGACCGCTTT' *.fastq > count/B2097_Fm0.txt
agrep -0 -c 'CAAATCAAGCCCCCTCG' *.fastq > count/B2098_Fm0.txt
agrep -0 -c 'TCATAGGGTAGAGCACG' *.fastq > count/B2099_Fm0.txt
agrep -0 -c 'GACCGGTTTTTCAGATT' *.fastq > count/B2100_Fm0.txt
agrep -0 -c 'CACCTTCAAAGTTGTAG' *.fastq > count/B2101_Fm0.txt
agrep -0 -c 'CACAGAGGGATAGGGCA' *.fastq > count/B2103_Fm0.txt
agrep -0 -c 'CTACGCAACGGACCTAC' *.fastq > count/B2104_Fm0.txt
agrep -0 -c 'TAACTGAAAGGGCCTTC' *.fastq > count/B2105_Fm0.txt
agrep -0 -c 'TCTGCTATACGAGAATC' *.fastq > count/B2106_Fm0.txt
agrep -0 -c 'AGCGTCTCAGTCGTTCG' *.fastq > count/B2107_Fm0.txt
agrep -0 -c 'GCGCTATATTGGGAAGC' *.fastq > count/B2108_Fm0.txt
agrep -0 -c 'GAGAAGCTAGAACCGCC' *.fastq > count/B2109_Fm0.txt
agrep -0 -c 'GTTAGAGCGCTTTGTTG' *.fastq > count/B2110_Fm0.txt
agrep -0 -c 'GCAATGGCGACTGATTC' *.fastq > count/B2111_Fm0.txt
agrep -0 -c 'TGAAGTATGTGGTCTTG' *.fastq > count/B2112_Fm0.txt
agrep -0 -c 'TATAGGCGTCAAGAGAC' *.fastq > count/B2114_Fm0.txt
agrep -0 -c 'GAGTAGGTTATGGGGTC' *.fastq > count/B2116_Fm0.txt
agrep -0 -c 'CCTCCCATAGTTTTGCG' *.fastq > count/B2118_Fm0.txt
agrep -0 -c 'GAGTCTATGCTCGTCAG' *.fastq > count/B2119_Fm0.txt
agrep -0 -c 'AAGCGTTTTGCTCATCG' *.fastq > count/B2120_Fm0.txt
agrep -0 -c 'CGTGCCGGATCATGATA' *.fastq > count/B2121_Fm0.txt
agrep -0 -c 'TAGACTGGCGCGATTCA' *.fastq > count/B2122_Fm0.txt
agrep -0 -c 'CGTATCCCTCACAGCCT' *.fastq > count/B2124_Fm0.txt
agrep -0 -c 'GACCAGGTCACTGCTAA' *.fastq > count/B2125_Fm0.txt
agrep -0 -c 'ACCGGGGGACCCTATTT' *.fastq > count/B2126_Fm0.txt
agrep -0 -c 'GATGTGCCTAATGGACG' *.fastq > count/B2127_Fm0.txt
agrep -0 -c 'ACGAGCCTTAGGGTAGC' *.fastq > count/B2128_Fm0.txt
agrep -0 -c 'AGTCTGTAGCGTGCAAC' *.fastq > count/B2129_Fm0.txt
agrep -0 -c 'TACAAAAACGCACTCAG' *.fastq > count/B2130_Fm0.txt
agrep -0 -c 'CGTACTAGCGTCTTTCT' *.fastq > count/B2131_Fm0.txt
agrep -0 -c 'TTGGTGCGTACCATCTC' *.fastq > count/B2132_Fm0.txt
agrep -0 -c 'TCTCGAACGAGGACTTC' *.fastq > count/B2135_Fm0.txt
agrep -0 -c 'ACGCACACGAAATCAAT' *.fastq > count/B2137_Fm0.txt
agrep -0 -c 'ATAAACGCGGTTATGCT' *.fastq > count/B2138_Fm0.txt
agrep -0 -c 'TAGGGACCGTGTGACTA' *.fastq > count/B2139_Fm0.txt
agrep -0 -c 'TCGGTGTTTACTACGTG' *.fastq > count/B2140_Fm0.txt
agrep -0 -c 'TAAACCATGTGCGCTAT' *.fastq > count/B2141_Fm0.txt
agrep -0 -c 'ATACGTATGCCCGTACT' *.fastq > count/B2142_Fm0.txt
agrep -0 -c 'ATGCCTCGCCCCTAACA' *.fastq > count/B2144_Fm0.txt
agrep -0 -c 'TACCAAGTGTAAGTTGG' *.fastq > count/B2145_Fm0.txt
agrep -0 -c 'TTGTAAGCGGTCAACTG' *.fastq > count/B2146_Fm0.txt
agrep -0 -c 'CAGCGCTGCAGATACCA' *.fastq > count/B2148_Fm0.txt
agrep -0 -c 'TTAGGATTCTGCCGCTA' *.fastq > count/B2149_Fm0.txt
agrep -0 -c 'ATGAGGTTCTTAAGTCG' *.fastq > count/B2150_Fm0.txt
agrep -0 -c 'ACATCCTCCTAGGCTAG' *.fastq > count/B2151_Fm0.txt
agrep -0 -c 'GCATACTATGCCTATGG' *.fastq > count/B2152_Fm0.txt
agrep -0 -c 'TTCACTCAATTCACCCG' *.fastq > count/B2153_Fm0.txt
agrep -0 -c 'CACATCTCCGTTGAACG' *.fastq > count/B2154_Fm0.txt
agrep -0 -c 'GTACCTAATTCTCGTAG' *.fastq > count/B2155_Fm0.txt
agrep -0 -c 'CGCTGTATCAAATTGCG' *.fastq > count/B2157_Fm0.txt
agrep -0 -c 'ACGACAGCGACTGTACC' *.fastq > count/B2158_Fm0.txt
agrep -0 -c 'CATTGGCCGTCAACTTT' *.fastq > count/B2159_Fm0.txt
agrep -0 -c 'GTCACCATTTCATCGCC' *.fastq > count/B2160_Fm0.txt
agrep -0 -c 'AGAGGTCGGTAGCTATA' *.fastq > count/B2161_Fm0.txt
agrep -0 -c 'TTAGTAAATCCCGCGGG' *.fastq > count/B2162_Fm0.txt
agrep -0 -c 'ATTACGCGTCCGAAGTC' *.fastq > count/B2163_Fm0.txt
agrep -0 -c 'CCGGCATTATTGCTTAA' *.fastq > count/B2164_Fm0.txt
agrep -0 -c 'TTGGAGCTGACCCCTTA' *.fastq > count/B2165_Fm0.txt
agrep -0 -c 'TTACATAAGGTCCCCAA' *.fastq > count/B2166_Fm0.txt
agrep -0 -c 'CGATGTCGTTAACGGAT' *.fastq > count/B2167_Fm0.txt
agrep -0 -c 'TAATGGCCGAGCCCATT' *.fastq > count/B2170_Fm0.txt
agrep -0 -c 'GCGCACTGTAATCGAAA' *.fastq > count/B2171_Fm0.txt
agrep -0 -c 'CAGACTTCCACCCCGTT' *.fastq > count/B2172_Fm0.txt
agrep -0 -c 'TTAATAGTCCGGGCTAG' *.fastq > count/B2173_Fm0.txt
agrep -0 -c 'CATGACGCGTTAGAGCG' *.fastq > count/B2174_Fm0.txt
agrep -0 -c 'CCTAAGCTGGCTGGGTA' *.fastq > count/B2175_Fm0.txt
agrep -0 -c 'TCGTTCATGAGTTGGAT' *.fastq > count/B2176_Fm0.txt
agrep -0 -c 'GAGGTTTCACCAGGTCA' *.fastq > count/B2177_Fm0.txt
agrep -0 -c 'CAGCAAAATGACCTACG' *.fastq > count/B2179_Fm0.txt
agrep -0 -c 'AACACTCTATGCGGAGT' *.fastq > count/B2180_Fm0.txt
agrep -0 -c 'AGGACTCTCGAGTGTCA' *.fastq > count/B2181_Fm0.txt
agrep -0 -c 'CCTGTGAATGAAACGCT' *.fastq > count/B2182_Fm0.txt
agrep -0 -c 'ATGCGTTATTCTCCACC' *.fastq > count/B2183_Fm0.txt
agrep -0 -c 'ACAATAAGGCGCTCATG' *.fastq > count/B2184_Fm0.txt
agrep -0 -c 'TTGGTATTGACTACCCT' *.fastq > count/B2185_Fm0.txt
agrep -0 -c 'GTAACCTCGGCACAATA' *.fastq > count/B2186_Fm0.txt
agrep -0 -c 'ATGGCCACTACCTTCCC' *.fastq > count/B2187_Fm0.txt
agrep -0 -c 'ATAGGACGATGGGATTC' *.fastq > count/B2188_Fm0.txt
agrep -0 -c 'AGCGCACAATCGTATCA' *.fastq > count/B2189_Fm0.txt
agrep -0 -c 'TCGGCAGCAGTTATTTT' *.fastq > count/B2190_Fm0.txt
agrep -0 -c 'GCAGCCATCCTAGAACC' *.fastq > count/B2192_Fm0.txt
agrep -0 -c 'TAAAGTAGGACGTAGAC' *.fastq > count/B2194_Fm0.txt
agrep -0 -c 'CCGATATAGTAGACGGA' *.fastq > count/B2195_Fm0.txt
agrep -0 -c 'ACTGATACCCAAGTCTG' *.fastq > count/B2196_Fm0.txt
agrep -0 -c 'TTAGCACGATTACCGAG' *.fastq > count/B2197_Fm0.txt
agrep -0 -c 'AGCTTGGCGCATTGTGA' *.fastq > count/B2198_Fm0.txt
agrep -0 -c 'AGAAGGGACGATGGATT' *.fastq > count/B2200_Fm0.txt
agrep -0 -c 'ACGGCAAAGTAAGTCCA' *.fastq > count/B2201_Fm0.txt
agrep -0 -c 'ATCACGATAATTCGTGC' *.fastq > count/B2202_Fm0.txt
agrep -0 -c 'ATAACTGTAAACGGTCC' *.fastq > count/B2203_Fm0.txt
agrep -0 -c 'TCGTGACCTATAACGCT' *.fastq > count/B2204_Fm0.txt
agrep -0 -c 'AATGCTTTACTGCGCCC' *.fastq > count/B2205_Fm0.txt
agrep -0 -c 'ACGCTCTGCAGTGAGCT' *.fastq > count/B2206_Fm0.txt
agrep -0 -c 'CCAAGAAACTGGACCGC' *.fastq > count/B2207_Fm0.txt
agrep -0 -c 'GATTCCGGTTGAACCAA' *.fastq > count/B2208_Fm0.txt
agrep -0 -c 'TCGATGCCCGTATCACC' *.fastq > count/B2209_Fm0.txt
agrep -0 -c 'CAATACATGCTCCTCAA' *.fastq > count/B2210_Fm0.txt
agrep -0 -c 'ATCATGCGGGGATTCTT' *.fastq > count/B2211_Fm0.txt
agrep -0 -c 'ACCATCGATATACTCTC' *.fastq > count/B2212_Fm0.txt
agrep -0 -c 'TTGGAAGGACTAGCGCA' *.fastq > count/B2213_Fm0.txt
agrep -0 -c 'GTGGCGACATTCCTGTT' *.fastq > count/B2214_Fm0.txt
agrep -0 -c 'CGGGACTTACTGTGTGG' *.fastq > count/B2215_Fm0.txt
agrep -0 -c 'CGTACCCGCGTGTAGAT' *.fastq > count/B2217_Fm0.txt
agrep -0 -c 'TAGAAGCTAAAGCGCAA' *.fastq > count/B2218_Fm0.txt
agrep -0 -c 'TCGGGGAACTTTATTCC' *.fastq > count/B2219_Fm0.txt
agrep -0 -c 'CTCCGGTTCTTTAAGAT' *.fastq > count/B2220_Fm0.txt
agrep -0 -c 'GTCACGCTCACAGATCA' *.fastq > count/B2221_Fm0.txt
agrep -0 -c 'ACGGACCATGTTGTCGA' *.fastq > count/B2222_Fm0.txt
agrep -0 -c 'CCACTCTTCAGACGCAC' *.fastq > count/B2223_Fm0.txt
agrep -0 -c 'CACAGTGACCCACGATT' *.fastq > count/B2225_Fm0.txt
agrep -0 -c 'CGACAGCTTATGGTTGC' *.fastq > count/B2227_Fm0.txt
agrep -0 -c 'TACTCGCATTATCCTAG' *.fastq > count/B2228_Fm0.txt
agrep -0 -c 'GCCAAATAACTTGGTCT' *.fastq > count/B2229_Fm0.txt
agrep -0 -c 'GCAACACTATTGGTCAA' *.fastq > count/B2230_Fm0.txt
agrep -0 -c 'TGTGCTCCTTAGCCACC' *.fastq > count/B2231_Fm0.txt
agrep -0 -c 'ACAAAGACCTGGACCTA' *.fastq > count/B2233_Fm0.txt
agrep -0 -c 'GGTTTGCTACGCCAGAT' *.fastq > count/B2234_Fm0.txt
agrep -0 -c 'AGTGAATCCATCTGTGT' *.fastq > count/B2235_Fm0.txt
agrep -0 -c 'AACCGCGATGGTACATT' *.fastq > count/B2236_Fm0.txt
agrep -0 -c 'TACTGGCTGCATCAAAA' *.fastq > count/B2237_Fm0.txt
agrep -0 -c 'CGATTTGGCGACCAATG' *.fastq > count/B2238_Fm0.txt
agrep -0 -c 'GAGATTGACACAAAGTG' *.fastq > count/B2239_Fm0.txt
agrep -0 -c 'CGGGGTACCGAGAACAA' *.fastq > count/B2240_Fm0.txt
agrep -0 -c 'GTGTGAATCAAGCAATC' *.fastq > count/B2241_Fm0.txt
agrep -0 -c 'GTAGTTTTAAGGGGCTA' *.fastq > count/B2242_Fm0.txt
agrep -0 -c 'CCAACTTTGGCACCTCG' *.fastq > count/B2243_Fm0.txt
agrep -0 -c 'TACCCTACCGATCGTTC' *.fastq > count/B2244_Fm0.txt
agrep -0 -c 'CTCGAACCGGGAATATT' *.fastq > count/B2245_Fm0.txt
agrep -0 -c 'TTCGTGTCGACAGCAAC' *.fastq > count/B2246_Fm0.txt
agrep -0 -c 'GCGATCACTTAGCCACA' *.fastq > count/B2247_Fm0.txt
agrep -0 -c 'AGTTAGGACGGGTCGGT' *.fastq > count/B2248_Fm0.txt
agrep -0 -c 'CTACGGAGGTGCTGCAA' *.fastq > count/B2249_Fm0.txt
agrep -0 -c 'CGATAGTCAGGATTAGA' *.fastq > count/B2250_Fm0.txt
agrep -0 -c 'CAAATTCACACGCCCTA' *.fastq > count/B2251_Fm0.txt
agrep -0 -c 'GTACGGCTCTGCTAGTG' *.fastq > count/B2252_Fm0.txt
agrep -0 -c 'GTCTGATAGCTTCGATA' *.fastq > count/B2254_Fm0.txt
agrep -0 -c 'TGCCAAAGTTGCACAAT' *.fastq > count/B2256_Fm0.txt
agrep -0 -c 'CATGACTATCATCACTC' *.fastq > count/B2258_Fm0.txt
agrep -0 -c 'TACGGTTTGTGGCATAG' *.fastq > count/B2260_Fm0.txt
agrep -0 -c 'ATTGTTAGGCCTTCGTG' *.fastq > count/B2261_Fm0.txt
agrep -0 -c 'CAGGTTTAACGGTTTAG' *.fastq > count/B2262_Fm0.txt
agrep -0 -c 'CGCGTGAAAAACTGTTA' *.fastq > count/B2263_Fm0.txt
agrep -0 -c 'AAGGAGGACTGGTGGTA' *.fastq > count/B2264_Fm0.txt
agrep -0 -c 'GTCTTCCTATGCTAGTT' *.fastq > count/B2265_Fm0.txt
agrep -0 -c 'TCGGGTATTCGTGTCGT' *.fastq > count/B2266_Fm0.txt
agrep -0 -c 'TGAATCGATCTACGGTC' *.fastq > count/B2267_Fm0.txt
agrep -0 -c 'CGCCTTATGACTGTGTG' *.fastq > count/B2268_Fm0.txt
agrep -0 -c 'TTCTGATGGGGGGCAAG' *.fastq > count/B2269_Fm0.txt
agrep -0 -c 'CTATGAGCTGGCGCTTA' *.fastq > count/B2270_Fm0.txt
agrep -0 -c 'ATGTCTGCGGGAAAGGC' *.fastq > count/B2272_Fm0.txt
agrep -0 -c 'TAGCCACCGGATCAGAT' *.fastq > count/B2273_Fm0.txt
agrep -0 -c 'TAAATTGTGGGACTCTC' *.fastq > count/B2274_Fm0.txt
agrep -0 -c 'AACGTATCGCATCCTGA' *.fastq > count/B2275_Fm0.txt
agrep -0 -c 'GATGTTGGGGACCTGTT' *.fastq > count/B2277_Fm0.txt
agrep -0 -c 'ATGTACGCGGATGCATG' *.fastq > count/B2278_Fm0.txt
agrep -0 -c 'GGAGTCCTGGTGCCTAA' *.fastq > count/B2279_Fm0.txt
agrep -0 -c 'CGTTAAATAGACGCCTG' *.fastq > count/B2280_Fm0.txt
agrep -0 -c 'TGATTGGCTACCGTAGA' *.fastq > count/B2281_Fm0.txt
agrep -0 -c 'TCGCGTGACCATTACAT' *.fastq > count/B2282_Fm0.txt
agrep -0 -c 'AGGAGATAGGCAAACCC' *.fastq > count/B2283_Fm0.txt
agrep -0 -c 'CTAGGACTTATCTCGCT' *.fastq > count/B2284_Fm0.txt
agrep -0 -c 'AGAGATGGTCATCGAAA' *.fastq > count/B2286_Fm0.txt
agrep -0 -c 'CCTGAGATCGTTCAATT' *.fastq > count/B2287_Fm0.txt
agrep -0 -c 'TTCACCCATGAGACCCA' *.fastq > count/B2288_Fm0.txt
agrep -0 -c 'CGCAAAGCTTTTACACT' *.fastq > count/B2289_Fm0.txt
agrep -0 -c 'AGGAGCTTATCCGTTCA' *.fastq > count/B2290_Fm0.txt
agrep -0 -c 'GATCGTGAGTCCGCTTA' *.fastq > count/B2291_Fm0.txt
agrep -0 -c 'TTAAGAAGAGCGGATGA' *.fastq > count/B2293_Fm0.txt
agrep -0 -c 'TAGTGAAAAGCCGGGTA' *.fastq > count/B2294_Fm0.txt
agrep -0 -c 'GCCTATATGTAACCTAG' *.fastq > count/B2295_Fm0.txt
agrep -0 -c 'GAGTTCCGTCCGTAACA' *.fastq > count/B2296_Fm0.txt
agrep -0 -c 'TCAGTTGATACACAGAG' *.fastq > count/B2297_Fm0.txt
agrep -0 -c 'TGACCGTTTCCTATCTC' *.fastq > count/B2298_Fm0.txt
agrep -0 -c 'CGAAGTTCTGCTATTGG' *.fastq > count/B2299_Fm0.txt
agrep -0 -c 'AGACGAACATGAGGACA' *.fastq > count/B2300_Fm0.txt
agrep -0 -c 'GTCACAACTAACAGGTC' *.fastq > count/B2303_Fm0.txt
agrep -0 -c 'TGAATGCTTGTTACCGG' *.fastq > count/B2304_Fm0.txt
agrep -0 -c 'TTGAGTCATACGACCCG' *.fastq > count/B2305_Fm0.txt
agrep -0 -c 'GCCTGTTTGATCTATGC' *.fastq > count/B2306_Fm0.txt
agrep -0 -c 'CCGGAGCGGATTAATAA' *.fastq > count/B2307_Fm0.txt
agrep -0 -c 'CTATTCCGCGTATGGTC' *.fastq > count/B2308_Fm0.txt
agrep -0 -c 'ATATACCCTCCGGGGTG' *.fastq > count/B2309_Fm0.txt
agrep -0 -c 'CGCACTTAGCCCAAACA' *.fastq > count/B2310_Fm0.txt
agrep -0 -c 'TGGACCAAGTTTTGCAT' *.fastq > count/B2311_Fm0.txt
agrep -0 -c 'CTGTGATACAGCGGGTT' *.fastq > count/B2312_Fm0.txt
agrep -0 -c 'ACGTGGAACGGCTTTAG' *.fastq > count/B2313_Fm0.txt
agrep -0 -c 'TAGTTGACGCGATTGAT' *.fastq > count/B2314_Fm0.txt
agrep -0 -c 'GACCGCAAGAAATCACG' *.fastq > count/B2315_Fm0.txt
agrep -0 -c 'ACAGGGATTGGGCGGAT' *.fastq > count/B2316_Fm0.txt
agrep -0 -c 'ATAAGAATACGGGCCTA' *.fastq > count/B2317_Fm0.txt
agrep -0 -c 'CGCAAACAGAGGGCAGT' *.fastq > count/B2318_Fm0.txt
agrep -0 -c 'CTTGGTTAGATAACCTG' *.fastq > count/B2319_Fm0.txt
agrep -0 -c 'CTTCAGTCGGGAAGTGG' *.fastq > count/B2320_Fm0.txt
agrep -0 -c 'GTACTTGGGTATTGCCC' *.fastq > count/B2321_Fm0.txt
agrep -0 -c 'ATAGATTCTGGCACTGG' *.fastq > count/B2322_Fm0.txt
agrep -0 -c 'AATGACGTAACCTCATC' *.fastq > count/B2323_Fm0.txt
agrep -0 -c 'ATGAATTGGGGGAGTAC' *.fastq > count/B2324_Fm0.txt
agrep -0 -c 'AGGTTTGGCTGGTGGTG' *.fastq > count/B2325_Fm0.txt
agrep -0 -c 'GTGCAGCCCTAGTACCT' *.fastq > count/B2326_Fm0.txt
agrep -0 -c 'GTATTGCCGACCAACCG' *.fastq > count/B2327_Fm0.txt
agrep -0 -c 'CGGGATCATAACTGCTC' *.fastq > count/B2328_Fm0.txt
agrep -0 -c 'AGCGCATATGAGCTAGG' *.fastq > count/B2329_Fm0.txt
agrep -0 -c 'TGATTTGGCGATAGTCA' *.fastq > count/B2330_Fm0.txt
agrep -0 -c 'AGTAGCCCACAAGCGTG' *.fastq > count/B2331_Fm0.txt
agrep -0 -c 'GTTTGAGGTCCGAGCTT' *.fastq > count/B2332_Fm0.txt
agrep -0 -c 'GCGCAATTTCCCAACCC' *.fastq > count/B2333_Fm0.txt
agrep -0 -c 'GTATCCAACTTGTAGTG' *.fastq > count/B2335_Fm0.txt
agrep -0 -c 'AGTCGTCGTTTGGGATA' *.fastq > count/B2336_Fm0.txt
agrep -0 -c 'AACTCAACCACAGTTCT' *.fastq > count/B2337_Fm0.txt
agrep -0 -c 'AGTGACCGCGTTACGGT' *.fastq > count/B2340_Fm0.txt
agrep -0 -c 'AACGCGGGCATGAAGCT' *.fastq > count/B2341_Fm0.txt
agrep -0 -c 'AATGACTGCCCCTTCGA' *.fastq > count/B2342_Fm0.txt
agrep -0 -c 'TTCCAATAGCCGTTACC' *.fastq > count/B2343_Fm0.txt
agrep -0 -c 'CAGAACTGACGGCCTTT' *.fastq > count/B2345_Fm0.txt
agrep -0 -c 'CTTCCTGTAGACGAATA' *.fastq > count/B2346_Fm0.txt
agrep -0 -c 'CTCAAGCCGCAGTTATC' *.fastq > count/B2347_Fm0.txt
agrep -0 -c 'GATATAGAGACTAGCGA' *.fastq > count/B2348_Fm0.txt
agrep -0 -c 'AACTGTTTCGCACCCAC' *.fastq > count/B2349_Fm0.txt
agrep -0 -c 'CTTGGTCACGAAGCCAG' *.fastq > count/B2350_Fm0.txt
agrep -0 -c 'AGCCTTTGTTTCGCGAT' *.fastq > count/B2351_Fm0.txt
agrep -0 -c 'GTGACAGTGAGTATATC' *.fastq > count/B2352_Fm0.txt
agrep -0 -c 'TACAGATTAGAGCGTGC' *.fastq > count/B2353_Fm0.txt
agrep -0 -c 'AGCAGTTATGGATCCCC' *.fastq > count/B2354_Fm0.txt
agrep -0 -c 'CGCCGTATGCAGATCGT' *.fastq > count/B2355_Fm0.txt
agrep -0 -c 'CGGGTGTTGGGACCTTT' *.fastq > count/B2356_Fm0.txt
agrep -0 -c 'ACCCTATCGCATGCATC' *.fastq > count/B2357_Fm0.txt
agrep -0 -c 'TGACTAACGCGGACATT' *.fastq > count/B2359_Fm0.txt
agrep -0 -c 'CCGCAAACAGTCCCAAA' *.fastq > count/B2360_Fm0.txt
agrep -0 -c 'TCGATGAGATAACGGTT' *.fastq > count/B2361_Fm0.txt
agrep -0 -c 'AGTAGCACGATCCATAC' *.fastq > count/B2362_Fm0.txt
agrep -0 -c 'GTTACCTAGGCGCCAGT' *.fastq > count/B2363_Fm0.txt
agrep -0 -c 'ACAATCCCTAACCGAAA' *.fastq > count/B2364_Fm0.txt
agrep -0 -c 'GATTGTGCGATGGACTC' *.fastq > count/B2366_Fm0.txt
agrep -0 -c 'TTATAGGAGTCGACATC' *.fastq > count/B2368_Fm0.txt
agrep -0 -c 'GGTCCCGTAAGCATTGC' *.fastq > count/B2369_Fm0.txt
agrep -0 -c 'AGTTTCGCGGAAGGCCA' *.fastq > count/B2370_Fm0.txt
agrep -0 -c 'TCAAGATCTGGGGGCCT' *.fastq > count/B2371_Fm0.txt
agrep -0 -c 'TGAAAAGCGTCACTGGG' *.fastq > count/B2372_Fm0.txt
agrep -0 -c 'CCTCGGAAGGAACAGAC' *.fastq > count/B2374_Fm0.txt
agrep -0 -c 'ACCGAAATCTTCTTGCC' *.fastq > count/B2375_Fm0.txt
agrep -0 -c 'AGTACACTTTATAGCGC' *.fastq > count/B2377_Fm0.txt
agrep -0 -c 'GAGAAGCATAGTGCGCT' *.fastq > count/B2378_Fm0.txt
agrep -0 -c 'CAGTTCACCGAGCGAAT' *.fastq > count/B2379_Fm0.txt
agrep -0 -c 'TCGGTCTCTGCAAAACC' *.fastq > count/B2380_Fm0.txt
agrep -0 -c 'CAGGGATCTAGTTCAGA' *.fastq > count/B2382_Fm0.txt
agrep -0 -c 'ACTTGAAAGGTTGGTAG' *.fastq > count/B2383_Fm0.txt
agrep -0 -c 'CCACCTTGTACAGCCGA' *.fastq > count/B2384_Fm0.txt
agrep -0 -c 'TGAGGGCACCAGTCTTA' *.fastq > count/B2385_Fm0.txt
agrep -0 -c 'TGAAATGCGGGGTAGAG' *.fastq > count/B2386_Fm0.txt
agrep -0 -c 'TGTTCTCAGGTGGAAGC' *.fastq > count/B2388_Fm0.txt
agrep -0 -c 'AGCGACCCTATACTCCA' *.fastq > count/B2389_Fm0.txt
agrep -0 -c 'AGGGCCCATTCTAGTCT' *.fastq > count/B2390_Fm0.txt
agrep -0 -c 'TGCCCTCCTACATATTT' *.fastq > count/B2392_Fm0.txt
agrep -0 -c 'CACTTTTCAACCATACC' *.fastq > count/B2393_Fm0.txt
agrep -0 -c 'GTGGAACGCATACATCG' *.fastq > count/B2395_Fm0.txt
agrep -0 -c 'ATGTAAGTCGCATCCCT' *.fastq > count/B2397_Fm0.txt
agrep -0 -c 'ACGCACCTTACGATCTC' *.fastq > count/B2398_Fm0.txt
agrep -0 -c 'TCTGGGTATATCGCCCT' *.fastq > count/B2399_Fm0.txt
agrep -0 -c 'GTATGGTAGACTGTGCT' *.fastq > count/B2400_Fm0.txt
agrep -0 -c 'GCACCTCGCGATATTAT' *.fastq > count/B2402_Fm0.txt
agrep -0 -c 'GGGGGTTTTTATGGCAA' *.fastq > count/B2403_Fm0.txt
agrep -0 -c 'CGCGAGCCTGACTAAAG' *.fastq > count/B2404_Fm0.txt
agrep -0 -c 'TCGATTTAGACAGCAAC' *.fastq > count/B2405_Fm0.txt
agrep -0 -c 'ATGGTAATGTAGGTAGG' *.fastq > count/B2406_Fm0.txt
agrep -0 -c 'CACCAGTGAATACCATT' *.fastq > count/B2407_Fm0.txt
agrep -0 -c 'GCGATTGATGCCCGCTA' *.fastq > count/B2408_Fm0.txt
agrep -0 -c 'TAGTTGGGAGTCTAAGT' *.fastq > count/B2410_Fm0.txt
agrep -0 -c 'GCCGTACGTAAAAGTAC' *.fastq > count/B2411_Fm0.txt
agrep -0 -c 'CAAGGGGACACAATTCC' *.fastq > count/B2412_Fm0.txt
agrep -0 -c 'ACGATCAAATATGCCCG' *.fastq > count/B2413_Fm0.txt
agrep -0 -c 'CTGGTGTCAATCGACAT' *.fastq > count/B2414_Fm0.txt
agrep -0 -c 'CTCGAACTTGCGTATAC' *.fastq > count/B2415_Fm0.txt
agrep -0 -c 'CCTAATTTTGTCAAGGG' *.fastq > count/B2416_Fm0.txt
agrep -0 -c 'GTAGCGACTTTATCCGT' *.fastq > count/B2417_Fm0.txt
agrep -0 -c 'AACTACGAGTCTTGCCC' *.fastq > count/B2418_Fm0.txt
agrep -0 -c 'TGCTTATGGCCTACGTA' *.fastq > count/B2420_Fm0.txt
agrep -0 -c 'TGTAAGAGTAATGCTCG' *.fastq > count/B2421_Fm0.txt
agrep -0 -c 'CAGTCCTATCTTCCAGC' *.fastq > count/B2422_Fm0.txt
agrep -0 -c 'GGCGCAGTATGACCATC' *.fastq > count/B2423_Fm0.txt
agrep -0 -c 'CGGGTCTTGGAGAACAG' *.fastq > count/B2424_Fm0.txt
agrep -0 -c 'TGATAAACGATCCCCGG' *.fastq > count/B2425_Fm0.txt
agrep -0 -c 'ATACCCAACTGATTACG' *.fastq > count/B2426_Fm0.txt
agrep -0 -c 'CGCACATAATAGCTACA' *.fastq > count/B2427_Fm0.txt
agrep -0 -c 'AAGCCCCGAGGTAAACG' *.fastq > count/B2428_Fm0.txt
agrep -0 -c 'GTTGACCCATCGTACTA' *.fastq > count/B2429_Fm0.txt
agrep -0 -c 'GTGACATAACGGATCCA' *.fastq > count/B2430_Fm0.txt
agrep -0 -c 'GTGTTTTCGAACTTGCG' *.fastq > count/B2431_Fm0.txt
agrep -0 -c 'CCAAATAGTGGGTGACG' *.fastq > count/B2432_Fm0.txt
agrep -0 -c 'CTCTTTCCCTGTCTATA' *.fastq > count/B2433_Fm0.txt
agrep -0 -c 'ACCCCGTTTCAGCAAAG' *.fastq > count/B2434_Fm0.txt
agrep -0 -c 'GTCATGAGGATGAGAGC' *.fastq > count/B2436_Fm0.txt
agrep -0 -c 'ATCATTAGTTCCGCCCT' *.fastq > count/B2437_Fm0.txt
agrep -0 -c 'AGTTTCGAAGCGTGCTG' *.fastq > count/B2438_Fm0.txt
agrep -0 -c 'GTCAGTCATCCCCTTTT' *.fastq > count/B2439_Fm0.txt
agrep -0 -c 'GGCGAACGTAGAATCCG' *.fastq > count/B2440_Fm0.txt
agrep -0 -c 'GACAGGCTAGGTCTATA' *.fastq > count/B2441_Fm0.txt
agrep -0 -c 'GACTCCGTTGCAATCCC' *.fastq > count/B2442_Fm0.txt
agrep -0 -c 'TAAGCCGCCTAGTTATA' *.fastq > count/B2443_Fm0.txt
agrep -0 -c 'TTCCACGCCTCTACAGA' *.fastq > count/B2445_Fm0.txt
agrep -0 -c 'CGTGGCTAGGAATATGT' *.fastq > count/B2446_Fm0.txt
agrep -0 -c 'CATAGGGTCGTGTCACA' *.fastq > count/B2447_Fm0.txt
agrep -0 -c 'TGTATAATCAGCGTGCA' *.fastq > count/B2448_Fm0.txt
agrep -0 -c 'CTAACTACTTCTGGAGG' *.fastq > count/B2449_Fm0.txt
agrep -0 -c 'GTACGTCTCGCTTGTCA' *.fastq > count/B2450_Fm0.txt
agrep -0 -c 'ATAATCATACTGGCTGG' *.fastq > count/B2452_Fm0.txt
agrep -0 -c 'TGCAGGGTGAAACTCCG' *.fastq > count/B2454_Fm0.txt
agrep -0 -c 'CAGGTTGAATATGGCTC' *.fastq > count/B2455_Fm0.txt
agrep -0 -c 'CCGTAGGGCATTTTGCA' *.fastq > count/B2456_Fm0.txt
agrep -0 -c 'CTTACCCCTGCTAGTTA' *.fastq > count/B2458_Fm0.txt
agrep -0 -c 'CAAACAACCGTATTGCC' *.fastq > count/B2460_Fm0.txt
agrep -0 -c 'TTAACCACACACATAGC' *.fastq > count/B2461_Fm0.txt
agrep -0 -c 'GGGTGACGGGTGTTTTT' *.fastq > count/B2462_Fm0.txt
agrep -0 -c 'TTACCCCGACTTCACAT' *.fastq > count/B2463_Fm0.txt
agrep -0 -c 'GCGGAACAGGAAAACTG' *.fastq > count/B2466_Fm0.txt
agrep -0 -c 'CTAAGAGAGCGCGTGGA' *.fastq > count/B2467_Fm0.txt
agrep -0 -c 'ATGGTGTTATGCTGAGA' *.fastq > count/B2469_Fm0.txt
agrep -0 -c 'ACTAGCCTCAGAGAGTG' *.fastq > count/B2470_Fm0.txt
agrep -0 -c 'GGCTTCTTGGTAAGCGA' *.fastq > count/B2471_Fm0.txt
agrep -0 -c 'TAAGCCTAGTCAGACTT' *.fastq > count/B2472_Fm0.txt
agrep -0 -c 'AGAACACGTGTCGCCCA' *.fastq > count/B2473_Fm0.txt
agrep -0 -c 'CTTGCCGCATAGCAGAA' *.fastq > count/B2474_Fm0.txt
agrep -0 -c 'TTCTACTGGGACTACCA' *.fastq > count/B2476_Fm0.txt
agrep -0 -c 'CGAGTTGGTGACAAGGT' *.fastq > count/B2477_Fm0.txt
agrep -0 -c 'TCTCCGAATTCTCCTAC' *.fastq > count/B2478_Fm0.txt
agrep -0 -c 'CGCCGCCTCAGTACTTT' *.fastq > count/B2479_Fm0.txt
agrep -0 -c 'GAGCACTCGGTGACGTT' *.fastq > count/B2480_Fm0.txt
agrep -0 -c 'GGTAGATTGACGGACGT' *.fastq > count/B2481_Fm0.txt
agrep -0 -c 'ATCGCGACTAGTTACTC' *.fastq > count/B2482_Fm0.txt
agrep -0 -c 'AACTACCTCCTCTGATG' *.fastq > count/B2483_Fm0.txt
agrep -0 -c 'ATCCGAGAGTTTAGGCT' *.fastq > count/B2484_Fm0.txt
agrep -0 -c 'CATCCGTAGTAGTCCAT' *.fastq > count/B2485_Fm0.txt
agrep -0 -c 'CACCCTTACATAGAGTT' *.fastq > count/B2486_Fm0.txt
agrep -0 -c 'CGCTACTTCTCGAGAAG' *.fastq > count/B2487_Fm0.txt
agrep -0 -c 'ACGTGCATAGGATTGCA' *.fastq > count/B2488_Fm0.txt
agrep -0 -c 'ATGGCCGAACTTTCTAG' *.fastq > count/B2489_Fm0.txt
agrep -0 -c 'AAGCTTTTGCGCTTGAC' *.fastq > count/B2490_Fm0.txt
agrep -0 -c 'ATCAGTAGCTTCAGCCC' *.fastq > count/B2491_Fm0.txt
agrep -0 -c 'ATATCTACATACGCGAG' *.fastq > count/B2492_Fm0.txt
agrep -0 -c 'TGCCACCCAGTATATTG' *.fastq > count/B2493_Fm0.txt
agrep -0 -c 'GGCTCGGGGGTTGATTT' *.fastq > count/B2494_Fm0.txt
agrep -0 -c 'CATCCTCAGCATAAGAC' *.fastq > count/B2495_Fm0.txt
agrep -0 -c 'GCGGAAGTTATGAGACT' *.fastq > count/B2496_Fm0.txt
agrep -0 -c 'TCTTTCGCATCTTCCTA' *.fastq > count/B2497_Fm0.txt
agrep -0 -c 'ATAAGAATCCGCCCACT' *.fastq > count/B2498_Fm0.txt
agrep -0 -c 'GTACCAGCAGCAGCTAG' *.fastq > count/B2500_Fm0.txt
agrep -0 -c 'GGAACCTCTGCGTCTTC' *.fastq > count/B2501_Fm0.txt
agrep -0 -c 'AGTACGGCCCTGGTCTT' *.fastq > count/B2502_Fm0.txt
agrep -0 -c 'GATGAGTATGGAAGTCT' *.fastq > count/B2503_Fm0.txt
agrep -0 -c 'ATGGATCAGTTGACTGC' *.fastq > count/B2504_Fm0.txt
agrep -0 -c 'TTATTGTGGGATGACGG' *.fastq > count/B2505_Fm0.txt
agrep -0 -c 'AGGATGGTGGCACGAAG' *.fastq > count/B2506_Fm0.txt
agrep -0 -c 'TTCGTACGGAGAGTTCC' *.fastq > count/B2507_Fm0.txt
agrep -0 -c 'ATGACTTAGTCATTGGC' *.fastq > count/B2508_Fm0.txt
agrep -0 -c 'AGAGAGCTTCGATAGTA' *.fastq > count/B2509_Fm0.txt
agrep -0 -c 'ATGGCCGTGACGACCTA' *.fastq > count/B2513_Fm0.txt
agrep -0 -c 'TAATCAGCGGTTGCAAA' *.fastq > count/B2514_Fm0.txt
agrep -0 -c 'GTAAGAGCTTGAGGATT' *.fastq > count/B2515_Fm0.txt
agrep -0 -c 'ACCATTGAAATGCTGGA' *.fastq > count/B2516_Fm0.txt
agrep -0 -c 'TTCTTAATCACACCGGT' *.fastq > count/B2517_Fm0.txt
agrep -0 -c 'AAGACTTAAAGGCACCC' *.fastq > count/B2518_Fm0.txt
agrep -0 -c 'ATTCCAGCCGGGAAAGA' *.fastq > count/B2519_Fm0.txt
agrep -0 -c 'TTCTTCACCCACAGTTG' *.fastq > count/B2520_Fm0.txt
agrep -0 -c 'AACTCGACCTGCCGACT' *.fastq > count/B2521_Fm0.txt
agrep -0 -c 'TCGGCTTCTCCGATCCA' *.fastq > count/B2522_Fm0.txt
agrep -0 -c 'TTAATCCAGCAGCGCCC' *.fastq > count/B2523_Fm0.txt
agrep -0 -c 'CCGGAAGACACGAATAC' *.fastq > count/B2524_Fm0.txt
agrep -0 -c 'AACGGTATACTGCGGTT' *.fastq > count/B2525_Fm0.txt
agrep -0 -c 'ACCGATGGACCAACACG' *.fastq > count/B2526_Fm0.txt
agrep -0 -c 'CGAATGCCCAAGAAGGC' *.fastq > count/B2528_Fm0.txt
agrep -0 -c 'TGGAATAACTCAAGCCT' *.fastq > count/B2529_Fm0.txt
agrep -0 -c 'TAACAACTGCAATCGGA' *.fastq > count/B2530_Fm0.txt
agrep -0 -c 'ATCGGGCAGCAGATGCA' *.fastq > count/B2531_Fm0.txt
agrep -0 -c 'CCGATATTTGACAGCGA' *.fastq > count/B2532_Fm0.txt
agrep -0 -c 'AGTCTGAATACGCTACA' *.fastq > count/B2533_Fm0.txt
agrep -0 -c 'GCACCTTTTAACGCCAA' *.fastq > count/B2534_Fm0.txt
agrep -0 -c 'GGTGCTTGTCTGGAGTA' *.fastq > count/B2535_Fm0.txt
agrep -0 -c 'TTGGCTTTGGGACGTAG' *.fastq > count/B2537_Fm0.txt
agrep -0 -c 'ACAACGGGTCGTAATCT' *.fastq > count/B2538_Fm0.txt
agrep -0 -c 'GTACGTTTTGGCGACGG' *.fastq > count/B2539_Fm0.txt
agrep -0 -c 'TATACAACGGGACTTCC' *.fastq > count/B2540_Fm0.txt
agrep -0 -c 'CGGCCCAATTCCCATTG' *.fastq > count/B2541_Fm0.txt
agrep -0 -c 'TAAACGAACTCGTTACC' *.fastq > count/B2543_Fm0.txt
agrep -0 -c 'GTTCCTTGAGAACGACG' *.fastq > count/B2544_Fm0.txt
agrep -0 -c 'ACATCCCTAAGTCGTCG' *.fastq > count/B2545_Fm0.txt
agrep -0 -c 'ATGACTCCCCGCAATCC' *.fastq > count/B2546_Fm0.txt
agrep -0 -c 'TGCTCCGATGGTGTTCG' *.fastq > count/B2547_Fm0.txt
agrep -0 -c 'TTAAGGTCGAGTTTACC' *.fastq > count/B2551_Fm0.txt
agrep -0 -c 'TAAGATATCACCGGCTG' *.fastq > count/B2552_Fm0.txt
agrep -0 -c 'CACTGCCGCGTCTAACT' *.fastq > count/B2553_Fm0.txt
agrep -0 -c 'GACCGACTAATATTGAC' *.fastq > count/B2554_Fm0.txt
agrep -0 -c 'CCTGAAACTTACCTGGT' *.fastq > count/B2555_Fm0.txt
agrep -0 -c 'GCTGGAGCCCTATTTGT' *.fastq > count/B2556_Fm0.txt
agrep -0 -c 'TGAGGCTCGAGCCACAA' *.fastq > count/B2557_Fm0.txt
agrep -0 -c 'GTGGGAGGTGTCCAATC' *.fastq > count/B2559_Fm0.txt
agrep -0 -c 'TGAGAAATACGTCTGCT' *.fastq > count/B2560_Fm0.txt
agrep -0 -c 'CGTAAACGAAAACGAGA' *.fastq > count/B2561_Fm0.txt
agrep -0 -c 'TCTTTCCCGTAATCTGT' *.fastq > count/B2562_Fm0.txt
agrep -0 -c 'AGAGCCGAATCATGGTC' *.fastq > count/B2563_Fm0.txt
agrep -0 -c 'GCCAGCTTCGGAGTAAT' *.fastq > count/B2564_Fm0.txt
agrep -0 -c 'TTAGCCTAGGTCGGTGC' *.fastq > count/B2565_Fm0.txt
agrep -0 -c 'GTGGTTCTGCGGTGGTA' *.fastq > count/B2566_Fm0.txt
agrep -0 -c 'ATATGCGCACCGTGATT' *.fastq > count/B2567_Fm0.txt
agrep -0 -c 'GGCTCGGCATGTCATAC' *.fastq > count/B2568_Fm0.txt
agrep -0 -c 'TAACCGACTAGGATGTC' *.fastq > count/B2569_Fm0.txt
agrep -0 -c 'CACGTACAGTGGTAACT' *.fastq > count/B2571_Fm0.txt
agrep -0 -c 'TGCGTTTCTGCTATTTG' *.fastq > count/B2572_Fm0.txt
agrep -0 -c 'CACATAATCCCGTTACG' *.fastq > count/B2573_Fm0.txt
agrep -0 -c 'CCATATCACCGGAACCC' *.fastq > count/B2574_Fm0.txt
agrep -0 -c 'TGCCTAGTGCAGACAAA' *.fastq > count/B2575_Fm0.txt
agrep -0 -c 'GTTCTAAACGAGGCGTT' *.fastq > count/B2576_Fm0.txt
agrep -0 -c 'AATCTCGCCTTGGCACG' *.fastq > count/B2577_Fm0.txt
agrep -0 -c 'TGATCGTAACTCCTGCT' *.fastq > count/B2580_Fm0.txt
agrep -0 -c 'CTTCGTTGCAATCGGAC' *.fastq > count/B2581_Fm0.txt
agrep -0 -c 'TACCTAGTCCACTCCTT' *.fastq > count/B2582_Fm0.txt
agrep -0 -c 'CCGAACTCGCGAATAAG' *.fastq > count/B2583_Fm0.txt
agrep -0 -c 'CAACCATCGACAGCGGT' *.fastq > count/B2584_Fm0.txt
agrep -0 -c 'CGAGACGTCGAGTCATA' *.fastq > count/B2585_Fm0.txt
agrep -0 -c 'AGTTCGCCAGCACTCGA' *.fastq > count/B2586_Fm0.txt
agrep -0 -c 'CGCGTGCAGAATCAAAA' *.fastq > count/B2587_Fm0.txt
agrep -0 -c 'ATACACGTTTAGAGACC' *.fastq > count/B2588_Fm0.txt
agrep -0 -c 'TTGATTCGTTGCATCAC' *.fastq > count/B2589_Fm0.txt
agrep -0 -c 'AATTCCGCCGATTTGCG' *.fastq > count/B2590_Fm0.txt
agrep -0 -c 'GCGCACCAGTTTTAAAG' *.fastq > count/B2593_Fm0.txt
agrep -0 -c 'CTACTAACTAAGGACTG' *.fastq > count/B2594_Fm0.txt
agrep -0 -c 'TGACACTGACATCCCAC' *.fastq > count/B2595_Fm0.txt
agrep -0 -c 'CAGACTCGAGAGTTGGT' *.fastq > count/B2596_Fm0.txt
agrep -0 -c 'CGATACAACAACGTGTG' *.fastq > count/B2597_Fm0.txt
agrep -0 -c 'GCATTGTGCTAGAGAAC' *.fastq > count/B2598_Fm0.txt
agrep -0 -c 'CCACCGGAGTTGCAAAG' *.fastq > count/B2599_Fm0.txt
agrep -0 -c 'TGTAGCAACTACATCGG' *.fastq > count/B2600_Fm0.txt
agrep -0 -c 'AAGTGAAGGCGTACCGA' *.fastq > count/B2601_Fm0.txt
agrep -0 -c 'CGATATTTATAGGGGGG' *.fastq > count/B2602_Fm0.txt
agrep -0 -c 'TGCACGAGCCTTCGTTG' *.fastq > count/B2603_Fm0.txt
agrep -0 -c 'GATTGATTACTCGCTGC' *.fastq > count/B2604_Fm0.txt
agrep -0 -c 'GTCAGGATCAAGCCTGG' *.fastq > count/B2605_Fm0.txt
agrep -0 -c 'GAGCGATTACGGACAGT' *.fastq > count/B2606_Fm0.txt
agrep -0 -c 'CAGATGTTTGAGGCATC' *.fastq > count/B2607_Fm0.txt
agrep -0 -c 'GCCTTAACGATCCACCA' *.fastq > count/B2609_Fm0.txt
agrep -0 -c 'CACGTTCGGAATGATCG' *.fastq > count/B2610_Fm0.txt
agrep -0 -c 'GCGTTACTAGTTAGACG' *.fastq > count/B2614_Fm0.txt
agrep -0 -c 'ACAGACTTGTAACCGGA' *.fastq > count/B2615_Fm0.txt
agrep -0 -c 'GGCTTTGCGGCCTAATG' *.fastq > count/B2616_Fm0.txt
agrep -0 -c 'TGATCGTCTGCTGTGCT' *.fastq > count/B2617_Fm0.txt
agrep -0 -c 'CTTCAATTAATTCCCGG' *.fastq > count/B2618_Fm0.txt
agrep -0 -c 'GTGAAGCCGTAGTTATG' *.fastq > count/B2619_Fm0.txt
agrep -0 -c 'GACTCTGCATAATGACC' *.fastq > count/B2620_Fm0.txt
agrep -0 -c 'CGTCTAGGATAAATCGG' *.fastq > count/B2621_Fm0.txt
agrep -0 -c 'ATAGATGCTGTCGTCGG' *.fastq > count/B2622_Fm0.txt
agrep -0 -c 'CTGACCGTTTCGCTAAA' *.fastq > count/B2623_Fm0.txt
agrep -0 -c 'GTACGGAGCGTATAGCG' *.fastq > count/B2624_Fm0.txt
agrep -0 -c 'TGTGAAGTTCGCGATAG' *.fastq > count/B2625_Fm0.txt
agrep -0 -c 'AAGGCGTCGACTTCTGC' *.fastq > count/B2626_Fm0.txt
agrep -0 -c 'CCGCCATAGAGTACGAG' *.fastq > count/B2627_Fm0.txt
agrep -0 -c 'CGTGCTCTTGATGCGAC' *.fastq > count/B2628_Fm0.txt
agrep -0 -c 'GCTATCTATTATCGCCC' *.fastq > count/B2629_Fm0.txt
agrep -0 -c 'GGCCGTTTCTGCTGGAT' *.fastq > count/B2630_Fm0.txt
agrep -0 -c 'TTGACTAACTGAGGACG' *.fastq > count/B2631_Fm0.txt
agrep -0 -c 'GGCCGGGTACTTCAAGT' *.fastq > count/B2633_Fm0.txt
agrep -0 -c 'AGCCGAGAAAGTTCACC' *.fastq > count/B2634_Fm0.txt
agrep -0 -c 'TCGCTGAATGGGGTTGG' *.fastq > count/B2635_Fm0.txt
agrep -0 -c 'CTCTTTTCTCCTACCGA' *.fastq > count/B2636_Fm0.txt
agrep -0 -c 'ATCTGATAAAGCCTCCC' *.fastq > count/B2637_Fm0.txt
agrep -0 -c 'CTGGGTTTTGCGCAATG' *.fastq > count/B2638_Fm0.txt
agrep -0 -c 'ACGAAAGACAGCGGATC' *.fastq > count/B2641_Fm0.txt
agrep -0 -c 'AGGCCAGTCAAACCCGA' *.fastq > count/B2642_Fm0.txt
agrep -0 -c 'GAGTACGATTCTTACCG' *.fastq > count/B2643_Fm0.txt
agrep -0 -c 'TTCAAATCGTGACACCC' *.fastq > count/B2644_Fm0.txt
agrep -0 -c 'TGCTGGTTGTCTTACCC' *.fastq > count/B2645_Fm0.txt
agrep -0 -c 'ACCTGCGCAAAAGATGA' *.fastq > count/B2646_Fm0.txt
agrep -0 -c 'CCGAGTGGTTAGATTAT' *.fastq > count/B2647_Fm0.txt
agrep -0 -c 'GATCGGATACTAACACT' *.fastq > count/B2648_Fm0.txt
agrep -0 -c 'GGACAGTATACTCAAGA' *.fastq > count/B2649_Fm0.txt
agrep -0 -c 'TAAGCGAAGTTCTTTCC' *.fastq > count/B2650_Fm0.txt
agrep -0 -c 'GCGCCTAAGAAATCTTG' *.fastq > count/B2651_Fm0.txt
agrep -0 -c 'TTGAGACTTTACCAAGG' *.fastq > count/B2652_Fm0.txt
agrep -0 -c 'GGTGACTATAGCCTCGG' *.fastq > count/B2653_Fm0.txt
agrep -0 -c 'AGCGAGACGTGCATATG' *.fastq > count/B2654_Fm0.txt
agrep -0 -c 'ATTAACGCGGCATTTCG' *.fastq > count/B2655_Fm0.txt
agrep -0 -c 'GGTCATAGCGGAAGAAA' *.fastq > count/B2656_Fm0.txt
agrep -0 -c 'AACTTAATCCACTGCGC' *.fastq > count/B2657_Fm0.txt
agrep -0 -c 'ACAACTGCCACTCAACG' *.fastq > count/B2658_Fm0.txt
agrep -0 -c 'GCACTCGTAATGGTAAG' *.fastq > count/B2659_Fm0.txt
agrep -0 -c 'CGTCTCTCGTCTTAAAC' *.fastq > count/B2660_Fm0.txt
agrep -0 -c 'AGAATCGTTGTCGCATC' *.fastq > count/B2661_Fm0.txt
agrep -0 -c 'GGACAGAGAGACTACCG' *.fastq > count/B2663_Fm0.txt
agrep -0 -c 'CGTTCCTAAGGACTTTC' *.fastq > count/B2664_Fm0.txt
agrep -0 -c 'AATGAGCGTCAGGTCCC' *.fastq > count/B2665_Fm0.txt
agrep -0 -c 'ACTTGGTCGTGTTTCGG' *.fastq > count/B2666_Fm0.txt
agrep -0 -c 'GGGAGTGTAAGGGTTGC' *.fastq > count/B2668_Fm0.txt
agrep -0 -c 'AGGTATTCCAAGGAGTA' *.fastq > count/B2670_Fm0.txt
agrep -0 -c 'TCGGATCGCTCACTTCA' *.fastq > count/B2671_Fm0.txt
agrep -0 -c 'ACGCACAGTCCCCTAGA' *.fastq > count/B2672_Fm0.txt
agrep -0 -c 'GGCACAAATGTGCGGGT' *.fastq > count/B2673_Fm0.txt
agrep -0 -c 'CGTGCCACTTCTAGCAT' *.fastq > count/B2674_Fm0.txt
agrep -0 -c 'CAAGTGCACGTGAGAGA' *.fastq > count/B2675_Fm0.txt
agrep -0 -c 'TGTAAAGGGTTGCCCCT' *.fastq > count/B2676_Fm0.txt
agrep -0 -c 'CGACACTTCTGGTAATA' *.fastq > count/B2677_Fm0.txt
agrep -0 -c 'TATTGCACGCCGGGCTA' *.fastq > count/B2679_Fm0.txt
agrep -0 -c 'GAGGCAAGGTAGTTTAC' *.fastq > count/B2680_Fm0.txt
agrep -0 -c 'CGACATGGAGCCGAAAC' *.fastq > count/B2681_Fm0.txt
agrep -0 -c 'TCAGAGAAATATCCGCG' *.fastq > count/B2682_Fm0.txt
agrep -0 -c 'AACCGTTCACTTGCAGG' *.fastq > count/B2683_Fm0.txt
agrep -0 -c 'TGTTGTTCTATCCGTCA' *.fastq > count/B2684_Fm0.txt
agrep -0 -c 'ACCGGTACTACTACCAG' *.fastq > count/B2685_Fm0.txt
agrep -0 -c 'AACTGTGTGATACGCCG' *.fastq > count/B2686_Fm0.txt
agrep -0 -c 'CGCTAGCACCGGTCTTA' *.fastq > count/B2688_Fm0.txt
agrep -0 -c 'CAACAGGTGTTAGCGGG' *.fastq > count/B2689_Fm0.txt
agrep -0 -c 'ACGCGTACTAGCGTCTG' *.fastq > count/B2691_Fm0.txt
agrep -0 -c 'ACAGCCCGGATACAGTC' *.fastq > count/B2692_Fm0.txt
agrep -0 -c 'GGATCCAGAATTAAGAC' *.fastq > count/B2693_Fm0.txt
agrep -0 -c 'CAGCAGTAAGTAGCAGG' *.fastq > count/B2694_Fm0.txt
agrep -0 -c 'ATTGAGCCGATTACTTC' *.fastq > count/B2696_Fm0.txt
agrep -0 -c 'TACATTGTAACCGTAGC' *.fastq > count/B2697_Fm0.txt
agrep -0 -c 'GGGGTTGCCATCCTATC' *.fastq > count/B2698_Fm0.txt
agrep -0 -c 'CAGCTTAACAGCACCTG' *.fastq > count/B2699_Fm0.txt
agrep -0 -c 'AAGAGTACATTCCGGGA' *.fastq > count/B2700_Fm0.txt
agrep -0 -c 'AATGAAGCGTACCTGAG' *.fastq > count/B2702_Fm0.txt
agrep -0 -c 'TAATTCGCAGCTTGTGT' *.fastq > count/B2703_Fm0.txt
agrep -0 -c 'TCACCTCTGATACAATG' *.fastq > count/B2704_Fm0.txt
agrep -0 -c 'AGCCAAGCGCCAGTTAT' *.fastq > count/B2705_Fm0.txt
agrep -0 -c 'TGAGACAGCAATATCAC' *.fastq > count/B2706_Fm0.txt
agrep -0 -c 'AAGCGCATTGTTGGCTG' *.fastq > count/B2707_Fm0.txt
agrep -0 -c 'AGTCGGTTCATAAAGTC' *.fastq > count/B2708_Fm0.txt
agrep -0 -c 'CGGTAGCGATGTATCTT' *.fastq > count/B2709_Fm0.txt
agrep -0 -c 'CCGAGACAGCGTAGGAT' *.fastq > count/B2710_Fm0.txt
agrep -0 -c 'CCGGGGAAATCGCCATA' *.fastq > count/B2712_Fm0.txt
agrep -0 -c 'GCGTGCATGCAACGTTA' *.fastq > count/B2713_Fm0.txt
agrep -0 -c 'TAGCATAGACGTGGCTC' *.fastq > count/B2714_Fm0.txt
agrep -0 -c 'TACTTCGGTAATACCGG' *.fastq > count/B2716_Fm0.txt
agrep -0 -c 'TATAGGGCGATAGTCCA' *.fastq > count/B2717_Fm0.txt
agrep -0 -c 'TGCGTGATGTGACGGAA' *.fastq > count/B2718_Fm0.txt
agrep -0 -c 'GCTCCGCCAATTTGAAA' *.fastq > count/B2719_Fm0.txt
agrep -0 -c 'AGGCGCGTTGATGGTTG' *.fastq > count/B2720_Fm0.txt
agrep -0 -c 'CACTAGGTTATAGTCCC' *.fastq > count/B2721_Fm0.txt
agrep -0 -c 'ACCTTACTTGCAATCCC' *.fastq > count/B2722_Fm0.txt
agrep -0 -c 'CGAACCCGTCTATGAAA' *.fastq > count/B2723_Fm0.txt
agrep -0 -c 'TATGATCGTCAAAGCCG' *.fastq > count/B2724_Fm0.txt
agrep -0 -c 'CGGGCTGAAATTGACCG' *.fastq > count/B2725_Fm0.txt
agrep -0 -c 'TATTAGCAGGGGACGTA' *.fastq > count/B2726_Fm0.txt
agrep -0 -c 'TTGACAGCAACTTAGCT' *.fastq > count/B2727_Fm0.txt
agrep -0 -c 'TAGGGTCCTGATAGTTT' *.fastq > count/B2728_Fm0.txt
agrep -0 -c 'ATGATCTGTCCTCCTAC' *.fastq > count/B2729_Fm0.txt
agrep -0 -c 'TCAGACACCTTCAAAGG' *.fastq > count/B2730_Fm0.txt
agrep -0 -c 'CTGTAATAAGCTGGTCC' *.fastq > count/B2731_Fm0.txt
agrep -0 -c 'GTTGACTTACGGCCAAC' *.fastq > count/B2732_Fm0.txt
agrep -0 -c 'GAGCGTATTCATCTTGT' *.fastq > count/B2734_Fm0.txt
agrep -0 -c 'GGGAGTTGCTACTGGTT' *.fastq > count/B2735_Fm0.txt
agrep -0 -c 'CAAGTCCCGGAGGCTAA' *.fastq > count/B2736_Fm0.txt
agrep -0 -c 'ATTATTGCAAGCAGGCC' *.fastq > count/B2737_Fm0.txt
agrep -0 -c 'AGGTTCCTGTTCGTGCT' *.fastq > count/B2740_Fm0.txt
agrep -0 -c 'CTCAACGTGAGTATCGA' *.fastq > count/B2741_Fm0.txt
agrep -0 -c 'CGTTCCAACTTAGATGA' *.fastq > count/B2743_Fm0.txt
agrep -0 -c 'AGCACATACCTGCGGAA' *.fastq > count/B2744_Fm0.txt
agrep -0 -c 'ACCAGAACATTAGTCAG' *.fastq > count/B2745_Fm0.txt
agrep -0 -c 'TGTCCCTGTCCAACCGT' *.fastq > count/B2746_Fm0.txt
agrep -0 -c 'TCTAGGTGCGGAATCCT' *.fastq > count/B2749_Fm0.txt
agrep -0 -c 'GTGCGATCTAACACCTC' *.fastq > count/B2750_Fm0.txt
agrep -0 -c 'CCGCATAACTTGGAGTC' *.fastq > count/B2751_Fm0.txt
agrep -0 -c 'GTTTGGCGCGCATAACT' *.fastq > count/B2752_Fm0.txt
agrep -0 -c 'GTCAAGCATGAATGCCG' *.fastq > count/B2753_Fm0.txt
agrep -0 -c 'TCGGCACCACTAGTCCT' *.fastq > count/B2754_Fm0.txt
agrep -0 -c 'TCACGGTTCTTAGTTAG' *.fastq > count/B2755_Fm0.txt
agrep -0 -c 'ATAGTCACCGGTCATGC' *.fastq > count/B2756_Fm0.txt
agrep -0 -c 'CCATTCGTCCCATGGAA' *.fastq > count/B2759_Fm0.txt
agrep -0 -c 'CGCAAATGTCGCATTGC' *.fastq > count/B2760_Fm0.txt
agrep -0 -c 'GAGTTCTCGACCTGGGT' *.fastq > count/B2761_Fm0.txt
agrep -0 -c 'TACTAATTCATGCCGAC' *.fastq > count/B2762_Fm0.txt
agrep -0 -c 'GTAGTTAGGCTCCACCA' *.fastq > count/B2763_Fm0.txt
agrep -0 -c 'TAGCCCGCAATAGCACT' *.fastq > count/B2764_Fm0.txt
agrep -0 -c 'TTCTAGAATGGATGACC' *.fastq > count/B2765_Fm0.txt
agrep -0 -c 'CTAAGCAAGTAGCATTG' *.fastq > count/B2766_Fm0.txt
agrep -0 -c 'AACGACGCCTGTTCCAC' *.fastq > count/B2767_Fm0.txt
agrep -0 -c 'TGCAGCATATTGCAGCG' *.fastq > count/B2768_Fm0.txt
agrep -0 -c 'TGATAGCCTACGATCCG' *.fastq > count/B2770_Fm0.txt
agrep -0 -c 'TCGTATAATCTTTGCGG' *.fastq > count/B2771_Fm0.txt
agrep -0 -c 'GTCATTGTTCGCGACCC' *.fastq > count/B2772_Fm0.txt
agrep -0 -c 'GCGTGTGGTAACTTGAC' *.fastq > count/B2774_Fm0.txt
agrep -0 -c 'AGTTCCGATGGTTCCGA' *.fastq > count/B2775_Fm0.txt
agrep -0 -c 'CTGACCGAACCGTAGAC' *.fastq > count/B2776_Fm0.txt
agrep -0 -c 'ATGCGTTCCCTCCTTCT' *.fastq > count/B2777_Fm0.txt
agrep -0 -c 'AGGTAAACCCCTCGTAG' *.fastq > count/B2778_Fm0.txt
agrep -0 -c 'CGCTCCTACAATTTTGC' *.fastq > count/B2781_Fm0.txt
agrep -0 -c 'ATTTGCGTTGAGTAGAC' *.fastq > count/B2782_Fm0.txt
agrep -0 -c 'ACCAGGTCTTAGTTGAA' *.fastq > count/B2783_Fm0.txt
agrep -0 -c 'CGAGCATCCCGGATGTT' *.fastq > count/B2784_Fm0.txt
agrep -0 -c 'GCAGTTGAAGGATGTTT' *.fastq > count/B2785_Fm0.txt
agrep -0 -c 'GGTCCATCCGAATGGTA' *.fastq > count/B2786_Fm0.txt
agrep -0 -c 'ACCGATCCCGTAGGTTT' *.fastq > count/B2788_Fm0.txt
agrep -0 -c 'CTGCTTAGCCGGCTCTA' *.fastq > count/B2789_Fm0.txt
agrep -0 -c 'CAACGCCGACTACTGCT' *.fastq > count/B2790_Fm0.txt
agrep -0 -c 'GTCCCATTCTAGAGATC' *.fastq > count/B2791_Fm0.txt
agrep -0 -c 'TTAGGAAGTCGGATACT' *.fastq > count/B2792_Fm0.txt
agrep -0 -c 'ATGGGGGTTCGATATCC' *.fastq > count/B2793_Fm0.txt
agrep -0 -c 'GCCCTGCTAATAACTTA' *.fastq > count/B2795_Fm0.txt
agrep -0 -c 'CACTGACTTGTCCGTGC' *.fastq > count/B2796_Fm0.txt
agrep -0 -c 'GATTTGATCCCAAGTTG' *.fastq > count/B2797_Fm0.txt
agrep -0 -c 'GTGGGTAGCGTGTCTGT' *.fastq > count/B2798_Fm0.txt
agrep -0 -c 'AGGCTCAATCGGAAGCT' *.fastq > count/B2799_Fm0.txt
agrep -0 -c 'CTGTTCGAGGTTTGACT' *.fastq > count/B2801_Fm0.txt
agrep -0 -c 'AGATGACTCGCAGGGTC' *.fastq > count/B2802_Fm0.txt
agrep -0 -c 'ACTTATCTTACCACCAG' *.fastq > count/B2803_Fm0.txt
agrep -0 -c 'CTCGGTTATACTCCAAG' *.fastq > count/B2804_Fm0.txt
agrep -0 -c 'CCGTGAACTGGCAGGAT' *.fastq > count/B2805_Fm0.txt
agrep -0 -c 'CGACCGTAACAGGTTGC' *.fastq > count/B2806_Fm0.txt
agrep -0 -c 'CGATTCACGAACACGTA' *.fastq > count/B2807_Fm0.txt
agrep -0 -c 'CGCGAAACCTTAGTATG' *.fastq > count/B2808_Fm0.txt
agrep -0 -c 'ATGCCCACGGATAAAAG' *.fastq > count/B2809_Fm0.txt
agrep -0 -c 'ACCTCAACAATTGTCCG' *.fastq > count/B2810_Fm0.txt
agrep -0 -c 'CCTCATCTTGATAGCAC' *.fastq > count/B2813_Fm0.txt
agrep -0 -c 'GCTATCGCTCATAGCTG' *.fastq > count/B2814_Fm0.txt
agrep -0 -c 'AGAGGCGCCATCTGAAC' *.fastq > count/B2815_Fm0.txt
agrep -0 -c 'CCGGTGCAAAACCGCAT' *.fastq > count/B2816_Fm0.txt
agrep -0 -c 'TAACTTGACCAACCGTA' *.fastq > count/B2818_Fm0.txt
agrep -0 -c 'GGGCCATCGAATCGTCT' *.fastq > count/B2819_Fm0.txt
agrep -0 -c 'TGCTGGACGTTATGGTG' *.fastq > count/B2820_Fm0.txt
agrep -0 -c 'ATACTACAAATGCAGCC' *.fastq > count/B2821_Fm0.txt
agrep -0 -c 'CTTGCAAGGACTCGTCG' *.fastq > count/B2822_Fm0.txt
agrep -0 -c 'GCGACAGAGTAAGGACA' *.fastq > count/B2823_Fm0.txt
agrep -0 -c 'AACGAGCATTACGTTGT' *.fastq > count/B2824_Fm0.txt
agrep -0 -c 'CACGTCAAGCTGCTGCT' *.fastq > count/B2825_Fm0.txt
agrep -0 -c 'GTAGCTTCTATCTCCTT' *.fastq > count/B2826_Fm0.txt
agrep -0 -c 'ATCGATTATGCTCTCGT' *.fastq > count/B2827_Fm0.txt
agrep -0 -c 'AGATCTAAGTGACTCGT' *.fastq > count/B2828_Fm0.txt
agrep -0 -c 'GATCAACGAGTAGCTAT' *.fastq > count/B2829_Fm0.txt
agrep -0 -c 'CAACTTGACAGAAGACA' *.fastq > count/B2831_Fm0.txt
agrep -0 -c 'TTCGATTCAGGCCTTTC' *.fastq > count/B2834_Fm0.txt
agrep -0 -c 'GCGAGACCTTGTAACTG' *.fastq > count/B2835_Fm0.txt
agrep -0 -c 'ATACATAACCATTGCGG' *.fastq > count/B2836_Fm0.txt
agrep -0 -c 'TGACTTATTCGTGGCAA' *.fastq > count/B2837_Fm0.txt
agrep -0 -c 'CTAGCCGCACAGAATAG' *.fastq > count/B2838_Fm0.txt
agrep -0 -c 'CCTGGATACACACGATT' *.fastq > count/B2839_Fm0.txt
agrep -0 -c 'AAGGTAGGTCCATCTGC' *.fastq > count/B2842_Fm0.txt
agrep -0 -c 'GTGGACATTCGATAACT' *.fastq > count/B2844_Fm0.txt
agrep -0 -c 'TCGCAGGTTTACGGGAA' *.fastq > count/B2845_Fm0.txt
agrep -0 -c 'ACGATTAGCCAGAGAGA' *.fastq > count/B2847_Fm0.txt
agrep -0 -c 'GAGTTTGATGGGCTAGG' *.fastq > count/B2848_Fm0.txt
agrep -0 -c 'TAGACTAGACTACACGG' *.fastq > count/B2850_Fm0.txt
agrep -0 -c 'ATCGGTCCACGAATACG' *.fastq > count/B2851_Fm0.txt
agrep -0 -c 'CCGCTGAAACACTAGAT' *.fastq > count/B2852_Fm0.txt
agrep -0 -c 'GGGTATTAGGTTGGGTT' *.fastq > count/B2853_Fm0.txt
agrep -0 -c 'ACGTTTGTCCCATTAAG' *.fastq > count/B2854_Fm0.txt
agrep -0 -c 'TGCCATTATGATCAGCC' *.fastq > count/B2855_Fm0.txt
agrep -0 -c 'CCGGTTGATCTTCCACA' *.fastq > count/B2857_Fm0.txt
agrep -0 -c 'AAGGTTGACCTTTTGCA' *.fastq > count/B2858_Fm0.txt
agrep -0 -c 'ACTGGTCGAGTCATCGG' *.fastq > count/B2859_Fm0.txt
agrep -0 -c 'TAACAGCGAGAGGAGTA' *.fastq > count/B2860_Fm0.txt
agrep -0 -c 'ACTCGTTGAATCCTGTC' *.fastq > count/B2861_Fm0.txt
agrep -0 -c 'ATCGCCCGAAATTAATC' *.fastq > count/B2862_Fm0.txt
agrep -0 -c 'CAGTTTACAACGTGCTG' *.fastq > count/B2863_Fm0.txt
agrep -0 -c 'ATCCACAAATATGCGCA' *.fastq > count/B2864_Fm0.txt
agrep -0 -c 'CTGCAAAACCCAGTTTT' *.fastq > count/B2865_Fm0.txt
agrep -0 -c 'CGAGTGTAACGCCAATA' *.fastq > count/B2866_Fm0.txt
agrep -0 -c 'ACATAAGGCAGGCGGTT' *.fastq > count/B2867_Fm0.txt
agrep -0 -c 'GAGGCATCCTTCTCGTA' *.fastq > count/B2869_Fm0.txt
agrep -0 -c 'CATCGGATATGGCTGGT' *.fastq > count/B2870_Fm0.txt
agrep -0 -c 'CACGTATGCCCTGGAAG' *.fastq > count/B2871_Fm0.txt
agrep -0 -c 'GTACACCACTAGTACGT' *.fastq > count/B2872_Fm0.txt
agrep -0 -c 'CGATGACGTCAGTGTGG' *.fastq > count/B2873_Fm0.txt
agrep -0 -c 'ACACTGGGGTTTGCGAC' *.fastq > count/B2874_Fm0.txt
agrep -0 -c 'AGGTGAACCTCACGTTC' *.fastq > count/B2875_Fm0.txt
agrep -0 -c 'TCACGAACTCTTGCGTA' *.fastq > count/B2876_Fm0.txt
agrep -0 -c 'CGTCTGAACGGCAGAAG' *.fastq > count/B2877_Fm0.txt
agrep -0 -c 'CGCGAATAACTTATGGG' *.fastq > count/B2878_Fm0.txt
agrep -0 -c 'CGGCAGAAGTCTACCTG' *.fastq > count/B2880_Fm0.txt
agrep -0 -c 'TACGTTACGCTCACCTT' *.fastq > count/B2881_Fm0.txt
agrep -0 -c 'TTCCATGGACTATCTTG' *.fastq > count/B2883_Fm0.txt
agrep -0 -c 'GGGAGGCTTGGTCGAAT' *.fastq > count/B2884_Fm0.txt
agrep -0 -c 'CCTACGACTGTCGGATC' *.fastq > count/B2885_Fm0.txt
agrep -0 -c 'CACGCAGTTTCAATAAC' *.fastq > count/B2886_Fm0.txt
agrep -0 -c 'ATAATCAACCGCACGGG' *.fastq > count/B2887_Fm0.txt
agrep -0 -c 'CAACCACCATAAATGAG' *.fastq > count/B2888_Fm0.txt
agrep -0 -c 'GTTCATGTCGATTCCGC' *.fastq > count/B2889_Fm0.txt
agrep -0 -c 'TCGTGGAACCAGACTGA' *.fastq > count/B2890_Fm0.txt
agrep -0 -c 'GTGCGTCCGATCATGCT' *.fastq > count/B2892_Fm0.txt
agrep -0 -c 'CATTATCTGAGACATGG' *.fastq > count/B2893_Fm0.txt
agrep -0 -c 'ATCCGTTATCGTAGGAT' *.fastq > count/B2894_Fm0.txt
agrep -0 -c 'AATTCGCAGGCGATACG' *.fastq > count/B2896_Fm0.txt
agrep -0 -c 'GTCTGTGCAGCATTGAC' *.fastq > count/B2897_Fm0.txt
agrep -0 -c 'TTGCGGCTACACTAGTC' *.fastq > count/B2898_Fm0.txt
agrep -0 -c 'ACCCCGTAGAATCGCAT' *.fastq > count/B2899_Fm0.txt
agrep -0 -c 'TCATTCCATCGCTTTCT' *.fastq > count/B2900_Fm0.txt
agrep -0 -c 'CCATTCAACAAGCTGCC' *.fastq > count/B2901_Fm0.txt
agrep -0 -c 'CCTCAGAACAACCCAGG' *.fastq > count/B2902_Fm0.txt
agrep -0 -c 'CCAGGCGCGTGTTATAC' *.fastq > count/B2903_Fm0.txt
agrep -0 -c 'TACAACTCGGCATCGGT' *.fastq > count/B2904_Fm0.txt
agrep -0 -c 'CACGCTGGATGGCTATT' *.fastq > count/B2905_Fm0.txt
agrep -0 -c 'TACGTGATTTTCATGCC' *.fastq > count/B2906_Fm0.txt
agrep -0 -c 'TATTTATGGTGGGCTTG' *.fastq > count/B2907_Fm0.txt
agrep -0 -c 'CGGAACTCCACGCAGAA' *.fastq > count/B2910_Fm0.txt
agrep -0 -c 'GCCGACTTACATTATGC' *.fastq > count/B2911_Fm0.txt
agrep -0 -c 'ACTACGGGTGGGTATCA' *.fastq > count/B2912_Fm0.txt
agrep -0 -c 'CTACCAGGTCGGTATCC' *.fastq > count/B2914_Fm0.txt
agrep -0 -c 'AGCTTTCGCTACTCAAA' *.fastq > count/B2915_Fm0.txt
agrep -0 -c 'CAACTTGCGCGTGACAA' *.fastq > count/B2917_Fm0.txt
agrep -0 -c 'CATATAGAGACGGGAGG' *.fastq > count/B2919_Fm0.txt
agrep -0 -c 'CAGCTTGCTGTCATAGA' *.fastq > count/B2920_Fm0.txt
agrep -0 -c 'ACGTACTGACTGCATAT' *.fastq > count/B2921_Fm0.txt
agrep -0 -c 'AATGGCAAAGGTGAGAC' *.fastq > count/B2922_Fm0.txt
agrep -0 -c 'ACCTCAATGTCAGGCCA' *.fastq > count/B2923_Fm0.txt
agrep -0 -c 'CAGCGCACAGCATGGTT' *.fastq > count/B2924_Fm0.txt
agrep -0 -c 'AACCAGGGCAGACTTAC' *.fastq > count/B2925_Fm0.txt
agrep -0 -c 'CATAGAGGCCCGGTGAA' *.fastq > count/B2926_Fm0.txt
agrep -0 -c 'CTTCAATATCACCGGGT' *.fastq > count/B2927_Fm0.txt
agrep -0 -c 'TAACAGCCACGGGCAAG' *.fastq > count/B2928_Fm0.txt
agrep -0 -c 'AGGTTAGCTACGGGTCA' *.fastq > count/B2930_Fm0.txt
agrep -0 -c 'CCGTCCAGATTTCGGTT' *.fastq > count/B2931_Fm0.txt
agrep -0 -c 'TCAGCTTCTAGGCGTTT' *.fastq > count/B2934_Fm0.txt
agrep -0 -c 'CCGGAGCCACTGTATGT' *.fastq > count/B2935_Fm0.txt
agrep -0 -c 'AGTTCTGTCCCTCCATA' *.fastq > count/B2936_Fm0.txt
agrep -0 -c 'CTCCAGCAGTGACATCG' *.fastq > count/B2937_Fm0.txt
agrep -0 -c 'AAGTTTCACTCGGTAAC' *.fastq > count/B2943_Fm0.txt
agrep -0 -c 'GTTAAGTCGGCTACATA' *.fastq > count/B2944_Fm0.txt
agrep -0 -c 'ACGGTTTCCAAGCTGCC' *.fastq > count/B2947_Fm0.txt
agrep -0 -c 'AACACCCCCATCCAAAG' *.fastq > count/B2948_Fm0.txt
agrep -0 -c 'CCACTAAGAGAATCCGC' *.fastq > count/B2949_Fm0.txt
agrep -0 -c 'TCAGCGTGCACCGATAC' *.fastq > count/B2950_Fm0.txt
agrep -0 -c 'GTACATTTATTCCCTGC' *.fastq > count/B2951_Fm0.txt
agrep -0 -c 'ACGCTTCACCCTACACG' *.fastq > count/B2952_Fm0.txt
agrep -0 -c 'GATGTCGCTCCTCCATA' *.fastq > count/B2954_Fm0.txt
agrep -0 -c 'GGTGTTCAGGTAATGGG' *.fastq > count/B2956_Fm0.txt
agrep -0 -c 'GTATGACTCTGGAGGTA' *.fastq > count/B2957_Fm0.txt
agrep -0 -c 'CGAGAGTACCCAAGACT' *.fastq > count/B2958_Fm0.txt
agrep -0 -c 'CACGAGACGATGTTTAC' *.fastq > count/B2959_Fm0.txt
agrep -0 -c 'TAATTGGAGAACCCAGA' *.fastq > count/B2960_Fm0.txt
agrep -0 -c 'CAAACCCAAGGATATAG' *.fastq > count/B2961_Fm0.txt
agrep -0 -c 'ACTCACGCACTCCATAA' *.fastq > count/B2963_Fm0.txt
agrep -0 -c 'ACCCCAAGTTTTCTACT' *.fastq > count/B2964_Fm0.txt
agrep -0 -c 'TCGGCCATGCGATACGA' *.fastq > count/B2965_Fm0.txt
agrep -0 -c 'CCATACGCGTCGGGATA' *.fastq > count/B2966_Fm0.txt
agrep -0 -c 'GAGTGGGCCAACTGATA' *.fastq > count/B2967_Fm0.txt
agrep -0 -c 'TATGGTATGCATGACAG' *.fastq > count/B2968_Fm0.txt
agrep -0 -c 'AGTAGCAGCTGGCACGA' *.fastq > count/B2969_Fm0.txt
agrep -0 -c 'GATGCACCAAATGTCAT' *.fastq > count/B2970_Fm0.txt
agrep -0 -c 'ATCGAACCTGAACAATC' *.fastq > count/B2971_Fm0.txt
agrep -0 -c 'AATCCTCACACTATACG' *.fastq > count/B2973_Fm0.txt
agrep -0 -c 'ACTGGTCCGGTATCTAG' *.fastq > count/B2974_Fm0.txt
agrep -0 -c 'GCAAGCTCCTGCCACAT' *.fastq > count/B2975_Fm0.txt
agrep -0 -c 'GAGCGAGTCGACGATGA' *.fastq > count/B2976_Fm0.txt
agrep -0 -c 'TTGTATAGTGGTAGACG' *.fastq > count/B2978_Fm0.txt
agrep -0 -c 'GCGTGCTATTCACGTTG' *.fastq > count/B2979_Fm0.txt
agrep -0 -c 'AACCCGTTGTGAAAAGA' *.fastq > count/B2980_Fm0.txt
agrep -0 -c 'TTAGCATCTAAGCGGAG' *.fastq > count/B2981_Fm0.txt
agrep -0 -c 'TAACCGCTGAGAAGAGT' *.fastq > count/B2982_Fm0.txt
agrep -0 -c 'CCGTAGGAACTAACGTT' *.fastq > count/B2983_Fm0.txt
agrep -0 -c 'AATACCTATGCGTTCGC' *.fastq > count/B2985_Fm0.txt
agrep -0 -c 'TAGGTCTGGCCCTAGTT' *.fastq > count/B2986_Fm0.txt
agrep -0 -c 'GCGGATTTCGGATATTT' *.fastq > count/B2987_Fm0.txt
agrep -0 -c 'ATATCGAACACGTCGAA' *.fastq > count/B2989_Fm0.txt
agrep -0 -c 'CTTGGTATTGACGACCG' *.fastq > count/B2990_Fm0.txt
agrep -0 -c 'TTCCCGAGAAGGTGATT' *.fastq > count/B2991_Fm0.txt
agrep -0 -c 'TATGCTGAATGGCACGA' *.fastq > count/B2992_Fm0.txt
agrep -0 -c 'CTAAAATTGACTCACGG' *.fastq > count/B2993_Fm0.txt
agrep -0 -c 'TGCGCCTACTGGATACG' *.fastq > count/B2994_Fm0.txt
agrep -0 -c 'TTCACGTTGTGCGGAAG' *.fastq > count/B2995_Fm0.txt
agrep -0 -c 'ATATCGTCAGAAGGGAA' *.fastq > count/B2996_Fm0.txt
agrep -0 -c 'GAGGAATCCCTGCAGTC' *.fastq > count/B2998_Fm0.txt
agrep -0 -c 'GGTTTTCTTGCGCACTA' *.fastq > count/B2999_Fm0.txt
agrep -0 -c 'ATAGGATAGATGACGAC' *.fastq > count/B3001_Fm0.txt
agrep -0 -c 'GCTATTGAAGAAACGCA' *.fastq > count/B3002_Fm0.txt
agrep -0 -c 'GCAGCTTCACTAGGTTT' *.fastq > count/B3003_Fm0.txt
agrep -0 -c 'GTACGACAGTTGCGACG' *.fastq > count/B3006_Fm0.txt
agrep -0 -c 'CTAAATGTGCCTTCTTC' *.fastq > count/B3007_Fm0.txt
agrep -0 -c 'GACGCTGTCTTGAGAGA' *.fastq > count/B3008_Fm0.txt
agrep -0 -c 'CAGTGTAACTCTCACGG' *.fastq > count/B3009_Fm0.txt
agrep -0 -c 'CGGCATGGGTTCTATCT' *.fastq > count/B3010_Fm0.txt
agrep -0 -c 'CTGATACCAGTCTGGTT' *.fastq > count/B3011_Fm0.txt
agrep -0 -c 'TACGCGGGATCGCTCAT' *.fastq > count/B3012_Fm0.txt
agrep -0 -c 'GTATCGCAGCAAACTTG' *.fastq > count/B3013_Fm0.txt
agrep -0 -c 'CCTTAAACAACGTCGGA' *.fastq > count/B3014_Fm0.txt
agrep -0 -c 'ATAGGAACCTCGTCTTT' *.fastq > count/B3015_Fm0.txt
agrep -0 -c 'CGTGATCAGCCCCTTAC' *.fastq > count/B3016_Fm0.txt
agrep -0 -c 'AGCGGGTAGCCACATCA' *.fastq > count/B3018_Fm0.txt
agrep -0 -c 'TGTAGGGGATCTAAGTG' *.fastq > count/B3019_Fm0.txt
agrep -0 -c 'TGATTCCGCTTCAAAAG' *.fastq > count/B3021_Fm0.txt
agrep -0 -c 'AATCTTAGAGCGTGGTC' *.fastq > count/B3022_Fm0.txt
agrep -0 -c 'TCACTCGTATACACCGG' *.fastq > count/B3023_Fm0.txt
agrep -0 -c 'ACGTAAACGCTCATGAC' *.fastq > count/B3024_Fm0.txt
agrep -0 -c 'TATTTCGACCGTTCACA' *.fastq > count/B3025_Fm0.txt
agrep -0 -c 'ACACACCCGTTGATACC' *.fastq > count/B3026_Fm0.txt
agrep -0 -c 'GCCGCCAGATCGATGTA' *.fastq > count/B3027_Fm0.txt
agrep -0 -c 'TCTTGCGTCTCCACAAG' *.fastq > count/B3028_Fm0.txt
agrep -0 -c 'CAGATACTAAAACCGTC' *.fastq > count/B3029_Fm0.txt
agrep -0 -c 'CGCAAGACCGAAAGCAA' *.fastq > count/B3030_Fm0.txt
agrep -0 -c 'GCGTAAACGGTTAACGG' *.fastq > count/B3031_Fm0.txt
agrep -0 -c 'CATCGCGTACACCGTGT' *.fastq > count/B3032_Fm0.txt
agrep -0 -c 'GTCTCTAGGTTACACGG' *.fastq > count/B3033_Fm0.txt
agrep -0 -c 'AGATGATGACCGAGTCT' *.fastq > count/B3034_Fm0.txt
agrep -0 -c 'TAAGGATTAGTGGGGGA' *.fastq > count/B3035_Fm0.txt
agrep -0 -c 'GCCGTTAGGGTCATAGT' *.fastq > count/B3037_Fm0.txt
agrep -0 -c 'ACTTCCGCATCACTATC' *.fastq > count/B3038_Fm0.txt
agrep -0 -c 'TATGACCCGTGAGGTTC' *.fastq > count/B3039_Fm0.txt
agrep -0 -c 'CGAACCTCAAGGCTCTC' *.fastq > count/B3041_Fm0.txt
agrep -0 -c 'CTAGCGCTCAAGGCCAA' *.fastq > count/B3042_Fm0.txt
agrep -0 -c 'CGCAATCACCCTCTCAC' *.fastq > count/B3043_Fm0.txt
agrep -0 -c 'AACATTTCAGCGACACC' *.fastq > count/B3044_Fm0.txt
agrep -0 -c 'ACCTCACGTCGAACTTC' *.fastq > count/B3046_Fm0.txt
agrep -0 -c 'GATTTCCCTCCACTTGT' *.fastq > count/B3047_Fm0.txt
agrep -0 -c 'TACATCATCCGACCTTC' *.fastq > count/B3048_Fm0.txt
agrep -0 -c 'CTGTGGGCGAGACTAAT' *.fastq > count/B3049_Fm0.txt
agrep -0 -c 'TTCGGTTGGGGTGCATG' *.fastq > count/B3051_Fm0.txt
agrep -0 -c 'GTCGGCATTTGCGTTTT' *.fastq > count/B3052_Fm0.txt
agrep -0 -c 'ATTTCCGTGCAACAAGC' *.fastq > count/B3053_Fm0.txt
agrep -0 -c 'CTGGCTCATCTAAGCCG' *.fastq > count/B3054_Fm0.txt
agrep -0 -c 'TAGAACTGCCCGATCCT' *.fastq > count/B3055_Fm0.txt
agrep -0 -c 'GGTGGCGAAATCTCTGA' *.fastq > count/B3056_Fm0.txt
agrep -0 -c 'GTCTCTATGTGTTCGCC' *.fastq > count/B3057_Fm0.txt
agrep -0 -c 'ACGATAGATCCCCGTGT' *.fastq > count/B3059_Fm0.txt
agrep -0 -c 'ACACCTTTACCGGAATA' *.fastq > count/B3060_Fm0.txt
agrep -0 -c 'AGCAGATTCTTGTCGAG' *.fastq > count/B3061_Fm0.txt
agrep -0 -c 'CTGTACTTAGTACAGGG' *.fastq > count/B3062_Fm0.txt
agrep -0 -c 'TTCTAAAGGGGGGCGAT' *.fastq > count/B3063_Fm0.txt
agrep -0 -c 'ACTAACACGGAGAGCCA' *.fastq > count/B3064_Fm0.txt
agrep -0 -c 'GGCCTATCCACGCAATC' *.fastq > count/B3067_Fm0.txt
agrep -0 -c 'CGCAAGTAACCAGGTCC' *.fastq > count/B3068_Fm0.txt
agrep -0 -c 'AATACTCTGAATGCGAG' *.fastq > count/B3069_Fm0.txt
agrep -0 -c 'TTCGCGCGATACATAAT' *.fastq > count/B3070_Fm0.txt
agrep -0 -c 'CAGAGGCTCACGAGCAA' *.fastq > count/B3071_Fm0.txt
agrep -0 -c 'ATAAGGCATAGAGAGTG' *.fastq > count/B3072_Fm0.txt
agrep -0 -c 'AATACAGGTTGACTGGT' *.fastq > count/B3073_Fm0.txt
agrep -0 -c 'TCTAACCATCGTTCCCG' *.fastq > count/B3074_Fm0.txt
agrep -0 -c 'TACGCCGGGAACATTCT' *.fastq > count/B3076_Fm0.txt
agrep -0 -c 'GCGTGCTATGCGATCTT' *.fastq > count/B3077_Fm0.txt
agrep -0 -c 'AGGAGGATTCCTACAGG' *.fastq > count/B3078_Fm0.txt
agrep -0 -c 'CCGGGGTGGAAACTTTT' *.fastq > count/B3079_Fm0.txt
agrep -0 -c 'GTAGTCGGAGTGCATTT' *.fastq > count/B3080_Fm0.txt
agrep -0 -c 'ACCTGCCCAGTTACTTT' *.fastq > count/B3083_Fm0.txt
agrep -0 -c 'GCCAACCATTTAACAGC' *.fastq > count/B3084_Fm0.txt
agrep -0 -c 'CCGATCCTGGATTCTTG' *.fastq > count/B3085_Fm0.txt
agrep -0 -c 'TCTCTGTTCGCACGTAG' *.fastq > count/B3086_Fm0.txt
agrep -0 -c 'GGTTGAAGCGAAGTTAT' *.fastq > count/B3087_Fm0.txt
agrep -0 -c 'ACTGGGGAAACCGTAGG' *.fastq > count/B3088_Fm0.txt
agrep -0 -c 'AGTCCCGTGGAGATTTG' *.fastq > count/B3089_Fm0.txt
agrep -0 -c 'GGTCCCCAACAAGAATA' *.fastq > count/B3090_Fm0.txt
agrep -0 -c 'GTCCTCTTTGTCGAACA' *.fastq > count/B3091_Fm0.txt
agrep -0 -c 'ACTTGCCAGCGCTTGAG' *.fastq > count/B3092_Fm0.txt
agrep -0 -c 'GTGCTACCTAACTACCC' *.fastq > count/B3094_Fm0.txt
agrep -0 -c 'CAGGTTGGTCATCTCCG' *.fastq > count/B3095_Fm0.txt
agrep -0 -c 'TTCGAGGGTATCAAAAG' *.fastq > count/B3096_Fm0.txt
agrep -0 -c 'TGCATCAGCCGTGACTG' *.fastq > count/B3097_Fm0.txt
agrep -0 -c 'CTATTACCCGTGTCCTT' *.fastq > count/B3098_Fm0.txt
agrep -0 -c 'AATGGGTTACCGTGGTT' *.fastq > count/B3099_Fm0.txt
agrep -0 -c 'CGTTACCTAATGTACTC' *.fastq > count/B3100_Fm0.txt
agrep -0 -c 'AGAGTTACGTGACACCA' *.fastq > count/B3101_Fm0.txt
agrep -0 -c 'TTACATCGCTGCAGGAG' *.fastq > count/B3102_Fm0.txt
agrep -0 -c 'GGTCGATGCGCGAATGA' *.fastq > count/B3103_Fm0.txt
agrep -0 -c 'GTGCTGTGAATCGAGAA' *.fastq > count/B3104_Fm0.txt
agrep -0 -c 'GCAACCAGAATTCGCGG' *.fastq > count/B3105_Fm0.txt
agrep -0 -c 'TAGAAAGCTCTGCGCTA' *.fastq > count/B3106_Fm0.txt
agrep -0 -c 'CAATACCGTCAAAATGG' *.fastq > count/B3107_Fm0.txt
agrep -0 -c 'TCTTATGCCCGTCCCAG' *.fastq > count/B3108_Fm0.txt
agrep -0 -c 'GTCTCGCGTGGGATAAT' *.fastq > count/B3109_Fm0.txt
agrep -0 -c 'TCATGCGGTGTCGGTAG' *.fastq > count/B3110_Fm0.txt
agrep -0 -c 'GGCACAGCTTTGTGCCA' *.fastq > count/B3112_Fm0.txt
agrep -0 -c 'CGTTCACTTAGGCACAG' *.fastq > count/B3113_Fm0.txt
agrep -0 -c 'ACCGGACTTGCAGTTGC' *.fastq > count/B3114_Fm0.txt
agrep -0 -c 'TAGTCCTTAAGCGTGTT' *.fastq > count/B3115_Fm0.txt
agrep -0 -c 'AGGGCGAAATGCCAGAT' *.fastq > count/B3116_Fm0.txt
agrep -0 -c 'GTCATCCTTCTATTGGC' *.fastq > count/B3117_Fm0.txt
agrep -0 -c 'TCAGGACTTGTAATCCG' *.fastq > count/B3118_Fm0.txt
agrep -0 -c 'TGATAGCGGCGTTAAGT' *.fastq > count/B3119_Fm0.txt
agrep -0 -c 'GGATTCCGGTACACGTG' *.fastq > count/B3120_Fm0.txt
agrep -0 -c 'ACCGCAGCCTCGGAAAA' *.fastq > count/B3121_Fm0.txt
agrep -0 -c 'ACTCGAACACGCACCGA' *.fastq > count/B3122_Fm0.txt
agrep -0 -c 'CTACCTATAGAAGGAAC' *.fastq > count/B3123_Fm0.txt
agrep -0 -c 'GCACATAGTTCAAGTGC' *.fastq > count/B3124_Fm0.txt
agrep -0 -c 'AGTCCGGTCTTTCTAAC' *.fastq > count/B3125_Fm0.txt
agrep -0 -c 'ACCTACTTCTGCGGAGA' *.fastq > count/B3126_Fm0.txt
agrep -0 -c 'AACCGCAGGATGTAAAG' *.fastq > count/B3127_Fm0.txt
agrep -0 -c 'ATGGTCGTAGGAGTGGA' *.fastq > count/B3128_Fm0.txt
agrep -0 -c 'AGGTCTGTACCCGTGGT' *.fastq > count/B3129_Fm0.txt
agrep -0 -c 'GCACTTCAGCTCTATCG' *.fastq > count/B3130_Fm0.txt
agrep -0 -c 'CACCCCACTGAGTATTG' *.fastq > count/B3131_Fm0.txt
agrep -0 -c 'CGGAACGTACTACTCGT' *.fastq > count/B3132_Fm0.txt
agrep -0 -c 'GTTCGGGTCATTAGGAA' *.fastq > count/B3133_Fm0.txt
agrep -0 -c 'ACCCCTCAAGACAGCGT' *.fastq > count/B3134_Fm0.txt
agrep -0 -c 'GTCCGTGCGGATTTAGT' *.fastq > count/B3135_Fm0.txt
agrep -0 -c 'AGCAGTACAATAGGATC' *.fastq > count/B3137_Fm0.txt
agrep -0 -c 'CGAAGGGCCAATCAATC' *.fastq > count/B3138_Fm0.txt
agrep -0 -c 'TTGCCCTAGCGAATACG' *.fastq > count/B3139_Fm0.txt
agrep -0 -c 'GCCACCCTATAATATGA' *.fastq > count/B3140_Fm0.txt
agrep -0 -c 'CCATACCATCTGTACTA' *.fastq > count/B3141_Fm0.txt
agrep -0 -c 'CCAGACCAGCATCGAAG' *.fastq > count/B3142_Fm0.txt
agrep -0 -c 'CACGATGTCCGTACTGT' *.fastq > count/B3145_Fm0.txt
agrep -0 -c 'TGTGGTATGTCCCCGAT' *.fastq > count/B3146_Fm0.txt
agrep -0 -c 'AGAGCCAAATGAGATTC' *.fastq > count/B3147_Fm0.txt
agrep -0 -c 'AGAATGTCGCTGTAGAG' *.fastq > count/B3148_Fm0.txt
agrep -0 -c 'GACTTCGATACACGGCC' *.fastq > count/B3149_Fm0.txt
agrep -0 -c 'CAATGGTCGTAGCAGTT' *.fastq > count/B3152_Fm0.txt
agrep -0 -c 'ATCCACGGTGCTTTAGA' *.fastq > count/B3153_Fm0.txt
agrep -0 -c 'CGAGTCATGTAGTGTTG' *.fastq > count/B3155_Fm0.txt
agrep -0 -c 'CTCTTGTAATCCCACGA' *.fastq > count/B3156_Fm0.txt
agrep -0 -c 'GCTAGCTACAACGTAAG' *.fastq > count/B3157_Fm0.txt
agrep -0 -c 'GGTAACAACGGCACTAC' *.fastq > count/B3158_Fm0.txt
agrep -0 -c 'ATAGCACCTCGTTAGCG' *.fastq > count/B3159_Fm0.txt
agrep -0 -c 'CGATTAAGGGAGGGCTA' *.fastq > count/B3161_Fm0.txt
agrep -0 -c 'AGGCGTCTACCTTACTA' *.fastq > count/B3163_Fm0.txt
agrep -0 -c 'CTAGTACCTCGACCCTA' *.fastq > count/B3164_Fm0.txt
agrep -0 -c 'TGCCCTAAAAGTCCCCG' *.fastq > count/B3165_Fm0.txt
agrep -0 -c 'ACGAAGCGAGTTGTCAG' *.fastq > count/B3166_Fm0.txt
agrep -0 -c 'GTACCCTTGTTTCGTTT' *.fastq > count/B3167_Fm0.txt
agrep -0 -c 'GACTGGATGAGTATGGT' *.fastq > count/B3168_Fm0.txt
agrep -0 -c 'CCGGATGTCTGCGAATT' *.fastq > count/B3169_Fm0.txt
agrep -0 -c 'TGGATACTTCGGAAGAC' *.fastq > count/B3170_Fm0.txt
agrep -0 -c 'CGTTCGTTATGGCAACC' *.fastq > count/B3171_Fm0.txt
agrep -0 -c 'CTGATACGGGATAGTTA' *.fastq > count/B3174_Fm0.txt
agrep -0 -c 'TCTCCGATAATTCGGAT' *.fastq > count/B3175_Fm0.txt
agrep -0 -c 'TCTTCCGATCCCAAATG' *.fastq > count/B3176_Fm0.txt
agrep -0 -c 'CGATTGGAATGCGTGAC' *.fastq > count/B3178_Fm0.txt
agrep -0 -c 'ACGGGGAGGGTTTAACC' *.fastq > count/B3181_Fm0.txt
agrep -0 -c 'ACTTAATGTTACCGAGC' *.fastq > count/B3182_Fm0.txt
agrep -0 -c 'TTCCCGCTTGTTCGTGG' *.fastq > count/B3183_Fm0.txt
agrep -0 -c 'CCTATAAAGCCTGTTCC' *.fastq > count/B3184_Fm0.txt
agrep -0 -c 'CTATTTGTCCTGTACGG' *.fastq > count/B3185_Fm0.txt
agrep -0 -c 'CAGGTATCCGCCAGTGA' *.fastq > count/B3186_Fm0.txt
agrep -0 -c 'GAGCGGAAGCAATTACC' *.fastq > count/B3187_Fm0.txt
agrep -0 -c 'GCGATAATGGCACCGTG' *.fastq > count/B3188_Fm0.txt
agrep -0 -c 'GCGTTATCCCTGCGCTA' *.fastq > count/B3189_Fm0.txt
agrep -0 -c 'ACGCGTAAGAACCAGGG' *.fastq > count/B3190_Fm0.txt
agrep -0 -c 'TTACGTGTACCAATGAC' *.fastq > count/B3191_Fm0.txt
agrep -0 -c 'AGACCTCCTATACTCGT' *.fastq > count/B3192_Fm0.txt
agrep -0 -c 'GACTGCGTCTAAATTGC' *.fastq > count/B3193_Fm0.txt
agrep -0 -c 'AAGGACGTATACACCCA' *.fastq > count/B3195_Fm0.txt
agrep -0 -c 'GGAATCCAGGGTTAAAA' *.fastq > count/B3197_Fm0.txt
agrep -0 -c 'AACCAGGGTATACTTCC' *.fastq > count/B3198_Fm0.txt
agrep -0 -c 'TGTGAACGCCAAGACAT' *.fastq > count/B3199_Fm0.txt
agrep -0 -c 'CCTAAGGCGAACGCTTA' *.fastq > count/B3202_Fm0.txt
agrep -0 -c 'GGACCGGTCGCAAATCT' *.fastq > count/B3203_Fm0.txt
agrep -0 -c 'CCTGCGAAGTGCATAAT' *.fastq > count/B3204_Fm0.txt
agrep -0 -c 'TGCCCCCGTACGATCTA' *.fastq > count/B3205_Fm0.txt
agrep -0 -c 'CTAGCTCGTAACAGGCA' *.fastq > count/B3206_Fm0.txt
agrep -0 -c 'CCGTTTGGATTTTCGTA' *.fastq > count/B3208_Fm0.txt
agrep -0 -c 'TGACGTTGTTCCTCCAT' *.fastq > count/B3210_Fm0.txt
agrep -0 -c 'TCTGTGACCGAAACGGC' *.fastq > count/B3211_Fm0.txt
agrep -0 -c 'GGGCACTCGGGATGTTA' *.fastq > count/B3212_Fm0.txt
agrep -0 -c 'CGATCAGCCATAATCCT' *.fastq > count/B3213_Fm0.txt
agrep -0 -c 'GTAGCTTCCCGTATAGA' *.fastq > count/B3215_Fm0.txt
agrep -0 -c 'ACGGTGTCTGGCATTTA' *.fastq > count/B3216_Fm0.txt
agrep -0 -c 'CGACGAGCCAGTGAATT' *.fastq > count/B3217_Fm0.txt
agrep -0 -c 'GTGATGGCTAAGCAATG' *.fastq > count/B3218_Fm0.txt
agrep -0 -c 'GCAGTGCTATCTCTTGA' *.fastq > count/B3220_Fm0.txt
agrep -0 -c 'CAAATCTCATCCGATGC' *.fastq > count/B3221_Fm0.txt
agrep -0 -c 'AATTGGCTCGCATGCTC' *.fastq > count/B3222_Fm0.txt
agrep -0 -c 'CGCTATCGACGCATAAC' *.fastq > count/B3223_Fm0.txt
agrep -0 -c 'AATGATTTTCGCCCCAC' *.fastq > count/B3224_Fm0.txt
agrep -0 -c 'TAACAGGTAGTAACGTC' *.fastq > count/B3226_Fm0.txt
agrep -0 -c 'CGGCCCGAGCTATACTT' *.fastq > count/B3227_Fm0.txt
agrep -0 -c 'AATGCGCAAGTGGGTTC' *.fastq > count/B3228_Fm0.txt
agrep -0 -c 'TGCTCTCGGGGTAGTCA' *.fastq > count/B3229_Fm0.txt
agrep -0 -c 'TGCTGCACACCATTAGA' *.fastq > count/B3230_Fm0.txt
agrep -0 -c 'CATAGGGAATGCTCCGC' *.fastq > count/B3231_Fm0.txt
agrep -0 -c 'ATATCGCTTGCATGGGT' *.fastq > count/B3233_Fm0.txt
agrep -0 -c 'TGCGCATGAACCAGTGC' *.fastq > count/B3234_Fm0.txt
agrep -0 -c 'CATTCGCGCGGAAAATC' *.fastq > count/B3235_Fm0.txt
agrep -0 -c 'GCGCGAATGATGTAGAA' *.fastq > count/B3236_Fm0.txt
agrep -0 -c 'TCAGTTCTGCGAACAAT' *.fastq > count/B3238_Fm0.txt
agrep -0 -c 'ATAAAGGCTGTTCCCCT' *.fastq > count/B3239_Fm0.txt
agrep -0 -c 'TGTGAATCAGCAGGGGA' *.fastq > count/B3240_Fm0.txt
agrep -0 -c 'ATCTGGCTTACTCCCTC' *.fastq > count/B3241_Fm0.txt
agrep -0 -c 'ACCGATAGTTCACCGTA' *.fastq > count/B3242_Fm0.txt
agrep -0 -c 'AGGTCCTAGTGCTTAAG' *.fastq > count/B3244_Fm0.txt
agrep -0 -c 'GTTAACCCAAGGGCATT' *.fastq > count/B3245_Fm0.txt
agrep -0 -c 'GCAGTCGAGATTACATT' *.fastq > count/B3246_Fm0.txt
agrep -0 -c 'CTACGGTGTGCCATGGA' *.fastq > count/B3247_Fm0.txt
agrep -0 -c 'TACAATCCGCCTTGATC' *.fastq > count/B3248_Fm0.txt
agrep -0 -c 'TTGCTTTCTCGGTGTAA' *.fastq > count/B3249_Fm0.txt
agrep -0 -c 'CGTCAACACACCTTCCC' *.fastq > count/B3250_Fm0.txt
agrep -0 -c 'CTATCGTGCAACCTCTT' *.fastq > count/B3252_Fm0.txt
agrep -0 -c 'GTATCAGGAGTCTACAG' *.fastq > count/B3253_Fm0.txt
agrep -0 -c 'ATCCAGCTCACATCGAT' *.fastq > count/B3254_Fm0.txt
agrep -0 -c 'TACGGTGGTGCCTATCG' *.fastq > count/B3255_Fm0.txt
agrep -0 -c 'CTCTGAGTATGATAAGC' *.fastq > count/B3256_Fm0.txt
agrep -0 -c 'ATAAGAGAGCGACACGA' *.fastq > count/B3257_Fm0.txt
agrep -0 -c 'GGATTTGGAGGGCAGCT' *.fastq > count/B3258_Fm0.txt
agrep -0 -c 'TTATACGGACTGCTGGC' *.fastq > count/B3259_Fm0.txt
agrep -0 -c 'AGCAAGTCGCGGTGTTC' *.fastq > count/B3260_Fm0.txt
agrep -0 -c 'GTTTACGTTGTGCAACA' *.fastq > count/B3261_Fm0.txt
agrep -0 -c 'TACGGACACAAGCTCGG' *.fastq > count/B3262_Fm0.txt
agrep -0 -c 'GTCCGAACATTAATGGT' *.fastq > count/B3263_Fm0.txt
agrep -0 -c 'CAGGTGAAGTTACTAAC' *.fastq > count/B3264_Fm0.txt
agrep -0 -c 'ATACCGTCCAACGGTTA' *.fastq > count/B3265_Fm0.txt
agrep -0 -c 'AGCTACAAAGCCATCCG' *.fastq > count/B3266_Fm0.txt
agrep -0 -c 'GGAGTATTGTCAGCGAC' *.fastq > count/B3267_Fm0.txt
agrep -0 -c 'GAGGGAGTACGTAGAGC' *.fastq > count/B3268_Fm0.txt
agrep -0 -c 'GCAGAAATGTCCCGTAG' *.fastq > count/B3269_Fm0.txt
agrep -0 -c 'ACTATCGGTGCTCGAGT' *.fastq > count/B3270_Fm0.txt
agrep -0 -c 'ACAGGACTTAGTGCCAG' *.fastq > count/B3271_Fm0.txt
agrep -0 -c 'CGGAATCCTTGTAAGTG' *.fastq > count/B3272_Fm0.txt
agrep -0 -c 'CGAATGGTCGCAAGCGT' *.fastq > count/B3273_Fm0.txt
agrep -0 -c 'TGTAGACCAGTGCGTAC' *.fastq > count/B3274_Fm0.txt
agrep -0 -c 'TAGCTCATTGATGCATG' *.fastq > count/B3275_Fm0.txt
agrep -0 -c 'TACAGGCAACTTGGAGA' *.fastq > count/B3276_Fm0.txt
agrep -0 -c 'TTGGTTTCCGGGAGCCT' *.fastq > count/B3277_Fm0.txt
agrep -0 -c 'CATAGATTGGGACTCTG' *.fastq > count/B3278_Fm0.txt
agrep -0 -c 'GCGATCTTGGGATCACG' *.fastq > count/B3280_Fm0.txt
agrep -0 -c 'TTAAGCATTCCGTGGGC' *.fastq > count/B3282_Fm0.txt
agrep -0 -c 'GGATCTTTGAGGCCAGT' *.fastq > count/B3283_Fm0.txt
agrep -0 -c 'AATAGATTGGGGGACGA' *.fastq > count/B3284_Fm0.txt
agrep -0 -c 'AGAGGTCTTCCTAGCCG' *.fastq > count/B3285_Fm0.txt
agrep -0 -c 'TTGTAACCTCTGCCAGT' *.fastq > count/B3287_Fm0.txt
agrep -0 -c 'ATATATTTGCTCCGAGC' *.fastq > count/B3289_Fm0.txt
agrep -0 -c 'CGTGACTACCCTGAGAA' *.fastq > count/B3290_Fm0.txt
agrep -0 -c 'ATCCGAATAGCCATTTG' *.fastq > count/B3292_Fm0.txt
agrep -0 -c 'CATTTATGCGCACTAGC' *.fastq > count/B3293_Fm0.txt
agrep -0 -c 'TGTCTTGTCTCGACCGG' *.fastq > count/B3294_Fm0.txt
agrep -0 -c 'CGGATGCTCTCACCTAA' *.fastq > count/B3295_Fm0.txt
agrep -0 -c 'ACCCAGTGCTGAACATG' *.fastq > count/B3296_Fm0.txt
agrep -0 -c 'GGCAAGTGCAACATGTG' *.fastq > count/B3297_Fm0.txt
agrep -0 -c 'TTCGACCAAGAGGCGGT' *.fastq > count/B3298_Fm0.txt
agrep -0 -c 'GCTTGCTATGTTCGATA' *.fastq > count/B3299_Fm0.txt
agrep -0 -c 'TTCCTCCAACTGCAGTC' *.fastq > count/B3300_Fm0.txt
agrep -0 -c 'TGTGCTTCGGGTTAGCA' *.fastq > count/B3301_Fm0.txt
agrep -0 -c 'GTGTTACAGAGCTTGCC' *.fastq > count/B3302_Fm0.txt
agrep -0 -c 'TGTTGCCCCAGAAAGGG' *.fastq > count/B3303_Fm0.txt
agrep -0 -c 'CTTAGTCGGAACCTACC' *.fastq > count/B3304_Fm0.txt
agrep -0 -c 'TACCTGCAGATAGAAGG' *.fastq > count/B3305_Fm0.txt
agrep -0 -c 'ACCAGGCGGCTAACAAA' *.fastq > count/B3306_Fm0.txt
agrep -0 -c 'TTGCACCTATTCGACGT' *.fastq > count/B3307_Fm0.txt
agrep -0 -c 'AGGGACCAGTTGAGCAA' *.fastq > count/B3309_Fm0.txt
agrep -0 -c 'GGATATCCGGTACTACC' *.fastq > count/B3310_Fm0.txt
agrep -0 -c 'GCCGCGCCTTCATTGTT' *.fastq > count/B3311_Fm0.txt
agrep -0 -c 'GGGCGTTATGTAGAGCT' *.fastq > count/B3312_Fm0.txt
agrep -0 -c 'GTATTATCCCTCGCCGC' *.fastq > count/B3313_Fm0.txt
agrep -0 -c 'CTACATGCCACCCCTTG' *.fastq > count/B3314_Fm0.txt
agrep -0 -c 'AATAGGCATACCCTAGA' *.fastq > count/B3315_Fm0.txt
agrep -0 -c 'GGTGGGGTAAAAACGTC' *.fastq > count/B3318_Fm0.txt
agrep -0 -c 'CGAGCGTTCGTAACGTG' *.fastq > count/B3319_Fm0.txt
agrep -0 -c 'CTCGGCACTGCTTATGT' *.fastq > count/B3320_Fm0.txt
agrep -0 -c 'ATTCGAATGGGACGGGA' *.fastq > count/B3321_Fm0.txt
agrep -0 -c 'TGAGATGGCAATAACCG' *.fastq > count/B3322_Fm0.txt
agrep -0 -c 'TTCTAGTAGCGCCTGCG' *.fastq > count/B3323_Fm0.txt
agrep -0 -c 'ACAGTACCCTCAGTACT' *.fastq > count/B3324_Fm0.txt
agrep -0 -c 'TAGGGTTACGCAGATCC' *.fastq > count/B3325_Fm0.txt
agrep -0 -c 'CAACAGGGACTTGTACG' *.fastq > count/B3326_Fm0.txt
agrep -0 -c 'CTAGTCGTGTGGATCAC' *.fastq > count/B3327_Fm0.txt
agrep -0 -c 'TGAGTAACAGAGTGCGT' *.fastq > count/B3328_Fm0.txt
agrep -0 -c 'GTAGAAAGGAAGTCGTA' *.fastq > count/B3329_Fm0.txt
agrep -0 -c 'TGTGCCGTCCGTTATTT' *.fastq > count/B3330_Fm0.txt
agrep -0 -c 'TAGATCACTGGGTCATC' *.fastq > count/B3331_Fm0.txt
agrep -0 -c 'GTGATATGGTGCGAGTG' *.fastq > count/B3332_Fm0.txt
agrep -0 -c 'GGAAGTAGTTCGGATAG' *.fastq > count/B3333_Fm0.txt
agrep -0 -c 'GCTTCGCATGATAGCTT' *.fastq > count/B3336_Fm0.txt
agrep -0 -c 'GCGCTCTAAGCCCATAC' *.fastq > count/B3337_Fm0.txt
agrep -0 -c 'CTCGTGTTTGTCTGCTC' *.fastq > count/B3338_Fm0.txt
agrep -0 -c 'CACTTACGCCGTGTGTC' *.fastq > count/B3339_Fm0.txt
agrep -0 -c 'AATTAACACCGTCCTTC' *.fastq > count/B3340_Fm0.txt
agrep -0 -c 'AAGTCACAACGCGATTA' *.fastq > count/B3341_Fm0.txt
agrep -0 -c 'TGCCGTATGCATGAAGA' *.fastq > count/B3342_Fm0.txt
agrep -0 -c 'TAGCCCTAGTCGCATGG' *.fastq > count/B3343_Fm0.txt
agrep -0 -c 'ACTTGACGCCCTTGCGA' *.fastq > count/B3344_Fm0.txt
agrep -0 -c 'TACGGTAACGCGACGAA' *.fastq > count/B3348_Fm0.txt
agrep -0 -c 'GGCTCGTATACTCCATC' *.fastq > count/B3349_Fm0.txt
agrep -0 -c 'ATAGAGAGGATTAGCGA' *.fastq > count/B3353_Fm0.txt
agrep -0 -c 'AGTCGATTGGTAGTCAT' *.fastq > count/B3355_Fm0.txt
agrep -0 -c 'CGTGGGGTGACGAACTT' *.fastq > count/B3357_Fm0.txt
agrep -0 -c 'GGAATCGAGTGTTTGGC' *.fastq > count/B3358_Fm0.txt
agrep -0 -c 'CAAGATAGCTGGGCGAC' *.fastq > count/B3359_Fm0.txt
agrep -0 -c 'CTGGCGACAGAGGTCTT' *.fastq > count/B3360_Fm0.txt
agrep -0 -c 'TGTACGCGGGTTCGATT' *.fastq > count/B3362_Fm0.txt
agrep -0 -c 'TAGACGAAAGCACTCGG' *.fastq > count/B3363_Fm0.txt
agrep -0 -c 'GCCCGATTGAAACTTCT' *.fastq > count/B3364_Fm0.txt
agrep -0 -c 'CTCCTTGGGAAGGTTCT' *.fastq > count/B3365_Fm0.txt
agrep -0 -c 'AAGTTATAGGTTAGCCG' *.fastq > count/B3366_Fm0.txt
agrep -0 -c 'AAGGGGTGAACCTCTTG' *.fastq > count/B3367_Fm0.txt
agrep -0 -c 'GGAGAGATAACCTAAGG' *.fastq > count/B3370_Fm0.txt
agrep -0 -c 'TGCACTTGTACGTATTG' *.fastq > count/B3371_Fm0.txt
agrep -0 -c 'CAATAGTCTAATGCACC' *.fastq > count/B3374_Fm0.txt
agrep -0 -c 'GCTAAATAGGTTACTCG' *.fastq > count/B3375_Fm0.txt
agrep -0 -c 'GATCACCTTCCTACGGT' *.fastq > count/B3376_Fm0.txt
agrep -0 -c 'TTATGGGTAGGACATAC' *.fastq > count/B3377_Fm0.txt
agrep -0 -c 'AACCAGTGACGCAGCTG' *.fastq > count/B3380_Fm0.txt
agrep -0 -c 'CTCCACGTGCTCTTATA' *.fastq > count/B3381_Fm0.txt
agrep -0 -c 'CGAACCACGGGCAGATT' *.fastq > count/B3382_Fm0.txt
agrep -0 -c 'TGTGTGGTCGTTTTAAC' *.fastq > count/B3383_Fm0.txt
agrep -0 -c 'GGGGTCTATGAGCGTTC' *.fastq > count/B3384_Fm0.txt
agrep -0 -c 'ATAATTGTCACCGTGGA' *.fastq > count/B3386_Fm0.txt
agrep -0 -c 'CGCGAGTAAGGGAGTCA' *.fastq > count/B3387_Fm0.txt
agrep -0 -c 'ATCTGCCTTCGGGAGCA' *.fastq > count/B3389_Fm0.txt
agrep -0 -c 'GACGTTACCAAGCAATC' *.fastq > count/B3390_Fm0.txt
agrep -0 -c 'CGAGCTTATGAAAGGAC' *.fastq > count/B3391_Fm0.txt
agrep -0 -c 'TGGCATTTAGGTTGTAG' *.fastq > count/B3392_Fm0.txt
agrep -0 -c 'TTCACAAAATCCCACCA' *.fastq > count/B3393_Fm0.txt
agrep -0 -c 'GTCGTCACAGTTATAGC' *.fastq > count/B3394_Fm0.txt
agrep -0 -c 'GAGCCGAGTCTAGATAT' *.fastq > count/B3395_Fm0.txt
agrep -0 -c 'GTGTGACTCGCAGTCGT' *.fastq > count/B3396_Fm0.txt
agrep -0 -c 'CGGGTATATGTTAACGT' *.fastq > count/B3397_Fm0.txt
agrep -0 -c 'GTAGTTTTCACGACGAG' *.fastq > count/B3398_Fm0.txt
agrep -0 -c 'ACTCAACACCTTTTCGG' *.fastq > count/B3399_Fm0.txt
agrep -0 -c 'ATTTGGAACCGTGGAAG' *.fastq > count/B3400_Fm0.txt
agrep -0 -c 'TTCAAGTTAAACGGCCC' *.fastq > count/B3401_Fm0.txt
agrep -0 -c 'CAGGGTGATCTAGAAGG' *.fastq > count/B3402_Fm0.txt
agrep -0 -c 'AGCTATAACCAGTGACT' *.fastq > count/B3403_Fm0.txt
agrep -0 -c 'AATGGCCACTTTGAAAC' *.fastq > count/B3404_Fm0.txt
agrep -0 -c 'ATACCTCAGTCGAGTCC' *.fastq > count/B3406_Fm0.txt
agrep -0 -c 'GTTTATGTGCAGTTCTC' *.fastq > count/B3407_Fm0.txt
agrep -0 -c 'GACCGGTTTCGATTAAA' *.fastq > count/B3408_Fm0.txt
agrep -0 -c 'CCTCTTTGAAGCCGAGA' *.fastq > count/B3410_Fm0.txt
agrep -0 -c 'CAGCAGATACGTGTCAC' *.fastq > count/B3412_Fm0.txt
agrep -0 -c 'CGGGGACATAATCTGAT' *.fastq > count/B3413_Fm0.txt
agrep -0 -c 'ACGTTTGGAACCCTCTT' *.fastq > count/B3414_Fm0.txt
agrep -0 -c 'CACTACAAACAGCAATC' *.fastq > count/B3415_Fm0.txt
agrep -0 -c 'TAGCACCTAGGTAGTTA' *.fastq > count/B3417_Fm0.txt
agrep -0 -c 'TCACAGACGCACTGTAG' *.fastq > count/B3418_Fm0.txt
agrep -0 -c 'TAAGTCTCGCGATAGTT' *.fastq > count/B3419_Fm0.txt
agrep -0 -c 'CGAGGAACTAGAGCAAT' *.fastq > count/B3420_Fm0.txt
agrep -0 -c 'GTTCGGCATGAATATTC' *.fastq > count/B3421_Fm0.txt
agrep -0 -c 'TATGCGTGTAGTACCCA' *.fastq > count/B3422_Fm0.txt
agrep -0 -c 'GGGGAGAATAGGTCATT' *.fastq > count/B3423_Fm0.txt
agrep -0 -c 'GATTAGGGAGCTATGGT' *.fastq > count/B3424_Fm0.txt
agrep -0 -c 'TTAAGATCACGGGCATC' *.fastq > count/B3426_Fm0.txt
agrep -0 -c 'GGCTTACAGCTCAATGC' *.fastq > count/B3427_Fm0.txt
agrep -0 -c 'CCTATACGAAGCCGAGC' *.fastq > count/B3428_Fm0.txt
agrep -0 -c 'ATAAGCAGTACTAGGGC' *.fastq > count/B3429_Fm0.txt
agrep -0 -c 'TTGTCGACTCTCGAGGC' *.fastq > count/B3430_Fm0.txt
agrep -0 -c 'CACACCTTGATCGATAG' *.fastq > count/B3431_Fm0.txt
agrep -0 -c 'CTAGCGGGGCTGATTTA' *.fastq > count/B3432_Fm0.txt
agrep -0 -c 'TAAGCTACTAGAGTGGG' *.fastq > count/B3433_Fm0.txt
agrep -0 -c 'GCAGATCCTCTACAAGG' *.fastq > count/B3434_Fm0.txt
agrep -0 -c 'ATGTTGAAAGTCGGTTC' *.fastq > count/B3435_Fm0.txt
agrep -0 -c 'TGCCGACAACGTTAGCT' *.fastq > count/B3436_Fm0.txt
agrep -0 -c 'CACCATTTACTCCCCGT' *.fastq > count/B3437_Fm0.txt
agrep -0 -c 'ACAGCCGTCGGTTTTAG' *.fastq > count/B3438_Fm0.txt
agrep -0 -c 'CTATGCGCGTGATCAAA' *.fastq > count/B3439_Fm0.txt
agrep -0 -c 'CAAGCGAACCGCCCTTA' *.fastq > count/B3440_Fm0.txt
agrep -0 -c 'AAGCCGCAAAACAACAT' *.fastq > count/B3441_Fm0.txt
agrep -0 -c 'TTGCATTGGGGATTCCG' *.fastq > count/B3442_Fm0.txt
agrep -0 -c 'CTAACCTGTATATGGGC' *.fastq > count/B3443_Fm0.txt
agrep -0 -c 'GATACCTTAATCGTACG' *.fastq > count/B3444_Fm0.txt
agrep -0 -c 'GCACTCGCACAACGATC' *.fastq > count/B3446_Fm0.txt
agrep -0 -c 'GCCCCTTAGCTGAACCT' *.fastq > count/B3447_Fm0.txt
agrep -0 -c 'CCGTTAAGGCGAAGGTA' *.fastq > count/B3448_Fm0.txt
agrep -0 -c 'TAAGGTAACCGTCCGTT' *.fastq > count/B3449_Fm0.txt
agrep -0 -c 'AGTCAGGGTGTCGCATG' *.fastq > count/B3451_Fm0.txt
agrep -0 -c 'GCGAAGCACTCTGTTCT' *.fastq > count/B3452_Fm0.txt
agrep -0 -c 'ATCGGGCTTGGTGAAGG' *.fastq > count/B3453_Fm0.txt
agrep -0 -c 'TCGGATCTACATCAGCG' *.fastq > count/B3454_Fm0.txt
agrep -0 -c 'GGTTTTGATCAGAGATG' *.fastq > count/B3456_Fm0.txt
agrep -0 -c 'CAAAGCGGACGTCTAGA' *.fastq > count/B3457_Fm0.txt
agrep -0 -c 'CTCCAGCTCAGACGTAC' *.fastq > count/B3458_Fm0.txt
agrep -0 -c 'AGGCATTCCACAATGGG' *.fastq > count/B3459_Fm0.txt
agrep -0 -c 'TATGCCTCACTGGTGTA' *.fastq > count/B3460_Fm0.txt
agrep -0 -c 'AGTGGCCTTTCTTCTCG' *.fastq > count/B3461_Fm0.txt
agrep -0 -c 'AGAACGCTGAACCCTAG' *.fastq > count/B3462_Fm0.txt
agrep -0 -c 'CTTCTACACCGGCTAAA' *.fastq > count/B3463_Fm0.txt
agrep -0 -c 'GCCCACTGTCACAAAAA' *.fastq > count/B3464_Fm0.txt
agrep -0 -c 'ACGGAGCTTAAGACCGA' *.fastq > count/B3465_Fm0.txt
agrep -0 -c 'GTACGACTTACATGTCA' *.fastq > count/B3467_Fm0.txt
agrep -0 -c 'AGGTATAGCTTGGCAGC' *.fastq > count/B3468_Fm0.txt
agrep -0 -c 'CAATAGTAAATGGCGGC' *.fastq > count/B3469_Fm0.txt
agrep -0 -c 'TATGGTCGTACAGTAAC' *.fastq > count/B3471_Fm0.txt
agrep -0 -c 'AGACCCCGCTTGCACTT' *.fastq > count/B3472_Fm0.txt
agrep -0 -c 'TTAAAGGTAGTGAAGGC' *.fastq > count/B3473_Fm0.txt
agrep -0 -c 'TGCACAACCTTCCGTAT' *.fastq > count/B3474_Fm0.txt
agrep -0 -c 'AGGCCGTCAGATCCTTC' *.fastq > count/B3475_Fm0.txt
agrep -0 -c 'CAGATCGTCCCGGAGTA' *.fastq > count/B3476_Fm0.txt
agrep -0 -c 'TTAGTCGGTCAGTCCGC' *.fastq > count/B3477_Fm0.txt
agrep -0 -c 'ATAAGTCAAATCCTGCC' *.fastq > count/B3479_Fm0.txt
agrep -0 -c 'GCGTAGCATAGTCACGT' *.fastq > count/B3480_Fm0.txt
agrep -0 -c 'GTCACTAAACTCCGGGA' *.fastq > count/B3481_Fm0.txt
agrep -0 -c 'TGTAGCTGCTGTGCAAT' *.fastq > count/B3482_Fm0.txt
agrep -0 -c 'TCGTACCGTCCAACACA' *.fastq > count/B3483_Fm0.txt
agrep -0 -c 'ACGGTCCGCGTAATTAC' *.fastq > count/B3484_Fm0.txt
agrep -0 -c 'TGTCCAAAGACATACGT' *.fastq > count/B3485_Fm0.txt
agrep -0 -c 'TTAGAACGGCTAGGCCT' *.fastq > count/B3486_Fm0.txt
agrep -0 -c 'TGCACTCCATTCCGACT' *.fastq > count/B3487_Fm0.txt
agrep -0 -c 'TTGGGTCAAGTCACCCG' *.fastq > count/B3488_Fm0.txt
agrep -0 -c 'AAGGTTTGTCAGCCACG' *.fastq > count/B3490_Fm0.txt
agrep -0 -c 'GATTCCAGATCCAGCAT' *.fastq > count/B3491_Fm0.txt
agrep -0 -c 'CCTCTCTCGTGAAAGGG' *.fastq > count/B3492_Fm0.txt
agrep -0 -c 'ATCCTAAAAACGGGCAG' *.fastq > count/B3493_Fm0.txt
agrep -0 -c 'CGTTGATGCTTAGGGTC' *.fastq > count/B3494_Fm0.txt
agrep -0 -c 'ACATCGATAGCACGTGT' *.fastq > count/B3495_Fm0.txt
agrep -0 -c 'TCAGATGTATGCACCAC' *.fastq > count/B3496_Fm0.txt
agrep -0 -c 'AGAGCGAACGCTCATCC' *.fastq > count/B3497_Fm0.txt
agrep -0 -c 'GTACTCTCTACGGTTTC' *.fastq > count/B3498_Fm0.txt
agrep -0 -c 'AGGCCTTGAGACATGTA' *.fastq > count/B3499_Fm0.txt
agrep -0 -c 'CGGCTTTCTCAACATCT' *.fastq > count/B3500_Fm0.txt
agrep -0 -c 'TGCCGGTGTATCTAATT' *.fastq > count/B3501_Fm0.txt
agrep -0 -c 'GGTGGGCGAAAATAAGA' *.fastq > count/B3502_Fm0.txt
agrep -0 -c 'ATTGTTGGGTCGAGTAC' *.fastq > count/B3503_Fm0.txt
agrep -0 -c 'TACGTGTACGGGTTCCT' *.fastq > count/B3504_Fm0.txt
agrep -0 -c 'GTAGGAGGTGATGTAGC' *.fastq > count/B3506_Fm0.txt
agrep -0 -c 'TAACGGTCAGCGAATAG' *.fastq > count/B3507_Fm0.txt
agrep -0 -c 'TGCACGTGCTCGTACTT' *.fastq > count/B3508_Fm0.txt
agrep -0 -c 'CAGGACCTTTCCAGTAC' *.fastq > count/B3509_Fm0.txt
agrep -0 -c 'TACCTGGGTGGTGATGC' *.fastq > count/B3510_Fm0.txt
agrep -0 -c 'TGCTGGTGGTCAAATAT' *.fastq > count/B3512_Fm0.txt
agrep -0 -c 'GGTGGCAACGGCTTTCA' *.fastq > count/B3513_Fm0.txt
agrep -0 -c 'CGTTCGCAAATGAGACA' *.fastq > count/B3515_Fm0.txt
agrep -0 -c 'ACGGCCTTATGTAGTCG' *.fastq > count/B3517_Fm0.txt
agrep -0 -c 'TGAGGCCAAAGTCCGTC' *.fastq > count/B3518_Fm0.txt
agrep -0 -c 'ACAATAGGAATCCGCCG' *.fastq > count/B3519_Fm0.txt
agrep -0 -c 'CTTGATATTGGGGATCG' *.fastq > count/B3520_Fm0.txt
agrep -0 -c 'TACCGAAGTCCGCAGAA' *.fastq > count/B3523_Fm0.txt
agrep -0 -c 'TAGGCCGCGAAGTGTGA' *.fastq > count/B3524_Fm0.txt
agrep -0 -c 'AGATAGAGATCATGCGC' *.fastq > count/B3525_Fm0.txt
agrep -0 -c 'GCGAGAACAGACGATGT' *.fastq > count/B3526_Fm0.txt
agrep -0 -c 'TCTATAGTGTTGGGTTG' *.fastq > count/B3527_Fm0.txt
agrep -0 -c 'TGATCGAGGGTCTATTC' *.fastq > count/B3528_Fm0.txt
agrep -0 -c 'CAAACTCGCCTTAGCTC' *.fastq > count/B3529_Fm0.txt
agrep -0 -c 'AGTGTTCTCAGGTCCCC' *.fastq > count/B3530_Fm0.txt
agrep -0 -c 'AGCGTTGAATCGGTTAA' *.fastq > count/B3531_Fm0.txt
agrep -0 -c 'GTGGGCGTTGGCAAACA' *.fastq > count/B3532_Fm0.txt
agrep -0 -c 'TGTCGAACCTAGAACTC' *.fastq > count/B3533_Fm0.txt
agrep -0 -c 'ATCCGCTTAGATCAAGT' *.fastq > count/B3534_Fm0.txt
agrep -0 -c 'CCATTGGGTCTGATTCC' *.fastq > count/B3535_Fm0.txt
agrep -0 -c 'TAGTCCCTCCGATATGT' *.fastq > count/B3536_Fm0.txt
agrep -0 -c 'TATAGTAAGGCGAGCAA' *.fastq > count/B3537_Fm0.txt
agrep -0 -c 'GCCAATAGCCGTAATTA' *.fastq > count/B3540_Fm0.txt
agrep -0 -c 'AGACACTCCGTCCCAAG' *.fastq > count/B3541_Fm0.txt
agrep -0 -c 'GTGAGTAGACATGGACT' *.fastq > count/B3542_Fm0.txt
agrep -0 -c 'CCTGACGTCATGCACGA' *.fastq > count/B3543_Fm0.txt
agrep -0 -c 'CCGTTTTCGGTGCCAGT' *.fastq > count/B3544_Fm0.txt
agrep -0 -c 'GTGCTTAACAGTGCGTT' *.fastq > count/B3545_Fm0.txt
agrep -0 -c 'GACGGTAGAACGATCCC' *.fastq > count/B3546_Fm0.txt
agrep -0 -c 'TGAAACCGCTATCTCCG' *.fastq > count/B3547_Fm0.txt
agrep -0 -c 'TCATTGGCAGTACCCAC' *.fastq > count/B3548_Fm0.txt
agrep -0 -c 'AGGCGGCATTACTGAAA' *.fastq > count/B3550_Fm0.txt
agrep -0 -c 'CTGGAGCTGGTACGAAG' *.fastq > count/B3551_Fm0.txt
agrep -0 -c 'ACGAGCTCTACAGGGAT' *.fastq > count/B3552_Fm0.txt
agrep -0 -c 'CCTTAAAGGGGTTGAAC' *.fastq > count/B3553_Fm0.txt
agrep -0 -c 'GCATGACAGTGGCATGA' *.fastq > count/B3554_Fm0.txt
agrep -0 -c 'AGTTAGTAAAGTACCGG' *.fastq > count/B3555_Fm0.txt
agrep -0 -c 'GCCAGTCAGAGAGACTC' *.fastq > count/B3557_Fm0.txt
agrep -0 -c 'CCTCCCACACGCGATAA' *.fastq > count/B3558_Fm0.txt
agrep -0 -c 'GTGGTGCATAGAGACTA' *.fastq > count/B3559_Fm0.txt
agrep -0 -c 'AGCCACAACTCCCAATG' *.fastq > count/B3560_Fm0.txt
agrep -0 -c 'TGCCAGATTATGCCGCC' *.fastq > count/B3561_Fm0.txt
agrep -0 -c 'GGTTAATGAATGCGGCC' *.fastq > count/B3562_Fm0.txt
agrep -0 -c 'AGTGAACGATCTCGATA' *.fastq > count/B3563_Fm0.txt
agrep -0 -c 'AGTCAGGCTGGAAGTAA' *.fastq > count/B3564_Fm0.txt
agrep -0 -c 'AGTCGGAAGTTATACCT' *.fastq > count/B3565_Fm0.txt
agrep -0 -c 'TATTTAGGACCAGAGCC' *.fastq > count/B3566_Fm0.txt
agrep -0 -c 'TACCCAATCACCTGAAG' *.fastq > count/B3567_Fm0.txt
agrep -0 -c 'AGTCGAGATGTTGTTGG' *.fastq > count/B3568_Fm0.txt
agrep -0 -c 'TTGTCGGACTTGCCTCC' *.fastq > count/B3569_Fm0.txt
agrep -0 -c 'CTACTTAGAAAGGGACT' *.fastq > count/B3570_Fm0.txt
agrep -0 -c 'CTGGTGAGTAGTCGGCA' *.fastq > count/B3571_Fm0.txt
agrep -0 -c 'GCCGTGGCGTCTATTCT' *.fastq > count/B3572_Fm0.txt
agrep -0 -c 'AGTGTGGTCGCCCACAA' *.fastq > count/B3573_Fm0.txt
agrep -0 -c 'TATCGTCTTCTGTGAGG' *.fastq > count/B3574_Fm0.txt
agrep -0 -c 'CACATTTCCGGAGGGAC' *.fastq > count/B3575_Fm0.txt
agrep -0 -c 'TTCTACTGGCCCGCATG' *.fastq > count/B3577_Fm0.txt
agrep -0 -c 'GTATCATCTAGGGACAT' *.fastq > count/B3578_Fm0.txt
agrep -0 -c 'GTCGGAAAGCGGTAGAA' *.fastq > count/B3579_Fm0.txt
agrep -0 -c 'ATCCCAACCTACCAAGG' *.fastq > count/B3580_Fm0.txt
agrep -0 -c 'AGTCTTGTTGGACAACG' *.fastq > count/B3582_Fm0.txt
agrep -0 -c 'ACTTACTCTCTCAATCG' *.fastq > count/B3583_Fm0.txt
agrep -0 -c 'GACTTGTACTAGCCCGT' *.fastq > count/B3584_Fm0.txt
agrep -0 -c 'CTCTTTTTCCTCAGAGT' *.fastq > count/B3585_Fm0.txt
agrep -0 -c 'TGCTCTTTGCGTCTCGT' *.fastq > count/B3586_Fm0.txt
agrep -0 -c 'GGAGGCGTATTCGCGAT' *.fastq > count/B3587_Fm0.txt
agrep -0 -c 'CTGTTACAAAAGAGGCT' *.fastq > count/B3589_Fm0.txt
agrep -0 -c 'GGGCATAGGTCATATAC' *.fastq > count/B3590_Fm0.txt
agrep -0 -c 'CGGGACCAATTGTCACG' *.fastq > count/B3591_Fm0.txt
agrep -0 -c 'GTTATCAGCGAGTTGCG' *.fastq > count/B3592_Fm0.txt
agrep -0 -c 'ACGGGAATCATCGTATA' *.fastq > count/B3594_Fm0.txt
agrep -0 -c 'CCACGCGCACATGTAAG' *.fastq > count/B3595_Fm0.txt
agrep -0 -c 'CCATGTGGTTGTAGTGA' *.fastq > count/B3596_Fm0.txt
agrep -0 -c 'GCAAGACATGCGTGCTG' *.fastq > count/B3597_Fm0.txt
agrep -0 -c 'ACGATGCTTCGAGATAG' *.fastq > count/B3598_Fm0.txt
agrep -0 -c 'CTCCCCCGAGTATTATC' *.fastq > count/B3599_Fm0.txt
agrep -0 -c 'ATGTTTTCCGGTAACCT' *.fastq > count/B3600_Fm0.txt
agrep -0 -c 'CTATGGCACGACCCGAT' *.fastq > count/B3601_Fm0.txt
agrep -0 -c 'TATGGTTAACAGACCCG' *.fastq > count/B3603_Fm0.txt
agrep -0 -c 'AACTCACTCCGGCATGA' *.fastq > count/B3605_Fm0.txt
agrep -0 -c 'TTGTATAACAGGGAGGA' *.fastq > count/B3606_Fm0.txt
agrep -0 -c 'AGATCATGAACGCGCCT' *.fastq > count/B3607_Fm0.txt
agrep -0 -c 'TGACCATGTCCGAGAAT' *.fastq > count/B3608_Fm0.txt
agrep -0 -c 'CGTTTACGCGACCTCTA' *.fastq > count/B3609_Fm0.txt
agrep -0 -c 'CCTCCTATCTGGGCATT' *.fastq > count/B3610_Fm0.txt
agrep -0 -c 'TTGCTTTATTTGCCGCT' *.fastq > count/B3611_Fm0.txt
agrep -0 -c 'GCGAAACCTCGAAAAAC' *.fastq > count/B3612_Fm0.txt
agrep -0 -c 'GGCTGGAATGTCGGTTC' *.fastq > count/B3614_Fm0.txt
agrep -0 -c 'ATTTGAAACCCGCACTC' *.fastq > count/B3616_Fm0.txt
agrep -0 -c 'CAACGAGACTAGGGTTG' *.fastq > count/B3617_Fm0.txt
agrep -0 -c 'TGACCAGAGTCAGTCTT' *.fastq > count/B3618_Fm0.txt
agrep -0 -c 'TGCTATTAACAGCACGT' *.fastq > count/B3619_Fm0.txt
agrep -0 -c 'TACCGCTTACCGTTCCA' *.fastq > count/B3620_Fm0.txt
agrep -0 -c 'GCTTCGACACCATTTCG' *.fastq > count/B3621_Fm0.txt
agrep -0 -c 'ATTGGGGCGATTGGATT' *.fastq > count/B3622_Fm0.txt
agrep -0 -c 'TGACGAGAAACGTGGGT' *.fastq > count/B3623_Fm0.txt
agrep -0 -c 'CAAGGGTGATGCGATGC' *.fastq > count/B3624_Fm0.txt
agrep -0 -c 'TCGCTCACAGCTTCTAG' *.fastq > count/B3626_Fm0.txt
agrep -0 -c 'CTCGACAGACTCTTTAA' *.fastq > count/B3627_Fm0.txt
agrep -0 -c 'CGGTTTAGGGATCTTTT' *.fastq > count/B3628_Fm0.txt
agrep -0 -c 'AAGCGGACCTTGAAGGC' *.fastq > count/B3629_Fm0.txt
agrep -0 -c 'GACCTGGGAATGCACGA' *.fastq > count/B3630_Fm0.txt
agrep -0 -c 'TACGACGCAGACTGTGG' *.fastq > count/B3631_Fm0.txt
agrep -0 -c 'GTCCATCGGAACGTAAA' *.fastq > count/B3632_Fm0.txt
agrep -0 -c 'CCATAAACCGCGTCAGC' *.fastq > count/B3633_Fm0.txt
agrep -0 -c 'AGAGCCCGCCAATCGTA' *.fastq > count/B3634_Fm0.txt
agrep -0 -c 'ATGACCAAGCTCGTGGC' *.fastq > count/B3635_Fm0.txt
agrep -0 -c 'GTCCAACTACAGCCGTA' *.fastq > count/B3636_Fm0.txt
agrep -0 -c 'CCGCTCATACCAGTTGG' *.fastq > count/B3637_Fm0.txt
agrep -0 -c 'ACTATTCCCGCTAATAC' *.fastq > count/B3638_Fm0.txt
agrep -0 -c 'ATATGGTGCGGGGATCT' *.fastq > count/B3639_Fm0.txt
agrep -0 -c 'TCAACGCAAGGTGTCAA' *.fastq > count/B3641_Fm0.txt
agrep -0 -c 'GTGGTCAGTAGGATCCG' *.fastq > count/B3642_Fm0.txt
agrep -0 -c 'CCACTAGCGCATTCAAT' *.fastq > count/B3643_Fm0.txt
agrep -0 -c 'ACGTAGATTAACTCCAC' *.fastq > count/B3644_Fm0.txt
agrep -0 -c 'TTAGTCCCTTGGAGGTG' *.fastq > count/B3645_Fm0.txt
agrep -0 -c 'TCTATGCGGAGGCTTTG' *.fastq > count/B3646_Fm0.txt
agrep -0 -c 'CAGGTATCTGTAGCCCC' *.fastq > count/B3647_Fm0.txt
agrep -0 -c 'GTGCTTAATTGAGCGTC' *.fastq > count/B3648_Fm0.txt
agrep -0 -c 'ATGCTTCACTTATAGGG' *.fastq > count/B3652_Fm0.txt
agrep -0 -c 'CTGATTTCGCAGCAAAT' *.fastq > count/B3653_Fm0.txt
agrep -0 -c 'TGCCGTGACGTGGATTA' *.fastq > count/B3654_Fm0.txt
agrep -0 -c 'TGTATCAGCGCAAACTC' *.fastq > count/B3655_Fm0.txt
agrep -0 -c 'GCCCCTCATCTTCAGAT' *.fastq > count/B3656_Fm0.txt
agrep -0 -c 'GATTTTTGCCTACGCAT' *.fastq > count/B3658_Fm0.txt
agrep -0 -c 'TTACCTCGTCAATCGGG' *.fastq > count/B3659_Fm0.txt
agrep -0 -c 'TACTGTCCCGTCAAATG' *.fastq > count/B3660_Fm0.txt
agrep -0 -c 'TGCCTTTTATAGTCCAG' *.fastq > count/B3661_Fm0.txt
agrep -0 -c 'ACCGTAGTCAGTCAATA' *.fastq > count/B3664_Fm0.txt
agrep -0 -c 'GATATTGCTGCTGATCG' *.fastq > count/B3666_Fm0.txt
agrep -0 -c 'TCTGCCGGATAACCCCT' *.fastq > count/B3668_Fm0.txt
agrep -0 -c 'CGGCGAACTTTAGCTCT' *.fastq > count/B3669_Fm0.txt
agrep -0 -c 'TGCTCAAACAATAGGCT' *.fastq > count/B3670_Fm0.txt
agrep -0 -c 'CCAAGTCGGCAGCAGTA' *.fastq > count/B3671_Fm0.txt
agrep -0 -c 'GCTCTTCCGCGTCGTTT' *.fastq > count/B3673_Fm0.txt
agrep -0 -c 'GTTAGAACCGTACAGGG' *.fastq > count/B3674_Fm0.txt
agrep -0 -c 'TTGTCCCTGCTAGACGA' *.fastq > count/B3675_Fm0.txt
agrep -0 -c 'CATCTGCCTTGATCGAT' *.fastq > count/B3676_Fm0.txt
agrep -0 -c 'GTCACAAGGAACGGCTC' *.fastq > count/B3677_Fm0.txt
agrep -0 -c 'ACGGCTTAGCCCTGGTT' *.fastq > count/B3678_Fm0.txt
agrep -0 -c 'GCGTCCGGTCAAATCCT' *.fastq > count/B3679_Fm0.txt
agrep -0 -c 'CAGAGTTGTGTTTCGGA' *.fastq > count/B3680_Fm0.txt
agrep -0 -c 'GTTCGTGCCACATGTCC' *.fastq > count/B3682_Fm0.txt
agrep -0 -c 'GTGCAAATATTCCCCTG' *.fastq > count/B3683_Fm0.txt
agrep -0 -c 'AGCTCAAAGCGGGCTAT' *.fastq > count/B3685_Fm0.txt
agrep -0 -c 'AACTAAACTCTCGTAGG' *.fastq > count/B3686_Fm0.txt
agrep -0 -c 'CGTCATGTTCGGATGCC' *.fastq > count/B3687_Fm0.txt
agrep -0 -c 'GCGAGGCAATGTTCCCA' *.fastq > count/B3688_Fm0.txt
agrep -0 -c 'GCTGGTGCCAGGATATG' *.fastq > count/B3689_Fm0.txt
agrep -0 -c 'CGACGTCTCGGTTGATG' *.fastq > count/B3691_Fm0.txt
agrep -0 -c 'ATCATCCCTGTTTCGCA' *.fastq > count/B3692_Fm0.txt
agrep -0 -c 'GATCCGGGTGTAATACC' *.fastq > count/B3693_Fm0.txt
agrep -0 -c 'TCGGGGTTAATTCCCCT' *.fastq > count/B3694_Fm0.txt
agrep -0 -c 'GTCCTAGGTCGATGCGA' *.fastq > count/B3695_Fm0.txt
agrep -0 -c 'GTTTCTCCTAATGGCCA' *.fastq > count/B3696_Fm0.txt
agrep -0 -c 'CTGTCGCCCAAATTATA' *.fastq > count/B3697_Fm0.txt
agrep -0 -c 'TAAGTTGGATCCCCCTT' *.fastq > count/B3698_Fm0.txt
agrep -0 -c 'CGATATGCAGGCGGAGT' *.fastq > count/B3699_Fm0.txt
agrep -0 -c 'CAGCGGGGGGACTTATT' *.fastq > count/B3700_Fm0.txt
agrep -0 -c 'AGTGAAAGGTAGTACAC' *.fastq > count/B3701_Fm0.txt
agrep -0 -c 'GAGTAGCTGTGACATCA' *.fastq > count/B3703_Fm0.txt
agrep -0 -c 'CGTCTACTCTTAGCACC' *.fastq > count/B3704_Fm0.txt
agrep -0 -c 'TCGCTACTTGATCCGAC' *.fastq > count/B3705_Fm0.txt
agrep -0 -c 'GGCAAGACACTCGACTC' *.fastq > count/B3706_Fm0.txt
agrep -0 -c 'GGCATAACAGTACGTGC' *.fastq > count/B3707_Fm0.txt
agrep -0 -c 'CAAGAACTGTAGTCGGA' *.fastq > count/B3708_Fm0.txt
agrep -0 -c 'CTGAGGCGAATTCTTCT' *.fastq > count/B3709_Fm0.txt
agrep -0 -c 'GCATTTTGATTCCGCTC' *.fastq > count/B3710_Fm0.txt
agrep -0 -c 'GGTTATCATCTGCATCC' *.fastq > count/B3711_Fm0.txt
agrep -0 -c 'ATATAAGCCCTTGCGCC' *.fastq > count/B3712_Fm0.txt
agrep -0 -c 'AGTCGACACTGCTCTTC' *.fastq > count/B3713_Fm0.txt
agrep -0 -c 'AATTAGCGTCCAGACCA' *.fastq > count/B3714_Fm0.txt
agrep -0 -c 'GGAACCGGCCATAGACT' *.fastq > count/B3715_Fm0.txt
agrep -0 -c 'AGCGGGTGAGTTCGTTC' *.fastq > count/B3716_Fm0.txt
agrep -0 -c 'ATCGCAGATGGGTGACT' *.fastq > count/B3717_Fm0.txt
agrep -0 -c 'GGTACGCTTACTGATGG' *.fastq > count/B3718_Fm0.txt
agrep -0 -c 'TAGGCGAGGTCCGCTTT' *.fastq > count/B3719_Fm0.txt
agrep -0 -c 'GGTGAGATAATCATGGG' *.fastq > count/B3720_Fm0.txt
agrep -0 -c 'CAGGTGGTCCTTCGGAT' *.fastq > count/B3721_Fm0.txt
agrep -0 -c 'AGGCGGGGATTAATGTC' *.fastq > count/B3722_Fm0.txt
agrep -0 -c 'GTACGTCATCCGATCCC' *.fastq > count/B3723_Fm0.txt
agrep -0 -c 'CAGACATTGCTAAAGCG' *.fastq > count/B3724_Fm0.txt
agrep -0 -c 'GACTAAAGAGCCTAAGT' *.fastq > count/B3725_Fm0.txt
agrep -0 -c 'GTGTCATATGCTCGGTA' *.fastq > count/B3726_Fm0.txt
agrep -0 -c 'TCGCACGATCAAAGGAC' *.fastq > count/B3727_Fm0.txt
agrep -0 -c 'CACGGCTACATCCGGAA' *.fastq > count/B3728_Fm0.txt
agrep -0 -c 'GGTTCTTGATAGTACAC' *.fastq > count/B3729_Fm0.txt
agrep -0 -c 'AAGGTCAGTGCAAACGG' *.fastq > count/B3730_Fm0.txt
agrep -0 -c 'CCTCTTTAGACGCTTCA' *.fastq > count/B3732_Fm0.txt
agrep -0 -c 'AGAAAATCTGGTCTTCC' *.fastq > count/B3733_Fm0.txt
agrep -0 -c 'CGAAACGCATTGTTAGA' *.fastq > count/B3734_Fm0.txt
agrep -0 -c 'TGGCTGGGCCAAGTTTC' *.fastq > count/B3735_Fm0.txt
agrep -0 -c 'TGCGGCAACGAATACGA' *.fastq > count/B3736_Fm0.txt
agrep -0 -c 'GTCTGACTGGCGGAATC' *.fastq > count/B3737_Fm0.txt
agrep -0 -c 'ACCGAGCCACGTTAACA' *.fastq > count/B3738_Fm0.txt
agrep -0 -c 'CTGCCACAAAGGATTCT' *.fastq > count/B3739_Fm0.txt
agrep -0 -c 'TTAGCCAGCACAAGTCC' *.fastq > count/B3740_Fm0.txt
agrep -0 -c 'CGATAGCATACTCTAAC' *.fastq > count/B3741_Fm0.txt
agrep -0 -c 'TTGCACGAGCTTGTCCA' *.fastq > count/B3742_Fm0.txt
agrep -0 -c 'GGCGGGAACTTGAAAGG' *.fastq > count/B3744_Fm0.txt
agrep -0 -c 'GGTATATCCATATTGCG' *.fastq > count/B3745_Fm0.txt
agrep -0 -c 'CATAGCGGTAGTCCGAA' *.fastq > count/B3746_Fm0.txt
agrep -0 -c 'GTTTGTCTATACATGCG' *.fastq > count/B3749_Fm0.txt
agrep -0 -c 'TATGGGTTAGTCCAAGT' *.fastq > count/B3750_Fm0.txt
agrep -0 -c 'TTCGTACACGTAATGGC' *.fastq > count/B3751_Fm0.txt
agrep -0 -c 'GCGTCTCGCTCTACTGA' *.fastq > count/B3752_Fm0.txt
agrep -0 -c 'ACCGATTTCGGATGGTT' *.fastq > count/B3753_Fm0.txt
agrep -0 -c 'AGCAGATCACGAACTCA' *.fastq > count/B3754_Fm0.txt
agrep -0 -c 'GGGTGGATACTTCTTAG' *.fastq > count/B3755_Fm0.txt
agrep -0 -c 'AGAGTATCGGCTACCTG' *.fastq > count/B3756_Fm0.txt
agrep -0 -c 'TAACCTATAGCCGGCCT' *.fastq > count/B3757_Fm0.txt
agrep -0 -c 'GCCGGGCAAGGTTTAGT' *.fastq > count/B3758_Fm0.txt
agrep -0 -c 'GCCGAAAATCGATTTCG' *.fastq > count/B3759_Fm0.txt
agrep -0 -c 'GACCCCGTTCTAGCTAC' *.fastq > count/B3760_Fm0.txt
agrep -0 -c 'TTGTCTTCGTTATCCAG' *.fastq > count/B3761_Fm0.txt
agrep -0 -c 'TAGAGGTCAGTAAAAGG' *.fastq > count/B3762_Fm0.txt
agrep -0 -c 'TTACTCCCAGAACGGGC' *.fastq > count/B3763_Fm0.txt
agrep -0 -c 'TAACTACCATAGAGGCC' *.fastq > count/B3764_Fm0.txt
agrep -0 -c 'CATTCTACTTCATCGCT' *.fastq > count/B3765_Fm0.txt
agrep -0 -c 'CGGATTCGGTGGAATCG' *.fastq > count/B3766_Fm0.txt
agrep -0 -c 'GTCTAACTACTGGTGTC' *.fastq > count/B3767_Fm0.txt
agrep -0 -c 'ACAGGTCGGGTATTCTT' *.fastq > count/B3768_Fm0.txt
agrep -0 -c 'TGTCCGTATCTACGCTG' *.fastq > count/B3769_Fm0.txt
agrep -0 -c 'ATTCGCCTCCCAAACAG' *.fastq > count/B3770_Fm0.txt
agrep -0 -c 'CACCCTGCTGGACAATG' *.fastq > count/B3771_Fm0.txt
agrep -0 -c 'CGGAGTATTGGGAGTCT' *.fastq > count/B3772_Fm0.txt
agrep -0 -c 'CAAACGTACGGGTTCCC' *.fastq > count/B3773_Fm0.txt
agrep -0 -c 'ATTCTGTTGTCATCGAC' *.fastq > count/B3774_Fm0.txt
agrep -0 -c 'CCTTGGCTTATGCGTTT' *.fastq > count/B3775_Fm0.txt
agrep -0 -c 'CGAATGCTCGACGGTCT' *.fastq > count/B3776_Fm0.txt
agrep -0 -c 'ATATCCGTACAGCTGCG' *.fastq > count/B3777_Fm0.txt
agrep -0 -c 'TATAGGGCCAGAGACGA' *.fastq > count/B3778_Fm0.txt
agrep -0 -c 'CAGTATTGCTCCTCGCC' *.fastq > count/B3779_Fm0.txt
agrep -0 -c 'CGCTAGTGACTTCTTCT' *.fastq > count/B3780_Fm0.txt
agrep -0 -c 'TTCTCTGAGGGATAGAA' *.fastq > count/B3781_Fm0.txt
agrep -0 -c 'ATATGGCATGTCCGATG' *.fastq > count/B3782_Fm0.txt
agrep -0 -c 'TCGATACCATGATGTGC' *.fastq > count/B3784_Fm0.txt
agrep -0 -c 'GTACATACCTACGGCCA' *.fastq > count/B3785_Fm0.txt
agrep -0 -c 'GGGGACTAATACTCAGT' *.fastq > count/B3786_Fm0.txt
agrep -0 -c 'CTGGTTTTGGGTCACGG' *.fastq > count/B3788_Fm0.txt
agrep -0 -c 'GGGACTGACGTACCTCA' *.fastq > count/B3789_Fm0.txt
agrep -0 -c 'CCTAACACCCGGGGAAT' *.fastq > count/B3790_Fm0.txt
agrep -0 -c 'GTTTCGCGCACGACTAC' *.fastq > count/B3791_Fm0.txt
agrep -0 -c 'TAGTCAACCTATCTTGC' *.fastq > count/B3792_Fm0.txt
agrep -0 -c 'TAGCATCATGCCTGCCG' *.fastq > count/B3794_Fm0.txt
agrep -0 -c 'CGGTTTCAGTATCACTT' *.fastq > count/B3795_Fm0.txt
agrep -0 -c 'AGCGCGATCGACATAGC' *.fastq > count/B3797_Fm0.txt
agrep -0 -c 'GTAAATTGCTTACAGGC' *.fastq > count/B3798_Fm0.txt
agrep -0 -c 'CATATTCCCACAAGAGT' *.fastq > count/B3799_Fm0.txt
agrep -0 -c 'GGTAAGCCGCATGCACT' *.fastq > count/B3800_Fm0.txt
agrep -0 -c 'CCGGTGCACTAGTAGAA' *.fastq > count/B3801_Fm0.txt
agrep -0 -c 'TATGAGGGCCAGTACGT' *.fastq > count/B3802_Fm0.txt
agrep -0 -c 'AGACCGAACTTACAGCG' *.fastq > count/B3803_Fm0.txt
agrep -0 -c 'CCGGACTTCTACTCCCA' *.fastq > count/B3804_Fm0.txt
agrep -0 -c 'TAACAACGAGTGGACAA' *.fastq > count/B3805_Fm0.txt
agrep -0 -c 'GTAAGCTTCCACCTACT' *.fastq > count/B3806_Fm0.txt
agrep -0 -c 'GTGCCCTCGCATTAATG' *.fastq > count/B3808_Fm0.txt
agrep -0 -c 'GGGACCGCGTCTATATT' *.fastq > count/B3810_Fm0.txt
agrep -0 -c 'AATTCATCAGCCCATTG' *.fastq > count/B3811_Fm0.txt
agrep -0 -c 'TCGGCATTGAAGACATC' *.fastq > count/B3813_Fm0.txt
agrep -0 -c 'TAATCCAAGGACCCCAT' *.fastq > count/B3814_Fm0.txt
agrep -0 -c 'CGTTTGAAATTCTTGGG' *.fastq > count/B3815_Fm0.txt
agrep -0 -c 'TGCGAGTTACCTGAACT' *.fastq > count/B3816_Fm0.txt
agrep -0 -c 'AGCCTCCATATTGTATG' *.fastq > count/B3817_Fm0.txt
agrep -0 -c 'TTCTAATGAGCGGCGAC' *.fastq > count/B3818_Fm0.txt
agrep -0 -c 'GACAAACGACGCTATGT' *.fastq > count/B3819_Fm0.txt
agrep -0 -c 'AAGGGGTTCCCTGAATA' *.fastq > count/B3820_Fm0.txt
agrep -0 -c 'ACGGACCACGGTTGCAT' *.fastq > count/B3821_Fm0.txt
agrep -0 -c 'GCAAGGGTTATCACTAT' *.fastq > count/B3825_Fm0.txt
agrep -0 -c 'GACGAGCTCAACACACC' *.fastq > count/B3826_Fm0.txt
agrep -0 -c 'GTGGTAAGTCCGGTAGG' *.fastq > count/B3827_Fm0.txt
agrep -0 -c 'CACTTCTTCGAGGTGAT' *.fastq > count/B3828_Fm0.txt
agrep -0 -c 'GTGAGACGCATTCCAGA' *.fastq > count/B3829_Fm0.txt
agrep -0 -c 'GGGCTCAGACCTTTTGT' *.fastq > count/B3830_Fm0.txt
agrep -0 -c 'CGCTCGTTATGTTGTAT' *.fastq > count/B3831_Fm0.txt
agrep -0 -c 'TCAGTGTCTTACAGCGA' *.fastq > count/B3832_Fm0.txt
agrep -0 -c 'CAGAAGCTTGTCGCGAT' *.fastq > count/B3833_Fm0.txt
agrep -0 -c 'CCTCACCGGTTCACATG' *.fastq > count/B3834_Fm0.txt
agrep -0 -c 'GGTCTTCAGATCCACCG' *.fastq > count/B3835_Fm0.txt
agrep -0 -c 'AGCTGCTGGATCTAGCT' *.fastq > count/B3836_Fm0.txt
agrep -0 -c 'ACAACTTTACACAGGAG' *.fastq > count/B3837_Fm0.txt
agrep -0 -c 'TTCGCGTCTTAGCAAGC' *.fastq > count/B3838_Fm0.txt
agrep -0 -c 'AACAGAGCTGCGGAAGC' *.fastq > count/B3839_Fm0.txt
agrep -0 -c 'TACACGATGATGTGCGC' *.fastq > count/B3840_Fm0.txt
agrep -0 -c 'TCGACTACAAAGGGTCA' *.fastq > count/B3841_Fm0.txt
agrep -0 -c 'ATGAAGCAGAAATGGTC' *.fastq > count/B3842_Fm0.txt
agrep -0 -c 'AACCTAGGCGTCCAGAT' *.fastq > count/B3843_Fm0.txt
agrep -0 -c 'GAGGCCCCCAGAGATTT' *.fastq > count/B3845_Fm0.txt
agrep -0 -c 'ACGTAACCGGTTCAGAT' *.fastq > count/B3846_Fm0.txt
agrep -0 -c 'CATTATGCTCTGCACAG' *.fastq > count/B3848_Fm0.txt
agrep -0 -c 'AATAGGTGCGGCGAATA' *.fastq > count/B3849_Fm0.txt
agrep -0 -c 'CTGTGAATGTTCCTCGT' *.fastq > count/B3850_Fm0.txt
agrep -0 -c 'ACGGTACCGGGTTGTTG' *.fastq > count/B3851_Fm0.txt
agrep -0 -c 'TGGCAAAGACGAGGAAC' *.fastq > count/B3852_Fm0.txt
agrep -0 -c 'CTTCAATTTGAGCCTGG' *.fastq > count/B3853_Fm0.txt
agrep -0 -c 'GGGAGGCTTACCGTATA' *.fastq > count/B3854_Fm0.txt
agrep -0 -c 'GGTACCCGGATCGATGT' *.fastq > count/B3855_Fm0.txt
agrep -0 -c 'CTCTAGTCCGAGTATGT' *.fastq > count/B3858_Fm0.txt
agrep -0 -c 'CGGTGCGAATTACCAAG' *.fastq > count/B3859_Fm0.txt
agrep -0 -c 'TGTCGAACACGCAGCTG' *.fastq > count/B3860_Fm0.txt
agrep -0 -c 'AGACATACGGAAGGGGC' *.fastq > count/B3862_Fm0.txt
agrep -0 -c 'TGATGCCGAGAGGGTAT' *.fastq > count/B3865_Fm0.txt
agrep -0 -c 'GTAGCTGTTCTAGGATT' *.fastq > count/B3866_Fm0.txt
agrep -0 -c 'GTAGGAGCTGTCTTAAA' *.fastq > count/B3867_Fm0.txt
agrep -0 -c 'GATCACATCCCAACTGT' *.fastq > count/B3868_Fm0.txt
agrep -0 -c 'GGGAAGCTCTCTGCGAA' *.fastq > count/B3869_Fm0.txt
agrep -0 -c 'ATACAGCTGTGCGCAAT' *.fastq > count/B3870_Fm0.txt
agrep -0 -c 'GTCTAAGCCTTATATCG' *.fastq > count/B3871_Fm0.txt
agrep -0 -c 'GGACCGTAATTCGGACC' *.fastq > count/B3872_Fm0.txt
agrep -0 -c 'GCGTATACCTTAACACC' *.fastq > count/B3873_Fm0.txt
agrep -0 -c 'TTAACATGACTTCTCGG' *.fastq > count/B3874_Fm0.txt
agrep -0 -c 'CCGGGAGCAAAAGGAAT' *.fastq > count/B3875_Fm0.txt
agrep -0 -c 'ACTCACAAGGACTGACA' *.fastq > count/B3876_Fm0.txt
agrep -0 -c 'TAGGTAAGCCGGGTAAA' *.fastq > count/B3877_Fm0.txt
agrep -0 -c 'TTAGGGCGCATTTTTCA' *.fastq > count/B3878_Fm0.txt
agrep -0 -c 'GGCACATCCCTACTTGG' *.fastq > count/B3879_Fm0.txt
agrep -0 -c 'CGATCCCCTCGTTAGGA' *.fastq > count/B3880_Fm0.txt
agrep -0 -c 'TCTGCAGCGTGAGTCTA' *.fastq > count/B3882_Fm0.txt
agrep -0 -c 'CCGTGGTTTTAGTCCCA' *.fastq > count/B3883_Fm0.txt
agrep -0 -c 'CTCTTCAAATGGCGCGA' *.fastq > count/B3884_Fm0.txt
agrep -0 -c 'TCTAGCCCAGACAGTTC' *.fastq > count/B3885_Fm0.txt
agrep -0 -c 'CAGGAGAGTCTTACTCC' *.fastq > count/B3886_Fm0.txt
agrep -0 -c 'TGACGTCCCGGGCTAAT' *.fastq > count/B3888_Fm0.txt
agrep -0 -c 'TCTAACGTGAGTCGCTA' *.fastq > count/B3889_Fm0.txt
agrep -0 -c 'AACTTCCTCCGTGGTAC' *.fastq > count/B3890_Fm0.txt
agrep -0 -c 'ATATACAGAGCGCGCGT' *.fastq > count/B3891_Fm0.txt
agrep -0 -c 'TCGGGGGGTTCAGAATG' *.fastq > count/B3892_Fm0.txt
agrep -0 -c 'GCAGTATGTGTGGGCAC' *.fastq > count/B3894_Fm0.txt
agrep -0 -c 'AATATAGCGGGAGAATC' *.fastq > count/B3895_Fm0.txt
agrep -0 -c 'CTGGCCTCCCATAAACT' *.fastq > count/B3897_Fm0.txt
agrep -0 -c 'GCCGGTATACTCATATC' *.fastq > count/B3898_Fm0.txt
agrep -0 -c 'GCGTTTCAGCCCGATGT' *.fastq > count/B3899_Fm0.txt
agrep -0 -c 'CTTACTGGGCCATATAG' *.fastq > count/B3900_Fm0.txt
agrep -0 -c 'GCGCATAAACGTTCATA' *.fastq > count/B3901_Fm0.txt
agrep -0 -c 'CTTCGACTCAGTTTCTA' *.fastq > count/B3903_Fm0.txt
agrep -0 -c 'TTCCACTGGCTTGGGCA' *.fastq > count/B3905_Fm0.txt
agrep -0 -c 'GTATCTCTCGAGGCAAT' *.fastq > count/B3906_Fm0.txt
agrep -0 -c 'GAGCAAGTTCGCATATC' *.fastq > count/B3907_Fm0.txt
agrep -0 -c 'CTCTGCCATATGAGGCG' *.fastq > count/B3908_Fm0.txt
agrep -0 -c 'CAGGCTCACTCTCATTT' *.fastq > count/B3909_Fm0.txt
agrep -0 -c 'GTAATAGGAAACCGCGA' *.fastq > count/B3910_Fm0.txt
agrep -0 -c 'ACGCAGAATTAACCGCG' *.fastq > count/B3911_Fm0.txt
agrep -0 -c 'CAGCGCCCAGAATAGCT' *.fastq > count/B3913_Fm0.txt
agrep -0 -c 'TCAGTGGACTTACCTGC' *.fastq > count/B3914_Fm0.txt
agrep -0 -c 'AACGGGCGTAATTTGAC' *.fastq > count/B3915_Fm0.txt
agrep -0 -c 'CCGATAAAGACAAAGCC' *.fastq > count/B3916_Fm0.txt
agrep -0 -c 'TCATTGAGGTCGCCTGA' *.fastq > count/B3917_Fm0.txt
agrep -0 -c 'CTCATGTAGGCACAGTC' *.fastq > count/B3918_Fm0.txt
agrep -0 -c 'TCGGAGTTCTACTGGGC' *.fastq > count/B3919_Fm0.txt
agrep -0 -c 'CCTGCGTAAACGCTAAC' *.fastq > count/B3920_Fm0.txt
agrep -0 -c 'ACCCGCTTAAGAGTGGT' *.fastq > count/B3921_Fm0.txt
agrep -0 -c 'CAAATTTGAACCGGGTG' *.fastq > count/B3922_Fm0.txt
agrep -0 -c 'ATAGCGACAAGCGAGGA' *.fastq > count/B3923_Fm0.txt
agrep -0 -c 'CGCCCGATCCTGCATTA' *.fastq > count/B3924_Fm0.txt
agrep -0 -c 'GAGTACTGGAGTACGGA' *.fastq > count/B3925_Fm0.txt
agrep -0 -c 'ATGAAGGGTGGTCCGTA' *.fastq > count/B3926_Fm0.txt
agrep -0 -c 'TGTGGCAATAGATGTTC' *.fastq > count/B3927_Fm0.txt
agrep -0 -c 'CCAAAGGGTTATGACCG' *.fastq > count/B3928_Fm0.txt
agrep -0 -c 'TGTAGGGTAGTCGCCCA' *.fastq > count/B3929_Fm0.txt
agrep -0 -c 'AAGGGGGCCTGAAAAGG' *.fastq > count/B3930_Fm0.txt
agrep -0 -c 'ACTTATTTTACCGCTCC' *.fastq > count/B3931_Fm0.txt
agrep -0 -c 'GCTTAAACTCGATTCCG' *.fastq > count/B3932_Fm0.txt
agrep -0 -c 'TCGCTCCAAGAAAGGTA' *.fastq > count/B3933_Fm0.txt
agrep -0 -c 'ACAGGGGCTTACTACCC' *.fastq > count/B3935_Fm0.txt
agrep -0 -c 'GCGTAAAGTTGGACGAT' *.fastq > count/B3936_Fm0.txt
agrep -0 -c 'GACCAGTTCGTCCGATT' *.fastq > count/B3937_Fm0.txt
agrep -0 -c 'ATTTTCGGAGCGTCGCG' *.fastq > count/B3938_Fm0.txt
agrep -0 -c 'GGTACGACAGAGGATTA' *.fastq > count/B3940_Fm0.txt
agrep -0 -c 'GTTTGGGCAGGATCGAG' *.fastq > count/B3941_Fm0.txt
agrep -0 -c 'TCAAGCTTGCTGCCTCG' *.fastq > count/B3942_Fm0.txt
agrep -0 -c 'GTTAAATTTGCACTGGC' *.fastq > count/B3943_Fm0.txt
agrep -0 -c 'TGTAAACCTCGGGAGGA' *.fastq > count/B3944_Fm0.txt
agrep -0 -c 'CGAAACATGTACGTGCC' *.fastq > count/B3945_Fm0.txt
agrep -0 -c 'TCATCGCAAAAATAGGG' *.fastq > count/B3946_Fm0.txt
agrep -0 -c 'GAGGGTTGTCTTGCGCA' *.fastq > count/B3947_Fm0.txt
agrep -0 -c 'ACCGCTAGCGTGTACAT' *.fastq > count/B3948_Fm0.txt
agrep -0 -c 'TTAGGCGTGCATGCGAA' *.fastq > count/B3949_Fm0.txt
agrep -0 -c 'CTGCTATCACTATGAGG' *.fastq > count/B3950_Fm0.txt
agrep -0 -c 'TTGCTGGATACGGTCTC' *.fastq > count/B3951_Fm0.txt
agrep -0 -c 'TCTGGACGTACAGAATA' *.fastq > count/B3952_Fm0.txt
agrep -0 -c 'CTAGGACCACTACAGTA' *.fastq > count/B3953_Fm0.txt
agrep -0 -c 'TCTTCGAAGGCACCGAT' *.fastq > count/B3954_Fm0.txt
agrep -0 -c 'GTATGGCGTCAAGTTAT' *.fastq > count/B3955_Fm0.txt
agrep -0 -c 'GGGGCAATGAATTAAAC' *.fastq > count/B3956_Fm0.txt
agrep -0 -c 'GCAACTGCTTACTGTGC' *.fastq > count/B3957_Fm0.txt
agrep -0 -c 'GTCCACATGTACACGAA' *.fastq > count/B3958_Fm0.txt
agrep -0 -c 'ATGCGTGTCCAGATGCT' *.fastq > count/B3962_Fm0.txt
agrep -0 -c 'AAGAAGTGCGGCTCATC' *.fastq > count/B3963_Fm0.txt
agrep -0 -c 'GGGTGAGCAAACGGTGT' *.fastq > count/B3964_Fm0.txt
agrep -0 -c 'GATGGACCATGCACGTC' *.fastq > count/B3965_Fm0.txt
agrep -0 -c 'TACGCTACGATCCTCAG' *.fastq > count/B3969_Fm0.txt
agrep -0 -c 'GTCAGCTCAAGAAGAAT' *.fastq > count/B3970_Fm0.txt
agrep -0 -c 'CGACGATTTAACCGCTC' *.fastq > count/B3972_Fm0.txt
agrep -0 -c 'GACGGCTAGTATGGATA' *.fastq > count/B3974_Fm0.txt
agrep -0 -c 'AATAACGAAATGCCGGC' *.fastq > count/B3975_Fm0.txt
agrep -0 -c 'ATTCGGGTAAACCTCAG' *.fastq > count/B3976_Fm0.txt
agrep -0 -c 'CCAGCGCCAGACTATTG' *.fastq > count/B3977_Fm0.txt
agrep -0 -c 'CGCAATAAGACATACGT' *.fastq > count/B3979_Fm0.txt
agrep -0 -c 'CATAAGGAACGGCTCAG' *.fastq > count/B3981_Fm0.txt
agrep -0 -c 'GTCTGGCGTATAACTGT' *.fastq > count/B3983_Fm0.txt
agrep -0 -c 'ACATGCTAGTTATGCGG' *.fastq > count/B3985_Fm0.txt
agrep -0 -c 'GGGTTTCCGCTGCAAAC' *.fastq > count/B3986_Fm0.txt
agrep -0 -c 'GTACAAGTCACCAGCCA' *.fastq > count/B3987_Fm0.txt
agrep -0 -c 'GTTCAACCGGGGAGATT' *.fastq > count/B3988_Fm0.txt
agrep -0 -c 'CGGGGATTTGCACTTTC' *.fastq > count/B3989_Fm0.txt
agrep -0 -c 'TCGCATAACCTGGGGTT' *.fastq > count/B3991_Fm0.txt
agrep -0 -c 'ATGAACACACACGCCAT' *.fastq > count/B3992_Fm0.txt
agrep -0 -c 'GCGCAGGTCAAACTCTT' *.fastq > count/B3993_Fm0.txt
agrep -0 -c 'CTTATGAGGACGATAGC' *.fastq > count/B3994_Fm0.txt
agrep -0 -c 'GCACTTGCATACATTGG' *.fastq > count/B3995_Fm0.txt
agrep -0 -c 'TAGCTTGGAGTGCCCCT' *.fastq > count/B3996_Fm0.txt
agrep -0 -c 'ACAAGGAGCATAAGCTG' *.fastq > count/B3997_Fm0.txt
agrep -0 -c 'ATGACTGAGTCACCCTA' *.fastq > count/B3998_Fm0.txt
agrep -0 -c 'TGACTTATGCTCGTGAG' *.fastq > count/B3999_Fm0.txt
agrep -0 -c 'CAATCGCAACAAACATC' *.fastq > count/B4000_Fm0.txt
agrep -0 -c 'TGTTGCCGGAGTAAATT' *.fastq > count/B4001_Fm0.txt
agrep -0 -c 'TACTTGGCGAAAGTGAT' *.fastq > count/B4002_Fm0.txt
agrep -0 -c 'GTGGCCCTTAAGAGGAA' *.fastq > count/B4003_Fm0.txt
agrep -0 -c 'ATGGACCAGTATCCCAC' *.fastq > count/B4004_Fm0.txt
agrep -0 -c 'GCGTGCTAGCGGATTGT' *.fastq > count/B4006_Fm0.txt
agrep -0 -c 'GGTACGCTGTAGACCTG' *.fastq > count/B4007_Fm0.txt
agrep -0 -c 'CCATAGGCTGCCCAATT' *.fastq > count/B4008_Fm0.txt
agrep -0 -c 'TCTATCGATTGCTGCCA' *.fastq > count/B4009_Fm0.txt
agrep -0 -c 'GGTCCAATCATTGGCAA' *.fastq > count/B4010_Fm0.txt
agrep -0 -c 'CGTTCGTTCAAGCAAGT' *.fastq > count/B4011_Fm0.txt
agrep -0 -c 'ATGGTTGAACGGAATTC' *.fastq > count/B4013_Fm0.txt
agrep -0 -c 'AGACTACACTGCAAACT' *.fastq > count/B4014_Fm0.txt
agrep -0 -c 'GTACACCCCAACACAAC' *.fastq > count/B4016_Fm0.txt
agrep -0 -c 'ACTGCAGCGTCTAATTG' *.fastq > count/B4017_Fm0.txt
agrep -0 -c 'CAGATTGTGGATCCCGG' *.fastq > count/B4018_Fm0.txt
agrep -0 -c 'GGGCAACCAACTGTTGA' *.fastq > count/B4019_Fm0.txt
agrep -0 -c 'TATATGTTCGCGGGTGG' *.fastq > count/B4020_Fm0.txt
agrep -0 -c 'TGAGTTGCAGGATCGCG' *.fastq > count/B4022_Fm0.txt
agrep -0 -c 'CCTTGACACGACGTTCC' *.fastq > count/B4023_Fm0.txt
agrep -0 -c 'GGGACATTACCTGTTGG' *.fastq > count/B4024_Fm0.txt
agrep -0 -c 'GTTAGATTGGCGTAAGG' *.fastq > count/B4025_Fm0.txt
agrep -0 -c 'GTTCTACAACCTTCGAA' *.fastq > count/B4027_Fm0.txt
agrep -0 -c 'ACCATCTGCAAGCTGCG' *.fastq > count/B4028_Fm0.txt
agrep -0 -c 'GCCGCATGGTAAATAGT' *.fastq > count/B4029_Fm0.txt
agrep -0 -c 'TGTAGGCGATACCCTTT' *.fastq > count/B4030_Fm0.txt
agrep -0 -c 'AGTAAAACCAAAACCGC' *.fastq > count/B4031_Fm0.txt
agrep -0 -c 'GTCGAGATTCCTGCAGA' *.fastq > count/B4032_Fm0.txt
agrep -0 -c 'TTGAAGGTACAGCAGAC' *.fastq > count/B4034_Fm0.txt
agrep -0 -c 'CCGATCATCGCCTATTC' *.fastq > count/B4035_Fm0.txt
agrep -0 -c 'ATTGGCACAGCCCTTCC' *.fastq > count/B4036_Fm0.txt
agrep -0 -c 'AACGCAATGGTCTAGCC' *.fastq > count/B4037_Fm0.txt
agrep -0 -c 'AATTGTGACTAGAGGAC' *.fastq > count/B4039_Fm0.txt
agrep -0 -c 'CCTTAACCACACCGAAA' *.fastq > count/B4040_Fm0.txt
agrep -0 -c 'ATTATGATGCGGGGGCC' *.fastq > count/B4041_Fm0.txt
agrep -0 -c 'TTGCGCTTGGTGAGCAC' *.fastq > count/B4042_Fm0.txt
agrep -0 -c 'GTGTTCCTTGCGTTGGT' *.fastq > count/B4043_Fm0.txt
agrep -0 -c 'TATCCAGCTCAAGATCT' *.fastq > count/B4046_Fm0.txt
agrep -0 -c 'TCGAACGTCCAGTAGGG' *.fastq > count/B4047_Fm0.txt
agrep -0 -c 'ACTCTTTCGTGGTACTT' *.fastq > count/B4048_Fm0.txt
agrep -0 -c 'TCGGTCGACAATCGCGA' *.fastq > count/B4049_Fm0.txt
agrep -0 -c 'ATCGCAGTGGGTCCTAA' *.fastq > count/B4051_Fm0.txt
agrep -0 -c 'TCTCTTGACCTGCGAGA' *.fastq > count/B4052_Fm0.txt
agrep -0 -c 'GGACCCCAGACATCAAT' *.fastq > count/B4053_Fm0.txt
agrep -0 -c 'CGCTAAATGTTCGAGAA' *.fastq > count/B4054_Fm0.txt
agrep -0 -c 'CATATGTCAGTGAGCCA' *.fastq > count/B4055_Fm0.txt
agrep -0 -c 'GTTCGCACTTACTTCCA' *.fastq > count/B4056_Fm0.txt
agrep -0 -c 'CCAATTGATTGTACCGT' *.fastq > count/B4057_Fm0.txt
agrep -0 -c 'CCAATGGGTCCCTTCGA' *.fastq > count/B4058_Fm0.txt
agrep -0 -c 'ATAGGTGACATCCGCCC' *.fastq > count/B4059_Fm0.txt
agrep -0 -c 'GTTGAACGGTTCTGAAT' *.fastq > count/B4060_Fm0.txt
agrep -0 -c 'AAGTCCATGCTCCCAAG' *.fastq > count/B4061_Fm0.txt
agrep -0 -c 'CACCAGTTGATCTCCCC' *.fastq > count/B4063_Fm0.txt
agrep -0 -c 'GGCCACAATGAGTAATC' *.fastq > count/B4066_Fm0.txt
agrep -0 -c 'ACTCAGAAACTAAGGAG' *.fastq > count/B4067_Fm0.txt
agrep -0 -c 'TGAAATGACACCCCAGC' *.fastq > count/B4068_Fm0.txt
agrep -0 -c 'CAGTGCCGAACCAATGA' *.fastq > count/B4069_Fm0.txt
agrep -0 -c 'TACCCATGTATGCCTCC' *.fastq > count/B4071_Fm0.txt
agrep -0 -c 'GCACAGAATAGAAGGGA' *.fastq > count/B4072_Fm0.txt
agrep -0 -c 'TGCCGTCGAATCATCAA' *.fastq > count/B4074_Fm0.txt
agrep -0 -c 'TGCAAAGTACGTGAAGA' *.fastq > count/B4075_Fm0.txt
agrep -0 -c 'CCAACCTTTGATTGTGG' *.fastq > count/B4076_Fm0.txt
agrep -0 -c 'AACCTTGCTCCTACTTG' *.fastq > count/B4077_Fm0.txt
agrep -0 -c 'CATGGTGGCGTTCTGTT' *.fastq > count/B4078_Fm0.txt
agrep -0 -c 'CCGCGACTTTCTACCAG' *.fastq > count/B4080_Fm0.txt
agrep -0 -c 'AATGCTCCCCAAATCGT' *.fastq > count/B4081_Fm0.txt
agrep -0 -c 'CACCTTGTTTACCCAGA' *.fastq > count/B4082_Fm0.txt
agrep -0 -c 'AGCCTAATGGTAAACCA' *.fastq > count/B4083_Fm0.txt
agrep -0 -c 'GACATATTTCGGTGGCC' *.fastq > count/B4084_Fm0.txt
agrep -0 -c 'GAGCCACCTAATGGGAA' *.fastq > count/B4085_Fm0.txt
agrep -0 -c 'AGACAGTTCTGACCACT' *.fastq > count/B4086_Fm0.txt
agrep -0 -c 'GCACTTAAGGCTGACCA' *.fastq > count/B4088_Fm0.txt
agrep -0 -c 'TTACAACTTACCTGCCA' *.fastq > count/B4089_Fm0.txt
agrep -0 -c 'GCGAGCCATCTTATCCC' *.fastq > count/B4090_Fm0.txt
agrep -0 -c 'TTGCAAAAATACCGCGG' *.fastq > count/B4091_Fm0.txt
agrep -0 -c 'TTGCTCGCGTTTAATGG' *.fastq > count/B4092_Fm0.txt
agrep -0 -c 'CCGTTCGCTCTAAGTGA' *.fastq > count/B4093_Fm0.txt
agrep -0 -c 'TGCAGCCTGACTCGAAA' *.fastq > count/B4094_Fm0.txt
agrep -0 -c 'GTGTTCACTCCACATAC' *.fastq > count/B4095_Fm0.txt
agrep -0 -c 'GTAAGATGACCTAGCTG' *.fastq > count/B4096_Fm0.txt
agrep -0 -c 'AGAAAATCGGTGCAAGC' *.fastq > count/B4097_Fm0.txt
agrep -0 -c 'GACCTTTATCGGGTATT' *.fastq > count/B4098_Fm0.txt
agrep -0 -c 'GTCTAAAGACCATGAGT' *.fastq > count/B4099_Fm0.txt
agrep -0 -c 'CCGAGCATACGCTTGCT' *.fastq > count/B4100_Fm0.txt
agrep -0 -c 'ATACAATCCAGCGCTTC' *.fastq > count/B4102_Fm0.txt
agrep -0 -c 'GGACACTCAGACAGCTC' *.fastq > count/B4103_Fm0.txt
agrep -0 -c 'TGTAAGGGGTTAAGTTG' *.fastq > count/B4104_Fm0.txt
agrep -0 -c 'AGATCACTCCGGAGTTT' *.fastq > count/B4105_Fm0.txt
agrep -0 -c 'TGACCCCTCTTATAGAC' *.fastq > count/B4106_Fm0.txt
agrep -0 -c 'GTCCACGAGTGGATTCC' *.fastq > count/B4107_Fm0.txt
agrep -0 -c 'AGTTCCGGACGAACTGT' *.fastq > count/B4108_Fm0.txt
agrep -0 -c 'AACGCACTTATTCGTAG' *.fastq > count/B4109_Fm0.txt
agrep -0 -c 'CTATCTCCGTACTGCGT' *.fastq > count/B4110_Fm0.txt
agrep -0 -c 'GCTAGTTCCTTCTCACC' *.fastq > count/B4111_Fm0.txt
agrep -0 -c 'GCAAGCAGCTCATTTCA' *.fastq > count/B4112_Fm0.txt
agrep -0 -c 'CGATTAACCCAAGAGAG' *.fastq > count/B4114_Fm0.txt
agrep -0 -c 'TACGTATCTCGCACAGT' *.fastq > count/B4115_Fm0.txt
agrep -0 -c 'GTATAGGATCTACATGG' *.fastq > count/B4117_Fm0.txt
agrep -0 -c 'GTCACGGACCTCTTTAC' *.fastq > count/B4118_Fm0.txt
agrep -0 -c 'GCTGTGAGGTGGAAACG' *.fastq > count/B4119_Fm0.txt
agrep -0 -c 'CGACCACCTGAATCCGA' *.fastq > count/B4120_Fm0.txt
agrep -0 -c 'ACGCGCTGTTCTCTTTC' *.fastq > count/B4121_Fm0.txt
agrep -0 -c 'GCGTTCATAGCTCAGAA' *.fastq > count/B4122_Fm0.txt
agrep -0 -c 'GTAGGTCAGAGTAGTCA' *.fastq > count/B4123_Fm0.txt
agrep -0 -c 'GTAGGTGTAACGACTCT' *.fastq > count/B4124_Fm0.txt
agrep -0 -c 'CCGAACAGTAGACAGAA' *.fastq > count/B4125_Fm0.txt
agrep -0 -c 'ACCGAAATTAGTAGGTC' *.fastq > count/B4126_Fm0.txt
agrep -0 -c 'ATACCACAGACGAGAGG' *.fastq > count/B4127_Fm0.txt
agrep -0 -c 'CATTGCGTCTACCTGCT' *.fastq > count/B4128_Fm0.txt
agrep -0 -c 'GCATTCGTCCCTCGTGA' *.fastq > count/B4129_Fm0.txt
agrep -0 -c 'CTCCCATACAGGTGGTG' *.fastq > count/B4130_Fm0.txt
agrep -0 -c 'TGACCCTCGCACCGATT' *.fastq > count/B4131_Fm0.txt
agrep -0 -c 'TCATGTGTCTATATCCG' *.fastq > count/B4132_Fm0.txt
agrep -0 -c 'AAGACCGATCCCCAATC' *.fastq > count/B4133_Fm0.txt
agrep -0 -c 'GACGTGTATTGGACATT' *.fastq > count/B4134_Fm0.txt
agrep -0 -c 'GGTTGTAGTCTGGGCTT' *.fastq > count/B4135_Fm0.txt
agrep -0 -c 'TAAACCGTTCACACGCC' *.fastq > count/B4137_Fm0.txt
agrep -0 -c 'GTGAAGGTCCACTCTCA' *.fastq > count/B4139_Fm0.txt
agrep -0 -c 'GCCCAAGCTAGGTTTTT' *.fastq > count/B4140_Fm0.txt
agrep -0 -c 'GGATGAGCATACAGGGC' *.fastq > count/B4141_Fm0.txt
agrep -0 -c 'GCTCTACAAGATCACAA' *.fastq > count/B4142_Fm0.txt
agrep -0 -c 'TTGGATTACAGCTCCAG' *.fastq > count/B4144_Fm0.txt
agrep -0 -c 'TGTTCCGCAGCTGACTA' *.fastq > count/B4145_Fm0.txt
agrep -0 -c 'ATAGAGTACAGTCTGTG' *.fastq > count/B4146_Fm0.txt
agrep -0 -c 'CGGTTGCATAGTGTCTG' *.fastq > count/B4147_Fm0.txt
agrep -0 -c 'TTCACTCATAGCCCCAC' *.fastq > count/B4148_Fm0.txt
agrep -0 -c 'TTAATCACCTGGTAGTG' *.fastq > count/B4149_Fm0.txt
agrep -0 -c 'CATGGCGGGATGTTTAG' *.fastq > count/B4150_Fm0.txt
agrep -0 -c 'AGCAGGAGTTACCGGGT' *.fastq > count/B4151_Fm0.txt
agrep -0 -c 'ACTCGTCAGGCTTAGGT' *.fastq > count/B4152_Fm0.txt
agrep -0 -c 'CCACCGTCAACGCATCA' *.fastq > count/B4153_Fm0.txt
agrep -0 -c 'AGTCAGGTTAATTAGCC' *.fastq > count/B4154_Fm0.txt
agrep -0 -c 'GCGGTTCCTCATATTCA' *.fastq > count/B4155_Fm0.txt
agrep -0 -c 'GGTGATCATGGCTGTTT' *.fastq > count/B4156_Fm0.txt
agrep -0 -c 'ATTTACCCGCACCGCCA' *.fastq > count/B4157_Fm0.txt
agrep -0 -c 'TATTTGGAGTCCGCCAT' *.fastq > count/B4158_Fm0.txt
agrep -0 -c 'GGCAACGCCATATCAAA' *.fastq > count/B4159_Fm0.txt
agrep -0 -c 'TGCCATTGATCTCACAT' *.fastq > count/B4160_Fm0.txt
agrep -0 -c 'TAGAACTTAACGACGGG' *.fastq > count/B4164_Fm0.txt
agrep -0 -c 'CCTCCCTTGGTTATACC' *.fastq > count/B4165_Fm0.txt
agrep -0 -c 'ATTAGCACTATCTGCCT' *.fastq > count/B4166_Fm0.txt
agrep -0 -c 'CGGCTATTAGGTCAAGC' *.fastq > count/B4167_Fm0.txt
agrep -0 -c 'GGTATTTCCTTACATGC' *.fastq > count/B4168_Fm0.txt
agrep -0 -c 'AACACTATTGCTCCCTA' *.fastq > count/B4169_Fm0.txt
agrep -0 -c 'GGGAGACGTGAAGTTTG' *.fastq > count/B4170_Fm0.txt
agrep -0 -c 'CGCAACTCTGGAACGAT' *.fastq > count/B4171_Fm0.txt
agrep -0 -c 'ATCCATTAGGAGCACTT' *.fastq > count/B4172_Fm0.txt
agrep -0 -c 'AGTATGGGACATTACCA' *.fastq > count/B4173_Fm0.txt
agrep -0 -c 'GTTGTGGATCCATACAA' *.fastq > count/B4174_Fm0.txt
agrep -0 -c 'CCACTTCGGCATTAAGG' *.fastq > count/B4175_Fm0.txt
agrep -0 -c 'GCGAAGTGATCGAAGAG' *.fastq > count/B4176_Fm0.txt
agrep -0 -c 'GTCGAGAACTGATCAGT' *.fastq > count/B4177_Fm0.txt
agrep -0 -c 'CAGAGCGCAACTGGGAA' *.fastq > count/B4178_Fm0.txt
agrep -0 -c 'AGCTTCGTCAAGATGAT' *.fastq > count/B4179_Fm0.txt
agrep -0 -c 'AGAACATCACATACCAG' *.fastq > count/B4181_Fm0.txt
agrep -0 -c 'TTGGTTACAGTCCACGT' *.fastq > count/B4182_Fm0.txt
agrep -0 -c 'CACTTCCCTTGGTGCAA' *.fastq > count/B4183_Fm0.txt
agrep -0 -c 'TAGTGGCCCTATAGTAC' *.fastq > count/B4184_Fm0.txt
agrep -0 -c 'GCATCTGGGGCCATATA' *.fastq > count/B4185_Fm0.txt
agrep -0 -c 'CTAATTAACTCCGGTGT' *.fastq > count/B4186_Fm0.txt
agrep -0 -c 'CGCCTCACAAGTATAGG' *.fastq > count/B4187_Fm0.txt
agrep -0 -c 'CCGACAATCACTAGCTA' *.fastq > count/B4188_Fm0.txt
agrep -0 -c 'TGACTATGGTGTTCTTG' *.fastq > count/B4190_Fm0.txt
agrep -0 -c 'TTGGACGGCCTTTCGAC' *.fastq > count/B4191_Fm0.txt
agrep -0 -c 'CGTCAGATTTGCCATTG' *.fastq > count/B4193_Fm0.txt
agrep -0 -c 'GCGGGTCTGGAGATTCA' *.fastq > count/B4194_Fm0.txt
agrep -0 -c 'ACCCGTTATATGTATCG' *.fastq > count/B4195_Fm0.txt
agrep -0 -c 'AACTGGACGAAACGACG' *.fastq > count/B4196_Fm0.txt
agrep -0 -c 'GTATTTCCTGTTCTACG' *.fastq > count/B4197_Fm0.txt
agrep -0 -c 'GGGAATTCGGGTGCACT' *.fastq > count/B4198_Fm0.txt
agrep -0 -c 'GTTAAAGGATCGTTGAC' *.fastq > count/B4199_Fm0.txt
agrep -0 -c 'GTGCTATAAGACTAGTC' *.fastq > count/B4200_Fm0.txt
agrep -0 -c 'CTAGCAGTGTAGCAACT' *.fastq > count/B4201_Fm0.txt
agrep -0 -c 'GCACAATACAACACGAA' *.fastq > count/B4202_Fm0.txt
agrep -0 -c 'CTACTTCATTACCGCCA' *.fastq > count/B4204_Fm0.txt
agrep -0 -c 'TACCGGTCCGGTACTTC' *.fastq > count/B4205_Fm0.txt
agrep -0 -c 'CACTGCATCAGATAGGT' *.fastq > count/B4206_Fm0.txt
agrep -0 -c 'GAGATCTCGGGCGTTAC' *.fastq > count/B4207_Fm0.txt
agrep -0 -c 'CGCTTCCCATACTGGTT' *.fastq > count/B4208_Fm0.txt
agrep -0 -c 'GTAGCATTCACGGGATC' *.fastq > count/B4209_Fm0.txt
agrep -0 -c 'CAACCAGCGACAATTTG' *.fastq > count/B4210_Fm0.txt
agrep -0 -c 'GTGAAGCTAACCAATTC' *.fastq > count/B4213_Fm0.txt
agrep -0 -c 'CGGATTTAAGCATGAAG' *.fastq > count/B4215_Fm0.txt
agrep -0 -c 'AGAAGGACGAACCGCGT' *.fastq > count/B4216_Fm0.txt
agrep -0 -c 'CAGCAACCGACGGTATC' *.fastq > count/B4217_Fm0.txt
agrep -0 -c 'ACCACTCCACGCAACGT' *.fastq > count/B4218_Fm0.txt
agrep -0 -c 'CGTGTAATGTCCGTCGA' *.fastq > count/B4219_Fm0.txt
agrep -0 -c 'GAGTGCTTAGTTTCAGA' *.fastq > count/B4220_Fm0.txt
agrep -0 -c 'GCTGTACGAAGCTAGGC' *.fastq > count/B4221_Fm0.txt
agrep -0 -c 'TTACGCGTATGGCAGAG' *.fastq > count/B4222_Fm0.txt
agrep -0 -c 'CAGCGTGACATTCCTAT' *.fastq > count/B4223_Fm0.txt
agrep -0 -c 'TTGCGCCGAACGCCAAT' *.fastq > count/B4224_Fm0.txt
agrep -0 -c 'ATAGGTGGAAACGGGAT' *.fastq > count/B4225_Fm0.txt
agrep -0 -c 'TTGTGTCAGGTCGCGTG' *.fastq > count/B4227_Fm0.txt
agrep -0 -c 'AGGGTTGGTTAAATCGA' *.fastq > count/B4228_Fm0.txt
agrep -0 -c 'TCTGTTTCCGCGTGTCA' *.fastq > count/B4229_Fm0.txt
agrep -0 -c 'TAGAGACGAAAGTCGGC' *.fastq > count/B4232_Fm0.txt
agrep -0 -c 'CAGGGCCGTTTTTACAG' *.fastq > count/B4234_Fm0.txt
agrep -0 -c 'GTAAGTCCTCAGACAGG' *.fastq > count/B4235_Fm0.txt
agrep -0 -c 'ATTCTGCGCGACCAAGG' *.fastq > count/B4237_Fm0.txt
agrep -0 -c 'GTTCTTTAACCCTCAAG' *.fastq > count/B4238_Fm0.txt
agrep -0 -c 'AGCGTACTCAAAAGTGC' *.fastq > count/B4239_Fm0.txt
agrep -0 -c 'ACACTGGATCGTAAAAC' *.fastq > count/B4240_Fm0.txt
agrep -0 -c 'TAATCTTCCGCTTAGAG' *.fastq > count/B4241_Fm0.txt
agrep -0 -c 'GTAATTACAGCGTCATC' *.fastq > count/B4243_Fm0.txt
agrep -0 -c 'CACAATATACCCGCATG' *.fastq > count/B4244_Fm0.txt
agrep -0 -c 'CTGGTAACACCCTGACT' *.fastq > count/B4245_Fm0.txt
agrep -0 -c 'GCGGAGCCCTCTTAAAT' *.fastq > count/B4246_Fm0.txt
agrep -0 -c 'TAGACATAACCTGGTGT' *.fastq > count/B4248_Fm0.txt
agrep -0 -c 'AGAATGACCATGCAGGC' *.fastq > count/B4250_Fm0.txt
agrep -0 -c 'AACTCTGTCTGAGCTCT' *.fastq > count/B4251_Fm0.txt
agrep -0 -c 'TTGACCGATACGCATAT' *.fastq > count/B4252_Fm0.txt
agrep -0 -c 'GAGCCTCCTATCGCTAC' *.fastq > count/B4253_Fm0.txt
agrep -0 -c 'ATCGCCGTAATAAGCGC' *.fastq > count/B4254_Fm0.txt
agrep -0 -c 'CTCAATCATGGTGTCGA' *.fastq > count/B4255_Fm0.txt
agrep -0 -c 'GGAGATCTTTCCGGCTA' *.fastq > count/B4256_Fm0.txt
agrep -0 -c 'TTCGAGCTCCGGTAAGC' *.fastq > count/B4257_Fm0.txt
agrep -0 -c 'TTAATTCGGTCCGGCTC' *.fastq > count/B4258_Fm0.txt
agrep -0 -c 'GCGGAATTGCGTGGTAC' *.fastq > count/B4259_Fm0.txt
agrep -0 -c 'GCCGCAGGTGAATGGTT' *.fastq > count/B4260_Fm0.txt
agrep -0 -c 'GTGTCTCGTCCATAGAA' *.fastq > count/B4261_Fm0.txt
agrep -0 -c 'ACGTCATCTGGTGTAAG' *.fastq > count/B4262_Fm0.txt
agrep -0 -c 'CTCAGTGTTCGACCTTG' *.fastq > count/B4263_Fm0.txt
agrep -0 -c 'ACTTTACGTAAGGGTGT' *.fastq > count/B4264_Fm0.txt
agrep -0 -c 'CTAGCCGGAAATACGTC' *.fastq > count/B4265_Fm0.txt
agrep -0 -c 'AGCCCAGGTTTATAAGC' *.fastq > count/B4266_Fm0.txt
agrep -0 -c 'CCACTCTTTAGAGGGTC' *.fastq > count/B4267_Fm0.txt
agrep -0 -c 'CCTATCTTGGTCGGTTT' *.fastq > count/B4268_Fm0.txt
agrep -0 -c 'TGCTGATTGCCACACGA' *.fastq > count/B4269_Fm0.txt
agrep -0 -c 'GGAGAATGTACCCGATC' *.fastq > count/B4270_Fm0.txt
agrep -0 -c 'GACACCGGGCAATGATA' *.fastq > count/B4271_Fm0.txt
agrep -0 -c 'ACATAGTAGCAGCGCGA' *.fastq > count/B4272_Fm0.txt
agrep -0 -c 'CGTATGCCGTCAGAGAC' *.fastq > count/B4274_Fm0.txt
agrep -0 -c 'ACAACGCCACACTTCTA' *.fastq > count/B4275_Fm0.txt
agrep -0 -c 'AATGCCGAAGGAGTCCT' *.fastq > count/B4276_Fm0.txt
agrep -0 -c 'CGCGGGTTTGCTTAACC' *.fastq > count/B4277_Fm0.txt
agrep -0 -c 'GTTAAGCCCTAGCAACC' *.fastq > count/B4278_Fm0.txt
agrep -0 -c 'ATGCCTTCATTGCGACG' *.fastq > count/B4279_Fm0.txt
agrep -0 -c 'CGGTTATGGAAACATTC' *.fastq > count/B4280_Fm0.txt
agrep -0 -c 'GCACACGAAGCAGCTTC' *.fastq > count/B4281_Fm0.txt
agrep -0 -c 'CACCAGCCTGTTGTGGT' *.fastq > count/B4282_Fm0.txt
agrep -0 -c 'TTCTCTCGCCTTACTCT' *.fastq > count/B4283_Fm0.txt
agrep -0 -c 'AACGAAGTGGAGAAGGT' *.fastq > count/B4284_Fm0.txt
agrep -0 -c 'TAGATATGACACGTCGG' *.fastq > count/B4285_Fm0.txt
agrep -0 -c 'AACAGTTGGGGTCAAAT' *.fastq > count/B4286_Fm0.txt
agrep -0 -c 'TCTCGGGATGAAAAATG' *.fastq > count/B4287_Fm0.txt
agrep -0 -c 'AGCTAGTGGTCCAGTTC' *.fastq > count/B4288_Fm0.txt
agrep -0 -c 'AGGTTACGATAAAGGTC' *.fastq > count/B4289_Fm0.txt
agrep -0 -c 'CCTTCGGCACCTGAATC' *.fastq > count/B4290_Fm0.txt
agrep -0 -c 'AATATCGTGTCGGTGAT' *.fastq > count/B4292_Fm0.txt
agrep -0 -c 'AGGTCCTTAACTACATC' *.fastq > count/B4295_Fm0.txt
agrep -0 -c 'AACCCGACCAGTCCTCT' *.fastq > count/B4296_Fm0.txt
agrep -0 -c 'TGGCATAACGGAGTCGT' *.fastq > count/B4297_Fm0.txt
agrep -0 -c 'TCTGTAGAGCATGCGTA' *.fastq > count/B4298_Fm0.txt
agrep -0 -c 'TTACGCTCAGCTCGACA' *.fastq > count/B4299_Fm0.txt
agrep -0 -c 'CCATGACAACCTGTGTA' *.fastq > count/B4300_Fm0.txt
agrep -0 -c 'CTCGACGAGTGGGAGAT' *.fastq > count/B4301_Fm0.txt
agrep -0 -c 'ACCGCACGTATATTCGA' *.fastq > count/B4303_Fm0.txt
agrep -0 -c 'GCATCCAGCAAGGTTAG' *.fastq > count/B4304_Fm0.txt
agrep -0 -c 'AATCGAACCTGCCCATG' *.fastq > count/B4305_Fm0.txt
agrep -0 -c 'CTCACCCGATTACAATG' *.fastq > count/B4308_Fm0.txt
agrep -0 -c 'CAGAATTGTTCCTCAGT' *.fastq > count/B4309_Fm0.txt
agrep -0 -c 'AACGAAAGTACGCCATC' *.fastq > count/B4311_Fm0.txt
agrep -0 -c 'TCAACCACGAAGTTAAG' *.fastq > count/B4313_Fm0.txt
agrep -0 -c 'GGAAGGCCGTGACATGA' *.fastq > count/B4314_Fm0.txt
agrep -0 -c 'AATACGCTATGATAGCG' *.fastq > count/B4315_Fm0.txt
agrep -0 -c 'TATAAGTCCGGCGAATC' *.fastq > count/B4316_Fm0.txt
agrep -0 -c 'CGTAGACCCCGGAGTTA' *.fastq > count/B4317_Fm0.txt
agrep -0 -c 'CTTCCGCCTATCTCTCC' *.fastq > count/B4318_Fm0.txt
agrep -0 -c 'ACGAGAAAGTGCTCATG' *.fastq > count/B4319_Fm0.txt
agrep -0 -c 'GGTCGCTATGGACTATC' *.fastq > count/B4320_Fm0.txt
agrep -0 -c 'CATACGCTTAAGGAGCC' *.fastq > count/B4321_Fm0.txt
agrep -0 -c 'CCTTTTCAGAATCGGTA' *.fastq > count/B4322_Fm0.txt
agrep -0 -c 'TCATATTGTGTACCGCG' *.fastq > count/B4323_Fm0.txt
agrep -0 -c 'ACCGCAATTTTCTGGAC' *.fastq > count/B4324_Fm0.txt
agrep -0 -c 'ATAACCCTTCGTTCACT' *.fastq > count/B4325_Fm0.txt
agrep -0 -c 'ATGAAGCACCTTAGGAA' *.fastq > count/B4326_Fm0.txt
agrep -0 -c 'ATTCGAGCCGCTAAACA' *.fastq > count/B4327_Fm0.txt
agrep -0 -c 'CCACGTCTACATAAGTT' *.fastq > count/B4328_Fm0.txt
agrep -0 -c 'AATCCGCCGACAGATCT' *.fastq > count/B4329_Fm0.txt
agrep -0 -c 'TGCGTCGTACATAATGT' *.fastq > count/B4330_Fm0.txt
agrep -0 -c 'GCGACGCTATTGCTTTA' *.fastq > count/B4331_Fm0.txt
agrep -0 -c 'CTCTGTATATCTCGAGC' *.fastq > count/B4332_Fm0.txt
agrep -0 -c 'TCGCATCCGACGTTCCT' *.fastq > count/B4333_Fm0.txt
agrep -0 -c 'AACCCGAATAAGTGAGT' *.fastq > count/B4335_Fm0.txt
agrep -0 -c 'TGAAACTACTCTGCCGG' *.fastq > count/B4336_Fm0.txt
agrep -0 -c 'TGACTCGATACAGATCA' *.fastq > count/B4337_Fm0.txt
agrep -0 -c 'CATATCCTGGTGCGATC' *.fastq > count/B4338_Fm0.txt
agrep -0 -c 'TACTACGGAAGAGCCCC' *.fastq > count/B4340_Fm0.txt
agrep -0 -c 'CCTATCTATTGCGCTGA' *.fastq > count/B4341_Fm0.txt
agrep -0 -c 'AACCGCCGAATTTAGTA' *.fastq > count/B4342_Fm0.txt
agrep -0 -c 'ATTTGACGCATGCAACT' *.fastq > count/B4343_Fm0.txt
agrep -0 -c 'TCACGTCCCGTTCATTG' *.fastq > count/B4344_Fm0.txt
agrep -0 -c 'TATGTCAGCTTGCTAGT' *.fastq > count/B4345_Fm0.txt
agrep -0 -c 'TAAAATATGGTGCCGGC' *.fastq > count/B4346_Fm0.txt
agrep -0 -c 'TTCGACAGCGGACTTGA' *.fastq > count/B4347_Fm0.txt
agrep -0 -c 'CGCAGGGTGCTAATGAG' *.fastq > count/B4348_Fm0.txt
agrep -0 -c 'CGGCTCTTGTTTCAAAC' *.fastq > count/B4350_Fm0.txt
agrep -0 -c 'ACTCGTTAACCCCATCT' *.fastq > count/B4351_Fm0.txt
agrep -0 -c 'TTAGTGAGTACTACGTC' *.fastq > count/B4352_Fm0.txt
agrep -0 -c 'ATGTCGCGATGGTCAAG' *.fastq > count/B4353_Fm0.txt
agrep -0 -c 'AACGGGCCCTGCTACTA' *.fastq > count/B4354_Fm0.txt
agrep -0 -c 'TTCGCAGCTCAAACGGT' *.fastq > count/B4356_Fm0.txt
agrep -0 -c 'CTGTCCCATGCTTAAAA' *.fastq > count/B4357_Fm0.txt
agrep -0 -c 'ATCTACGAACGGGTCGC' *.fastq > count/B4358_Fm0.txt
agrep -0 -c 'CGGAACTTGAGTAGCAT' *.fastq > count/B4359_Fm0.txt
agrep -0 -c 'TTGTGATGAATCGCACG' *.fastq > count/B4360_Fm0.txt
agrep -0 -c 'ATCCACACCGTTGACCC' *.fastq > count/B4362_Fm0.txt
agrep -0 -c 'GATGCGGTCGAACTTAT' *.fastq > count/B4363_Fm0.txt
agrep -0 -c 'ACGACTAGTTTGGCACC' *.fastq > count/B4364_Fm0.txt
agrep -0 -c 'GGGTAGTGCGGATAGAA' *.fastq > count/B4365_Fm0.txt
agrep -0 -c 'CCTGATTAGGCCTCCTT' *.fastq > count/B4366_Fm0.txt
agrep -0 -c 'TTAGACACCCAGATTGC' *.fastq > count/B4367_Fm0.txt
agrep -0 -c 'TAGTACCCTTACGCCCA' *.fastq > count/B4368_Fm0.txt
agrep -0 -c 'GTAATGCTTGGTGTTCC' *.fastq > count/B4369_Fm0.txt
agrep -0 -c 'TAGTATCGAATCTTCCC' *.fastq > count/B4370_Fm0.txt
agrep -0 -c 'AACGCTTTTCGTCCAAT' *.fastq > count/B4372_Fm0.txt
agrep -0 -c 'CCTGCCTCAGGTACAAA' *.fastq > count/B4374_Fm0.txt
agrep -0 -c 'CTAACCACAATTGCGTA' *.fastq > count/B4375_Fm0.txt
agrep -0 -c 'GTGAGTAGCCGTCACAC' *.fastq > count/B4376_Fm0.txt
agrep -0 -c 'ATCCTCGATAGTCTCCG' *.fastq > count/B4377_Fm0.txt
agrep -0 -c 'TCGATTAACCTGAGCCC' *.fastq > count/B4378_Fm0.txt
agrep -0 -c 'GGGGCCTCATCAAGTTT' *.fastq > count/B4379_Fm0.txt
agrep -0 -c 'TCGTTACCGTCATGTCT' *.fastq > count/B4380_Fm0.txt
agrep -0 -c 'CGCGAAAGATCAGCTTG' *.fastq > count/B4381_Fm0.txt
agrep -0 -c 'CAACGGAAGAACCGCTG' *.fastq > count/B4382_Fm0.txt
agrep -0 -c 'CAGCGGGACAACCTTCA' *.fastq > count/B4383_Fm0.txt
agrep -0 -c 'AACCTGCGCTATGCCAG' *.fastq > count/B4384_Fm0.txt
agrep -0 -c 'GTCGCATTTCGCAAAGA' *.fastq > count/B4386_Fm0.txt
agrep -0 -c 'GACCGAGATGTTCTAAA' *.fastq > count/B4387_Fm0.txt
agrep -0 -c 'ATTGAGTCCGTCAGCAG' *.fastq > count/B4388_Fm0.txt
agrep -0 -c 'CACTTTTTATGCTCAGG' *.fastq > count/B4389_Fm0.txt
agrep -0 -c 'GAGTTCAGATTTGGCCG' *.fastq > count/B4390_Fm0.txt
agrep -0 -c 'TCACCGTTGGTGATTTA' *.fastq > count/B4391_Fm0.txt
agrep -0 -c 'TCGGGGCTCTATGTGTA' *.fastq > count/B4393_Fm0.txt
agrep -0 -c 'TATCACCCGCAAGCTAT' *.fastq > count/B4394_Fm0.txt
agrep -0 -c 'GCGTGGCGTACTCTATA' *.fastq > count/B4396_Fm0.txt
agrep -0 -c 'GGCAACGGATGCTAATC' *.fastq > count/B4398_Fm0.txt
agrep -0 -c 'TTACAAGTAGCATCGAG' *.fastq > count/B4399_Fm0.txt
agrep -0 -c 'ACGTCAAGCCCGAATGG' *.fastq > count/B4400_Fm0.txt
agrep -0 -c 'GTTGAACCGCCCAGAAG' *.fastq > count/B4401_Fm0.txt
agrep -0 -c 'TCGTCCGTTCTGCAAAA' *.fastq > count/B4404_Fm0.txt
agrep -0 -c 'AGACCGCTCAAAGCAGG' *.fastq > count/B4405_Fm0.txt
agrep -0 -c 'GCAAGGCTTTGATCACA' *.fastq > count/B4406_Fm0.txt
agrep -0 -c 'GTATGAGCGCATCACCC' *.fastq > count/B4407_Fm0.txt
agrep -0 -c 'TACGTGCGACAGCTATT' *.fastq > count/B4409_Fm0.txt
agrep -0 -c 'GCCCGACACACAACAAA' *.fastq > count/B4411_Fm0.txt
agrep -0 -c 'ATTTGCGCTGGGGTAGC' *.fastq > count/B4412_Fm0.txt
agrep -0 -c 'CATCACCAACGGTATCG' *.fastq > count/B4415_Fm0.txt
agrep -0 -c 'CTCGCATACAATCCAGC' *.fastq > count/B4416_Fm0.txt
agrep -0 -c 'GACATGAGTCATCCGGG' *.fastq > count/B4417_Fm0.txt
agrep -0 -c 'CGGTGGTCATATAGTTT' *.fastq > count/B4418_Fm0.txt
agrep -0 -c 'ATGCCTGTACGCGAATA' *.fastq > count/B4419_Fm0.txt
agrep -0 -c 'CTGGAATGGCTACATCG' *.fastq > count/B4420_Fm0.txt
agrep -0 -c 'GTAGTGCAGGAGGCCTT' *.fastq > count/B4421_Fm0.txt
agrep -0 -c 'AGGCAAGTATCGAATCG' *.fastq > count/B4422_Fm0.txt
agrep -0 -c 'AGGACACAATCACAAGT' *.fastq > count/B4424_Fm0.txt
agrep -0 -c 'TATCATCTGACGTGTCA' *.fastq > count/B4426_Fm0.txt
agrep -0 -c 'CGCCACGTAATGCTGGA' *.fastq > count/B4427_Fm0.txt
agrep -0 -c 'TGACGCTAGTCCGATAC' *.fastq > count/B4429_Fm0.txt
agrep -0 -c 'AATTCGGCGGTCTTTAA' *.fastq > count/B4430_Fm0.txt
agrep -0 -c 'GGCTAATTGCCGAAAAT' *.fastq > count/B4431_Fm0.txt
agrep -0 -c 'TCAAGCCGGGAGCCAAT' *.fastq > count/B4432_Fm0.txt
agrep -0 -c 'GCCTTCTGGGGTTTATC' *.fastq > count/B4433_Fm0.txt
agrep -0 -c 'GTTAGGTCGGTATTGCC' *.fastq > count/B4434_Fm0.txt
agrep -0 -c 'ACGTTTCACCGTAAGTT' *.fastq > count/B4435_Fm0.txt
agrep -0 -c 'AATTGGTTCAGGCTCCG' *.fastq > count/B4437_Fm0.txt
agrep -0 -c 'CCTGTTTGTCCGACCGT' *.fastq > count/B4438_Fm0.txt
agrep -0 -c 'ATCCATTCCTGATAGCC' *.fastq > count/B4440_Fm0.txt
agrep -0 -c 'GTAATCTCCGGTAAAAC' *.fastq > count/B4441_Fm0.txt
agrep -0 -c 'CCAGATGGATATGAGTC' *.fastq > count/B4442_Fm0.txt
agrep -0 -c 'CAAGTAGTCAGTCAACA' *.fastq > count/B4443_Fm0.txt
agrep -0 -c 'AGTATTTTCAACCGCTC' *.fastq > count/B4444_Fm0.txt
agrep -0 -c 'AGGCACTCAAGACTCCC' *.fastq > count/B4445_Fm0.txt
agrep -0 -c 'AGACAATGGTGAAGCAT' *.fastq > count/B4446_Fm0.txt
agrep -0 -c 'ACTGCAATCACGTCTTA' *.fastq > count/B4447_Fm0.txt
agrep -0 -c 'GGTCAACACTATTGACA' *.fastq > count/B4448_Fm0.txt
agrep -0 -c 'GTCGAGCTGAGTACTTC' *.fastq > count/B4449_Fm0.txt
agrep -0 -c 'TGGAACCAAGACCGGAT' *.fastq > count/B4450_Fm0.txt
agrep -0 -c 'GCGTCTTGCTACGTTCT' *.fastq > count/B4452_Fm0.txt
agrep -0 -c 'AAGAACTAACCGTTCCC' *.fastq > count/B4453_Fm0.txt
agrep -0 -c 'CACCCTGTTGGGCTATC' *.fastq > count/B4455_Fm0.txt
agrep -0 -c 'TACTCGCATTGTGTCCT' *.fastq > count/B4456_Fm0.txt
agrep -0 -c 'ATGATCGCCCTTTCGTG' *.fastq > count/B4457_Fm0.txt
agrep -0 -c 'GCGTTGACAAAGTGTTA' *.fastq > count/B4458_Fm0.txt
agrep -0 -c 'ACAAGCGTTGCGGTCAA' *.fastq > count/B4459_Fm0.txt
agrep -0 -c 'CCACCTTTCGTGGTATA' *.fastq > count/B4460_Fm0.txt
agrep -0 -c 'TATGAAAGTCTCGGCGT' *.fastq > count/B4461_Fm0.txt
agrep -0 -c 'TCAGAGGGATACGATGG' *.fastq > count/B4462_Fm0.txt
agrep -0 -c 'ACAGGTGTACCAGTACG' *.fastq > count/B4463_Fm0.txt
agrep -0 -c 'TTCAATCGTTCGTTCCA' *.fastq > count/B4465_Fm0.txt
agrep -0 -c 'ACACCGATTACGACCGA' *.fastq > count/B4467_Fm0.txt
agrep -0 -c 'GTCCTCACGCGACTGAA' *.fastq > count/B4468_Fm0.txt
agrep -0 -c 'TATTGGTCAGGCAGCCT' *.fastq > count/B4469_Fm0.txt
agrep -0 -c 'ACCCCCATCAAGGTCTA' *.fastq > count/B4470_Fm0.txt
agrep -0 -c 'GTCTCTCGTATTTACGA' *.fastq > count/B4471_Fm0.txt
agrep -0 -c 'TTATCTGATGTGCCGAG' *.fastq > count/B4472_Fm0.txt
agrep -0 -c 'CTCGAAGGGTGGTTTCC' *.fastq > count/B4473_Fm0.txt
agrep -0 -c 'AGGCTCAAGCGTAAGAA' *.fastq > count/B4474_Fm0.txt
agrep -0 -c 'AGCCTGCCGACCACTAA' *.fastq > count/B4475_Fm0.txt
agrep -0 -c 'CCGCACCGTTCAAGAGT' *.fastq > count/B4476_Fm0.txt
agrep -0 -c 'ATGAGCACCGGGATACT' *.fastq > count/B4477_Fm0.txt
agrep -0 -c 'CAACTCCGCCGAAAAAG' *.fastq > count/B4478_Fm0.txt
agrep -0 -c 'TCGTACTACGCCTAGAT' *.fastq > count/B4479_Fm0.txt
agrep -0 -c 'GTATAGCGAGGACGGCA' *.fastq > count/B4481_Fm0.txt
agrep -0 -c 'CTCAGTTCATGAGTGAG' *.fastq > count/B4482_Fm0.txt
agrep -0 -c 'CAGAGTTGAACCAAACG' *.fastq > count/B4483_Fm0.txt
agrep -0 -c 'GCGCTAATCGAGCTTGG' *.fastq > count/B4485_Fm0.txt
agrep -0 -c 'AACTCCTCGTTAACCTG' *.fastq > count/B4487_Fm0.txt
agrep -0 -c 'ACTGCAAGAATCCCGGT' *.fastq > count/B4488_Fm0.txt
agrep -0 -c 'TGGATTACTCCGCAAGT' *.fastq > count/B4489_Fm0.txt
agrep -0 -c 'CTTGCCAGCGGGCAAAT' *.fastq > count/B4491_Fm0.txt
agrep -0 -c 'TTCTCCGTATTACTTCG' *.fastq > count/B4492_Fm0.txt
agrep -0 -c 'GTTTCCCGCATGTTCGG' *.fastq > count/B4494_Fm0.txt
agrep -0 -c 'AATCATGCCGCCCGTCT' *.fastq > count/B4495_Fm0.txt
agrep -0 -c 'ATGCGCTATAACCTTGG' *.fastq > count/B4496_Fm0.txt
agrep -0 -c 'GTCGTGTTATTTCCTAC' *.fastq > count/B4497_Fm0.txt
agrep -0 -c 'CCTAAAATGGGGTCGTT' *.fastq > count/B4498_Fm0.txt
agrep -0 -c 'CATTACTTCGGGCTAGA' *.fastq > count/B4499_Fm0.txt
agrep -0 -c 'AAGGGGAACTGCTCGCA' *.fastq > count/B4500_Fm0.txt
agrep -0 -c 'GTAATAGAGTATAGCCC' *.fastq > count/B4501_Fm0.txt
agrep -0 -c 'AACCGCTTGGCTTCAAA' *.fastq > count/B4502_Fm0.txt
agrep -0 -c 'ACAGCTTCTCCTCAACG' *.fastq > count/B4504_Fm0.txt
agrep -0 -c 'GCGTCGTTCGGTCTACA' *.fastq > count/B4505_Fm0.txt
agrep -0 -c 'GGTAAGACATCAAACGA' *.fastq > count/B4506_Fm0.txt
agrep -0 -c 'GTCCGATGAATCCCCAG' *.fastq > count/B4508_Fm0.txt
agrep -0 -c 'GTGGGACACTCGTAGTA' *.fastq > count/B4509_Fm0.txt
agrep -0 -c 'CTACGATAACCTGATAC' *.fastq > count/B4510_Fm0.txt
agrep -0 -c 'TCGATTGTTGCCAGAGG' *.fastq > count/B4512_Fm0.txt
agrep -0 -c 'CGTTGCTCAGTACGTCC' *.fastq > count/B4514_Fm0.txt
agrep -0 -c 'ATAAAGAGTTGGTCCGC' *.fastq > count/B4515_Fm0.txt
agrep -0 -c 'ACGCTTCGAGTACGTGT' *.fastq > count/B4517_Fm0.txt
agrep -0 -c 'GGTTCTTTGAACGCGTT' *.fastq > count/B4518_Fm0.txt
agrep -0 -c 'CTATCAAACGATAGGGT' *.fastq > count/B4519_Fm0.txt
agrep -0 -c 'CCTGTCCTTATAGATCC' *.fastq > count/B4520_Fm0.txt
agrep -0 -c 'ACTCCACTATGTCTTCT' *.fastq > count/B4522_Fm0.txt
agrep -0 -c 'TTACACCTAACCCGCCT' *.fastq > count/B4523_Fm0.txt
agrep -0 -c 'ACCCCAAAGCTAACGAG' *.fastq > count/B4525_Fm0.txt
agrep -0 -c 'TACCCTTATGGATCGGC' *.fastq > count/B4526_Fm0.txt
agrep -0 -c 'CCTTGGAGCTAATAGAG' *.fastq > count/B4527_Fm0.txt
agrep -0 -c 'CCATCAAATCAGACCAC' *.fastq > count/B4528_Fm0.txt
agrep -0 -c 'GTAAGTGCTAAACGCTG' *.fastq > count/B4529_Fm0.txt
agrep -0 -c 'GTAACGGTAACCCCTGT' *.fastq > count/B4530_Fm0.txt
agrep -0 -c 'CCTTCATGTAAAGGAGC' *.fastq > count/B4531_Fm0.txt
agrep -0 -c 'TCGGTCAGCTAATGCCA' *.fastq > count/B4532_Fm0.txt
agrep -0 -c 'GGATGCGCGGTTTGTAT' *.fastq > count/B4533_Fm0.txt
agrep -0 -c 'ACGTCGTCAGGTCGACA' *.fastq > count/B4534_Fm0.txt
agrep -0 -c 'CAAGAGAAAACAGTCTG' *.fastq > count/B4535_Fm0.txt
agrep -0 -c 'GTTCGCCAGTAGTAGCC' *.fastq > count/B4536_Fm0.txt
agrep -0 -c 'CTTGCCCAACAGGGCAT' *.fastq > count/B4537_Fm0.txt
agrep -0 -c 'ACTATGCATAGGTGTGG' *.fastq > count/B4538_Fm0.txt
agrep -0 -c 'GTACCAGCCTTATGCTG' *.fastq > count/B4539_Fm0.txt
agrep -0 -c 'AGTACAACGTGGCACTA' *.fastq > count/B4540_Fm0.txt
agrep -0 -c 'ATATAGTCTCCACGGTT' *.fastq > count/B4541_Fm0.txt
agrep -0 -c 'GGCGGTTATCATTTGCC' *.fastq > count/B4542_Fm0.txt
agrep -0 -c 'TAACCATCTCGCACCGA' *.fastq > count/B4543_Fm0.txt
agrep -0 -c 'ATATGTGACCACCGGCA' *.fastq > count/B4544_Fm0.txt
agrep -0 -c 'GGAACTTGGTAGAGCCT' *.fastq > count/B4545_Fm0.txt
agrep -0 -c 'CGGTTGACTGAACAGGC' *.fastq > count/B4547_Fm0.txt
agrep -0 -c 'TTATGAACGCCTAGGAT' *.fastq > count/B4548_Fm0.txt
agrep -0 -c 'CTCCGGACACATTTTAC' *.fastq > count/B4549_Fm0.txt
agrep -0 -c 'AATGCGACATCACGCGC' *.fastq > count/B4551_Fm0.txt
agrep -0 -c 'GCGTCCAGATTCAAAAA' *.fastq > count/B4553_Fm0.txt
agrep -0 -c 'CTACAGTACGAAGTTAC' *.fastq > count/B4554_Fm0.txt
agrep -0 -c 'CACCGCAGTACCTCCAT' *.fastq > count/B4556_Fm0.txt
agrep -0 -c 'TACCCGAGTCCAGTTAC' *.fastq > count/B4558_Fm0.txt
agrep -0 -c 'GGATACCATAGAGTGCG' *.fastq > count/B4559_Fm0.txt
agrep -0 -c 'CACTATGGTATCCCCAT' *.fastq > count/B4561_Fm0.txt
agrep -0 -c 'AACCGTGCCTACCGTTG' *.fastq > count/B4562_Fm0.txt
agrep -0 -c 'TTGTACTCGCGCGAAAT' *.fastq > count/B4563_Fm0.txt
agrep -0 -c 'GTACCGCGGTACGAATA' *.fastq > count/B4564_Fm0.txt
agrep -0 -c 'TGAAGGAACTCCAACAT' *.fastq > count/B4566_Fm0.txt
agrep -0 -c 'GATGGTTCGGTATAACT' *.fastq > count/B4567_Fm0.txt
agrep -0 -c 'GCAGTACAGCAAGGTCG' *.fastq > count/B4568_Fm0.txt
agrep -0 -c 'ATGCCATGCGCGTTAAA' *.fastq > count/B4569_Fm0.txt
agrep -0 -c 'GAGTCTGAAGAAGGTCC' *.fastq > count/B4570_Fm0.txt
agrep -0 -c 'GTTTCAGCGGATCGGTA' *.fastq > count/B4572_Fm0.txt
agrep -0 -c 'CAGGGGTGTCTTCACTT' *.fastq > count/B4574_Fm0.txt
agrep -0 -c 'CGAACGTTGGTAATGCA' *.fastq > count/B4575_Fm0.txt
agrep -0 -c 'CATGGATTATTCGAACG' *.fastq > count/B4576_Fm0.txt
agrep -0 -c 'CGATTCGGCAAACTATG' *.fastq > count/B4580_Fm0.txt
agrep -0 -c 'GATCGAGTATTCGCAGT' *.fastq > count/B4581_Fm0.txt
agrep -0 -c 'AATACGTGCGCACTTGA' *.fastq > count/B4583_Fm0.txt
agrep -0 -c 'ACTAGATTACTCAGACC' *.fastq > count/B4584_Fm0.txt
agrep -0 -c 'GGTCGTTCGTGATTCGG' *.fastq > count/B4585_Fm0.txt
agrep -0 -c 'AATTTGGGTCCTCGAAC' *.fastq > count/B4586_Fm0.txt
agrep -0 -c 'TCGTTGGACTACGCTGA' *.fastq > count/B4587_Fm0.txt
agrep -0 -c 'GACTTACTCGTGGGTTC' *.fastq > count/B4589_Fm0.txt
agrep -0 -c 'TATTAGCGCCTCGCCAC' *.fastq > count/B4590_Fm0.txt
agrep -0 -c 'ACTGATAGGAAGGCCTA' *.fastq > count/B4591_Fm0.txt
agrep -0 -c 'TGCTAACATTCACGTCG' *.fastq > count/B4592_Fm0.txt
agrep -0 -c 'CACGATCCAACGTGCCT' *.fastq > count/B4593_Fm0.txt
agrep -0 -c 'CGCAAACCGTGTACCAA' *.fastq > count/B4594_Fm0.txt
agrep -0 -c 'CGTCGCCGAAGCCTAAA' *.fastq > count/B4595_Fm0.txt
agrep -0 -c 'GATACGTACCGCGTATT' *.fastq > count/B4596_Fm0.txt
agrep -0 -c 'GTGCCCCATTGGTATTC' *.fastq > count/B4597_Fm0.txt
agrep -0 -c 'ACGGTAGAGGTAAGTAT' *.fastq > count/B4600_Fm0.txt
agrep -0 -c 'AATGTCATGACGAGGCC' *.fastq > count/B4601_Fm0.txt
agrep -0 -c 'TATATCTCAAAGCTCGC' *.fastq > count/B4602_Fm0.txt
agrep -0 -c 'TTCTTGGTATCTGGCGA' *.fastq > count/B4604_Fm0.txt
agrep -0 -c 'ACACAAAGGCCTATCTC' *.fastq > count/B4605_Fm0.txt
agrep -0 -c 'GTAGACACTGTCCTCAG' *.fastq > count/B4606_Fm0.txt
agrep -0 -c 'TTACACTAAGGCTACGG' *.fastq > count/B4607_Fm0.txt
agrep -0 -c 'AAGGGATGCAAGGACAA' *.fastq > count/B4609_Fm0.txt
agrep -0 -c 'CTGAGCACTGATCGCAG' *.fastq > count/B4610_Fm0.txt
agrep -0 -c 'ATCTAGCCCAGTTGTTT' *.fastq > count/B4611_Fm0.txt
agrep -0 -c 'CGATTGGAGTAGCTGAG' *.fastq > count/B4612_Fm0.txt
agrep -0 -c 'TGCGGCATATCGGCTAC' *.fastq > count/B4613_Fm0.txt
agrep -0 -c 'CCATTTGGATTGCCGAG' *.fastq > count/B4614_Fm0.txt
agrep -0 -c 'AGGCCGAGGAATTGAAC' *.fastq > count/B4616_Fm0.txt
agrep -0 -c 'AGCCTCGGAAGACTTAC' *.fastq > count/B4617_Fm0.txt
agrep -0 -c 'GTACCCTTGGCTTACCC' *.fastq > count/B4618_Fm0.txt
agrep -0 -c 'TCTTCTACGGATGTCAT' *.fastq > count/B4619_Fm0.txt
agrep -0 -c 'TCGAAAGTCGAAGTGTT' *.fastq > count/B4620_Fm0.txt
agrep -0 -c 'GCCCCGAAGGTAGATTT' *.fastq > count/B4621_Fm0.txt
agrep -0 -c 'GAGTGTTCGCCTACATC' *.fastq > count/B4622_Fm0.txt
agrep -0 -c 'TGAGGCATGCGTATCAA' *.fastq > count/B4623_Fm0.txt
agrep -0 -c 'ACGCGTCACGCTGGATT' *.fastq > count/B4624_Fm0.txt
agrep -0 -c 'ACCTACAGGCGATATGC' *.fastq > count/B4625_Fm0.txt
agrep -0 -c 'TGCCAAGCATTCGAGAG' *.fastq > count/B4626_Fm0.txt
agrep -0 -c 'CGACTGGAATACGACGT' *.fastq > count/B4627_Fm0.txt
agrep -0 -c 'AAGAGGATACCCAAGGT' *.fastq > count/B4628_Fm0.txt
agrep -0 -c 'GCAGTTTGCTAATGCTT' *.fastq > count/B4629_Fm0.txt
agrep -0 -c 'GCCGCCGAAGTAACTTT' *.fastq > count/B4630_Fm0.txt
agrep -0 -c 'AACAGCGATTATCCGTA' *.fastq > count/B4631_Fm0.txt
agrep -0 -c 'CATCTTTTGACGAAGAG' *.fastq > count/B4633_Fm0.txt
agrep -0 -c 'CGGGAGGCAGCAATATT' *.fastq > count/B4634_Fm0.txt
agrep -0 -c 'AGATTTTCCTGGGTGTC' *.fastq > count/B4635_Fm0.txt
agrep -0 -c 'ACTGATGGTTACCCGCT' *.fastq > count/B4636_Fm0.txt
agrep -0 -c 'CCGGTTACAGACTATAA' *.fastq > count/B4637_Fm0.txt
agrep -0 -c 'CTTCTTCCCGAATGTAC' *.fastq > count/B4639_Fm0.txt
agrep -0 -c 'ATCCAGTTGGCGGACTA' *.fastq > count/B4642_Fm0.txt
agrep -0 -c 'GTAATGTCGGACCTGGG' *.fastq > count/B4643_Fm0.txt
agrep -0 -c 'AGAGTGACGACATGGGG' *.fastq > count/B4644_Fm0.txt
agrep -0 -c 'CTCCGTAATGTCTCCGT' *.fastq > count/B4645_Fm0.txt
agrep -0 -c 'GGCCTTAGTTTTGGCCT' *.fastq > count/B4646_Fm0.txt
agrep -0 -c 'CTCGGAATATCTGCAGA' *.fastq > count/B4647_Fm0.txt
agrep -0 -c 'AGTCGGAGTTGGCATCA' *.fastq > count/B4648_Fm0.txt
agrep -0 -c 'CTGGTTACGAAGTCCGG' *.fastq > count/B4649_Fm0.txt
agrep -0 -c 'ATCATCGGGATCCACGG' *.fastq > count/B4650_Fm0.txt
agrep -0 -c 'TATCTCTAATCTTGGCG' *.fastq > count/B4651_Fm0.txt
agrep -0 -c 'AAGATGTCGGGTACCCA' *.fastq > count/B4653_Fm0.txt
agrep -0 -c 'TTAAGTCCGACGGAAAT' *.fastq > count/B4654_Fm0.txt
agrep -0 -c 'GCTCCTTAAACCAGGAC' *.fastq > count/B4655_Fm0.txt
agrep -0 -c 'ATTCATAGGCTCGACGT' *.fastq > count/B4656_Fm0.txt
agrep -0 -c 'CGTTTCACTGAGTTGAT' *.fastq > count/B4657_Fm0.txt
agrep -0 -c 'GAGATATATCTCCTGCT' *.fastq > count/B4659_Fm0.txt
agrep -0 -c 'CATCCTTGGAGTTGCGA' *.fastq > count/B4660_Fm0.txt
agrep -0 -c 'CGTCGCAACCAGATCTT' *.fastq > count/B4661_Fm0.txt
agrep -0 -c 'GATACTCCCGGTACAAT' *.fastq > count/B4662_Fm0.txt
agrep -0 -c 'TGTCAGGACCTTGATAG' *.fastq > count/B4663_Fm0.txt
agrep -0 -c 'GGAGGCATTATTCGAGT' *.fastq > count/B4665_Fm0.txt
agrep -0 -c 'TTATCGCTGGGAACATG' *.fastq > count/B4666_Fm0.txt
agrep -0 -c 'TTAACCAGTCGCACGCA' *.fastq > count/B4668_Fm0.txt
agrep -0 -c 'TGAGAATGAGTTCACGC' *.fastq > count/B4669_Fm0.txt
agrep -0 -c 'TAAAGGCAGTGGCACAC' *.fastq > count/B4671_Fm0.txt
agrep -0 -c 'GTGGAATCCATTGGGTT' *.fastq > count/B4672_Fm0.txt
agrep -0 -c 'GTTGAGGTCCGGTCCAA' *.fastq > count/B4673_Fm0.txt
agrep -0 -c 'CTTATCCCCCATGGTCA' *.fastq > count/B4674_Fm0.txt
agrep -0 -c 'ATCACTGCCGCTTTTGT' *.fastq > count/B4675_Fm0.txt
agrep -0 -c 'TGACGTCTAACTGAGGC' *.fastq > count/B4676_Fm0.txt
agrep -0 -c 'AGCGATCGCGGATTGCA' *.fastq > count/B4677_Fm0.txt
agrep -0 -c 'CGGTCATGCCGTTATGC' *.fastq > count/B4678_Fm0.txt
agrep -0 -c 'ATCTTGACCTACCTAGA' *.fastq > count/B4679_Fm0.txt
agrep -0 -c 'CTTGACTTAGGGTGGGC' *.fastq > count/B4680_Fm0.txt
agrep -0 -c 'CATGACGGACTCGTGTT' *.fastq > count/B4681_Fm0.txt
agrep -0 -c 'ACCTGTCGTTCGAGAAA' *.fastq > count/B4682_Fm0.txt
agrep -0 -c 'CGGACGTAAAGCAAGTA' *.fastq > count/B4683_Fm0.txt
agrep -0 -c 'TGTAATTGTTGGCAGCG' *.fastq > count/B4684_Fm0.txt
agrep -0 -c 'TGGCTGTGAATTAATGC' *.fastq > count/B4685_Fm0.txt
agrep -0 -c 'CCGACTGTTGGTATTGC' *.fastq > count/B4687_Fm0.txt
agrep -0 -c 'TGACTCAAACGGATCCC' *.fastq > count/B4689_Fm0.txt
agrep -0 -c 'GCTACGTACTCGTTAAA' *.fastq > count/B4690_Fm0.txt
agrep -0 -c 'TTGTAGCACTCTAGCCA' *.fastq > count/B4691_Fm0.txt
agrep -0 -c 'TAGCCCTCATGATGCAA' *.fastq > count/B4693_Fm0.txt
agrep -0 -c 'GTACACTCTGTCCAGAC' *.fastq > count/B4694_Fm0.txt
agrep -0 -c 'AAGTTATACCGCCCGTT' *.fastq > count/B4695_Fm0.txt
agrep -0 -c 'GACTTGGCTGACGGTTA' *.fastq > count/B4696_Fm0.txt
agrep -0 -c 'AATCGGGAAACGCCGAA' *.fastq > count/B4697_Fm0.txt
agrep -0 -c 'CGTCTTATTTCGCGTGC' *.fastq > count/B4698_Fm0.txt
agrep -0 -c 'CGACAATTTCTCCAATC' *.fastq > count/B4699_Fm0.txt
agrep -0 -c 'CATTCGTACGGCCACAA' *.fastq > count/B4700_Fm0.txt
agrep -0 -c 'CGCAATAAACATTCGGA' *.fastq > count/B4701_Fm0.txt
agrep -0 -c 'GGGCAAACTAGTCATCA' *.fastq > count/B4702_Fm0.txt
agrep -0 -c 'GCCATTTGTCTATGTAC' *.fastq > count/B4704_Fm0.txt
agrep -0 -c 'AGACATTGGCGCCACGA' *.fastq > count/B4705_Fm0.txt
agrep -0 -c 'GCTGCGAAAGGCATGGT' *.fastq > count/B4706_Fm0.txt
agrep -0 -c 'TGAGCGCCGTTCGTACA' *.fastq > count/B4707_Fm0.txt
agrep -0 -c 'CTACAATAGGTCACCAT' *.fastq > count/B4708_Fm0.txt
agrep -0 -c 'GCCGTTACTCAAAGACT' *.fastq > count/B4710_Fm0.txt
agrep -0 -c 'GAGTTAAGACCCCTCTA' *.fastq > count/B4712_Fm0.txt
agrep -0 -c 'CGTCAGATAATGCGAAT' *.fastq > count/B4713_Fm0.txt
agrep -0 -c 'ATTAGCGTAGCCATCTT' *.fastq > count/B4714_Fm0.txt
agrep -0 -c 'GCTAGGAGCTAACGTAC' *.fastq > count/B4716_Fm0.txt
agrep -0 -c 'GTATGGATGCGTATTGG' *.fastq > count/B4717_Fm0.txt
agrep -0 -c 'ACAAGCGACAACTCTAA' *.fastq > count/B4719_Fm0.txt
agrep -0 -c 'CGGACTTTTAGGTTTTG' *.fastq > count/B4720_Fm0.txt
agrep -0 -c 'TCTACTGACTCGCGTCA' *.fastq > count/B4721_Fm0.txt
agrep -0 -c 'TGAGCCCCAATAGATAT' *.fastq > count/B4722_Fm0.txt
agrep -0 -c 'ACGAGCGTTCTATCTAG' *.fastq > count/B4723_Fm0.txt
agrep -0 -c 'GGAGCTTTGCTAAGATA' *.fastq > count/B4724_Fm0.txt
agrep -0 -c 'TAGGAACGGTATCTTCA' *.fastq > count/B4726_Fm0.txt
agrep -0 -c 'CCGTGTCAAACTTTTCC' *.fastq > count/B4727_Fm0.txt
agrep -0 -c 'GGAATGGTGTTGGTAAA' *.fastq > count/B4728_Fm0.txt
agrep -0 -c 'CAAAGAGGCGTACCAGA' *.fastq > count/B4729_Fm0.txt
agrep -0 -c 'AAGAGACTAGTTCCATC' *.fastq > count/B4731_Fm0.txt
agrep -0 -c 'CCAGCAAATTTTAGGCG' *.fastq > count/B4732_Fm0.txt
agrep -0 -c 'CATCGCAATTGAAGTAG' *.fastq > count/B4733_Fm0.txt
agrep -0 -c 'GGATATCAGCACCACGG' *.fastq > count/B4734_Fm0.txt
agrep -0 -c 'TTGACTTCACGAACAAG' *.fastq > count/B4735_Fm0.txt
agrep -0 -c 'CATCTCATAAAGTGGCG' *.fastq > count/B4736_Fm0.txt
agrep -0 -c 'TAACCGTCCCAGACGAC' *.fastq > count/B4737_Fm0.txt
agrep -0 -c 'CACGGTTGGAGTATATG' *.fastq > count/B4738_Fm0.txt
agrep -0 -c 'GAGTATAATGGCGGCCG' *.fastq > count/B4739_Fm0.txt
agrep -0 -c 'TTAGGAAGACTTTGAGC' *.fastq > count/B4740_Fm0.txt
agrep -0 -c 'CGTTGCGCTAGCTGATA' *.fastq > count/B4741_Fm0.txt
agrep -0 -c 'GCGTAACTCTAGACCCA' *.fastq > count/B4742_Fm0.txt
agrep -0 -c 'TGCAACCGAACGTCGAG' *.fastq > count/B4743_Fm0.txt
agrep -0 -c 'AGATCACGCCAATTTTG' *.fastq > count/B4744_Fm0.txt
agrep -0 -c 'GGTAGGAAGAAGCTGAT' *.fastq > count/B4745_Fm0.txt
agrep -0 -c 'GCCTCCTAGTGTGTTGG' *.fastq > count/B4746_Fm0.txt
agrep -0 -c 'CTTCAGAGTTACACTCT' *.fastq > count/B4747_Fm0.txt
agrep -0 -c 'GATAAGTAGGGATCAAC' *.fastq > count/B4750_Fm0.txt
agrep -0 -c 'TTACGCCTTCCACTTAC' *.fastq > count/B4752_Fm0.txt
agrep -0 -c 'CGATAGGATTTGGGTCT' *.fastq > count/B4753_Fm0.txt
agrep -0 -c 'CGATCAGAACGAAGCAC' *.fastq > count/B4757_Fm0.txt
agrep -0 -c 'AATACCTGCACGTAGGT' *.fastq > count/B4758_Fm0.txt
agrep -0 -c 'CTGGTCTTGGTCGGTGA' *.fastq > count/B4759_Fm0.txt
agrep -0 -c 'AGATTCAAGGTGGCGTT' *.fastq > count/B4762_Fm0.txt
agrep -0 -c 'GCGACAGTGATTTTCTG' *.fastq > count/B4763_Fm0.txt
agrep -0 -c 'CGGTAAATCAACGCGTC' *.fastq > count/B4765_Fm0.txt
agrep -0 -c 'ACGTGTTGTCACCTTGT' *.fastq > count/B4766_Fm0.txt
agrep -0 -c 'GACCACCTGACCACATC' *.fastq > count/B4769_Fm0.txt
agrep -0 -c 'ATCGGGTTTAGCACCAT' *.fastq > count/B4770_Fm0.txt
agrep -0 -c 'GTACGGGCAAAATTCTG' *.fastq > count/B4771_Fm0.txt
agrep -0 -c 'GATTGCCTACTCTGGTT' *.fastq > count/B4772_Fm0.txt
agrep -0 -c 'GGCAAAGCACGTGATTA' *.fastq > count/B4773_Fm0.txt
agrep -0 -c 'GCGTTCTATCTGTGACA' *.fastq > count/B4774_Fm0.txt
agrep -0 -c 'TCAACTCCCGCGCTTAA' *.fastq > count/B4775_Fm0.txt
agrep -0 -c 'TACCCGGAGGAAACACA' *.fastq > count/B4776_Fm0.txt
agrep -0 -c 'GGAACCGAACTTATCAA' *.fastq > count/B4780_Fm0.txt
agrep -0 -c 'AGATGAATCGGAAACGA' *.fastq > count/B4781_Fm0.txt
agrep -0 -c 'TTCCAATCGTGGCACTG' *.fastq > count/B4782_Fm0.txt
agrep -0 -c 'CCGGGCGTGAAAGTATG' *.fastq > count/B4783_Fm0.txt
agrep -0 -c 'AGTCCGTACTTATGAGA' *.fastq > count/B4784_Fm0.txt
agrep -0 -c 'GATTCCCAACCTAGGAT' *.fastq > count/B4785_Fm0.txt
agrep -0 -c 'CTCAAAGGGCTGAACCC' *.fastq > count/B4786_Fm0.txt
agrep -0 -c 'ATCACATCGTATCCGTG' *.fastq > count/B4787_Fm0.txt
agrep -0 -c 'GACCTGCGCGGAAAATT' *.fastq > count/B4789_Fm0.txt
agrep -0 -c 'AGAGTTCCGAATAGCTT' *.fastq > count/B4790_Fm0.txt
agrep -0 -c 'GCGCTGCGAGGAAAATA' *.fastq > count/B4791_Fm0.txt
agrep -0 -c 'ATAGTAGTGTTCTGAGC' *.fastq > count/B4792_Fm0.txt
agrep -0 -c 'ATCTTGAACTCGGCTGC' *.fastq > count/B4793_Fm0.txt
agrep -0 -c 'GACACTTACCCGCCTCA' *.fastq > count/B4794_Fm0.txt
agrep -0 -c 'CTTCGGCGTCTTATACT' *.fastq > count/B4795_Fm0.txt
agrep -0 -c 'ATAACTTCTCCTGCCTT' *.fastq > count/B4796_Fm0.txt
agrep -0 -c 'AGTCACGCACCAATCCA' *.fastq > count/B4797_Fm0.txt
agrep -0 -c 'GCCTGTTCATTGAGTCC' *.fastq > count/B4799_Fm0.txt
agrep -0 -c 'CGTCCAGCACCTTCGTA' *.fastq > count/B4800_Fm0.txt
agrep -0 -c 'CGGATCGCTGCACCTTT' *.fastq > count/B4801_Fm0.txt
agrep -0 -c 'GGGAGGTTGCGAATTGT' *.fastq > count/B4802_Fm0.txt
agrep -0 -c 'GACTAAGGCAATGTTGT' *.fastq > count/B4803_Fm0.txt
agrep -0 -c 'GGAGCGACAAACACATA' *.fastq > count/B4804_Fm0.txt
agrep -0 -c 'CAACTGCCATTTCTTGA' *.fastq > count/B4806_Fm0.txt
agrep -0 -c 'CGCTTCTTAGGACGATT' *.fastq > count/B4807_Fm0.txt
agrep -0 -c 'GGTATGCAAGATACCAT' *.fastq > count/B4809_Fm0.txt
agrep -0 -c 'GGTCTACATCATGGCTT' *.fastq > count/B4810_Fm0.txt
agrep -0 -c 'GTTGGACTAGTCCCATT' *.fastq > count/B4811_Fm0.txt
agrep -0 -c 'AACGTGGGCACTAACTT' *.fastq > count/B4812_Fm0.txt
agrep -0 -c 'GTAGCATCATGGGAGAC' *.fastq > count/B4816_Fm0.txt
agrep -0 -c 'TATTCGACCATGGGCAC' *.fastq > count/B4817_Fm0.txt
agrep -0 -c 'AGTGATCCAACGGGGGA' *.fastq > count/B4818_Fm0.txt
agrep -0 -c 'CGAGCAGTATGCTCACT' *.fastq > count/B4819_Fm0.txt
agrep -0 -c 'GGAACGTAGAATACGAT' *.fastq > count/B4820_Fm0.txt
agrep -0 -c 'CACGTTGTGAAGTCCAA' *.fastq > count/B4821_Fm0.txt
agrep -0 -c 'GTTAGAAGTTGCCTTTG' *.fastq > count/B4822_Fm0.txt
agrep -0 -c 'GCAGGAGAAGTGAATAT' *.fastq > count/B4823_Fm0.txt
agrep -0 -c 'ATTTCGCCAACCTAAGA' *.fastq > count/B4825_Fm0.txt
agrep -0 -c 'GCAACTTTTTCCTCGTG' *.fastq > count/B4826_Fm0.txt
agrep -0 -c 'CTATAGGACGCCCGATA' *.fastq > count/B4827_Fm0.txt
agrep -0 -c 'TGAGCGATCTTAAATCG' *.fastq > count/B4828_Fm0.txt
agrep -0 -c 'AGACGGTTAACCTGGGG' *.fastq > count/B4829_Fm0.txt
agrep -0 -c 'AACGGCAGTCAGCACTA' *.fastq > count/B4830_Fm0.txt
agrep -0 -c 'GCGCAGAGAACCGTAAG' *.fastq > count/B4831_Fm0.txt
agrep -0 -c 'GCGTCTTCGTTACTAGA' *.fastq > count/B4833_Fm0.txt
agrep -0 -c 'AGGCCGAAACGGATCGA' *.fastq > count/B4834_Fm0.txt
agrep -0 -c 'GCTGTCCAAAAAACTGT' *.fastq > count/B4835_Fm0.txt
agrep -0 -c 'AATGCCGGGGGTAAGGA' *.fastq > count/B4836_Fm0.txt
agrep -0 -c 'GATCGGTAGCATTGATT' *.fastq > count/B4838_Fm0.txt
agrep -0 -c 'AACGCTATGCAACTCTC' *.fastq > count/B4839_Fm0.txt
agrep -0 -c 'AAGGTCGAAAATCGACC' *.fastq > count/B4840_Fm0.txt
agrep -0 -c 'ATATCCCCTAACGCATT' *.fastq > count/B4841_Fm0.txt
agrep -0 -c 'ACTCGACCTAACTTATG' *.fastq > count/B4842_Fm0.txt
agrep -0 -c 'GTGTCCATCGTTCAACG' *.fastq > count/B4844_Fm0.txt
agrep -0 -c 'CAAAGCGATTATACGCG' *.fastq > count/B4845_Fm0.txt
agrep -0 -c 'GTGGTGACTACACAGTA' *.fastq > count/B4846_Fm0.txt
agrep -0 -c 'CCAGGTTGTCCAAGAGG' *.fastq > count/B4847_Fm0.txt
agrep -0 -c 'CGCCAGTGTTACGTTCG' *.fastq > count/B4848_Fm0.txt
agrep -0 -c 'CCGTCGGATAAGGACGT' *.fastq > count/B4850_Fm0.txt
agrep -0 -c 'TACTTATGCCTTCCTCG' *.fastq > count/B4851_Fm0.txt
agrep -0 -c 'ATTACGGACAGGATTCG' *.fastq > count/B4852_Fm0.txt
agrep -0 -c 'TTCACGCGGTTTGTCGA' *.fastq > count/B4854_Fm0.txt
agrep -0 -c 'TATCAGAGCGAGAAGTG' *.fastq > count/B4855_Fm0.txt
agrep -0 -c 'GCCAGCTCACGATGAGT' *.fastq > count/B4856_Fm0.txt
agrep -0 -c 'CGGAACCCAATATACGG' *.fastq > count/B4857_Fm0.txt
agrep -0 -c 'CCGAATCTAAGTGGGGG' *.fastq > count/B4859_Fm0.txt
agrep -0 -c 'CGAGGGACAAAATTGGA' *.fastq > count/B4860_Fm0.txt
agrep -0 -c 'CTAGTGACCCTTACCGT' *.fastq > count/B4861_Fm0.txt
agrep -0 -c 'GATCGATAGCCATGACG' *.fastq > count/B4862_Fm0.txt
agrep -0 -c 'CGCACTGAGATAGGAAC' *.fastq > count/B4863_Fm0.txt
agrep -0 -c 'ACAACGAGAGTTGTTTC' *.fastq > count/B4865_Fm0.txt
agrep -0 -c 'CAATAGGTGGCTACTTT' *.fastq > count/B4866_Fm0.txt
agrep -0 -c 'CTCGGCTACTAAGTACG' *.fastq > count/B4867_Fm0.txt
agrep -0 -c 'CTCCGGTTGCTGTTTCC' *.fastq > count/B4868_Fm0.txt
agrep -0 -c 'TCAGGTCTAGTAGATGT' *.fastq > count/B4869_Fm0.txt
agrep -0 -c 'GTCTCCGATCTGGAGTA' *.fastq > count/B4871_Fm0.txt
agrep -0 -c 'TTAGGAGGAACTGGTGT' *.fastq > count/B4872_Fm0.txt
agrep -0 -c 'ACATTCCGCTCCCATCA' *.fastq > count/B4873_Fm0.txt
agrep -0 -c 'CAGCGGTTTAACGAAGA' *.fastq > count/B4875_Fm0.txt
agrep -0 -c 'GCTACCTTAATACAAGC' *.fastq > count/B4876_Fm0.txt
agrep -0 -c 'GTGTCACCTGTACGAGC' *.fastq > count/B4877_Fm0.txt
agrep -0 -c 'CCAAACGCTCCCTCACT' *.fastq > count/B4878_Fm0.txt
agrep -0 -c 'GGAGAATTACGCTTCAG' *.fastq > count/B4879_Fm0.txt
agrep -0 -c 'GCGAATCCATGTCAGGT' *.fastq > count/B4881_Fm0.txt
agrep -0 -c 'GGGTCTACTGAATCCTA' *.fastq > count/B4882_Fm0.txt
agrep -0 -c 'ACTCATATAGAGGCGTT' *.fastq > count/B4883_Fm0.txt
agrep -0 -c 'AGGAAGATGCACTCTAA' *.fastq > count/B4884_Fm0.txt
agrep -0 -c 'GTTCCGAACGGGATTTT' *.fastq > count/B4885_Fm0.txt
agrep -0 -c 'GTCACGAGTAGTTCCCA' *.fastq > count/B4886_Fm0.txt
agrep -0 -c 'TGACTCGACTAGATTCG' *.fastq > count/B4887_Fm0.txt
agrep -0 -c 'TCATACACGATCTGATC' *.fastq > count/B4888_Fm0.txt
agrep -0 -c 'AAGAACGTCGCGGTTTG' *.fastq > count/B4889_Fm0.txt
agrep -0 -c 'GTGGATTCCACGAGAGC' *.fastq > count/B4890_Fm0.txt
agrep -0 -c 'AGCCAATAAGTCCGGAT' *.fastq > count/B4891_Fm0.txt
agrep -0 -c 'CTTGGCAGTCTCGATCC' *.fastq > count/B4892_Fm0.txt
agrep -0 -c 'GGCCCGCGTTGACTTAT' *.fastq > count/B4894_Fm0.txt
agrep -0 -c 'TCTAACTTTATCGAGCG' *.fastq > count/B4896_Fm0.txt
agrep -0 -c 'AATTCCCCGTAGCAGCT' *.fastq > count/B4897_Fm0.txt
agrep -0 -c 'CTGACGCAGAGAATCGC' *.fastq > count/B4898_Fm0.txt
agrep -0 -c 'CTTCCACTCTCAGAACG' *.fastq > count/B4899_Fm0.txt
agrep -0 -c 'CACATGGGATACAACAC' *.fastq > count/B4900_Fm0.txt
agrep -0 -c 'GTTGACACTCGTGAGAA' *.fastq > count/B4901_Fm0.txt
agrep -0 -c 'GCGAGACGGTCTAGTTT' *.fastq > count/B4902_Fm0.txt
agrep -0 -c 'GTTCGTCGTCAGCTTAG' *.fastq > count/B4904_Fm0.txt
agrep -0 -c 'ACGTAAACAACGGCGTT' *.fastq > count/B4905_Fm0.txt
agrep -0 -c 'AGCCATGTAAAACCAAG' *.fastq > count/B4906_Fm0.txt
agrep -0 -c 'TAAGACGTCCATGAATG' *.fastq > count/B4907_Fm0.txt
agrep -0 -c 'CCGAGCACGTAACAGGT' *.fastq > count/B4909_Fm0.txt
agrep -0 -c 'CCTAGTACCCAATTGTA' *.fastq > count/B4910_Fm0.txt
agrep -0 -c 'TAGAGTTGCTTTCGGCC' *.fastq > count/B4911_Fm0.txt
agrep -0 -c 'CACCAGTTATAAGCGCA' *.fastq > count/B4912_Fm0.txt
agrep -0 -c 'TGTGGCTGCATTATACT' *.fastq > count/B4913_Fm0.txt
agrep -0 -c 'ATACGTTCAGCGACGAC' *.fastq > count/B4914_Fm0.txt
agrep -0 -c 'TACTCCCGGTCCTTCTC' *.fastq > count/B4915_Fm0.txt
agrep -0 -c 'ACGGCGGCAATCCTACT' *.fastq > count/B4916_Fm0.txt
agrep -0 -c 'GTACCTCGCAATACTAT' *.fastq > count/B4918_Fm0.txt
agrep -0 -c 'AACGATGACAGGGCCGA' *.fastq > count/B4919_Fm0.txt
agrep -0 -c 'CGTCGAATGCAGCCTTT' *.fastq > count/B4920_Fm0.txt
agrep -0 -c 'ACAGCATCCCCGCTTTT' *.fastq > count/B4921_Fm0.txt
agrep -0 -c 'CAAGGCGAATTAGTTGC' *.fastq > count/B4922_Fm0.txt
agrep -0 -c 'GCCCGTTTCCTAAAGCA' *.fastq > count/B4923_Fm0.txt
agrep -0 -c 'TAGCGCACACCTGTTCG' *.fastq > count/B4924_Fm0.txt
agrep -0 -c 'AACCGCCAGTGTATCGA' *.fastq > count/B4925_Fm0.txt
agrep -0 -c 'TTGCCTTACCCGGTAAT' *.fastq > count/B4926_Fm0.txt
agrep -0 -c 'CACAATCTACCAGCCTC' *.fastq > count/B4927_Fm0.txt
agrep -0 -c 'GGTTAGATTGCTGCAAT' *.fastq > count/B4928_Fm0.txt
agrep -0 -c 'CTAACCTGCATTCCTAT' *.fastq > count/B4929_Fm0.txt
agrep -0 -c 'GGCATGATCGGACTCAA' *.fastq > count/B4930_Fm0.txt
agrep -0 -c 'TGATGACCTTGTAGATG' *.fastq > count/B4932_Fm0.txt
agrep -0 -c 'CTTGGTCACTTAGCGGA' *.fastq > count/B4933_Fm0.txt
agrep -0 -c 'GGTGACTACCTAATGTG' *.fastq > count/B4936_Fm0.txt
agrep -0 -c 'CAACACCGTAAGAAGGC' *.fastq > count/B4937_Fm0.txt
agrep -0 -c 'GTCGAACCCTCCCTCAT' *.fastq > count/B4938_Fm0.txt
agrep -0 -c 'CAGTCTCACAACATCCC' *.fastq > count/B4939_Fm0.txt
agrep -0 -c 'AGTTGGACGTCCTGTGT' *.fastq > count/B4940_Fm0.txt
agrep -0 -c 'TCTTAAGTTCCTTCCCG' *.fastq > count/B4941_Fm0.txt
agrep -0 -c 'TCAAAAAATTCCGGCGA' *.fastq > count/B4942_Fm0.txt
agrep -0 -c 'TGAGCTCAGTTTGGCAA' *.fastq > count/B4943_Fm0.txt
agrep -0 -c 'AACCGTGAGATATGTAC' *.fastq > count/B4944_Fm0.txt
agrep -0 -c 'GGAGTACCATTGCGTAC' *.fastq > count/B4945_Fm0.txt
agrep -0 -c 'CAACTAGCGAAGGAAAT' *.fastq > count/B4946_Fm0.txt
agrep -0 -c 'AGCAACTAGTCCCGGTC' *.fastq > count/B4947_Fm0.txt
agrep -0 -c 'ACCTCCCACTTGATGGT' *.fastq > count/B4948_Fm0.txt
agrep -0 -c 'GACTCATCGTATGGATT' *.fastq > count/B4950_Fm0.txt
agrep -0 -c 'CAGAAGAATGCCCGGAT' *.fastq > count/B4951_Fm0.txt
agrep -0 -c 'ATACTGACCGCCGTCAA' *.fastq > count/B4952_Fm0.txt
agrep -0 -c 'CGACTTGCCTAGTAGCT' *.fastq > count/B4954_Fm0.txt
agrep -0 -c 'CAATTCATGAAGAGCAG' *.fastq > count/B4955_Fm0.txt
agrep -0 -c 'ACGTTGAAAGAGGCTAG' *.fastq > count/B4956_Fm0.txt
agrep -0 -c 'TGCAAGCGAAACTAGTG' *.fastq > count/B4958_Fm0.txt
agrep -0 -c 'AGTACGTACGACATGTT' *.fastq > count/B4959_Fm0.txt
agrep -0 -c 'ACTTGGCTTATGTGGTC' *.fastq > count/B4960_Fm0.txt
agrep -0 -c 'TCGTGAAAGATTGACGT' *.fastq > count/B4962_Fm0.txt
agrep -0 -c 'ACAACTTGAGCGATCCG' *.fastq > count/B4963_Fm0.txt
agrep -0 -c 'CAGCCCGTGCTAACGTT' *.fastq > count/B4964_Fm0.txt
agrep -0 -c 'GTTCCCAATCCTACTGC' *.fastq > count/B4965_Fm0.txt
agrep -0 -c 'TCTGCGGAAATGCGTAT' *.fastq > count/B4966_Fm0.txt
agrep -0 -c 'GGATTGGCACTACGTAA' *.fastq > count/B4968_Fm0.txt
agrep -0 -c 'CGCTACAAAATATCACG' *.fastq > count/B4969_Fm0.txt
agrep -0 -c 'CGACTTTCTTTTTGCCC' *.fastq > count/B4970_Fm0.txt
agrep -0 -c 'GATGTACATCGGACTTA' *.fastq > count/B4971_Fm0.txt
agrep -0 -c 'AGTAAAAGGGCGTCGGC' *.fastq > count/B4972_Fm0.txt
agrep -0 -c 'GCCAAGTCAATGAATGC' *.fastq > count/B4974_Fm0.txt
agrep -0 -c 'AGTGCCTGATCGGCCTA' *.fastq > count/B4975_Fm0.txt
agrep -0 -c 'TTGGCACACTTCTATCA' *.fastq > count/B4976_Fm0.txt
agrep -0 -c 'TATTACCCGCGAGCAGA' *.fastq > count/B4977_Fm0.txt
agrep -0 -c 'CTAGCTATGATCGTGTA' *.fastq > count/B4978_Fm0.txt
agrep -0 -c 'AACCCACGACAAAGACT' *.fastq > count/B4979_Fm0.txt
agrep -0 -c 'TGTAGGCTTGGTCTTTG' *.fastq > count/B4980_Fm0.txt
agrep -0 -c 'GCGACTCGTAATGTCCG' *.fastq > count/B4981_Fm0.txt
agrep -0 -c 'GTACTGGCTTCGTATAG' *.fastq > count/B4983_Fm0.txt
agrep -0 -c 'CCTAAGTTGACAAACCT' *.fastq > count/B4986_Fm0.txt
agrep -0 -c 'AGCAATCGTCCTCAGTT' *.fastq > count/B4987_Fm0.txt
agrep -0 -c 'CACGGCCAATCGTCAAA' *.fastq > count/B4989_Fm0.txt
agrep -0 -c 'TGCTATCCACAGAAGTA' *.fastq > count/B4990_Fm0.txt
agrep -0 -c 'ATGGATAACGGAGTCCT' *.fastq > count/B4991_Fm0.txt
agrep -0 -c 'CACAGACCCTAATGGGC' *.fastq > count/B4992_Fm0.txt
agrep -0 -c 'AACGTATCGCTGGGTCG' *.fastq > count/B4993_Fm0.txt
agrep -0 -c 'ACGGCTACAGTTCCAGG' *.fastq > count/B4995_Fm0.txt
agrep -0 -c 'CTCACAAGTGCTTCTCA' *.fastq > count/B4996_Fm0.txt
agrep -0 -c 'TAAACCTGGCCCCGTAG' *.fastq > count/B4997_Fm0.txt
agrep -0 -c 'CGGAATCGCTTCACATT' *.fastq > count/B4998_Fm0.txt
agrep -0 -c 'ATTGGTATCGTCACCAT' *.fastq > count/B4999_Fm0.txt
agrep -0 -c 'GGAATGTGAAAGGCCGC' *.fastq > count/B5001_Fm0.txt
agrep -0 -c 'GCCTCGCAAGTTTGATC' *.fastq > count/B5002_Fm0.txt
agrep -0 -c 'CACAGGTAGGGTCGTAC' *.fastq > count/B5003_Fm0.txt
agrep -0 -c 'CCTTCAAGGTATCTGCA' *.fastq > count/B5004_Fm0.txt
agrep -0 -c 'CCAACTTCTCAACTAGT' *.fastq > count/B5005_Fm0.txt
agrep -0 -c 'TGCCCTCGTTCGGGTAA' *.fastq > count/B5007_Fm0.txt
agrep -0 -c 'TACGACAAATGACACGA' *.fastq > count/B5009_Fm0.txt
agrep -0 -c 'ATGGTTCCCATCCCTAA' *.fastq > count/B5010_Fm0.txt
agrep -0 -c 'GATAGACTTGACAGGTT' *.fastq > count/B5011_Fm0.txt
agrep -0 -c 'CGCCAGATGCAACGCTA' *.fastq > count/B5012_Fm0.txt
agrep -0 -c 'GGTCGTTTTAGTACCTT' *.fastq > count/B5013_Fm0.txt
agrep -0 -c 'GTTCCGGGAGCAAGATG' *.fastq > count/B5014_Fm0.txt
agrep -0 -c 'GGATAAGCCTCCTCAGT' *.fastq > count/B5019_Fm0.txt
agrep -0 -c 'CCGGATGCTATGGCAGA' *.fastq > count/B5020_Fm0.txt
agrep -0 -c 'CTACCCGTAATCTCCGG' *.fastq > count/B5021_Fm0.txt
agrep -0 -c 'ACTCGATCCTCTCGGTA' *.fastq > count/B5022_Fm0.txt
agrep -0 -c 'GCAAGTCCAACTTCAGG' *.fastq > count/B5023_Fm0.txt
agrep -0 -c 'GTCTCATAGTGACAGTG' *.fastq > count/B5025_Fm0.txt
agrep -0 -c 'GTTCGTAATCAGTAACG' *.fastq > count/B5026_Fm0.txt
agrep -0 -c 'ATATCTCCGTGATTCGT' *.fastq > count/B5027_Fm0.txt
agrep -0 -c 'TTCCGTCTCCTTAGCTG' *.fastq > count/B5028_Fm0.txt
agrep -0 -c 'AGCTATGTAGCTGCTAC' *.fastq > count/B5029_Fm0.txt
agrep -0 -c 'GGATTCCGTAAGCCTGT' *.fastq > count/B5030_Fm0.txt
agrep -0 -c 'CTTCGCCAGTACCAGAG' *.fastq > count/B5031_Fm0.txt
agrep -0 -c 'GCGCCTATACAATTCAG' *.fastq > count/B5032_Fm0.txt
agrep -0 -c 'CCTTCACCTAAGTAGGC' *.fastq > count/B5034_Fm0.txt
agrep -0 -c 'ACGGGTAAATTCCTCTG' *.fastq > count/B5035_Fm0.txt
agrep -0 -c 'ATCCGGTTCAAGTCGAG' *.fastq > count/B5037_Fm0.txt
agrep -0 -c 'AAGGCGTACATGGTAAG' *.fastq > count/B5038_Fm0.txt
agrep -0 -c 'AGGGTCCTTCAGGCAAT' *.fastq > count/B5040_Fm0.txt
agrep -0 -c 'GAGAAACCGTACAAAAG' *.fastq > count/B5041_Fm0.txt
agrep -0 -c 'TCTGGGCCCCGAATAAA' *.fastq > count/B5042_Fm0.txt
agrep -0 -c 'TTCAGAAGGTGTCCATG' *.fastq > count/B5045_Fm0.txt
agrep -0 -c 'GCTCGGTCACTACAGGT' *.fastq > count/B5046_Fm0.txt
agrep -0 -c 'GTTAGCATGTGTAATGC' *.fastq > count/B5048_Fm0.txt
agrep -0 -c 'GACACGCTTATGCAATT' *.fastq > count/B5049_Fm0.txt
agrep -0 -c 'ATGCTCCTTATGTCCCC' *.fastq > count/B5050_Fm0.txt
agrep -0 -c 'GATTGTGGTTTCTGGAC' *.fastq > count/B5051_Fm0.txt
agrep -0 -c 'TGATACGAGAGCTTCCT' *.fastq > count/B5052_Fm0.txt
agrep -0 -c 'GAGGGCAGGACCAATTT' *.fastq > count/B5053_Fm0.txt
agrep -0 -c 'GACTAAGATCCGGCAGA' *.fastq > count/B5054_Fm0.txt
agrep -0 -c 'ACTGGGTGTCGTTGGCA' *.fastq > count/B5055_Fm0.txt
agrep -0 -c 'AGTGCTGGGCCAAAAAA' *.fastq > count/B5058_Fm0.txt
agrep -0 -c 'TAGTTCGATAAAGCCGC' *.fastq > count/B5062_Fm0.txt
agrep -0 -c 'CGCGATTACTAAGTCGA' *.fastq > count/B5063_Fm0.txt
agrep -0 -c 'CTAATGCGTCGATTACC' *.fastq > count/B5064_Fm0.txt
agrep -0 -c 'AGTTAAGCGGGCCTTCT' *.fastq > count/B5065_Fm0.txt
agrep -0 -c 'TTAACCCACTGGCAGTA' *.fastq > count/B5066_Fm0.txt
agrep -0 -c 'TGTACGTGCCCTCAGAC' *.fastq > count/B5067_Fm0.txt
agrep -0 -c 'GTCGGGTGTTATGTTGC' *.fastq > count/B5068_Fm0.txt
agrep -0 -c 'GATGGAACGGGAACCAC' *.fastq > count/B5069_Fm0.txt
agrep -0 -c 'CACGGGAAATGCAAGAT' *.fastq > count/B5070_Fm0.txt
agrep -0 -c 'CATCAGTCATCCAGCGA' *.fastq > count/B5071_Fm0.txt
agrep -0 -c 'AGAGTCCACTCGGTACA' *.fastq > count/B5072_Fm0.txt
agrep -0 -c 'ATGCGTAACAGGGCGGA' *.fastq > count/B5075_Fm0.txt
agrep -0 -c 'TGAGTACAGTACCTGGG' *.fastq > count/B5076_Fm0.txt
agrep -0 -c 'GAGGTAGTACTTCGGCT' *.fastq > count/B5077_Fm0.txt
agrep -0 -c 'ACGATGTACACCCTGCG' *.fastq > count/B5078_Fm0.txt
agrep -0 -c 'GCAATAACGTGTCCAAG' *.fastq > count/B5079_Fm0.txt
agrep -0 -c 'CAGATGGATACGTGCAA' *.fastq > count/B5080_Fm0.txt
agrep -0 -c 'CGTGCCAGGTCCACAAA' *.fastq > count/B5081_Fm0.txt
agrep -0 -c 'AAGCATATCTCTCGGGT' *.fastq > count/B5083_Fm0.txt
agrep -0 -c 'TGACCGGGAGGAATACT' *.fastq > count/B5084_Fm0.txt
agrep -0 -c 'ATCTACCACTCCCTATT' *.fastq > count/B5085_Fm0.txt
agrep -0 -c 'GGAATGCCTCCGATCAC' *.fastq > count/B5086_Fm0.txt
agrep -0 -c 'CCTACGGATAGTACCAG' *.fastq > count/B5087_Fm0.txt
agrep -0 -c 'CGCATGGTTTAATTCCT' *.fastq > count/B5088_Fm0.txt
agrep -0 -c 'TCTTGATGCATAGCTGG' *.fastq > count/B5089_Fm0.txt
agrep -0 -c 'GGCAACTTTAGGATCTG' *.fastq > count/B5090_Fm0.txt
agrep -0 -c 'TATTCGCTTTCCGGACA' *.fastq > count/B5091_Fm0.txt
agrep -0 -c 'CGGGAATCCAGTTATAC' *.fastq > count/B5092_Fm0.txt
agrep -0 -c 'CTTCGACCGCATAGTCG' *.fastq > count/B5093_Fm0.txt
agrep -0 -c 'TTATTCGCTGAAGCCTG' *.fastq > count/B5094_Fm0.txt
agrep -0 -c 'GTGAACTGAGCGGAGTT' *.fastq > count/B5095_Fm0.txt
agrep -0 -c 'CTGTATCGCAGGACGAC' *.fastq > count/B5096_Fm0.txt
agrep -0 -c 'CCGTTCCCAAACCCCTT' *.fastq > count/B5097_Fm0.txt
agrep -0 -c 'GCCGATGTCCTCGACTA' *.fastq > count/B5098_Fm0.txt
agrep -0 -c 'CACATTAGATCCGCGGC' *.fastq > count/B5099_Fm0.txt
agrep -0 -c 'GTCATCGATTCTAGCTC' *.fastq > count/B5100_Fm0.txt
agrep -0 -c 'TTATGGGGTCTTCATCG' *.fastq > count/B5101_Fm0.txt
agrep -0 -c 'TGCACTTTGCTTGGAAG' *.fastq > count/B5102_Fm0.txt
agrep -0 -c 'CGTGATCTTGCATATAC' *.fastq > count/B5103_Fm0.txt
agrep -0 -c 'GTTGAATCCGCACTACG' *.fastq > count/B5104_Fm0.txt
agrep -0 -c 'ACGGTGCGGAGAAAGTT' *.fastq > count/B5106_Fm0.txt
agrep -0 -c 'ATTAGGCGAGAAGTGCG' *.fastq > count/B5107_Fm0.txt
agrep -0 -c 'GTACTGACAGAGCACTC' *.fastq > count/B5108_Fm0.txt
agrep -0 -c 'TAGGGCGAATGACGTGC' *.fastq > count/B5109_Fm0.txt
agrep -0 -c 'CTAATCCTTCCGAGTAG' *.fastq > count/B5110_Fm0.txt
agrep -0 -c 'TATCGAGACGATACACA' *.fastq > count/B5111_Fm0.txt
agrep -0 -c 'GACGGCGCATAGGCAAA' *.fastq > count/B5113_Fm0.txt
agrep -0 -c 'TTACGGCTAGCGAGGTT' *.fastq > count/B5114_Fm0.txt
agrep -0 -c 'CGAGAACGTCTTAGCCG' *.fastq > count/B5115_Fm0.txt
agrep -0 -c 'TACTGGAATCGCGTTGC' *.fastq > count/B5116_Fm0.txt
agrep -0 -c 'ACGTGATGTCCGCAGCT' *.fastq > count/B5118_Fm0.txt
agrep -0 -c 'TTCAGTTGCGGAAACCT' *.fastq > count/B5119_Fm0.txt
agrep -0 -c 'CACATGAACAGCCGCGA' *.fastq > count/B5120_Fm0.txt
agrep -0 -c 'TCGAGTCACGGTAATGG' *.fastq > count/B5121_Fm0.txt
agrep -0 -c 'AGCCAGCTCCCTAATGT' *.fastq > count/B5122_Fm0.txt
agrep -0 -c 'TTCCGGTTTCGCTACGA' *.fastq > count/B5123_Fm0.txt
agrep -0 -c 'CACGGTGTTCGAAGGAT' *.fastq > count/B5124_Fm0.txt
agrep -0 -c 'CTCGGGCGCTTGTATTG' *.fastq > count/B5125_Fm0.txt
agrep -0 -c 'TCTATTTGCGCGGGAGT' *.fastq > count/B5126_Fm0.txt
agrep -0 -c 'TATAGCCGAGCTCCCTG' *.fastq > count/B5127_Fm0.txt
agrep -0 -c 'TCTTCACAGGCTGCAAG' *.fastq > count/B5128_Fm0.txt
agrep -0 -c 'GTATCTATTCGTTGGAG' *.fastq > count/B5129_Fm0.txt
agrep -0 -c 'AAGTACCGTACTCGAAG' *.fastq > count/B5131_Fm0.txt
agrep -0 -c 'AGAGCGTACCTTATGTT' *.fastq > count/B5132_Fm0.txt
agrep -0 -c 'TACACGATCTGGTCTAG' *.fastq > count/B5133_Fm0.txt
agrep -0 -c 'ATATTAGCTGTCCGATG' *.fastq > count/B5134_Fm0.txt
agrep -0 -c 'CAAATACAGATCGATGC' *.fastq > count/B5135_Fm0.txt
agrep -0 -c 'TAACAACCCGAAGTACT' *.fastq > count/B5136_Fm0.txt
agrep -0 -c 'CGCATATCTAGGGTCGT' *.fastq > count/B5137_Fm0.txt
agrep -0 -c 'GTGTGGTCTCATTGGCG' *.fastq > count/B5138_Fm0.txt
agrep -0 -c 'CGATCCCGTTATCGGAA' *.fastq > count/B5139_Fm0.txt
agrep -0 -c 'AGCCAGTTTGGAGTCGG' *.fastq > count/B5140_Fm0.txt
agrep -0 -c 'GGTTATTGACGACGAAA' *.fastq > count/B5141_Fm0.txt
agrep -0 -c 'GGTATAATACATGCCAC' *.fastq > count/B5142_Fm0.txt
agrep -0 -c 'ACAGTCGGTGATCTTTC' *.fastq > count/B5144_Fm0.txt
agrep -0 -c 'GCGCAGTGCTATATCGG' *.fastq > count/B5147_Fm0.txt
agrep -0 -c 'CAAAAGCCGACCCAACG' *.fastq > count/B5148_Fm0.txt
agrep -0 -c 'ACTGCAGTGATGTACGA' *.fastq > count/B5149_Fm0.txt
agrep -0 -c 'ACACGGGTGCTCAAATG' *.fastq > count/B5150_Fm0.txt
agrep -0 -c 'CCTGTGTAGTCTCACGT' *.fastq > count/B5151_Fm0.txt
agrep -0 -c 'CTAACACGGTGTCGTAC' *.fastq > count/B5152_Fm0.txt
agrep -0 -c 'CCTTGCCCGCTACTGAA' *.fastq > count/B5153_Fm0.txt
agrep -0 -c 'AGCGCGTGAGGAAGTAT' *.fastq > count/B5154_Fm0.txt
agrep -0 -c 'TTGTCGCTCTGTAGTAA' *.fastq > count/B5155_Fm0.txt
agrep -0 -c 'CTGTATCGAACTGCGGG' *.fastq > count/B5156_Fm0.txt
agrep -0 -c 'TAATAACGATGCCGGTC' *.fastq > count/B5158_Fm0.txt
agrep -0 -c 'TTAGCCCGCCATTTAAC' *.fastq > count/B5159_Fm0.txt
agrep -0 -c 'GTTAGTCTCCCCAACGG' *.fastq > count/B5160_Fm0.txt
agrep -0 -c 'ACGTTGACCGGTACGCA' *.fastq > count/B5161_Fm0.txt
agrep -0 -c 'GTAGGCTTGACCCAAGT' *.fastq > count/B5163_Fm0.txt
agrep -0 -c 'TAGAAACCTCGCTCATG' *.fastq > count/B5164_Fm0.txt
agrep -0 -c 'GGCTTGTCTGCGGTGTT' *.fastq > count/B5166_Fm0.txt
agrep -0 -c 'GTCACAGACGTATAGTG' *.fastq > count/B5168_Fm0.txt
agrep -0 -c 'TATTGATGGTGGAGCTC' *.fastq > count/B5169_Fm0.txt
agrep -0 -c 'ACCCATGCGGCAATCTA' *.fastq > count/B5170_Fm0.txt
agrep -0 -c 'TGAACTTGCCTTAGATC' *.fastq > count/B5171_Fm0.txt
agrep -0 -c 'GATACCCGGACTGCACA' *.fastq > count/B5172_Fm0.txt
agrep -0 -c 'TTACCGACTAATACCTG' *.fastq > count/B5174_Fm0.txt
agrep -0 -c 'ACCGACTCTAAGGATGA' *.fastq > count/B5176_Fm0.txt
agrep -0 -c 'GTTTCTAGAACTGGTCT' *.fastq > count/B5177_Fm0.txt
agrep -0 -c 'GCACAGTTACAACGAGT' *.fastq > count/B5180_Fm0.txt
agrep -0 -c 'GCGGGTCTTTCAAATCC' *.fastq > count/B5182_Fm0.txt
agrep -0 -c 'TGACCAGTACGTTGCTT' *.fastq > count/B5183_Fm0.txt
agrep -0 -c 'TACTCCAAGACCAACAG' *.fastq > count/B5184_Fm0.txt
agrep -0 -c 'ACACAAAGTGTCCTGGC' *.fastq > count/B5185_Fm0.txt
agrep -0 -c 'GAGCTGACAACTTTGCA' *.fastq > count/B5188_Fm0.txt
agrep -0 -c 'TGACGGAGATTACGTGT' *.fastq > count/B5189_Fm0.txt
agrep -0 -c 'CTTACTAGTCAGTACGG' *.fastq > count/B5190_Fm0.txt
agrep -0 -c 'CGGGCTTCGATATCAAT' *.fastq > count/B5191_Fm0.txt
agrep -0 -c 'AACCTTCAAAGTACGTC' *.fastq > count/B5192_Fm0.txt
agrep -0 -c 'TTACACGCATGCTGGAC' *.fastq > count/B5194_Fm0.txt
agrep -0 -c 'CATTTATCTCACCGGAG' *.fastq > count/B5195_Fm0.txt
agrep -0 -c 'TCATCAGTTGGTGTAAG' *.fastq > count/B5196_Fm0.txt
agrep -0 -c 'GTTTCGGACTTCAGCTT' *.fastq > count/B5197_Fm0.txt
agrep -0 -c 'GGTGTAACCCAGTATAG' *.fastq > count/B5198_Fm0.txt
agrep -0 -c 'AGACGTGTTCCATAGAT' *.fastq > count/B5199_Fm0.txt
agrep -0 -c 'AGGTCGTGGTCTGTTCC' *.fastq > count/B5200_Fm0.txt
agrep -0 -c 'GGGAGGACATCAGCACA' *.fastq > count/B5201_Fm0.txt
agrep -0 -c 'AATCTGCACGGGGTGGA' *.fastq > count/B5202_Fm0.txt
agrep -0 -c 'AGGGGACGCTGTCTAGT' *.fastq > count/B5203_Fm0.txt
agrep -0 -c 'GATACTACCTGAACGAA' *.fastq > count/B5204_Fm0.txt
agrep -0 -c 'GTATACGACCATACGCG' *.fastq > count/B5205_Fm0.txt
agrep -0 -c 'ATCATGAGATGACGGTC' *.fastq > count/B5206_Fm0.txt
agrep -0 -c 'CAGACGGACCTGAAACT' *.fastq > count/B5207_Fm0.txt
agrep -0 -c 'AGAGTAGTGAGCATTCA' *.fastq > count/B5209_Fm0.txt
agrep -0 -c 'CCTGGCATCGTTCCAAT' *.fastq > count/B5210_Fm0.txt
agrep -0 -c 'TTGGACTCTAATTTCGC' *.fastq > count/B5211_Fm0.txt
agrep -0 -c 'TGCCTCTTAACCGGCAA' *.fastq > count/B5212_Fm0.txt
agrep -0 -c 'CACGATATTCTGAGTTG' *.fastq > count/B5214_Fm0.txt
agrep -0 -c 'CGTTTGCTCCCAGAGTC' *.fastq > count/B5215_Fm0.txt
agrep -0 -c 'AAGGTAAGTGTAGACAG' *.fastq > count/B5216_Fm0.txt
agrep -0 -c 'AGAGACTAGATGTTGTG' *.fastq > count/B5217_Fm0.txt
agrep -0 -c 'ACGGGGTAGACAATAAA' *.fastq > count/B5218_Fm0.txt
agrep -0 -c 'CAGCATAGGACGGGAAC' *.fastq > count/B5220_Fm0.txt
agrep -0 -c 'TGACACTCTAAACACGT' *.fastq > count/B5221_Fm0.txt
agrep -0 -c 'ATAGGGTGGCACCAAGT' *.fastq > count/B5222_Fm0.txt
agrep -0 -c 'GGGCAAATCTACAGGTG' *.fastq > count/B5224_Fm0.txt
agrep -0 -c 'TGCATTCCCGGACTTAT' *.fastq > count/B5225_Fm0.txt
agrep -0 -c 'GGGAGTTTAGCGTTTAT' *.fastq > count/B5227_Fm0.txt
agrep -0 -c 'GATCCATACGTCCAGAT' *.fastq > count/B5228_Fm0.txt
agrep -0 -c 'CAAGCTAGTACGCGCAT' *.fastq > count/B5229_Fm0.txt
agrep -0 -c 'AGTTCAAGCGACGGTCA' *.fastq > count/B5230_Fm0.txt
agrep -0 -c 'CAGGCACGACTATCTTT' *.fastq > count/B5231_Fm0.txt
agrep -0 -c 'ATCGACATTCTAACCCG' *.fastq > count/B5232_Fm0.txt
agrep -0 -c 'CGATGGATAGCTAGGGA' *.fastq > count/B5233_Fm0.txt
agrep -0 -c 'AGTGGGGTACCGTCATT' *.fastq > count/B5234_Fm0.txt
agrep -0 -c 'CCGACAAATCGGCATTA' *.fastq > count/B5236_Fm0.txt
agrep -0 -c 'GATAGGGTAACGGTTAG' *.fastq > count/B5239_Fm0.txt
agrep -0 -c 'TAGCCCCCATTCGCGAA' *.fastq > count/B5240_Fm0.txt
agrep -0 -c 'ACCACCCTACTGAGGCT' *.fastq > count/B5241_Fm0.txt
agrep -0 -c 'TCGTAGGCGTAACCTGA' *.fastq > count/B5243_Fm0.txt
agrep -0 -c 'TGACGGATGCCAAATTG' *.fastq > count/B5244_Fm0.txt
agrep -0 -c 'GACCGTTCCCCTCAACT' *.fastq > count/B5245_Fm0.txt
agrep -0 -c 'GGCCTCCTCATTACCAG' *.fastq > count/B5246_Fm0.txt
agrep -0 -c 'GACTATCACATCGGGCG' *.fastq > count/B5248_Fm0.txt
agrep -0 -c 'GTACTGACCAATAAGCT' *.fastq > count/B5249_Fm0.txt
agrep -0 -c 'GACGCACTTGTCTATAT' *.fastq > count/B5250_Fm0.txt
agrep -0 -c 'GAGAGAATTCCGTACGC' *.fastq > count/B5252_Fm0.txt
agrep -0 -c 'TCACACAATTGTGTCAG' *.fastq > count/B5253_Fm0.txt
agrep -0 -c 'CGCCCGATCACTTAGTC' *.fastq > count/B5254_Fm0.txt
agrep -0 -c 'ACGGTAAACCTTGGTTC' *.fastq > count/B5255_Fm0.txt
agrep -0 -c 'GGGTGGCATCTAGATAT' *.fastq > count/B5256_Fm0.txt
agrep -0 -c 'ATGACATATGTGCGTAG' *.fastq > count/B5257_Fm0.txt
agrep -0 -c 'TCGGAAACAGGCACTGG' *.fastq > count/B5258_Fm0.txt
agrep -0 -c 'ACTGGGCAATCACCTTA' *.fastq > count/B5259_Fm0.txt
agrep -0 -c 'TCTGGTATCTATCAGGA' *.fastq > count/B5261_Fm0.txt
agrep -0 -c 'TGCGGTGTGGGTAGTTA' *.fastq > count/B5265_Fm0.txt
agrep -0 -c 'ACCTCTCATGTCAAGCG' *.fastq > count/B5266_Fm0.txt
agrep -0 -c 'CAGGAAGATATGGGCGG' *.fastq > count/B5267_Fm0.txt
agrep -0 -c 'TACCGAAAAGGTTGGTT' *.fastq > count/B5268_Fm0.txt
agrep -0 -c 'CCTAATCTTAATGCCCT' *.fastq > count/B5269_Fm0.txt
agrep -0 -c 'AGAGGCGAAATCGCAGA' *.fastq > count/B5270_Fm0.txt
agrep -0 -c 'CGTGGGCGGACAGATAA' *.fastq > count/B5271_Fm0.txt
agrep -0 -c 'TTGTTAACGGCGCTCTA' *.fastq > count/B5272_Fm0.txt
agrep -0 -c 'GATACGAAGGTCCTGCT' *.fastq > count/B5273_Fm0.txt
agrep -0 -c 'CACCGAACTTACGTATC' *.fastq > count/B5274_Fm0.txt
agrep -0 -c 'AAGAGACGATGTCTCTT' *.fastq > count/B5275_Fm0.txt
agrep -0 -c 'GCGTGTGGCAGCTAAGA' *.fastq > count/B5276_Fm0.txt
agrep -0 -c 'GTATACCCGAGATACTA' *.fastq > count/B5278_Fm0.txt
agrep -0 -c 'GCTCACATTCATACTCC' *.fastq > count/B5280_Fm0.txt
agrep -0 -c 'GTAATGACATGTTCAGC' *.fastq > count/B5281_Fm0.txt
agrep -0 -c 'TAGTCAAGCGCCAATAT' *.fastq > count/B5282_Fm0.txt
agrep -0 -c 'TCTTCTGGTTAGTTCCG' *.fastq > count/B5283_Fm0.txt
agrep -0 -c 'CGATTCCCTAGTTGGCG' *.fastq > count/B5284_Fm0.txt
agrep -0 -c 'GTCGTATTACCTCCGGG' *.fastq > count/B5285_Fm0.txt
agrep -0 -c 'CGAGGCAGCACCTTTAC' *.fastq > count/B5286_Fm0.txt
agrep -0 -c 'GTGTAATTCGGGGTGGA' *.fastq > count/B5289_Fm0.txt
agrep -0 -c 'TATCGATCTGGGCCAGC' *.fastq > count/B5290_Fm0.txt
agrep -0 -c 'GGTGGAGCGTGACATTA' *.fastq > count/B5294_Fm0.txt
agrep -0 -c 'CCAACTCTGCCACTGTA' *.fastq > count/B5295_Fm0.txt
agrep -0 -c 'CTAGGGGCTCTCAATTC' *.fastq > count/B5296_Fm0.txt
agrep -0 -c 'GCGTTTGGCTTCCTATT' *.fastq > count/B5297_Fm0.txt
agrep -0 -c 'TTAGACGGTATCAGGGG' *.fastq > count/B5298_Fm0.txt
agrep -0 -c 'CCGTATGGATCGATGAA' *.fastq > count/B5299_Fm0.txt
agrep -0 -c 'CACTTTCTTTTTGGTCG' *.fastq > count/B5300_Fm0.txt
agrep -0 -c 'CGGCGGGCTTTTAACCA' *.fastq > count/B5301_Fm0.txt
agrep -0 -c 'TCGATGGACTTCTCGGC' *.fastq > count/B5302_Fm0.txt
agrep -0 -c 'TTCCCTCCATTGCTAGC' *.fastq > count/B5303_Fm0.txt
agrep -0 -c 'ACCCGTTGCGCACTTGA' *.fastq > count/B5304_Fm0.txt
agrep -0 -c 'CGCCAGTTTCTACAGGG' *.fastq > count/B5305_Fm0.txt
agrep -0 -c 'GGCCTTAACAACGACGC' *.fastq > count/B5306_Fm0.txt
agrep -0 -c 'CGTCTGAACTGTGAGCG' *.fastq > count/B5307_Fm0.txt
agrep -0 -c 'GCATGCACCTGCTGAAG' *.fastq > count/B5308_Fm0.txt
agrep -0 -c 'AGAGCTGGCTTTGGCGA' *.fastq > count/B5310_Fm0.txt
agrep -0 -c 'AGCTTTAGTTACGTTCC' *.fastq > count/B5311_Fm0.txt
agrep -0 -c 'GAGCGTAGGTAAGCGTT' *.fastq > count/B5312_Fm0.txt
agrep -0 -c 'CACGATCGTGACGGAGA' *.fastq > count/B5313_Fm0.txt
agrep -0 -c 'TCTTGTGCCACTCATGA' *.fastq > count/B5314_Fm0.txt
agrep -0 -c 'ACAGCATAGTCCAGGAC' *.fastq > count/B5315_Fm0.txt
agrep -0 -c 'AGACGCTAGTTCCTATT' *.fastq > count/B5316_Fm0.txt
agrep -0 -c 'CCTCGTTCAGGAAGTAT' *.fastq > count/B5317_Fm0.txt
agrep -0 -c 'CACGTTAGTGCATACGA' *.fastq > count/B5318_Fm0.txt
agrep -0 -c 'GTACGTCCCGACTGTCA' *.fastq > count/B5320_Fm0.txt
agrep -0 -c 'CTAACTCTCGCTCCGGA' *.fastq > count/B5323_Fm0.txt
agrep -0 -c 'ATTAGCGAGTGACTGGA' *.fastq > count/B5324_Fm0.txt
agrep -0 -c 'GCGAGACAATGGTCAAC' *.fastq > count/B5325_Fm0.txt
agrep -0 -c 'CCTGTTTAAGAGTAGTC' *.fastq > count/B5326_Fm0.txt
agrep -0 -c 'CGCAGTATACTTTCAGA' *.fastq > count/B5327_Fm0.txt
agrep -0 -c 'ACTGACCCTTCGTTTGT' *.fastq > count/B5330_Fm0.txt
agrep -0 -c 'CGTCATTCCCGTACATA' *.fastq > count/B5331_Fm0.txt
agrep -0 -c 'GATTGAATGGTAACCGA' *.fastq > count/B5332_Fm0.txt
agrep -0 -c 'GCGCGCTCTCACTTTCT' *.fastq > count/B5333_Fm0.txt
agrep -0 -c 'AATACGTCGAGAGGACT' *.fastq > count/B5334_Fm0.txt
agrep -0 -c 'CGGTCCGACATATCATA' *.fastq > count/B5335_Fm0.txt
agrep -0 -c 'GGTTAGCCATTTGACCA' *.fastq > count/B5338_Fm0.txt
agrep -0 -c 'ATGCCAAGGCAGTCCAC' *.fastq > count/B5340_Fm0.txt
agrep -0 -c 'GTGATCTTGACTTCTCA' *.fastq > count/B5341_Fm0.txt
agrep -0 -c 'CATCCCATCCGGTAAAT' *.fastq > count/B5342_Fm0.txt
agrep -0 -c 'AGGTATTGGAAGCGATT' *.fastq > count/B5343_Fm0.txt
agrep -0 -c 'TTCGAGGCTAGCAGAAC' *.fastq > count/B5344_Fm0.txt
agrep -0 -c 'GACCATTGTGCAAGAAA' *.fastq > count/B5345_Fm0.txt
agrep -0 -c 'ATGCTTCAGTGTGATCT' *.fastq > count/B5347_Fm0.txt
agrep -0 -c 'TAGCAATGGAGCGCAGG' *.fastq > count/B5348_Fm0.txt
agrep -0 -c 'CCTAAAGACGAGTAACA' *.fastq > count/B5349_Fm0.txt
agrep -0 -c 'GTGATGATTGACACCCC' *.fastq > count/B5350_Fm0.txt
agrep -0 -c 'AATGGTTCGTACCGTGT' *.fastq > count/B5352_Fm0.txt
agrep -0 -c 'ACCTTGATCGCTGCACA' *.fastq > count/B5353_Fm0.txt
agrep -0 -c 'TCTTGCAACCCTCCAGT' *.fastq > count/B5355_Fm0.txt
agrep -0 -c 'AAGTACCCCTTGATTCC' *.fastq > count/B5356_Fm0.txt
agrep -0 -c 'ACGGACGTCCACTACTG' *.fastq > count/B5357_Fm0.txt
agrep -0 -c 'AACTTCTTCGCTTGCAG' *.fastq > count/B5358_Fm0.txt
agrep -0 -c 'TAAGACGGAACGCCTTT' *.fastq > count/B5359_Fm0.txt
agrep -0 -c 'ACATGAATTTTGGGCGA' *.fastq > count/B5360_Fm0.txt
agrep -0 -c 'TGGACGCGAATGGCTAA' *.fastq > count/B5361_Fm0.txt
agrep -0 -c 'GTGACATTGTCGATTAC' *.fastq > count/B5362_Fm0.txt
agrep -0 -c 'TCGCCGTTTCATATCGG' *.fastq > count/B5363_Fm0.txt
agrep -0 -c 'GCTCCACAAGAGCTCTT' *.fastq > count/B5365_Fm0.txt
agrep -0 -c 'CTTGCGATCAACTTCCC' *.fastq > count/B5366_Fm0.txt
agrep -0 -c 'TAGCTCCGCTAACCTTC' *.fastq > count/B5367_Fm0.txt
agrep -0 -c 'GCGTGATATGCAGTACT' *.fastq > count/B5368_Fm0.txt
agrep -0 -c 'CGGTATAGAATCGGGTG' *.fastq > count/B5369_Fm0.txt
agrep -0 -c 'ATGGAAGCTCGCCTACT' *.fastq > count/B5370_Fm0.txt
agrep -0 -c 'ACGTACAACTCAAACCG' *.fastq > count/B5371_Fm0.txt
agrep -0 -c 'TCGACAGCGAATCTCTT' *.fastq > count/B5372_Fm0.txt
agrep -0 -c 'CCTAAATAAGTGGCCAG' *.fastq > count/B5374_Fm0.txt
agrep -0 -c 'ACTCCCACTTGACTAAG' *.fastq > count/B5375_Fm0.txt
agrep -0 -c 'GTACTGCCAGAGCTGAC' *.fastq > count/B5376_Fm0.txt
agrep -0 -c 'CACGTTGGTACCAATGG' *.fastq > count/B5377_Fm0.txt
agrep -0 -c 'GATGTTGATGACGTTTC' *.fastq > count/B5378_Fm0.txt
agrep -0 -c 'ATATGGCCCCAGCCTAC' *.fastq > count/B5379_Fm0.txt
agrep -0 -c 'GGACTCCAGCCATGAAA' *.fastq > count/B5380_Fm0.txt
agrep -0 -c 'GACTCTAAATGACCCCG' *.fastq > count/B5381_Fm0.txt
agrep -0 -c 'AGCAGATGACTCCACGT' *.fastq > count/B5382_Fm0.txt
agrep -0 -c 'TTCATGTGCCTCGTTAC' *.fastq > count/B5384_Fm0.txt
agrep -0 -c 'TAGGAATGTCCCGGGGT' *.fastq > count/B5385_Fm0.txt
agrep -0 -c 'CACATCCACTAGATCTA' *.fastq > count/B5386_Fm0.txt
agrep -0 -c 'TATCGTAAGAAGGCGTA' *.fastq > count/B5387_Fm0.txt
agrep -0 -c 'CGAAGGAGACCTATCGA' *.fastq > count/B5388_Fm0.txt
agrep -0 -c 'CGCCAAAATGTACCGAT' *.fastq > count/B5389_Fm0.txt
agrep -0 -c 'TAGACGTCTGTTAGCTC' *.fastq > count/B5390_Fm0.txt
agrep -0 -c 'GGCGAACTCGATTTTGC' *.fastq > count/B5391_Fm0.txt
agrep -0 -c 'TAAGACCCTTCCTAATC' *.fastq > count/B5393_Fm0.txt
agrep -0 -c 'AGACGTACCTCGCGGAA' *.fastq > count/B5394_Fm0.txt
agrep -0 -c 'CGCTCATAATTGGATCT' *.fastq > count/B5395_Fm0.txt
agrep -0 -c 'AACTGCCGCATAGGTAG' *.fastq > count/B5397_Fm0.txt
agrep -0 -c 'CGTTATAATTGACCTCC' *.fastq > count/B5398_Fm0.txt
agrep -0 -c 'TAATGCCAGTGATCCTG' *.fastq > count/B5399_Fm0.txt
agrep -0 -c 'GCTTGGTACCGATGAAG' *.fastq > count/B5400_Fm0.txt
agrep -0 -c 'GACGATGTGCCTCACAA' *.fastq > count/B5401_Fm0.txt
agrep -0 -c 'TCTGTTGTCGTTGATCC' *.fastq > count/B5402_Fm0.txt
agrep -0 -c 'CGACAACACGCAATCAC' *.fastq > count/B5405_Fm0.txt
agrep -0 -c 'ACACTTCGTTGGACTAC' *.fastq > count/B5406_Fm0.txt
agrep -0 -c 'CCAGGCCAATTGTCGTC' *.fastq > count/B5407_Fm0.txt
agrep -0 -c 'ATCCTCTTATCGTAGGT' *.fastq > count/B5408_Fm0.txt
agrep -0 -c 'CGACCGGAAAACTCGCT' *.fastq > count/B5410_Fm0.txt
agrep -0 -c 'TGTAACCGCTAACCGAG' *.fastq > count/B5411_Fm0.txt
agrep -0 -c 'TGATGAGGTCCGCTAAC' *.fastq > count/B5412_Fm0.txt
agrep -0 -c 'GCCATTCATCGATCTTC' *.fastq > count/B5413_Fm0.txt
agrep -0 -c 'AGTGTATCTGTGCTTCG' *.fastq > count/B5414_Fm0.txt
agrep -0 -c 'TCGTCCTTTAGTACAGA' *.fastq > count/B5415_Fm0.txt
agrep -0 -c 'GTCACAGGTCGATGTTT' *.fastq > count/B5416_Fm0.txt
agrep -0 -c 'AAGGGATGAACACACGC' *.fastq > count/B5417_Fm0.txt
agrep -0 -c 'ATATGTCTAAGGGGTCA' *.fastq > count/B5418_Fm0.txt
agrep -0 -c 'TACTCTTTTACGTGCCA' *.fastq > count/B5420_Fm0.txt
agrep -0 -c 'GCAGTTTGCAATTCGCG' *.fastq > count/B5421_Fm0.txt
agrep -0 -c 'GGATAACGACATCGAAG' *.fastq > count/B5422_Fm0.txt
agrep -0 -c 'CGGTTATAGCGCAACGA' *.fastq > count/B5423_Fm0.txt
agrep -0 -c 'ATGGTGTTCAAGACCCG' *.fastq > count/B5424_Fm0.txt
agrep -0 -c 'CCTATCACGACCGCAAA' *.fastq > count/B5425_Fm0.txt
agrep -0 -c 'TGAAGTACCAGTTTAGC' *.fastq > count/B5426_Fm0.txt
agrep -0 -c 'ACGCGTTTACCACTCCC' *.fastq > count/B5429_Fm0.txt
agrep -0 -c 'GACGTAAATTCGGCCTT' *.fastq > count/B5431_Fm0.txt
agrep -0 -c 'TGAATCTGGCGAGAATC' *.fastq > count/B5432_Fm0.txt
agrep -0 -c 'ACGTTGCGTGCTCTGGA' *.fastq > count/B5433_Fm0.txt
agrep -0 -c 'CAATTCGTTCCTCTCAC' *.fastq > count/B5435_Fm0.txt
agrep -0 -c 'CTGAGTTACCTCAGAGG' *.fastq > count/B5436_Fm0.txt
agrep -0 -c 'CATTAGTTCCCACGGAT' *.fastq > count/B5437_Fm0.txt
agrep -0 -c 'GCTTCAATTGTGGAAGG' *.fastq > count/B5439_Fm0.txt
agrep -0 -c 'ACGAGATTCATTGGCGT' *.fastq > count/B5440_Fm0.txt
agrep -0 -c 'CCGAGTATTGTCCCTCT' *.fastq > count/B5442_Fm0.txt
agrep -0 -c 'ATCGCCCTAGCATAGAG' *.fastq > count/B5443_Fm0.txt
agrep -0 -c 'CGCAAATCACAACGCAG' *.fastq > count/B5444_Fm0.txt
agrep -0 -c 'TACCATACCCCGGATAG' *.fastq > count/B5445_Fm0.txt
agrep -0 -c 'TCATCCGCTTCAGCATA' *.fastq > count/B5447_Fm0.txt
agrep -0 -c 'AAGTTTATGCAGACCGC' *.fastq > count/B5448_Fm0.txt
agrep -0 -c 'CAGGTAAGTTAGAAAGG' *.fastq > count/B5449_Fm0.txt
agrep -0 -c 'TAGCATTCAAGTCGCTC' *.fastq > count/B5450_Fm0.txt
agrep -0 -c 'TTCTGGTTCGTGCAGTT' *.fastq > count/B5452_Fm0.txt
agrep -0 -c 'ATCCACGAATGTCGATC' *.fastq > count/B5453_Fm0.txt
agrep -0 -c 'AGGGAGACACACCTCGT' *.fastq > count/B5454_Fm0.txt
agrep -0 -c 'TGCCCACTCGATAGAGT' *.fastq > count/B5455_Fm0.txt
agrep -0 -c 'ATGGTTTAAAGGTGCGC' *.fastq > count/B5456_Fm0.txt
agrep -0 -c 'CTACTCACGTAGCCAGT' *.fastq > count/B5457_Fm0.txt
agrep -0 -c 'TATCCGCTATTGCCAAA' *.fastq > count/B5458_Fm0.txt
agrep -0 -c 'ATGGGAGTGACCACGTC' *.fastq > count/B5459_Fm0.txt
agrep -0 -c 'CATAAACGTGCTCCGGC' *.fastq > count/B5460_Fm0.txt
agrep -0 -c 'CTAAGAGCAGCGGAGAA' *.fastq > count/B5462_Fm0.txt
agrep -0 -c 'ACACAAGAGGGTTTCAT' *.fastq > count/B5463_Fm0.txt
agrep -0 -c 'ATTTGGAACTGCGGCAG' *.fastq > count/B5464_Fm0.txt
agrep -0 -c 'CGCTATTACTGAATCCG' *.fastq > count/B5465_Fm0.txt
agrep -0 -c 'CTCAACCGACTTTCCTT' *.fastq > count/B5466_Fm0.txt
agrep -0 -c 'TGAATGTCCCGAATCCG' *.fastq > count/B5467_Fm0.txt
agrep -0 -c 'GAGCTGTAACTCCCGCT' *.fastq > count/B5468_Fm0.txt
agrep -0 -c 'ACGCGTTAAGCCTCCAC' *.fastq > count/B5470_Fm0.txt
agrep -0 -c 'GCTGGATATTCAATGCT' *.fastq > count/B5471_Fm0.txt
agrep -0 -c 'ACGATCGGATCCTAGTC' *.fastq > count/B5472_Fm0.txt
agrep -0 -c 'TTAGCCGAGCGATAGGC' *.fastq > count/B5473_Fm0.txt
agrep -0 -c 'TAACGTAATCCGGTCCG' *.fastq > count/B5474_Fm0.txt
agrep -0 -c 'ACTTCGGAGTGTCCGAA' *.fastq > count/B5475_Fm0.txt
agrep -0 -c 'AAGTTGAAGTCCCTTCA' *.fastq > count/B5476_Fm0.txt
agrep -0 -c 'CTTGCTAGCGTGGGAAC' *.fastq > count/B5478_Fm0.txt
agrep -0 -c 'GCTAGGACACGACTTCC' *.fastq > count/B5479_Fm0.txt
agrep -0 -c 'ATCGCAAAACTAGGGTT' *.fastq > count/B5480_Fm0.txt
agrep -0 -c 'TTACCGGAGAGCAGACC' *.fastq > count/B5482_Fm0.txt
agrep -0 -c 'GCAGGTAACTATCGTTA' *.fastq > count/B5483_Fm0.txt
agrep -0 -c 'CAAAATGGTCACCGATT' *.fastq > count/B5484_Fm0.txt
agrep -0 -c 'CCACTGACGGTTTATCG' *.fastq > count/B5485_Fm0.txt
agrep -0 -c 'GACAATCAGGAAGAACT' *.fastq > count/B5486_Fm0.txt
agrep -0 -c 'CAGTGTAGAGATGCCGC' *.fastq > count/B5487_Fm0.txt
agrep -0 -c 'ACCTTAACTCCGTAGAA' *.fastq > count/B5488_Fm0.txt
agrep -0 -c 'CAGGTCAATACGGCATA' *.fastq > count/B5489_Fm0.txt
agrep -0 -c 'ACTTGCTGCCATATGGG' *.fastq > count/B5491_Fm0.txt
agrep -0 -c 'AGTCCCTGTTGGCATAT' *.fastq > count/B5492_Fm0.txt
agrep -0 -c 'GCCTGATGGTATGGAAC' *.fastq > count/B5493_Fm0.txt
agrep -0 -c 'ATACGATGGCTAAATGC' *.fastq > count/B5494_Fm0.txt
agrep -0 -c 'CAGTGTCGCACTAGATA' *.fastq > count/B5496_Fm0.txt
agrep -0 -c 'AGCCTGGTGGATGCGTA' *.fastq > count/B5497_Fm0.txt
agrep -0 -c 'TCGGCAATCGATTGGGT' *.fastq > count/B5500_Fm0.txt
agrep -0 -c 'CTTCACACCGGTTACGG' *.fastq > count/B5501_Fm0.txt
agrep -0 -c 'AGGTGTCAGTGCAAGCC' *.fastq > count/B5502_Fm0.txt
agrep -0 -c 'GAGCCGTAAAATCCAGA' *.fastq > count/B5503_Fm0.txt
agrep -0 -c 'TAAACTAAGGACGAGAG' *.fastq > count/B5504_Fm0.txt
agrep -0 -c 'CAATCCCTTCCGTATTT' *.fastq > count/B5505_Fm0.txt
agrep -0 -c 'CGGTTACCGCTATGTAC' *.fastq > count/B5507_Fm0.txt
agrep -0 -c 'GGTCAATAGCATACATG' *.fastq > count/B5508_Fm0.txt
agrep -0 -c 'TCGGATAAAGACGCTAT' *.fastq > count/B5510_Fm0.txt
agrep -0 -c 'GGTGCCCATTGATCATG' *.fastq > count/B5511_Fm0.txt
agrep -0 -c 'GGATCCTTCGGTGAAGC' *.fastq > count/B5512_Fm0.txt
agrep -0 -c 'CAATCATAGCAACACTC' *.fastq > count/B5513_Fm0.txt
agrep -0 -c 'GTCCAGCGTAAGACCTA' *.fastq > count/B5514_Fm0.txt
agrep -0 -c 'CGAATTCCTCGACACTT' *.fastq > count/B5515_Fm0.txt
agrep -0 -c 'CACCGAGTTGTGTAACG' *.fastq > count/B5516_Fm0.txt
agrep -0 -c 'GTGATTTAGTTCCCCAA' *.fastq > count/B5518_Fm0.txt
agrep -0 -c 'CATGTCAATCACTCCGC' *.fastq > count/B5520_Fm0.txt
agrep -0 -c 'TTCGTATCTACGCACGA' *.fastq > count/B5522_Fm0.txt
agrep -0 -c 'CTAAGGATTCGCATTGC' *.fastq > count/B5523_Fm0.txt
agrep -0 -c 'GTGCTGAAATCAATGCG' *.fastq > count/B5524_Fm0.txt
agrep -0 -c 'TATGTCTGTCGTAAGGG' *.fastq > count/B5525_Fm0.txt
agrep -0 -c 'CATGTAGTAAGTCTGGA' *.fastq > count/B5526_Fm0.txt
agrep -0 -c 'CGTGCAGCGAGACGTAA' *.fastq > count/B5527_Fm0.txt
agrep -0 -c 'CGGGCTGATTATCAAAA' *.fastq > count/B5530_Fm0.txt
agrep -0 -c 'GTTCGGGTAACGGGCAA' *.fastq > count/B5531_Fm0.txt
agrep -0 -c 'GTGAGCTTTAGTGAATC' *.fastq > count/B5532_Fm0.txt
agrep -0 -c 'AATCGGCTAGCGCAGCT' *.fastq > count/B5533_Fm0.txt
agrep -0 -c 'TACCAGGTGCTGATGTA' *.fastq > count/B5537_Fm0.txt
agrep -0 -c 'TTACACCTATCGAGGCC' *.fastq > count/B5538_Fm0.txt
agrep -0 -c 'GTCATGTTAGCGCATCC' *.fastq > count/B5539_Fm0.txt
agrep -0 -c 'CTGGGAGTCACTATGAT' *.fastq > count/B5540_Fm0.txt
agrep -0 -c 'AATAGGTTCCACCGCAT' *.fastq > count/B5541_Fm0.txt
agrep -0 -c 'CATAGTCACTCAATCCA' *.fastq > count/B5542_Fm0.txt
agrep -0 -c 'GATTAGGGCGCTTTACG' *.fastq > count/B5543_Fm0.txt
agrep -0 -c 'GTCGCCAACTCAAGTGG' *.fastq > count/B5544_Fm0.txt
agrep -0 -c 'ATCGGCCAGATAGCATC' *.fastq > count/B5545_Fm0.txt
agrep -0 -c 'CACGGCGAATTCACTGA' *.fastq > count/B5546_Fm0.txt
agrep -0 -c 'ATAGTTGCCGACTCTAG' *.fastq > count/B5549_Fm0.txt
agrep -0 -c 'AGCCAACACGCTTACTG' *.fastq > count/B5550_Fm0.txt
agrep -0 -c 'GTGCCTTCGTGTGTCTG' *.fastq > count/B5551_Fm0.txt
agrep -0 -c 'TAATGGTACGTATCCCG' *.fastq > count/B5552_Fm0.txt
agrep -0 -c 'GGAAGGTAATAGACGGC' *.fastq > count/B5553_Fm0.txt
agrep -0 -c 'GGCGGGTGTCTATCAAC' *.fastq > count/B5554_Fm0.txt
agrep -0 -c 'CAACCTCCAACACGGTG' *.fastq > count/B5555_Fm0.txt
agrep -0 -c 'GATCTATTCCTAGCCCA' *.fastq > count/B5556_Fm0.txt
agrep -0 -c 'ACTGCATGTTCTCCAAC' *.fastq > count/B5557_Fm0.txt
agrep -0 -c 'CTGAGTAGTGGTCAGGC' *.fastq > count/B5559_Fm0.txt
agrep -0 -c 'AGACACGAGTCCTACAA' *.fastq > count/B5560_Fm0.txt
agrep -0 -c 'TGCCCTTCGATTTAGAG' *.fastq > count/B5562_Fm0.txt
agrep -0 -c 'TGATCTAGTTACCCTCT' *.fastq > count/B5563_Fm0.txt
agrep -0 -c 'CCTAGCCATCGTCAATG' *.fastq > count/B5564_Fm0.txt
agrep -0 -c 'GCGAGAAAGAATTGCAG' *.fastq > count/B5565_Fm0.txt
agrep -0 -c 'AGCCTAGGATGCCTTTC' *.fastq > count/B5566_Fm0.txt
agrep -0 -c 'CTATGACACGGTATCTC' *.fastq > count/B5567_Fm0.txt
agrep -0 -c 'AGATTTCGAGCAGATAC' *.fastq > count/B5571_Fm0.txt
agrep -0 -c 'CTTCGTCCCACCACTAT' *.fastq > count/B5572_Fm0.txt
agrep -0 -c 'ACGTGTGAACTAATAGG' *.fastq > count/B5573_Fm0.txt
agrep -0 -c 'AGGATAGCAGCGTGACT' *.fastq > count/B5574_Fm0.txt
agrep -0 -c 'TATCGTTAGGTAACTGC' *.fastq > count/B5576_Fm0.txt
agrep -0 -c 'CCGAATTCTACATGCAC' *.fastq > count/B5577_Fm0.txt
agrep -0 -c 'TCAGTGTGGTGTCCGAA' *.fastq > count/B5578_Fm0.txt
agrep -0 -c 'ATTTATGCGGCCTCTCC' *.fastq > count/B5579_Fm0.txt
agrep -0 -c 'TGTTCCCGCGACTTAAT' *.fastq > count/B5580_Fm0.txt
agrep -0 -c 'AGCGATGCTATGGACAA' *.fastq > count/B5581_Fm0.txt
agrep -0 -c 'CTTCTCGTGCATGAGTA' *.fastq > count/B5582_Fm0.txt
agrep -0 -c 'TTGCGGTCCTCCCACAT' *.fastq > count/B5585_Fm0.txt
agrep -0 -c 'GGTCGTACGCTGACGTA' *.fastq > count/B5586_Fm0.txt
agrep -0 -c 'CTTCGCAACCATTAGGC' *.fastq > count/B5587_Fm0.txt
agrep -0 -c 'CACCGTTAATTTCGCAC' *.fastq > count/B5588_Fm0.txt
agrep -0 -c 'TTAATTCCGCCTACCAG' *.fastq > count/B5591_Fm0.txt
agrep -0 -c 'CTGCCGCGATCGTTATA' *.fastq > count/B5592_Fm0.txt
agrep -0 -c 'ATCTAGATGCGGGATGG' *.fastq > count/B5593_Fm0.txt
agrep -0 -c 'TACCTAGTATACAGAGG' *.fastq > count/B5595_Fm0.txt
agrep -0 -c 'GCTGAAGGGGGTTTCTC' *.fastq > count/B5596_Fm0.txt
agrep -0 -c 'CCACCTCCCGTCATTAA' *.fastq > count/B5597_Fm0.txt
agrep -0 -c 'AGGTGCTGTACAACTCC' *.fastq > count/B5599_Fm0.txt
agrep -0 -c 'TACCCCAGAAATCCGAG' *.fastq > count/B5600_Fm0.txt
agrep -0 -c 'TAGTTGGAAATTCCCCT' *.fastq > count/B5601_Fm0.txt
agrep -0 -c 'CTCCACTGTACTAGCTA' *.fastq > count/B5602_Fm0.txt
agrep -0 -c 'AGTGTAACTCGCGAGAA' *.fastq > count/B5603_Fm0.txt
agrep -0 -c 'AGGTGGTTACATACACG' *.fastq > count/B5604_Fm0.txt
agrep -0 -c 'TATGGTCTCCGTGTCCG' *.fastq > count/B5606_Fm0.txt
agrep -0 -c 'AAGCCATGGTGTCTACG' *.fastq > count/B5607_Fm0.txt
agrep -0 -c 'TTCAGTCCACAGGATTA' *.fastq > count/B5608_Fm0.txt
agrep -0 -c 'GCACGAACGTGAATTCT' *.fastq > count/B5609_Fm0.txt
agrep -0 -c 'AGCATGGAAGACTGGTC' *.fastq > count/B5610_Fm0.txt
agrep -0 -c 'GGAGGTTCTTAAGGGCA' *.fastq > count/B5611_Fm0.txt
agrep -0 -c 'CCGGTCATCTATCGTCA' *.fastq > count/B5612_Fm0.txt
agrep -0 -c 'ATCAATTGTGGCCCGTG' *.fastq > count/B5614_Fm0.txt
agrep -0 -c 'GTGAAATTCATCGGAAC' *.fastq > count/B5615_Fm0.txt
agrep -0 -c 'TTGTGCGTAGTCGGAGA' *.fastq > count/B5616_Fm0.txt
agrep -0 -c 'ACCCGAATAGAGCATCT' *.fastq > count/B5617_Fm0.txt
agrep -0 -c 'ATGGAAGTGGGACGTTA' *.fastq > count/B5618_Fm0.txt
agrep -0 -c 'CCATGTCTAACCCGTAC' *.fastq > count/B5621_Fm0.txt
agrep -0 -c 'GTGTCCGTTTGCGACAG' *.fastq > count/B5622_Fm0.txt
agrep -0 -c 'TGTAGGGGGAGTATTTG' *.fastq > count/B5623_Fm0.txt
agrep -0 -c 'GTGGACTCGTTCGGTTT' *.fastq > count/B5624_Fm0.txt
agrep -0 -c 'ACAACGCGGGTATTTTA' *.fastq > count/B5625_Fm0.txt
agrep -0 -c 'GGCCACCAGCACAATAA' *.fastq > count/B5626_Fm0.txt
agrep -0 -c 'TAAAGTGAAGTAGTGCG' *.fastq > count/B5627_Fm0.txt
agrep -0 -c 'ACCCGATCCATTCATTA' *.fastq > count/B5629_Fm0.txt
agrep -0 -c 'CAGTGTTTCGATCATCA' *.fastq > count/B5630_Fm0.txt
agrep -0 -c 'AGCACGGGCTTATCGTG' *.fastq > count/B5631_Fm0.txt
agrep -0 -c 'TTCGCGGCCCATTCCAT' *.fastq > count/B5632_Fm0.txt
agrep -0 -c 'TGTGCGCGAATTCCTGC' *.fastq > count/B5634_Fm0.txt
agrep -0 -c 'CACCAGTTTCTTGTACG' *.fastq > count/B5635_Fm0.txt
agrep -0 -c 'CGCATCCATTAGTGATT' *.fastq > count/B5636_Fm0.txt
agrep -0 -c 'CAGCTGGTTGTATGCGA' *.fastq > count/B5637_Fm0.txt
agrep -0 -c 'TCAGACCACCTTATGTC' *.fastq > count/B5638_Fm0.txt
agrep -0 -c 'GGTGTGAAGGAATAGGT' *.fastq > count/B5639_Fm0.txt
agrep -0 -c 'TAGAGGTCATGCACTGA' *.fastq > count/B5641_Fm0.txt
agrep -0 -c 'AACCGTAACCTGGGTCG' *.fastq > count/B5642_Fm0.txt
agrep -0 -c 'CTACTTTCCTAGTGTGC' *.fastq > count/B5643_Fm0.txt
agrep -0 -c 'CGTAACTCATTTACCTG' *.fastq > count/B5645_Fm0.txt
agrep -0 -c 'TAGTGAGCTCGGGTAAC' *.fastq > count/B5646_Fm0.txt
agrep -0 -c 'GCATGGTCGCTTGTGAA' *.fastq > count/B5647_Fm0.txt
agrep -0 -c 'CAGCCGGCAAATAAGGG' *.fastq > count/B5648_Fm0.txt
agrep -0 -c 'CAGGCATGAGCCGTTAT' *.fastq > count/B5649_Fm0.txt
agrep -0 -c 'TCGGATAGCGTCAAGGT' *.fastq > count/B5650_Fm0.txt
agrep -0 -c 'CGACGCTCTAGTCTCCT' *.fastq > count/B5651_Fm0.txt
agrep -0 -c 'GATTGCACCATCATTCC' *.fastq > count/B5652_Fm0.txt
agrep -0 -c 'GGGGCCGACACTATGAA' *.fastq > count/B5653_Fm0.txt
agrep -0 -c 'CATGACTCGTATCGATG' *.fastq > count/B5655_Fm0.txt
agrep -0 -c 'GCCCTAACAGTCGATGC' *.fastq > count/B5658_Fm0.txt
agrep -0 -c 'CCATTCAAGCTCGAATT' *.fastq > count/B5659_Fm0.txt
agrep -0 -c 'GTAGAAAGTTACAGCGC' *.fastq > count/B5660_Fm0.txt
agrep -0 -c 'CCGTACGCCTTCGTCTA' *.fastq > count/B5661_Fm0.txt
agrep -0 -c 'CTCTACAAACTAGTTGC' *.fastq > count/B5662_Fm0.txt
agrep -0 -c 'CAGTTTTTAGGAGGTGT' *.fastq > count/B5663_Fm0.txt
agrep -0 -c 'TTATGGAGTTGATTGCC' *.fastq > count/B5664_Fm0.txt
agrep -0 -c 'TAGTGAAAGAGTGGCAA' *.fastq > count/B5665_Fm0.txt
agrep -0 -c 'AGAACGGTGACTAGAGT' *.fastq > count/B5666_Fm0.txt
agrep -0 -c 'ACTACGCTAGGTTGGTA' *.fastq > count/B5668_Fm0.txt
agrep -0 -c 'TGCACACGACCTCATAC' *.fastq > count/B5669_Fm0.txt
agrep -0 -c 'CGAGCTCACTCAAACAG' *.fastq > count/B5670_Fm0.txt
agrep -0 -c 'CTGTTACCCGATTCGTG' *.fastq > count/B5671_Fm0.txt
agrep -0 -c 'AGACCACTACCTGTGTA' *.fastq > count/B5672_Fm0.txt
agrep -0 -c 'ACAGCGACGCTTCAAAG' *.fastq > count/B5673_Fm0.txt
agrep -0 -c 'TGGCTCAGGTCGGGTTT' *.fastq > count/B5674_Fm0.txt
agrep -0 -c 'CCACCGATTCCTACTTG' *.fastq > count/B5675_Fm0.txt
agrep -0 -c 'CGTATTGCAACGACCCT' *.fastq > count/B5676_Fm0.txt
agrep -0 -c 'TACACAGATAGCGGGCT' *.fastq > count/B5677_Fm0.txt
agrep -0 -c 'AACCCTCTGACGCCACT' *.fastq > count/B5678_Fm0.txt
agrep -0 -c 'AGCCCACTATCTTGCTC' *.fastq > count/B5679_Fm0.txt
agrep -0 -c 'TTCCGCTTCTGGCCATC' *.fastq > count/B5680_Fm0.txt
agrep -0 -c 'CCATGGCCGAGCAAAAC' *.fastq > count/B5681_Fm0.txt
agrep -0 -c 'GCAGGAGGAAAGCGATA' *.fastq > count/B5683_Fm0.txt
agrep -0 -c 'CCGGATACATCGTTGTA' *.fastq > count/B5684_Fm0.txt
agrep -0 -c 'CGAGTGTAGAATCTCCA' *.fastq > count/B5687_Fm0.txt
agrep -0 -c 'TCGGAGAGATCCTTCCA' *.fastq > count/B5689_Fm0.txt
agrep -0 -c 'GCCCCTGTGCAATTTTG' *.fastq > count/B5690_Fm0.txt
agrep -0 -c 'CGACTATTGATAATGCC' *.fastq > count/B5691_Fm0.txt
agrep -0 -c 'TAAGTCTGTGCCCAGCA' *.fastq > count/B5692_Fm0.txt
agrep -0 -c 'CCGGTGTCACATCAACA' *.fastq > count/B5693_Fm0.txt
agrep -0 -c 'GGCTGTCCCATTAGTCG' *.fastq > count/B5694_Fm0.txt
agrep -0 -c 'CCTACTATGCGAGGAGG' *.fastq > count/B5697_Fm0.txt
agrep -0 -c 'ACAGGTATGAAACCCGG' *.fastq > count/B5698_Fm0.txt
agrep -0 -c 'GGACCCTTGAGAATTGG' *.fastq > count/B5699_Fm0.txt
agrep -0 -c 'GGATTATAGTCACGATC' *.fastq > count/B5700_Fm0.txt
agrep -0 -c 'GGGACACAAGAACTCCA' *.fastq > count/B5701_Fm0.txt
agrep -0 -c 'GGCATGCCTTGGGGATA' *.fastq > count/B5702_Fm0.txt
agrep -0 -c 'AACTAGATTGGTCCGGG' *.fastq > count/B5703_Fm0.txt
agrep -0 -c 'TGGCCTCAGTGTCGCTA' *.fastq > count/B5704_Fm0.txt
agrep -0 -c 'CGTGTACTCCCATGGCT' *.fastq > count/B5705_Fm0.txt
agrep -0 -c 'TATACGGGGAAGACTCC' *.fastq > count/B5706_Fm0.txt
agrep -0 -c 'GCGGAAAACTGCAGGCA' *.fastq > count/B5707_Fm0.txt
agrep -0 -c 'TCGACCTCAATCAGTTA' *.fastq > count/B5709_Fm0.txt
agrep -0 -c 'AGTTGCGAGTTAGGCGT' *.fastq > count/B5710_Fm0.txt
agrep -0 -c 'GTCAGGTATTCCATGGC' *.fastq > count/B5712_Fm0.txt
agrep -0 -c 'GCACCGTGTGATGGTTA' *.fastq > count/B5714_Fm0.txt
agrep -0 -c 'GGCACACTGTCGAAACC' *.fastq > count/B5715_Fm0.txt
agrep -0 -c 'TAACCGCTCTATACTCC' *.fastq > count/B5716_Fm0.txt
agrep -0 -c 'ACGCATTCTCTCAAGGT' *.fastq > count/B5717_Fm0.txt
agrep -0 -c 'AACGAGCGCAGGGATTT' *.fastq > count/B5718_Fm0.txt
agrep -0 -c 'ACCCCGCTTATTCCGTT' *.fastq > count/B5719_Fm0.txt
agrep -0 -c 'TCATCGCGGCGTTTTAA' *.fastq > count/B5720_Fm0.txt
agrep -0 -c 'TAGTTGTGATCATCTCG' *.fastq > count/B5721_Fm0.txt
agrep -0 -c 'CCAGTCCGCACGTAATC' *.fastq > count/B5722_Fm0.txt
agrep -0 -c 'GGTATAATACCTGCTCT' *.fastq > count/B5723_Fm0.txt
agrep -0 -c 'CCGTAACTCAGCCTCGA' *.fastq > count/B5725_Fm0.txt
agrep -0 -c 'TGCGTCATCAAAAGCTT' *.fastq > count/B5726_Fm0.txt
agrep -0 -c 'CCTCATTACCAGGCGCT' *.fastq > count/B5727_Fm0.txt
agrep -0 -c 'CGATCAGCTTGACATCG' *.fastq > count/B5728_Fm0.txt
agrep -0 -c 'ACCTAATTGAACTGCCG' *.fastq > count/B5729_Fm0.txt
agrep -0 -c 'GGTTTATCTTCTGCATG' *.fastq > count/B5730_Fm0.txt
agrep -0 -c 'CGAAACCAGCGGCAATC' *.fastq > count/B5731_Fm0.txt
agrep -0 -c 'TTAGCGCTCTATCGCTT' *.fastq > count/B5732_Fm0.txt
agrep -0 -c 'TCTTACTACTGGTGTCC' *.fastq > count/B5734_Fm0.txt
agrep -0 -c 'CGAAAAGGTCGTTGCTC' *.fastq > count/B5735_Fm0.txt
agrep -0 -c 'TTCACCCCGGTAACTGC' *.fastq > count/B5736_Fm0.txt
agrep -0 -c 'CACCCTAGCACGTCGAT' *.fastq > count/B5737_Fm0.txt
agrep -0 -c 'AGAAGAAACGCTCCCAC' *.fastq > count/B5738_Fm0.txt
agrep -0 -c 'AGCATTCGTCTTAACAC' *.fastq > count/B5739_Fm0.txt
agrep -0 -c 'GGCTTACAACGTATTCA' *.fastq > count/B5740_Fm0.txt
agrep -0 -c 'GTCCAGCGGAGTGTTAC' *.fastq > count/B5741_Fm0.txt
agrep -0 -c 'TAATCGCCCGAGATTGA' *.fastq > count/B5742_Fm0.txt
agrep -0 -c 'GGATAGGAACCAATCCA' *.fastq > count/B5743_Fm0.txt
agrep -0 -c 'GCGTTGGATCGAGTTTG' *.fastq > count/B5746_Fm0.txt
agrep -0 -c 'TGATATCCGTGATCAGG' *.fastq > count/B5747_Fm0.txt
agrep -0 -c 'GGATTTCGACTATAGGG' *.fastq > count/B5749_Fm0.txt
agrep -0 -c 'AATGCCAGAGAGGAGTG' *.fastq > count/B5750_Fm0.txt
agrep -0 -c 'GTCCGCTCTTGGGGTTA' *.fastq > count/B5751_Fm0.txt
agrep -0 -c 'TTCCGGCGATCGACGTT' *.fastq > count/B5752_Fm0.txt
agrep -0 -c 'TAGTTGAGTCGGCCTAT' *.fastq > count/B5753_Fm0.txt
agrep -0 -c 'ATTCGCCAATAAGTCGG' *.fastq > count/B5754_Fm0.txt
agrep -0 -c 'AATCGCTTGTGGCCAAG' *.fastq > count/B5755_Fm0.txt
agrep -0 -c 'ATAGCTAGCCTGGTGAT' *.fastq > count/B5756_Fm0.txt
agrep -0 -c 'ATGAGCGTACCTTCTCC' *.fastq > count/B5757_Fm0.txt
agrep -0 -c 'AAGGCGCTTTCGTTGAT' *.fastq > count/B5758_Fm0.txt
agrep -0 -c 'TGAACCTGCCGCTATTC' *.fastq > count/B5759_Fm0.txt
agrep -0 -c 'ACCAGGATAGACGCCTT' *.fastq > count/B5760_Fm0.txt
agrep -0 -c 'GAGACAAGGACGGTATC' *.fastq > count/B5762_Fm0.txt
agrep -0 -c 'GTGTCTTGCCGTAAGTC' *.fastq > count/B5763_Fm0.txt
agrep -0 -c 'ACTAGTAGACCAGGAGG' *.fastq > count/B5764_Fm0.txt
agrep -0 -c 'AGCCATAACCCAAGAAA' *.fastq > count/B5765_Fm0.txt
agrep -0 -c 'TCTGTTCGTCTTTAGCC' *.fastq > count/B5767_Fm0.txt
agrep -0 -c 'AATGAGTCACCACAGGA' *.fastq > count/B5768_Fm0.txt
agrep -0 -c 'TCGGTCACATACAAGAC' *.fastq > count/B5769_Fm0.txt
agrep -0 -c 'TAACCGAATGTCTAGCT' *.fastq > count/B5770_Fm0.txt
agrep -0 -c 'AGTATGGGTACCTCCTA' *.fastq > count/B5771_Fm0.txt
agrep -0 -c 'CCATATCTCACGCGGCA' *.fastq > count/B5772_Fm0.txt
agrep -0 -c 'TCACATCCAGCTCGGAA' *.fastq > count/B5773_Fm0.txt
agrep -0 -c 'TAATCTAAGGCCGTGGG' *.fastq > count/B5774_Fm0.txt
agrep -0 -c 'TTGTAGGGGATCAAGAA' *.fastq > count/B5775_Fm0.txt
agrep -0 -c 'ACTGTTGCTCAGAGGTA' *.fastq > count/B5776_Fm0.txt
agrep -0 -c 'CCGTTGCTTGCGCCTTT' *.fastq > count/B5777_Fm0.txt
agrep -0 -c 'CCTCGCCCTTTGATGGA' *.fastq > count/B5778_Fm0.txt
agrep -0 -c 'TCACTACCACGGCGCAT' *.fastq > count/B5779_Fm0.txt
agrep -0 -c 'ACCTCTCCCTATCGGTT' *.fastq > count/B5780_Fm0.txt
agrep -0 -c 'GGAACTGGTTCGCTTAC' *.fastq > count/B5781_Fm0.txt
agrep -0 -c 'TTATCAGGATCCGACTG' *.fastq > count/B5782_Fm0.txt
agrep -0 -c 'CTTACAAAAGCACTACG' *.fastq > count/B5783_Fm0.txt
agrep -0 -c 'ATAAAGCGAATGGGCAT' *.fastq > count/B5785_Fm0.txt
agrep -0 -c 'GAGGAACCCGTATCTCC' *.fastq > count/B5786_Fm0.txt
agrep -0 -c 'TTCTAACCTTCGCGACG' *.fastq > count/B5787_Fm0.txt
agrep -0 -c 'ACAAAACGATGGTTAGG' *.fastq > count/B5789_Fm0.txt
agrep -0 -c 'CGTGGACGTTTGAAATC' *.fastq > count/B5790_Fm0.txt
agrep -0 -c 'GGGCAGACACTAGTGTT' *.fastq > count/B5792_Fm0.txt
agrep -0 -c 'GGGCCTCTGGAAAACGT' *.fastq > count/B5793_Fm0.txt
agrep -0 -c 'CGAGACTTTGTCTGCCT' *.fastq > count/B5795_Fm0.txt
agrep -0 -c 'TATCAAAGCCTTAGCGA' *.fastq > count/B5797_Fm0.txt
agrep -0 -c 'TACCGCTATTCCCTAGC' *.fastq > count/B5798_Fm0.txt
agrep -0 -c 'TCGTAGAGATTTCAGGG' *.fastq > count/B5799_Fm0.txt
agrep -0 -c 'CAAGTGCGATCAAGCGA' *.fastq > count/B5800_Fm0.txt
agrep -0 -c 'CTTGAATACCCGGCAAC' *.fastq > count/B5801_Fm0.txt
agrep -0 -c 'GCGATCATTGCTCTTGG' *.fastq > count/B5802_Fm0.txt
agrep -0 -c 'AACCTAGCAGTCAACGT' *.fastq > count/B5803_Fm0.txt
agrep -0 -c 'ACCACATAATACTACCG' *.fastq > count/B5804_Fm0.txt
agrep -0 -c 'TCTCCGCGCAGTAGATA' *.fastq > count/B5805_Fm0.txt
agrep -0 -c 'CTAGACCCTAGCTGCAA' *.fastq > count/B5806_Fm0.txt
agrep -0 -c 'TAGGCTAAGCAACACGG' *.fastq > count/B5807_Fm0.txt
agrep -0 -c 'AGGACCTCTCTGACCTC' *.fastq > count/B5808_Fm0.txt
agrep -0 -c 'AATGCCACTCATTATCG' *.fastq > count/B5809_Fm0.txt
agrep -0 -c 'CATTGGGTGCCCTGTTC' *.fastq > count/B5810_Fm0.txt
agrep -0 -c 'TCTGGGACTGAATCGTT' *.fastq > count/B5811_Fm0.txt
agrep -0 -c 'CCGTTGTATACGGATCA' *.fastq > count/B5812_Fm0.txt
agrep -0 -c 'TTGATTCCAACGTCAGG' *.fastq > count/B5813_Fm0.txt
agrep -0 -c 'AGTCCTTATGCAATGAG' *.fastq > count/B5814_Fm0.txt
agrep -0 -c 'CGGTCTAAACCGTGTCG' *.fastq > count/B5815_Fm0.txt
agrep -0 -c 'CCGGCTCGTTTCTGAGT' *.fastq > count/B5816_Fm0.txt
agrep -0 -c 'ATGCGGCGACTTTATGC' *.fastq > count/B5817_Fm0.txt
agrep -0 -c 'TAACATGTCTCGTTCAG' *.fastq > count/B5818_Fm0.txt
agrep -0 -c 'GATTGTTAGGTATGCGC' *.fastq > count/B5821_Fm0.txt
agrep -0 -c 'GGCCTAAATTGTTGGAA' *.fastq > count/B5822_Fm0.txt
agrep -0 -c 'CCAGATATCGATGTGGG' *.fastq > count/B5823_Fm0.txt
agrep -0 -c 'TAAGTTCTGCGTCGAGA' *.fastq > count/B5824_Fm0.txt
agrep -0 -c 'TCGCACTCGAAACGAGT' *.fastq > count/B5825_Fm0.txt
agrep -0 -c 'TCGTCCAGCCAACTGAA' *.fastq > count/B5826_Fm0.txt
agrep -0 -c 'GTCAGTGCCGCTTGATG' *.fastq > count/B5827_Fm0.txt
agrep -0 -c 'CTCTAAGGTATCGGCTC' *.fastq > count/B5828_Fm0.txt
agrep -0 -c 'TATTGGCAACCAGTGTA' *.fastq > count/B5829_Fm0.txt
agrep -0 -c 'TAGCGTTGGAACCACGG' *.fastq > count/B5830_Fm0.txt
agrep -0 -c 'ATTGTGACGAACTCTGC' *.fastq > count/B5831_Fm0.txt
agrep -0 -c 'GTCACCGTAACAGACCT' *.fastq > count/B5833_Fm0.txt
agrep -0 -c 'CCGAGTAATTTCACCAC' *.fastq > count/B5834_Fm0.txt
agrep -0 -c 'GCGGCGATGGATCCTAA' *.fastq > count/B5835_Fm0.txt
agrep -0 -c 'CCGTTCGTTCGAAATGA' *.fastq > count/B5836_Fm0.txt
agrep -0 -c 'TCTCAACAGGGACATAT' *.fastq > count/B5837_Fm0.txt
agrep -0 -c 'CGAAGATGGTTACCGAC' *.fastq > count/B5838_Fm0.txt
agrep -0 -c 'CTGAATCCCCATATCCG' *.fastq > count/B5839_Fm0.txt
agrep -0 -c 'CCGCCTGATATCGACTT' *.fastq > count/B5840_Fm0.txt
agrep -0 -c 'CGACTTCAACACGTATT' *.fastq > count/B5841_Fm0.txt
agrep -0 -c 'TGCCAAATGTACCCTAA' *.fastq > count/B5842_Fm0.txt
agrep -0 -c 'TATCTCACGGTCAGGTG' *.fastq > count/B5843_Fm0.txt
agrep -0 -c 'CGATCCCAATATGATGT' *.fastq > count/B5844_Fm0.txt
agrep -0 -c 'GCATGTGAGACAATCAG' *.fastq > count/B5847_Fm0.txt
agrep -0 -c 'TCGACGTAATCCTACTC' *.fastq > count/B5848_Fm0.txt
agrep -0 -c 'GTTGGAGTCCAGCCTAC' *.fastq > count/B5849_Fm0.txt
agrep -0 -c 'GGATGCGAACTAGCTCA' *.fastq > count/B5850_Fm0.txt
agrep -0 -c 'ATGCGATAGATTCCGGG' *.fastq > count/B5851_Fm0.txt
agrep -0 -c 'GCAAACAGCCCGGTATA' *.fastq > count/B5852_Fm0.txt
agrep -0 -c 'TGACCGCCCATTTGATG' *.fastq > count/B5853_Fm0.txt
agrep -0 -c 'TACTATACCCGCCTAAG' *.fastq > count/B5854_Fm0.txt
agrep -0 -c 'ATCAGCCTTGCCAACAT' *.fastq > count/B5855_Fm0.txt
agrep -0 -c 'CTAACACAGACAGAGAT' *.fastq > count/B5856_Fm0.txt
agrep -0 -c 'GCTCCGTCCTCTAATTA' *.fastq > count/B5858_Fm0.txt
agrep -0 -c 'CTTCTCGTGGGTTCCTG' *.fastq > count/B5859_Fm0.txt
agrep -0 -c 'GTATTTAGGGTGCCCCC' *.fastq > count/B5860_Fm0.txt
agrep -0 -c 'AGTGAGTCAACAAAGTG' *.fastq > count/B5861_Fm0.txt
agrep -0 -c 'CCAGGTCCATGAATCAG' *.fastq > count/B5862_Fm0.txt
agrep -0 -c 'GAGCCGGGTTTATGTCA' *.fastq > count/B5863_Fm0.txt
agrep -0 -c 'TCAACTCATTGACCCGG' *.fastq > count/B5864_Fm0.txt
agrep -0 -c 'CGCGCATTCATGATTGC' *.fastq > count/B5865_Fm0.txt
agrep -0 -c 'ACTGCACCATGAATAGT' *.fastq > count/B5866_Fm0.txt
agrep -0 -c 'AATGCCTAGCCATGACA' *.fastq > count/B5867_Fm0.txt
agrep -0 -c 'GTTTCTAAGACTGGCTC' *.fastq > count/B5868_Fm0.txt
agrep -0 -c 'AGTCGTTAAACGCGCGC' *.fastq > count/B5869_Fm0.txt
agrep -0 -c 'CGAATACAGAACGGGAT' *.fastq > count/B5870_Fm0.txt
agrep -0 -c 'TCAAATCCAACACAGCG' *.fastq > count/B5871_Fm0.txt
agrep -0 -c 'CGCGCTCTACTCTTATA' *.fastq > count/B5873_Fm0.txt
agrep -0 -c 'AAGTAAGGGATTCGGGA' *.fastq > count/B5874_Fm0.txt
agrep -0 -c 'CGGTTTTGACTGAGACT' *.fastq > count/B5876_Fm0.txt
agrep -0 -c 'GAGTTGACTAGCTGATT' *.fastq > count/B5877_Fm0.txt
agrep -0 -c 'GTTCTAGCGATCACTGT' *.fastq > count/B5878_Fm0.txt
agrep -0 -c 'CATCCGTAGCATCTAGA' *.fastq > count/B5879_Fm0.txt
agrep -0 -c 'ATCCGTAGCAAGTCCGT' *.fastq > count/B5880_Fm0.txt
agrep -0 -c 'CGTGGCTATTTTCAGTG' *.fastq > count/B5882_Fm0.txt
agrep -0 -c 'AGTTATATCGCCCGCTT' *.fastq > count/B5884_Fm0.txt
agrep -0 -c 'CAGCAGGTAGTATCGTC' *.fastq > count/B5885_Fm0.txt
agrep -0 -c 'CTCGCAGCAAGCATACC' *.fastq > count/B5886_Fm0.txt
agrep -0 -c 'ATGCGTGGCTCCATTGT' *.fastq > count/B5887_Fm0.txt
agrep -0 -c 'AAGTCTGTAGCAGGCAT' *.fastq > count/B5888_Fm0.txt
agrep -0 -c 'CACCTGAACGCAAAAGG' *.fastq > count/B5889_Fm0.txt
agrep -0 -c 'TCTCTAACCATGGCCGT' *.fastq > count/B5890_Fm0.txt
agrep -0 -c 'GTCTAGTCTGCGTCCAG' *.fastq > count/B5891_Fm0.txt
agrep -0 -c 'GCCTGACTATGGTGTGC' *.fastq > count/B5893_Fm0.txt
agrep -0 -c 'TCGACAAGTCAATGCTT' *.fastq > count/B5894_Fm0.txt
agrep -0 -c 'CAGCGAAATATATGGCG' *.fastq > count/B5895_Fm0.txt
agrep -0 -c 'AGGCGTGGTTGACAGTA' *.fastq > count/B5897_Fm0.txt
agrep -0 -c 'GTAAGTTACGTCAGTCG' *.fastq > count/B5898_Fm0.txt
agrep -0 -c 'TCGGAGCGTATTAATTG' *.fastq > count/B5899_Fm0.txt
agrep -0 -c 'CGCTGTCATTTCAAAAC' *.fastq > count/B5900_Fm0.txt
agrep -0 -c 'TCGTGCTAAGTAGTCAA' *.fastq > count/B5901_Fm0.txt
agrep -0 -c 'ATTCAGTTGTAGCGTTC' *.fastq > count/B5902_Fm0.txt
agrep -0 -c 'GGCATTCCGTTACTCAG' *.fastq > count/B5903_Fm0.txt
agrep -0 -c 'CTACCGTACCCTAAGCG' *.fastq > count/B5904_Fm0.txt
agrep -0 -c 'TAGTAATGACTTGTGCG' *.fastq > count/B5905_Fm0.txt
agrep -0 -c 'ACTCCGCGGCTCCTTAA' *.fastq > count/B5906_Fm0.txt
agrep -0 -c 'ACATGTTGAAGCCACTT' *.fastq > count/B5907_Fm0.txt
agrep -0 -c 'CGTTGAGTGTCTGTGTG' *.fastq > count/B5908_Fm0.txt
agrep -0 -c 'ACAGCCCCTTACCATAA' *.fastq > count/B5909_Fm0.txt
agrep -0 -c 'AGCCATGAAAGCTTCCC' *.fastq > count/B5910_Fm0.txt
agrep -0 -c 'GTTCTCTAGTACCAGGA' *.fastq > count/B5911_Fm0.txt
agrep -0 -c 'ATACCACCCAGGGGGAA' *.fastq > count/B5912_Fm0.txt
agrep -0 -c 'GCGGAGTACTAATGCCC' *.fastq > count/B5913_Fm0.txt
agrep -0 -c 'GCTAGGGTGGGATCGTT' *.fastq > count/B5915_Fm0.txt
agrep -0 -c 'CGCGCTGAGTGTCAACT' *.fastq > count/B5917_Fm0.txt
agrep -0 -c 'GACTTCAGAACCAAGAA' *.fastq > count/B5918_Fm0.txt
agrep -0 -c 'CGTTACTGGCTACAGGT' *.fastq > count/B5919_Fm0.txt
agrep -0 -c 'AACTGCTCTGACGACTT' *.fastq > count/B5920_Fm0.txt
agrep -0 -c 'CTGCCGTTCCATCCCAT' *.fastq > count/B5921_Fm0.txt
agrep -0 -c 'AGCTGGCCTACGAACAA' *.fastq > count/B5922_Fm0.txt
agrep -0 -c 'CGCTACCGGGATACACT' *.fastq > count/B5924_Fm0.txt
agrep -0 -c 'CTATCTTGCATTGAGCC' *.fastq > count/B5926_Fm0.txt
agrep -0 -c 'AATAGTTCGCGGGCTAA' *.fastq > count/B5927_Fm0.txt
agrep -0 -c 'AAGGGGCAATATCGTTT' *.fastq > count/B5928_Fm0.txt
agrep -0 -c 'CGACTCTAACTGTGAAC' *.fastq > count/B5929_Fm0.txt
agrep -0 -c 'TCTGGTCTACCTATACG' *.fastq > count/B5930_Fm0.txt
agrep -0 -c 'TAACCCACTCTGTCGGA' *.fastq > count/B5931_Fm0.txt
agrep -0 -c 'GCTCGGCTTTATGCCGT' *.fastq > count/B5932_Fm0.txt
agrep -0 -c 'ACGGGTCCTAACGCAAT' *.fastq > count/B5933_Fm0.txt
agrep -0 -c 'TTCCGGAGTTTAGGGAC' *.fastq > count/B5934_Fm0.txt
agrep -0 -c 'AGCATAGTTGCTCCCAA' *.fastq > count/B5935_Fm0.txt
agrep -0 -c 'GATATCACATCCCGACC' *.fastq > count/B5936_Fm0.txt
agrep -0 -c 'ATTGGCTTTAGCTCCAT' *.fastq > count/B5937_Fm0.txt
agrep -0 -c 'GGCATCAGAGTTACTCG' *.fastq > count/B5939_Fm0.txt
agrep -0 -c 'AATCACCGAGTAGCCGC' *.fastq > count/B5940_Fm0.txt
agrep -0 -c 'TTCGACGGCTAATTCGT' *.fastq > count/B5941_Fm0.txt
agrep -0 -c 'CTAGCTGTTGCTTTTCG' *.fastq > count/B5943_Fm0.txt
agrep -0 -c 'AGCTTCACTCGCAAATG' *.fastq > count/B5944_Fm0.txt
agrep -0 -c 'CTATATCACCCCAAACG' *.fastq > count/B5945_Fm0.txt
agrep -0 -c 'CAGCGCACACTCCTTAT' *.fastq > count/B5946_Fm0.txt
agrep -0 -c 'GTAGTCGACACGGAATT' *.fastq > count/B5949_Fm0.txt
agrep -0 -c 'TATGTGATGCGTGTCGT' *.fastq > count/B5950_Fm0.txt
agrep -0 -c 'TGCGCACTCGATCGCAA' *.fastq > count/B5952_Fm0.txt
agrep -0 -c 'TGCTAATGCTGTCTTTG' *.fastq > count/B5953_Fm0.txt
agrep -0 -c 'ACCGCGAATGTGGGCTT' *.fastq > count/B5954_Fm0.txt
agrep -0 -c 'CTCAGATAACGGAAGCT' *.fastq > count/B5955_Fm0.txt
agrep -0 -c 'GCCATCATCACGCGCAA' *.fastq > count/B5957_Fm0.txt
agrep -0 -c 'ATACACATGGGGGGTGC' *.fastq > count/B5958_Fm0.txt
agrep -0 -c 'CGAAAACGACCCCCTTC' *.fastq > count/B5959_Fm0.txt
agrep -0 -c 'TTGGTGTGGTACAGACC' *.fastq > count/B5960_Fm0.txt
agrep -0 -c 'TCGCCCGTGCATAAGAG' *.fastq > count/B5961_Fm0.txt
agrep -0 -c 'AGTTTGGCTCCTCTACC' *.fastq > count/B5962_Fm0.txt
agrep -0 -c 'CACCTCCAACCTACGTT' *.fastq > count/B5963_Fm0.txt
agrep -0 -c 'GTTAAATTCCCCCGCCT' *.fastq > count/B5964_Fm0.txt
agrep -0 -c 'TCAAACCGGAGTACTGA' *.fastq > count/B5968_Fm0.txt
agrep -0 -c 'TGAGCGTGGACTGAATC' *.fastq > count/B5970_Fm0.txt
agrep -0 -c 'CTGCCATGAGGAGCACT' *.fastq > count/B5971_Fm0.txt
agrep -0 -c 'CAACAATTTGATCGGTC' *.fastq > count/B5972_Fm0.txt
agrep -0 -c 'AGACACAGTCCTGTTTT' *.fastq > count/B5973_Fm0.txt
agrep -0 -c 'AGCCGTGTACTAATTCA' *.fastq > count/B5974_Fm0.txt
agrep -0 -c 'TAGTCCGTCTTAGCTAA' *.fastq > count/B5975_Fm0.txt
agrep -0 -c 'TAAGGTCACGCTGAACA' *.fastq > count/B5976_Fm0.txt
agrep -0 -c 'GAGTGAGTTCTACGTAT' *.fastq > count/B5977_Fm0.txt
agrep -0 -c 'ATTCAGAGATGCTATCG' *.fastq > count/B5978_Fm0.txt
agrep -0 -c 'TCAGCATAGGGGCACTA' *.fastq > count/B5979_Fm0.txt
agrep -0 -c 'GCATATATGTATCCGCA' *.fastq > count/B5980_Fm0.txt
agrep -0 -c 'TCACTCTTTGCGACCAG' *.fastq > count/B5981_Fm0.txt
agrep -0 -c 'TGTCCTTTACTCTTGTG' *.fastq > count/B5982_Fm0.txt
agrep -0 -c 'TCGTGTTCTCAGCGGTT' *.fastq > count/B5983_Fm0.txt
agrep -0 -c 'GACGCTCGAGGTTATGC' *.fastq > count/B5984_Fm0.txt
agrep -0 -c 'GAGTGTCCAATGATCGA' *.fastq > count/B5985_Fm0.txt
agrep -0 -c 'ACTGCCGGAGGTACACT' *.fastq > count/B5986_Fm0.txt
agrep -0 -c 'TGAGACAAGCCTCCTGA' *.fastq > count/B5987_Fm0.txt
agrep -0 -c 'CATGCCCTACAGTTTAA' *.fastq > count/B5988_Fm0.txt
agrep -0 -c 'TCGCGACAGGAAGATAC' *.fastq > count/B5989_Fm0.txt
agrep -0 -c 'CAAGCTACATAACCTGA' *.fastq > count/B5990_Fm0.txt
agrep -0 -c 'TAAGACAAGCCTGAGTC' *.fastq > count/B5991_Fm0.txt
agrep -0 -c 'GTTAGTAGCAACGTGCA' *.fastq > count/B5993_Fm0.txt
agrep -0 -c 'TCTGGACAATGGGTCGT' *.fastq > count/B5994_Fm0.txt
agrep -0 -c 'CTAGGAATACCCTAAGA' *.fastq > count/B5995_Fm0.txt
agrep -0 -c 'CCATTTGCCCGCGTTAA' *.fastq > count/B5996_Fm0.txt
agrep -0 -c 'GGGCCGGCAATAACAAT' *.fastq > count/B5999_Fm0.txt
agrep -0 -c 'AGAGGTGGTATCTCTCC' *.fastq > count/B6000_Fm0.txt
agrep -0 -c 'AATATATTGCCGGAGGA' *.fastq > count/B6001_Fm0.txt
agrep -0 -c 'CCAGATTGGCACCTGGA' *.fastq > count/B6002_Fm0.txt
agrep -0 -c 'AATCGGGTGCGCATGTT' *.fastq > count/B6003_Fm0.txt
agrep -0 -c 'CCATTGATTGCCTACTT' *.fastq > count/B6004_Fm0.txt
agrep -0 -c 'CATCGCGACATATAACT' *.fastq > count/B6005_Fm0.txt
agrep -0 -c 'CAGGATCATGCACAAGC' *.fastq > count/B6007_Fm0.txt
agrep -0 -c 'GCAAATGTCAATGAGGG' *.fastq > count/B6008_Fm0.txt
agrep -0 -c 'GATACCATTACCAGGGC' *.fastq > count/B6009_Fm0.txt
agrep -0 -c 'GCTATGAAGTAGATGCC' *.fastq > count/B6011_Fm0.txt
agrep -0 -c 'TTGCCTAGAGAACCTGC' *.fastq > count/B6012_Fm0.txt
agrep -0 -c 'ATCCGCAGGTTTGAGTA' *.fastq > count/B6013_Fm0.txt
agrep -0 -c 'CCGACCATTCGAGTTAA' *.fastq > count/B6014_Fm0.txt
agrep -0 -c 'TGTCGCTGCCATCTACG' *.fastq > count/B6015_Fm0.txt
agrep -0 -c 'CTCCAAGCCTCCTTAAC' *.fastq > count/B6018_Fm0.txt
agrep -0 -c 'CGCAAGATTCCCGAACC' *.fastq > count/B6019_Fm0.txt
agrep -0 -c 'GGAGGAGCTCATAGGTC' *.fastq > count/B6021_Fm0.txt
agrep -0 -c 'TGTACCAGCGGAGAAAG' *.fastq > count/B6022_Fm0.txt
agrep -0 -c 'GGGGGGACAAATTTGTA' *.fastq > count/B6024_Fm0.txt
agrep -0 -c 'TGCTCTGCCAATGGACT' *.fastq > count/B6025_Fm0.txt
agrep -0 -c 'CGAACCTACGATTTCCT' *.fastq > count/B6026_Fm0.txt
agrep -0 -c 'CTTATGACAAACGCTCC' *.fastq > count/B6027_Fm0.txt
agrep -0 -c 'GGATATAGTCGACGCAA' *.fastq > count/B6028_Fm0.txt
agrep -0 -c 'ACCTGACCACGATGACC' *.fastq > count/B6030_Fm0.txt
agrep -0 -c 'ATGATTGCGATCGGCCA' *.fastq > count/B6031_Fm0.txt
agrep -0 -c 'CATACTCATTGCGAGAT' *.fastq > count/B6032_Fm0.txt
agrep -0 -c 'CGCAATGGGGGCTTATC' *.fastq > count/B6034_Fm0.txt
agrep -0 -c 'TTCATCGGGTAGCCATG' *.fastq > count/B6036_Fm0.txt
agrep -0 -c 'CGGTCTAGGGTTTGGTC' *.fastq > count/B6037_Fm0.txt
agrep -0 -c 'ATGAAGAGGCGAATGCG' *.fastq > count/B6038_Fm0.txt
agrep -0 -c 'AGGTTTCTCGCTATGGT' *.fastq > count/B6040_Fm0.txt
agrep -0 -c 'AACGGTGACGCAATCGA' *.fastq > count/B6041_Fm0.txt
agrep -0 -c 'CTCGTTAACACTGCTTG' *.fastq > count/B6042_Fm0.txt
agrep -0 -c 'AAGAATGCCGTACACTC' *.fastq > count/B6043_Fm0.txt
agrep -0 -c 'TATGGCACGCAAAGTGC' *.fastq > count/B6044_Fm0.txt
agrep -0 -c 'ATGTGATGACCGGGAAC' *.fastq > count/B6045_Fm0.txt
agrep -0 -c 'AACCTAGTCAACTGCTT' *.fastq > count/B6046_Fm0.txt
agrep -0 -c 'ATCTGTTCTCCGTCATG' *.fastq > count/B6047_Fm0.txt
agrep -0 -c 'CGGTCGTATCACTATTG' *.fastq > count/B6048_Fm0.txt
agrep -0 -c 'GTAGAGCGTCGAAGGTA' *.fastq > count/B6049_Fm0.txt
agrep -0 -c 'CGACTTTCTGATTTAGC' *.fastq > count/B6050_Fm0.txt
agrep -0 -c 'TGATAGACCCTATCCGC' *.fastq > count/B6051_Fm0.txt
agrep -0 -c 'TTATTTCGCTCGCATTC' *.fastq > count/B6052_Fm0.txt
agrep -0 -c 'AATGGAGCACACGTTGG' *.fastq > count/B6053_Fm0.txt
agrep -0 -c 'TCTACTGAGTCTACGAA' *.fastq > count/B6054_Fm0.txt
agrep -0 -c 'GTCCTATTCCTTACACG' *.fastq > count/B6055_Fm0.txt
agrep -0 -c 'GTGCACATGGGCAATAC' *.fastq > count/B6056_Fm0.txt
agrep -0 -c 'GGCTCGTATTTCCCGGT' *.fastq > count/B6057_Fm0.txt
agrep -0 -c 'AGTAGTTCTGAGTCTCT' *.fastq > count/B6058_Fm0.txt
agrep -0 -c 'CGCAAGAGAAGTAATGG' *.fastq > count/B6059_Fm0.txt
agrep -0 -c 'ATCTGCGTCATAAAAGG' *.fastq > count/B6060_Fm0.txt
agrep -0 -c 'GTATAGCTCTGTCACCA' *.fastq > count/B6061_Fm0.txt
agrep -0 -c 'TGGATAATGATCCAGGC' *.fastq > count/B6062_Fm0.txt
agrep -0 -c 'CATGAGGGTGCACGAAG' *.fastq > count/B6063_Fm0.txt
agrep -0 -c 'CGGGGCAGTTTTTATAC' *.fastq > count/B6064_Fm0.txt
agrep -0 -c 'AGACCCATGGAATCACA' *.fastq > count/B6065_Fm0.txt
agrep -0 -c 'ACCTAACGATACACTGG' *.fastq > count/B6066_Fm0.txt
agrep -0 -c 'TTCGGTCTTACTCTGTG' *.fastq > count/B6068_Fm0.txt
agrep -0 -c 'TAACAGAGTGACACACC' *.fastq > count/B6069_Fm0.txt
agrep -0 -c 'AAGCCATGAAACGTTGA' *.fastq > count/B6070_Fm0.txt
agrep -0 -c 'GCAATCGTTCACGAGTG' *.fastq > count/B6072_Fm0.txt
agrep -0 -c 'GTAGATTTCGTATCCGG' *.fastq > count/B6073_Fm0.txt
agrep -0 -c 'CAAGTGAGATCGATCGT' *.fastq > count/B6074_Fm0.txt
agrep -0 -c 'AACACAGGTCAAGGCCG' *.fastq > count/B6075_Fm0.txt
agrep -0 -c 'ATCGCCGTCATTACACG' *.fastq > count/B6076_Fm0.txt
agrep -0 -c 'CATCCTTCCCAAGTTAC' *.fastq > count/B6077_Fm0.txt
agrep -0 -c 'CTTCTAAAGTCAGGATC' *.fastq > count/B6079_Fm0.txt
agrep -0 -c 'AGTCATGGTGACTACCG' *.fastq > count/B6080_Fm0.txt
agrep -0 -c 'CATTACTACGCCACACC' *.fastq > count/B6081_Fm0.txt
agrep -0 -c 'GCGATTCAGGATTCGGA' *.fastq > count/B6083_Fm0.txt
agrep -0 -c 'AGGATTAACCACACCGG' *.fastq > count/B6084_Fm0.txt
agrep -0 -c 'GTGACGACGAATGTACT' *.fastq > count/B6085_Fm0.txt
agrep -0 -c 'TGCGCAGTAAGGGTGAG' *.fastq > count/B6086_Fm0.txt
agrep -0 -c 'TCACGTTTCCGAGAAAC' *.fastq > count/B6087_Fm0.txt
agrep -0 -c 'AATAGACAACACCCCCT' *.fastq > count/B6090_Fm0.txt
agrep -0 -c 'ATACTCGGGACCACAGA' *.fastq > count/B6091_Fm0.txt
agrep -0 -c 'ATGCTCCGAACATCGAC' *.fastq > count/B6093_Fm0.txt
agrep -0 -c 'GACTCTACTCACGCTGC' *.fastq > count/B6094_Fm0.txt
agrep -0 -c 'GCAGCGGCAACGTTTGA' *.fastq > count/B6095_Fm0.txt
agrep -0 -c 'TACCACTCACAGGTAAA' *.fastq > count/B6096_Fm0.txt
agrep -0 -c 'GGGACATTAGAATGTCT' *.fastq > count/B6097_Fm0.txt
agrep -0 -c 'GGAGGATCGACTTCGTG' *.fastq > count/B6098_Fm0.txt
agrep -0 -c 'ACTCGGTCGAACGCAAC' *.fastq > count/B6099_Fm0.txt
agrep -0 -c 'CTCAGTTGTTCAGGCGC' *.fastq > count/B6100_Fm0.txt
agrep -0 -c 'CTGCCTACACTAAAGGT' *.fastq > count/B6103_Fm0.txt
agrep -0 -c 'CATAAGGCGTCCCTCTA' *.fastq > count/B6104_Fm0.txt
agrep -0 -c 'TGTCCATCAGATACGGC' *.fastq > count/B6105_Fm0.txt
agrep -0 -c 'ACTGTTCGAGTGTGCAT' *.fastq > count/B6106_Fm0.txt
agrep -0 -c 'GTCTGTGGCGTATTAAA' *.fastq > count/B6107_Fm0.txt
agrep -0 -c 'GCGCTAACACCTTGACT' *.fastq > count/B6110_Fm0.txt
agrep -0 -c 'TACACGTATCGCAGTCA' *.fastq > count/B6111_Fm0.txt
agrep -0 -c 'GCTTCTGGGACGTCTAC' *.fastq > count/B6112_Fm0.txt
agrep -0 -c 'CTATAGGACGCTGTTCG' *.fastq > count/B6113_Fm0.txt
agrep -0 -c 'TCTAGTTGTTGGGCCGC' *.fastq > count/B6114_Fm0.txt
agrep -0 -c 'GTGGAAGCAACTTCATC' *.fastq > count/B6115_Fm0.txt
agrep -0 -c 'TTGCGTTCACAGGTCGC' *.fastq > count/B6116_Fm0.txt
agrep -0 -c 'ACACAACGAACCTGCCC' *.fastq > count/B6117_Fm0.txt
agrep -0 -c 'GTTAGCCACCTATGTAC' *.fastq > count/B6118_Fm0.txt
agrep -0 -c 'AGCCCTCTTTTTATGCA' *.fastq > count/B6119_Fm0.txt
agrep -0 -c 'GGTGGCGTTGTTAACGT' *.fastq > count/B6121_Fm0.txt
agrep -0 -c 'GTCGAGAGTTCGTTCTT' *.fastq > count/B6122_Fm0.txt
agrep -0 -c 'CTTAACAGACCTCTCAT' *.fastq > count/B6123_Fm0.txt
agrep -0 -c 'GCGTACTAGTCATGTGA' *.fastq > count/B6124_Fm0.txt
agrep -0 -c 'GGATACGGAGGCTACTC' *.fastq > count/B6126_Fm0.txt
agrep -0 -c 'CATAGCGTTGTTGTAAG' *.fastq > count/B6127_Fm0.txt
agrep -0 -c 'CGGTGTATTGCTATAGG' *.fastq > count/B6128_Fm0.txt
agrep -0 -c 'TGTACTATGTCGCTAGG' *.fastq > count/B6129_Fm0.txt
agrep -0 -c 'AAGGAGTTGCGGACGTA' *.fastq > count/B6131_Fm0.txt
agrep -0 -c 'TACCCATTGGTCGATGA' *.fastq > count/B6132_Fm0.txt
agrep -0 -c 'CGTACTCTTGCGTCACG' *.fastq > count/B6133_Fm0.txt
agrep -0 -c 'AGCCTGTCAGTGCTCGT' *.fastq > count/B6134_Fm0.txt
agrep -0 -c 'TGGAAAGGCAAGGGCGT' *.fastq > count/B6135_Fm0.txt
agrep -0 -c 'GGACCTAAAAAGCGTCA' *.fastq > count/B6136_Fm0.txt
agrep -0 -c 'GTTACCCATGGCAAATA' *.fastq > count/B6137_Fm0.txt
agrep -0 -c 'AGTCTACAAGTATGGTG' *.fastq > count/B6138_Fm0.txt
agrep -0 -c 'CCGTTGACATGCAATGT' *.fastq > count/B6139_Fm0.txt
agrep -0 -c 'CCAGACGGCATGCTGAA' *.fastq > count/B6140_Fm0.txt
agrep -0 -c 'TGACACAGACCGATCAT' *.fastq > count/B6141_Fm0.txt
agrep -0 -c 'AGTTACACACGACTGAA' *.fastq > count/B6143_Fm0.txt
agrep -0 -c 'TAGTGTACTGTCAAACG' *.fastq > count/B6144_Fm0.txt
agrep -0 -c 'GGTCCTATCAACCACAT' *.fastq > count/B6145_Fm0.txt
agrep -0 -c 'CACTCTAGGTGTCTACC' *.fastq > count/B6146_Fm0.txt
agrep -0 -c 'CTGTCATACGTGTGCAG' *.fastq > count/B6147_Fm0.txt
agrep -0 -c 'CCTCACCCCCATAGCAA' *.fastq > count/B6148_Fm0.txt
agrep -0 -c 'GACCTTTGTGGGCATCG' *.fastq > count/B6149_Fm0.txt
agrep -0 -c 'TCTTGGCGAACTGCTCT' *.fastq > count/B6150_Fm0.txt
agrep -0 -c 'CCGTCTAGTATGCAACG' *.fastq > count/B6151_Fm0.txt
agrep -0 -c 'TCTAGTATCCCCATGGT' *.fastq > count/B6152_Fm0.txt
agrep -0 -c 'GGCAACTCCTAGACCAT' *.fastq > count/B6153_Fm0.txt
agrep -0 -c 'AAGGACCACGGCTCACT' *.fastq > count/B6155_Fm0.txt
agrep -0 -c 'CGCGTTTAGCGTTATGT' *.fastq > count/B6156_Fm0.txt
agrep -0 -c 'CCAACTCCTAACGCCGA' *.fastq > count/B6157_Fm0.txt
agrep -0 -c 'TCGGGTTATACCTGCGG' *.fastq > count/B6158_Fm0.txt
agrep -0 -c 'GGCAGTCCCATCAGATC' *.fastq > count/B6160_Fm0.txt
agrep -0 -c 'AACGACAGCGTACTAGA' *.fastq > count/B6161_Fm0.txt
agrep -0 -c 'TGCTTATATGGGCTATC' *.fastq > count/B6162_Fm0.txt
agrep -0 -c 'TATAACGGCATCCGTTT' *.fastq > count/B6163_Fm0.txt
agrep -0 -c 'TTCTGGGCTTAAAATCC' *.fastq > count/B6164_Fm0.txt
agrep -0 -c 'GCTGTCGACGTACATCG' *.fastq > count/B6165_Fm0.txt
agrep -0 -c 'TCGAACCACGCTACAGC' *.fastq > count/B6166_Fm0.txt
agrep -0 -c 'TGCACCCGAAGGTGACT' *.fastq > count/B6167_Fm0.txt
agrep -0 -c 'AACTAAGGGTTGACTGC' *.fastq > count/B6168_Fm0.txt
agrep -0 -c 'GTCAAAACGCATACCCC' *.fastq > count/B6169_Fm0.txt
agrep -0 -c 'GGCAATATGAGTGACAA' *.fastq > count/B6170_Fm0.txt
agrep -0 -c 'ATTGTCGGGCTTTTTCG' *.fastq > count/B6171_Fm0.txt
agrep -0 -c 'GCAGGTGCTGTAAGCGT' *.fastq > count/B6172_Fm0.txt
agrep -0 -c 'CGAAGTACAGGAAGCGA' *.fastq > count/B6173_Fm0.txt
agrep -0 -c 'CGATCTCGGTGGACATC' *.fastq > count/B6174_Fm0.txt
agrep -0 -c 'TTCGACGAATACGGACG' *.fastq > count/B6175_Fm0.txt
agrep -0 -c 'ACACGTGGGTATGTGTA' *.fastq > count/B6177_Fm0.txt
agrep -0 -c 'CGCTACATGTCGACGTT' *.fastq > count/B6178_Fm0.txt
agrep -0 -c 'TAATCGGATTCATGCCC' *.fastq > count/B6180_Fm0.txt
agrep -0 -c 'AAGTTATCGCCTTGTAC' *.fastq > count/B6182_Fm0.txt
agrep -0 -c 'AGTTTTCACTATGCACG' *.fastq > count/B6183_Fm0.txt
agrep -0 -c 'TGCAGGCATCGCTGGTA' *.fastq > count/B6184_Fm0.txt
agrep -0 -c 'GATCCGACCCGAAAGGA' *.fastq > count/B6185_Fm0.txt
agrep -0 -c 'AGAAATGAAACAGCGGC' *.fastq > count/B6186_Fm0.txt
agrep -0 -c 'AGACTTCAACACATGCG' *.fastq > count/B6187_Fm0.txt
agrep -0 -c 'TGAAGGGCGTTAGACCG' *.fastq > count/B6188_Fm0.txt
agrep -0 -c 'TGAACACTGGTAAACGA' *.fastq > count/B6189_Fm0.txt
agrep -0 -c 'ATAGATTGCGCGACCGG' *.fastq > count/B6190_Fm0.txt
agrep -0 -c 'GGATAACGGATTCCGAT' *.fastq > count/B6191_Fm0.txt
agrep -0 -c 'AGGCATCATAGCTTTCT' *.fastq > count/B6193_Fm0.txt
agrep -0 -c 'ATATCGGCAGGATTAGT' *.fastq > count/B6194_Fm0.txt
agrep -0 -c 'CGTAGGCCGCTTACTTC' *.fastq > count/B6195_Fm0.txt
agrep -0 -c 'GATAGTTACGGGGGCCT' *.fastq > count/B6196_Fm0.txt
agrep -0 -c 'TCAACCGATATGACACA' *.fastq > count/B6197_Fm0.txt
agrep -0 -c 'GATTCGACCTCTACGCT' *.fastq > count/B6198_Fm0.txt
agrep -0 -c 'TAGTACTAAGCCGTTAG' *.fastq > count/B6199_Fm0.txt
agrep -0 -c 'CTCTTACCGCACCTGTC' *.fastq > count/B6200_Fm0.txt
agrep -0 -c 'ATGGGGCTAATCGTTCC' *.fastq > count/B6201_Fm0.txt
agrep -0 -c 'CTGGTAATCTGAATCTG' *.fastq > count/B6202_Fm0.txt
agrep -0 -c 'TCAAGGGTAGGATGTCC' *.fastq > count/B6203_Fm0.txt
agrep -0 -c 'GTCGTCTTCAAGTAGGC' *.fastq > count/B6204_Fm0.txt
agrep -0 -c 'AGTGATGAACACGGCGT' *.fastq > count/B6205_Fm0.txt
agrep -0 -c 'GCAGCTTACTGTGAATC' *.fastq > count/B6206_Fm0.txt
agrep -0 -c 'AAGATCTCTTACACGCG' *.fastq > count/B6208_Fm0.txt
agrep -0 -c 'AACGGGCAACATCGAAA' *.fastq > count/B6209_Fm0.txt
agrep -0 -c 'CTGATAGTCCGGACCAG' *.fastq > count/B6212_Fm0.txt
agrep -0 -c 'GCAGCGATAAATTTCAC' *.fastq > count/B6213_Fm0.txt
agrep -0 -c 'CGCCATTCAAACCATCC' *.fastq > count/B6214_Fm0.txt
agrep -0 -c 'TTAGAATGAACGCAACG' *.fastq > count/B6215_Fm0.txt
agrep -0 -c 'AGTCGCAGTGAGAATAA' *.fastq > count/B6216_Fm0.txt
agrep -0 -c 'TGTAAAGCTAGTCCGTA' *.fastq > count/B6217_Fm0.txt
agrep -0 -c 'TGTACTAACCCTTCTCG' *.fastq > count/B6219_Fm0.txt
agrep -0 -c 'ATCCTCGGTAGTAACTT' *.fastq > count/B6220_Fm0.txt
agrep -0 -c 'TATACGCCCAAGGTAAG' *.fastq > count/B6224_Fm0.txt
agrep -0 -c 'TCTATAGGGGTTTACTG' *.fastq > count/B6225_Fm0.txt
agrep -0 -c 'CATCGTCGCAACTTACC' *.fastq > count/B6226_Fm0.txt
agrep -0 -c 'ACCAACACCGCTTGACG' *.fastq > count/B6227_Fm0.txt
agrep -0 -c 'TCGTAGCGGGTAAAAAA' *.fastq > count/B6228_Fm0.txt
agrep -0 -c 'GGCGATGTACTAAAGCC' *.fastq > count/B6229_Fm0.txt
agrep -0 -c 'GGGGCTATGTAGCTCCA' *.fastq > count/B6230_Fm0.txt
agrep -0 -c 'CACCTGGCATAGAGCGA' *.fastq > count/B6231_Fm0.txt
agrep -0 -c 'TTCTGATACCACTCGCG' *.fastq > count/B6232_Fm0.txt
agrep -0 -c 'ATGCGGATGCCAGTTAA' *.fastq > count/B6233_Fm0.txt
agrep -0 -c 'GATTTATGCCAAGGCAT' *.fastq > count/B6234_Fm0.txt
agrep -0 -c 'TTGCCCCACTAGACCTA' *.fastq > count/B6235_Fm0.txt
agrep -0 -c 'GGGAACAGTACTCACAC' *.fastq > count/B6238_Fm0.txt
agrep -0 -c 'AGAGTCCTCGGACAACT' *.fastq > count/B6239_Fm0.txt
agrep -0 -c 'GTCGGCGAAGCACACAT' *.fastq > count/B6240_Fm0.txt
agrep -0 -c 'CCTCCAGATGGCGTCTT' *.fastq > count/B6241_Fm0.txt
agrep -0 -c 'TAGTTCTCGTACCCTGG' *.fastq > count/B6242_Fm0.txt
agrep -0 -c 'ATTCAATCCGGACCAAT' *.fastq > count/B6243_Fm0.txt
agrep -0 -c 'CGCAATCATGCTGTTGG' *.fastq > count/B6244_Fm0.txt
agrep -0 -c 'CGTGTTCATGATCGTAT' *.fastq > count/B6246_Fm0.txt
agrep -0 -c 'ATTGACCAGCTACCGCG' *.fastq > count/B6247_Fm0.txt
agrep -0 -c 'AGGGAGATGAACGCCGA' *.fastq > count/B6248_Fm0.txt
agrep -0 -c 'GAGATTGACTCACTACG' *.fastq > count/B6249_Fm0.txt
agrep -0 -c 'AGTGCATCCGGTAGTGT' *.fastq > count/B6250_Fm0.txt
agrep -0 -c 'CCACAAAGTAATGGCAG' *.fastq > count/B6251_Fm0.txt
agrep -0 -c 'ATTTCACCTGCTACCCA' *.fastq > count/B6252_Fm0.txt
agrep -0 -c 'CCTTTGGGAACTCTTAA' *.fastq > count/B6253_Fm0.txt
agrep -0 -c 'GATCAATGCGAACAATC' *.fastq > count/B6255_Fm0.txt
agrep -0 -c 'TACACGCACGGACCTGA' *.fastq > count/B6256_Fm0.txt
agrep -0 -c 'TACGAAGTACTGGAAGT' *.fastq > count/B6257_Fm0.txt
agrep -0 -c 'CGCCGAAAACTGTGTCG' *.fastq > count/B6258_Fm0.txt
agrep -0 -c 'GTCAAATTGTGTCCGGT' *.fastq > count/B6259_Fm0.txt
agrep -0 -c 'CAAACGTTTGGCTAGGC' *.fastq > count/B6260_Fm0.txt
agrep -0 -c 'GGTGACGAGTATGCTAC' *.fastq > count/B6261_Fm0.txt
agrep -0 -c 'GATCTTTGCCGTGCACG' *.fastq > count/B6263_Fm0.txt
agrep -0 -c 'TTGAAGGACTCCATGGA' *.fastq > count/B6264_Fm0.txt
agrep -0 -c 'CGCCAACCTCCACCAAA' *.fastq > count/B6265_Fm0.txt
agrep -0 -c 'ACGGTAGATTATCCAGG' *.fastq > count/B6266_Fm0.txt
agrep -0 -c 'AGCGTAGTCTACTCCCC' *.fastq > count/B6267_Fm0.txt
agrep -0 -c 'CTCTTGCGGTAGCCAAA' *.fastq > count/B6268_Fm0.txt
agrep -0 -c 'GTGAGCCTATTGAAAGG' *.fastq > count/B6270_Fm0.txt
agrep -0 -c 'GATTCCGATGAGCTAGA' *.fastq > count/B6271_Fm0.txt
agrep -0 -c 'TGGCAACTCATGAACAA' *.fastq > count/B6272_Fm0.txt
agrep -0 -c 'CGCTCCTATTTCGTCAT' *.fastq > count/B6273_Fm0.txt
agrep -0 -c 'CGTATAGTAGCCTAGCG' *.fastq > count/B6274_Fm0.txt
agrep -0 -c 'CACGAGTACCGCCTTAG' *.fastq > count/B6275_Fm0.txt
agrep -0 -c 'GGGTTAACCATAGATCA' *.fastq > count/B6276_Fm0.txt
agrep -0 -c 'GTTCAAGGTCGACGTGT' *.fastq > count/B6277_Fm0.txt
agrep -0 -c 'ATGGAAGAGTCGTTTGA' *.fastq > count/B6278_Fm0.txt
agrep -0 -c 'CAAATTCATCATGTCCG' *.fastq > count/B6279_Fm0.txt
agrep -0 -c 'GAGTAAGTCTCCAGGTC' *.fastq > count/B6280_Fm0.txt
agrep -0 -c 'TCTAATAGGGGTATGTC' *.fastq > count/B6281_Fm0.txt
agrep -0 -c 'CTGCCGCACAATACGTT' *.fastq > count/B6282_Fm0.txt
agrep -0 -c 'GGGGCTCGTTATTACAG' *.fastq > count/B6283_Fm0.txt
agrep -0 -c 'CGAATGCTAGGGTTCAC' *.fastq > count/B6285_Fm0.txt
agrep -0 -c 'GTAAGACACCGGTTCAA' *.fastq > count/B6286_Fm0.txt
agrep -0 -c 'GTGGTCGATTTCTGAAG' *.fastq > count/B6288_Fm0.txt
agrep -0 -c 'AGTCAAAAGCGGACACG' *.fastq > count/B6289_Fm0.txt
agrep -0 -c 'CGAAAGATGTATAGCGA' *.fastq > count/B6290_Fm0.txt
agrep -0 -c 'TCAAGACCCCTCACAGG' *.fastq > count/B6291_Fm0.txt
agrep -0 -c 'GCGCCACCGAAAAGATC' *.fastq > count/B6293_Fm0.txt
agrep -0 -c 'AAGGGCACGGAGTAAAT' *.fastq > count/B6295_Fm0.txt
agrep -0 -c 'GTCTGATCTCATGCATC' *.fastq > count/B6296_Fm0.txt
agrep -0 -c 'ATACTGCCGTTTCTGGT' *.fastq > count/B6297_Fm0.txt
agrep -0 -c 'ATTTTCGGCTTTTGGGT' *.fastq > count/B6299_Fm0.txt
agrep -0 -c 'ATCTGTGATAAGTCGGT' *.fastq > count/B6301_Fm0.txt
agrep -0 -c 'TAGTTTCGCACCGCAGT' *.fastq > count/B6302_Fm0.txt
agrep -0 -c 'CATGTCGGAGAGGTGTC' *.fastq > count/B6304_Fm0.txt
agrep -0 -c 'AACCCGTTATAGGGGTT' *.fastq > count/B6306_Fm0.txt
agrep -0 -c 'GCAAGCGAGCTAGACTA' *.fastq > count/B6307_Fm0.txt
agrep -0 -c 'TAGACGTTAAGGACCGA' *.fastq > count/B6308_Fm0.txt
agrep -0 -c 'GACTCGTCAAGCAAAGG' *.fastq > count/B6309_Fm0.txt
agrep -0 -c 'GTTGACGCGGTCCTACA' *.fastq > count/B6310_Fm0.txt
agrep -0 -c 'TTCAGTCCGTGCCTTTT' *.fastq > count/B6311_Fm0.txt
agrep -0 -c 'AAGTCCTCGTCATTTCT' *.fastq > count/B6312_Fm0.txt
agrep -0 -c 'TGAAAGGGGGCACAATT' *.fastq > count/B6313_Fm0.txt
agrep -0 -c 'GAGGGTTCATAGTAAGG' *.fastq > count/B6314_Fm0.txt
agrep -0 -c 'TCATGTGGGGGTTAATA' *.fastq > count/B6315_Fm0.txt
agrep -0 -c 'GTGAGCTTGTCGCACAC' *.fastq > count/B6316_Fm0.txt
agrep -0 -c 'AGTCTTAGCGTCGGGGT' *.fastq > count/B6317_Fm0.txt
agrep -0 -c 'TGACCTAGCGGTGAGGA' *.fastq > count/B6318_Fm0.txt
agrep -0 -c 'GCTGGCTATCAAATGGA' *.fastq > count/B6319_Fm0.txt
agrep -0 -c 'CGCAAATCTTCAGTAGC' *.fastq > count/B6320_Fm0.txt
agrep -0 -c 'TTCCAGCATAAGAGGAA' *.fastq > count/B6321_Fm0.txt
agrep -0 -c 'GTTAAGATTCCGTTGCG' *.fastq > count/B6322_Fm0.txt
agrep -0 -c 'CCTATCGTAGTTCATGT' *.fastq > count/B6323_Fm0.txt
agrep -0 -c 'TGCCAACCTTCGACAAT' *.fastq > count/B6324_Fm0.txt
agrep -0 -c 'ACATAAAGTAACCCTCG' *.fastq > count/B6325_Fm0.txt
agrep -0 -c 'GATGATTAGGTACTCGA' *.fastq > count/B6326_Fm0.txt
agrep -0 -c 'AGCTTCATACCAAGTCT' *.fastq > count/B6327_Fm0.txt
agrep -0 -c 'CTTCGACCAATAACAGC' *.fastq > count/B6328_Fm0.txt
agrep -0 -c 'AAGTTACATGTGCCTGA' *.fastq > count/B6329_Fm0.txt
agrep -0 -c 'TATTCTAAAGGGAAGGG' *.fastq > count/B6331_Fm0.txt
agrep -0 -c 'CTACGCCAACATCTCTT' *.fastq > count/B6332_Fm0.txt
agrep -0 -c 'CGATATCCTGTCATGGG' *.fastq > count/B6333_Fm0.txt
agrep -0 -c 'CGGAAACTAGTAGGTCC' *.fastq > count/B6334_Fm0.txt
agrep -0 -c 'CGCACACGGAGCATTTT' *.fastq > count/B6335_Fm0.txt
agrep -0 -c 'ATAATGCACTTGGCCGC' *.fastq > count/B6336_Fm0.txt
agrep -0 -c 'CGGCGACATCGTTAAAA' *.fastq > count/B6338_Fm0.txt
agrep -0 -c 'ACCTGGCATTTGTTCGT' *.fastq > count/B6339_Fm0.txt
agrep -0 -c 'TCGCGGAAGATCTGCCT' *.fastq > count/B6340_Fm0.txt
agrep -0 -c 'GACTTATGGAGTGTCGA' *.fastq > count/B6341_Fm0.txt
agrep -0 -c 'TCTATGAACCGGCCTAT' *.fastq > count/B6342_Fm0.txt
agrep -0 -c 'CTGCTAGCCTAATGGTC' *.fastq > count/B6343_Fm0.txt
agrep -0 -c 'GTAGGGCACTAATCCCG' *.fastq > count/B6344_Fm0.txt
agrep -0 -c 'CAAAGAAACGACCCGTT' *.fastq > count/B6345_Fm0.txt
agrep -0 -c 'TACACTAACTAGCCACC' *.fastq > count/B6346_Fm0.txt
agrep -0 -c 'TTCGACGCCATAGGAGT' *.fastq > count/B6347_Fm0.txt
agrep -0 -c 'AGAGAACCCGCACAGTC' *.fastq > count/B6348_Fm0.txt
agrep -0 -c 'GTTTTATGTGCGCTAGC' *.fastq > count/B6349_Fm0.txt
agrep -0 -c 'TAGGTCCCGGACATAAT' *.fastq > count/B6350_Fm0.txt
agrep -0 -c 'ACAAGTGATCACGCCAG' *.fastq > count/B6351_Fm0.txt
agrep -0 -c 'TACTCGGACTGTAGACA' *.fastq > count/B6352_Fm0.txt
agrep -0 -c 'TATCATTACTAGGCCAG' *.fastq > count/B6353_Fm0.txt
agrep -0 -c 'CCTCCGTAAGGGATCGT' *.fastq > count/B6354_Fm0.txt
agrep -0 -c 'AATAATCGTGCCACCAG' *.fastq > count/B6355_Fm0.txt
agrep -0 -c 'GGTCAAAGCGCAATAAT' *.fastq > count/B6356_Fm0.txt
agrep -0 -c 'ACCACCGGCTAGATGTG' *.fastq > count/B6359_Fm0.txt
agrep -0 -c 'GGCAACTGCGCGTTTAA' *.fastq > count/B6361_Fm0.txt
agrep -0 -c 'TATCCATGCATGCACGT' *.fastq > count/B6362_Fm0.txt
agrep -0 -c 'AAGGGCTCTGCCTAAAC' *.fastq > count/B6363_Fm0.txt
agrep -0 -c 'ACCAAGGCTTAAGGACA' *.fastq > count/B6364_Fm0.txt
agrep -0 -c 'GGTATGTGACGCGGATA' *.fastq > count/B6365_Fm0.txt
agrep -0 -c 'CCATGTTAACGACAAGA' *.fastq > count/B6366_Fm0.txt
agrep -0 -c 'CTTGGAAAGAGATGTCT' *.fastq > count/B6368_Fm0.txt
agrep -0 -c 'GGTCAAGGATGTGGACA' *.fastq > count/B6369_Fm0.txt
agrep -0 -c 'GGGAATGATGCTTCGAG' *.fastq > count/B6370_Fm0.txt
agrep -0 -c 'CGATAGCATGCTCGAGG' *.fastq > count/B6372_Fm0.txt
agrep -0 -c 'GATGCGCTTTTCCGGTT' *.fastq > count/B6373_Fm0.txt
agrep -0 -c 'TGGAATGTGGCTCGACG' *.fastq > count/B6374_Fm0.txt
agrep -0 -c 'TCAGTCGGCGTCTCGTA' *.fastq > count/B6375_Fm0.txt
agrep -0 -c 'GCAGTCTGAATGTCACC' *.fastq > count/B6376_Fm0.txt
agrep -0 -c 'CTACGAGCCGAATATGT' *.fastq > count/B6377_Fm0.txt
agrep -0 -c 'ATCTTAGTAGCGCGTAG' *.fastq > count/B6379_Fm0.txt
agrep -0 -c 'AGGTGACATTAGGTCGC' *.fastq > count/B6380_Fm0.txt
agrep -0 -c 'CCGTGGAGGACTTAAGG' *.fastq > count/B6381_Fm0.txt
agrep -0 -c 'CCGGCCAGAAATGTCTC' *.fastq > count/B6382_Fm0.txt
agrep -0 -c 'TCGGGAGGTGTTTTAGA' *.fastq > count/B6384_Fm0.txt
agrep -0 -c 'TAGCGATATCTGTGGCC' *.fastq > count/B6385_Fm0.txt
agrep -0 -c 'GGCGTAGTCTTTACGCG' *.fastq > count/B6386_Fm0.txt
agrep -0 -c 'TATTCGACTTCCATGTC' *.fastq > count/B6388_Fm0.txt
agrep -0 -c 'GGCGGCTAGTAATAGCC' *.fastq > count/B6389_Fm0.txt
agrep -0 -c 'TTGACCAGCGGACACTT' *.fastq > count/B6390_Fm0.txt
agrep -0 -c 'AAGTATTCTCCCCGACG' *.fastq > count/B6391_Fm0.txt
agrep -0 -c 'TATCGTCGAGGCGCATT' *.fastq > count/B6392_Fm0.txt
agrep -0 -c 'AAGTCGTCACGGTATGC' *.fastq > count/B6393_Fm0.txt
agrep -0 -c 'GGAGTATCAACAAAGTC' *.fastq > count/B6394_Fm0.txt
agrep -0 -c 'ACAAACCCCTTAAAGCC' *.fastq > count/B6395_Fm0.txt
agrep -0 -c 'TTGAGTGGTCGTTGATC' *.fastq > count/B6396_Fm0.txt
agrep -0 -c 'CCGGTGCGAGCTTTGAA' *.fastq > count/B6397_Fm0.txt
agrep -0 -c 'GGGAACACTGTACTCTC' *.fastq > count/B6398_Fm0.txt
agrep -0 -c 'GTTTGGCGATTCGCCTC' *.fastq > count/B6399_Fm0.txt
agrep -0 -c 'GTAGCCTAGTATCCACG' *.fastq > count/B6401_Fm0.txt
agrep -0 -c 'GACACCCATGGTCGATT' *.fastq > count/B6402_Fm0.txt
agrep -0 -c 'AGGTGACTCCATACTCG' *.fastq > count/B6403_Fm0.txt
agrep -0 -c 'GCGAACGCTTGAGTCTT' *.fastq > count/B6404_Fm0.txt
agrep -0 -c 'TTATATTGATCTGGCGG' *.fastq > count/B6405_Fm0.txt
agrep -0 -c 'GTTCCTCATGTGCAGCG' *.fastq > count/B6407_Fm0.txt
agrep -0 -c 'AGCTGATACAGTGCGCA' *.fastq > count/B6408_Fm0.txt
agrep -0 -c 'CGTCGCTAAATCTCACC' *.fastq > count/B6409_Fm0.txt
agrep -0 -c 'AACAGCAAGCCATAGTC' *.fastq > count/B6411_Fm0.txt
agrep -0 -c 'TGTCGGACTAGATGATT' *.fastq > count/B6412_Fm0.txt
agrep -0 -c 'GGCTAAAGTTATGAACG' *.fastq > count/B6413_Fm0.txt
agrep -0 -c 'AACGGAACAGGCTATTG' *.fastq > count/B6414_Fm0.txt
agrep -0 -c 'ATGAGCTAGCTTTTCCC' *.fastq > count/B6415_Fm0.txt
agrep -0 -c 'TTCCTATAAGCGCGTGA' *.fastq > count/B6416_Fm0.txt
agrep -0 -c 'AGGGACCTAGTGGATAC' *.fastq > count/B6418_Fm0.txt
agrep -0 -c 'AGCCTTTCAGCGCCAAC' *.fastq > count/B6420_Fm0.txt
agrep -0 -c 'CTTATCCAATCGGAGAT' *.fastq > count/B6421_Fm0.txt
agrep -0 -c 'TGTGTAAGGTCCAGCGC' *.fastq > count/B6422_Fm0.txt
agrep -0 -c 'CTCTGAACGTCGTACGC' *.fastq > count/B6423_Fm0.txt
agrep -0 -c 'GCGTTATATGCTGCCAA' *.fastq > count/B6424_Fm0.txt
agrep -0 -c 'AAGTAGGAAAGGCATTG' *.fastq > count/B6425_Fm0.txt
agrep -0 -c 'GTCTGAAGTAGTACGCT' *.fastq > count/B6426_Fm0.txt
agrep -0 -c 'GATTGCAAGCTTAACGG' *.fastq > count/B6427_Fm0.txt
agrep -0 -c 'CATTCTAGGTTTTCGCT' *.fastq > count/B6429_Fm0.txt
agrep -0 -c 'ATCGATAGGGTCTCTCG' *.fastq > count/B6430_Fm0.txt
agrep -0 -c 'ATCGTGCTTCATCGCAA' *.fastq > count/B6432_Fm0.txt
agrep -0 -c 'AGCACGAACTATCAAGT' *.fastq > count/B6433_Fm0.txt
agrep -0 -c 'GACTAGCAACTGCCGTA' *.fastq > count/B6435_Fm0.txt
agrep -0 -c 'TCTGTCCGCTGGAATCG' *.fastq > count/B6436_Fm0.txt
agrep -0 -c 'CACTTTACGTCCTCTCT' *.fastq > count/B6437_Fm0.txt
agrep -0 -c 'GTGGTCGTTACTGCGTG' *.fastq > count/B6439_Fm0.txt
agrep -0 -c 'TGCCGCAATAGGGAGAC' *.fastq > count/B6440_Fm0.txt
agrep -0 -c 'GGGGGACTTCCGTTTTT' *.fastq > count/B6441_Fm0.txt
agrep -0 -c 'GTCGCGTCATTGTAGCA' *.fastq > count/B6442_Fm0.txt
agrep -0 -c 'CGAGCATGTTGGATATT' *.fastq > count/B6443_Fm0.txt
agrep -0 -c 'TGAGGCCCTTCGATACG' *.fastq > count/B6444_Fm0.txt
agrep -0 -c 'CTGATCGTACGCATATG' *.fastq > count/B6445_Fm0.txt
agrep -0 -c 'GAGTCCAGGCGTGCTAT' *.fastq > count/B6446_Fm0.txt
agrep -0 -c 'GTTTCGCGCGTTACCAG' *.fastq > count/B6447_Fm0.txt
agrep -0 -c 'TAAGCTCCTGAAGGTTT' *.fastq > count/B6449_Fm0.txt
agrep -0 -c 'TATCATGGCCGGTATAT' *.fastq > count/B6450_Fm0.txt
agrep -0 -c 'ACCATGCGTGCAAACAC' *.fastq > count/B6451_Fm0.txt
agrep -0 -c 'GCGATCTTCGGCGCAAT' *.fastq > count/B6452_Fm0.txt
agrep -0 -c 'TTACAGTCGCCGAAGTG' *.fastq > count/B6453_Fm0.txt
agrep -0 -c 'GGTGTCAGCCTCAACTG' *.fastq > count/B6454_Fm0.txt
agrep -0 -c 'GCCCGAGGACTATTACG' *.fastq > count/B6455_Fm0.txt
agrep -0 -c 'CCGTTTGGTATTGGGCA' *.fastq > count/B6457_Fm0.txt
agrep -0 -c 'GGATCAAACGGAAAAGG' *.fastq > count/B6458_Fm0.txt
agrep -0 -c 'AGGAACTCTAGACCCAA' *.fastq > count/B6459_Fm0.txt
agrep -0 -c 'AGATGTAGACAAACGTG' *.fastq > count/B6460_Fm0.txt
agrep -0 -c 'TAAGAACCTCGGTAGCG' *.fastq > count/B6461_Fm0.txt
agrep -0 -c 'TATCTAACGTGGGCAAG' *.fastq > count/B6462_Fm0.txt
agrep -0 -c 'GCTAAGGATCTTGGCGT' *.fastq > count/B6463_Fm0.txt
agrep -0 -c 'GTCTTAGCGCCCAGTTC' *.fastq > count/B6464_Fm0.txt
agrep -0 -c 'CAGGCTCGTACGAAATC' *.fastq > count/B6465_Fm0.txt
agrep -0 -c 'CCAGACCGAGACAACAT' *.fastq > count/B6466_Fm0.txt
agrep -0 -c 'CAGGTATCCCTTAAACT' *.fastq > count/B6468_Fm0.txt
agrep -0 -c 'CAAACAAGGAACGACAT' *.fastq > count/B6469_Fm0.txt
agrep -0 -c 'GAGTGAATCGTTACATC' *.fastq > count/B6471_Fm0.txt
agrep -0 -c 'TTGCCCTTGTAAACAGT' *.fastq > count/B6472_Fm0.txt
agrep -0 -c 'GCAAAGATACTGCCTAT' *.fastq > count/B6473_Fm0.txt
agrep -0 -c 'CTACGAAAGGACCCGAT' *.fastq > count/B6474_Fm0.txt
agrep -0 -c 'CGCCCATATTGAAACCA' *.fastq > count/B6477_Fm0.txt
agrep -0 -c 'ACAGTCGTAAGGAGTTC' *.fastq > count/B6478_Fm0.txt
agrep -0 -c 'CTGTCCGCAGCATACGA' *.fastq > count/B6479_Fm0.txt
agrep -0 -c 'CTACGATATGACCCTGA' *.fastq > count/B6480_Fm0.txt
agrep -0 -c 'GGAACCGATATGAATAC' *.fastq > count/B6483_Fm0.txt
agrep -0 -c 'GGAGCCATTTATGTCGT' *.fastq > count/B6484_Fm0.txt
agrep -0 -c 'CATCATAGCACGCTGAA' *.fastq > count/B6485_Fm0.txt
agrep -0 -c 'AGTAAGCGAGCAGACTC' *.fastq > count/B6486_Fm0.txt
agrep -0 -c 'GGAAGGGGGGGAGATAA' *.fastq > count/B6487_Fm0.txt
agrep -0 -c 'AGACGATGTTTGACTGG' *.fastq > count/B6488_Fm0.txt
agrep -0 -c 'GGTCTAAACTTTCAGGC' *.fastq > count/B6489_Fm0.txt
agrep -0 -c 'CCTGTCGGGACAGTCTT' *.fastq > count/B6490_Fm0.txt
agrep -0 -c 'TGAATACAGCGACAACA' *.fastq > count/B6492_Fm0.txt
agrep -0 -c 'AGCCTAGGACCTTTTCA' *.fastq > count/B6493_Fm0.txt
agrep -0 -c 'ACCTCGCTGTCACACAG' *.fastq > count/B6494_Fm0.txt
agrep -0 -c 'AGACCCAATCCCGACAG' *.fastq > count/B6495_Fm0.txt
agrep -0 -c 'ACATACGTGCTATGTCA' *.fastq > count/B6496_Fm0.txt
agrep -0 -c 'AGCACAGGTTAAGTGAA' *.fastq > count/B6497_Fm0.txt
agrep -0 -c 'AGGGTTACTGGGTCTGC' *.fastq > count/B6498_Fm0.txt
agrep -0 -c 'TGGATACAATGGGCTCT' *.fastq > count/B6499_Fm0.txt
agrep -0 -c 'AGGGCCTACCTATCACA' *.fastq > count/B6500_Fm0.txt
agrep -0 -c 'GTGAGCGCAAGACGATA' *.fastq > count/B6501_Fm0.txt
agrep -0 -c 'TTGCAGTTGTAACCCTA' *.fastq > count/B6502_Fm0.txt
agrep -0 -c 'GATCCGTTCACCATAAG' *.fastq > count/B6504_Fm0.txt
agrep -0 -c 'AAGACCTGAGGGTTGCT' *.fastq > count/B6505_Fm0.txt
agrep -0 -c 'CAGTGCACTGCTCCAGA' *.fastq > count/B6506_Fm0.txt
agrep -0 -c 'TAACTCCGCACAGGCAT' *.fastq > count/B6508_Fm0.txt
agrep -0 -c 'TCTGAAGCGATGCCAAC' *.fastq > count/B6509_Fm0.txt
agrep -0 -c 'GGTCACGTCTCAAAGCG' *.fastq > count/B6510_Fm0.txt
agrep -0 -c 'TATGTGCTATGGGCAAA' *.fastq > count/B6511_Fm0.txt
agrep -0 -c 'TGCGAAGTCACTACTTA' *.fastq > count/B6514_Fm0.txt
agrep -0 -c 'CTTACGCGGAGGTCGAA' *.fastq > count/B6515_Fm0.txt
agrep -0 -c 'ATTCTAGTTAGCGCGAA' *.fastq > count/B6516_Fm0.txt
agrep -0 -c 'CTTAAAAGTGTGTGCCC' *.fastq > count/B6517_Fm0.txt
agrep -0 -c 'TGAGTGTTACACAGTTC' *.fastq > count/B6518_Fm0.txt
agrep -0 -c 'CATACGTCCACTGCTAC' *.fastq > count/B6520_Fm0.txt
agrep -0 -c 'GTTAGCTGGGATGCCAT' *.fastq > count/B6521_Fm0.txt
agrep -0 -c 'TATGGCACACTTGGCGG' *.fastq > count/B6522_Fm0.txt
agrep -0 -c 'CGAGTATCTTTCTTCCC' *.fastq > count/B6523_Fm0.txt
agrep -0 -c 'TAGAGGTGTCGCACAGT' *.fastq > count/B6524_Fm0.txt
agrep -0 -c 'ACCATAATTCAACGTGC' *.fastq > count/B6525_Fm0.txt
agrep -0 -c 'ACTCGACTCAGCGGCTT' *.fastq > count/B6526_Fm0.txt
agrep -0 -c 'TATGATGCCAACCAGAG' *.fastq > count/B6527_Fm0.txt
agrep -0 -c 'ATACGGTTCCGAGACGT' *.fastq > count/B6528_Fm0.txt
agrep -0 -c 'GCACTAACCCTACAGAG' *.fastq > count/B6529_Fm0.txt
agrep -0 -c 'ACGATACTTAACGGCCG' *.fastq > count/B6531_Fm0.txt
agrep -0 -c 'TTCCGTCGCGAGGAAAC' *.fastq > count/B6532_Fm0.txt
agrep -0 -c 'TGTAGTCTTTGGTTGCC' *.fastq > count/B6534_Fm0.txt
agrep -0 -c 'GGTTGGCCCTTGATCTC' *.fastq > count/B6536_Fm0.txt
agrep -0 -c 'GACAACAGCATTAAGCA' *.fastq > count/B6537_Fm0.txt
agrep -0 -c 'TTCCTTTAGGGTGACCA' *.fastq > count/B6538_Fm0.txt
agrep -0 -c 'CGAGGACTGCGATATGG' *.fastq > count/B6540_Fm0.txt
agrep -0 -c 'TAGTCTGCCATTAGACA' *.fastq > count/B6541_Fm0.txt
agrep -0 -c 'TACGCCCTAGGTTTGAG' *.fastq > count/B6542_Fm0.txt
agrep -0 -c 'GGTTCAGAACTGCGCAG' *.fastq > count/B6543_Fm0.txt
agrep -0 -c 'ATTGGGTGATCGTTTGT' *.fastq > count/B6544_Fm0.txt
agrep -0 -c 'TACTGTCCGAGACGCAA' *.fastq > count/B6545_Fm0.txt
agrep -0 -c 'TGCCGTTCATGTTACGA' *.fastq > count/B6546_Fm0.txt
agrep -0 -c 'CTGCGATACTGCAAGCA' *.fastq > count/B6547_Fm0.txt
agrep -0 -c 'GTCCTTACCAAAAGTAG' *.fastq > count/B6548_Fm0.txt
agrep -0 -c 'TTACGTTGCAGTGGAAA' *.fastq > count/B6549_Fm0.txt
agrep -0 -c 'CAGACCACGTACACATC' *.fastq > count/B6550_Fm0.txt
agrep -0 -c 'AAGAGCTGTTACGAGCA' *.fastq > count/B6552_Fm0.txt
agrep -0 -c 'AAGTACGTTCGTAAGGG' *.fastq > count/B6553_Fm0.txt
agrep -0 -c 'GATACCGCTTGGAATAC' *.fastq > count/B6555_Fm0.txt
agrep -0 -c 'ACTGAGCGATGGCTCCT' *.fastq > count/B6556_Fm0.txt
agrep -0 -c 'TGCAAAAATAGGCGTCG' *.fastq > count/B6557_Fm0.txt
agrep -0 -c 'CGACTGATTTTGATCAG' *.fastq > count/B6558_Fm0.txt
agrep -0 -c 'GCAATGTACCTGCAATA' *.fastq > count/B6559_Fm0.txt
agrep -0 -c 'GGTGGACAAACTGTTAC' *.fastq > count/B6560_Fm0.txt
agrep -0 -c 'CATTCCCGTGATCCCCA' *.fastq > count/B6561_Fm0.txt
agrep -0 -c 'ACCATGTCCCCTTCAGG' *.fastq > count/B6562_Fm0.txt
agrep -0 -c 'GCCTCGGCTAACTTGTA' *.fastq > count/B6563_Fm0.txt
agrep -0 -c 'TTGCTTGACGCACATTC' *.fastq > count/B6564_Fm0.txt
agrep -0 -c 'GCAGGGAATTGTCTCGC' *.fastq > count/B6565_Fm0.txt
agrep -0 -c 'TTCGTCCCACGCGTTTT' *.fastq > count/B6567_Fm0.txt
agrep -0 -c 'TTCAGGCAACGTGTATC' *.fastq > count/B6568_Fm0.txt
agrep -0 -c 'CTTGTAGCGCCTTTGAT' *.fastq > count/B6569_Fm0.txt
agrep -0 -c 'GGACATAGGTTTGTAGC' *.fastq > count/B6570_Fm0.txt
agrep -0 -c 'GCTCAAAGTGGTTACGC' *.fastq > count/B6571_Fm0.txt
agrep -0 -c 'ACCTCTAAATCGAACCT' *.fastq > count/B6572_Fm0.txt
agrep -0 -c 'GATGTCATGGACGTTCA' *.fastq > count/B6573_Fm0.txt
agrep -0 -c 'TACCCAGTGCTGGACTA' *.fastq > count/B6575_Fm0.txt
agrep -0 -c 'CCACCTTAGGCACTCTG' *.fastq > count/B6576_Fm0.txt
agrep -0 -c 'ATCGTCTTAGCCAACCA' *.fastq > count/B6577_Fm0.txt
agrep -0 -c 'GCGTTTTCCTGTGGTGG' *.fastq > count/B6578_Fm0.txt
agrep -0 -c 'TGTGAGTCTTTGTGCGA' *.fastq > count/B6579_Fm0.txt
agrep -0 -c 'GGAGCGAAGTTAACGCT' *.fastq > count/B6580_Fm0.txt
agrep -0 -c 'GTTTTGGGCCCACAAGA' *.fastq > count/B6581_Fm0.txt
agrep -0 -c 'CAACCCACCAGTTTTTA' *.fastq > count/B6585_Fm0.txt
agrep -0 -c 'CCGCATTTGATTTGGCT' *.fastq > count/B6586_Fm0.txt
agrep -0 -c 'ATACCGTTTTAGGGCTT' *.fastq > count/B6587_Fm0.txt
agrep -0 -c 'TATAGTATAGGCGCGTA' *.fastq > count/B6589_Fm0.txt
agrep -0 -c 'CCTATGACGCAAACATC' *.fastq > count/B6591_Fm0.txt
agrep -0 -c 'ACCGCCAATCCAAAACC' *.fastq > count/B6592_Fm0.txt
agrep -0 -c 'AAGTGTAGTACGGACCG' *.fastq > count/B6593_Fm0.txt
agrep -0 -c 'ACTGGGTTGACGTATCC' *.fastq > count/B6594_Fm0.txt
agrep -0 -c 'GCTGGGTGGTGATCCAA' *.fastq > count/B6597_Fm0.txt
agrep -0 -c 'ACGACTGACCTTGCATT' *.fastq > count/B6598_Fm0.txt
agrep -0 -c 'AATCCTCGTGAGAACTG' *.fastq > count/B6599_Fm0.txt
agrep -0 -c 'GACGGAATGGGCTTCGT' *.fastq > count/B6600_Fm0.txt
agrep -0 -c 'TGCTTAGGGTCGATTCC' *.fastq > count/B6601_Fm0.txt
agrep -0 -c 'AGATACTCTAGCATGTG' *.fastq > count/B6602_Fm0.txt
agrep -0 -c 'CCAGTTACCTTGCAAGG' *.fastq > count/B6604_Fm0.txt
agrep -0 -c 'AGGCTCCGCGTGCATAA' *.fastq > count/B6606_Fm0.txt
agrep -0 -c 'AGACACTTACCCCAGGG' *.fastq > count/B6607_Fm0.txt
agrep -0 -c 'AAGGCAAAAACTAGTCG' *.fastq > count/B6608_Fm0.txt
agrep -0 -c 'TGTCATCGACGATACGT' *.fastq > count/B6609_Fm0.txt
agrep -0 -c 'AGACGTGGACCCATGAT' *.fastq > count/B6610_Fm0.txt
agrep -0 -c 'CAAAGCCATCCAGAGGT' *.fastq > count/B6611_Fm0.txt
agrep -0 -c 'CAGTGTTCTTGACTACG' *.fastq > count/B6612_Fm0.txt
agrep -0 -c 'GCGAAAGTATCCCTCTA' *.fastq > count/B6613_Fm0.txt
agrep -0 -c 'CAACGTTGAATAGTAGC' *.fastq > count/B6614_Fm0.txt
agrep -0 -c 'AGGAGCGAAAAGCGGGA' *.fastq > count/B6615_Fm0.txt
agrep -0 -c 'GACATCTCTTGAGACGA' *.fastq > count/B6617_Fm0.txt
agrep -0 -c 'CTAGCTGGTTAGCAGGG' *.fastq > count/B6619_Fm0.txt
agrep -0 -c 'ACGATCCCGATTGACCC' *.fastq > count/B6621_Fm0.txt
agrep -0 -c 'CGAGACAAGTGACCAGC' *.fastq > count/B6624_Fm0.txt
agrep -0 -c 'AGGGAGCCGACTGAATT' *.fastq > count/B6625_Fm0.txt
agrep -0 -c 'ACATGGTGTAGCATCGT' *.fastq > count/B6626_Fm0.txt
agrep -0 -c 'TTCCGCTGACTACGTGC' *.fastq > count/B6627_Fm0.txt
agrep -0 -c 'GCATACAGCTGATGGAA' *.fastq > count/B6628_Fm0.txt
agrep -0 -c 'CATGGTTGAACAAGCTT' *.fastq > count/B6629_Fm0.txt
agrep -0 -c 'GGGGCCTGCTTTAAGTC' *.fastq > count/B6630_Fm0.txt
agrep -0 -c 'AGTGCGCGGGATTTCGA' *.fastq > count/B6633_Fm0.txt
agrep -0 -c 'CGGAAATGCTAAAGACT' *.fastq > count/B6634_Fm0.txt
agrep -0 -c 'TTCATTGCCCTTAGCCC' *.fastq > count/B6635_Fm0.txt
agrep -0 -c 'GTTGATACACATCGATC' *.fastq > count/B6636_Fm0.txt
agrep -0 -c 'GCTAGACGCCGTAAACT' *.fastq > count/B6637_Fm0.txt
agrep -0 -c 'AACTGCGTGCGGGTCTA' *.fastq > count/B6639_Fm0.txt
agrep -0 -c 'ATCCGTACTAGGCGTAT' *.fastq > count/B6641_Fm0.txt
agrep -0 -c 'TCAAGATCACCTGCGTG' *.fastq > count/B6642_Fm0.txt
agrep -0 -c 'TGTAGCTACGCGTAGCC' *.fastq > count/B6645_Fm0.txt
agrep -0 -c 'GAGAAATACCGCAGCTG' *.fastq > count/B6646_Fm0.txt
agrep -0 -c 'GGTGTACGCTTTGGTTC' *.fastq > count/B6647_Fm0.txt
agrep -0 -c 'CACTGTACGTTTGCATA' *.fastq > count/B6648_Fm0.txt
agrep -0 -c 'AACGGAGGCTAAGGGAT' *.fastq > count/B6649_Fm0.txt
agrep -0 -c 'GATTATTGTACCTCAGC' *.fastq > count/B6650_Fm0.txt
agrep -0 -c 'CGGTTTGTAGGAGCCTC' *.fastq > count/B6652_Fm0.txt
agrep -0 -c 'AACACTATGCGTCATCA' *.fastq > count/B6653_Fm0.txt
agrep -0 -c 'TCAGAGACATGCTACCG' *.fastq > count/B6655_Fm0.txt
agrep -0 -c 'TAGCGCGAGGGTAAACA' *.fastq > count/B6657_Fm0.txt
agrep -0 -c 'ACCATATCGTCAACTGC' *.fastq > count/B6659_Fm0.txt
agrep -0 -c 'TGCCGGTACATTCGTAG' *.fastq > count/B6660_Fm0.txt
agrep -0 -c 'CGAGACAAAATAGAGGT' *.fastq > count/B6661_Fm0.txt
agrep -0 -c 'AGTGCAGTGATGCGTGA' *.fastq > count/B6662_Fm0.txt
agrep -0 -c 'CTCGGAGACAAGCTATC' *.fastq > count/B6663_Fm0.txt
agrep -0 -c 'CGTAGGCATACTCCCAC' *.fastq > count/B6664_Fm0.txt
agrep -0 -c 'GACCACGGGACAATCAA' *.fastq > count/B6666_Fm0.txt
agrep -0 -c 'TGTGACTTCGCGGAGCA' *.fastq > count/B6667_Fm0.txt
agrep -0 -c 'GTATAGGCTCGTATTGT' *.fastq > count/B6668_Fm0.txt
agrep -0 -c 'CAGTGGGGCCTTCTATG' *.fastq > count/B6669_Fm0.txt
agrep -0 -c 'GAGTTACCGTGGAACAG' *.fastq > count/B6670_Fm0.txt
agrep -0 -c 'GTCTTAGTAAAGGCTTG' *.fastq > count/B6671_Fm0.txt
agrep -0 -c 'CATACCTTCGATTCCGC' *.fastq > count/B6672_Fm0.txt
agrep -0 -c 'GATGCACTCGAGATTCA' *.fastq > count/B6673_Fm0.txt
agrep -0 -c 'CGACTTATCGAGCGCTG' *.fastq > count/B6674_Fm0.txt
agrep -0 -c 'TCGGGATGATCAGCGAA' *.fastq > count/B6675_Fm0.txt
agrep -0 -c 'GCTGCAATCCCAGAACA' *.fastq > count/B6676_Fm0.txt
agrep -0 -c 'CGTTTGCGGGTCACTAT' *.fastq > count/B6677_Fm0.txt
agrep -0 -c 'TAACACCCTATCATTGG' *.fastq > count/B6678_Fm0.txt
agrep -0 -c 'GTATCACCTTGGCGGTT' *.fastq > count/B6679_Fm0.txt
agrep -0 -c 'GGTAGTGAGGTTAGTAG' *.fastq > count/B6680_Fm0.txt
agrep -0 -c 'AGAGTCTGTCTATGGCA' *.fastq > count/B6681_Fm0.txt
agrep -0 -c 'TATAACTCGGATCCACA' *.fastq > count/B6682_Fm0.txt
agrep -0 -c 'AGTAAAGCGGTGACGGT' *.fastq > count/B6683_Fm0.txt
agrep -0 -c 'GTGCTTGGGTCGAGCTA' *.fastq > count/B6684_Fm0.txt
agrep -0 -c 'GATAATGGTAGCAACGG' *.fastq > count/B6685_Fm0.txt
agrep -0 -c 'TCGGGCCTACCAGCAAA' *.fastq > count/B6687_Fm0.txt
agrep -0 -c 'CATAATTATGGCACGGC' *.fastq > count/B6688_Fm0.txt
agrep -0 -c 'TACCCGAATGGATGAGG' *.fastq > count/B6689_Fm0.txt
agrep -0 -c 'AGCCTCCATTCGTACCA' *.fastq > count/B6690_Fm0.txt
agrep -0 -c 'GTGCGTCCACGTACTTG' *.fastq > count/B6692_Fm0.txt
agrep -0 -c 'TGAGGAAGATCGTAGAG' *.fastq > count/B6693_Fm0.txt
agrep -0 -c 'GGGATTCTTATCTTGGC' *.fastq > count/B6694_Fm0.txt
agrep -0 -c 'TGAAGGCATACCTTGCT' *.fastq > count/B6696_Fm0.txt
agrep -0 -c 'CGGCCATTTACACAATC' *.fastq > count/B6697_Fm0.txt
agrep -0 -c 'CTGGAAAAGACGTAGGG' *.fastq > count/B6698_Fm0.txt
agrep -0 -c 'CACCTGATCACACGTAC' *.fastq > count/B6699_Fm0.txt
agrep -0 -c 'CGAAGTTGCTTGACAAG' *.fastq > count/B6700_Fm0.txt
agrep -0 -c 'CCAATATCCACCGGCTT' *.fastq > count/B6701_Fm0.txt
agrep -0 -c 'GGCTACTATGTATGTGG' *.fastq > count/B6702_Fm0.txt
agrep -0 -c 'CAGTCACTGACTTCCCG' *.fastq > count/B6703_Fm0.txt
agrep -0 -c 'TGCATGTCACGTTATTG' *.fastq > count/B6704_Fm0.txt
agrep -0 -c 'AGCCTAACTCGGTGCGA' *.fastq > count/B6705_Fm0.txt
agrep -0 -c 'CTAGAGGTATACATAGG' *.fastq > count/B6706_Fm0.txt
agrep -0 -c 'TTGATTACCCCATGAAC' *.fastq > count/B6707_Fm0.txt
agrep -0 -c 'CGGGCATAGTTTTGAGA' *.fastq > count/B6708_Fm0.txt
agrep -0 -c 'AGCAGGTACAATTCGAT' *.fastq > count/B6709_Fm0.txt
agrep -0 -c 'AAGACATTCGGTATCTG' *.fastq > count/B6710_Fm0.txt
agrep -0 -c 'GTATGGCGCTTCATCTC' *.fastq > count/B6711_Fm0.txt
agrep -0 -c 'CATTGGTTTCCAACGGC' *.fastq > count/B6712_Fm0.txt
agrep -0 -c 'TTGTATAACCTGACGCA' *.fastq > count/B6713_Fm0.txt
agrep -0 -c 'TCGTCGCCAGTATTGAC' *.fastq > count/B6714_Fm0.txt
agrep -0 -c 'ATGCTCACAAACCGTGC' *.fastq > count/B6715_Fm0.txt
agrep -0 -c 'GTAACAGCATTGCTCTA' *.fastq > count/B6717_Fm0.txt
agrep -0 -c 'GGAAGCTTGGCAAGAAG' *.fastq > count/B6718_Fm0.txt
agrep -0 -c 'CCTTTGAAAGCTGCATT' *.fastq > count/B6719_Fm0.txt
agrep -0 -c 'TACGGCCATGCGGCTTT' *.fastq > count/B6720_Fm0.txt
agrep -0 -c 'TAGAGCCGTTTGATAGG' *.fastq > count/B6721_Fm0.txt
agrep -0 -c 'CTCCGAGGATAGAACCT' *.fastq > count/B6722_Fm0.txt
agrep -0 -c 'CAGCAGGACTAACCGAG' *.fastq > count/B6723_Fm0.txt
agrep -0 -c 'GGCGATATTCGATTACC' *.fastq > count/B6724_Fm0.txt
agrep -0 -c 'CAGCGATTGGGGTCCTT' *.fastq > count/B6726_Fm0.txt
agrep -0 -c 'CGCTTTACTGTGACAAC' *.fastq > count/B6727_Fm0.txt
agrep -0 -c 'GTGTTACCTTCCACACT' *.fastq > count/B6728_Fm0.txt
agrep -0 -c 'CTCCCGCCCAAGGATAT' *.fastq > count/B6729_Fm0.txt
agrep -0 -c 'AAGTTCGCCAGTGATGC' *.fastq > count/B6730_Fm0.txt
agrep -0 -c 'TCGGATTTGAGACTGTC' *.fastq > count/B6731_Fm0.txt
agrep -0 -c 'ATGTCGTAAAGGCGCGG' *.fastq > count/B6733_Fm0.txt
agrep -0 -c 'CAACTCTGATGCCCCTT' *.fastq > count/B6734_Fm0.txt
agrep -0 -c 'ACCCCCGCAACTATACA' *.fastq > count/B6735_Fm0.txt
agrep -0 -c 'TTGCTCACCCTACTCCT' *.fastq > count/B6736_Fm0.txt
agrep -0 -c 'AGATGGCGGTAGAGTCC' *.fastq > count/B6738_Fm0.txt
agrep -0 -c 'CTCGGGCGGTATTCATG' *.fastq > count/B6739_Fm0.txt
agrep -0 -c 'GCCTTGGCTCTGATCTG' *.fastq > count/B6741_Fm0.txt
agrep -0 -c 'ATTGGCGATCAGCAGTC' *.fastq > count/B6742_Fm0.txt
agrep -0 -c 'CCATCAGAAATTTTCCG' *.fastq > count/B6743_Fm0.txt
agrep -0 -c 'ATGACGCTGCACGAGGA' *.fastq > count/B6745_Fm0.txt
agrep -0 -c 'GTAAGGAGCCACACCCT' *.fastq > count/B6746_Fm0.txt
agrep -0 -c 'GCCGCTTTGGTTCAGCT' *.fastq > count/B6747_Fm0.txt
agrep -0 -c 'GCGGACTCAACCGGTTT' *.fastq > count/B6748_Fm0.txt
agrep -0 -c 'GTTTAGTCCGGGATGAA' *.fastq > count/B6749_Fm0.txt
agrep -0 -c 'GATCGCTGTGACAGTCT' *.fastq > count/B6750_Fm0.txt
agrep -0 -c 'GTAATCTATGAGGGTAC' *.fastq > count/B6751_Fm0.txt
agrep -0 -c 'GCATTTGGCCTGAGCGA' *.fastq > count/B6752_Fm0.txt
agrep -0 -c 'CATGACCACCGATACTG' *.fastq > count/B6753_Fm0.txt
agrep -0 -c 'CAATGCGTACGTAAGGC' *.fastq > count/B6754_Fm0.txt
agrep -0 -c 'GATCGGGAGATCGTCCA' *.fastq > count/B6755_Fm0.txt
agrep -0 -c 'ATATTCGCGAATCAGGG' *.fastq > count/B6756_Fm0.txt
agrep -0 -c 'TCTAGCGAGACCAAACG' *.fastq > count/B6758_Fm0.txt
agrep -0 -c 'ATCGCCTAGCAGTTATC' *.fastq > count/B6759_Fm0.txt
agrep -0 -c 'GATGTGATGATGCTCCC' *.fastq > count/B6760_Fm0.txt
agrep -0 -c 'CAGTCATGATCTCGACC' *.fastq > count/B6761_Fm0.txt
agrep -0 -c 'CAACGGCAACACTAGGT' *.fastq > count/B6762_Fm0.txt
agrep -0 -c 'TTCACGACCATTGTAGG' *.fastq > count/B6763_Fm0.txt
agrep -0 -c 'TTCTAGTCCAAAACCGC' *.fastq > count/B6764_Fm0.txt
agrep -0 -c 'AGTCGCCTGCTTACCCT' *.fastq > count/B6765_Fm0.txt
agrep -0 -c 'AGGGTACCATCACGTTA' *.fastq > count/B6766_Fm0.txt
agrep -0 -c 'GATGTGGCCGTTAGGCT' *.fastq > count/B6767_Fm0.txt
agrep -0 -c 'CGCATCTAATGGCACAT' *.fastq > count/B6768_Fm0.txt
agrep -0 -c 'CGTATACGACTCTGTCA' *.fastq > count/B6769_Fm0.txt
agrep -0 -c 'TGGCGTTGTTGCCCAGA' *.fastq > count/B6770_Fm0.txt
agrep -0 -c 'AAGGATGGCCTATCGGC' *.fastq > count/B6771_Fm0.txt
agrep -0 -c 'GGCACAACTGACAGGTG' *.fastq > count/B6772_Fm0.txt
agrep -0 -c 'TTAGAATACATCGCCCG' *.fastq > count/B6773_Fm0.txt
agrep -0 -c 'TCACAGTGGAAACGGGG' *.fastq > count/B6775_Fm0.txt
agrep -0 -c 'ACCTACGCTCAAGAGCG' *.fastq > count/B6776_Fm0.txt
agrep -0 -c 'GTATCCGGTCGTTTATA' *.fastq > count/B6778_Fm0.txt
agrep -0 -c 'GGAAGTGTGTTCGTCAT' *.fastq > count/B6779_Fm0.txt
agrep -0 -c 'CTGTAGACTACATCTGA' *.fastq > count/B6780_Fm0.txt
agrep -0 -c 'TGCCCCGACATTGATGC' *.fastq > count/B6781_Fm0.txt
agrep -0 -c 'GGACATAATCACGACAT' *.fastq > count/B6782_Fm0.txt
agrep -0 -c 'CAATTGTGGGAGTAAAG' *.fastq > count/B6784_Fm0.txt
agrep -0 -c 'GCGTTTGTCCCCACCTT' *.fastq > count/B6787_Fm0.txt
agrep -0 -c 'CAAGAGCGAAATGTTCT' *.fastq > count/B6788_Fm0.txt
agrep -0 -c 'GCACGCTAAAAACGGGC' *.fastq > count/B6789_Fm0.txt
agrep -0 -c 'ACGCTACGATGAGGTAG' *.fastq > count/B6790_Fm0.txt
agrep -0 -c 'TTCTAGCAATGGGTCCG' *.fastq > count/B6791_Fm0.txt
agrep -0 -c 'CCATCGCCCAAGTCCAA' *.fastq > count/B6792_Fm0.txt
agrep -0 -c 'TACACGTGATGTCGAGA' *.fastq > count/B6793_Fm0.txt
agrep -0 -c 'CTAGGATTGGAGCCGTT' *.fastq > count/B6794_Fm0.txt
agrep -0 -c 'CAAACTCGCTTACAGCG' *.fastq > count/B6795_Fm0.txt
agrep -0 -c 'GGTCGATAAAGTTACTC' *.fastq > count/B6796_Fm0.txt
agrep -0 -c 'TTGACGTTGCGGGTAGT' *.fastq > count/B6800_Fm0.txt
agrep -0 -c 'TCGAAGGTCGTTCGTAC' *.fastq > count/B6801_Fm0.txt
agrep -0 -c 'CGCCTTGTTATGTCATA' *.fastq > count/B6802_Fm0.txt
agrep -0 -c 'TAAGAGTTGGTCTGACC' *.fastq > count/B6803_Fm0.txt
agrep -0 -c 'CCTTATTCGGTGTCGCC' *.fastq > count/B6804_Fm0.txt
agrep -0 -c 'TGCAGTTCCGATGTGAC' *.fastq > count/B6806_Fm0.txt
agrep -0 -c 'TATACACCGTGTGCAGG' *.fastq > count/B6807_Fm0.txt
agrep -0 -c 'CTGACATTGAGACGGCT' *.fastq > count/B6810_Fm0.txt
agrep -0 -c 'TCTGTTTGGCGCTTAGC' *.fastq > count/B6811_Fm0.txt
agrep -0 -c 'ACAAGCGCAGTATTTCT' *.fastq > count/B6812_Fm0.txt
agrep -0 -c 'CGGAATACTTCTGCGCT' *.fastq > count/B6813_Fm0.txt
agrep -0 -c 'GTATCGAATGCACTGGA' *.fastq > count/B6814_Fm0.txt
agrep -0 -c 'AGTGGGTACAAACGAAT' *.fastq > count/B6815_Fm0.txt
agrep -0 -c 'TTGCTCACAGAAAGCGG' *.fastq > count/B6816_Fm0.txt
agrep -0 -c 'CGTAACAGTCCTGGATT' *.fastq > count/B6817_Fm0.txt
agrep -0 -c 'GCCAACAGGCGATCTTG' *.fastq > count/B6818_Fm0.txt
agrep -0 -c 'GGCTCATGTTTGTCTTC' *.fastq > count/B6819_Fm0.txt
agrep -0 -c 'ACGCTTTGTAGCAAGCT' *.fastq > count/B6821_Fm0.txt
agrep -0 -c 'AGGTCGGACTAACGCAA' *.fastq > count/B6822_Fm0.txt
agrep -0 -c 'TGAAAAAGGGGAAACGT' *.fastq > count/B6824_Fm0.txt
agrep -0 -c 'AGTCCGAACTGACACCT' *.fastq > count/B6826_Fm0.txt
agrep -0 -c 'CAGGCCGTAACACATCC' *.fastq > count/B6828_Fm0.txt
agrep -0 -c 'GTCCATAATTGGCGCCA' *.fastq > count/B6830_Fm0.txt
agrep -0 -c 'AGCCGCTATTCTAACGG' *.fastq > count/B6831_Fm0.txt
agrep -0 -c 'ACAGCTCGGAGTCGAGT' *.fastq > count/B6832_Fm0.txt
agrep -0 -c 'TACGAAATGACCTGTGG' *.fastq > count/B6833_Fm0.txt
agrep -0 -c 'CACGCCCATTCAGGTCT' *.fastq > count/B6835_Fm0.txt
agrep -0 -c 'CAATCCTACTACGCCAC' *.fastq > count/B6837_Fm0.txt
agrep -0 -c 'CCGCAGTACGAGCATGA' *.fastq > count/B6838_Fm0.txt
agrep -0 -c 'GGATAGTTTAGGGCGCC' *.fastq > count/B6839_Fm0.txt
agrep -0 -c 'CTATGTGCGAGACTGGG' *.fastq > count/B6841_Fm0.txt
agrep -0 -c 'AGTGTTTTCACGGACCG' *.fastq > count/B6842_Fm0.txt
agrep -0 -c 'CGTTGCTCGAAACATCC' *.fastq > count/B6843_Fm0.txt
agrep -0 -c 'AGATCGAAAATTGCCGT' *.fastq > count/B6844_Fm0.txt
agrep -0 -c 'TGATGCAAAACGCTGTA' *.fastq > count/B6845_Fm0.txt
agrep -0 -c 'GCAATAAGCGATCTGCA' *.fastq > count/B6846_Fm0.txt
agrep -0 -c 'CGCGTTCTGACCTCGAA' *.fastq > count/B6847_Fm0.txt
agrep -0 -c 'ACAGGAGTAGTCCGAAG' *.fastq > count/B6848_Fm0.txt
agrep -0 -c 'AACAATGGCTCGAATTC' *.fastq > count/B6849_Fm0.txt
agrep -0 -c 'GTCCCCATAACGCGTCA' *.fastq > count/B6851_Fm0.txt
agrep -0 -c 'GTGGTGGAACAATGTAA' *.fastq > count/B6853_Fm0.txt
agrep -0 -c 'TACGCTTAGATCCATCA' *.fastq > count/B6854_Fm0.txt
agrep -0 -c 'TACCTTAGGCAATAGCC' *.fastq > count/B6855_Fm0.txt
agrep -0 -c 'AGGGAATAAAATCCGTC' *.fastq > count/B6856_Fm0.txt
agrep -0 -c 'CACGTGCACGCAGACTA' *.fastq > count/B6857_Fm0.txt
agrep -0 -c 'GACGTTGAACTTGCGAG' *.fastq > count/B6858_Fm0.txt
agrep -0 -c 'CTTGACCAACAGCACGT' *.fastq > count/B6859_Fm0.txt
agrep -0 -c 'ACAGCCATATCGACGTT' *.fastq > count/B6861_Fm0.txt
agrep -0 -c 'CTGGATTAGTCCCTGAT' *.fastq > count/B6862_Fm0.txt
agrep -0 -c 'AGTATAATCAGCCCGGA' *.fastq > count/B6863_Fm0.txt
agrep -0 -c 'ACACCATGGAGTCGAGC' *.fastq > count/B6865_Fm0.txt
agrep -0 -c 'AACATTAATTACCCCGC' *.fastq > count/B6866_Fm0.txt
agrep -0 -c 'TATACTGACTGGCGTTG' *.fastq > count/B6867_Fm0.txt
agrep -0 -c 'CCATCTTTCGCAGCTAC' *.fastq > count/B6868_Fm0.txt
agrep -0 -c 'CAACATGGATTGTGACG' *.fastq > count/B6869_Fm0.txt
agrep -0 -c 'CGCTTGGACATACTATG' *.fastq > count/B6871_Fm0.txt
agrep -0 -c 'ATCACTGAATGGGCCAA' *.fastq > count/B6872_Fm0.txt
agrep -0 -c 'GGGATTAGAGGGCCAAA' *.fastq > count/B6873_Fm0.txt
agrep -0 -c 'ATGAAAGAGGGTTTGCA' *.fastq > count/B6874_Fm0.txt
agrep -0 -c 'TCTCGCACATATAGAGG' *.fastq > count/B6875_Fm0.txt
agrep -0 -c 'TTCGAGGAGAACACATG' *.fastq > count/B6876_Fm0.txt
agrep -0 -c 'TGGCTTTTGACCTCCAA' *.fastq > count/B6878_Fm0.txt
agrep -0 -c 'ATGTTACTACCCGGACG' *.fastq > count/B6880_Fm0.txt
agrep -0 -c 'ATTCCACTTGTGCAGTA' *.fastq > count/B6881_Fm0.txt
agrep -0 -c 'GGTCCGCTACGCTTTAA' *.fastq > count/B6883_Fm0.txt
agrep -0 -c 'CTCCGATCGCGAATGTG' *.fastq > count/B6884_Fm0.txt
agrep -0 -c 'GGACTGGCGATCTAAGT' *.fastq > count/B6886_Fm0.txt
agrep -0 -c 'CACTTGATCCGAGGGAG' *.fastq > count/B6887_Fm0.txt
agrep -0 -c 'CTTGAGACTAATGAAGG' *.fastq > count/B6888_Fm0.txt
agrep -0 -c 'TACGACTTCAGGTTTGT' *.fastq > count/B6889_Fm0.txt
agrep -0 -c 'TCGTGTTATGCCCATCG' *.fastq > count/B6890_Fm0.txt
agrep -0 -c 'AGAGTTCTCACATGCCA' *.fastq > count/B6891_Fm0.txt
agrep -0 -c 'TAAGGTAGACGAAAGAG' *.fastq > count/B6893_Fm0.txt
agrep -0 -c 'TCACGGATTCTATGTGG' *.fastq > count/B6894_Fm0.txt
agrep -0 -c 'CGCAGAGACAATTGTAA' *.fastq > count/B6896_Fm0.txt
agrep -0 -c 'CGTCTGACGATATACTA' *.fastq > count/B6898_Fm0.txt
agrep -0 -c 'GTTATAGTACCACTAGC' *.fastq > count/B6899_Fm0.txt
agrep -0 -c 'GACACTCCTGGCGAAGT' *.fastq > count/B6900_Fm0.txt
agrep -0 -c 'AGCGCGTTGAACCTGGT' *.fastq > count/B6902_Fm0.txt
agrep -0 -c 'TTGCACATTCGCGACTG' *.fastq > count/B6903_Fm0.txt
agrep -0 -c 'TCTACGCTCGGTAATGC' *.fastq > count/B6905_Fm0.txt
agrep -0 -c 'TCGAACGCTCGACCTGA' *.fastq > count/B6907_Fm0.txt
agrep -0 -c 'TTCTGTTAAACCTTCCG' *.fastq > count/B6908_Fm0.txt
agrep -0 -c 'GAGACTTCGTAAACCTG' *.fastq > count/B6909_Fm0.txt
agrep -0 -c 'GTCCGTCGCAGAATGCA' *.fastq > count/B6910_Fm0.txt
agrep -0 -c 'ATTCCTTTGGTCTAGAG' *.fastq > count/B6911_Fm0.txt
agrep -0 -c 'CAACTATGGGCTTGTTC' *.fastq > count/B6912_Fm0.txt
agrep -0 -c 'ATGTACCGGGTCGTTTC' *.fastq > count/B6913_Fm0.txt
agrep -0 -c 'TAAAACGGACATCCGCG' *.fastq > count/B6914_Fm0.txt
agrep -0 -c 'ATGGATTGCGGACACAC' *.fastq > count/B6915_Fm0.txt
agrep -0 -c 'GGTCCGTGGATATTGTG' *.fastq > count/B6916_Fm0.txt
agrep -0 -c 'AGAACGGTGGGTTGCTT' *.fastq > count/B6917_Fm0.txt
agrep -0 -c 'GGTTGTGGGCCATACTC' *.fastq > count/B6918_Fm0.txt
agrep -0 -c 'GCGTGGCAAATTTGCAG' *.fastq > count/B6919_Fm0.txt
agrep -0 -c 'AAGCTGCTAGAGTTCGA' *.fastq > count/B6920_Fm0.txt
agrep -0 -c 'AGAATCGGCCGATTTTA' *.fastq > count/B6921_Fm0.txt
agrep -0 -c 'ATCCTGGAGCTGGTATA' *.fastq > count/B6922_Fm0.txt
agrep -0 -c 'CGATTTCCTAAGTCCGT' *.fastq > count/B6923_Fm0.txt
agrep -0 -c 'TAACCCCTTTCTGTACG' *.fastq > count/B6924_Fm0.txt
agrep -0 -c 'CCTTAAGACCCATTGCA' *.fastq > count/B6925_Fm0.txt
agrep -0 -c 'CACGAGTACCAAGTGGA' *.fastq > count/B6926_Fm0.txt
agrep -0 -c 'GGTGCTTCAGATCTAGA' *.fastq > count/B6927_Fm0.txt
agrep -0 -c 'CATCTGCCACAATCGCT' *.fastq > count/B6928_Fm0.txt
agrep -0 -c 'CACCAATCGCATGAATC' *.fastq > count/B6929_Fm0.txt
agrep -0 -c 'GGCCAATCTCACAACGT' *.fastq > count/B6930_Fm0.txt
agrep -0 -c 'GTCAATCGGACAACAAT' *.fastq > count/B6931_Fm0.txt
agrep -0 -c 'ATATACGGCTCAGGATT' *.fastq > count/B6932_Fm0.txt
agrep -0 -c 'TGCGATACTTAGGCTTG' *.fastq > count/B6933_Fm0.txt
agrep -0 -c 'ATGAACTGACCCAAGTT' *.fastq > count/B6934_Fm0.txt
agrep -0 -c 'CTCCCATTGTGCAATCG' *.fastq > count/B6938_Fm0.txt
agrep -0 -c 'GGAGATGAGTTGACTAC' *.fastq > count/B6939_Fm0.txt
agrep -0 -c 'ATGCGAGGCCTATGTCT' *.fastq > count/B6942_Fm0.txt
agrep -0 -c 'CAGATGCTTGTCACGGC' *.fastq > count/B6943_Fm0.txt
agrep -0 -c 'GCCTCGTCTCACTGTAC' *.fastq > count/B6944_Fm0.txt
agrep -0 -c 'GTAGTCTGGTCACGAAT' *.fastq > count/B6946_Fm0.txt
agrep -0 -c 'TCACTCAACGCTCTGCA' *.fastq > count/B6947_Fm0.txt
agrep -0 -c 'CAGTGGCTCGAACATGA' *.fastq > count/B6948_Fm0.txt
agrep -0 -c 'GTACCGACGAACTGAAC' *.fastq > count/B6950_Fm0.txt
agrep -0 -c 'GTGGCTCCGAAATGTTT' *.fastq > count/B6951_Fm0.txt
agrep -0 -c 'CTCACGCTATGAGCGAA' *.fastq > count/B6952_Fm0.txt
agrep -0 -c 'GGAAGTTCCACTCACCC' *.fastq > count/B6953_Fm0.txt
agrep -0 -c 'ATACGGGGGTAATGGTA' *.fastq > count/B6954_Fm0.txt
agrep -0 -c 'ATAGAACCTGCTCTTCG' *.fastq > count/B6955_Fm0.txt
agrep -0 -c 'GAGATTCGAAGAAGTTC' *.fastq > count/B6957_Fm0.txt
agrep -0 -c 'TAATGCGTCTAGGAGTC' *.fastq > count/B6958_Fm0.txt
agrep -0 -c 'CTCGGAGTATCTAAACA' *.fastq > count/B6959_Fm0.txt
agrep -0 -c 'CTCCCTTACCCAGTAGG' *.fastq > count/B6960_Fm0.txt
agrep -0 -c 'TGCTGCTTTTTAGGTTC' *.fastq > count/B6961_Fm0.txt
agrep -0 -c 'ATGTAGACGATGCTGTA' *.fastq > count/B6962_Fm0.txt
agrep -0 -c 'TAGGTTGCACAACCGGC' *.fastq > count/B6963_Fm0.txt
agrep -0 -c 'TGTATGTAACAGTGCCG' *.fastq > count/B6964_Fm0.txt
agrep -0 -c 'TGTGAACCCTGGAGCAC' *.fastq > count/B6967_Fm0.txt
agrep -0 -c 'AATCCCGCACTTTGAAC' *.fastq > count/B6968_Fm0.txt
agrep -0 -c 'CTGCCACCAGCCAACTA' *.fastq > count/B6969_Fm0.txt
agrep -0 -c 'CAATCACGCCTGGATGT' *.fastq > count/B6970_Fm0.txt
agrep -0 -c 'CTGGGGTGATCCTAGGA' *.fastq > count/B6971_Fm0.txt
agrep -0 -c 'GCAGGTCTTTTCGAGAC' *.fastq > count/B6972_Fm0.txt
agrep -0 -c 'CTGCTTACCGATGTAAC' *.fastq > count/B6974_Fm0.txt
agrep -0 -c 'GGTTCCATAGAGCCACG' *.fastq > count/B6975_Fm0.txt
agrep -0 -c 'TACACTTCGACCTTCGA' *.fastq > count/B6976_Fm0.txt
agrep -0 -c 'GCCGTGCAGCAGATTTG' *.fastq > count/B6977_Fm0.txt
agrep -0 -c 'TCGCACGTCAGCAAACC' *.fastq > count/B6979_Fm0.txt
agrep -0 -c 'GGCATGGCAGGCTTAAT' *.fastq > count/B6980_Fm0.txt
agrep -0 -c 'GATACAGCGTCGAGTAC' *.fastq > count/B6981_Fm0.txt
agrep -0 -c 'GCGAACTCTAGGATGAG' *.fastq > count/B6982_Fm0.txt
agrep -0 -c 'GATTACGGCACCGTGTC' *.fastq > count/B6983_Fm0.txt
agrep -0 -c 'GATCTTGTTACAGTCGA' *.fastq > count/B6984_Fm0.txt
agrep -0 -c 'AGTCGAAGTGGCTTTCT' *.fastq > count/B6985_Fm0.txt
agrep -0 -c 'GATATCAGCATACGGTC' *.fastq > count/B6986_Fm0.txt
agrep -0 -c 'AGTCCTGGTAGAAGTGG' *.fastq > count/B6989_Fm0.txt
agrep -0 -c 'CGTTTAACCTCTGGACT' *.fastq > count/B6991_Fm0.txt
agrep -0 -c 'CGACTTCTTCTACTAGA' *.fastq > count/B6992_Fm0.txt
agrep -0 -c 'CAACGTACACACGAATA' *.fastq > count/B6993_Fm0.txt
agrep -0 -c 'TTAAGAGGTTACCCGGT' *.fastq > count/B6994_Fm0.txt
agrep -0 -c 'TGCCCTGGTTGTGATAC' *.fastq > count/B6996_Fm0.txt
agrep -0 -c 'ATTAGGTCAGCGACTTC' *.fastq > count/B6997_Fm0.txt
agrep -0 -c 'CCGAGACCCCTGAACTT' *.fastq > count/B7000_Fm0.txt
agrep -0 -c 'GCACCCCCATCAAACTT' *.fastq > count/B7001_Fm0.txt
agrep -0 -c 'AGGTGCCGCGTTTGCAA' *.fastq > count/B7002_Fm0.txt
agrep -0 -c 'TATCGTATACGTAGTCG' *.fastq > count/B7003_Fm0.txt
agrep -0 -c 'TGACACAGAAATCCTGG' *.fastq > count/B7004_Fm0.txt
agrep -0 -c 'ATATCGCTAGCTCTAGC' *.fastq > count/B7007_Fm0.txt
agrep -0 -c 'TGCCGATGAGAGCAACA' *.fastq > count/B7011_Fm0.txt
agrep -0 -c 'CAATGCTCTAGGCCTAT' *.fastq > count/B7012_Fm0.txt
agrep -0 -c 'GACCAATGGCGATTTGC' *.fastq > count/B7013_Fm0.txt
agrep -0 -c 'CCGCCAGCAATGTTCCA' *.fastq > count/B7014_Fm0.txt
agrep -0 -c 'CTATCCTAGAGTCCTAA' *.fastq > count/B7015_Fm0.txt
agrep -0 -c 'CAGTATCAGGGGAATCG' *.fastq > count/B7016_Fm0.txt
agrep -0 -c 'TTAGCCTCCGAGACCAG' *.fastq > count/B7017_Fm0.txt
agrep -0 -c 'ATCGGCGGAACATTCCG' *.fastq > count/B7018_Fm0.txt
agrep -0 -c 'GGCTACCGGAGTATACA' *.fastq > count/B7020_Fm0.txt
agrep -0 -c 'GCCCAATCAGTGAAGTG' *.fastq > count/B7021_Fm0.txt
agrep -0 -c 'GGGTCGAGCACATCCAA' *.fastq > count/B7022_Fm0.txt
agrep -0 -c 'AGTCAAGCCTCCTTCAA' *.fastq > count/B7023_Fm0.txt
agrep -0 -c 'GGTCGTTTTAAGCCCAC' *.fastq > count/B7024_Fm0.txt
agrep -0 -c 'ACGTAGAAGGCTTGTGC' *.fastq > count/B7026_Fm0.txt
agrep -0 -c 'ACCATGCTTGTACTTTG' *.fastq > count/B7027_Fm0.txt
agrep -0 -c 'GGGCTTTAATCGATCTC' *.fastq > count/B7029_Fm0.txt
agrep -0 -c 'ATCCCGACGACAACGTG' *.fastq > count/B7031_Fm0.txt
agrep -0 -c 'CGTGAGACCAAATGCCG' *.fastq > count/B7032_Fm0.txt
agrep -0 -c 'AGTAGTCTGAAGCGTCC' *.fastq > count/B7034_Fm0.txt
agrep -0 -c 'CTAATATCGGCGCAGGC' *.fastq > count/B7035_Fm0.txt
agrep -0 -c 'AACGAGCCCCAGATTCT' *.fastq > count/B7036_Fm0.txt
agrep -0 -c 'GCGAAGTTATCGTCAAT' *.fastq > count/B7037_Fm0.txt
agrep -0 -c 'GGAAGCATGATAACCTT' *.fastq > count/B7038_Fm0.txt
agrep -0 -c 'GGTAATCTCAGGTCGTG' *.fastq > count/B7039_Fm0.txt
agrep -0 -c 'GGAGGAGATTGTGTCGG' *.fastq > count/B7040_Fm0.txt
agrep -0 -c 'TCAGAGAAGCATGCTGA' *.fastq > count/B7041_Fm0.txt
agrep -0 -c 'CCTGCGAATTAGGAGGA' *.fastq > count/B7042_Fm0.txt
agrep -0 -c 'CGAGTGAAGAGAAAAGT' *.fastq > count/B7043_Fm0.txt
agrep -0 -c 'TTCTACCAGACGGAAGA' *.fastq > count/B7044_Fm0.txt
agrep -0 -c 'TTGGACCGGTGCTACTC' *.fastq > count/B7046_Fm0.txt
agrep -0 -c 'GAGCGACCATCACCAGT' *.fastq > count/B7048_Fm0.txt
agrep -0 -c 'CGCAACAAGACTGGGAG' *.fastq > count/B7049_Fm0.txt
agrep -0 -c 'ATTGGAGAAACGACTGG' *.fastq > count/B7050_Fm0.txt
agrep -0 -c 'GTTCAAGCGTTCGATAT' *.fastq > count/B7051_Fm0.txt
agrep -0 -c 'GGTCACACTATACACGC' *.fastq > count/B7052_Fm0.txt
agrep -0 -c 'AAGGGATAACGGCCGAT' *.fastq > count/B7053_Fm0.txt
agrep -0 -c 'AATGATGACGTGTCCCC' *.fastq > count/B7054_Fm0.txt
agrep -0 -c 'TGTCTCACCTGACAACC' *.fastq > count/B7055_Fm0.txt
agrep -0 -c 'AGGGCTTGACTAAGTGT' *.fastq > count/B7056_Fm0.txt
agrep -0 -c 'CCGTCTCGGATGTAATA' *.fastq > count/B7057_Fm0.txt
agrep -0 -c 'ATAGACAGGACACTCCC' *.fastq > count/B7059_Fm0.txt
agrep -0 -c 'CATCATGCTAAACCGAC' *.fastq > count/B7061_Fm0.txt
agrep -0 -c 'ATAGAGCCCAAGTTAGT' *.fastq > count/B7062_Fm0.txt
agrep -0 -c 'CAACAGCTCTCATCGTC' *.fastq > count/B7063_Fm0.txt
agrep -0 -c 'GGTACGAGTAAGGATGC' *.fastq > count/B7064_Fm0.txt
agrep -0 -c 'GAGAGGGGGATACACGT' *.fastq > count/B7065_Fm0.txt
agrep -0 -c 'ACCTAGGGCTTGAGAAT' *.fastq > count/B7066_Fm0.txt
agrep -0 -c 'GCACTTCGGTGTTAATC' *.fastq > count/B7067_Fm0.txt
agrep -0 -c 'GTCGGGTAGAATTCACA' *.fastq > count/B7068_Fm0.txt
agrep -0 -c 'GGCACTACCTTGACCTC' *.fastq > count/B7070_Fm0.txt
agrep -0 -c 'CGCGACATTTTCGATAG' *.fastq > count/B7071_Fm0.txt
agrep -0 -c 'AGTGAGTATTGACAGAC' *.fastq > count/B7072_Fm0.txt
agrep -0 -c 'GGCACTATTACCTCTTT' *.fastq > count/B7075_Fm0.txt
agrep -0 -c 'CACGAGAACGCGAGTTT' *.fastq > count/B7076_Fm0.txt
agrep -0 -c 'ACTTAGGGGGCATGCTA' *.fastq > count/B7077_Fm0.txt
agrep -0 -c 'CTTACAAACCGCGTGTC' *.fastq > count/B7078_Fm0.txt
agrep -0 -c 'AATGGAAGGTACGATAG' *.fastq > count/B7079_Fm0.txt
agrep -0 -c 'AAGGTAATCAGTGCGGG' *.fastq > count/B7080_Fm0.txt
agrep -0 -c 'AATTCAGCGGATTGCTA' *.fastq > count/B7081_Fm0.txt
agrep -0 -c 'TTGTATGGCCCTTCGTG' *.fastq > count/B7082_Fm0.txt
agrep -0 -c 'CGACGAGCAACTCAAAA' *.fastq > count/B7083_Fm0.txt
agrep -0 -c 'AAGAGCCATAGGTAGCC' *.fastq > count/B7085_Fm0.txt
agrep -0 -c 'CTTGCCTAATCTGAGGG' *.fastq > count/B7086_Fm0.txt
agrep -0 -c 'GGTTATAGTCCGTAGAC' *.fastq > count/B7087_Fm0.txt
agrep -0 -c 'CGGCAGCTATGGACTTA' *.fastq > count/B7088_Fm0.txt
agrep -0 -c 'ACGCTAGAGTGTTTGAG' *.fastq > count/B7089_Fm0.txt
agrep -0 -c 'GCCGCCGGTTGCTATTA' *.fastq > count/B7090_Fm0.txt
agrep -0 -c 'TAAGAGACTGGCACAAC' *.fastq > count/B7091_Fm0.txt
agrep -0 -c 'CACACTCGAACCACTTG' *.fastq > count/B7092_Fm0.txt
agrep -0 -c 'ACGCCTGCCCGATATCA' *.fastq > count/B7093_Fm0.txt
agrep -0 -c 'GTGTCAACCACTGGGAA' *.fastq > count/B7094_Fm0.txt
agrep -0 -c 'GTAATAGTACTAGTCGC' *.fastq > count/B7095_Fm0.txt
agrep -0 -c 'TAGCACCTTACCAGTGA' *.fastq > count/B7097_Fm0.txt
agrep -0 -c 'TTACTACATTGTGCCGA' *.fastq > count/B7098_Fm0.txt
agrep -0 -c 'TATGTCAGGTGCGGTTG' *.fastq > count/B7099_Fm0.txt
agrep -0 -c 'TCGTACTCCAAGACCTA' *.fastq > count/B7100_Fm0.txt
agrep -0 -c 'ACAGTCGGCTACACTCG' *.fastq > count/B7101_Fm0.txt
agrep -0 -c 'TTGCATGGTGAAAGTCT' *.fastq > count/B7103_Fm0.txt
agrep -0 -c 'CCACACGTTACCCTAGT' *.fastq > count/B7104_Fm0.txt
agrep -0 -c 'CTAGTTCTTTCACGCCA' *.fastq > count/B7105_Fm0.txt
agrep -0 -c 'GCAGGTATCGGGTCACT' *.fastq > count/B7106_Fm0.txt
agrep -0 -c 'AGTGGCGAGCAGTACAG' *.fastq > count/B7107_Fm0.txt
agrep -0 -c 'ACCATTGTCGGTATCGT' *.fastq > count/B7108_Fm0.txt
agrep -0 -c 'AGAGCGGCGCACTTATG' *.fastq > count/B7109_Fm0.txt
agrep -0 -c 'GGGAGAAAGGTGTAACC' *.fastq > count/B7110_Fm0.txt
agrep -0 -c 'ACGAAGCTGGAACTCCT' *.fastq > count/B7111_Fm0.txt
agrep -0 -c 'TTAATCAGCATCTGAGG' *.fastq > count/B7112_Fm0.txt
agrep -0 -c 'TGACTCATTAAGGTGAG' *.fastq > count/B7113_Fm0.txt
agrep -0 -c 'CATAGATTGGAACGCCA' *.fastq > count/B7114_Fm0.txt
agrep -0 -c 'CAACGTGTATCACTGCT' *.fastq > count/B7115_Fm0.txt
agrep -0 -c 'CGAGGCTAATAACTAGG' *.fastq > count/B7116_Fm0.txt
agrep -0 -c 'TCGACAAAGAGGGGAGT' *.fastq > count/B7117_Fm0.txt
agrep -0 -c 'GTGATACTATGGCGTTC' *.fastq > count/B7118_Fm0.txt
agrep -0 -c 'CCGAAGACTGACTCCTA' *.fastq > count/B7119_Fm0.txt
agrep -0 -c 'TGCGATAGTTCAGAAGG' *.fastq > count/B7120_Fm0.txt
agrep -0 -c 'GGCTCGTGGGATCAAAA' *.fastq > count/B7121_Fm0.txt
agrep -0 -c 'ACCGATCTCCAATTCCA' *.fastq > count/B7122_Fm0.txt
agrep -0 -c 'GGGTAAGATATGGGTTC' *.fastq > count/B7123_Fm0.txt
agrep -0 -c 'CACCTTTGTAACTACCC' *.fastq > count/B7124_Fm0.txt
agrep -0 -c 'CATAGTCTATAGACTCG' *.fastq > count/B7125_Fm0.txt
agrep -0 -c 'CTCTCAGCGTACCGATG' *.fastq > count/B7126_Fm0.txt
agrep -0 -c 'ACGCTTGGCCGTACTAG' *.fastq > count/B7127_Fm0.txt
agrep -0 -c 'TTACTCCTTGTGCGATC' *.fastq > count/B7128_Fm0.txt
agrep -0 -c 'GTAGAGGTCTATTCAGA' *.fastq > count/B7131_Fm0.txt
agrep -0 -c 'ACCTGGGCCATTTCATT' *.fastq > count/B7132_Fm0.txt
agrep -0 -c 'AGCACAAGTGGACAACC' *.fastq > count/B7133_Fm0.txt
agrep -0 -c 'ATGTGGGCTCTTAACCT' *.fastq > count/B7134_Fm0.txt
agrep -0 -c 'CCATCTCACCATGATCC' *.fastq > count/B7135_Fm0.txt
agrep -0 -c 'ATTATCGAGACCCACCA' *.fastq > count/B7136_Fm0.txt
agrep -0 -c 'ATTCCTTGACATGCTGA' *.fastq > count/B7137_Fm0.txt
agrep -0 -c 'CCTCCTAATGCAATCAC' *.fastq > count/B7138_Fm0.txt
agrep -0 -c 'TGATCCACACTTGCGCC' *.fastq > count/B7140_Fm0.txt
agrep -0 -c 'GGCTGGCGTATATTCCC' *.fastq > count/B7141_Fm0.txt
agrep -0 -c 'GGTACACGTACTGAAGT' *.fastq > count/B7142_Fm0.txt
agrep -0 -c 'CTGCCAACTACATACTG' *.fastq > count/B7143_Fm0.txt
agrep -0 -c 'CGCGGTAACGTATGGTG' *.fastq > count/B7144_Fm0.txt
agrep -0 -c 'GTACACTTCGTTCGTGT' *.fastq > count/B7145_Fm0.txt
agrep -0 -c 'GCGACCATGTACGAACG' *.fastq > count/B7146_Fm0.txt
agrep -0 -c 'CAATGGAATGTGGCCGG' *.fastq > count/B7148_Fm0.txt
agrep -0 -c 'CTCCGTTAAGACTTGAC' *.fastq > count/B7149_Fm0.txt
agrep -0 -c 'GTCATCACATCAGGGGA' *.fastq > count/B7150_Fm0.txt
agrep -0 -c 'AGGGTGAAAGCTTACAC' *.fastq > count/B7151_Fm0.txt
agrep -0 -c 'AGAATCGCTGATGCCTA' *.fastq > count/B7153_Fm0.txt
agrep -0 -c 'TAACGCTACTTTTTCGC' *.fastq > count/B7154_Fm0.txt
agrep -0 -c 'TACAGGACCGTAACCCT' *.fastq > count/B7155_Fm0.txt
agrep -0 -c 'CGCTGGTAATCCTAACT' *.fastq > count/B7156_Fm0.txt
agrep -0 -c 'CAGCGGTGGACAGAATT' *.fastq > count/B7158_Fm0.txt
agrep -0 -c 'AGTGTTCCGTAGTCACC' *.fastq > count/B7159_Fm0.txt
agrep -0 -c 'ACTACATACCGTACCAC' *.fastq > count/B7160_Fm0.txt
agrep -0 -c 'CCACACCGTCCAGAAGT' *.fastq > count/B7161_Fm0.txt
agrep -0 -c 'TATCACCGTTTATGGAG' *.fastq > count/B7162_Fm0.txt
agrep -0 -c 'TCTACTCTCAGAAGACT' *.fastq > count/B7163_Fm0.txt
agrep -0 -c 'CTCTCACAGGGTATAGT' *.fastq > count/B7164_Fm0.txt
agrep -0 -c 'ATTCAACCCTTCCTACA' *.fastq > count/B7165_Fm0.txt
agrep -0 -c 'GCGCCCAGGGTATACTT' *.fastq > count/B7167_Fm0.txt
agrep -0 -c 'GGACTCAACGCTAGACT' *.fastq > count/B7169_Fm0.txt
agrep -0 -c 'TCTTTTAGGTCAACGAG' *.fastq > count/B7170_Fm0.txt
agrep -0 -c 'TCGAGCGTCACCTAATA' *.fastq > count/B7171_Fm0.txt
agrep -0 -c 'TTAGAAAACTTCCGAGG' *.fastq > count/B7173_Fm0.txt
agrep -0 -c 'ATGCCGTCGCGTTGAGA' *.fastq > count/B7175_Fm0.txt
agrep -0 -c 'TTGTTCAGGGCTGGAGG' *.fastq > count/B7176_Fm0.txt
agrep -0 -c 'ATATGTCTACGCTCGGT' *.fastq > count/B7177_Fm0.txt
agrep -0 -c 'CGGGAAGGTTACTCTTC' *.fastq > count/B7178_Fm0.txt
agrep -0 -c 'TGTCGCGCATATCCAGG' *.fastq > count/B7180_Fm0.txt
agrep -0 -c 'ATCGGGTCCTTTCTAGC' *.fastq > count/B7181_Fm0.txt
agrep -0 -c 'AATAAGTGACGCAGTCA' *.fastq > count/B7183_Fm0.txt
agrep -0 -c 'GTGTAAAGTCCTGTTCG' *.fastq > count/B7185_Fm0.txt
agrep -0 -c 'GCATGACGTACCCTAAA' *.fastq > count/B7186_Fm0.txt
agrep -0 -c 'CAACACAAGGCAAGCGG' *.fastq > count/B7187_Fm0.txt
agrep -0 -c 'GTTCCCGATGTCAAGCA' *.fastq > count/B7188_Fm0.txt
agrep -0 -c 'GCCAGCTCGGAAATCCT' *.fastq > count/B7189_Fm0.txt
agrep -0 -c 'ATACGTGCGAATTGTTC' *.fastq > count/B7190_Fm0.txt
agrep -0 -c 'CAGAGGACCCTCTGACA' *.fastq > count/B7191_Fm0.txt
agrep -0 -c 'GACTCAAAGTCTACGGA' *.fastq > count/B7193_Fm0.txt
agrep -0 -c 'CAACTAGCTCATGCCCT' *.fastq > count/B7194_Fm0.txt
agrep -0 -c 'GCCACAGCCCCTAATTA' *.fastq > count/B7195_Fm0.txt
agrep -0 -c 'TATGGTCTGAACACGAC' *.fastq > count/B7196_Fm0.txt
agrep -0 -c 'ATTACCGGCTAGCCCGA' *.fastq > count/B7197_Fm0.txt
agrep -0 -c 'ACAGGTCTTTAACCAGA' *.fastq > count/B7200_Fm0.txt
agrep -0 -c 'GGGATACATGTGCACTG' *.fastq > count/B7201_Fm0.txt
agrep -0 -c 'TCTTTGACCACATTGGG' *.fastq > count/B7202_Fm0.txt
agrep -0 -c 'AGGACTTATCGCAGCCA' *.fastq > count/B7203_Fm0.txt
agrep -0 -c 'CATACAGGGCTACAACT' *.fastq > count/B7204_Fm0.txt
agrep -0 -c 'AGCGTGGGCTAAGTCAG' *.fastq > count/B7205_Fm0.txt
agrep -0 -c 'CGATCGAGCTGGCTGAT' *.fastq > count/B7207_Fm0.txt
agrep -0 -c 'ATTAAAACCCCAACCAG' *.fastq > count/B7208_Fm0.txt
agrep -0 -c 'ATACTCCACCACGTTGA' *.fastq > count/B7210_Fm0.txt
agrep -0 -c 'CTGATCCAGCGAGCGTT' *.fastq > count/B7211_Fm0.txt
agrep -0 -c 'GCCAGATATTGTGGCCC' *.fastq > count/B7214_Fm0.txt
agrep -0 -c 'AGCGTTCAGTTGCAGGA' *.fastq > count/B7215_Fm0.txt
agrep -0 -c 'CCGCCCCTATACATTAC' *.fastq > count/B7217_Fm0.txt
agrep -0 -c 'TCGATCGATAAGCAGTC' *.fastq > count/B7218_Fm0.txt
agrep -0 -c 'TCGAACGTCTTTCACCG' *.fastq > count/B7219_Fm0.txt
agrep -0 -c 'ACCATCTGAAATGTGCT' *.fastq > count/B7220_Fm0.txt
agrep -0 -c 'GCTGTCGTATCGACATA' *.fastq > count/B7222_Fm0.txt
agrep -0 -c 'CTCGAACTGTGAATACC' *.fastq > count/B7223_Fm0.txt
agrep -0 -c 'GTCTGACTTAACCCAGC' *.fastq > count/B7224_Fm0.txt
agrep -0 -c 'TGGCTGTTGTGCTATTG' *.fastq > count/B7226_Fm0.txt
agrep -0 -c 'CGTCATTGAAGGGTTGC' *.fastq > count/B7228_Fm0.txt
agrep -0 -c 'TTACTACGCCGGCGTTT' *.fastq > count/B7229_Fm0.txt
agrep -0 -c 'CGGTAAGCTAGTTCCCC' *.fastq > count/B7230_Fm0.txt
agrep -0 -c 'CTCACGGCAGATACTGA' *.fastq > count/B7232_Fm0.txt
agrep -0 -c 'AACCAGAACGATCAAGT' *.fastq > count/B7233_Fm0.txt
agrep -0 -c 'CGAGGCGATTGCAATGA' *.fastq > count/B7234_Fm0.txt
agrep -0 -c 'GTAGAGGAGCAAGTCCG' *.fastq > count/B7236_Fm0.txt
agrep -0 -c 'ACCCTCCCAGGCTATTC' *.fastq > count/B7237_Fm0.txt
agrep -0 -c 'ACAGGCTACGGGATTTT' *.fastq > count/B7239_Fm0.txt
agrep -0 -c 'CCTGGCGGATGTGAATG' *.fastq > count/B7240_Fm0.txt
agrep -0 -c 'TGCGTGACCGATAGGTG' *.fastq > count/B7241_Fm0.txt
agrep -0 -c 'TACCGGCGTCGGAATTC' *.fastq > count/B7242_Fm0.txt
agrep -0 -c 'GACTCTACACTACCACG' *.fastq > count/B7243_Fm0.txt
agrep -0 -c 'ATTTGGGCTTAGCCCCA' *.fastq > count/B7244_Fm0.txt
agrep -0 -c 'TTCGTGCGGTGAACCCA' *.fastq > count/B7245_Fm0.txt
agrep -0 -c 'ACCCTAGACGCTAGGAC' *.fastq > count/B7246_Fm0.txt
agrep -0 -c 'GCTTATGCTTGGCAACA' *.fastq > count/B7247_Fm0.txt
agrep -0 -c 'TGAGTTGCTTTGGCCGG' *.fastq > count/B7248_Fm0.txt
agrep -0 -c 'GCGGTACGGAGTACTCA' *.fastq > count/B7249_Fm0.txt
agrep -0 -c 'TGTTCATGAGTGGCAGA' *.fastq > count/B7250_Fm0.txt
agrep -0 -c 'GCGTCAGAACGCCTACA' *.fastq > count/B7251_Fm0.txt
agrep -0 -c 'AGGACATTGTAACCTCA' *.fastq > count/B7252_Fm0.txt
agrep -0 -c 'CTATTATGTCTCGGGTC' *.fastq > count/B7253_Fm0.txt
agrep -0 -c 'TAAGCGGAGAAGATCAA' *.fastq > count/B7255_Fm0.txt
agrep -0 -c 'AGGACGGTGAGATGAGA' *.fastq > count/B7256_Fm0.txt
agrep -0 -c 'ACGGTGAATGCACTCAG' *.fastq > count/B7257_Fm0.txt
agrep -0 -c 'GTTAGGTAGGCGCAATT' *.fastq > count/B7258_Fm0.txt
agrep -0 -c 'TGTGCCGTCTTCCTAAA' *.fastq > count/B7259_Fm0.txt
agrep -0 -c 'TGCCTCTACTGGACCGA' *.fastq > count/B7260_Fm0.txt
agrep -0 -c 'CGAATTACTTCAGCAGT' *.fastq > count/B7261_Fm0.txt
agrep -0 -c 'TATTAGCGTGCCGAATC' *.fastq > count/B7262_Fm0.txt
agrep -0 -c 'AGGTGCTACCGCTGATT' *.fastq > count/B7263_Fm0.txt
agrep -0 -c 'GAGGGACTAGGTACGAA' *.fastq > count/B7264_Fm0.txt
agrep -0 -c 'GATCACTACTGATCCAT' *.fastq > count/B7265_Fm0.txt
agrep -0 -c 'CTCGACCTTTTCCAAAC' *.fastq > count/B7266_Fm0.txt
agrep -0 -c 'GTAATCGAGTATCGATG' *.fastq > count/B7267_Fm0.txt
agrep -0 -c 'TCGCCTAAAAGCCGCAA' *.fastq > count/B7268_Fm0.txt
agrep -0 -c 'AAGAAGGGCTCGAAATG' *.fastq > count/B7269_Fm0.txt
agrep -0 -c 'AGTAAGCGAGGCCAAGC' *.fastq > count/B7270_Fm0.txt
agrep -0 -c 'TTGAGGGAATTAACTGC' *.fastq > count/B7271_Fm0.txt
agrep -0 -c 'GATCTGAACGCTGGAAG' *.fastq > count/B7272_Fm0.txt
agrep -0 -c 'ATACGGTCCCAACAAGC' *.fastq > count/B7273_Fm0.txt
agrep -0 -c 'CACGGTCTTAAAGACCT' *.fastq > count/B7274_Fm0.txt
agrep -0 -c 'GATTGATCGCCTTTCAA' *.fastq > count/B7275_Fm0.txt
agrep -0 -c 'TAGGACTACTAGAGGAG' *.fastq > count/B7276_Fm0.txt
agrep -0 -c 'TTACCGGAGCTGTCGTG' *.fastq > count/B7277_Fm0.txt
agrep -0 -c 'TGATACACCAAGGACTG' *.fastq > count/B7278_Fm0.txt
agrep -0 -c 'TCAGCATCGCGTCTGGA' *.fastq > count/B7279_Fm0.txt
agrep -0 -c 'AGGCGGGGTGAATTATG' *.fastq > count/B7280_Fm0.txt
agrep -0 -c 'CCTTGCACAAGGTATGG' *.fastq > count/B7281_Fm0.txt
agrep -0 -c 'GTGCCTGCGATAGTCCA' *.fastq > count/B7282_Fm0.txt
agrep -0 -c 'GTTAGACGGTTGGGGCT' *.fastq > count/B7283_Fm0.txt
agrep -0 -c 'GTACTGCTAAGCAGCGA' *.fastq > count/B7284_Fm0.txt
agrep -0 -c 'GACGGAGGTTGTAGTTA' *.fastq > count/B7285_Fm0.txt
agrep -0 -c 'AGGAGAACCCTTTTGCT' *.fastq > count/B7286_Fm0.txt
agrep -0 -c 'AGGGGCGGCAATACGTA' *.fastq > count/B7287_Fm0.txt
agrep -0 -c 'AGAAGCCCTTACCGTTA' *.fastq > count/B7288_Fm0.txt
agrep -0 -c 'GGTTCTAGTTAGCGTCG' *.fastq > count/B7289_Fm0.txt
agrep -0 -c 'TGAGTTGAACATGAAGG' *.fastq > count/B7291_Fm0.txt
agrep -0 -c 'TGGACTAACATGGAATC' *.fastq > count/B7292_Fm0.txt
agrep -0 -c 'GCAGGGAATTAGGATGT' *.fastq > count/B7293_Fm0.txt
agrep -0 -c 'TTGCCGTCGGAAGCCAT' *.fastq > count/B7295_Fm0.txt
agrep -0 -c 'CTATTGTGGCTTCTATC' *.fastq > count/B7296_Fm0.txt
agrep -0 -c 'ATCCCCAACTGATCTTA' *.fastq > count/B7299_Fm0.txt
agrep -0 -c 'ATCGTATATGGCGGGGT' *.fastq > count/B7300_Fm0.txt
agrep -0 -c 'TGGACTTCTTGGCATAG' *.fastq > count/B7302_Fm0.txt
agrep -0 -c 'TTACTGTGCTCAGAATC' *.fastq > count/B7303_Fm0.txt
agrep -0 -c 'GGGGTTGGCTAGAACCT' *.fastq > count/B7304_Fm0.txt
agrep -0 -c 'ACTCCCAGCAGTCCTAG' *.fastq > count/B7305_Fm0.txt
agrep -0 -c 'AGGATCGAGTGACGTGT' *.fastq > count/B7306_Fm0.txt
agrep -0 -c 'TTCTGATCTCCGTCGAG' *.fastq > count/B7309_Fm0.txt
agrep -0 -c 'GCCGGTAATCTAGCAAT' *.fastq > count/B7311_Fm0.txt
agrep -0 -c 'TCGACTAGAATCAGGCT' *.fastq > count/B7312_Fm0.txt
agrep -0 -c 'GTACACCAGATGTGGGG' *.fastq > count/B7313_Fm0.txt
agrep -0 -c 'TGGACCTGTTTGCTGAC' *.fastq > count/B7314_Fm0.txt
agrep -0 -c 'GTCTTGAAAAAGAGCCT' *.fastq > count/B7315_Fm0.txt
agrep -0 -c 'ATTTCGAGTGCGAGCGC' *.fastq > count/B7318_Fm0.txt
agrep -0 -c 'AGAAGAGCGACCCAAGA' *.fastq > count/B7319_Fm0.txt
agrep -0 -c 'GTCGTTCAGAGTAGTCC' *.fastq > count/B7320_Fm0.txt
agrep -0 -c 'GCTCGATACGCATGCAA' *.fastq > count/B7321_Fm0.txt
agrep -0 -c 'GTAAGGCTATGGGGACG' *.fastq > count/B7322_Fm0.txt
agrep -0 -c 'CTAGCGTGAACTGCGGA' *.fastq > count/B7323_Fm0.txt
agrep -0 -c 'CAGGATTTCGACTCGTG' *.fastq > count/B7324_Fm0.txt
agrep -0 -c 'CCAGCCTAGTAGTATGC' *.fastq > count/B7325_Fm0.txt
agrep -0 -c 'GGAATTGAGCCTATCCC' *.fastq > count/B7326_Fm0.txt
agrep -0 -c 'AGGGGATAAACTGTTAG' *.fastq > count/B7328_Fm0.txt
agrep -0 -c 'GTGCACGAAGGGTAGGT' *.fastq > count/B7329_Fm0.txt
agrep -0 -c 'GCTAGCAGTCCGAACCA' *.fastq > count/B7330_Fm0.txt
agrep -0 -c 'TTGCTACGTCGGTGTCT' *.fastq > count/B7333_Fm0.txt
agrep -0 -c 'AGTCCAAGCAGGGTCGT' *.fastq > count/B7334_Fm0.txt
agrep -0 -c 'AGCTTTCGCGGTATGGC' *.fastq > count/B7335_Fm0.txt
agrep -0 -c 'TCAGCTAATCGCGCGCT' *.fastq > count/B7337_Fm0.txt
agrep -0 -c 'AAGGTCGTAACAAGCCG' *.fastq > count/B7338_Fm0.txt
agrep -0 -c 'ATACGGATCGGCGTACT' *.fastq > count/B7339_Fm0.txt
agrep -0 -c 'CCAAATTGGACGGGCCA' *.fastq > count/B7340_Fm0.txt
agrep -0 -c 'GGAAGAAATTGGGCGCT' *.fastq > count/B7341_Fm0.txt
agrep -0 -c 'GTATTCCAGAACCGACA' *.fastq > count/B7343_Fm0.txt
agrep -0 -c 'ATAAACCTGTTGGCGAA' *.fastq > count/B7346_Fm0.txt
agrep -0 -c 'ATGTTTGTAGCGCTAGG' *.fastq > count/B7347_Fm0.txt
agrep -0 -c 'TTGAATGTAATCGCCCG' *.fastq > count/B7348_Fm0.txt
agrep -0 -c 'GCGCTATATAAGGAAGT' *.fastq > count/B7350_Fm0.txt
agrep -0 -c 'TACCCGAAGATCCGTGT' *.fastq > count/B7351_Fm0.txt
agrep -0 -c 'CATGCTCATCGTACTAG' *.fastq > count/B7352_Fm0.txt
agrep -0 -c 'TCATCAGGTGAGATCCC' *.fastq > count/B7353_Fm0.txt
agrep -0 -c 'TTCTGTAGTCACCCCGC' *.fastq > count/B7354_Fm0.txt
agrep -0 -c 'TTCGCAATGTCGTTTCC' *.fastq > count/B7355_Fm0.txt
agrep -0 -c 'GGATAGGGTAGTAAGGA' *.fastq > count/B7356_Fm0.txt
agrep -0 -c 'TGGAGTAATTCGTTCGG' *.fastq > count/B7357_Fm0.txt
agrep -0 -c 'ATAGAAGTTAGGGGGCA' *.fastq > count/B7358_Fm0.txt
agrep -0 -c 'CGAAGCCCTTGAAAGGT' *.fastq > count/B7359_Fm0.txt
agrep -0 -c 'GCGTTCTCTACCTGAAT' *.fastq > count/B7360_Fm0.txt
agrep -0 -c 'ATGGCCGTTAGCACTAC' *.fastq > count/B7361_Fm0.txt
agrep -0 -c 'AGAATTTACTGAGGGCC' *.fastq > count/B7362_Fm0.txt
agrep -0 -c 'TAGTACGCCTAGGGAAT' *.fastq > count/B7363_Fm0.txt
agrep -0 -c 'ATCGCAACGATTTCAAG' *.fastq > count/B7364_Fm0.txt
agrep -0 -c 'TACTTCTGTGTTCAGGT' *.fastq > count/B7365_Fm0.txt
agrep -0 -c 'TAGCGCCCACATCGACA' *.fastq > count/B7368_Fm0.txt
agrep -0 -c 'TGAGGCAGTACGGAGTC' *.fastq > count/B7369_Fm0.txt
agrep -0 -c 'CATCAGGTGTACCACTC' *.fastq > count/B7370_Fm0.txt
agrep -0 -c 'TTGGATCAAAAAGTCGC' *.fastq > count/B7371_Fm0.txt
agrep -0 -c 'TCACGTCAATAGACCGA' *.fastq > count/B7373_Fm0.txt
agrep -0 -c 'TGAGTGGCTTGCGGCTA' *.fastq > count/B7374_Fm0.txt
agrep -0 -c 'AGCATGGTGTAGTGGTG' *.fastq > count/B7375_Fm0.txt
agrep -0 -c 'GATTGTCAGAAATCTCG' *.fastq > count/B7378_Fm0.txt
agrep -0 -c 'CATAGTTGATAGCCGAC' *.fastq > count/B7379_Fm0.txt
agrep -0 -c 'GCATCAGATTCGCAGTG' *.fastq > count/B7380_Fm0.txt
agrep -0 -c 'TGATCGCCCTTAGCACA' *.fastq > count/B7381_Fm0.txt
agrep -0 -c 'TGATGCAATTTTCGCTG' *.fastq > count/B7382_Fm0.txt
agrep -0 -c 'CTACTCTGCTTCACCGA' *.fastq > count/B7383_Fm0.txt
agrep -0 -c 'GAGCGCTGTAAACCCTA' *.fastq > count/B7384_Fm0.txt
agrep -0 -c 'CTCGAAGGTTTTTGTCC' *.fastq > count/B7385_Fm0.txt
agrep -0 -c 'CGCACGATAGTCTCGAC' *.fastq > count/B7386_Fm0.txt
agrep -0 -c 'CTTGGTCTATACGGAGA' *.fastq > count/B7387_Fm0.txt
agrep -0 -c 'CCGGATTCGAGTTGTAG' *.fastq > count/B7388_Fm0.txt
agrep -0 -c 'ACGACAGTAGCACGTTA' *.fastq > count/B7389_Fm0.txt
agrep -0 -c 'CGCCGTTATTTACGATA' *.fastq > count/B7390_Fm0.txt
agrep -0 -c 'GTGTCAAGTGACAAGAC' *.fastq > count/B7391_Fm0.txt
agrep -0 -c 'CCATGCACACGGGAATC' *.fastq > count/B7392_Fm0.txt
agrep -0 -c 'CGAGAAGCGTGCACAAT' *.fastq > count/B7393_Fm0.txt
agrep -0 -c 'GTGGTTTCGAAAGTCAC' *.fastq > count/B7394_Fm0.txt
agrep -0 -c 'CACTTCTAAGTAATCGC' *.fastq > count/B7395_Fm0.txt
agrep -0 -c 'ACTTTGCCATCTCCGGA' *.fastq > count/B7397_Fm0.txt
agrep -0 -c 'GAGGGTCCATTCCATAG' *.fastq > count/B7399_Fm0.txt
agrep -0 -c 'CACTCCCAGGGACTTAT' *.fastq > count/B7402_Fm0.txt
agrep -0 -c 'TCAGAGGGAGCCACATA' *.fastq > count/B7403_Fm0.txt
agrep -0 -c 'GACGTCTTTTCCAGGAG' *.fastq > count/B7404_Fm0.txt
agrep -0 -c 'TGAGACACCCTATGGCC' *.fastq > count/B7406_Fm0.txt
agrep -0 -c 'TAAACGTCAGCATACCG' *.fastq > count/B7407_Fm0.txt
agrep -0 -c 'TCGCTACTACGCTACTC' *.fastq > count/B7408_Fm0.txt
agrep -0 -c 'GGGTTAACCGCGTTACT' *.fastq > count/B7409_Fm0.txt
agrep -0 -c 'CAACTACACGCTCCCAT' *.fastq > count/B7410_Fm0.txt
agrep -0 -c 'ATCGATCCACGACTCTG' *.fastq > count/B7411_Fm0.txt
agrep -0 -c 'GTGTGGTAAGTCTTCAA' *.fastq > count/B7412_Fm0.txt
agrep -0 -c 'CGAATCAGAGTTACCCG' *.fastq > count/B7413_Fm0.txt
agrep -0 -c 'CGCCAGATCGGTCTTTA' *.fastq > count/B7415_Fm0.txt
agrep -0 -c 'TCGTGGTCTTTTGACTT' *.fastq > count/B7416_Fm0.txt
agrep -0 -c 'CACTCCATGGGAAAGGA' *.fastq > count/B7417_Fm0.txt
agrep -0 -c 'CTGAATCTCTGAATGCC' *.fastq > count/B7418_Fm0.txt
agrep -0 -c 'GACAAAGGCTCAACTGC' *.fastq > count/B7419_Fm0.txt
agrep -0 -c 'CACTTGTGTACGGTATT' *.fastq > count/B7420_Fm0.txt
agrep -0 -c 'TAGTCATCGGCCACCTA' *.fastq > count/B7422_Fm0.txt
agrep -0 -c 'TTAGCAAATCAACGCGC' *.fastq > count/B7423_Fm0.txt
agrep -0 -c 'GGTCCTCACTTGTCGTG' *.fastq > count/B7424_Fm0.txt
agrep -0 -c 'GAGCTATACAACTACTG' *.fastq > count/B7426_Fm0.txt
agrep -0 -c 'TGTCTTTTTGATGTCCC' *.fastq > count/B7427_Fm0.txt
agrep -0 -c 'CTCACAGGTCTCATCTA' *.fastq > count/B7428_Fm0.txt
agrep -0 -c 'AATGTTAATAGCCGAGC' *.fastq > count/B7429_Fm0.txt
agrep -0 -c 'TTCACTGTATGGGTACG' *.fastq > count/B7430_Fm0.txt
agrep -0 -c 'GCGGCTTACGAGTGGTA' *.fastq > count/B7431_Fm0.txt
agrep -0 -c 'ATTGCCGAAGGTCCTTT' *.fastq > count/B7432_Fm0.txt
agrep -0 -c 'AGTAGCCGCTTCGTTCC' *.fastq > count/B7433_Fm0.txt
agrep -0 -c 'GTAGAGGTGTCGCGGTT' *.fastq > count/B7434_Fm0.txt
agrep -0 -c 'CGCTCTCGCTTTTTCAG' *.fastq > count/B7435_Fm0.txt
agrep -0 -c 'TAGGGAGGCGTTCAGAG' *.fastq > count/B7437_Fm0.txt
agrep -0 -c 'TTGTAGCGATCTCCGAC' *.fastq > count/B7438_Fm0.txt
agrep -0 -c 'CTCTTGAGATTGACCGT' *.fastq > count/B7439_Fm0.txt
agrep -0 -c 'CTTGGGGCCAGTATTCG' *.fastq > count/B7443_Fm0.txt
agrep -0 -c 'GCCAGTGGTATACTCCG' *.fastq > count/B7444_Fm0.txt
agrep -0 -c 'GTGACGATCCAACTCGC' *.fastq > count/B7445_Fm0.txt
agrep -0 -c 'ATCCAATCCCGCGGAGT' *.fastq > count/B7447_Fm0.txt
agrep -0 -c 'GTTCTCGGGCTTGCAAA' *.fastq > count/B7448_Fm0.txt
agrep -0 -c 'GGCAATACTCTATTCGT' *.fastq > count/B7449_Fm0.txt
agrep -0 -c 'GGATAGCTGGGGTCGTT' *.fastq > count/B7450_Fm0.txt
agrep -0 -c 'TTAGCCAGTCATGACAT' *.fastq > count/B7451_Fm0.txt
agrep -0 -c 'TCACTGTTAACCTGGGC' *.fastq > count/B7452_Fm0.txt
agrep -0 -c 'AGGTGTTTTTCGATGCA' *.fastq > count/B7453_Fm0.txt
agrep -0 -c 'AGCTGCAGTAGTGACAC' *.fastq > count/B7454_Fm0.txt
agrep -0 -c 'GTCCGACGTTGCTACAC' *.fastq > count/B7455_Fm0.txt
agrep -0 -c 'AATGCAAGCTACTTGCA' *.fastq > count/B7456_Fm0.txt
agrep -0 -c 'TTCAGTCGTTGACATAC' *.fastq > count/B7457_Fm0.txt
agrep -0 -c 'ACTAAATGGCTCTACTG' *.fastq > count/B7458_Fm0.txt
agrep -0 -c 'TTGTCGGTAGCGCTATC' *.fastq > count/B7459_Fm0.txt
agrep -0 -c 'CACCTATCCAGATAGTT' *.fastq > count/B7460_Fm0.txt
agrep -0 -c 'TGGCACATATATCGCAT' *.fastq > count/B7461_Fm0.txt
agrep -0 -c 'AGAGAGTTCGTCGAACT' *.fastq > count/B7462_Fm0.txt
agrep -0 -c 'CTTCACCGGTTAGGAGG' *.fastq > count/B7463_Fm0.txt
agrep -0 -c 'ATGGCTGTCGCTCTAAA' *.fastq > count/B7464_Fm0.txt
agrep -0 -c 'ATCGAACGGTTGGCATG' *.fastq > count/B7466_Fm0.txt
agrep -0 -c 'AACGACACCGTGCTTTT' *.fastq > count/B7467_Fm0.txt
agrep -0 -c 'GACGAAGATAGACTGGC' *.fastq > count/B7468_Fm0.txt
agrep -0 -c 'GGGTGTCGACGTTTTTA' *.fastq > count/B7469_Fm0.txt
agrep -0 -c 'AGGGCCTAAGGCCACAA' *.fastq > count/B7470_Fm0.txt
agrep -0 -c 'CATTAGACCAACGTGGC' *.fastq > count/B7471_Fm0.txt
agrep -0 -c 'CCACACGTGTGTTAGCG' *.fastq > count/B7472_Fm0.txt
agrep -0 -c 'ACGATACTTCCTGACTA' *.fastq > count/B7473_Fm0.txt
agrep -0 -c 'TAGACCCATATGACTGT' *.fastq > count/B7474_Fm0.txt
agrep -0 -c 'AAGATCTGGCCGACGAG' *.fastq > count/B7475_Fm0.txt
agrep -0 -c 'CTACCTTACTCTTAGCC' *.fastq > count/B7476_Fm0.txt
agrep -0 -c 'CCTGGACAACCGTTCTA' *.fastq > count/B7477_Fm0.txt
agrep -0 -c 'GGGGTACTCTGCTTTTG' *.fastq > count/B7478_Fm0.txt
agrep -0 -c 'GTCAGACAAACACTGAT' *.fastq > count/B7479_Fm0.txt
agrep -0 -c 'GGGATACGAAGGCACAC' *.fastq > count/B7480_Fm0.txt
agrep -0 -c 'AGGAAGCTTGGTCGCAT' *.fastq > count/B7481_Fm0.txt
agrep -0 -c 'GTGGTAAGCGGTCAGTG' *.fastq > count/B7482_Fm0.txt
agrep -0 -c 'CCGTTGATAACAATCTG' *.fastq > count/B7483_Fm0.txt
agrep -0 -c 'AGCGCTAATAACTCTTG' *.fastq > count/B7484_Fm0.txt
agrep -0 -c 'GCGATAACTGTAACATG' *.fastq > count/B7485_Fm0.txt
agrep -0 -c 'CCGTCTTAATGCAATCT' *.fastq > count/B7487_Fm0.txt
agrep -0 -c 'GCGCTTTAGTTCAAACC' *.fastq > count/B7488_Fm0.txt
agrep -0 -c 'CATAGCAGCGACTGATA' *.fastq > count/B7489_Fm0.txt
agrep -0 -c 'GGTCGAGGAATTAAAAG' *.fastq > count/B7490_Fm0.txt
agrep -0 -c 'CAATGGTGTCCGATTAC' *.fastq > count/B7491_Fm0.txt
agrep -0 -c 'TGCTCGCGAGACCGAAA' *.fastq > count/B7492_Fm0.txt
agrep -0 -c 'TGCCAACAGGACGAGTC' *.fastq > count/B7493_Fm0.txt
agrep -0 -c 'CTAAAGCATCGATGGGG' *.fastq > count/B7496_Fm0.txt
agrep -0 -c 'AAGAGCTCCTATGACGA' *.fastq > count/B7497_Fm0.txt
agrep -0 -c 'GGAGCATTCTTTCGGGC' *.fastq > count/B7498_Fm0.txt
agrep -0 -c 'CATCCCATTGCCCTACG' *.fastq > count/B7501_Fm0.txt
agrep -0 -c 'ATGGGCGACTGTATTGG' *.fastq > count/B7502_Fm0.txt
agrep -0 -c 'TAGCGCGCCATTACGTC' *.fastq > count/B7503_Fm0.txt
agrep -0 -c 'TCGACAGAGATGCGAGA' *.fastq > count/B7504_Fm0.txt
agrep -0 -c 'CCGTTTACGTACAATGG' *.fastq > count/B7507_Fm0.txt
agrep -0 -c 'TAATAGGGAACGTAACC' *.fastq > count/B7508_Fm0.txt
agrep -0 -c 'TCGTCTATAACAAGTGC' *.fastq > count/B7509_Fm0.txt
agrep -0 -c 'GGAGGGAATCTGTTAAT' *.fastq > count/B7510_Fm0.txt
agrep -0 -c 'ACCGCTGTGGTCAGAAT' *.fastq > count/B7511_Fm0.txt
agrep -0 -c 'CCTTTGGGAACCTTATG' *.fastq > count/B7512_Fm0.txt
agrep -0 -c 'ACAAGAGATCGGGTCTT' *.fastq > count/B7513_Fm0.txt
agrep -0 -c 'AATGTGGGCCCTTCGAC' *.fastq > count/B7514_Fm0.txt
agrep -0 -c 'GTTGCGATCTTTGCTTG' *.fastq > count/B7515_Fm0.txt
agrep -0 -c 'TATAGTTCTAGGACCCT' *.fastq > count/B7516_Fm0.txt
agrep -0 -c 'TGGAAACATTGGAGTAG' *.fastq > count/B7517_Fm0.txt
agrep -0 -c 'TCGATGACGGCCTATAC' *.fastq > count/B7518_Fm0.txt
agrep -0 -c 'GATTCAGACGGTAAAAC' *.fastq > count/B7519_Fm0.txt
agrep -0 -c 'GTCTGTATTCACTAGGG' *.fastq > count/B7521_Fm0.txt
agrep -0 -c 'TTACTCGGCCATCTTAA' *.fastq > count/B7522_Fm0.txt
agrep -0 -c 'GCTATGGTTTGACACTT' *.fastq > count/B7523_Fm0.txt
agrep -0 -c 'TCAAACCCTTAATTGCC' *.fastq > count/B7524_Fm0.txt
agrep -0 -c 'GCTCATACGTAGAACAC' *.fastq > count/B7525_Fm0.txt
agrep -0 -c 'GTTCCGAATCGAAGTTC' *.fastq > count/B7526_Fm0.txt
agrep -0 -c 'GCCTAATAGCGGGTAGT' *.fastq > count/B7527_Fm0.txt
agrep -0 -c 'GTCACTTTTAGGCATGA' *.fastq > count/B7528_Fm0.txt
agrep -0 -c 'ACGCGGAGCTAAAGTGA' *.fastq > count/B7529_Fm0.txt
agrep -0 -c 'ATACTCATCGCTAACCT' *.fastq > count/B7531_Fm0.txt
agrep -0 -c 'AGAGGGTACCTCCTCCT' *.fastq > count/B7532_Fm0.txt
agrep -0 -c 'TACTGACTCACGCAGAA' *.fastq > count/B7533_Fm0.txt
agrep -0 -c 'AGAGTAACCAGCGATGC' *.fastq > count/B7534_Fm0.txt
agrep -0 -c 'CCTGACAAATTGCCGGA' *.fastq > count/B7536_Fm0.txt
agrep -0 -c 'TTGTACCGAGCAAGCGA' *.fastq > count/B7537_Fm0.txt
agrep -0 -c 'ATCGAGGGGTAAGCGCA' *.fastq > count/B7538_Fm0.txt
agrep -0 -c 'CGTTATCGGTACACTTA' *.fastq > count/B7539_Fm0.txt
agrep -0 -c 'ACTAACGGGAGGCATCA' *.fastq > count/B7540_Fm0.txt
agrep -0 -c 'GCTAGTCAGTGCCCACT' *.fastq > count/B7541_Fm0.txt
agrep -0 -c 'GAGAGTTTTACCCCGGG' *.fastq > count/B7542_Fm0.txt
agrep -0 -c 'CAGCGGGTTGAATTGCC' *.fastq > count/B7543_Fm0.txt
agrep -0 -c 'TTCAGAGTGACACTCTA' *.fastq > count/B7544_Fm0.txt
agrep -0 -c 'TCAGGCAATCGTTCCTT' *.fastq > count/B7546_Fm0.txt
agrep -0 -c 'CTGGCGTATCACACGCT' *.fastq > count/B7547_Fm0.txt
agrep -0 -c 'ACGGACCATACAGTCTT' *.fastq > count/B7548_Fm0.txt
agrep -0 -c 'ATGCGAAAACGGAGGTT' *.fastq > count/B7550_Fm0.txt
agrep -0 -c 'TGACCGCCTAACATTCG' *.fastq > count/B7551_Fm0.txt
agrep -0 -c 'CAAAAAGCCGGACACAC' *.fastq > count/B7552_Fm0.txt
agrep -0 -c 'ATATCGCGGAGGTGGGT' *.fastq > count/B7553_Fm0.txt
agrep -0 -c 'ACTGGGGCGATCAACCT' *.fastq > count/B7555_Fm0.txt
agrep -0 -c 'TATTTACAGGCGCACCC' *.fastq > count/B7556_Fm0.txt
agrep -0 -c 'GACCAAACGGAGATCAC' *.fastq > count/B7561_Fm0.txt
agrep -0 -c 'TTCCCGTAGTCGTTAGA' *.fastq > count/B7562_Fm0.txt
agrep -0 -c 'ACTCAAAGGGAACTCAG' *.fastq > count/B7563_Fm0.txt
agrep -0 -c 'CCTGAGCGTTTAGAATG' *.fastq > count/B7565_Fm0.txt
agrep -0 -c 'AAGCAGCGAACTTATGT' *.fastq > count/B7566_Fm0.txt
agrep -0 -c 'CCGATTGGGCTCGTATT' *.fastq > count/B7568_Fm0.txt
agrep -0 -c 'GTTAAGCCGTCAAGATA' *.fastq > count/B7569_Fm0.txt
agrep -0 -c 'ATTCGTTCTCTTCCTTC' *.fastq > count/B7570_Fm0.txt
agrep -0 -c 'GGGTACTCATCTAGGAC' *.fastq > count/B7571_Fm0.txt
agrep -0 -c 'ATCAACGACGGCCACGA' *.fastq > count/B7572_Fm0.txt
agrep -0 -c 'CTAAGAACATGCGTTTG' *.fastq > count/B7573_Fm0.txt
agrep -0 -c 'GTTAGGTCCATGCATAC' *.fastq > count/B7574_Fm0.txt
agrep -0 -c 'AACATTCTGACGCCATA' *.fastq > count/B7576_Fm0.txt
agrep -0 -c 'TTAGCCCCTGAGGTAAG' *.fastq > count/B7578_Fm0.txt
agrep -0 -c 'CGAATTCAGGACTTACC' *.fastq > count/B7579_Fm0.txt
agrep -0 -c 'GTCTCCAGCGGAATCTC' *.fastq > count/B7580_Fm0.txt
agrep -0 -c 'TGGAGTGTATGACTCTA' *.fastq > count/B7582_Fm0.txt
agrep -0 -c 'GACGCGATTTGTCACCC' *.fastq > count/B7583_Fm0.txt
agrep -0 -c 'TACTAAACCGGCGGAAG' *.fastq > count/B7585_Fm0.txt
agrep -0 -c 'TATGCGACTGCCGAAGA' *.fastq > count/B7586_Fm0.txt
agrep -0 -c 'GGCCTTACTATCTGAAA' *.fastq > count/B7588_Fm0.txt
agrep -0 -c 'CGTCACAACAATGCCGG' *.fastq > count/B7589_Fm0.txt
agrep -0 -c 'ACCAGCGCCCTCGTTAA' *.fastq > count/B7590_Fm0.txt
agrep -0 -c 'AACCGAAGCGTGCTCTT' *.fastq > count/B7591_Fm0.txt
agrep -0 -c 'ATGGCTCAGACCCTTCG' *.fastq > count/B7592_Fm0.txt
agrep -0 -c 'TGTCGCCACTTTGATTC' *.fastq > count/B7593_Fm0.txt
agrep -0 -c 'CATAAAGTCGACATGAG' *.fastq > count/B7594_Fm0.txt
agrep -0 -c 'CGACTGAATGATGATCC' *.fastq > count/B7595_Fm0.txt
agrep -0 -c 'AATTATAGCAGGGCTGT' *.fastq > count/B7596_Fm0.txt
agrep -0 -c 'GTATACCCTGTGGTATC' *.fastq > count/B7597_Fm0.txt
agrep -0 -c 'GCCCTAACTGTCTCGGT' *.fastq > count/B7598_Fm0.txt
agrep -0 -c 'CGAGCTGTAGTTGAAGC' *.fastq > count/B7599_Fm0.txt
agrep -0 -c 'ACCCTGAGAAGGTCAGC' *.fastq > count/B7600_Fm0.txt
agrep -0 -c 'GTAATTACGTCACGTAC' *.fastq > count/B7601_Fm0.txt
agrep -0 -c 'TCGACGCCTCTCTAATC' *.fastq > count/B7602_Fm0.txt
agrep -0 -c 'GATGTGACTGACCCGTT' *.fastq > count/B7605_Fm0.txt
agrep -0 -c 'CCAAAGGATTACCCAAG' *.fastq > count/B7606_Fm0.txt
agrep -0 -c 'GTGAGTTGTGTGCCCCT' *.fastq > count/B7607_Fm0.txt
agrep -0 -c 'TGCCGGATTACGATTGG' *.fastq > count/B7608_Fm0.txt
agrep -0 -c 'GGTCCTGCAACTTGAAG' *.fastq > count/B7609_Fm0.txt
agrep -0 -c 'TGGCATCAACAATACGC' *.fastq > count/B7610_Fm0.txt
agrep -0 -c 'TTACGCGCCATTTCAGG' *.fastq > count/B7611_Fm0.txt
agrep -0 -c 'TCACGACGAGAAGTGAC' *.fastq > count/B7612_Fm0.txt
agrep -0 -c 'GGTGTGGCTGGCATCAT' *.fastq > count/B7613_Fm0.txt
agrep -0 -c 'GCAGATGGGCCATATAA' *.fastq > count/B7614_Fm0.txt
agrep -0 -c 'GATTCACGGCCCGAATC' *.fastq > count/B7616_Fm0.txt
agrep -0 -c 'CCGAAGTTGCTATCGCT' *.fastq > count/B7619_Fm0.txt
agrep -0 -c 'TGAGTTTCGGATCCGTG' *.fastq > count/B7620_Fm0.txt
agrep -0 -c 'CATGGACAACCGAATAG' *.fastq > count/B7621_Fm0.txt
agrep -0 -c 'ACGTTGCAACCAAGTTG' *.fastq > count/B7622_Fm0.txt
agrep -0 -c 'AGTAAGTCCCGGAGTGT' *.fastq > count/B7623_Fm0.txt
agrep -0 -c 'TCGGTCAGAGGATCAAT' *.fastq > count/B7625_Fm0.txt
agrep -0 -c 'TAGACTGTTCATGCGTC' *.fastq > count/B7626_Fm0.txt
agrep -0 -c 'TGCAGTCGGGAATCACG' *.fastq > count/B7627_Fm0.txt
agrep -0 -c 'CCGGTGAGACAAATACA' *.fastq > count/B7628_Fm0.txt
agrep -0 -c 'ACCTCGTCGCAAAGCGA' *.fastq > count/B7629_Fm0.txt
agrep -0 -c 'GGCCAAACATCTGCGTG' *.fastq > count/B7630_Fm0.txt
agrep -0 -c 'CTAGTTTCGATCCCACT' *.fastq > count/B7631_Fm0.txt
agrep -0 -c 'TCACGTATGGGAGATTC' *.fastq > count/B7632_Fm0.txt
agrep -0 -c 'CTCTGCTGACCGAAAAG' *.fastq > count/B7634_Fm0.txt
agrep -0 -c 'TCGCATGCCTTACCCTT' *.fastq > count/B7635_Fm0.txt
agrep -0 -c 'TAGCATAGGATCGATGA' *.fastq > count/B7636_Fm0.txt
agrep -0 -c 'TAAGGCGGCAATTGACT' *.fastq > count/B7637_Fm0.txt
agrep -0 -c 'ATGACGGTTTGTGACAT' *.fastq > count/B7638_Fm0.txt
agrep -0 -c 'CTAATATCAGAACGGGC' *.fastq > count/B7640_Fm0.txt
agrep -0 -c 'GTTAGTAGGGATTAGGA' *.fastq > count/B7641_Fm0.txt
agrep -0 -c 'GCAGTACCTATGTCTTG' *.fastq > count/B7643_Fm0.txt
agrep -0 -c 'TTAGGTGGGTATAGGCA' *.fastq > count/B7644_Fm0.txt
agrep -0 -c 'GACACCGGTGATTTTTA' *.fastq > count/B7647_Fm0.txt
agrep -0 -c 'AACAATCCTTTGGTGAG' *.fastq > count/B7648_Fm0.txt
agrep -0 -c 'ACACGTCTTACTACCTG' *.fastq > count/B7649_Fm0.txt
agrep -0 -c 'GTGGGTTAGTTCATGGA' *.fastq > count/B7650_Fm0.txt
agrep -0 -c 'GCGCCTCGTTTAATGTA' *.fastq > count/B7652_Fm0.txt
agrep -0 -c 'CCAATGCCCAATGGCCA' *.fastq > count/B7653_Fm0.txt
agrep -0 -c 'CAGAAGAGGCTTACGGA' *.fastq > count/B7654_Fm0.txt
agrep -0 -c 'CGGGGAAATCAGGATGA' *.fastq > count/B7655_Fm0.txt
agrep -0 -c 'TTGGTATTACCCCGGTT' *.fastq > count/B7656_Fm0.txt
agrep -0 -c 'CCTTTGTTCACTGGGCG' *.fastq > count/B7657_Fm0.txt
agrep -0 -c 'TTAGTTCCGCACCCGTT' *.fastq > count/B7658_Fm0.txt
agrep -0 -c 'GCTAGATCTACCCAATA' *.fastq > count/B7659_Fm0.txt
agrep -0 -c 'CACGCTAGTAAGCTGTT' *.fastq > count/B7660_Fm0.txt
agrep -0 -c 'ACCGGATATGGCAGCTA' *.fastq > count/B7661_Fm0.txt
agrep -0 -c 'AAGTTGCATACTCGTGC' *.fastq > count/B7662_Fm0.txt
agrep -0 -c 'TGACCGCCTCTGGATCA' *.fastq > count/B7663_Fm0.txt
agrep -0 -c 'GGAAGCGGCGGAAATAG' *.fastq > count/B7664_Fm0.txt
agrep -0 -c 'TGAATGAGATCAACTGG' *.fastq > count/B7665_Fm0.txt
agrep -0 -c 'GATTCCTGCAGGGCGAA' *.fastq > count/B7666_Fm0.txt
agrep -0 -c 'CCGGCAGGTACTGAAAT' *.fastq > count/B7667_Fm0.txt
agrep -0 -c 'CATCGATTTGGGTAAGC' *.fastq > count/B7668_Fm0.txt
agrep -0 -c 'ATAATGATGTACCTGCG' *.fastq > count/B7669_Fm0.txt
agrep -0 -c 'GGTCGTGGCCCTGAAAT' *.fastq > count/B7671_Fm0.txt
agrep -0 -c 'TATCGTTTTAGGCCAAC' *.fastq > count/B7673_Fm0.txt
agrep -0 -c 'ACGGTGGTCCAGTAACG' *.fastq > count/B7674_Fm0.txt
agrep -0 -c 'GCAGGAAAGGCGGATGT' *.fastq > count/B7676_Fm0.txt
agrep -0 -c 'ATACTGGCCTGTTCGTT' *.fastq > count/B7677_Fm0.txt
agrep -0 -c 'ACGTCGTAAGGATCGAA' *.fastq > count/B7678_Fm0.txt
agrep -0 -c 'ACTGCGATGTCGATTGG' *.fastq > count/B7679_Fm0.txt
agrep -0 -c 'CTCACCACATTCCGATT' *.fastq > count/B7680_Fm0.txt
agrep -0 -c 'GACCGCGATTCATGGAC' *.fastq > count/B7681_Fm0.txt
agrep -0 -c 'CGCAATTCAGAGGGTCT' *.fastq > count/B7682_Fm0.txt
agrep -0 -c 'CTTGCTCCCATGTTCGC' *.fastq > count/B7683_Fm0.txt
agrep -0 -c 'CGCAGATTTGCGGATGT' *.fastq > count/B7684_Fm0.txt
agrep -0 -c 'GTCTTTGCAGAGTCTAG' *.fastq > count/B7685_Fm0.txt
agrep -0 -c 'TTAGTTAACGGACGGCC' *.fastq > count/B7687_Fm0.txt
agrep -0 -c 'ACGGTTAGGGCGCAACT' *.fastq > count/B7688_Fm0.txt
agrep -0 -c 'AGCGGCCATCTTCATGA' *.fastq > count/B7689_Fm0.txt
agrep -0 -c 'ATCGAGTTCCAACGGCG' *.fastq > count/B7690_Fm0.txt
agrep -0 -c 'TACAACGTGTCCGCGGA' *.fastq > count/B7692_Fm0.txt
agrep -0 -c 'AATGCCCTAATACCACG' *.fastq > count/B7693_Fm0.txt
agrep -0 -c 'CCTGCAACTTGATAACG' *.fastq > count/B7694_Fm0.txt
agrep -0 -c 'TCGAACACGTTAGGTAA' *.fastq > count/B7695_Fm0.txt
agrep -0 -c 'GGGTCCAGGCATAATGC' *.fastq > count/B7696_Fm0.txt
agrep -0 -c 'CACGAGACTTGCCGGTT' *.fastq > count/B7698_Fm0.txt
agrep -0 -c 'CTACGTCAGGGAACTCG' *.fastq > count/B7699_Fm0.txt
agrep -0 -c 'ACACTTGGGTTCTGACT' *.fastq > count/B7701_Fm0.txt
agrep -0 -c 'GTAGGGCTTAACCAGTT' *.fastq > count/B7702_Fm0.txt
agrep -0 -c 'AGCTTCATCTTTGGCGA' *.fastq > count/B7703_Fm0.txt
agrep -0 -c 'AGTTAACGTCCCTTGGG' *.fastq > count/B7704_Fm0.txt
agrep -0 -c 'AGAAGATACGTGTGGGA' *.fastq > count/B7706_Fm0.txt
agrep -0 -c 'TAGGTCTGGCAATGAGC' *.fastq > count/B7707_Fm0.txt
agrep -0 -c 'CGCTTAGCAGGGAACGT' *.fastq > count/B7708_Fm0.txt
agrep -0 -c 'GGGTAGGAAAACCTTCA' *.fastq > count/B7711_Fm0.txt
agrep -0 -c 'GGAGTAACTAACGACTT' *.fastq > count/B7712_Fm0.txt
agrep -0 -c 'TGATGGCGGTCGTAGCT' *.fastq > count/B7713_Fm0.txt
agrep -0 -c 'AGAAGGTCCAGTTTGTA' *.fastq > count/B7714_Fm0.txt
agrep -0 -c 'CATTAGGGGGCTAGCTA' *.fastq > count/B7715_Fm0.txt
agrep -0 -c 'ATTATGAACATGGTCGG' *.fastq > count/B7716_Fm0.txt
agrep -0 -c 'CATTGCATTTCGCCCAT' *.fastq > count/B7717_Fm0.txt
agrep -0 -c 'GATCGTTCACAGTCGAG' *.fastq > count/B7718_Fm0.txt
agrep -0 -c 'CTCGTAACCCCGTATAT' *.fastq > count/B7719_Fm0.txt
agrep -0 -c 'TCTGGAGCATCGAACTA' *.fastq > count/B7720_Fm0.txt
agrep -0 -c 'ATGTCGCCGTAACGGTC' *.fastq > count/B7721_Fm0.txt
agrep -0 -c 'GCTCATCATCTACTGGA' *.fastq > count/B7723_Fm0.txt
agrep -0 -c 'TCTTACCGGTAGGTGAC' *.fastq > count/B7724_Fm0.txt
agrep -0 -c 'TGATGCTAAGTCAGAGC' *.fastq > count/B7725_Fm0.txt
agrep -0 -c 'GGTTAATAGAAGCCGTT' *.fastq > count/B7728_Fm0.txt
agrep -0 -c 'ATCTGCTCAAGAAGTCA' *.fastq > count/B7730_Fm0.txt
agrep -0 -c 'GAGGATTGTTCTAAGCC' *.fastq > count/B7731_Fm0.txt
agrep -0 -c 'ACGCGGGACACGCATTT' *.fastq > count/B7732_Fm0.txt
agrep -0 -c 'GCCTAGAGGAGATTGCC' *.fastq > count/B7733_Fm0.txt
agrep -0 -c 'CCATCAAATGTGGGGAA' *.fastq > count/B7734_Fm0.txt
agrep -0 -c 'GTGCGGGTGGAATCTTT' *.fastq > count/B7736_Fm0.txt
agrep -0 -c 'CACGCCACACAAGATTA' *.fastq > count/B7737_Fm0.txt
agrep -0 -c 'GTCCAGAATGGCTTAAT' *.fastq > count/B7738_Fm0.txt
agrep -0 -c 'TCAATGCTAAGGGGAGA' *.fastq > count/B7739_Fm0.txt
agrep -0 -c 'AAGTTTTTGTCCCGACG' *.fastq > count/B7740_Fm0.txt
agrep -0 -c 'GCCGCTATTCCATAGTT' *.fastq > count/B7741_Fm0.txt
agrep -0 -c 'AAGTGTCTTCCGGAGCT' *.fastq > count/B7742_Fm0.txt
agrep -0 -c 'GGTGGATAACCTGACTG' *.fastq > count/B7743_Fm0.txt
agrep -0 -c 'AGGAAACGGTACGTTAA' *.fastq > count/B7744_Fm0.txt
agrep -0 -c 'CTGCGACACTGATGACT' *.fastq > count/B7745_Fm0.txt
agrep -0 -c 'CGCGGATTTTTCCGTCT' *.fastq > count/B7746_Fm0.txt
agrep -0 -c 'GGATTTCAGGATACGGC' *.fastq > count/B7747_Fm0.txt
agrep -0 -c 'ACCCGTAATGATGAGCC' *.fastq > count/B7748_Fm0.txt
agrep -0 -c 'GAGCGGGCAGATAAAGG' *.fastq > count/B7749_Fm0.txt
agrep -0 -c 'CAGTCAACGGCCTATGT' *.fastq > count/B7750_Fm0.txt
agrep -0 -c 'CTCGCAGGTGAGAGGAA' *.fastq > count/B7751_Fm0.txt
agrep -0 -c 'TCAATGTTGCCTTACGT' *.fastq > count/B7752_Fm0.txt
agrep -0 -c 'CGTTGTCAGCCGTTCGA' *.fastq > count/B7753_Fm0.txt
agrep -0 -c 'AGATGGAAGGGCTTTAT' *.fastq > count/B7754_Fm0.txt
agrep -0 -c 'ACGGATAAGGCTTGATG' *.fastq > count/B7755_Fm0.txt
agrep -0 -c 'CACGCCTCATATGGAGT' *.fastq > count/B7758_Fm0.txt
agrep -0 -c 'TATTGCACCCGGGGTTG' *.fastq > count/B7759_Fm0.txt
agrep -0 -c 'ACCAAGTAGTATCTCTG' *.fastq > count/B7760_Fm0.txt
agrep -0 -c 'CAAGTTCTTGCTACCTG' *.fastq > count/B7761_Fm0.txt
agrep -0 -c 'GGGTTGCATATTGGGGG' *.fastq > count/B7762_Fm0.txt
agrep -0 -c 'TCGAACAACGTTTACGG' *.fastq > count/B7763_Fm0.txt
agrep -0 -c 'CCTGGGAGCGTATACGA' *.fastq > count/B7764_Fm0.txt
agrep -0 -c 'CAGACAGGGTTTACACG' *.fastq > count/B7765_Fm0.txt
agrep -0 -c 'TCACGGAGTCGCACTAT' *.fastq > count/B7766_Fm0.txt
agrep -0 -c 'ACATTTGTAGCGCGTGG' *.fastq > count/B7767_Fm0.txt
agrep -0 -c 'AACTTGTTCAGGTCGCG' *.fastq > count/B7768_Fm0.txt
agrep -0 -c 'GGAATCCTAACCAAGTA' *.fastq > count/B7769_Fm0.txt
agrep -0 -c 'ACGAGCTAGTTCTTGCC' *.fastq > count/B7770_Fm0.txt
agrep -0 -c 'GCAACGTGCTTTACTAC' *.fastq > count/B7771_Fm0.txt
agrep -0 -c 'ATCGCACGCGGTTGACT' *.fastq > count/B7772_Fm0.txt
agrep -0 -c 'CTGGTCCCCGTCTGTTA' *.fastq > count/B7773_Fm0.txt
agrep -0 -c 'ATACGATCTTTGCGGGT' *.fastq > count/B7774_Fm0.txt
agrep -0 -c 'CCTGTCGGCTATTTTTC' *.fastq > count/B7775_Fm0.txt
agrep -0 -c 'ACGTTATCTACGATCTC' *.fastq > count/B7778_Fm0.txt
agrep -0 -c 'GTGGACCGTCGGAACTA' *.fastq > count/B7779_Fm0.txt
agrep -0 -c 'CCGGACACGTAGCAAAG' *.fastq > count/B7780_Fm0.txt
agrep -0 -c 'TGCGTCGGGTACACACT' *.fastq > count/B7781_Fm0.txt
agrep -0 -c 'TAACAGTGCACGAAGCG' *.fastq > count/B7784_Fm0.txt
agrep -0 -c 'AAGTGGATTGCACATGG' *.fastq > count/B7785_Fm0.txt
agrep -0 -c 'TCTTGCTAAGGTACCGT' *.fastq > count/B7786_Fm0.txt
agrep -0 -c 'GGATACACCGGCAATTA' *.fastq > count/B7787_Fm0.txt
agrep -0 -c 'TTATCCTCTCGACTACT' *.fastq > count/B7788_Fm0.txt
agrep -0 -c 'CGACCTAAGAGCGGTCT' *.fastq > count/B7789_Fm0.txt
```
