## Supplementary_file_3 for "Bar-seq strategies for the LeishGEdit toolbox": pivot_table.py

```
#!/usr/bin/python

import pandas as pd

#pivot table for excel
column_names = ['File', 'Count', 'ID']
df = pd.read_csv('all_counts.csv', header=None, names = column_names)


df2 = df.pivot('ID', 'File', 'Count')

df2.drop(df.ID[:1], inplace=True)

df2.to_csv('0mismatch_all_barcodes_count.csv')


print ("Barcode count done! All files exported!")
```
